# Mathematical modeling of the influence of *ACE I/D* polymorphism on blood pressure and antihypertensive therapy

**DOI:** 10.1101/2024.01.09.574774

**Authors:** Elena Kutumova, Anna Kovaleva, Ruslan Sharipov, Galina Lifshits, Fedor Kolpakov

## Abstract

The angiotensin converting enzyme (ACE) gene (*ACE*) insertion/deletion (*I/D*) polymorphism has attracted much attention in recent years, as it raises the hope of personalizing ACE inhibitor therapy to optimize its efficiency and reduce side effects for genetically distinct subgroups. However, the extent of its influence among these subgroups remains inconclusive. Therefore, we extended our computational model of blood pressure regulation to investigate the effect of the *ACE I/D* polymorphism on hemodynamic parameters in humans and antihypertensive therapy. The model showed that the dependence of blood pressure on serum ACE activity is a function of saturation. Hence, a possible reason for the lack of association between *ACE I/D* and blood pressure levels could be a fairly high ACE activity in populations. Additionally, in an extended model simulating the effects of different classes of antihypertensive drugs, we explored the relationship between *ACE I/D* and the efficacy of inhibitors of the renin-angiotensin-aldosterone system. The model predicted that the response of cardiovascular and renal parameters to treatment directly depends on ACE activity. However, significant differences in parameter changes were observed only between groups with high and low ACE levels, while *ACE I/D* genotypes within the same group had similar changes in absolute values. We conclude that a single genetic variant is responsible for only a small fraction of heredity in treatment success, so its predictive value is limited.

## 1 Introduction

The human angiotensin-converting enzyme (ACE) gene (*ACE*) contains 26 exons interrupted by 25 introns and maps to chromosome 17q23 (Hubert et al., 1991; Crisan and Carr, 2000; Rigat et al., 1992). The *ACE* insertion/deletion (*I/D*) polymorphism (rs1799752, rs4340, rs13447447, or rs4646994), identified in 1990 by Rigat and colleagues (Rigat et al., 1990), is one of the most studied polymorphisms (Scharplatz et al., 2004). It is characterized by the presence or absence of a 287-bp *Alu* repetitive sequence within intron 16 (Rigat et al., 1992), resulting in three genotypes: *DD* and *II* homozygotes and *ID* heterozygotes (Samani et al., 1996; Gard, 2010). The *ACE D* allele is associated with the risk of coronary heart disease (Cambien et al., 1992) and myocardial infarction (Seckin et al., 2006), which is also confirmed by the meta-analysis data (Staessen et al., 1997). However, meta-analyses of small and large studies did not support the association of *ACE* genotype with blood pressure, and only supported an association with myocardial infarction and ischemic heart disease in small studies (Agerholm-Larsen et al., 2000). Some experiments have shown an association between the *DD* genotype and a higher prevalence of left ventricular hypertrophy (Montgomery et al., 1997; Cosenso-Martin et al., 2015; Bahramali et al., 2016), which was not found in other studies (Staessen et al., 1997). The relationship between *ACE I/D* polymorphism and the pathogenesis of essential hypertension is also controversial. Results of meta-analyses indicated that the *ACE D* allele is associated with hypertension susceptibility in Asian, Caucasian, and mixed populations (Liu et al., 2021), as well as in the Chinese population (Li, 2012), but this association was inconclusive in people of West African descent (Reiter et al., 2016). Heterogeneous data were also obtained for individual population groups. Thus, Barley et al. (1996) revealed that in white people of European descent there was no significant association between *ACE* genotype and high blood pressure, while black people of Afro-Caribbean descent showed a positive association between the frequency of the *D* allele and increasing blood pressure. O”Donnell et al. (1998) examined a sample of Framingham Heart Study participants and found consistent evidence for genetic linkage of the *ACE* locus with hypertension and blood pressure in men but not in women. Ned et al. (2012) reported that among non-Hispanic black people, the *D* allele was linked with increased systolic blood pressure (SBP) in additive and dominant covariate-adjusted models, and was also associated with increased diastolic blood pressure (DBP) in dominant models when participants taking ACE inhibitors were excluded from the analyses. However, significant genotype-sex interactions were detected only among Mexican Americans (positive associations with SBP and hypertension in women, but not in men), while this was not observed for non-Hispanic white persons and non-Hispanic black persons. Multivariate regression analysis by Han et al. (2017) demonstrated that the *ACE DD* genotype was independently associated with 24□hour SBP, 24□hour DBP, central SBP, and central augmentation index in Chinese patients recruited at Shandong Provincial Hospital in the Jinan area. Thus, it can be concluded that the influence of variation in the *ACE* gene on interindividual variation in blood pressure is dependent on contexts that are indexed by gender, age, and measures of body size (Turner et al., 1999). At the same time, ethnic predisposition cannot be ruled out. In addition to the above, it has been shown that *ACE I/D* polymorphism contribute to the development of hypertension in Swedes (Stefansson et al., 2000); North Indians (Sameer et al., 2010; Srivastava et al., 2012; Singh et al., 2016; Rana et al., 2018); Asian Indians (Das et al., 2008); residents of Gujarat, Western India (Patel et al., 2022); Saudi subjects (Ali et al., 2013); Russians (Kovaleva et al., 2019); indigenous ethnic group of Mountain Shoria, Russia (Barbarash et al., 2017); Japanese (Yoshida et al., 2000); Punjabi population from Faisalabad, Pakistan (Hussain et al., 2018); population of Burkina Faso, West Africa (Tchelougou et al., 2015); Ethiopian population, East Africa (Birhan et al., 2022); as well as Han, Kazakh, Tibetan, and Zhuang Chinese populations (Li, 2012; Sun et al., 2018). In contrast, the association between *ACE I/D* genotypes and hypertension has not been established in Slovenians (Glavnik and Petrovic, 2007); Buryats (Kovaleva et al., 2019); Thais (Charoen et al., 2019); Romany subjects and Slovaks (Danková et al., 2009); Cuban population, primarily of European and African ancestry, living in Havana (Nápoles et al., 2007); Algerian population from Oran (Meroufel et al., 2014); and some Chinese minorities, including Mongolians, Uyghurs, Yugurs, Koreans, and others (Li, 2012).

Despite controversy regarding the relationship of *ACE I/D* variants with cardiovascular diseases, experimental studies agree that the *ACE* polymorphism affects plasma ACE concentration and activity, which are increased with the number of *D* alleles (Rigat et al., 1990; Tiret et al., 1992; Staessen et al., 1997; Brown et al., 1998; Woods et al., 2000; Agerholm-Larsen et al., 2000; Lee et al., 2009; Mehri et al., 2010; Dai et al., 2019; Bánhegyi et al., 2021). A plausible mechanism for the lack of effect of *ACE I/D* on the cardiovascular system could be that subjects with the *DD* genotype, despite elevated plasma levels of ACE, do not necessarily produce increased amounts of angiotensin II (Agerholm-Larsen et al., 2000). On the other hand, a number of experimental investigations noted that increased ACE activity can lead to higher levels of angiotensin II (Danser et al., 1995; Brown et al., 1998). Probably, in this case, there is an association between the *ACE* genotype and cardiovascular risk.

The results of numerous studies confirm that genotype-based antihypertensive therapies are the most effective (Figure 1) and may help to avoid the occurrence of major adverse events, as well as decrease the costs of treatment (Rysz et al., 2020). It has been suggested that differences in circulating ACE might affect the therapeutic response of ACE inhibitors, explaining interindividual variability in cardiovascular or renal response to equivalent doses of drugs. Several studies have examined the extent of influence of *ACE* genotype on the effectiveness of ACE inhibitors in various conditions, such as hypertension, diabetic nephropathy and coronary artery disease (Scharplatz et al., 2004). However, the question of whether *ACE* genotyping helps in predicting the success of ACE inhibition is remains unresolved (Danser et al., 2007).

**Figure 1.**
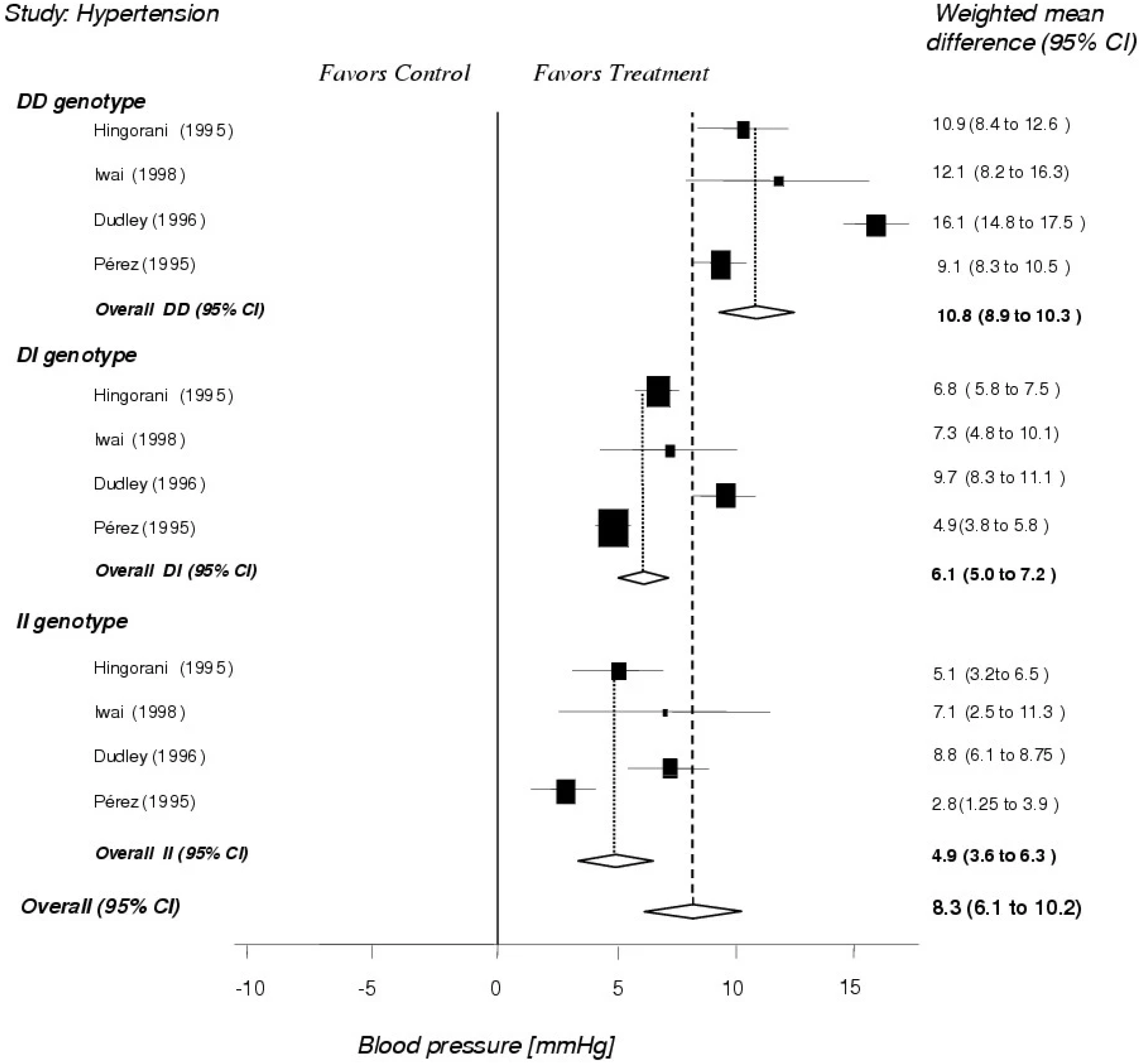
Differences of the ACE inhibitor effect in hypertensive patients with different *ACE I/D* genotypes. The diamonds below each of the three genotypes indicate the pooled results. The lowest (fourth) diamond reflects the overall effect of ACE inhibitors across all genotypes. In this example, the *DD* genotype shows the largest ACE inhibitor effect and the *II* genotype shows the smallest effect. The size of the box is related to the number of studied patients. Reprinted with permission from (Scharplatz et al., 2004).

To explore the association of *ACE I/D* polymorphism with the development of arterial hypertension *in silico*, we used a previously created mathematical model of the human cardiovascular and renal systems (Kutumova et al., 2021). Primary model incorporated the processes of blood circulation and the cardiac cycle, neurohumoral regulation, oxygen exchange between blood and tissue, the renin-angiotensin-aldosterone system (RAAS), renal microcirculation and sodium transport along the nephron, renal sympathetic nerve activity, and regulation of water and sodium balance. The model was further improved to account for the effects of various antihypertensive agents, including RAAS blockers such as the direct renin inhibitor aliskiren, the ACE inhibitor enalapril, and the angiotensin II receptor blocker losartan, as well as drugs with other mechanisms of action, such as the β-blocker bisoprolol, the calcium channel blocker amlodipine, and the thiazide diuretic hydrochlorothiazide (Kutumova et al., 2022).

Using this model, in the present study, we investigated why *ACE I/D* genotypes are associated with the development of hypertension in some ethnic groups but not in others, and whether they are associated with the success of antihypertensive therapy with RAAS blockers. For computational analysis, we used the BioUML software, an extensible open source platform that adapts a visual modeling approach to formally describe and simulate complex biological systems (Kolpakov et al., 2019; Kolpakov et al., 2022).

## 2 Results

### 2.1 Modelling the dependence of blood pressure on ACE activity

In humans, angiotensin (Ang) I is converted to Ang II by ACE and chymase (Urata et al., 1996). In our model (Kutumova et al., 2021), we defined this process according to Hallow et al. (2014) as two first-order biochemical reactions with the following rate constants:

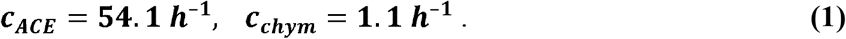

Ang II participates in blood pressure regulation through vasoconstriction and control of fluid-electrolyte balance (Hall, 1991; Yang et al., 2011). Therefore, the first computational experiment was to analyze the dependence of arterial pressure on ACE activity (*c*_*ACE*_).

We generated a virtual normotensive subject, i.e. found an equilibrium parameterizations of the model with indicators of a healthy person (for more details, see Materials and Methods), and analyzed the change in the equilibrium values of SBP and DBP with varying the parameter *c*_*ACE*_. As can be seen from Figure 2, blood pressure-ACE activity curves were similar to the saturation functions. Since serum ACE activity for the three genotypes (*II, ID*, and *DD*) increases with the number of *D* alleles (Staessen et al., 1997; Brown et al., 1998; Agerholm-Larsen et al., 2000; Lee et al., 2009; Mehri et al., 2010; Bánhegyi et al., 2021), we invoked the following hypotheses.

**Figure 2.**
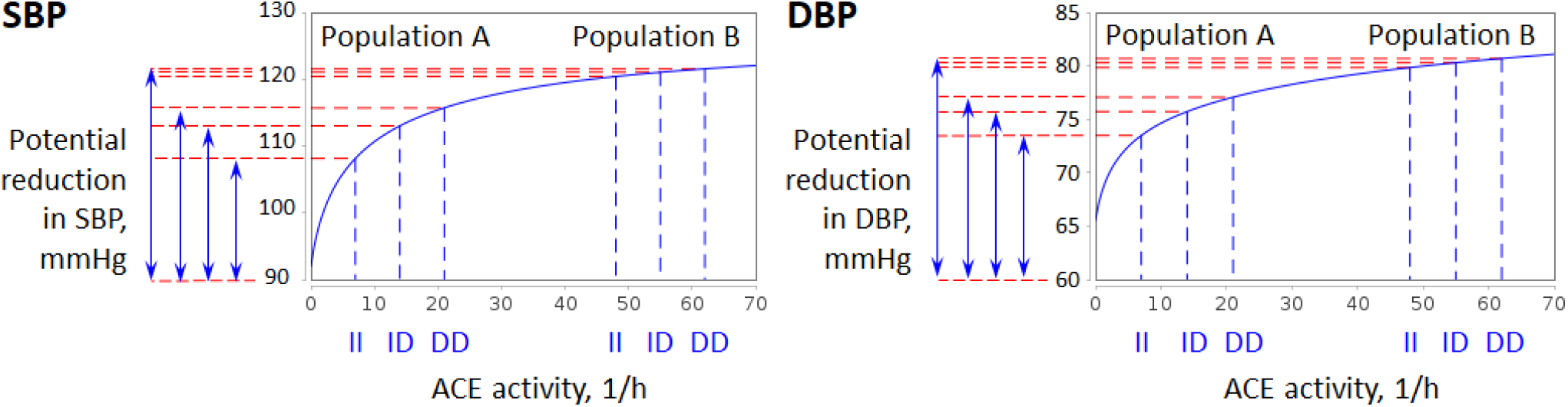
Simulated dependence of SBP and DBP on serum ACE activity (solid blue lines). Since ACE activity increases with the number of *D* alleles in the *ACE* gene, the modeled curves suggest that *ACE* polymorphism could be a marker of hypertension in population *A* with low ACE activity, while there is no such association for population *B* with high ACE activity. In addition, the potential reduction in blood pressure with ACE blockers is greater in population *B* than in population *A*.

#### Hypothesis 1

A possible reason for the lack of association between *ACE I/D* genotypes and blood pressure levels could be a fairly high ACE activity in the ethnic group (population *B* in Figure 2). In contrast, ethnic groups with such an association are likely to have low ACE activity (population *A* in Figure 2).

#### Hypothesis 2

The potential reduction in blood pressure with ACE blockers is greater in populations without the specified association than in populations with it.

An interactive implementation of the simulation experiment shown in Figure 2 is available online in the BioUML software (see Data Availability).

### 2.2 Generation and analysis of populations with different ACE activity

We generated two types of virtual hypertensive populations with low and high ACE activity to study the relationship between *ACE I/D* polymorphism and the success of RAAS inhibitor therapy. The choice of *c*_*ACE*_ parameter values in both cases and for different *ACE* genotypes was based on the following considerations.

Primary *c*_*ACE*_ and *c*_*chym*_ values (1) were taken from the renin-angiotensin system model developed by Lo et al. (2010), where a number of assumptions were made to estimate them.

1. The activity of ACE and chymase in converting Ang I to Ang II was assumed proportional to the concentration of the substrate (Ang I).
2. ACE was assumed to be responsible for > 95% of the conversion of Ang I to Ang II.

To be precise, the rate constants (1) correspond to 98% of the ACE contribution to the formation of Ang II. However, some experimental studies have reported different results. For example, analysis of angiotensin metabolism in human kidney homogenates showed a 93% contribution of ACE to Ang II (Domenig, 2016), while in a group of orthotopic heart transplant recipients with normal LV function, ACE mediated 89% of the conversion of Ang I to Ang II across the myocardial circulation *in vivo* (Zisman et al., 1995).

Therefore, we regarded the 98% contribution of ACE to Ang II as a high ACE activity and the 89% contribution as a low ACE activity with the corresponding values of *c*_*ACE*_:

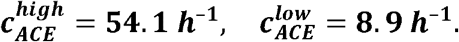

Assuming that these constants are the means of the three *ACE I/D* variants, we took into account that plasma ACE activity is increased by an average of 28% and 56% in *ID* and *DD* genotypes compared to *ACE II* (Agerholm-Larsen et al., 2000). Thus, we obtained the following ACE activities for different cases:

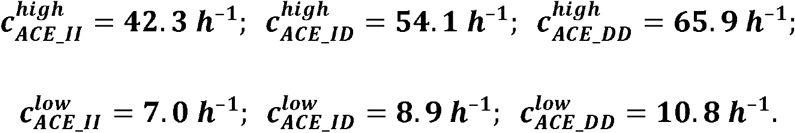

Using this data, we generated six populations (one for each *c*_*ACE*_ value) of 100 virtual patients with arterial hypertension (see the generation algorithm in Materials and Methods). The baseline characteristics of the populations are given in Table 1. Distribution plots for all parameters are shown in Supplementary File 1, Figures S1-S9 and are available in the Jupyter document on the web version of BioUML (see Data Availability). There were no significant differences between populations in most parameters (Table 1 and Supplementary File 1, Table S1). However, with increasing ACE activity, there was a trend towards higher Ang II (*P* ≤ 0.001 for all populations compared to each other), lower Ang I (*P* < 0.001 for any high *c*_*ACE*_ value *vs*. all other cases), and lower plasma renin activity (*P* < 0.001 for low *vs*. high *c*_*ACE*_). In addition to the RAAS components, intraglomerular hemodynamic parameters were also sensitive to changes in the *c*_*ACE*_ value. In populations with high ACE activity, the afferent arteriolar diameter was larger than in populations with low ACE activity (*P* < 0.001 for all pairs of such populations). The glomerular hydrostatic pressure slightly increased along with the *c*_*ACE*_ value (*P* < 0.001 for low ACE activities compared to *c*_*ACE*_ = 65.9 h^−1^), while the afferent arteriolar resistance tended to decrease (*P* = 0.001 for *c*_*ACE*_= 7.0 h^−1^ *vs. c*_*ACE*_ = 65.9 h^−1^). Note that for all populations, we used the same generation algorithm that utilized a random number function for SBP, DBP, heart rate, stroke volume, body weight, and body mass index with the same means and standard deviations (160 ± 10 mmHg, 100 ± 10 mmHg, 75 ± 10 beats/min, 70 ± 10 mL, 80 ± 20 kg, 29 ± 5 kg/m^2^, respectively). Thus, the resulting differences in populations were associated with the maintenance of the same level of these parameters with variations in ACE activity.

**Table 1.**
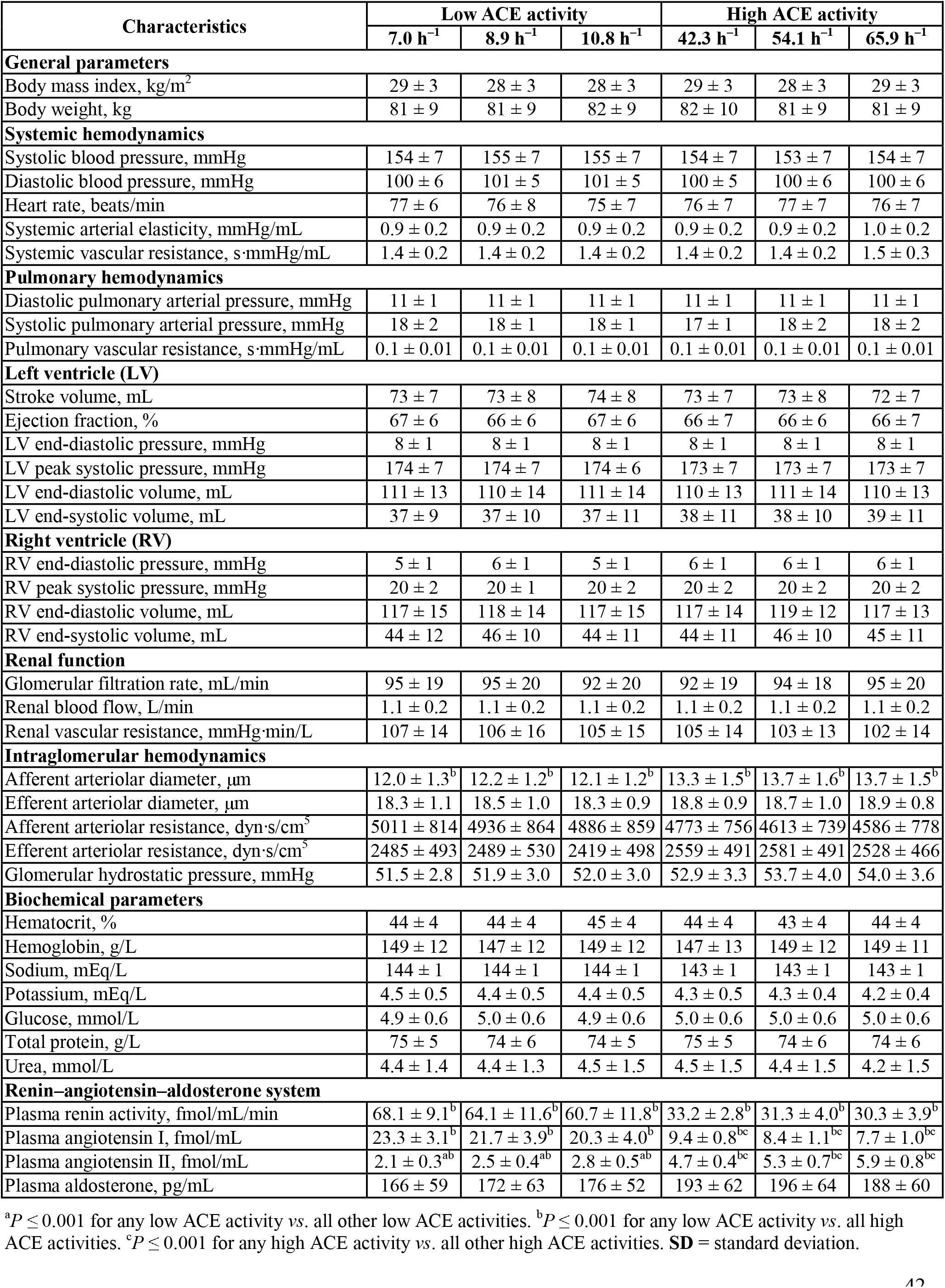
Baseline characteristics of virtual populations (*n* = 100). Data are shown as mean ± SD.

### 2.3 Simulation of antihypertensive treatment

To analyze the antihypertensive efficacy of RAAS blockers at various ACE activities, we simulated a 4-week treatment of the created virtual populations with enalapril 20 mg, losartan 100 mg, and aliskiren 300 mg. We also considered amlodipine 5 mg, bisoprolol 5 mg and hydrochlorothiazide 12.5 mg as additional drugs for combination therapy. For computational experiments at this stage of the work, we used an extension of the primary model, including the pharmacodynamic functions of these drugs in the indicated dosages (Kutumova et al., 2022). Note that in the current study, the bisoprolol and hydrochlorothiazide models were improved to better fit the clinical data (see Materials and Methods for details). As before, therapeutic parameters were estimated for the case of *c*_*ACE*_ = 54.1 h^−1^. A detailed analysis of the predicted response of the cardiovascular and renal systems to antihypertensive therapy for this case, as well as an agreement of the predictions with experimental studies, was carried out earlier (Kutumova et al., 2022). Therefore, in the current research, we mainly focused on comparing simulation results obtained with different ACE activity.

Figure 3 shows a simulated decrease in SBP and DBP with antihypertensive mono- and combination therapy. Figures S10-S35 in Supplementary File 1 demonstrate similar plots for all the physiological characteristics in Table 1 that were significantly altered by drugs. In addition, Tables S2-S57 (Supplementary File 1) contain numerical data for these characteristics, including values at baseline and after treatment, absolute parameter changes, and *P*-values calculated using the Kolmogorov-Smirnov test for endpoint *vs*. baseline and for parameter changes in populations treated with the same regimens. An interactive script that calculates and visualizes all statistics is implemented online (see Data Availability).

**Figure 3.**
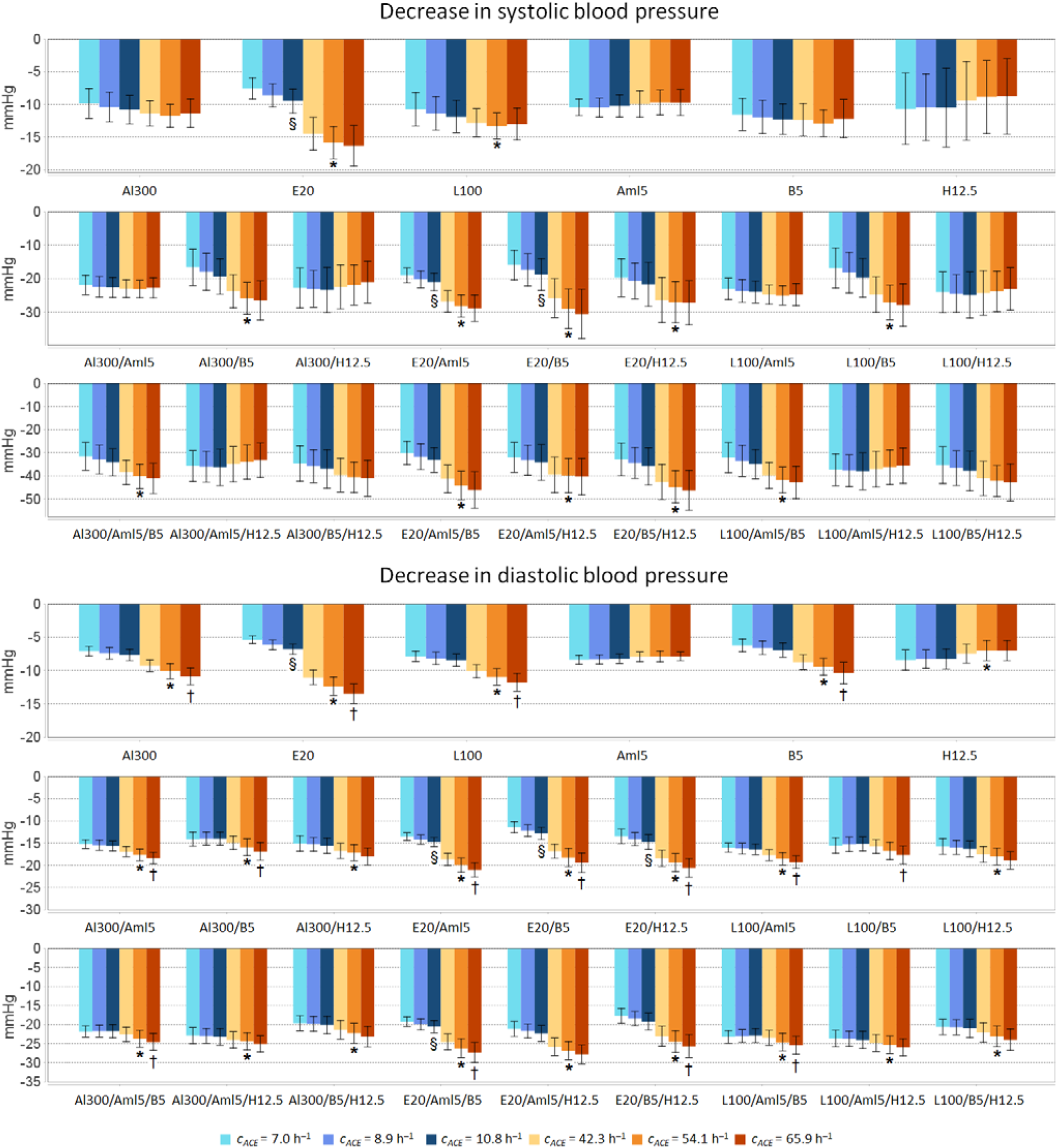
Simulated decrease in systolic and diastolic blood pressure from baseline to week 4 with aliskiren 300 mg (Al300), enalapril 20 mg (E20), losartan 100 mg (L100), amlodipine 5 mg (Aml5), bisoprolol 5 mg (B5), hydrochlorothiazide 12.5 mg (H12.5), as well as double and triple combinations of these drugs in populations (*n* = 100) with different ACE activity (*c*_*A C E*_). \**P* < 10 –5 *νs. c_A C E_* = 8.9 1/h. ^§^ *P* < 10^−5^ *νs. c*_*A C E*_ = 7.0 1/h. ^†^*P* < 10^−5^ *νs. c*_*A C E*_ = 42.3 1/h. Data are mean ± SD.

#### 2.3.1 Systemic hemodynamics

All drugs produced a statistically significant (*P* < 10^−5^) drop in SBP and DBP in all populations (Supplementary File 1, Tables S2 and S4). As can be seen from Figure 3 and Tables S3 and S5 (Supplementary File 1), a statistically greater decrease in SBP with high *vs*. low ACE activity was observed for all enalapril-based regimens, for losartan alone, and for combinations of aliskiren-bisoprolol, losartan-bisoprolol, aliskiren-amlodipine-bisoprolol, and losartan-amlodipine-bisoprolol. In addition, all therapies except amlodipine and the concomitant use of losartan and bisoprolol showed a clear difference in DBP reduction between high and low ACE populations. RAAS blockers (aliskiren, enalapril, losartan) and bisoprolol, whose ability to suppress renin secretion (Laragh and Sealey, 2011) was reproduced in the model, as well as all double and triple combinations of drugs, were better at lowering DBP with high ACE activity, while hydrochlorothiazide monotherapy showed the opposite trend.

There were no significant differences in SBP reduction between populations with high ACE activity, whereas in populations with low ACE activity, enalapril-based regimens without hydrochlorothiazide showed a statistically greater decrease in SBP for *ACE DD* compared to *ACE II* (*P* < 10^−4^). For DBP, statistical differences between *DD* and *II* variants were more frequently observed for high ACE activity than for low ACE activity.

Among single drugs, only bisoprolol led to a significant change (*P* < 10^−4^) in heart rate at all *c*_*ACE*_ values (Supplementary File 1, Figure S10 and Table S6), because its effect was modeled on the fact that selective β1-blockers have a negative chronotropic effect (Bazroon and Alrashidi, 2022). The model predicted a direct relationship between heart rate reduction with bisoprolol and *c*_*ACE*_ values in populations with high ACE activity, which also showed a significantly greater decrease in heart rate than the low ACE group (Supplementary File 1, Table S7). Co-administration of a RAAS blocker and amlodipine increased heart rate, with aliskiren- and losartan-based regimens resulting in a smaller increase with greater ACE activity. Similar dynamics were also observed in dual combinations with hydrochlorothiazide, but the difference between populations with low and high *c*_*ACE*_ values for aliskiren- and losartan-based regimens was more pronounced. Interestingly, among dual combinations with bisoprolol, a statistically significant decrease in heart rate was observed only for the case of *c*_*ACE*_ = 7.0 h^−1^ (*P* < 10^−4^), while it was practically absent at high levels of ACE activity. All triple drug combinations contributed to the increase in heart rate, which was a consequence of a strong decrease in blood pressure in the model. A significant difference between the *ACE ID* variants in high and low ACE groups was found in the populations treated with aliskiren-amlodipine-hydrochlorothiazide, enalapril-amlodipine-bisoprolol, losartan-amlodipine-bisoprolol, and losartan-amlodipine-hydrochlorothiazide (Supplementary File 1, Table S7).

In the current research, we extended the pharmacodynamic model of bisoprolol to take into account the decrease in systemic arterial elasticity during treatment (see Materials and Methods). As a result, all bisoprolol-based regimens reduced this variable (*P* < 0.005, Supplementary File 1, Figure S11 and Table S8). However, there were no statistically significant differences between populations (Supplementary File 1, Table S9).

All medications led to a decrease in systemic vascular resistance (Supplementary File 1, Figure S12), although in the case of monotherapy it was weak (on average, 3-9% of baseline, *P*-values from 0.00025 to 0.37, Supplementary File 1, Table S10). There was a statistically significantly smaller decrease for *c*_*ACE*_ = 54.1 h^−1^ *vs. c*_*ACE*_ = 8.9 h^−1^ for aliskiren or losartan in double combination with bisoprolol or in triple combination with bisoprolol and amlodipine (*P*_25_ in Supplementary File 1, Table S11). In contrast, enalapril (alone or in combination with amlodipine or hydrochlorothiazide) in this case showed a statistically significantly greater decrease. In addition, enalapril was the only drug that demonstrated significant difference between populations with low ACE activity (*P*_13_ in Supplementary File 1, Table S11).

#### 2.3.2 Pulmonary hemodynamics

Bisoprolol caused a small but statistically significant increase in diastolic and systolic pulmonary arterial pressure (Supplementary File 1, Figures S13-S14 and Tables S12, S14), which was practically independent of ACE activity (Supplementary File 1, Tables S13, S15). However, dependency appeared with the addition of RAAS blockers, which, in dual combinations with bisoprolol, prevented the increase in pulmonary arterial pressure in populations with high *c*_*ACE*_ values, but enhanced it in populations with low *c*_*ACE*_ values. Moreover, a significant difference between these two groups of populations remained in triple combinations with bisoprolol.

#### 2.3.3 Left ventricle

The model showed an increase in stroke volume with bisoprolol (Supplementary File 1, Figure S15). There were no significant differences within the groups with high and low ACE activity, but in the high ACE group, stroke volume increased more than in the low ACE group (Supplementary File 1, Tables S16-S17). For other monodrugs, no statistically significant changes in stroke volume were observed. The difference in the dynamics for high *vs*. low ACE groups persisted with the co-use of bisoprolol and RAAS blockers, as well as with the subsequent addition of amlodipine, while the addition of hydrochlorothiazide reduced it.

Ejection fraction was significantly decreased in all populations with only triple drug combinations (Supplementary File 1, Figure S16 and Table S18). For amlodipine-bisoprolol-based regimens, there was a trend towards a higher decrease with increasing *c*_*ACE*_ value (*P* < 0.001 for *ACE II vs. ACE DD* in the low ACE group and *P* < 10^−5^ between *ACE ID* variants in the groups with low and high ACE levels, Supplementary File 1, Table S19).

The simulated changes in left ventricular end-diastolic pressure for all populations and treatment regimens (Supplementary File 1, Figure S17 and Tables S20-S21) were similar to those for diastolic pulmonary arterial pressure (Supplementary File 1, Figure S13 and Tables S12-S13). At the same time, left ventricular peak systolic pressure was closely related to systolic blood pressure, namely, the plots showing the response of these variables to antihypertensive therapy were almost the same (Figure 3 and Supplementary File 1, Figure S18), as were the conclusions about the statistical significance of the changes (Supplementary File 1, Tables S2-S3 and S22-S23).

Another relationship demonstrated by the model was observed between left ventricular end-diastolic volume and stroke volume according to the Frank-Starling law (Braunwald et al., 2001). Within physiologic limits, the heart pumps all the blood that returns to it by the way of the veins (Hall, 2011). As venous return increases, end-diastolic volume also increases, and due to the length-tension relationship in the ventricles, stroke volume increases accordingly (Costanzo, 2014). Implementation of the Frank-Starling mechanism in the model (Kutumova et al., 2022) caused the same dynamics of changes in end-diastolic volume (Supplementary File 1, Figure S19 and Tables S24-S25) as described above for stroke volume. The modeling of bisoprolol in the present study included the negative inotropic effect of β1-blockers (Bazroon and Alrashidi, 2022), characterized by a decrease in stroke volume at a given end-diastolic volume. Therefore, bisoprolol-based regimens induced an increase in end-systolic volume (Supplementary File 1, Figure S20 and Table S26). However, statistically significant differences between populations were observed only for the combination of enalapril, amlodipine and bisoprolol (Supplementary File 1, Table S27).

#### 2.3.4 Right ventricle

The simulated response of right ventricular end-diastolic pressure, peak systolic pressure, end-diastolic and end-systolic volumes to antihypertensive therapy in all populations is presented in Supplementary File 1, Figures S21-S24 and Tables S28-S35. The dynamics of changes in end-diastolic pressure and volume with varying the value of *c*_*ACE*_ turned out to be the same as for the left ventricle. The response of right ventricular peak systolic pressure to treatment was the same as for systolic pulmonary arterial pressure. In addition, end-systolic volume increased with bisoprolol-based regimens due to the negative inotropic effect of the drug. However, in contrast to the left ventricle, there was a statistically significantly greater increase in end-systolic volume for *c*_*ACE*_ = 8.9 h^−1^ *vs. c*_*ACE*_ = 54.1 h^−1^ during therapy with double and triple combinations of bisoprolol, RAAS blockers and amlodipine (*P* < 0.0005).

#### 2.3.5 Renal function and intraglomerular hemodynamics

As follows from Supplementary File 1, Figure S25 and Table S36, glomerular filtration rate remained unchanged with aliskiren, enalapril, losartan, and bisoprolol, but slightly increased with amlodipine-based regimens and slightly decreased with hydrochlorothiazide-based regimens (without amlodipine). However, these changes were not statistically different between populations (Supplementary File 1, Table S37).

In our example, all drugs increased renal blood flow (Supplementary File 1, Figure S26). Although in the case of monodrugs and double combinations with bisoprolol, as well as all double combinations in patients with low ACE activity, this increase tended to be insignificant (Supplementary File 1, Table S38). For almost all drugs, the response of renal blood flow significantly depended on the *c*_*ACE*_ value, namely, high ACE activity resulted in a greater increase in this parameter (Supplementary File 1, Table S39).

All antihypertensive regimens contributed to a decrease in afferent and efferent arteriolar resistance and, consequently, to a decrease in renal vascular resistance (Supplementary File 1, Figures S27-S29 and Tables S40, S42, S44). RAAS blockers and bisoprolol administered alone resulted in a statistically stronger decrease in these resistances with high ACE activity, while the opposite trend was observed for hydrochlorothiazide (Supplementary File 1, Tables S41, S43, S45). Interestingly, in the baseline populations, afferent arteriolar resistance tended to decrease with increasing *c*_*ACE*_ value (Table 1). For amlodipine, there were no significant differences between populations. In this regard, the addition of hydrochlorothiazide and amlodipine to RAAS blockers reduced the difference between groups with high and low ACE activity.

Glomerular hydrostatic pressure was statistically significantly reduced in all treatment regimens, except for monotherapy with amlodipine and hydrochlorothiazide. At the same time, a direct relationship was observed between the magnitude of the reduction and the value of *c*_*ACE*_ for enalapril-based regimens (Supplementary File 1, Figure S30 and Tables S46-S47).

#### 2.3.6 Biochemical parameters

All blood counts listed in Table 1, except for sodium, are constant in the model. To account for the fact that thiazide diuretics reduce sodium reabsorption (Duarte and Cooper-DeHoff, 2010; Akbari and Khorasani-Zadeh, 2023), we previously included the corresponding effect of hydrochlorothiazide in the model (Kutumova et al., 2022). As a result, in the current study, we observed a slight decrease in circulating sodium levels with hydrochlorothiazide-based therapy (Supplementary File 1, Figure S31 and Table S48). There were no statistically significant differences between populations in these treatment regimens (Supplementary File 1, Table S49).

#### 2.3.7 Renin–angiotensin–aldosterone system

The simulated response of RAAS parameters (plasma renin activity, PRA; Ang I; Ang II; and aldosterone) to antihypertensive therapy was determined by the physiological targets of each individual class of drugs (see Materials and Methods). Taking cross-treatment effects into account, the model demonstrated the dynamics shown in Supplementary File 1, Figures S32-S35 and Tables S50, S52, S54, and S56. Aliskiren and all combinations with it reduced PRA, Ang I, Ang II, and aldosterone. With low ACE activity, bisoprolol eliminated the increase in PRA and Ang I when used together with enalapril and amlodipine, while other enalapril-based regimens led to a statistically significant increase in these parameters. All combinations with enalapril contributed to the reduction of Ang II and aldosterone. Losartan-based regimens induced a rise in PRA, Ang I, Ang II and a fall in aldosterone.

All predicted changes in RAAS parameters were highly dependent on the *c*_*ACE*_ value (Supplementary File 1, Tables S51, S53, S55, and S57). Moreover, the model showed a positive correlation between baseline levels of PRA, Ang I, Ang II and DBP reduction for RAAS-acting agents (aliskiren, enalapril, losartan, and bisoprolol), with a correlation coefficient that was directly related to ACE activity (increased along with the *c*_*ACE*_ value, Supplementary File 1, Table S58). The same trend persisted for dual combinations of RAAS blockers with amlodipine. Although for aldosterone, such a relationship was not observed. For systolic pressure, this trend was also absent (Supplementary File 1, Table S59).

## 3 Discussion

The *ACE I/D* polymorphism is one of the most studied polymorphisms (Scharplatz et al., 2004). However, its association with cardiovascular disease (particularly hypertension) remains controversial, as does its ability to predict the therapeutic effect of ACE inhibitors. Therefore, the aim of our study was to find out why *ACE I/D* genotypes are associated with the development of hypertension in some ethnic groups but not in others, using a previously created computational model of the human cardiorenal system. Computational experiments suggested that, firstly, a possible reason for the lack of association between *ACE I/D* and blood pressure levels could be a sufficiently high ACE activity in populations, and secondly, the potential decrease in blood pressure with ACE inhibitors is greater in populations without this association than in populations with it. We have not been able to find experimental confirmation or refutation of these hypotheses in the scientific literature, so they need experimental verification. To our knowledge, our study is the first attempt to model the influence of genetic factors on blood pressure regulation.

We also examined the relationship between the *ACE I/D* polymorphism and the success of RAAS inhibitor therapy. To do this, we considered two types of virtual populations of hypertensive patients with low and high ACE activity. Since plasma ACE levels have been reported to increase with the number of *ACE D* alleles, we assigned a specific value for the ACE activity parameter in the model for each genetic variant in both population types. Summarizing the above, we can formulate the following model-based conclusions:

1. An increase in ACE activity is accompanied by a rise in circulating Ang II. Under these conditions, maintaining blood pressure at the same level is achieved through the parameters of intraglomerular hemodynamics, namely, through the larger afferent arteriolar diameter, which entails a decrease in the afferent arteriolar resistance and an increase in the glomerular hydrostatic pressure.
2. In an ethnic group with high ACE activity, the association between different *ACE I/D* genotypes and blood pressure levels is erased, but this group is generally more susceptible to developing hypertension than the ACE-low population, where the *ACE DD* variant becomes a risk factor.
3. The response of cardiovascular and renal parameters to treatment with RAAS blockers directly depends on ACE activity. However, significant variations in parameter changes are observed only with pronounced differences in ACE levels, while the *ACE I/D* polymorphisms within the same ethnic group give similar changes in absolute values.

With regard to the latter conclusion, as an example, we note that a statistically significant difference in the effect of the ACE inhibitor enalapril on SBP between *ACE II* and *DD* variants was observed only in populations with low ACE activity (Supplementary File 1, Table S3). However, this difference was about 2 mmHg, i.e. only 1.3% of the baseline SBP. Thus, our results suggest that any single genetic variant explains only a very small fraction of heritability in the success of treating hypertension, which is consistent with a similar statement about the development of the disease (Zhang et al., 2010; Waken et al., 2017). Nevertheless, one should take into account the fact that the adjustment of the pharmacodynamic parameters of drugs using experimental data was carried out for the case of high ACE activity (*c*_*ACE*_ = 54.1 h^−1^). It is possible that refitting these parameters with lower ACE activity (assuming such populations could be recruited for clinical trials) will result in a more significant difference in blood pressure reduction between various *ACE I/D* variants.

The correspondence of the simulated response of cardiovascular and renal parameters to single-drug treatment with experimental observations was carried out in our previous study on modeling antihypertensive therapy (Kutumova et al., 2022). The dynamics of model parameters in the case of double and triple drug combinations in the present research were a consequence of cross-drug effects. However, some findings of the model should be discussed. The model predicted significant reductions in ejection fraction in all populations receiving triple drug combinations (Supplementary File 1, Figure S16). This finding is unexpected considering the use of RAAS inhibitors and β-blockers as anti-remodeling drugs (Cohn et al., 2000; Frigerio and Roubina, 2005; Ferrario, 2016). To clarify this contradiction, we should turn to the pathophysiological aspects of cardiac remodeling and the role of the Frank-Starling mechanism in heart failure. The Frank-Starling relationship is an intrinsic property of myocardium by which increased ventricular volume results in enhanced performance during the subsequent contraction (Braunwald et al., 1967; Braunwald et al., 2001; Moss and Fitzsimons, 2002). Changes in myocardial contractility (or inotropism) affect Frank-Starling curves (Hillegass, 2011; Costanzo, 2014). Namely, positive inotropic agents cause an increase in left ventricular stroke volume (SV) for a given end-diastolic volume (EDV) and therefore an increase in ejection fraction (EF), calculated as

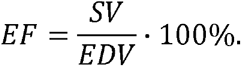

Negative inotropic agents have the opposite effect (Costanzo, 2014). Heart failure is also often associated with impaired contractility (Braunwald et al., 2001).

In the current study, we considered virtual patients with uncomplicated essential hypertension. The model showed an increase in SV with bisoprolol (Supplementary File 1, Figure S15), which is consistent with clinical data for this group of patients (Serg et al., 2014; Suojanen et al., 2017). Consequently, the implementation of the Frank-Starling law in the model led to an increase in EDV (Supplementary File 1, Figure S19). Because bisoprolol has a negative inotropic effect, the SV/EDV ratio decreased, as did EF (although this decrease was only ~2% and was not statistically significant; Supplementary File 1, Figure S16). With the use of other RAAS inhibitors, a slight decrease in EF by ~1% was also observed. Thus, with triple combinations of drugs (especially with the inclusion of bisoprolol), these effects were cumulative and the reduction in EF reached an average of 6-10%, becoming statistically significant.

In patients with heart failure, bisoprolol reduces left ventricular EDV and practically does not change SV (Dubach et al., 2002; Lee et al., 2016). This also corresponds to the Frank-Starling law (Kutumova et al., 2021), but leads to the fact that bisoprolol improves EF in patients with reduced values of the parameter (Dubach et al., 2002; Lee et al., 2016). The ability of the model to simulate the appropriate Frank-Starling curves in patients with heart failure has been previously validated (Kutumova et al., 2021). Therefore, the model is expected to show an improvement in EF for RAAS inhibitors in this group of patients. However, testing this hypothesis is beyond the scope of this study and requires further confirmation.

It is also worth noting that the model showed a strong relationship between left ventricular end-diastolic pressure (Supplementary File 1, Figure S17) and diastolic pulmonary arterial pressure (Supplementary File 1, Figure S13). This result is consistent with the fact that these values correlate with each other and with pulmonary artery wedge pressure in patients with normal left ventricular function (Falicov and Resnekov, 1970; Bouchard et al., 1971). At the same time, changes in end-diastolic pressure and volume of the right ventricle with varying *c*_*ACE*_ values turned out to be the same as for the left ventricle (Supplementary File 1, Figures S17, S19, S21, and S23). This is supported by the fact that left ventricular end-diastolic pressure is highly correlated with right ventricular end-diastolic pressure (Unverferth et al., 1981; Imamura and Kinugawa, 2017; Cortez et al., 2017), which is also typical for ventricular end-diastolic volumes (Kraut et al., 1997; Luecke et al., 2004).

### 3.1 Limitations of the study

In the present study, we relied on the fact that *ACE I/D* genotypes affect plasma concentration and activity of ACE and calculated the value of this activity (*c*_*ACE*_) relative to the value of chymase activity (*c*_*chym*_), which was taken from the model by Hallow et al. (2014). However, *ACE I/D* polymorphisms may also influence the circulating quantity of chymase (Hristova et al., 2019). This inference was not included in the model. In addition, we did not consider the inactivation of bradykinin by ACE in the model, although the inhibition of the bradykinin binding site on ACE is a promising target for developing antihypertensive drugs (Widodo et al., 2017), since this peptide is a potent endothelium-dependent vasodilator (Witherow et al., 2001).

Finally, it should be noted that we investigated the influence of only one of the polymorphic loci that affect the main pathophysiological systems of blood pressure regulation in humans. Due to the polygenic nature of arterial hypertension, the development of this disease is difficult to interpret for individual SNPs. However, our research is a very important step towards study of the multiple genetic factors that create a systemic environment for the occurrence of hypertension.

## 4 Materials and Methods

### 4.1 Mathematical model

We used a computational model of the human cardiovascular and renal systems developed by Kutumova et al. (2021) and available in the BioModels database (Malik-Sheriff et al., 2020) with ID MODEL22021600011^1^. The model is hybrid, i.e., discrete-continuous (Stéphanou and Volpert, 2016), and consists of a system of ordinary differential equations that includes several discrete events corresponding to instantaneous changes in the simulated dynamics (for example, the transition from systole to diastole). Lists of functions, equations, parameters, and variables of the model are given in Supplementary File 2, Tables S1-S4, respectively. The rationale for all formulas and changes made to the baseline models is provided in a supplementary file to our primary study (Kutumova et al., 2021).

### 4.2 Equilibrium states

We considered a model to be in equilibrium if all its variables either did not change (e.g., systolic/diastolic blood pressure) or oscillated steadily with an amplitude equal to the duration of the cardiac cycle (e.g., systemic arterial pressure).

### 4.3 Numerical solution of the model

The model covers significantly different time scales from fractions of a second (heart work) to days and weeks (regulation of water and sodium balance). Therefore, we used an agent-based co-simulation methodology (Hernández et al., 2009) to speed up numerical calculations. Briefly, we divided the model equations and events into two sets (modules) representing the functioning of the cardiovascular system (with units in seconds) and the renal system (with units in minutes). In our interpretation, the combination of a module and a suitable numerical solver was an agent. The agents were simulated independently for a given time (several minutes per time step), and then exchanged the values of common variables. Since the cardiovascular agent quickly reached equilibrium after each exchange, the acceleration of calculations was achieved by stopping its subsequent simulation until the next exchange. To simulate agents, we used a version of the CVODE solver (Hindmarsh et al., 2005) ported to Java and adapted to the BioUML software interface. For more information, see our agent-based modelling approach (Kutumova et al., 2021).

### 4.4 Simulation of antihypertensive treatment

For computational experiments in the current research, we used RAAS blockers with different mechanisms of action, including the direct renin inhibitor aliskiren (300 mg), the ACE inhibitor enalapril (20 mg), and the angiotensin II receptor blocker losartan (100 mg). In addition, we considered the following representatives of other classes of antihypertensive medications to analyze their combined action with RAAS inhibitors: the β-blocker bisoprolol (5 mg), the calcium channel blocker amlodipine (5 mg), and the thiazide diuretic hydrochlorothiazide (12.5 mg). An extension of the primary model to account for the effects of these drugs was described previously (Kutumova et al., 2022). Briefly, we identified target variables in the model for all listed medications. The pharmacodynamic effects of therapy were determined by multiplying each variable υ by the influence function:

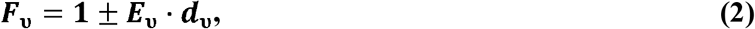

where *E*_*υ*_ is the amount of target stimulation (plus sign) or target inhibition (minus sign), and *d* _*υ*_ is the drug indicator of 1 or 0 depending on whether treatment with this drug is simulated or not. In the case of monotherapy, only one indicator in the model was equal to 1, while in combination therapy, several indicators took on non-zero values.

The pharmacodynamic functions of aliskiren, enalapril, losartan, and amlodipine were taken unchanged from the study by Kutumova et al. (2022). We also retained the hydrochlorothiazide targets in the model, but re-estimated the kinetic parameters in the influence functions to better fit clinical trials (MacKay et al., 1996). The pharmacodynamic model of bisoprolol was improved. For this drug, we previously considered a negative chronotropic effect due to blocking the access of catecholamines to their receptors (Frishman, 2003), and suppression of renal secretion of renin (Laragh and Sealey, 2011). In this study, we additionally took into account the facts that bisoprolol has a negative inotropic effect (Bazroon and Alrashidi, 2022), and reduces arterial stiffness as demonstrated, in particular, by a decrease in pulse wave velocity (Asmar et al., 1991; Kahonen et al., 2000; Palmieri et al., 2004; Ong et al., 2011; Zhou et al., 2013; Eguchi et al., 2015). A complete list of target points for all antihypertensive drugs, including actual *E* _*υ*_-values used in the formula (2), is provided in the Supplementary File 2, Table S5. An extended model comprising all pharmacodynamic functions is available online (see Data Availability).

The effect of each individual drug on the RAAS parameters corresponded to the following pharmacological properties.

- Direct renin inhibitors (aliskiren) provide a reduction in PRA, Ang I and Ang II (Shafiq et al., 2008).
- ACE inhibitors (enalapril) lead to an increase in PRA and Ang I, while Ang II decreases (Brown and Vaughan, 1998).
- Angiotensin II receptor blockers (losartan) stimulate the growth of PRA, Ang I and Ang II (Shafiq et al., 2008).
- RAAS inhibitors (aliskiren, enalapril, losartan) can lead to a decrease in plasma aldosterone levels (McInnes, 2007).
- Calcium antagonists (amlodipine) do not affect the RAAS parameters (Shafiq et al., 2008; Licata et al., 1993; Higashi et al., 1998; Bokuda et al., 2018).
- β-blockers (bisoprolol) cause a decrease in PRA, Ang I, Ang II, and aldosterone (Campbell, 2009; Savvatis et al., 2010).
- Thiazide diuretics induce activation of the RAAS (Tamargo et al., 2014), increasing PRA, Ang I and Ang II (Savvatis et al., 2010), while aldosterone levels can rise (Lijnen et al., 1981) or do not change significantly (Morales et al., 2015).

### 4.5 Virtual patient

We defined a virtual patient as a single equilibrium parameterization of the model within physiological ranges. The problem of generating a virtual patient can be formulated as minimizing the function of normalized distances between the desired 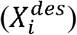 and simulated equilibrium (*X*_i_) values of the model variables (blood pressure, heart rate, etc.):

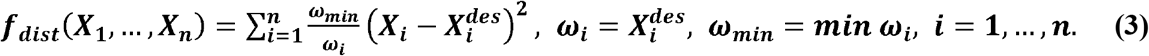

A list of fitting parameters, including the limits of the search space in hypertensive and normotensive cases, is presented in Supplementary File 2, Table S6. In addition, we took into account a set of physiological constraints 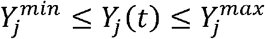 for *j* = 1, …, *m*. which were imposed on the model variables (Supplementary File 2, Table S7), and determined the penalty function of the optimization problem:

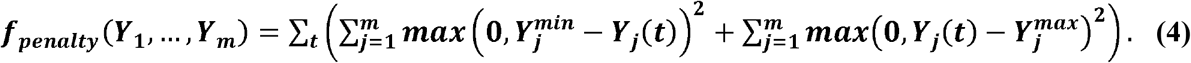

To solve such an optimization problem, we used a stochastic ranking evolution strategy (Runarsson and Yao, 2000) suitable for constrained global optimization. To accelerate the calculations, we optimized each of the model agents separately, and then combined the found equilibrium states.

In order to cut off decisions with unrealistic physiological behavior, we tested each virtual patient for resistance to an increase in sodium load. The computational experiment consisted of instantly changing the sodium intake (parameter Φ_*sodin*_ in the model) from the patient”s normal value in the range of 0.028 – 0.209 mEq/min to the elevated value of 0.243 mEq/min (He et al., 2001), and subsequently observing the dynamics of the model variables. In the case of convergence to a new equilibrium state with SBP increased by no more than 25 mmHg and slightly elevated pulmonary venous pressure, we included the virtual patient in further analysis. Other patients who demonstrated an acute increase in pulmonary venous pressure due to accumulation of blood in the pulmonary veins and, as a result, pulmonary edema (Burkhoff and Tyberg, 1993), were not taken into consideration.

### 4.6 Virtual hypertensive population

To generate a population of unique virtual hypertensive patients with varying values of SBP, DBP, heart rate (HR), stroke volume (SV), body weight (BW), and body mass index (BMI), we used a function that produces random numbers with the following means and standard deviations:

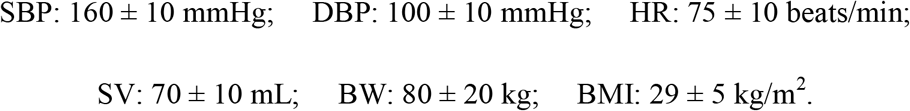

Body height (BH) in centimeters was calculated as

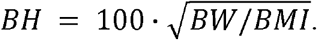

We considered the following inclusion criteria: essential hypertension (SBP ≥ 140 mmHg and/or DBP ≥ 90 mmHg; SBP > 130 mmHg; DBP > 80 mmHg), BMI > 22 kg/m^2^, and average height 160 – 180 cm. Exclusion criteria: severe hypertension (SBP > 179.5 mmHg or DBP > 109.5 mmHg), abnormal heart rate (HR < 60 or HR > 90 beats/min), and severe obesity (BMI > 36 kg/m^2^).

The population generation algorithm included the following steps:

1. Randomly generate SBP, DBP, HR, SV, BW, and BMI values using means and standard deviations within the inclusion and exclusion criteria above; select the gender of the patient (male or female) with a probability of ½.
2. Substitute the value of BW into the model. Use BW, BH, and gender to estimate ranges for baseline total body water (line 50 in Supplementary File 2, Table S6) and total blood volume (line 43 in Supplementary File 2, Table S7).
3. Generate a virtual patient by solving an optimization problem with fitting parameters from Supplementary File 2, Table S6; objective function (3) defined for the selected values of SBP, DBP, HR, and SV; and penalty function (4) determined by constraints from Supplementary File 2, Table S7.
4. If a solution to the optimization problem is found, check it with the sodium loading test and include it in the final population if the test is successful.

### 4.7 Modeling platform

Here, we used the BioUML software (https://sirius-web.org/bioumlweb/; https://ict.biouml.org/), an open-source Java-based integrated environment for systems biology (Kolpakov et al., 2019; Kolpakov et al., 2022) that supports a wide range of bioinformatic formats and mathematical methods for analyzing computational models of biological systems, including visual modeling tools.

## 6 Author Contributions

Elena Kutumova: Methodology, Software, Validation, Formal analysis, Investigation, Data curation, Writing – original draft preparation, Visualisation;

Anna Kovaleva: Writing – review & editing;

Ruslan Sharipov: Writing – review & editing;

Galina Lifshits: Writing – review & editing, Supervision;

Fedor Kolpakov: Methodology, Software, Resources, Writing – review & editing, Supervision, Project administration, Funding acquisition.

## 8 Supplementary Material

*Supplementary File 1*. Analysis of the model.

*Supplementary File 2*. Description of the model.

## 9 Data Availability

The data presented in this study can be found in the online repository at https://gitlab.sirius-web.org/virtual-patient/ace-polymorphism-in-cardiorenal-modeling.

## Footnotes

1 https://www.ebi.ac.uk/biomodels/MODEL2202160001

