## Supplementary File 1 for "Mathematical modeling of the influence of *ACE I/D* polymorphism on blood pressure and antihypertensive therapy"

*Supplementary File 1 – Model Analysis*

### Mathematical modeling of the influence of *ACE I/D* polymorphism on blood pressure and antihypertensive therapy

**Elena Kutumova*, Anna Kovaleva, Ruslan Sharipov, Galina Lifshits, Fedor Kolpakov**

**Figure S1.** Distribution of general parameters in baseline virtual hypertensive populations ($n$ = 100)

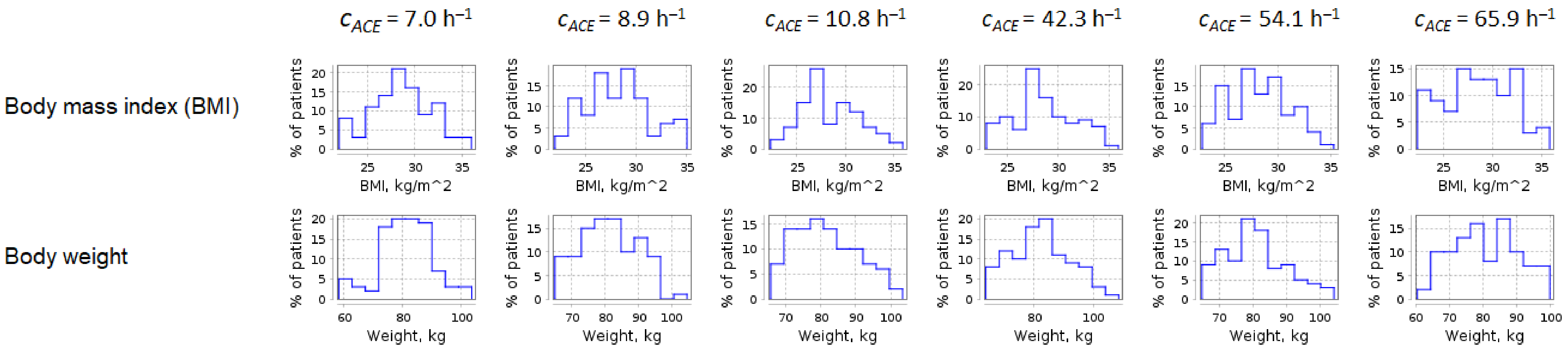

**Figure S2.** Distribution of systemic hemodynamic parameters in baseline virtual hypertensive populations ($n$ = 100)

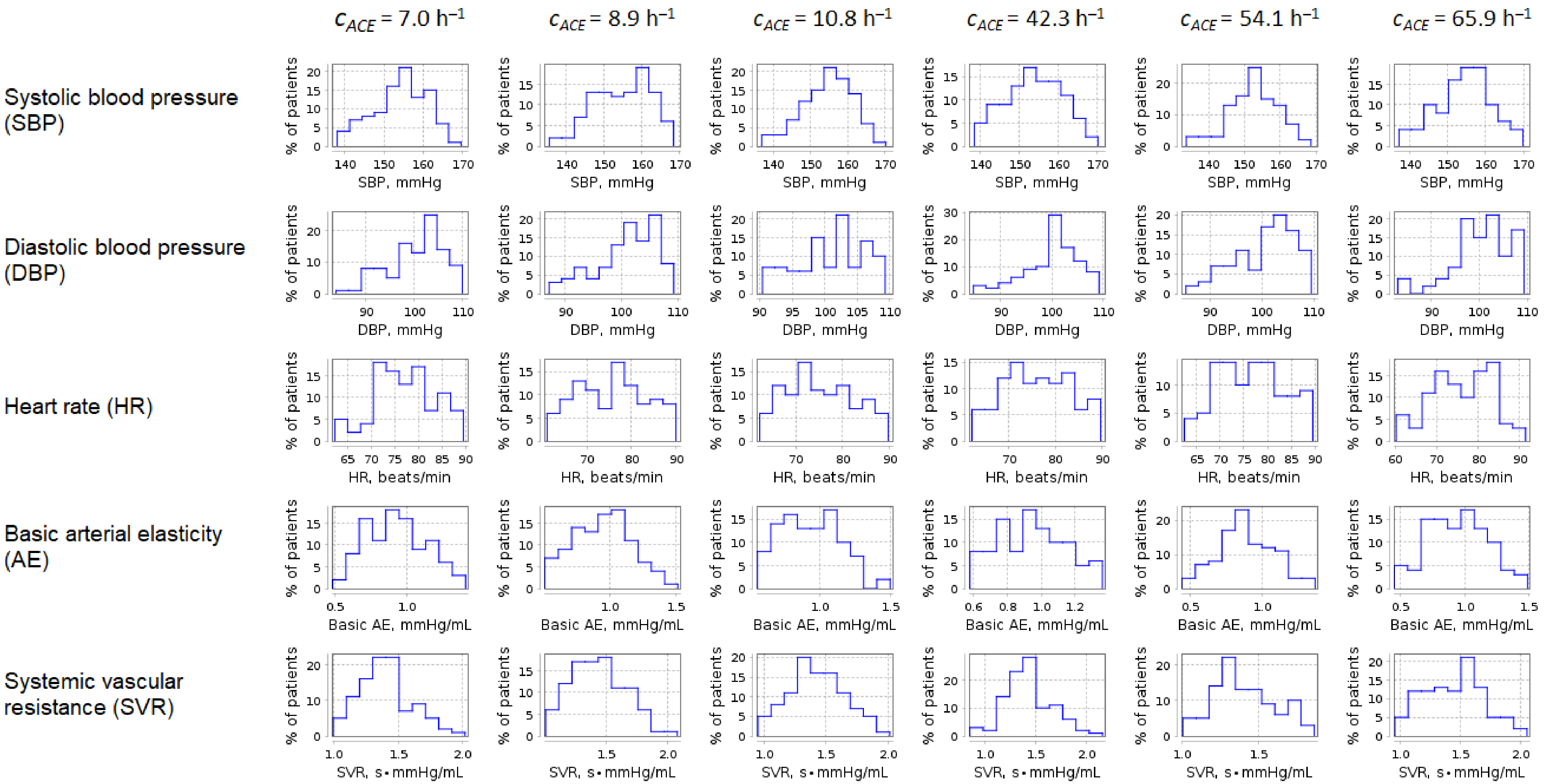

**Figure S3.** Distribution of pulmonary hemodynamic parameters in baseline virtual hypertensive populations ($n$ = 100)

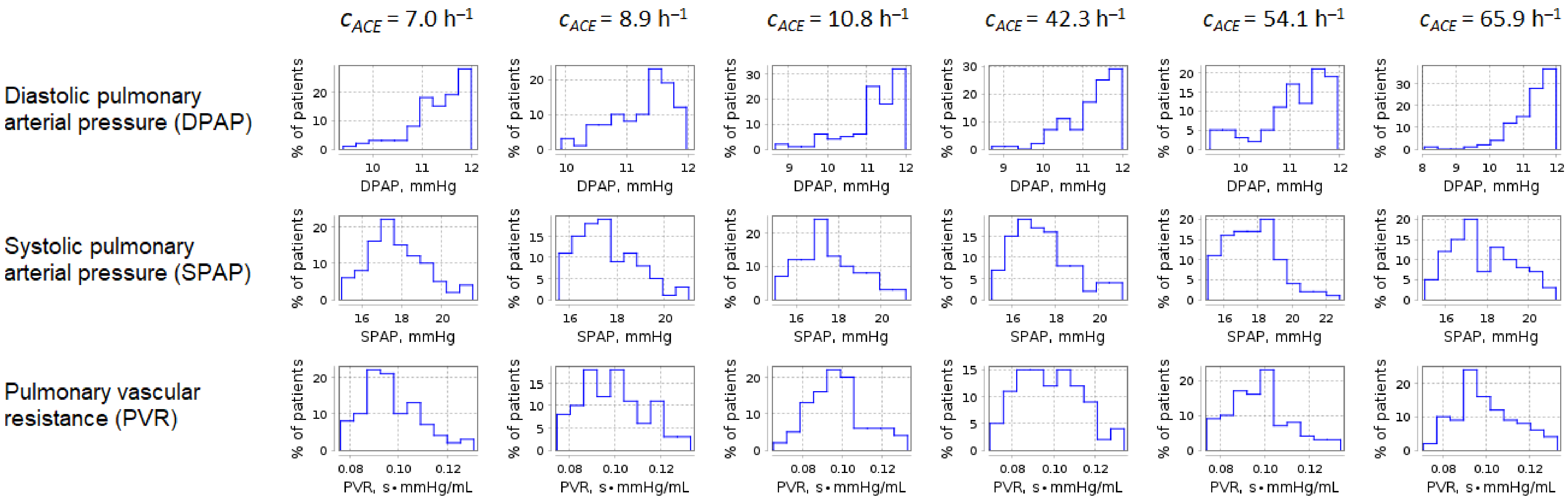

**Figure S4.** Distribution of left ventricular parameters in baseline virtual hypertensive populations ($n$ = 100)

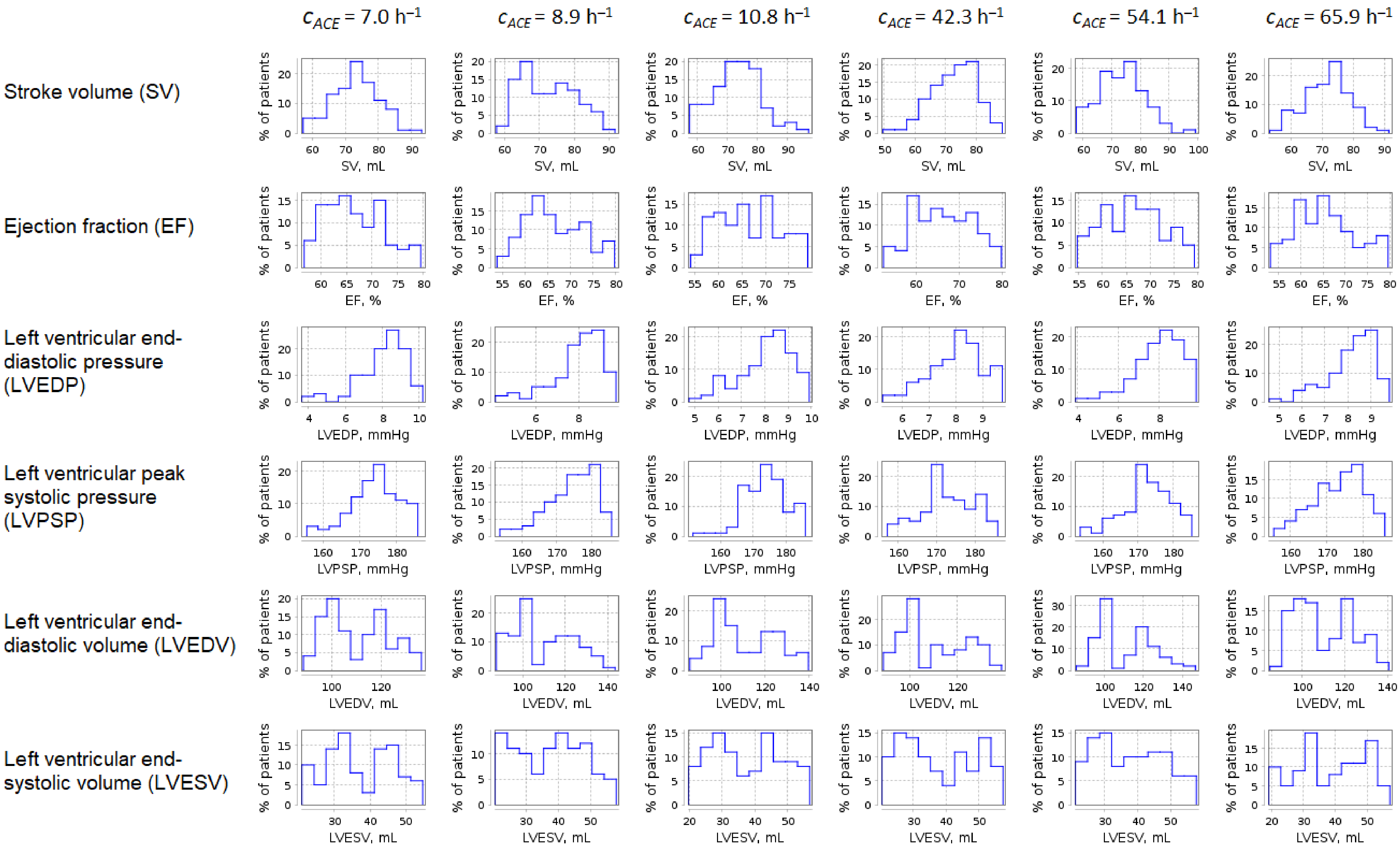

**Figure S5.** Distribution of right ventricular parameters in baseline virtual hypertensive populations ($n$ = 100)

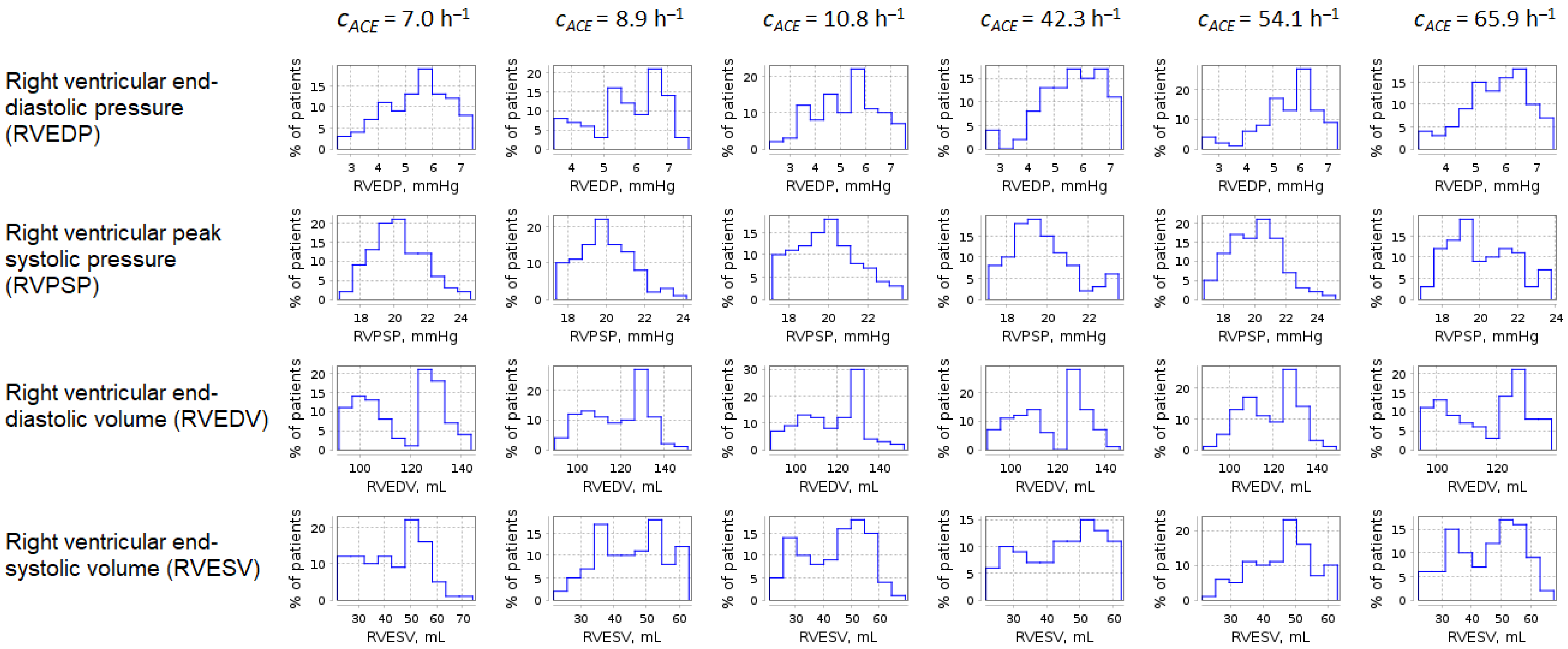

**Figure S6.** Distribution of renal function parameters in baseline virtual hypertensive populations ($n$ = 100)

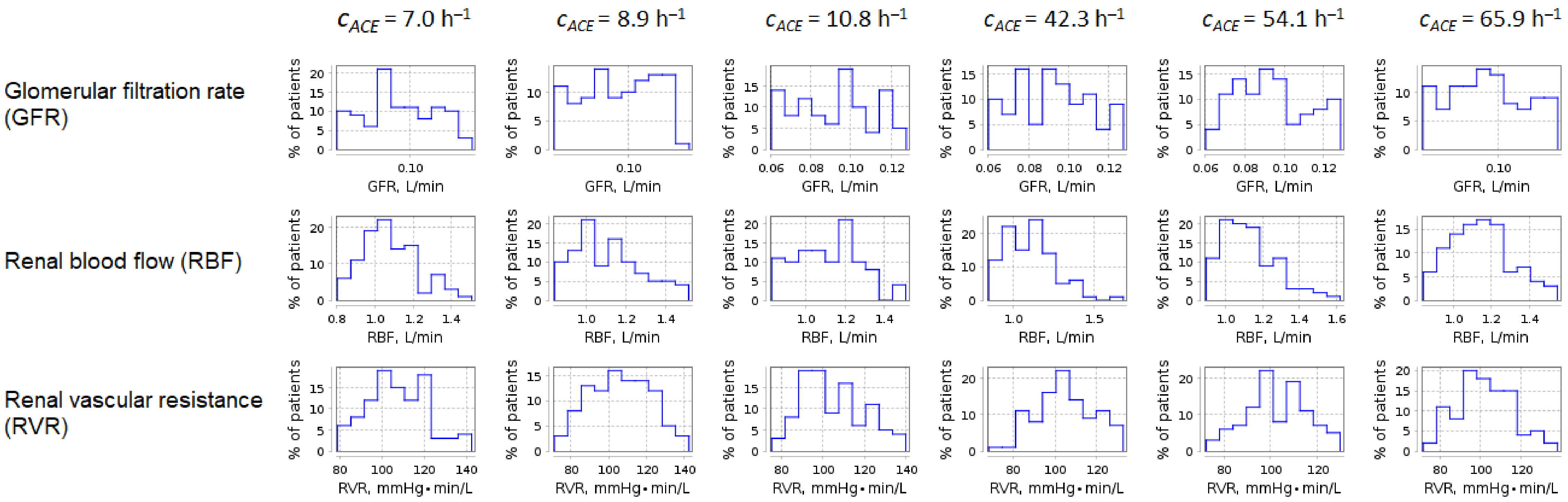

**Figure S7.** Distribution of intraglomerular hemodynamic parameters in baseline virtual hypertensive populations ($n$ = 100)

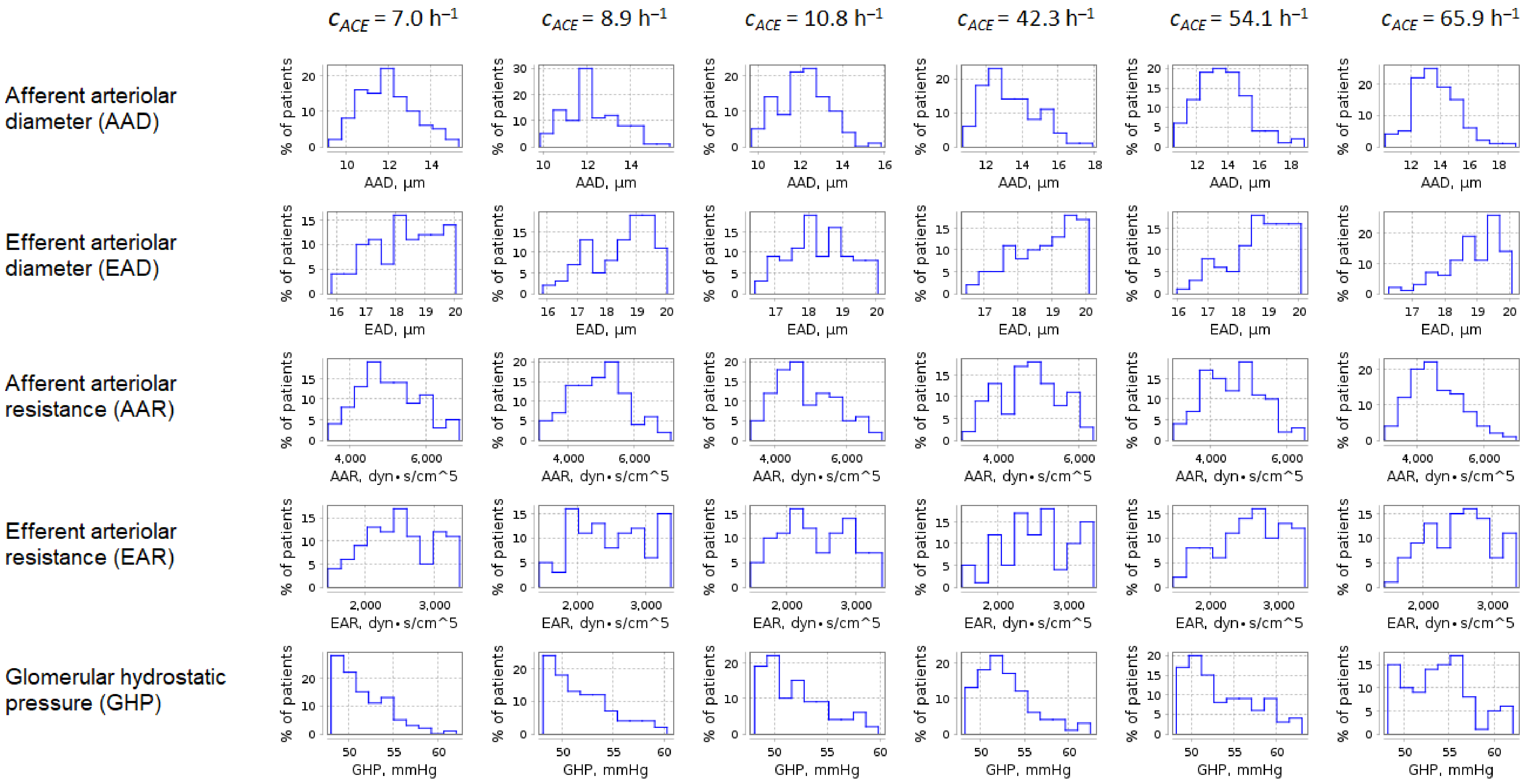

**Figure S8.** Distribution of blood parameters in baseline virtual hypertensive populations ($n$ = 100)

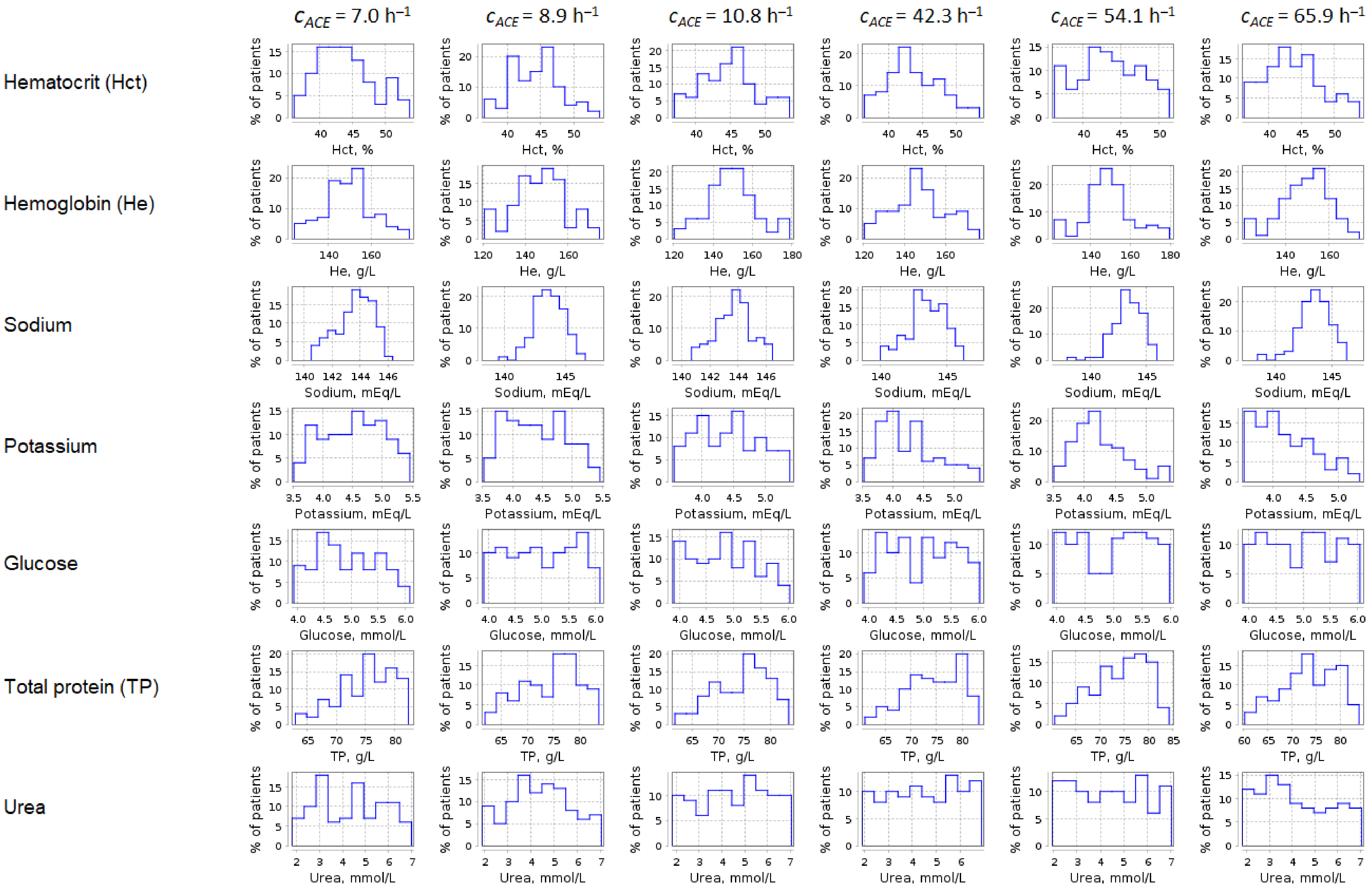

**Figure S9.** Distribution of renin-angiotensin-aldosterone system parameters in baseline virtual hypertensive populations ($n$ = 100)

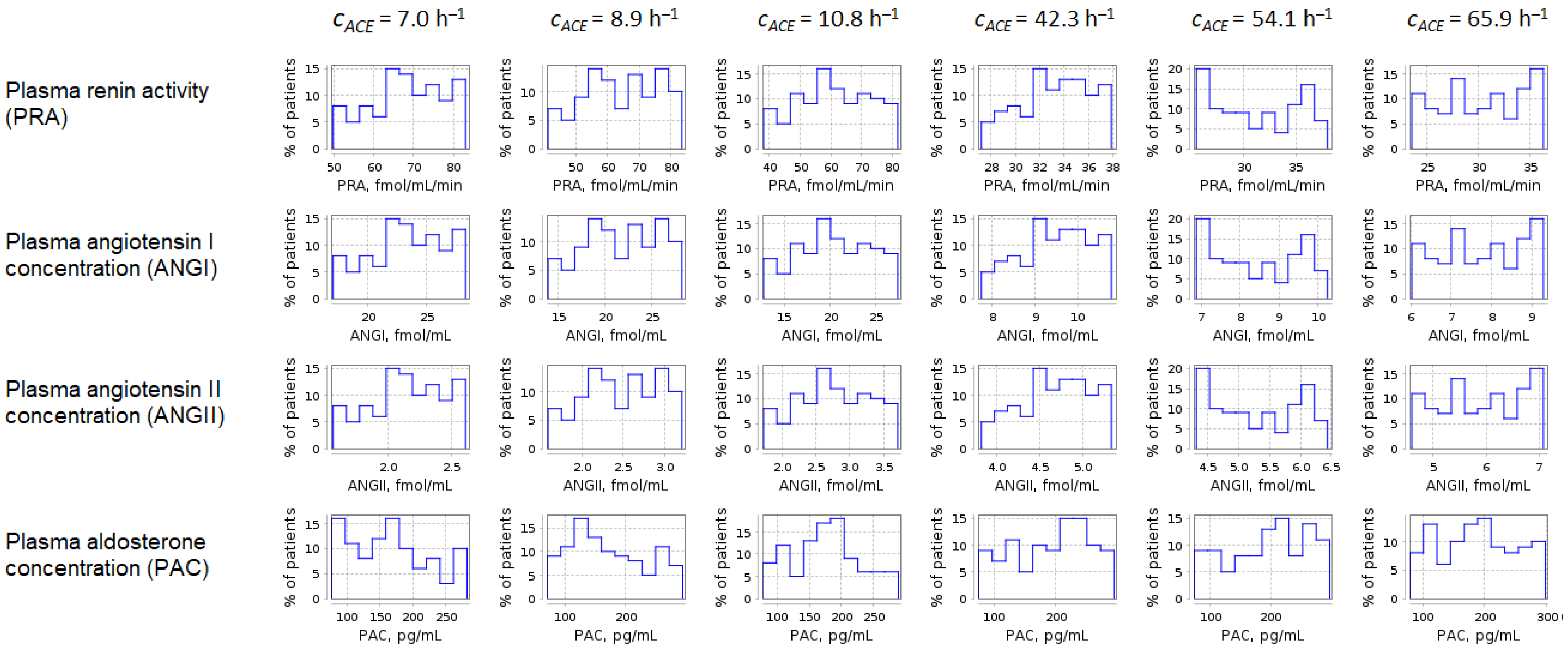

**Table S1.** *P*-values calculated using the Kolmogorov-Smirnov test for different cases of ACE activity: 7.0 $h^{-1}$ (population 1), 8.9 $h^{-1}$ (population 2), 10.8 $h^{-1}$ (population 3), 42.3 $h^{-1}$ (population 4), 54.1 $h^{-1}$ (population 5), and 65.9 $h^{-1}$ (population 6); *P*-value for population *i vs*. population *j* is denoted $P_{ij}$. Size of populations: $n$ = 100.

| **Characteristics** | **Low ACE activity *vs.* low ACE activity** | | | **Low ACE activity *vs.* high ACE activity** | | | | | | | | | **High ACE activity *vs.* high ACE activity** | | |
|---|---|---|---|---|---|---|---|---|---|---|---|---|---|---|---|
| | $P_{12}$ | $P_{13}$ | $P_{23}$ | $P_{14}$ | $P_{15}$ | $P_{16}$ | $P_{24}$ | $P_{25}$ | $P_{26}$ | $P_{34}$ | $P_{35}$ | $P_{36}$ | $P_{45}$ | $P_{46}$ | $P_{56}$ |
| Body mass index | 0.281 | 0.211 | 0.699 | 0.813 | 0.367 | 0.813 | 0.211 | 0.813 | 0.367 | 0.367 | 0.468 | 0.367 | 0.155 | 0.699 | 0.367 |
| Body weight | 0.581 | 0.813 | 0.994 | 0.468 | 0.581 | 0.367 | 0.468 | 0.967 | 0.813 | 0.581 | 0.699 | 0.813 | 0.367 | 0.367 | 0.468 |
| Systolic blood pressure | 0.367 | 0.967 | 0.367 | 0.813 | 0.111 | 1.000 | 0.468 | 0.024 | 0.367 | 0.813 | 0.054 | 0.699 | 0.468 | 0.906 | 0.111 |
| Diastolic blood pressure | 0.581 | 0.813 | 0.967 | 0.367 | 0.906 | 0.813 | 0.155 | 0.813 | 0.581 | 0.281 | 0.906 | 0.906 | 0.281 | 0.581 | 0.699 |
| Heart rate | 0.024 | 0.037 | 0.699 | 0.111 | 0.281 | 0.281 | 0.813 | 0.468 | 0.468 | 0.699 | 0.468 | 0.468 | 0.906 | 0.967 | 0.699 |
| Basic systemic arterial elasticity | 0.699 | 0.906 | 0.813 | 0.699 | 0.468 | 0.468 | 0.967 | 0.078 | 0.906 | 0.813 | 0.367 | 0.906 | 0.078 | 0.699 | 0.037 |
| Systemic vascular resistance | 0.211 | 0.281 | 0.813 | 0.581 | 0.906 | 0.006 | 0.906 | 0.468 | 0.281 | 0.813 | 0.699 | 0.211 | 0.468 | 0.078 | 0.078 |
| Diastolic pulmonary arterial pressure | 0.468 | 0.813 | 0.367 | 0.367 | 0.211 | 0.581 | 0.581 | 0.155 | 0.699 | 0.967 | 0.281 | 0.699 | 0.468 | 0.967 | 0.111 |
| Systolic pulmonary arterial pressure | 0.699 | 0.581 | 0.906 | 0.037 | 0.906 | 0.813 | 0.468 | 0.367 | 0.281 | 0.468 | 0.699 | 0.281 | 0.111 | 0.024 | 0.906 |
| Pulmonary vascular resistance | 0.281 | 0.906 | 0.468 | 0.155 | 0.813 | 0.367 | 0.699 | 0.699 | 0.967 | 0.367 | 0.813 | 0.699 | 0.281 | 0.211 | 0.581 |
| Stroke volume | 0.155 | 0.581 | 0.211 | 0.699 | 0.111 | 0.468 | 0.367 | 0.367 | 0.468 | 0.994 | 0.813 | 0.281 | 0.699 | 0.367 | 0.024 |
| Ejection fraction | 0.813 | 0.581 | 0.813 | 0.281 | 0.468 | 0.281 | 0.581 | 0.813 | 0.581 | 0.994 | 0.967 | 0.211 | 0.813 | 0.367 | 0.281 |
| Left ventricular end-diastolic pressure | 0.699 | 0.906 | 1.000 | 0.468 | 0.813 | 0.468 | 0.699 | 0.967 | 0.468 | 0.581 | 0.906 | 0.468 | 0.906 | 0.155 | 0.211 |
| Left ventricular peak systolic pressure | 0.581 | 0.906 | 0.468 | 0.078 | 0.211 | 0.813 | 0.054 | 0.078 | 0.581 | 0.581 | 0.699 | 0.468 | 0.906 | 0.468 | 0.468 |
| Left ventricular end-diastolic volume | 0.581 | 0.813 | 0.699 | 0.967 | 0.906 | 1.000 | 0.906 | 0.468 | 0.813 | 0.994 | 0.813 | 0.813 | 0.699 | 0.906 | 0.699 |
| Left ventricular end-systolic volume | 0.468 | 0.699 | 0.813 | 0.367 | 0.813 | 0.155 | 0.581 | 0.906 | 0.581 | 0.813 | 0.906 | 0.699 | 0.813 | 0.581 | 0.699 |
| Right ventricular end-diastolic pressure | 0.054 | 0.699 | 0.006 | 0.468 | 0.281 | 0.155 | 0.581 | 0.111 | 0.211 | 0.054 | 0.037 | 0.024 | 0.813 | 0.994 | 0.906 |
| Right ventricular peak systolic pressure | 0.468 | 0.281 | 0.906 | 0.078 | 0.967 | 0.581 | 0.155 | 0.468 | 0.111 | 0.468 | 0.211 | 0.581 | 0.078 | 0.155 | 0.699 |
| Right ventricular end-diastolic volume | 0.468 | 0.581 | 0.906 | 0.906 | 0.111 | 0.699 | 0.813 | 0.581 | 0.813 | 0.967 | 0.211 | 0.581 | 0.281 | 0.906 | 0.281 |
| Right ventricular end-systolic volume | 0.155 | 0.994 | 0.468 | 0.906 | 0.155 | 0.581 | 0.367 | 0.813 | 0.699 | 0.994 | 0.367 | 0.581 | 0.367 | 0.813 | 0.468 |
| Glomerular filtration rate | 0.906 | 0.367 | 0.211 | 0.581 | 0.699 | 0.994 | 0.367 | 0.367 | 0.906 | 0.813 | 0.367 | 0.468 | 0.906 | 0.699 | 0.581 |
| Renal blood flow | 0.367 | 0.078 | 0.581 | 0.211 | 0.111 | 0.010 | 0.581 | 0.367 | 0.155 | 0.367 | 0.581 | 0.699 | 0.367 | 0.281 | 0.581 |
| Renal vascular resistance | 0.281 | 0.078 | 0.581 | 0.468 | 0.211 | 0.024 | 0.581 | 0.155 | 0.111 | 0.581 | 0.367 | 0.367 | 0.468 | 0.078 | 0.468 |
| Afferent arteriolar diameter | 0.281 | 0.367 | 0.813 | SS | SS | SS | SS | SS | SS | SS | SS | SS | 0.111 | 0.024 | 0.813 |
| Efferent arteriolar diameter | 0.111 | 0.468 | 0.054 | 0.010 | 0.016 | SS | 0.111 | 0.581 | 0.037 | 0.004 | 0.024 | SS | 0.699 | 0.281 | 0.281 |
| Afferent arteriolar resistance | 0.699 | 0.281 | 0.468 | 0.155 | 0.016 | 0.001 | 0.281 | 0.024 | 0.002 | 0.581 | 0.211 | 0.155 | 0.155 | 0.037 | 0.699 |
| Efferent arteriolar resistance | 0.813 | 0.281 | 0.699 | 0.581 | 0.367 | 0.699 | 0.155 | 0.078 | 0.468 | 0.054 | 0.016 | 0.078 | 0.906 | 0.581 | 0.699 |
| Glomerular hydrostatic pressure | 0.967 | 0.813 | 1.000 | 0.010 | 0.001 | SS | 0.037 | 0.010 | SS | 0.037 | 0.016 | SS | 0.078 | 0.010 | 0.155 |
| Hematocrit | 0.468 | 0.281 | 0.967 | 0.906 | 0.699 | 0.967 | 0.367 | 0.281 | 0.281 | 0.078 | 0.111 | 0.111 | 0.813 | 0.906 | 0.813 |
| Hemoglobin | 0.281 | 1.000 | 0.367 | 0.367 | 0.468 | 0.906 | 0.813 | 0.054 | 0.111 | 0.468 | 0.813 | 0.906 | 0.211 | 0.155 | 0.367 |
| Plasma sodium | 0.699 | 0.813 | 0.813 | 0.367 | 0.281 | 0.699 | 0.699 | 0.468 | 0.699 | 0.281 | 0.211 | 0.211 | 0.906 | 0.994 | 0.906 |
| Plasma potassium | 0.281 | 0.211 | 0.906 | SS | SS | SS | 0.054 | 0.024 | 0.024 | 0.024 | 0.010 | 0.024 | 0.699 | 0.281 | 0.155 |
| Plasma glucose | 0.468 | 0.468 | 0.155 | 0.813 | 0.581 | 0.813 | 0.967 | 0.967 | 0.967 | 0.211 | 0.211 | 0.367 | 0.967 | 0.994 | 0.967 |
| Plasma total protein | 0.367 | 0.581 | 0.967 | 0.468 | 0.367 | 0.078 | 0.906 | 1.000 | 0.367 | 0.906 | 0.994 | 0.281 | 0.906 | 0.813 | 0.699 |
| Plasma urea | 0.468 | 0.581 | 0.367 | 0.699 | 0.967 | 0.211 | 0.281 | 0.468 | 0.111 | 0.994 | 0.813 | 0.111 | 0.906 | 0.211 | 0.468 |
| Plasma renin activity | 0.024 | SS | 0.367 | SS | SS | SS | SS | SS | SS | SS | SS | SS | SS | SS | 0.078 |
| Plasma angiotensin I | 0.010 | SS | 0.281 | SS | SS | SS | SS | SS | SS | SS | SS | SS | SS | SS | SS |
| Plasma angiotensin II | SS | SS | 0.001 | SS | SS | SS | SS | SS | SS | SS | SS | SS | SS | SS | SS |
| Plasma aldosterone concentration | 0.813 | 0.281 | 0.078 | 0.001 | SS | 0.037 | 0.024 | 0.006 | 0.078 | 0.002 | 0.001 | 0.111 | 0.906 | 0.581 | 0.581 |

**SS** = statistically significant ($P$ < 0.001)

**Table S2.** Simulated response of <u>systolic blood pressure</u> to antihypertensive therapy in virtual hypertensive populations (*n* = 100) with different ACE activity, including *P*-values (Kolmogorov-Smirnov test) for endpoint *vs*. baseline; data are presented as mean ± SD in mmHg

| **Regimens** | **$c_{ACE}$ = 7.0 h$^{-1}$** | | | **$c_{ACE}$ = 8.9 h$^{-1}$** | | | **$c_{ACE}$ = 10.8 h$^{-1}$** | | | **$c_{ACE}$ = 42.3 h$^{-1}$** | | | **$c_{ACE}$ = 54.1 h$^{-1}$** | | | **$c_{ACE}$ = 65.9 h$^{-1}$** | | |
|---|---|---|---|---|---|---|---|---|---|---|---|---|---|---|---|---|---|---|
| | **Value** | **Change** | ***P*** | **Value** | **Change** | ***P*** | **Value** | **Change** | ***P*** | **Value** | **Change** | ***P*** | **Value** | **Change** | ***P*** | **Value** | **Change** | ***P*** |
| Baseline | 154.1 ± 6.9 | – | – | 154.9 ± 7.4 | – | – | 154.5 ± 6.5 | – | – | 153.7 ± 7.2 | – | – | 152.5 ± 6.8 | – | – | 154.2 ± 7.2 | – | – |
| Al300 | 144.2 ± 6.7 | -9.8 ± 2.3 | SS | 144.5 ± 7.6 | -10.4 ± 2.3 | SS | 143.7 ± 6.9 | -10.8 ± 2.2 | SS | 142.4 ± 7.7 | -11.4 ± 1.9 | SS | 140.8 ± 6.9 | -11.7 ± 1.7 | SS | 142.8 ± 7.6 | -11.3 ± 2.1 | SS |
| E20 | 146.5 ± 6.6 | -7.6 ± 1.6 | SS | 146.3 ± 7.5 | -8.6 ± 1.8 | SS | 145.1 ± 6.8 | -9.5 ± 1.8 | SS | 139.3 ± 7.9 | -14.5 ± 2.5 | SS | 136.6 ± 7.0 | -15.9 ± 2.5 | SS | 137.9 ± 7.8 | -16.3 ± 3.1 | SS |
| L100 | 143.3 ± 6.8 | -10.7 ± 2.6 | SS | 143.5 ± 7.8 | -11.4 ± 2.6 | SS | 142.7 ± 7.0 | -11.9 ± 2.5 | SS | 140.9 ± 7.8 | -12.8 ± 2.2 | SS | 139.2 ± 7.0 | -13.3 ± 2.0 | SS | 141.2 ± 7.7 | -13.0 ± 2.4 | SS |
| Aml5 | 143.6 ± 6.6 | -10.4 ± 1.2 | SS | 144.4 ± 7.2 | -10.5 ± 1.4 | SS | 144.3 ± 6.3 | -10.2 ± 1.7 | SS | 143.8 ± 7.2 | -9.9 ± 2.0 | SS | 142.8 ± 6.1 | -9.7 ± 1.9 | SS | 144.5 ± 6.9 | -9.7 ± 2.0 | SS |
| B5 | 142.5 ± 7.0 | -11.5 ± 2.5 | SS | 143.0 ± 7.9 | -11.9 ± 2.5 | SS | 142.3 ± 6.9 | -12.2 ± 2.3 | SS | 141.4 ± 8.1 | -12.3 ± 2.5 | SS | 139.6 ± 7.1 | -12.9 ± 2.1 | SS | 142.0 ± 8.2 | -12.2 ± 2.9 | SS |
| H12.5 | 143.4 ± 8.0 | -10.7 ± 5.4 | SS | 144.4 ± 8.9 | -10.4 ± 5.1 | SS | 144.0 ± 8.5 | -10.5 ± 6.0 | SS | 144.3 ± 9.6 | -9.5 ± 6.0 | SS | 143.6 ± 8.0 | -8.9 ± 5.6 | SS | 145.4 ± 8.1 | -8.8 ± 5.8 | SS |
| Al300<br>Aml5 | 132.2 ± 6.6 | -21.9 ± 2.9 | SS | 132.5 ± 7.8 | -22.4 ± 3.0 | SS | 131.9 ± 6.9 | -22.6 ± 2.9 | SS | 130.8 ± 7.7 | -23.0 ± 2.6 | SS | 129.4 ± 6.5 | -23.1 ± 2.6 | SS | 131.5 ± 7.5 | -22.7 ± 3.0 | SS |
| Al300<br>B5 | 137.5 ± 9.0 | -16.6 ± 5.4 | SS | 137.0 ± 9.8 | -17.9 ± 5.5 | SS | 135.2 ± 8.8 | -19.3 ± 5.3 | SS | 130.0 ± 9.2 | -23.7 ± 4.9 | SS | 126.7 ± 8.0 | -25.8 ± 4.7 | SS | 127.8 ± 9.1 | -26.4 ± 5.8 | SS |
| Al300<br>H12.5 | 131.3 ± 7.7 | -22.8 ± 5.9 | SS | 131.9 ± 9.0 | -23.0 ± 5.5 | SS | 131.2 ± 9.0 | -23.4 ± 6.7 | SS | 131.3 ± 10.1 | -22.4 ± 6.5 | SS | 130.6 ± 8.2 | -21.9 ± 5.9 | SS | 133.2 ± 8.6 | -21.0 ± 6.2 | SS |
| E20<br>Aml5 | 135.0 ± 6.5 | -19.0 ± 2.2 | SS | 134.7 ± 7.6 | -20.2 ± 2.5 | SS | 133.5 ± 6.8 | -21.0 ± 2.6 | SS | 127.0 ± 7.9 | -26.8 ± 3.2 | SS | 124.4 ± 6.7 | -28.1 ± 3.3 | SS | 125.4 ± 7.6 | -28.7 ± 3.9 | SS |
| E20<br>B5 | 138.1 ± 8.1 | -15.9 ± 4.4 | SS | 137.5 ± 9.2 | -17.4 ± 4.8 | SS | 135.8 ± 8.4 | -18.8 ± 4.7 | SS | 128.0 ± 9.8 | -25.8 ± 5.8 | SS | 123.5 ± 8.7 | -29.0 ± 5.9 | SS | 123.7 ± 10.1 | -30.5 ± 7.3 | SS |
| E20<br>H12.5 | 134.4 ± 7.7 | -19.7 ± 5.6 | SS | 134.2 ± 8.9 | -20.7 ± 5.3 | SS | 132.9 ± 8.9 | -21.7 ± 6.5 | SS | 127.4 ± 10.2 | -26.3 ± 6.7 | SS | 125.5 ± 8.2 | -27.0 ± 6.1 | SS | 127.0 ± 8.7 | -27.1 ± 6.5 | SS |
| L100<br>Aml5 | 131.1 ± 6.8 | -23.0 ± 3.2 | SS | 131.2 ± 7.9 | -23.7 ± 3.4 | SS | 130.6 ± 7.1 | -24.0 ± 3.2 | SS | 129.0 ± 7.8 | -24.7 ± 2.8 | SS | 127.6 ± 6.6 | -24.9 ± 2.8 | SS | 129.5 ± 7.5 | -24.7 ± 3.2 | SS |
| L100<br>B5 | 137.2 ± 9.5 | -16.8 ± 6.0 | SS | 136.7 ± 10.3 | -18.2 ± 6.1 | SS | 134.8 ± 9.2 | -19.7 ± 5.7 | SS | 129.0 ± 9.4 | -24.7 ± 5.3 | SS | 125.5 ± 8.2 | -27.0 ± 5.1 | SS | 126.3 ± 9.4 | -27.8 ± 6.3 | SS |
| L100<br>H12.5 | 130.1 ± 7.8 | -24.0 ± 6.0 | SS | 130.5 ± 9.1 | -24.4 ± 5.6 | SS | 129.7 ± 9.1 | -24.8 ± 6.8 | SS | 129.5 ± 10.1 | -24.2 ± 6.6 | SS | 128.7 ± 8.2 | -23.8 ± 6.0 | SS | 131.2 ± 8.7 | -23.0 ± 6.3 | SS |
| Al300<br>Aml5/B5 | 122.4 ± 9.0 | -31.6 ± 6.0 | SS | 121.9 ± 10.1 | -33.0 ± 6.3 | SS | 120.3 ± 9.0 | -34.2 ± 5.9 | SS | 115.2 ± 8.9 | -38.5 ± 5.2 | SS | 112.3 ± 7.6 | -40.2 ± 5.2 | SS | 113.1 ± 8.9 | -41.1 ± 6.5 | SS |
| Al300<br>Aml5/H12.5 | 118.3 ± 7.9 | -35.8 ± 6.7 | SS | 118.8 ± 9.6 | -36.1 ± 6.7 | SS | 118.3 ± 9.7 | -36.3 ± 7.8 | SS | 118.7 ± 10.6 | -35.0 ± 7.7 | SS | 118.4 ± 8.8 | -34.1 ± 7.4 | SS | 120.9 ± 9.0 | -33.3 ± 7.5 | SS |
| Al300<br>B5/H12.5 | 119.4 ± 8.9 | -34.7 ± 7.7 | SS | 119.0 ± 10.1 | -35.9 ± 7.1 | SS | 117.5 ± 10.1 | -37.0 ± 8.3 | SS | 114.0 ± 10.1 | -39.7 ± 7.2 | SS | 111.9 ± 7.9 | -40.6 ± 6.5 | SS | 113.1 ± 9.0 | -41.1 ± 7.7 | SS |
| E20<br>Aml5/B5 | 123.9 ± 8.0 | -30.1 ± 5.0 | SS | 123.1 ± 9.5 | -31.8 ± 5.6 | SS | 121.3 ± 8.5 | -33.2 ± 5.4 | SS | 112.4 ± 9.4 | -41.3 ± 6.0 | SS | 108.3 ± 8.2 | -44.2 ± 6.1 | SS | 108.1 ± 9.7 | -46.1 ± 7.8 | SS |
| E20<br>Aml5/H12.5 | 122.0 ± 7.9 | -32.1 ± 6.5 | SS | 121.6 ± 9.5 | -33.3 ± 6.5 | SS | 120.3 ± 9.6 | -34.2 ± 7.7 | SS | 114.2 ± 10.4 | -39.6 ± 7.6 | SS | 112.6 ± 8.5 | -39.9 ± 7.3 | SS | 113.8 ± 8.8 | -40.4 ± 7.8 | SS |
| E20<br>B5/H12.5 | 121.1 ± 8.3 | -32.9 ± 7.1 | SS | 120.5 ± 9.7 | -34.4 ± 6.7 | SS | 118.7 ± 9.9 | -35.9 ± 8.0 | SS | 111.0 ± 10.3 | -42.7 ± 7.6 | SS | 107.8 ± 8.0 | -44.7 ± 6.9 | SS | 107.9 ± 9.4 | -46.2 ± 8.6 | SS |
| L100<br>Aml5/B5 | 121.9 ± 9.5 | -32.2 ± 6.5 | SS | 121.3 ± 10.6 | -33.6 ± 6.8 | SS | 119.5 ± 9.4 | -35.0 ± 6.3 | SS | 113.9 ± 9.1 | -39.8 ± 5.5 | SS | 110.7 ± 7.8 | -41.8 ± 5.6 | SS | 111.3 ± 9.2 | -42.9 ± 6.9 | SS |
| L100<br>Aml5/H12.5 | 116.7 ± 7.9 | -37.4 ± 6.8 | SS | 117.1 ± 9.6 | -37.8 ± 6.8 | SS | 116.5 ± 9.7 | -38.0 ± 7.9 | SS | 116.6 ± 10.5 | -37.1 ± 7.7 | SS | 116.3 ± 8.7 | -36.2 ± 7.4 | SS | 118.6 ± 8.9 | -35.6 ± 7.6 | SS |
| L100<br>B5/H12.5 | 118.7 ± 9.2 | -35.4 ± 8.0 | SS | 118.2 ± 10.4 | -36.6 ± 7.4 | SS | 116.7 ± 10.4 | -37.9 ± 8.6 | SS | 112.6 ± 10.2 | -41.1 ± 7.4 | SS | 110.3 ± 7.9 | -42.2 ± 6.7 | SS | 111.3 ± 9.1 | -42.9 ± 8.0 | SS |

**Al300 =** aliskiren 300 mg; **Aml5** = amlodipine 5 mg; **B5** = bisoprolol 5 mg; **E20** = enalapril 20 mg; **H12.5** = hydrochlorothiazide 12.5 mg; **L100** = losartan 100 mg; **SS** = statistically significant ($P$ < 0.00001)

**Table S3.** *P*-values calculated using the Kolmogorov-Smirnov test for changes in systolic blood pressure in populations ($n$ = 100) with different ACE activity receiving the same regimens (case 1: $c_{ACE}$ = 7.0 $h^{-1}$, case 2: $c_{ACE}$ = 8.9 $h^{-1}$, case 3: $c_{ACE}$ = 10.8 $h^{-1}$, case 4: $c_{ACE}$ = 42.3 $h^{-1}$, case 5: $c_{ACE}$ = 54.1 $h^{-1}$, case 6: $c_{ACE}$ = 65.9 $h^{-1}$; *P*-value for case *i* *vs*. case *j* is denoted $P_{ij}$)

| **Regimens** | $P_{12}$ | $P_{13}$ | $P_{23}$ | $P_{14}$ | $P_{15}$ | $P_{16}$ | $P_{24}$ | $P_{25}$ | $P_{26}$ | $P_{34}$ | $P_{35}$ | $P_{36}$ | $P_{45}$ | $P_{46}$ | $P_{56}$ |
|---|---|---|---|---|---|---|---|---|---|---|---|---|---|---|---|
| Al300 | 0.21055 | 0.01581 | 0.28093 | 0.00002 | SS | SS | 0.00232 | 0.00007 | 0.00386 | 0.11113 | 0.00232 | 0.07832 | 0.21055 | 0.69937 | 0.21055 |
| E20 | 0.00079 | SS | 0.03663 | SS | SS | SS | SS | SS | SS | SS | SS | SS | 0.00079 | 0.00013 | 0.07832 |
| L100 | 0.11113 | 0.00630 | 0.36672 | SS | SS | SS | 0.00045 | SS | 0.00007 | 0.02431 | 0.00025 | 0.01008 | 0.15454 | 0.69937 | 0.36672 |
| Aml5 | 0.81275 | 0.46756 | 0.69937 | 0.11113 | 0.00136 | 0.00232 | 0.07832 | 0.00045 | 0.00386 | 0.36672 | 0.00386 | 0.03663 | 0.11113 | 0.15454 | 0.96707 |
| B5 | 0.46756 | 0.07832 | 0.21055 | 0.07832 | 0.00232 | 0.07832 | 0.28093 | 0.05410 | 0.36672 | 0.96707 | 0.15454 | 0.81275 | 0.36672 | 0.69937 | 0.11113 |
| H12.5 | 0.58062 | 0.69937 | 0.36672 | 0.46756 | 0.15454 | 0.05410 | 0.28093 | 0.01581 | 0.00232 | 0.28093 | 0.05410 | 0.01581 | 0.36672 | 0.36672 | 0.96707 |
| Al300<br>Aml5 | 0.11113 | 0.01581 | 0.81275 | 0.00232 | 0.01581 | 0.01008 | 0.21055 | 0.46756 | 0.36672 | 0.28093 | 0.58062 | 0.90621 | 0.81275 | 0.69937 | 0.96707 |
| Al300<br>B5 | 0.21055 | 0.00045 | 0.15454 | SS | SS | SS | SS | SS | SS | 0.00004 | SS | SS | 0.00630 | 0.02431 | 0.46756 |
| Al300<br>H12.5 | 0.81275 | 0.07832 | 0.58062 | 0.81275 | 0.46756 | 0.07832 | 0.36672 | 0.11113 | 0.02431 | 0.46756 | 0.02431 | 0.00630 | 0.36672 | 0.15454 | 0.69937 |
| E20<br>Aml5 | 0.00045 | SS | 0.05410 | SS | SS | SS | SS | SS | SS | SS | SS | SS | 0.11113 | 0.00232 | 0.15454 |
| E20<br>B5 | 0.05410 | SS | 0.07832 | SS | SS | SS | SS | SS | SS | SS | SS | SS | 0.00232 | 0.00045 | 0.11113 |
| E20<br>H12.5 | 0.36672 | 0.00386 | 0.07832 | SS | SS | SS | SS | SS | SS | 0.00004 | SS | SS | 0.69937 | 0.36672 | 0.81275 |
| L100<br>Aml5 | 0.07832 | 0.00630 | 0.69937 | 0.00013 | 0.00013 | 0.00007 | 0.03663 | 0.03663 | 0.02431 | 0.07832 | 0.07832 | 0.28093 | 0.81275 | 0.90621 | 0.81275 |
| L100<br>B5 | 0.21055 | 0.00136 | 0.15454 | SS | SS | SS | SS | SS | SS | 0.00004 | SS | SS | 0.00630 | 0.01008 | 0.36672 |
| L100<br>H12.5 | 0.36672 | 0.15454 | 0.36672 | 0.46756 | 0.96707 | 0.58062 | 0.46756 | 0.21055 | 0.15454 | 0.58062 | 0.11113 | 0.07832 | 0.58062 | 0.28093 | 0.90621 |
| Al300<br>Aml5/B5 | 0.07832 | 0.00232 | 0.28093 | SS | SS | SS | SS | SS | SS | 0.00004 | SS | SS | 0.11113 | 0.00386 | 0.15454 |
| Al300<br>Aml5/H12.5 | 0.58062 | 0.58062 | 0.58062 | 0.58062 | 0.21055 | 0.11113 | 0.36672 | 0.05410 | 0.02431 | 0.21055 | 0.03663 | 0.02431 | 0.58062 | 0.15454 | 0.36672 |
| Al300<br>B5/H12.5 | 0.21055 | 0.05410 | 0.28093 | 0.00004 | SS | SS | 0.01581 | 0.00079 | 0.00013 | 0.02431 | 0.00386 | 0.01008 | 0.46756 | 0.15454 | 0.69937 |
| E20<br>Aml5/B5 | 0.01581 | 0.00002 | 0.21055 | SS | SS | SS | SS | SS | SS | SS | SS | SS | 0.01581 | 0.00045 | 0.07832 |
| E20<br>Aml5/H12.5 | 0.15454 | 0.02431 | 0.21055 | SS | SS | SS | SS | SS | SS | 0.00079 | 0.00013 | SS | 0.90621 | 0.36672 | 0.58062 |
| E20<br>B5/H12.5 | 0.07832 | 0.01581 | 0.15454 | SS | SS | SS | SS | SS | SS | 0.00002 | SS | SS | 0.21055 | 0.00232 | 0.21055 |
| L100<br>Aml5/B5 | 0.07832 | 0.00232 | 0.28093 | SS | SS | SS | SS | SS | SS | 0.00002 | SS | SS | 0.11113 | 0.00386 | 0.15454 |
| L100<br>Aml5/H12.5 | 0.46756 | 0.46756 | 0.58062 | 0.90621 | 0.28093 | 0.36672 | 0.58062 | 0.07832 | 0.11113 | 0.58062 | 0.15454 | 0.07832 | 0.69937 | 0.21055 | 0.46756 |
| L100<br>B5/H12.5 | 0.21055 | 0.02431 | 0.15454 | SS | SS | SS | 0.00232 | 0.00013 | 0.00002 | 0.01581 | 0.00079 | 0.00232 | 0.46756 | 0.05410 | 0.69937 |

**Al300 =** aliskiren 300 mg; **Aml5** = amlodipine 5 mg; **B5** = bisoprolol 5 mg; **E20** = enalapril 20 mg; **H12.5** = hydrochlorothiazide 12.5 mg; **L100** = losartan 100 mg; **SS** = statistically significant ($P < 0.00001$)

**Table S4.** Simulated response of diastolic blood pressure to antihypertensive therapy in virtual hypertensive populations ($n$ = 100) with different ACE activity, including $P$-values (Kolmogorov-Smirnov test) for endpoint *vs*. baseline; data are presented as mean ± SD in mmHg

| **Regimens** | $c_{ACE}$ **= 7.0 h**$^{-1}$ | | | $c_{ACE}$ **= 8.9 h**$^{-1}$ | | | $c_{ACE}$ **= 10.8 h**$^{-1}$ | | | $c_{ACE}$ **= 42.3 h**$^{-1}$ | | | $c_{ACE}$ **= 54.1 h**$^{-1}$ | | | $c_{ACE}$ **= 65.9 h**$^{-1}$ | | |
|---|---|---|---|---|---|---|---|---|---|---|---|---|---|---|---|---|---|---|
| | **Value** | **Change** | ***P*** | **Value** | **Change** | ***P*** | **Value** | **Change** | ***P*** | **Value** | **Change** | ***P*** | **Value** | **Change** | ***P*** | **Value** | **Change** | ***P*** |
| Baseline | 100.5 ± 5.6 | – | – | 101.0 ± 5.3 | – | – | 100.9 ± 5.1 | – | – | 100.1 ± 5.3 | – | – | 100.5 ± 5.8 | – | – | 100.5 ± 5.9 | – | – |
| Al300 | 93.4 ± 5.4 | -7.1 ± 0.7 | SS | 93.5 ± 5.1 | -7.4 ± 0.9 | SS | 93.3 ± 4.7 | -7.7 ± 0.9 | SS | 90.8 ± 5.1 | -9.3 ± 0.9 | SS | 90.4 ± 5.4 | -10.1 ± 1.1 | SS | 89.6 ± 5.8 | -10.9 ± 1.2 | SS |
| E20 | 95.1 ± 5.5 | -5.4 ± 0.6 | SS | 94.8 ± 5.1 | -6.2 ± 0.7 | SS | 94.1 ± 4.8 | -6.8 ± 0.8 | SS | 89.1 ± 5.0 | -11.0 ± 1.1 | SS | 88.2 ± 5.3 | -12.3 ± 1.4 | SS | 87.0 ± 5.8 | -13.5 ± 1.5 | SS |
| L100 | 92.6 ± 5.4 | -7.9 ± 0.8 | SS | 92.8 ± 5.1 | -8.2 ± 0.9 | SS | 92.5 ± 4.7 | -8.5 ± 0.9 | SS | 90.0 ± 5.1 | -10.1 ± 1.0 | SS | 89.5 ± 5.4 | -11.0 ± 1.2 | SS | 88.7 ± 5.8 | -11.8 ± 1.3 | SS |
| Aml5 | 92.1 ± 5.2 | -8.4 ± 0.7 | SS | 92.6 ± 5.0 | -8.3 ± 0.7 | SS | 92.7 ± 4.7 | -8.3 ± 0.7 | SS | 92.2 ± 4.9 | -7.9 ± 0.7 | SS | 92.6 ± 5.4 | -7.9 ± 0.8 | SS | 92.6 ± 5.6 | -7.9 ± 0.7 | SS |
| B5 | 94.2 ± 5.3 | -6.2 ± 0.9 | SS | 94.4 ± 5.0 | -6.6 ± 1.0 | SS | 94.0 ± 4.6 | -6.9 ± 1.1 | SS | 91.3 ± 5.0 | -8.8 ± 1.1 | SS | 91.0 ± 5.3 | -9.4 ± 1.2 | SS | 90.1 ± 5.7 | -10.4 ± 1.6 | SS |
| H12.5 | 92.0 ± 5.4 | -8.4 ± 1.5 | SS | 92.7 ± 5.2 | -8.3 ± 1.4 | SS | 92.7 ± 5.0 | -8.3 ± 1.5 | SS | 92.6 ± 5.0 | -7.5 ± 1.4 | SS | 93.5 ± 5.5 | -7.0 ± 1.5 | SS | 93.5 ± 5.6 | -7.0 ± 1.5 | SS |
| Al300<br>Aml5 | 85.2 ± 5.2 | -15.3 ± 1.0 | SS | 85.5 ± 4.9 | -15.5 ± 1.2 | SS | 85.3 ± 4.5 | -15.6 ± 1.1 | SS | 83.2 ± 4.8 | -16.9 ± 1.1 | SS | 82.8 ± 5.1 | -17.7 ± 1.3 | SS | 82.1 ± 5.7 | -18.4 ± 1.3 | SS |
| Al300<br>B5 | 86.4 ± 5.4 | -14.1 ± 1.6 | SS | 87.0 ± 5.1 | -14.0 ± 1.5 | SS | 86.9 ± 4.6 | -14.0 ± 1.5 | SS | 85.2 ± 4.9 | -14.9 ± 1.5 | SS | 84.5 ± 5.1 | -15.9 ± 1.9 | SS | 83.6 ± 5.7 | -16.9 ± 2.0 | SS |
| Al300<br>H12.5 | 85.4 ± 5.4 | -15.1 ± 1.7 | SS | 85.7 ± 5.0 | -15.3 ± 1.5 | SS | 85.3 ± 4.8 | -15.6 ± 1.7 | SS | 83.3 ± 4.9 | -16.8 ± 1.7 | SS | 83.3 ± 5.2 | -17.2 ± 1.8 | SS | 82.5 ± 5.7 | -18.0 ± 1.9 | SS |
| E20<br>Aml5 | 86.9 ± 5.2 | -13.5 ± 0.9 | SS | 86.8 ± 4.9 | -14.2 ± 1.0 | SS | 86.2 ± 4.5 | -14.8 ± 1.0 | SS | 81.4 ± 4.8 | -18.7 ± 1.4 | SS | 80.5 ± 5.1 | -20.0 ± 1.6 | SS | 79.4 ± 5.8 | -21.0 ± 1.6 | SS |
| E20<br>B5 | 89.0 ± 5.3 | -11.4 ± 1.3 | SS | 88.8 ± 5.0 | -12.2 ± 1.3 | SS | 88.1 ± 4.5 | -12.8 ± 1.4 | SS | 83.3 ± 4.9 | -16.8 ± 1.6 | SS | 82.2 ± 5.1 | -18.3 ± 2.1 | SS | 81.0 ± 5.8 | -19.4 ± 2.2 | SS |
| E20<br>H12.5 | 87.0 ± 5.4 | -13.5 ± 1.7 | SS | 86.9 ± 5.0 | -14.1 ± 1.5 | SS | 86.2 ± 4.8 | -14.8 ± 1.7 | SS | 81.7 ± 4.9 | -18.4 ± 1.9 | SS | 81.1 ± 5.1 | -19.4 ± 2.0 | SS | 79.9 ± 5.7 | -20.6 ± 2.1 | SS |
| L100<br>Aml5 | 84.5 ± 5.2 | -16.0 ± 1.0 | SS | 84.7 ± 4.9 | -16.2 ± 1.2 | SS | 84.5 ± 4.5 | -16.4 ± 1.2 | SS | 82.4 ± 4.8 | -17.7 ± 1.2 | SS | 81.9 ± 5.1 | -18.5 ± 1.4 | SS | 81.2 ± 5.7 | -19.3 ± 1.4 | SS |
| L100<br>B5 | 84.9 ± 5.4 | -15.5 ± 1.7 | SS | 85.7 ± 5.1 | -15.3 ± 1.6 | SS | 85.8 ± 4.6 | -15.2 ± 1.6 | SS | 84.3 ± 4.9 | -15.8 ± 1.5 | SS | 83.7 ± 5.1 | -16.8 ± 2.0 | SS | 82.8 ± 5.7 | -17.7 ± 2.1 | SS |
| L100<br>H12.5 | 84.7 ± 5.4 | -15.8 ± 1.8 | SS | 85.0 ± 5.0 | -16.0 ± 1.6 | SS | 84.6 ± 4.8 | -16.3 ± 1.8 | SS | 82.6 ± 4.9 | -17.5 ± 1.8 | SS | 82.5 ± 5.2 | -18.0 ± 1.9 | SS | 81.6 ± 5.7 | -18.9 ± 2.0 | SS |
| Al300<br>Aml5/B5 | 78.6 ± 5.2 | -21.8 ± 1.5 | SS | 79.2 ± 4.9 | -21.7 ± 1.6 | SS | 79.2 ± 4.5 | -21.7 ± 1.7 | SS | 77.5 ± 4.8 | -22.6 ± 1.9 | SS | 76.8 ± 5.0 | -23.7 ± 2.2 | SS | 75.9 ± 5.7 | -24.6 ± 2.2 | SS |
| Al300<br>Aml5/H12.5 | 77.6 ± 5.4 | -22.9 ± 2.1 | SS | 78.0 ± 5.0 | -23.0 ± 1.9 | SS | 77.7 ± 4.9 | -23.2 ± 2.2 | SS | 76.1 ± 4.9 | -24.0 ± 2.1 | SS | 76.1 ± 5.2 | -24.4 ± 2.2 | SS | 75.4 ± 5.7 | -25.1 ± 2.1 | SS |
| Al300<br>B5/H12.5 | 80.8 ± 5.4 | -19.7 ± 2.0 | SS | 81.2 ± 5.0 | -19.8 ± 2.0 | SS | 80.8 ± 4.8 | -20.1 ± 2.3 | SS | 78.8 ± 5.0 | -21.3 ± 2.4 | SS | 78.2 ± 5.0 | -22.3 ± 2.6 | SS | 77.3 ± 5.7 | -23.2 ± 2.7 | SS |
| E20<br>Aml5/B5 | 81.2 ± 5.1 | -19.2 ± 1.3 | SS | 81.0 ± 4.9 | -20.0 ± 1.4 | SS | 80.4 ± 4.4 | -20.6 ± 1.6 | SS | 75.6 ± 4.9 | -24.5 ± 2.1 | SS | 74.3 ± 5.1 | -26.2 ± 2.5 | SS | 73.1 ± 5.9 | -27.3 ± 2.7 | SS |
| E20<br>Aml5/H12.5 | 79.3 ± 5.3 | -21.2 ± 2.0 | SS | 79.3 ± 5.0 | -21.7 ± 1.8 | SS | 78.6 ± 4.9 | -22.3 ± 2.1 | SS | 74.3 ± 5.0 | -25.8 ± 2.4 | SS | 73.6 ± 5.2 | -26.9 ± 2.4 | SS | 72.7 ± 5.8 | -27.8 ± 2.5 | SS |
| E20<br>B5/H12.5 | 82.7 ± 5.3 | -17.7 ± 2.0 | SS | 82.5 ± 4.9 | -18.4 ± 1.9 | SS | 81.7 ± 4.7 | -19.2 ± 2.2 | SS | 77.1 ± 5.1 | -23.0 ± 2.6 | SS | 76.0 ± 5.1 | -24.5 ± 2.9 | SS | 74.8 ± 5.9 | -25.7 ± 3.1 | SS |
| L100<br>Aml5/B5 | 77.2 ± 5.3 | -23.2 ± 1.6 | SS | 78.0 ± 5.0 | -23.0 ± 1.7 | SS | 78.1 ± 4.5 | -22.9 ± 1.8 | SS | 76.7 ± 4.8 | -23.4 ± 2.0 | SS | 75.9 ± 5.0 | -24.6 ± 2.3 | SS | 75.0 ± 5.7 | -25.4 ± 2.4 | SS |
| L100<br>Aml5/H12.5 | 76.8 ± 5.4 | -23.6 ± 2.2 | SS | 77.2 ± 5.1 | -23.7 ± 2.0 | SS | 76.9 ± 5.0 | -24.0 ± 2.2 | SS | 75.2 ± 4.9 | -24.9 ± 2.2 | SS | 75.1 ± 5.2 | -25.3 ± 2.3 | SS | 74.5 ± 5.7 | -26.0 ± 2.3 | SS |
| L100<br>B5/H12.5 | 79.8 ± 5.4 | -20.6 ± 2.0 | SS | 80.3 ± 5.0 | -20.7 ± 2.0 | SS | 79.9 ± 4.8 | -21.0 ± 2.4 | SS | 78.0 ± 5.0 | -22.1 ± 2.5 | SS | 77.4 ± 5.0 | -23.1 ± 2.7 | SS | 76.5 ± 5.7 | -24.0 ± 2.8 | SS |

**Al300 =** aliskiren 300 mg; **Aml5** = amlodipine 5 mg; **B5** = bisoprolol 5 mg; **E20** = enalapril 20 mg; **H12.5** = hydrochlorothiazide 12.5 mg; **L100** = losartan 100 mg; **SS** = statistically significant ($P$ < 0.00001)

**Table S5.** *P*-values calculated using the Kolmogorov-Smirnov test for changes in diastolic blood pressure in populations ($n$ = 100) with different ACE activity receiving the same regimens (case 1: $c_{ACE}$ = 7.0 $h^{-1}$, case 2: $c_{ACE}$ = 8.9 $h^{-1}$, case 3: $c_{ACE}$ = 10.8 $h^{-1}$, case 4: $c_{ACE}$ = 42.3 $h^{-1}$, case 5: $c_{ACE}$ = 54.1 $h^{-1}$, case 6: $c_{ACE}$ = 65.9 $h^{-1}$; *P*-value for case *i* *vs*. case *j* is denoted $P_{ij}$)

| **Regimens** | $P_{12}$ | $P_{13}$ | $P_{23}$ | $P_{14}$ | $P_{15}$ | $P_{16}$ | $P_{24}$ | $P_{25}$ | $P_{26}$ | $P_{34}$ | $P_{35}$ | $P_{36}$ | $P_{45}$ | $P_{46}$ | $P_{56}$ |
|---|---|---|---|---|---|---|---|---|---|---|---|---|---|---|---|
| Al300 | 0.01581 | 0.00007 | 0.11113 | SS | SS | SS | SS | SS | SS | SS | SS | SS | 0.00045 | SS | 0.00025 |
| E20 | SS | SS | SS | SS | SS | SS | SS | SS | SS | SS | SS | SS | SS | SS | 0.00002 |
| L100 | 0.03663 | 0.00013 | 0.15454 | SS | SS | SS | SS | SS | SS | SS | SS | SS | 0.00025 | SS | 0.00079 |
| Aml5 | 0.81275 | 0.36672 | 0.46756 | SS | 0.00007 | SS | 0.00013 | 0.00013 | 0.00002 | 0.00386 | 0.00630 | 0.00136 | 0.90621 | 0.81275 | 0.69937 |
| B5 | 0.07832 | 0.00025 | 0.07832 | SS | SS | SS | SS | SS | SS | SS | SS | SS | 0.00013 | SS | 0.00630 |
| H12.5 | 0.90621 | 0.90621 | 0.90621 | 0.00007 | SS | SS | 0.00025 | SS | SS | 0.00045 | SS | SS | 0.11113 | 0.11113 | 0.99376 |
| Al300 Aml5 | 0.11113 | 0.02431 | 0.46756 | SS | SS | SS | SS | SS | SS | SS | SS | SS | 0.00232 | SS | 0.01581 |
| Al300 B5 | 0.81275 | 0.99376 | 0.96707 | 0.00136 | SS | SS | 0.00079 | SS | SS | 0.00386 | SS | SS | 0.00386 | SS | 0.00630 |
| Al300 H12.5 | 0.28093 | 0.05410 | 0.21055 | SS | SS | SS | SS | SS | SS | 0.00079 | SS | SS | 0.03663 | 0.00136 | 0.00630 |
| E20 Aml5 | SS | SS | 0.00079 | SS | SS | SS | SS | SS | SS | SS | SS | SS | SS | SS | 0.00136 |
| E20 B5 | 0.00136 | SS | 0.02431 | SS | SS | SS | SS | SS | SS | SS | SS | SS | 0.00025 | SS | 0.00025 |
| E20 H12.5 | 0.00232 | SS | 0.00630 | SS | SS | SS | SS | SS | SS | SS | SS | SS | 0.00025 | SS | 0.00136 |
| L100 Aml5 | 0.05410 | 0.02431 | 0.28093 | SS | SS | SS | SS | SS | SS | SS | SS | SS | 0.00136 | SS | 0.02431 |
| L100 B5 | 0.69937 | 0.58062 | 0.69937 | 0.58062 | 0.00025 | SS | 0.11113 | 0.00025 | SS | 0.15454 | SS | SS | 0.00630 | SS | 0.00630 |
| L100 H12.5 | 0.21055 | 0.05410 | 0.36672 | SS | SS | SS | SS | SS | SS | 0.00045 | SS | SS | 0.02431 | 0.00079 | 0.00630 |
| Al300 Aml5/B5 | 0.90621 | 0.36672 | 0.46756 | 0.03663 | SS | SS | 0.02431 | SS | SS | 0.02431 | SS | SS | 0.00386 | SS | 0.03663 |
| Al300 Aml5/H12.5 | 0.58062 | 0.15454 | 0.28093 | 0.01581 | SS | SS | 0.00232 | SS | SS | 0.15454 | 0.01581 | 0.00013 | 0.15454 | 0.01581 | 0.07832 |
| Al300 B5/H12.5 | 0.28093 | 0.11113 | 0.58062 | SS | SS | SS | 0.00079 | SS | SS | 0.01008 | SS | SS | 0.07832 | 0.00025 | 0.02431 |
| E20 Aml5/B5 | 0.00025 | SS | 0.05410 | SS | SS | SS | SS | SS | SS | SS | SS | SS | 0.00002 | SS | 0.02431 |
| E20 Aml5/H12.5 | 0.02431 | 0.00045 | 0.01581 | SS | SS | SS | SS | SS | SS | SS | SS | SS | 0.00136 | 0.00004 | 0.03663 |
| E20 B5/H12.5 | 0.01581 | 0.00002 | 0.11113 | SS | SS | SS | SS | SS | SS | SS | SS | SS | 0.00630 | SS | 0.01581 |
| L100 Aml5/B5 | 0.58062 | 0.28093 | 0.69937 | 0.21055 | 0.00007 | SS | 0.15454 | SS | SS | 0.28093 | SS | SS | 0.00232 | SS | 0.07832 |
| L100 Aml5/H12.5 | 0.58062 | 0.11113 | 0.28093 | 0.01008 | SS | SS | 0.00136 | SS | SS | 0.15454 | 0.00136 | 0.00002 | 0.05410 | 0.01581 | 0.07832 |
| L100 B5/H12.5 | 0.46756 | 0.28093 | 0.46756 | 0.00007 | SS | SS | 0.00386 | SS | SS | 0.05410 | SS | SS | 0.05410 | 0.00045 | 0.02431 |

**Al300 =** aliskiren 300 mg; **Aml5** = amlodipine 5 mg; **B5** = bisoprolol 5 mg; **E20** = enalapril 20 mg; **H12.5** = hydrochlorothiazide 12.5 mg; **L100** = losartan 100 mg; **SS** = statistically significant ($P$ < 0.00001)

**Figure S10.** Simulated change in heart rate from baseline to week 4 (mean ± SD, $n$ = 100).

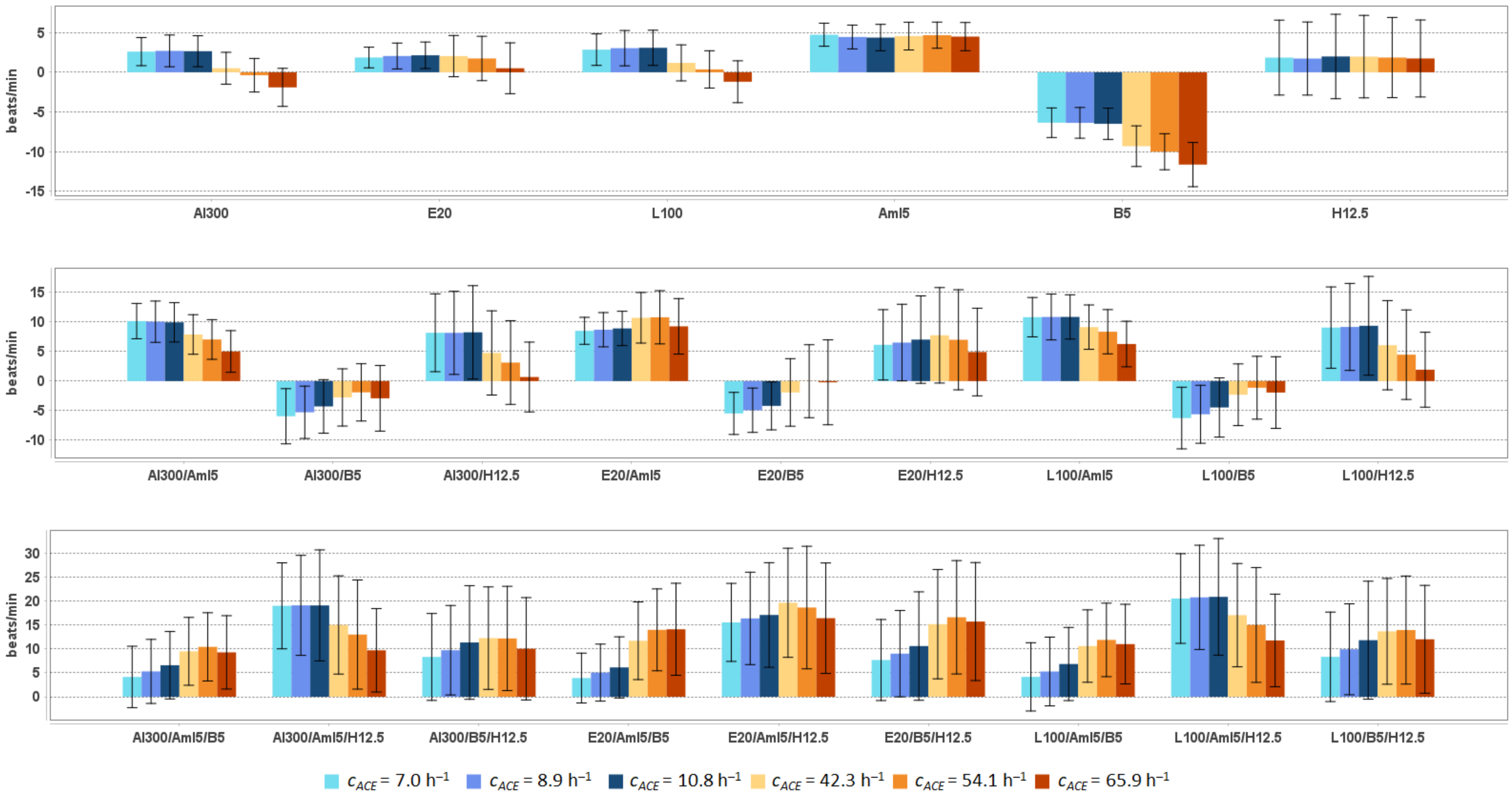


**Al300 =** aliskiren 300 mg; **Aml5** = amlodipine 5 mg; **B5** = bisoprolol 5 mg; **E20** = enalapril 20 mg; **H12.5** = hydrochlorothiazide 12.5 mg; **L100** = losartan 100 mg

**Table S6.** Simulated response of <u>heart rate</u> to antihypertensive therapy in virtual hypertensive populations ($n$ = 100) with different ACE activity, including $P$-values (Kolmogorov-Smirnov test) for endpoint *vs*. baseline; data are presented as mean ± SD in beats/min

| Regimens | $c_{ACE}$ = 7.0 h$^{-1}$ | | | $c_{ACE}$ = 8.9 h$^{-1}$ | | | $c_{ACE}$ = 10.8 h$^{-1}$ | | | $c_{ACE}$ = 42.3 h$^{-1}$ | | | $c_{ACE}$ = 54.1 h$^{-1}$ | | | $c_{ACE}$ = 65.9 h$^{-1}$ | | |
|---|---|---|---|---|---|---|---|---|---|---|---|---|---|---|---|---|---|---|
| | **Value** | **Change** | ***P*** | **Value** | **Change** | ***P*** | **Value** | **Change** | ***P*** | **Value** | **Change** | ***P*** | **Value** | **Change** | ***P*** | **Value** | **Change** | ***P*** |
| Baseline | 77.2 ± 6.4 | – | – | 75.7 ± 7.6 | – | – | 75.2 ± 7.1 | – | – | 75.9 ± 7.1 | – | – | 76.6 ± 6.8 | – | – | 76.0 ± 7.4 | – | – |
| Al300 | 79.8 ± 6.6 | 2.6 ± 1.8 | 0.05410 | 78.4 ± 8.1 | 2.7 ± 2.0 | 0.15454 | 77.9 ± 7.6 | 2.7 ± 2.0 | 0.15454 | 76.5 ± 7.7 | 0.5 ± 2.0 | 0.90621 | 76.2 ± 7.0 | -0.4 ± 2.1 | 0.96707 | 74.1 ± 8.2 | -1.9 ± 2.4 | 0.28093 |
| E20 | 79.1 ± 6.5 | 1.9 ± 1.3 | 0.21055 | 77.7 ± 8.0 | 2.0 ± 1.6 | 0.28093 | 77.4 ± 7.5 | 2.1 ± 1.7 | 0.21055 | 78.0 ± 8.0 | 2.0 ± 2.6 | 0.36672 | 78.3 ± 7.2 | 1.7 ± 2.8 | 0.28093 | 76.5 ± 8.6 | 0.5 ± 3.2 | 0.69937 |
| L100 | 80.1 ± 6.6 | 2.8 ± 2.0 | 0.02431 | 78.7 ± 8.2 | 3.0 ± 2.2 | 0.15454 | 78.3 ± 7.7 | 3.1 ± 2.2 | 0.11113 | 77.1 ± 7.8 | 1.2 ± 2.3 | 0.69937 | 77.0 ± 7.1 | 0.4 ± 2.3 | 0.90621 | 74.8 ± 8.3 | -1.2 ± 2.6 | 0.46756 |
| Aml5 | 82.0 ± 6.6 | 4.7 ± 1.5 | 0.00013 | 80.1 ± 7.8 | 4.4 ± 1.5 | 0.00136 | 79.6 ± 7.1 | 4.4 ± 1.7 | 0.00232 | 80.5 ± 7.6 | 4.6 ± 1.7 | 0.00630 | 81.3 ± 7.2 | 4.7 ± 1.6 | 0.00025 | 80.5 ± 7.8 | 4.5 ± 1.8 | 0.00136 |
| B5 | 70.9 ± 6.5 | -6.4 ± 1.9 | SS | 69.3 ± 7.9 | -6.4 ± 1.9 | 0.00004 | 68.8 ± 7.6 | -6.5 ± 2.0 | SS | 66.6 ± 7.4 | -9.3 ± 2.6 | SS | 66.6 ± 6.9 | -10.0 ± 2.3 | SS | 64.3 ± 8.1 | -11.6 ± 2.8 | SS |
| H12.5 | 79.1 ± 7.9 | 1.8 ± 4.7 | 0.21055 | 77.4 ± 8.8 | 1.7 ± 4.6 | 0.36672 | 77.2 ± 9.0 | 2.0 ± 5.3 | 0.21055 | 77.9 ± 8.8 | 2.0 ± 5.2 | 0.58062 | 78.4 ± 8.1 | 1.9 ± 5.0 | 0.28093 | 77.7 ± 9.4 | 1.7 ± 4.8 | 0.03663 |
| Al300<br>Aml5 | 87.4 ± 6.9 | 10.1 ± 3.0 | SS | 85.7 ± 8.8 | 10.0 ± 3.5 | SS | 85.2 ± 8.0 | 9.9 ± 3.3 | SS | 83.8 ± 8.6 | 7.9 ± 3.3 | SS | 83.6 ± 7.6 | 7.0 ± 3.4 | SS | 81.0 ± 8.9 | 5.0 ± 3.5 | 0.00025 |
| Al300<br>B5 | 71.3 ± 7.4 | -5.9 ± 4.7 | SS | 70.4 ± 8.6 | -5.3 ± 4.4 | 0.00386 | 71.0 ± 8.6 | -4.3 ± 4.5 | 0.00136 | 73.2 ± 8.9 | -2.8 ± 4.8 | 0.01581 | 74.7 ± 8.1 | -1.9 ± 4.8 | 0.21055 | 73.1 ± 9.6 | -2.9 ± 5.6 | 0.02431 |
| Al300<br>H12.5 | 85.4 ± 9.0 | 8.2 ± 6.6 | SS | 83.8 ± 10.7 | 8.2 ± 7.0 | SS | 83.5 ± 11.0 | 8.2 ± 7.9 | SS | 80.7 ± 10.4 | 4.8 ± 7.1 | 0.05410 | 79.7 ± 9.3 | 3.1 ± 7.1 | 0.05410 | 76.7 ± 10.6 | 0.7 ± 5.9 | 0.21055 |
| E20<br>Aml5 | 85.7 ± 6.7 | 8.5 ± 2.3 | SS | 84.4 ± 8.5 | 8.7 ± 2.9 | SS | 84.1 ± 7.8 | 8.9 ± 2.9 | SS | 86.7 ± 9.1 | 10.7 ± 4.3 | SS | 87.4 ± 8.0 | 10.8 ± 4.5 | SS | 85.2 ± 9.4 | 9.3 ± 4.7 | SS |
| E20<br>B5 | 71.8 ± 6.9 | -5.5 ± 3.6 | 0.00002 | 70.7 ± 8.4 | -4.9 ± 3.7 | 0.00386 | 71.1 ± 8.4 | -4.2 ± 4.1 | 0.00136 | 74.0 ± 9.4 | -1.9 ± 5.7 | 0.15454 | 76.6 ± 8.9 | -0.0 ± 6.2 | 0.46756 | 75.8 ± 10.5 | -0.2 ± 7.2 | 0.21055 |
| E20<br>H12.5 | 83.4 ± 8.6 | 6.1 ± 5.9 | 0.00002 | 82.2 ± 10.3 | 6.5 ± 6.5 | 0.00013 | 82.3 ± 10.7 | 7.0 ± 7.4 | 0.00007 | 83.7 ± 11.2 | 7.7 ± 8.1 | 0.00013 | 83.6 ± 10.2 | 7.0 ± 8.5 | SS | 80.9 ± 11.7 | 4.9 ± 7.4 | 0.00007 |
| L100<br>Aml5 | 88.0 ± 7.0 | 10.8 ± 3.3 | SS | 86.5 ± 9.0 | 10.8 ± 3.9 | SS | 86.1 ± 8.2 | 10.8 ± 3.7 | SS | 85.1 ± 8.8 | 9.1 ± 3.7 | SS | 84.9 ± 7.7 | 8.3 ± 3.7 | SS | 82.2 ± 9.0 | 6.3 ± 3.9 | 0.00004 |
| L100<br>B5 | 71.0 ± 7.8 | -6.2 ± 5.2 | SS | 70.1 ± 8.8 | -5.6 ± 4.9 | 0.00136 | 70.8 ± 8.9 | -4.5 ± 5.0 | 0.00025 | 73.6 ± 9.1 | -2.3 ± 5.2 | 0.03663 | 75.5 ± 8.3 | -1.1 ± 5.3 | 0.28093 | 74.0 ± 9.9 | -1.9 ± 6.1 | 0.07832 |
| L100<br>H12.5 | 86.3 ± 9.2 | 9.0 ± 6.9 | SS | 84.8 ± 11.0 | 9.2 ± 7.4 | SS | 84.6 ± 11.4 | 9.4 ± 8.4 | SS | 82.0 ± 10.7 | 6.1 ± 7.5 | 0.00630 | 81.1 ± 9.6 | 4.5 ± 7.6 | 0.00630 | 77.9 ± 10.9 | 1.9 ± 6.3 | 0.01581 |
| Al300<br>Aml5/B5 | 81.4 ± 8.6 | 4.1 ± 6.4 | 0.00045 | 81.0 ± 10.5 | 5.3 ± 6.7 | 0.00136 | 81.8 ± 10.3 | 6.6 ± 7.0 | 0.00007 | 85.4 ± 11.0 | 9.5 ± 7.1 | SS | 87.0 ± 9.5 | 10.4 ± 7.1 | SS | 85.2 ± 11.1 | 9.3 ± 7.7 | SS |
| Al300<br>Aml5/H12.5 | 96.2 ± 10.8 | 19.0 ± 9.0 | SS | 94.8 ± 13.0 | 19.1 ± 10.4 | SS | 94.3 ± 13.7 | 19.1 ± 11.6 | SS | 90.9 ± 13.0 | 15.0 ± 10.3 | SS | 89.6 ± 12.6 | 13.0 ± 11.4 | SS | 85.7 ± 12.6 | 9.7 ± 8.7 | SS |
| Al300<br>B5/H12.5 | 85.6 ± 10.8 | 8.3 ± 9.1 | SS | 85.4 ± 12.5 | 9.7 ± 9.3 | SS | 86.6 ± 14.3 | 11.3 ± 11.9 | SS | 88.2 ± 13.5 | 12.2 ± 10.7 | SS | 88.8 ± 11.9 | 12.2 ± 10.9 | SS | 86.0 ± 14.0 | 10.0 ± 10.7 | SS |
| E20<br>Aml5/B5 | 81.1 ± 7.9 | 3.9 ± 5.2 | 0.00136 | 80.7 ± 10.1 | 5.0 ± 6.0 | 0.00386 | 81.4 ± 9.9 | 6.1 ± 6.4 | 0.00079 | 87.6 ± 11.7 | 11.7 ± 8.1 | SS | 90.6 ± 10.5 | 14.0 ± 8.6 | SS | 90.1 ± 12.2 | 14.1 ± 9.6 | SS |
| E20<br>Aml5/H12.5 | 92.8 ± 10.2 | 15.5 ± 8.1 | SS | 92.0 ± 12.4 | 16.4 ± 9.7 | SS | 92.3 ± 13.1 | 17.1 ± 10.9 | SS | 95.6 ± 14.0 | 19.6 ± 11.4 | SS | 95.2 ± 13.6 | 18.6 ± 12.8 | SS | 92.4 ± 14.7 | 16.4 ± 11.5 | SS |
| E20<br>B5/H12.5 | 84.9 ± 10.3 | 7.7 ± 8.5 | SS | 84.7 ± 12.3 | 9.0 ± 9.0 | SS | 85.8 ± 13.9 | 10.6 ± 11.3 | SS | 91.1 ± 14.1 | 15.2 ± 11.4 | SS | 93.2 ± 12.6 | 16.6 ± 11.8 | SS | 91.7 ± 15.1 | 15.7 ± 12.3 | SS |
| L100<br>Aml5/B5 | 81.4 ± 9.2 | 4.2 ± 7.1 | 0.00025 | 81.0 ± 10.8 | 5.3 ± 7.2 | 0.00079 | 82.1 ± 10.7 | 6.8 ± 7.6 | 0.00007 | 86.5 ± 11.3 | 10.6 ± 7.6 | SS | 88.5 ± 9.9 | 11.9 ± 7.7 | SS | 87.0 ± 11.4 | 11.0 ± 8.3 | SS |
| L100<br>Aml5/H12.5 | 97.8 ± 11.1 | 20.5 ± 9.4 | SS | 96.4 ± 13.3 | 20.8 ± 10.9 | SS | 96.1 ± 14.2 | 20.9 ± 12.2 | SS | 93.0 ± 13.4 | 17.0 ± 10.8 | SS | 91.6 ± 13.0 | 15.0 ± 12.0 | SS | 87.7 ± 13.3 | 11.8 ± 9.7 | SS |
| L100<br>B5/H12.5 | 85.6 ± 11.0 | 8.3 ± 9.3 | SS | 85.6 ± 12.6 | 9.9 ± 9.5 | SS | 87.1 ± 14.6 | 11.8 ± 12.3 | SS | 89.6 ± 13.8 | 13.7 ± 11.0 | SS | 90.5 ± 12.2 | 13.9 ± 11.3 | SS | 88.0 ± 14.4 | 12.0 ± 11.3 | SS |

**Al300 =** aliskiren 300 mg; **Aml5** = amlodipine 5 mg; **B5** = bisoprolol 5 mg; **E20** = enalapril 20 mg; **H12.5** = hydrochlorothiazide 12.5 mg; **L100** = losartan 100 mg; **SS** = statistically significant ($P$ < 0.00001)

**Table S7.** *P*-values calculated using the Kolmogorov-Smirnov test for changes in heart rate in populations ($n$ = 100) with different ACE activity receiving the same regimens (case 1: $c_{ACE}$ = 7.0 $h^{-1}$, case 2: $c_{ACE}$ = 8.9 $h^{-1}$, case 3: $c_{ACE}$ = 10.8 $h^{-1}$, case 4: $c_{ACE}$ = 42.3 $h^{-1}$, case 5: $c_{ACE}$ = 54.1 $h^{-1}$, case 6: $c_{ACE}$ = 65.9 $h^{-1}$; *P*-value for case *i* *vs*. case *j* is denoted $P_{ij}$)

| Regimens | $P_{12}$ | $P_{13}$ | $P_{23}$ | $P_{14}$ | $P_{15}$ | $P_{16}$ | $P_{24}$ | $P_{25}$ | $P_{26}$ | $P_{34}$ | $P_{35}$ | $P_{36}$ | $P_{45}$ | $P_{46}$ | $P_{56}$ |
|---|---|---|---|---|---|---|---|---|---|---|---|---|---|---|---|
| Al300 | 0.36672 | 0.58062 | 0.36672 | SS | SS | SS | SS | SS | SS | SS | SS | SS | 0.02431 | SS | 0.00013 |
| E20 | 0.15454 | 0.15454 | 0.46756 | 0.00630 | 0.00013 | SS | 0.11113 | 0.00630 | SS | 0.11113 | 0.01008 | SS | 0.46756 | 0.00386 | 0.02431 |
| L100 | 0.36672 | 0.21055 | 0.69937 | 0.00004 | SS | SS | 0.00007 | SS | SS | 0.00013 | SS | SS | 0.05410 | SS | 0.00045 |
| Aml5 | 0.69937 | 0.58062 | 0.99963 | 0.81275 | 0.90621 | 0.46756 | 0.69937 | 0.36672 | 0.46756 | 0.81275 | 0.36672 | 0.81275 | 0.69937 | 0.99376 | 0.46756 |
| B5 | 0.96707 | 0.81275 | 0.81275 | SS | SS | SS | SS | SS | SS | SS | SS | SS | 0.02431 | SS | 0.00045 |
| H12.5 | 0.58062 | 0.81275 | 0.58062 | 0.58062 | 0.90621 | 0.90621 | 0.11113 | 0.21055 | 0.36672 | 0.81275 | 0.58062 | 0.58062 | 0.90621 | 0.81275 | 0.99376 |
| Al300 Aml5 | 0.69937 | 0.69937 | 0.81275 | 0.00007 | SS | SS | 0.00007 | SS | SS | 0.00232 | SS | SS | 0.11113 | SS | 0.00386 |
| Al300 B5 | 0.69937 | 0.05410 | 0.21055 | 0.00025 | SS | 0.00013 | 0.00232 | 0.00013 | 0.00232 | 0.05410 | 0.03663 | 0.21055 | 0.69937 | 0.58062 | 0.28093 |
| Al300 H12.5 | 0.81275 | 0.58062 | 0.36672 | 0.00079 | SS | SS | 0.00045 | SS | SS | 0.01008 | 0.00025 | SS | 0.11113 | 0.00025 | 0.15454 |
| E20 Aml5 | 0.36672 | 0.05410 | 0.81275 | SS | 0.00007 | 0.00630 | 0.00045 | 0.00386 | 0.21055 | 0.00386 | 0.00386 | 0.28093 | 0.69937 | 0.07832 | 0.15454 |
| E20 B5 | 0.81275 | 0.02431 | 0.28093 | SS | SS | SS | 0.00025 | SS | SS | 0.00136 | 0.00004 | 0.00025 | 0.11113 | 0.21055 | 0.36672 |
| E20 H12.5 | 0.96707 | 0.28093 | 0.46756 | 0.07832 | 0.15454 | 0.02431 | 0.21055 | 0.28093 | 0.01008 | 0.96707 | 0.46756 | 0.07832 | 0.36672 | 0.05410 | 0.28093 |
| L100 Aml5 | 0.69937 | 0.28093 | 0.90621 | 0.02431 | SS | SS | 0.01008 | SS | SS | 0.01008 | 0.00007 | SS | 0.15454 | 0.00013 | 0.00630 |
| L100 B5 | 0.81275 | 0.07832 | 0.21055 | 0.00004 | SS | SS | 0.00025 | SS | 0.00013 | 0.01008 | 0.00630 | 0.03663 | 0.36672 | 0.69937 | 0.28093 |
| L100 H12.5 | 0.96707 | 0.58062 | 0.58062 | 0.00386 | SS | SS | 0.00630 | SS | SS | 0.01581 | 0.00079 | SS | 0.15454 | 0.00136 | 0.11113 |
| Al300 Aml5/B5 | 0.36672 | 0.03663 | 0.28093 | SS | SS | SS | 0.00079 | 0.00025 | 0.00045 | 0.03663 | 0.00386 | 0.05410 | 0.28093 | 0.58062 | 0.58062 |
| Al300 Aml5/H12.5 | 0.69937 | 0.58062 | 0.81275 | 0.00386 | SS | SS | 0.01581 | 0.00002 | SS | 0.07832 | 0.00045 | SS | 0.21055 | 0.00079 | 0.07832 |
| Al300 B5/H12.5 | 0.11113 | 0.01008 | 0.36672 | 0.01581 | 0.01008 | 0.15454 | 0.28093 | 0.36672 | 0.99376 | 0.28093 | 0.21055 | 0.46756 | 0.46756 | 0.36672 | 0.36672 |
| E20 Aml5/B5 | 0.15454 | 0.01581 | 0.36672 | SS | SS | SS | SS | SS | SS | SS | SS | SS | 0.11113 | 0.28093 | 0.81275 |
| E20 Aml5/H12.5 | 0.69937 | 0.15454 | 0.58062 | 0.00630 | 0.03663 | 0.81275 | 0.03663 | 0.05410 | 0.90621 | 0.36672 | 0.69937 | 0.81275 | 0.46756 | 0.07832 | 0.21055 |
| E20 B5/H12.5 | 0.15454 | 0.00630 | 0.21055 | SS | SS | SS | 0.00045 | 0.00013 | 0.00004 | 0.00386 | 0.00025 | 0.01008 | 0.69937 | 0.58062 | 0.81275 |
| L100 Aml5/B5 | 0.46756 | 0.05410 | 0.21055 | SS | SS | SS | 0.00013 | SS | SS | 0.00630 | 0.00013 | 0.00136 | 0.15454 | 0.69937 | 0.69937 |
| L100 Aml5/H12.5 | 0.69937 | 0.46756 | 0.81275 | 0.01581 | SS | SS | 0.02431 | 0.00004 | SS | 0.15454 | 0.00079 | SS | 0.21055 | 0.00045 | 0.07832 |
| L100 B5/H12.5 | 0.21055 | 0.00630 | 0.28093 | 0.00079 | 0.00232 | 0.00630 | 0.11113 | 0.07832 | 0.58062 | 0.21055 | 0.07832 | 0.81275 | 0.58062 | 0.69937 | 0.46756 |

**Al300 =** aliskiren 300 mg; **Aml5** = amlodipine 5 mg; **B5** = bisoprolol 5 mg; **E20** = enalapril 20 mg; **H12.5** = hydrochlorothiazide 12.5 mg; **L100** = losartan 100 mg; **SS** = statistically significant ($P$ < 0.00001)

**Figure S11.** Simulated change in systemic arterial elasticity from baseline to week 4 (mean ± SD, $n$ = 100)

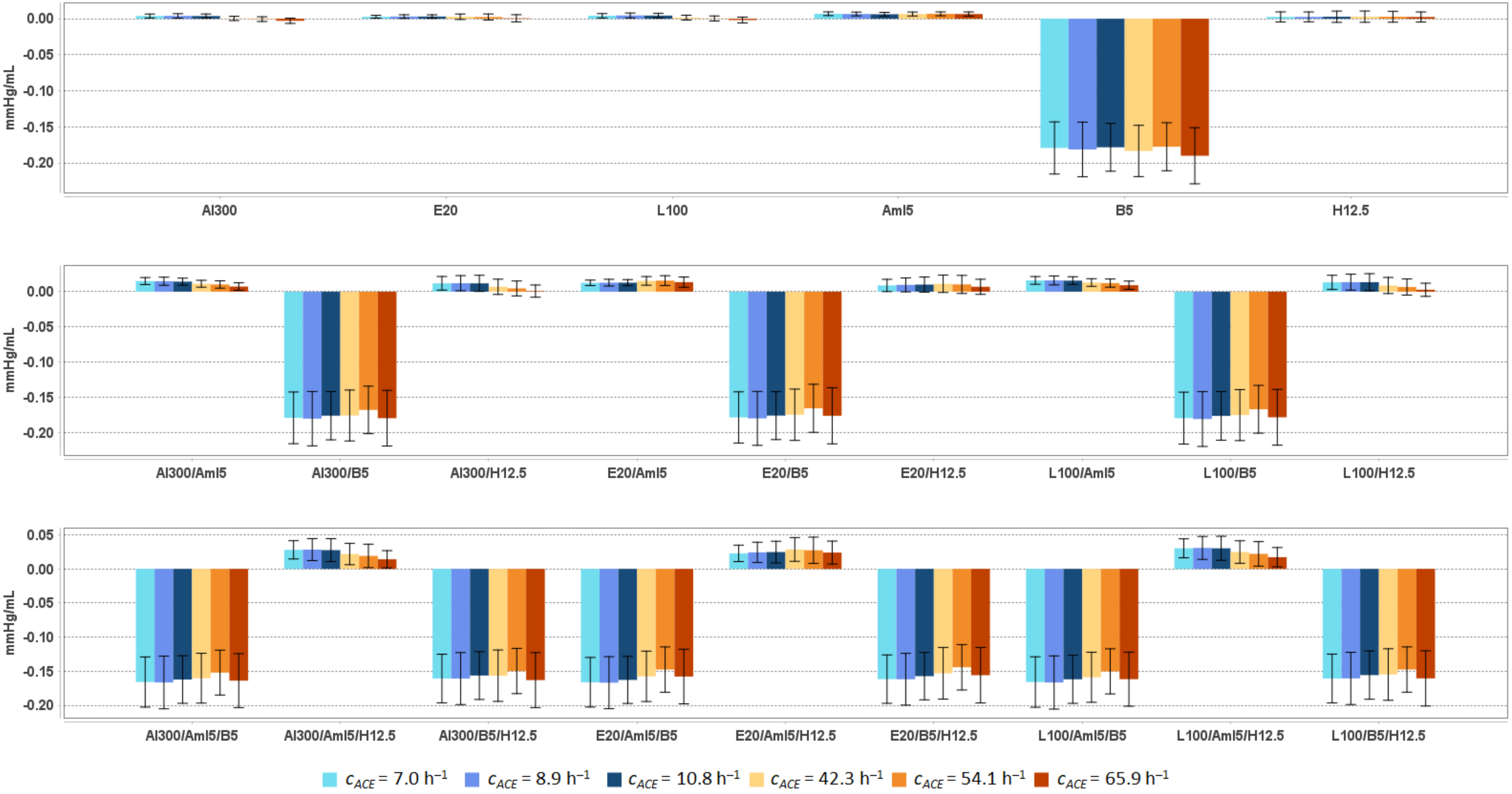


**Al300** = aliskiren 300 mg; **Aml5** = amlodipine 5 mg; **B5** = bisoprolol 5 mg; **E20** = enalapril 20 mg; **H12.5** = hydrochlorothiazide 12.5 mg; **L100** = losartan 100 mg

**Table S8.** Simulated response of <u>systemic arterial elasticity</u> to antihypertensive therapy in virtual hypertensive populations ($n$ = 100) with different ACE activity, including $P$-values (Kolmogorov-Smirnov test) for endpoint *vs*. baseline; data are presented as mean ± SD in mmHg/mL

| Regimens | $c_{ACE}$ = 7.0 h$^{-1}$ | | | $c_{ACE}$ = 8.9 h$^{-1}$ | | | $c_{ACE}$ = 10.8 h$^{-1}$ | | | $c_{ACE}$ = 42.3 h$^{-1}$ | | | $c_{ACE}$ = 54.1 h$^{-1}$ | | | $c_{ACE}$ = 65.9 h$^{-1}$ | | |
|---|---|---|---|---|---|---|---|---|---|---|---|---|---|---|---|---|---|---|
| | **Value** | **Change** | ***P*** | **Value** | **Change** | ***P*** | **Value** | **Change** | ***P*** | **Value** | **Change** | ***P*** | **Value** | **Change** | ***P*** | **Value** | **Change** | ***P*** |
| Baseline | 1.149 ± 0.240 | – | – | 1.163 ± 0.250 | – | – | 1.142 ± 0.220 | – | – | 1.153 ± 0.230 | – | – | 1.108 ± 0.220 | – | – | 1.177 ± 0.254 | – | – |
| Al300 | 1.153 ± 0.240 | 0.004 ± 0.003 | 0.99963 | 1.167 ± 0.251 | 0.004 ± 0.003 | 1.00000 | 1.146 ± 0.220 | 0.004 ± 0.003 | 1.00000 | 1.154 ± 0.229 | 0.000 ± 0.003 | 1.00000 | 1.107 ± 0.221 | -0.001 ± 0.003 | 1.00000 | 1.174 ± 0.254 | -0.003 ± 0.004 | 1.00000 |
| E20 | 1.152 ± 0.240 | 0.003 ± 0.002 | 1.00000 | 1.166 ± 0.251 | 0.003 ± 0.002 | 1.00000 | 1.145 ± 0.220 | 0.003 ± 0.002 | 1.00000 | 1.156 ± 0.229 | 0.003 ± 0.004 | 1.00000 | 1.110 ± 0.221 | 0.002 ± 0.004 | 1.00000 | 1.178 ± 0.254 | 0.001 ± 0.005 | 1.00000 |
| L100 | 1.153 ± 0.240 | 0.004 ± 0.003 | 0.99963 | 1.167 ± 0.251 | 0.005 ± 0.003 | 1.00000 | 1.147 ± 0.220 | 0.004 ± 0.003 | 1.00000 | 1.155 ± 0.229 | 0.002 ± 0.003 | 1.00000 | 1.108 ± 0.220 | 0.000 ± 0.003 | 1.00000 | 1.175 ± 0.254 | -0.002 ± 0.004 | 1.00000 |
| Aml5 | 1.156 ± 0.240 | 0.007 ± 0.003 | 0.99376 | 1.169 ± 0.250 | 0.007 ± 0.003 | 1.00000 | 1.149 ± 0.220 | 0.006 ± 0.003 | 0.99963 | 1.160 ± 0.230 | 0.007 ± 0.003 | 1.00000 | 1.114 ± 0.221 | 0.007 ± 0.003 | 1.00000 | 1.184 ± 0.255 | 0.007 ± 0.003 | 0.99963 |
| B5 | 0.970 ± 0.204 | -0.179 ± 0.036 | 0.00002 | 0.981 ± 0.212 | -0.181 ± 0.038 | 0.00002 | 0.964 ± 0.187 | -0.178 ± 0.033 | SS | 0.970 ± 0.195 | -0.183 ± 0.036 | 0.00004 | 0.930 ± 0.187 | -0.178 ± 0.033 | SS | 0.987 ± 0.216 | -0.190 ± 0.039 | 0.00002 |
| H12.5 | 1.151 ± 0.241 | 0.003 ± 0.007 | 1.00000 | 1.165 ± 0.251 | 0.003 ± 0.007 | 1.00000 | 1.145 ± 0.221 | 0.003 ± 0.008 | 1.00000 | 1.156 ± 0.230 | 0.003 ± 0.008 | 1.00000 | 1.110 ± 0.221 | 0.003 ± 0.008 | 1.00000 | 1.180 ± 0.254 | 0.002 ± 0.007 | 1.00000 |
| Al300 Aml5 | 1.164 ± 0.240 | 0.015 ± 0.005 | 0.90621 | 1.178 ± 0.251 | 0.015 ± 0.006 | 0.99376 | 1.157 ± 0.220 | 0.014 ± 0.005 | 0.96707 | 1.164 ± 0.229 | 0.011 ± 0.005 | 1.00000 | 1.118 ± 0.221 | 0.010 ± 0.005 | 0.99963 | 1.184 ± 0.254 | 0.007 ± 0.005 | 0.99963 |
| Al300 B5 | 0.970 ± 0.204 | -0.179 ± 0.037 | 0.00002 | 0.983 ± 0.212 | -0.180 ± 0.038 | 0.00004 | 0.967 ± 0.186 | -0.176 ± 0.034 | SS | 0.978 ± 0.195 | -0.175 ± 0.036 | 0.00007 | 0.940 ± 0.187 | -0.167 ± 0.034 | 0.00004 | 0.998 ± 0.216 | -0.179 ± 0.039 | 0.00002 |
| Al300 H12.5 | 1.161 ± 0.241 | 0.012 ± 0.010 | 0.90621 | 1.175 ± 0.252 | 0.012 ± 0.011 | 0.99963 | 1.154 ± 0.221 | 0.012 ± 0.011 | 0.96707 | 1.160 ± 0.229 | 0.007 ± 0.011 | 1.00000 | 1.112 ± 0.222 | 0.005 ± 0.011 | 0.99963 | 1.178 ± 0.254 | 0.001 ± 0.009 | 1.00000 |
| E20 Aml5 | 1.161 ± 0.240 | 0.013 ± 0.004 | 0.96707 | 1.176 ± 0.251 | 0.013 ± 0.005 | 0.99376 | 1.155 ± 0.220 | 0.013 ± 0.004 | 0.96707 | 1.168 ± 0.229 | 0.015 ± 0.006 | 0.99376 | 1.123 ± 0.222 | 0.016 ± 0.007 | 0.99376 | 1.191 ± 0.254 | 0.013 ± 0.007 | 0.99376 |
| E20 B5 | 0.971 ± 0.204 | -0.178 ± 0.036 | 0.00002 | 0.983 ± 0.212 | -0.179 ± 0.038 | 0.00004 | 0.967 ± 0.187 | -0.176 ± 0.034 | SS | 0.979 ± 0.195 | -0.174 ± 0.036 | 0.00007 | 0.942 ± 0.187 | -0.165 ± 0.034 | 0.00004 | 1.001 ± 0.216 | -0.176 ± 0.040 | 0.00004 |
| E20 H12.5 | 1.158 ± 0.241 | 0.009 ± 0.009 | 0.99376 | 1.172 ± 0.251 | 0.010 ± 0.010 | 1.00000 | 1.152 ± 0.221 | 0.010 ± 0.011 | 0.99376 | 1.164 ± 0.229 | 0.011 ± 0.012 | 0.99963 | 1.118 ± 0.223 | 0.010 ± 0.013 | 0.99963 | 1.184 ± 0.254 | 0.007 ± 0.011 | 1.00000 |
| L100 Aml5 | 1.165 ± 0.241 | 0.016 ± 0.005 | 0.90621 | 1.179 ± 0.251 | 0.016 ± 0.006 | 0.99376 | 1.158 ± 0.220 | 0.016 ± 0.006 | 0.96707 | 1.166 ± 0.229 | 0.013 ± 0.005 | 0.99376 | 1.120 ± 0.221 | 0.012 ± 0.006 | 0.99376 | 1.186 ± 0.254 | 0.009 ± 0.006 | 0.99963 |
| L100 B5 | 0.970 ± 0.204 | -0.179 ± 0.037 | 0.00002 | 0.982 ± 0.212 | -0.180 ± 0.039 | 0.00004 | 0.966 ± 0.186 | -0.176 ± 0.034 | SS | 0.978 ± 0.195 | -0.175 ± 0.036 | 0.00007 | 0.941 ± 0.187 | -0.167 ± 0.034 | 0.00004 | 0.999 ± 0.216 | -0.178 ± 0.039 | 0.00004 |
| L100 H12.5 | 1.162 ± 0.242 | 0.013 ± 0.010 | 0.90621 | 1.176 ± 0.252 | 0.013 ± 0.011 | 0.99963 | 1.156 ± 0.222 | 0.013 ± 0.012 | 0.96707 | 1.162 ± 0.229 | 0.009 ± 0.011 | 0.99963 | 1.114 ± 0.222 | 0.007 ± 0.011 | 0.99963 | 1.180 ± 0.254 | 0.003 ± 0.009 | 1.00000 |
| Al300 Aml5/B5 | 0.983 ± 0.204 | -0.166 ± 0.037 | 0.00004 | 0.996 ± 0.213 | -0.167 ± 0.039 | 0.00025 | 0.980 ± 0.186 | -0.162 ± 0.035 | SS | 0.993 ± 0.195 | -0.160 ± 0.037 | 0.00079 | 0.956 ± 0.189 | -0.152 ± 0.033 | 0.00025 | 1.013 ± 0.216 | -0.164 ± 0.040 | 0.00004 |
| Al300 Aml5/H12.5 | 1.177 ± 0.243 | 0.028 ± 0.013 | 0.58062 | 1.191 ± 0.253 | 0.028 ± 0.016 | 0.90621 | 1.170 ± 0.222 | 0.028 ± 0.017 | 0.58062 | 1.175 ± 0.230 | 0.022 ± 0.016 | 0.96707 | 1.127 ± 0.224 | 0.019 ± 0.017 | 0.90621 | 1.191 ± 0.255 | 0.014 ± 0.013 | 0.99376 |
| Al300 B5/H12.5 | 0.988 ± 0.206 | -0.161 ± 0.036 | 0.00007 | 1.002 ± 0.214 | -0.161 ± 0.038 | 0.00025 | 0.986 ± 0.189 | -0.156 ± 0.035 | 0.00013 | 0.997 ± 0.195 | -0.157 ± 0.038 | 0.00079 | 0.958 ± 0.191 | -0.150 ± 0.033 | 0.00045 | 1.014 ± 0.217 | -0.163 ± 0.040 | 0.00004 |
| E20 Aml5/B5 | 0.983 ± 0.204 | -0.166 ± 0.036 | 0.00004 | 0.996 ± 0.213 | -0.167 ± 0.038 | 0.00025 | 0.980 ± 0.187 | -0.163 ± 0.035 | SS | 0.996 ± 0.195 | -0.158 ± 0.037 | 0.00079 | 0.960 ± 0.189 | -0.148 ± 0.033 | 0.00045 | 1.019 ± 0.217 | -0.158 ± 0.040 | 0.00007 |
| E20 Aml5/H12.5 | 1.172 ± 0.242 | 0.023 ± 0.012 | 0.69937 | 1.187 ± 0.253 | 0.024 ± 0.015 | 0.90621 | 1.167 ± 0.222 | 0.025 ± 0.016 | 0.58062 | 1.182 ± 0.231 | 0.029 ± 0.017 | 0.90621 | 1.135 ± 0.225 | 0.027 ± 0.019 | 0.81275 | 1.201 ± 0.256 | 0.024 ± 0.017 | 0.90621 |
| E20 B5/H12.5 | 0.987 ± 0.206 | -0.162 ± 0.035 | 0.00007 | 1.001 ± 0.215 | -0.162 ± 0.038 | 0.00025 | 0.985 ± 0.189 | -0.157 ± 0.035 | 0.00013 | 1.000 ± 0.196 | -0.153 ± 0.038 | 0.00136 | 0.963 ± 0.191 | -0.144 ± 0.033 | 0.00045 | 1.021 ± 0.217 | -0.156 ± 0.041 | 0.00007 |
| L100 Aml5/B5 | 0.983 ± 0.204 | -0.166 ± 0.037 | 0.00007 | 0.996 ± 0.213 | -0.167 ± 0.039 | 0.00013 | 0.980 ± 0.186 | -0.162 ± 0.035 | SS | 0.994 ± 0.195 | -0.159 ± 0.037 | 0.00079 | 0.957 ± 0.189 | -0.150 ± 0.033 | 0.00045 | 1.016 ± 0.216 | -0.162 ± 0.040 | 0.00007 |
| L100 Aml5/H12.5 | 1.179 ± 0.243 | 0.030 ± 0.014 | 0.58062 | 1.194 ± 0.254 | 0.031 ± 0.017 | 0.90621 | 1.173 ± 0.223 | 0.030 ± 0.018 | 0.58062 | 1.178 ± 0.230 | 0.025 ± 0.017 | 0.96707 | 1.130 ± 0.224 | 0.022 ± 0.018 | 0.90621 | 1.194 ± 0.256 | 0.017 ± 0.014 | 0.99376 |
| L100 B5/H12.5 | 0.988 ± 0.206 | -0.161 ± 0.036 | 0.00007 | 1.002 ± 0.214 | -0.161 ± 0.038 | 0.00025 | 0.987 ± 0.189 | -0.156 ± 0.035 | 0.00013 | 0.998 ± 0.196 | -0.155 ± 0.038 | 0.00136 | 0.960 ± 0.191 | -0.148 ± 0.033 | 0.00045 | 1.017 ± 0.217 | -0.161 ± 0.040 | 0.00007 |

**Al300 =** aliskiren 300 mg; **Aml5** = amlodipine 5 mg; **B5** = bisoprolol 5 mg; **E20** = enalapril 20 mg; **H12.5** = hydrochlorothiazide 12.5 mg; **L100** = losartan 100 mg; **SS** = statistically significant ($P$ < 0.00001)

**Table S9.** *P*-values calculated using the Kolmogorov-Smirnov test for changes in systemic arterial elasticity in populations ($n$ = 100) with different ACE activity receiving the same regimens (case 1: $c_{ACE}$ = 7.0 h$^{-1}$, case 2: $c_{ACE}$ = 8.9 h$^{-1}$, case 3: $c_{ACE}$ = 10.8 h$^{-1}$, case 4: $c_{ACE}$ = 42.3 h$^{-1}$, case 5: $c_{ACE}$ = 54.1 h$^{-1}$, case 6: $c_{ACE}$ = 65.9 h$^{-1}$; $P$-value for case $i$ *vs.* case $j$ is denoted $P_{ij}$)

| **Regimens** | $P_{12}$ | $P_{13}$ | $P_{23}$ | $P_{14}$ | $P_{15}$ | $P_{16}$ | $P_{24}$ | $P_{25}$ | $P_{26}$ | $P_{34}$ | $P_{35}$ | $P_{36}$ | $P_{45}$ | $P_{46}$ | $P_{56}$ |
|---|---|---|---|---|---|---|---|---|---|---|---|---|---|---|---|
| Al300 | 0.46756 | 0.58062 | 0.58062 | SS | SS | SS | SS | SS | SS | SS | SS | SS | 0.01581 | SS | 0.00013 |
| E20 | 0.21055 | 0.15454 | 0.58062 | 0.00386 | 0.00025 | SS | 0.05410 | 0.00386 | SS | 0.05410 | 0.00386 | SS | 0.46756 | 0.00630 | 0.05410 |
| L100 | 0.21055 | 0.46756 | 0.58062 | 0.00013 | SS | SS | 0.00004 | SS | SS | SS | SS | SS | 0.03663 | SS | 0.00079 |
| Aml5 | 0.81275 | 0.28093 | 0.46756 | 0.81275 | 0.90621 | 0.58062 | 0.96707 | 0.90621 | 0.99376 | 0.36672 | 0.46756 | 0.46756 | 0.81275 | 0.99963 | 0.69937 |
| B5 | 0.69937 | 0.69937 | 0.81275 | 0.69937 | 0.69937 | 0.07832 | 0.69937 | 0.36672 | 0.36672 | 0.36672 | 0.81275 | 0.07832 | 0.36672 | 0.46756 | 0.01581 |
| H12.5 | 0.69937 | 0.96707 | 0.46756 | 0.69937 | 0.90621 | 0.69937 | 0.28093 | 0.21055 | 0.28093 | 0.81275 | 0.58062 | 0.46756 | 0.81275 | 0.90621 | 0.99376 |
| Al300<br>Aml5 | 0.81275 | 0.58062 | 0.58062 | 0.00002 | SS | SS | 0.00025 | SS | SS | 0.00136 | SS | SS | 0.11113 | 0.00002 | 0.00386 |
| Al300<br>B5 | 0.90621 | 0.69937 | 0.69937 | 0.90621 | 0.28093 | 0.96707 | 0.58062 | 0.05410 | 0.99963 | 0.90621 | 0.15454 | 0.69937 | 0.28093 | 0.81275 | 0.03663 |
| Al300<br>H12.5 | 0.46756 | 0.46756 | 0.28093 | 0.00079 | SS | SS | 0.00025 | SS | SS | 0.01581 | 0.00025 | SS | 0.15454 | 0.00079 | 0.15454 |
| E20<br>Aml5 | 0.46756 | 0.58062 | 0.96707 | 0.00045 | 0.00079 | 0.07832 | 0.01581 | 0.01008 | 0.36672 | 0.02431 | 0.00136 | 0.15454 | 0.58062 | 0.15454 | 0.07832 |
| E20<br>B5 | 0.90621 | 0.69937 | 0.69937 | 0.69937 | 0.15454 | 0.58062 | 0.69937 | 0.02431 | 0.90621 | 0.96707 | 0.07832 | 0.96707 | 0.15454 | 0.90621 | 0.05410 |
| E20<br>H12.5 | 0.96707 | 0.46756 | 0.58062 | 0.11113 | 0.15454 | 0.02431 | 0.28093 | 0.21055 | 0.01581 | 0.90621 | 0.46756 | 0.07832 | 0.36672 | 0.11113 | 0.15454 |
| L100<br>Aml5 | 0.81275 | 0.90621 | 0.81275 | 0.00386 | 0.00002 | SS | 0.00630 | 0.00004 | SS | 0.01008 | 0.00007 | SS | 0.11113 | 0.00045 | 0.01581 |
| L100<br>B5 | 0.90621 | 0.69937 | 0.69937 | 0.81275 | 0.21055 | 0.90621 | 0.58062 | 0.01581 | 0.96707 | 0.90621 | 0.07832 | 0.81275 | 0.21055 | 0.81275 | 0.05410 |
| L100<br>H12.5 | 0.58062 | 0.46756 | 0.69937 | 0.00386 | SS | SS | 0.00630 | SS | SS | 0.01581 | 0.00079 | SS | 0.21055 | 0.00386 | 0.21055 |
| Al300<br>Aml5/B5 | 0.90621 | 0.81275 | 0.81275 | 0.58062 | 0.11113 | 0.90621 | 0.46756 | 0.01008 | 0.90621 | 0.96707 | 0.07832 | 0.90621 | 0.21055 | 0.69937 | 0.02431 |
| Al300<br>Aml5/H12.5 | 0.69937 | 0.46756 | 0.69937 | 0.00136 | SS | SS | 0.00386 | 0.00002 | SS | 0.07832 | 0.00079 | SS | 0.21055 | 0.00386 | 0.05410 |
| Al300<br>B5/H12.5 | 0.99376 | 0.90621 | 0.69937 | 0.46756 | 0.11113 | 0.58062 | 0.46756 | 0.05410 | 0.90621 | 0.90621 | 0.21055 | 0.36672 | 0.36672 | 0.36672 | 0.02431 |
| E20<br>Aml5/B5 | 0.90621 | 0.90621 | 0.69937 | 0.28093 | 0.01581 | 0.28093 | 0.21055 | 0.00136 | 0.46756 | 0.69937 | 0.00386 | 0.81275 | 0.11113 | 0.99376 | 0.03663 |
| E20<br>Aml5/H12.5 | 0.46756 | 0.36672 | 0.58062 | 0.01581 | 0.03663 | 0.36672 | 0.11113 | 0.07832 | 0.90621 | 0.46756 | 0.69937 | 0.69937 | 0.58062 | 0.07832 | 0.11113 |
| E20<br>B5/H12.5 | 0.96707 | 0.96707 | 0.69937 | 0.15454 | 0.00630 | 0.36672 | 0.11113 | 0.01008 | 0.58062 | 0.58062 | 0.03663 | 0.81275 | 0.36672 | 0.69937 | 0.02431 |
| L100<br>Aml5/B5 | 0.90621 | 0.90621 | 0.81275 | 0.58062 | 0.03663 | 0.58062 | 0.46756 | 0.00630 | 0.90621 | 0.90621 | 0.03663 | 0.96707 | 0.21055 | 0.81275 | 0.02431 |
| L100<br>Aml5/H12.5 | 0.69937 | 0.58062 | 0.69937 | 0.00630 | 0.00004 | SS | 0.03663 | 0.00013 | SS | 0.15454 | 0.00232 | SS | 0.28093 | 0.00386 | 0.05410 |
| L100<br>B5/H12.5 | 0.99376 | 0.81275 | 0.69937 | 0.21055 | 0.07832 | 0.81275 | 0.46756 | 0.03663 | 0.96707 | 0.90621 | 0.21055 | 0.81275 | 0.36672 | 0.46756 | 0.03663 |

**Al300 =** aliskiren 300 mg; **Aml5** = amlodipine 5 mg; **B5** = bisoprolol 5 mg; **E20** = enalapril 20 mg; **H12.5** = hydrochlorothiazide 12.5 mg; **L100** = losartan 100 mg; **SS** = statistically significant ($P$ < 0.00001)

**Figure S12.** Simulated change in systemic vascular resistance from baseline to week 4 (mean ± SD, $n$ = 100)

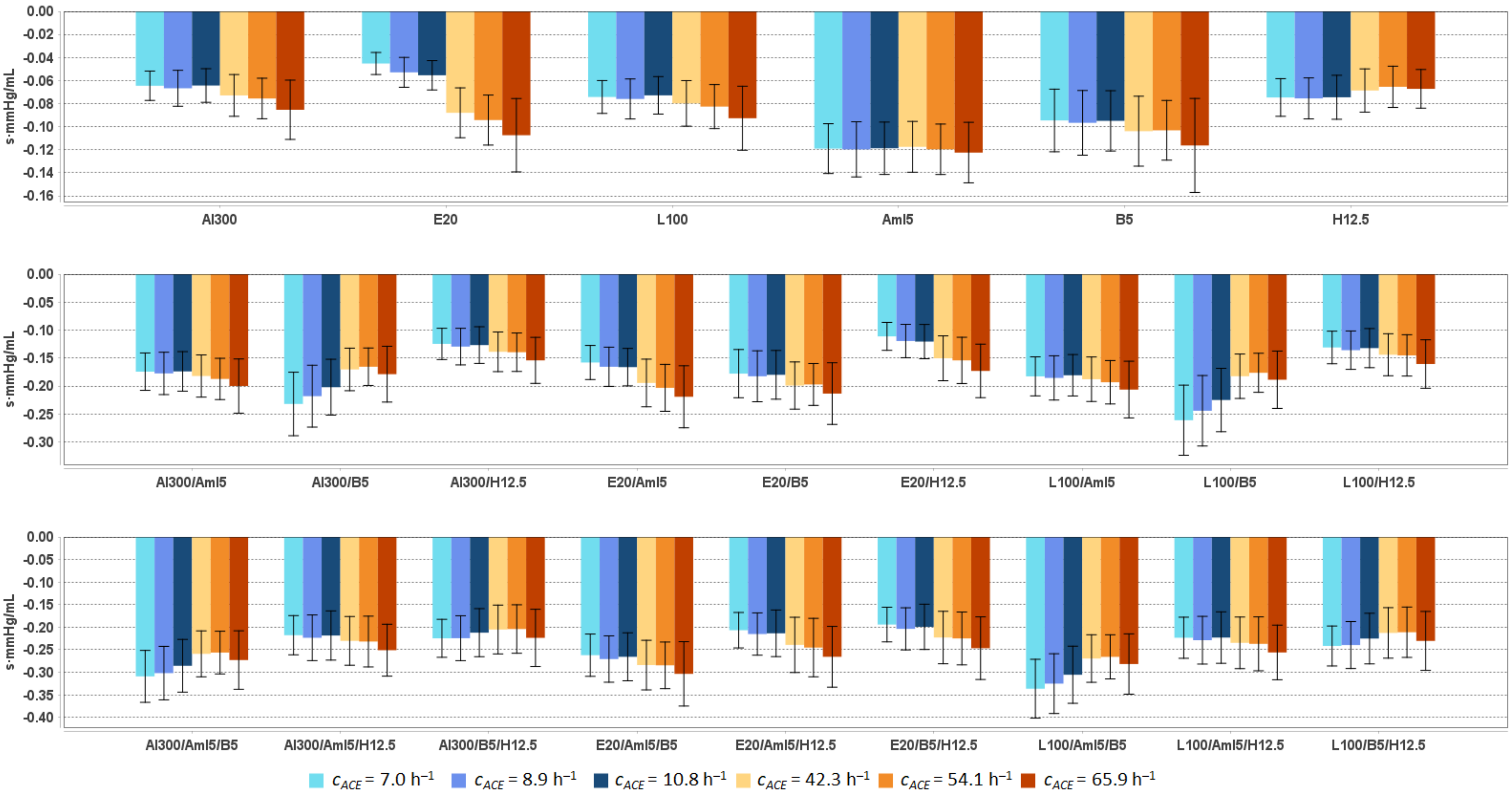


**Al300** = aliskiren 300 mg; **Aml5** = amlodipine 5 mg; **B5** = bisoprolol 5 mg; **E20** = enalapril 20 mg; **H12.5** = hydrochlorothiazide 12.5 mg; **L100** = losartan 100 mg

**Table S10.** Simulated response of <u>systemic vascular resistance</u> to antihypertensive therapy in virtual hypertensive populations ($n$ = 100) with different ACE activity, including $P$-values (Kolmogorov-Smirnov test) for endpoint *vs*. baseline; data are presented as mean ± SD in s·mmHg/mL

| **Regimens** | $c_{ACE}$ = 7.0 $h^{-1}$ | | | $c_{ACE}$ = 8.9 $h^{-1}$ | | | $c_{ACE}$ = 10.8 $h^{-1}$ | | | $c_{ACE}$ = 42.3 $h^{-1}$ | | | $c_{ACE}$ = 54.1 $h^{-1}$ | | | $c_{ACE}$ = 65.9 $h^{-1}$ | | |
|---|---|---|---|---|---|---|---|---|---|---|---|---|---|---|---|---|---|---|
| | **Value** | **Change** | ***P*** | **Value** | **Change** | ***P*** | **Value** | **Change** | ***P*** | **Value** | **Change** | ***P*** | **Value** | **Change** | ***P*** | **Value** | **Change** | ***P*** |
| Baseline | 1.39 ± 0.20 | – | – | 1.44 ± 0.22 | – | – | 1.42 ± 0.23 | – | – | 1.43 ± 0.22 | – | – | 1.41 ± 0.20 | – | – | 1.45 ± 0.25 | – | – |
| Al300 | 1.33 ± 0.19 | -0.06 ± 0.01 | 0.07832 | 1.37 ± 0.20 | -0.07 ± 0.02 | 0.03663 | 1.36 ± 0.22 | -0.06 ± 0.01 | 0.28093 | 1.36 ± 0.21 | -0.07 ± 0.02 | 0.05410 | 1.33 ± 0.19 | -0.08 ± 0.02 | 0.02431 | 1.37 ± 0.22 | -0.09 ± 0.03 | 0.02431 |
| E20 | 1.35 ± 0.19 | -0.05 ± 0.01 | 0.36672 | 1.38 ± 0.21 | -0.05 ± 0.01 | 0.11113 | 1.36 ± 0.22 | -0.06 ± 0.01 | 0.28093 | 1.34 ± 0.20 | -0.09 ± 0.02 | 0.02431 | 1.31 ± 0.18 | -0.09 ± 0.02 | 0.01581 | 1.35 ± 0.22 | -0.11 ± 0.03 | 0.00386 |
| L100 | 1.32 ± 0.19 | -0.07 ± 0.01 | 0.02431 | 1.36 ± 0.20 | -0.08 ± 0.02 | 0.01581 | 1.35 ± 0.22 | -0.07 ± 0.02 | 0.15454 | 1.35 ± 0.21 | -0.08 ± 0.02 | 0.03663 | 1.32 ± 0.19 | -0.08 ± 0.02 | 0.01581 | 1.36 ± 0.22 | -0.09 ± 0.03 | 0.01008 |
| Aml5 | 1.27 ± 0.18 | -0.12 ± 0.02 | 0.00025 | 1.32 ± 0.20 | -0.12 ± 0.02 | 0.00136 | 1.30 ± 0.21 | -0.12 ± 0.02 | 0.00232 | 1.31 ± 0.21 | -0.12 ± 0.02 | 0.00079 | 1.29 ± 0.18 | -0.12 ± 0.02 | 0.00136 | 1.33 ± 0.22 | -0.12 ± 0.03 | 0.00079 |
| B5 | 1.30 ± 0.18 | -0.09 ± 0.03 | 0.00630 | 1.34 ± 0.19 | -0.10 ± 0.03 | 0.00630 | 1.33 ± 0.21 | -0.09 ± 0.03 | 0.02431 | 1.33 ± 0.20 | -0.10 ± 0.03 | 0.00232 | 1.30 ± 0.18 | -0.10 ± 0.03 | 0.01008 | 1.34 ± 0.21 | -0.12 ± 0.04 | 0.00136 |
| H12.5 | 1.32 ± 0.19 | -0.07 ± 0.02 | 0.02431 | 1.36 ± 0.20 | -0.08 ± 0.02 | 0.02431 | 1.35 ± 0.21 | -0.07 ± 0.02 | 0.11113 | 1.36 ± 0.21 | -0.07 ± 0.02 | 0.11113 | 1.34 ± 0.19 | -0.07 ± 0.02 | 0.05410 | 1.39 ± 0.23 | -0.07 ± 0.02 | 0.15454 |
| Al300 Aml5 | 1.22 ± 0.17 | -0.17 ± 0.03 | SS | 1.26 ± 0.19 | -0.18 ± 0.04 | SS | 1.25 ± 0.20 | -0.17 ± 0.04 | 0.00002 | 1.25 ± 0.19 | -0.18 ± 0.04 | SS | 1.22 ± 0.17 | -0.19 ± 0.04 | SS | 1.25 ± 0.20 | -0.20 ± 0.05 | SS |
| Al300 B5 | 1.16 ± 0.16 | -0.23 ± 0.06 | SS | 1.22 ± 0.18 | -0.22 ± 0.06 | SS | 1.22 ± 0.19 | -0.20 ± 0.05 | SS | 1.26 ± 0.19 | -0.17 ± 0.04 | SS | 1.24 ± 0.17 | -0.17 ± 0.03 | SS | 1.28 ± 0.20 | -0.18 ± 0.05 | SS |
| Al300 H12.5 | 1.27 ± 0.18 | -0.12 ± 0.03 | 0.00004 | 1.31 ± 0.19 | -0.13 ± 0.03 | 0.00025 | 1.29 ± 0.21 | -0.13 ± 0.03 | 0.00232 | 1.29 ± 0.19 | -0.14 ± 0.04 | 0.00002 | 1.27 ± 0.18 | -0.14 ± 0.03 | 0.00025 | 1.30 ± 0.21 | -0.15 ± 0.04 | 0.00004 |
| E20 Aml5 | 1.24 ± 0.18 | -0.16 ± 0.03 | SS | 1.27 ± 0.19 | -0.17 ± 0.04 | 0.00002 | 1.25 ± 0.20 | -0.17 ± 0.03 | 0.00004 | 1.24 ± 0.19 | -0.19 ± 0.04 | SS | 1.20 ± 0.17 | -0.20 ± 0.04 | SS | 1.24 ± 0.20 | -0.22 ± 0.06 | SS |
| E20 B5 | 1.22 ± 0.17 | -0.18 ± 0.04 | SS | 1.25 ± 0.18 | -0.18 ± 0.05 | SS | 1.24 ± 0.19 | -0.18 ± 0.04 | SS | 1.23 ± 0.18 | -0.20 ± 0.04 | SS | 1.21 ± 0.17 | -0.20 ± 0.04 | SS | 1.24 ± 0.19 | -0.21 ± 0.06 | SS |
| E20 H12.5 | 1.28 ± 0.18 | -0.11 ± 0.02 | 0.00136 | 1.32 ± 0.19 | -0.12 ± 0.03 | 0.00079 | 1.30 ± 0.21 | -0.12 ± 0.03 | 0.00386 | 1.28 ± 0.19 | -0.15 ± 0.04 | SS | 1.25 ± 0.18 | -0.15 ± 0.04 | 0.00002 | 1.28 ± 0.20 | -0.17 ± 0.05 | 0.00002 |
| L100 Aml5 | 1.21 ± 0.17 | -0.18 ± 0.03 | SS | 1.25 ± 0.18 | -0.19 ± 0.04 | SS | 1.24 ± 0.20 | -0.18 ± 0.04 | 0.00002 | 1.24 ± 0.19 | -0.19 ± 0.04 | SS | 1.21 ± 0.17 | -0.19 ± 0.04 | SS | 1.25 ± 0.20 | -0.21 ± 0.05 | SS |
| L100 B5 | 1.13 ± 0.16 | -0.26 ± 0.06 | SS | 1.19 ± 0.18 | -0.24 ± 0.06 | SS | 1.20 ± 0.19 | -0.22 ± 0.06 | SS | 1.25 ± 0.19 | -0.18 ± 0.04 | SS | 1.23 ± 0.17 | -0.18 ± 0.03 | SS | 1.27 ± 0.20 | -0.19 ± 0.05 | SS |
| L100 H12.5 | 1.26 ± 0.18 | -0.13 ± 0.03 | 0.00002 | 1.30 ± 0.19 | -0.14 ± 0.03 | 0.00025 | 1.29 ± 0.21 | -0.13 ± 0.03 | 0.00136 | 1.29 ± 0.19 | -0.14 ± 0.04 | SS | 1.26 ± 0.18 | -0.15 ± 0.04 | 0.00013 | 1.29 ± 0.21 | -0.16 ± 0.04 | 0.00004 |
| Al300 Aml5/B5 | 1.08 ± 0.15 | -0.31 ± 0.06 | SS | 1.13 ± 0.17 | -0.30 ± 0.06 | SS | 1.13 ± 0.18 | -0.29 ± 0.06 | SS | 1.17 ± 0.18 | -0.26 ± 0.05 | SS | 1.15 ± 0.16 | -0.26 ± 0.05 | SS | 1.18 ± 0.19 | -0.27 ± 0.06 | SS |
| Al300 Aml5/H12.5 | 1.18 ± 0.17 | -0.22 ± 0.04 | SS | 1.21 ± 0.18 | -0.22 ± 0.05 | SS | 1.20 ± 0.20 | -0.22 ± 0.05 | SS | 1.20 ± 0.19 | -0.23 ± 0.05 | SS | 1.17 ± 0.17 | -0.23 ± 0.06 | SS | 1.20 ± 0.20 | -0.25 ± 0.06 | SS |
| Al300 B5/H12.5 | 1.17 ± 0.16 | -0.22 ± 0.04 | SS | 1.21 ± 0.18 | -0.22 ± 0.05 | SS | 1.21 ± 0.19 | -0.21 ± 0.05 | SS | 1.23 ± 0.18 | -0.21 ± 0.05 | SS | 1.20 ± 0.17 | -0.20 ± 0.05 | SS | 1.23 ± 0.19 | -0.22 ± 0.06 | SS |
| E20 Aml5/B5 | 1.13 ± 0.16 | -0.26 ± 0.05 | SS | 1.16 ± 0.17 | -0.27 ± 0.05 | SS | 1.15 ± 0.18 | -0.27 ± 0.05 | SS | 1.15 ± 0.17 | -0.28 ± 0.05 | SS | 1.12 ± 0.16 | -0.28 ± 0.05 | SS | 1.15 ± 0.18 | -0.30 ± 0.07 | SS |
| E20 Aml5/H12.5 | 1.19 ± 0.17 | -0.21 ± 0.04 | SS | 1.22 ± 0.18 | -0.22 ± 0.05 | SS | 1.21 ± 0.20 | -0.21 ± 0.05 | SS | 1.19 ± 0.19 | -0.24 ± 0.06 | SS | 1.16 ± 0.17 | -0.25 ± 0.06 | SS | 1.19 ± 0.20 | -0.27 ± 0.07 | SS |
| E20 B5/H12.5 | 1.20 ± 0.17 | -0.19 ± 0.04 | SS | 1.23 ± 0.18 | -0.20 ± 0.05 | SS | 1.22 ± 0.19 | -0.20 ± 0.05 | SS | 1.21 ± 0.18 | -0.22 ± 0.06 | SS | 1.18 ± 0.17 | -0.22 ± 0.06 | SS | 1.21 ± 0.19 | -0.25 ± 0.07 | SS |
| L100 Aml5/B5 | 1.06 ± 0.15 | -0.34 ± 0.07 | SS | 1.11 ± 0.17 | -0.33 ± 0.07 | SS | 1.11 ± 0.18 | -0.31 ± 0.06 | SS | 1.16 ± 0.18 | -0.27 ± 0.05 | SS | 1.14 ± 0.16 | -0.27 ± 0.05 | SS | 1.17 ± 0.19 | -0.28 ± 0.07 | SS |
| L100 Aml5/H12.5 | 1.17 ± 0.17 | -0.22 ± 0.05 | SS | 1.21 ± 0.18 | -0.23 ± 0.05 | SS | 1.20 ± 0.20 | -0.22 ± 0.06 | SS | 1.20 ± 0.19 | -0.23 ± 0.06 | SS | 1.17 ± 0.17 | -0.24 ± 0.06 | SS | 1.20 ± 0.20 | -0.26 ± 0.06 | SS |
| L100 B5/H12.5 | 1.15 ± 0.16 | -0.24 ± 0.04 | SS | 1.20 ± 0.18 | -0.24 ± 0.05 | SS | 1.20 ± 0.19 | -0.23 ± 0.06 | SS | 1.22 ± 0.18 | -0.21 ± 0.06 | SS | 1.19 ± 0.17 | -0.21 ± 0.06 | SS | 1.22 ± 0.19 | -0.23 ± 0.07 | SS |

**Al300 =** aliskiren 300 mg; **Aml5** = amlodipine 5 mg; **B5** = bisoprolol 5 mg; **E20** = enalapril 20 mg; **H12.5** = hydrochlorothiazide 12.5 mg; **L100** = losartan 100 mg; **SS** = statistically significant ($P$ < 0.00001)

**Table S11.** *P*-values calculated using the Kolmogorov-Smirnov test for changes in systemic vascular resistance in populations ($n$ = 100) with different ACE activity receiving the same regimens (case 1: $c_{ACE}$ = 7.0 $h^{-1}$, case 2: $c_{ACE}$ = 8.9 $h^{-1}$, case 3: $c_{ACE}$ = 10.8 $h^{-1}$, case 4: $c_{ACE}$ = 42.3 $h^{-1}$, case 5: $c_{ACE}$ = 54.1 $h^{-1}$, case 6: $c_{ACE}$ = 65.9 $h^{-1}$; *P*-value for case *i vs.* case *j* is denoted $P_{ij}$)

| Regimens | $P_{12}$ | $P_{13}$ | $P_{23}$ | $P_{14}$ | $P_{15}$ | $P_{16}$ | $P_{24}$ | $P_{25}$ | $P_{26}$ | $P_{34}$ | $P_{35}$ | $P_{36}$ | $P_{45}$ | $P_{46}$ | $P_{56}$ |
|---|---|---|---|---|---|---|---|---|---|---|---|---|---|---|---|
| Al300 | 0.21055 | 0.69937 | 0.46756 | 0.00079 | 0.00004 | SS | 0.03663 | 0.01581 | SS | 0.00630 | 0.00079 | SS | 0.69937 | 0.00136 | 0.01581 |
| E20 | 0.00013 | SS | 0.36672 | SS | SS | SS | SS | SS | SS | SS | SS | SS | 0.15454 | 0.00007 | 0.01581 |
| L100 | 0.46756 | 0.36672 | 0.58062 | 0.01581 | 0.00630 | SS | 0.28093 | 0.15454 | 0.00007 | 0.01581 | 0.00630 | SS | 0.58062 | 0.00232 | 0.01008 |
| Aml5 | 0.81275 | 0.90621 | 0.99376 | 0.99376 | 0.69937 | 0.11113 | 0.90621 | 0.90621 | 0.58062 | 0.58062 | 0.90621 | 0.58062 | 0.36672 | 0.07832 | 0.21055 |
| B5 | 0.58062 | 0.46756 | 0.96707 | 0.02431 | 0.01008 | SS | 0.07832 | 0.21055 | 0.00079 | 0.03663 | 0.07832 | 0.00079 | 0.96707 | 0.02431 | 0.07832 |
| H12.5 | 0.69937 | 0.46756 | 0.90621 | 0.01008 | 0.00079 | 0.01581 | 0.01581 | 0.00630 | 0.03663 | 0.02431 | 0.00232 | 0.11113 | 0.28093 | 0.81275 | 0.36672 |
| Al300<br>Aml5 | 0.90621 | 0.46756 | 0.81275 | 0.15454 | 0.01008 | 0.00025 | 0.46756 | 0.11113 | 0.00232 | 0.28093 | 0.11113 | 0.00079 | 0.28093 | 0.01581 | 0.05410 |
| Al300<br>B5 | 0.07832 | 0.00079 | 0.15454 | SS | SS | SS | SS | SS | 0.00045 | 0.00013 | 0.00007 | 0.01581 | 0.58062 | 0.07832 | 0.05410 |
| Al300<br>H12.5 | 0.21055 | 0.15454 | 0.46756 | 0.00386 | 0.01581 | SS | 0.07832 | 0.28093 | 0.00136 | 0.07832 | 0.05410 | 0.00002 | 0.90621 | 0.00630 | 0.03663 |
| E20<br>Aml5 | 0.21055 | 0.05410 | 0.90621 | SS | SS | SS | SS | SS | SS | 0.00004 | SS | SS | 0.21055 | 0.00630 | 0.01581 |
| E20<br>B5 | 0.15454 | 0.36672 | 0.99376 | 0.00002 | 0.00007 | SS | 0.01581 | 0.02431 | 0.00232 | 0.00386 | 0.00386 | 0.00013 | 0.90621 | 0.05410 | 0.03663 |
| E20<br>H12.5 | 0.07832 | 0.00630 | 0.81275 | SS | SS | SS | SS | SS | SS | SS | SS | SS | 0.58062 | 0.00232 | 0.02431 |
| L100<br>Aml5 | 0.90621 | 0.58062 | 0.81275 | 0.36672 | 0.03663 | 0.00045 | 0.81275 | 0.21055 | 0.00630 | 0.36672 | 0.15454 | 0.00079 | 0.36672 | 0.01581 | 0.03663 |
| L100<br>B5 | 0.05410 | 0.00025 | 0.21055 | SS | SS | SS | SS | SS | SS | SS | SS | 0.00136 | 0.46756 | 0.21055 | 0.07832 |
| L100<br>H12.5 | 0.21055 | 0.28093 | 0.46756 | 0.01581 | 0.02431 | SS | 0.15454 | 0.36672 | 0.00232 | 0.11113 | 0.05410 | 0.00004 | 0.90621 | 0.01008 | 0.02431 |
| Al300<br>Aml5/B5 | 0.21055 | 0.00630 | 0.15454 | SS | SS | SS | SS | SS | 0.00386 | 0.00386 | 0.00630 | 0.07832 | 0.81275 | 0.03663 | 0.05410 |
| Al300<br>Aml5/H12.5 | 0.28093 | 0.28093 | 0.69937 | 0.15454 | 0.07832 | 0.00013 | 0.28093 | 0.36672 | 0.03663 | 0.21055 | 0.36672 | 0.00232 | 0.69937 | 0.01581 | 0.11113 |
| Al300<br>B5/H12.5 | 0.28093 | 0.11113 | 0.21055 | 0.01008 | 0.00386 | 0.02431 | 0.02431 | 0.02431 | 0.58062 | 0.28093 | 0.11113 | 0.21055 | 0.81275 | 0.01581 | 0.05410 |
| E20<br>Aml5/B5 | 0.07832 | 0.58062 | 0.46756 | 0.00232 | 0.00386 | 0.00025 | 0.11113 | 0.36672 | 0.00232 | 0.07832 | 0.11113 | 0.00232 | 0.81275 | 0.02431 | 0.03663 |
| E20<br>Aml5/H12.5 | 0.07832 | 0.03663 | 0.96707 | SS | SS | SS | 0.00386 | 0.00136 | 0.00002 | 0.01581 | 0.00232 | SS | 0.81275 | 0.00630 | 0.11113 |
| E20<br>B5/H12.5 | 0.11113 | 0.28093 | 0.69937 | 0.00045 | 0.00025 | SS | 0.01581 | 0.05410 | 0.00013 | 0.02431 | 0.01581 | 0.00007 | 0.81275 | 0.02431 | 0.03663 |
| L100<br>Aml5/B5 | 0.15454 | 0.00386 | 0.15454 | SS | SS | SS | SS | SS | 0.00045 | 0.00079 | 0.00079 | 0.01581 | 0.81275 | 0.05410 | 0.07832 |
| L100<br>Aml5/H12.5 | 0.36672 | 0.28093 | 0.69937 | 0.15454 | 0.11113 | 0.00079 | 0.46756 | 0.58062 | 0.05410 | 0.36672 | 0.36672 | 0.00232 | 0.81275 | 0.01008 | 0.11113 |
| L100<br>B5/H12.5 | 0.36672 | 0.07832 | 0.15454 | 0.00045 | 0.00025 | 0.00232 | 0.00386 | 0.00630 | 0.21055 | 0.07832 | 0.03663 | 0.36672 | 0.90621 | 0.03663 | 0.05410 |

**Al300 =** aliskiren 300 mg; **Aml5** = amlodipine 5 mg; **B5** = bisoprolol 5 mg; **E20** = enalapril 20 mg; **H12.5** = hydrochlorothiazide 12.5 mg; **L100** = losartan 100 mg; **SS** = statistically significant ($P$ < 0.00001)

**Figure S13.** Simulated change in diastolic pulmonary arterial pressure from baseline to week 4 (mean ± SD, $n$ = 100)

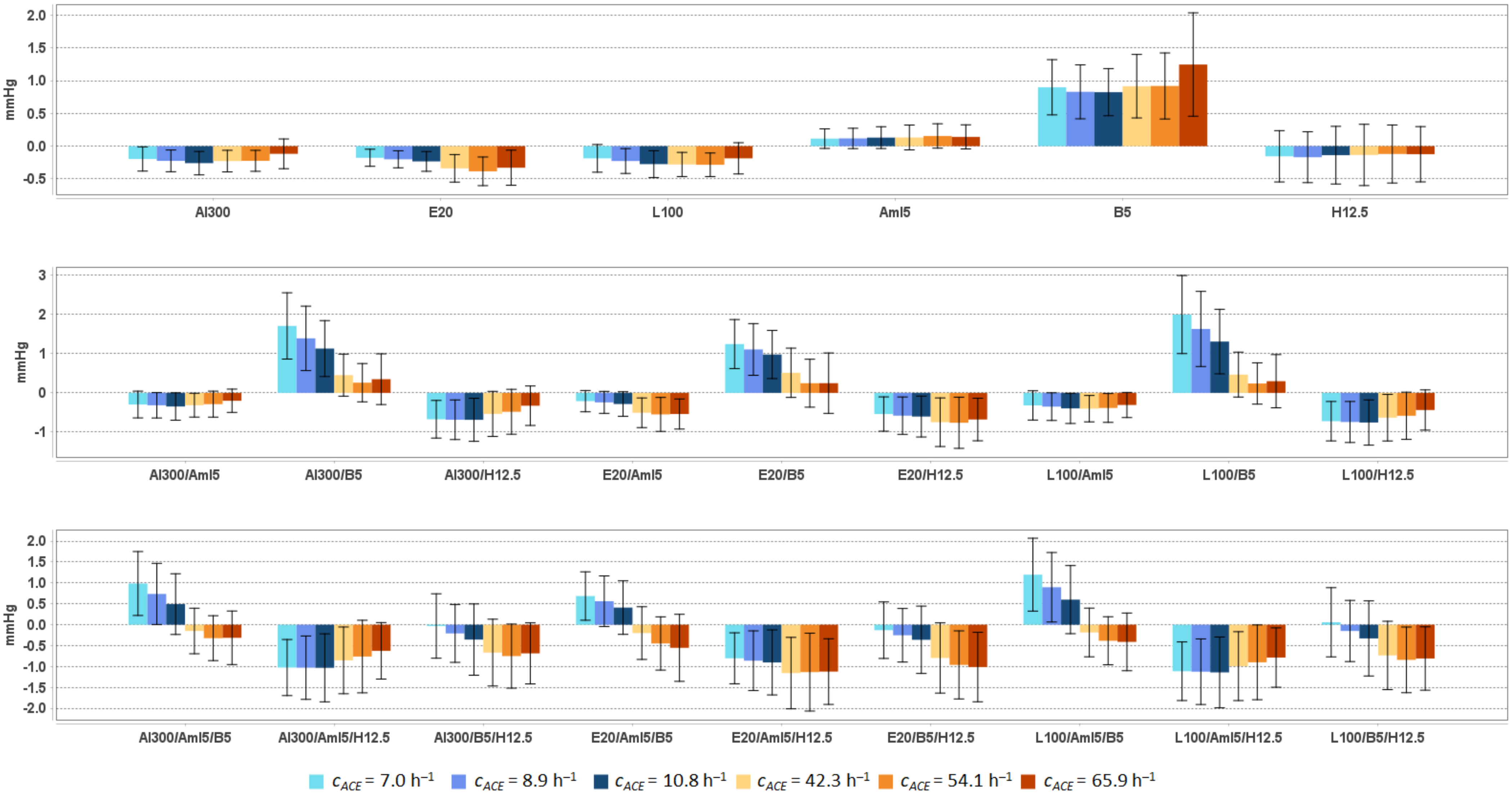


**Al300** = aliskiren 300 mg; **Aml5** = amlodipine 5 mg; **B5** = bisoprolol 5 mg; **E20** = enalapril 20 mg; **H12.5** = hydrochlorothiazide 12.5 mg; **L100** = losartan 100 mg

**Table S12.** Simulated response of <u>diastolic pulmonary arterial pressure</u> to antihypertensive therapy in virtual hypertensive populations (*n* = 100) with different ACE activity, including *P*-values (Kolmogorov-Smirnov test) for endpoint *vs*. baseline; data are presented as mean ± SD in mmHg

| Regimens | $c_{ACE}$ = 7.0 h$^{-1}$ | | | $c_{ACE}$ = 8.9 h$^{-1}$ | | | $c_{ACE}$ = 10.8 h$^{-1}$ | | | $c_{ACE}$ = 42.3 h$^{-1}$ | | | $c_{ACE}$ = 54.1 h$^{-1}$ | | | $c_{ACE}$ = 65.9 h$^{-1}$ | | |
|---|---|---|---|---|---|---|---|---|---|---|---|---|---|---|---|---|---|---|
| | **Value** | **Change** | ***P*** | **Value** | **Change** | ***P*** | **Value** | **Change** | ***P*** | **Value** | **Change** | ***P*** | **Value** | **Change** | ***P*** | **Value** | **Change** | ***P*** |
| Baseline | 11.3 ± 0.6 | – | – | 11.3 ± 0.5 | – | – | 11.2 ± 0.7 | – | – | 11.2 ± 0.6 | – | – | 11.1 ± 0.7 | – | – | 11.3 ± 0.6 | – | – |
| Al300 | 11.1 ± 0.6 | -0.2 ± 0.2 | 0.01008 | 11.0 ± 0.5 | -0.2 ± 0.2 | 0.00386 | 10.9 ± 0.7 | -0.3 ± 0.2 | 0.00232 | 11.0 ± 0.6 | -0.2 ± 0.2 | 0.00232 | 10.9 ± 0.7 | -0.2 ± 0.2 | 0.01008 | 11.1 ± 0.7 | -0.1 ± 0.2 | 0.15454 |
| E20 | 11.1 ± 0.6 | -0.2 ± 0.1 | 0.03663 | 11.1 ± 0.5 | -0.2 ± 0.1 | 0.01581 | 11.0 ± 0.7 | -0.2 ± 0.2 | 0.00630 | 10.9 ± 0.6 | -0.3 ± 0.2 | 0.00004 | 10.7 ± 0.7 | -0.4 ± 0.2 | 0.00013 | 10.9 ± 0.7 | -0.3 ± 0.3 | SS |
| L100 | 11.1 ± 0.6 | -0.2 ± 0.2 | 0.01581 | 11.0 ± 0.5 | -0.2 ± 0.2 | 0.00630 | 10.9 ± 0.7 | -0.3 ± 0.2 | 0.00136 | 10.9 ± 0.6 | -0.3 ± 0.2 | 0.00045 | 10.8 ± 0.7 | -0.3 ± 0.2 | 0.00232 | 11.1 ± 0.7 | -0.2 ± 0.2 | 0.00630 |
| Aml5 | 11.4 ± 0.6 | 0.1 ± 0.1 | 0.21055 | 11.4 ± 0.5 | 0.1 ± 0.2 | 0.15454 | 11.3 ± 0.7 | 0.1 ± 0.2 | 0.05410 | 11.4 ± 0.6 | 0.1 ± 0.2 | 0.11113 | 11.3 ± 0.7 | 0.2 ± 0.2 | 0.11113 | 11.4 ± 0.6 | 0.1 ± 0.2 | 0.21055 |
| B5 | 12.2 ± 0.8 | 0.9 ± 0.4 | SS | 12.1 ± 0.7 | 0.8 ± 0.4 | SS | 12.0 ± 0.9 | 0.8 ± 0.4 | SS | 12.1 ± 0.9 | 0.9 ± 0.5 | SS | 12.0 ± 0.9 | 0.9 ± 0.5 | SS | 12.5 ± 1.1 | 1.2 ± 0.8 | SS |
| H12.5 | 11.2 ± 0.7 | -0.2 ± 0.4 | 0.07832 | 11.1 ± 0.6 | -0.2 ± 0.4 | 0.05410 | 11.1 ± 0.8 | -0.1 ± 0.4 | 0.15454 | 11.1 ± 0.7 | -0.1 ± 0.5 | 0.07832 | 11.0 ± 0.8 | -0.1 ± 0.4 | 0.11113 | 11.1 ± 0.7 | -0.1 ± 0.4 | 0.15454 |
| Al300<br>Aml5 | 11.0 ± 0.6 | -0.3 ± 0.3 | 0.00013 | 11.0 ± 0.6 | -0.3 ± 0.3 | 0.00013 | 10.8 ± 0.7 | -0.3 ± 0.3 | 0.00045 | 10.9 ± 0.6 | -0.3 ± 0.3 | 0.00045 | 10.8 ± 0.7 | -0.3 ± 0.3 | 0.00232 | 11.1 ± 0.7 | -0.2 ± 0.3 | 0.00136 |
| Al300<br>B5 | 13.0 ± 1.2 | 1.7 ± 0.8 | SS | 12.7 ± 1.1 | 1.4 ± 0.8 | SS | 12.3 ± 1.1 | 1.1 ± 0.7 | SS | 11.7 ± 0.9 | 0.5 ± 0.5 | SS | 11.4 ± 0.9 | 0.3 ± 0.5 | 0.00136 | 11.6 ± 1.0 | 0.3 ± 0.6 | SS |
| Al300<br>H12.5 | 10.6 ± 0.7 | -0.7 ± 0.5 | SS | 10.6 ± 0.7 | -0.7 ± 0.5 | SS | 10.5 ± 0.8 | -0.7 ± 0.5 | SS | 10.7 ± 0.8 | -0.5 ± 0.6 | SS | 10.6 ± 0.9 | -0.5 ± 0.6 | SS | 10.9 ± 0.8 | -0.3 ± 0.5 | 0.00136 |
| E20<br>Aml5 | 11.1 ± 0.6 | -0.2 ± 0.3 | 0.00386 | 11.0 ± 0.5 | -0.2 ± 0.3 | 0.00136 | 10.9 ± 0.7 | -0.3 ± 0.3 | 0.00386 | 10.7 ± 0.7 | -0.5 ± 0.4 | SS | 10.6 ± 0.7 | -0.5 ± 0.4 | SS | 10.7 ± 0.7 | -0.5 ± 0.4 | SS |
| E20<br>B5 | 12.6 ± 1.0 | 1.2 ± 0.6 | SS | 12.4 ± 0.9 | 1.1 ± 0.7 | SS | 12.2 ± 1.1 | 1.0 ± 0.6 | SS | 11.7 ± 0.9 | 0.5 ± 0.6 | SS | 11.4 ± 1.0 | 0.2 ± 0.6 | 0.00079 | 11.5 ± 1.0 | 0.2 ± 0.8 | 0.00025 |
| E20<br>H12.5 | 10.8 ± 0.7 | -0.5 ± 0.4 | SS | 10.7 ± 0.6 | -0.6 ± 0.5 | SS | 10.6 ± 0.8 | -0.6 ± 0.5 | SS | 10.5 ± 0.8 | -0.7 ± 0.6 | SS | 10.4 ± 0.9 | -0.8 ± 0.7 | SS | 10.6 ± 0.8 | -0.7 ± 0.5 | SS |
| L100<br>Aml5 | 11.0 ± 0.7 | -0.3 ± 0.4 | 0.00013 | 10.9 ± 0.6 | -0.3 ± 0.4 | 0.00004 | 10.8 ± 0.7 | -0.4 ± 0.4 | 0.00007 | 10.8 ± 0.6 | -0.4 ± 0.3 | 0.00004 | 10.7 ± 0.7 | -0.4 ± 0.4 | 0.00013 | 10.9 ± 0.7 | -0.3 ± 0.3 | 0.00004 |
| L100<br>B5 | 13.3 ± 1.3 | 2.0 ± 1.0 | SS | 12.9 ± 1.2 | 1.6 ± 1.0 | SS | 12.5 ± 1.2 | 1.3 ± 0.8 | SS | 11.7 ± 0.9 | 0.5 ± 0.6 | SS | 11.4 ± 0.9 | 0.2 ± 0.5 | 0.00079 | 11.6 ± 1.0 | 0.3 ± 0.7 | 0.00004 |
| L100<br>H12.5 | 10.6 ± 0.7 | -0.7 ± 0.5 | SS | 10.5 ± 0.7 | -0.7 ± 0.5 | SS | 10.4 ± 0.8 | -0.8 ± 0.6 | SS | 10.6 ± 0.8 | -0.6 ± 0.6 | SS | 10.5 ± 0.9 | -0.6 ± 0.6 | SS | 10.8 ± 0.8 | -0.4 ± 0.5 | SS |
| Al300<br>Aml5/B5 | 12.3 ± 1.1 | 1.0 ± 0.8 | SS | 12.0 ± 1.0 | 0.7 ± 0.7 | SS | 11.7 ± 1.1 | 0.5 ± 0.7 | SS | 11.1 ± 0.8 | -0.1 ± 0.5 | 0.05410 | 10.8 ± 0.8 | -0.3 ± 0.5 | 0.00045 | 10.9 ± 0.9 | -0.3 ± 0.6 | 0.00079 |
| Al300<br>Aml5/H12.5 | 10.3 ± 0.8 | -1.0 ± 0.7 | SS | 10.2 ± 0.8 | -1.0 ± 0.8 | SS | 10.2 ± 0.9 | -1.0 ± 0.8 | SS | 10.4 ± 0.9 | -0.9 ± 0.8 | SS | 10.4 ± 1.0 | -0.8 ± 0.9 | SS | 10.6 ± 0.8 | -0.6 ± 0.7 | SS |
| Al300<br>B5/H12.5 | 11.3 ± 1.0 | -0.0 ± 0.8 | 0.02431 | 11.1 ± 0.9 | -0.2 ± 0.7 | 0.00630 | 10.8 ± 1.1 | -0.4 ± 0.9 | 0.00025 | 10.6 ± 0.9 | -0.7 ± 0.8 | SS | 10.4 ± 1.0 | -0.8 ± 0.8 | SS | 10.6 ± 0.9 | -0.7 ± 0.7 | SS |
| E20<br>Aml5/B5 | 12.0 ± 0.9 | 0.7 ± 0.6 | SS | 11.8 ± 0.8 | 0.6 ± 0.6 | SS | 11.6 ± 1.0 | 0.4 ± 0.6 | SS | 11.0 ± 0.9 | -0.2 ± 0.6 | 0.05410 | 10.7 ± 0.9 | -0.4 ± 0.6 | SS | 10.7 ± 1.0 | -0.6 ± 0.8 | SS |
| E20<br>Aml5/H12.5 | 10.5 ± 0.8 | -0.8 ± 0.6 | SS | 10.4 ± 0.8 | -0.9 ± 0.7 | SS | 10.3 ± 0.9 | -0.9 ± 0.8 | SS | 10.1 ± 0.9 | -1.2 ± 0.9 | SS | 10.0 ± 1.1 | -1.1 ± 0.9 | SS | 10.1 ± 0.9 | -1.1 ± 0.8 | SS |
| E20<br>B5/H12.5 | 11.2 ± 0.9 | -0.1 ± 0.7 | 0.07832 | 11.0 ± 0.8 | -0.3 ± 0.6 | 0.01008 | 10.8 ± 1.0 | -0.4 ± 0.8 | 0.00013 | 10.4 ± 1.0 | -0.8 ± 0.8 | SS | 10.2 ± 1.0 | -1.0 ± 0.8 | SS | 10.2 ± 0.9 | -1.0 ± 0.8 | SS |
| L100<br>Aml5/B5 | 12.5 ± 1.2 | 1.2 ± 0.9 | SS | 12.2 ± 1.1 | 0.9 ± 0.8 | SS | 11.8 ± 1.1 | 0.6 ± 0.8 | SS | 11.0 ± 0.8 | -0.2 ± 0.6 | 0.05410 | 10.7 ± 0.9 | -0.4 ± 0.6 | 0.00002 | 10.8 ± 0.9 | -0.4 ± 0.7 | 0.00007 |
| L100<br>Aml5/H12.5 | 10.2 ± 0.9 | -1.1 ± 0.7 | SS | 10.1 ± 0.9 | -1.1 ± 0.8 | SS | 10.0 ± 1.0 | -1.1 ± 0.8 | SS | 10.2 ± 0.9 | -1.0 ± 0.8 | SS | 10.2 ± 1.0 | -0.9 ± 0.9 | SS | 10.5 ± 0.8 | -0.8 ± 0.7 | SS |
| L100<br>B5/H12.5 | 11.4 ± 1.1 | 0.1 ± 0.8 | 0.00630 | 11.1 ± 0.9 | -0.1 ± 0.7 | 0.01008 | 10.9 ± 1.1 | -0.3 ± 0.9 | 0.00079 | 10.5 ± 0.9 | -0.7 ± 0.8 | SS | 10.3 ± 1.0 | -0.8 ± 0.8 | SS | 10.4 ± 0.9 | -0.8 ± 0.8 | SS |

**Al300 =** aliskiren 300 mg; **Aml5** = amlodipine 5 mg; **B5** = bisoprolol 5 mg; **E20** = enalapril 20 mg; **H12.5** = hydrochlorothiazide 12.5 mg; **L100** = losartan 100 mg; **SS** = statistically significant (*P* < 0.00001)

**Table S13.** *P*-values calculated using the Kolmogorov-Smirnov test for changes in diastolic pulmonary arterial pressure in populations ($n$ = 100) with different ACE activity receiving the same regimens (case 1: $c_{ACE}$ = 7.0 h$^{-1}$, case 2: $c_{ACE}$ = 8.9 h$^{-1}$, case 3: $c_{ACE}$ = 10.8 h$^{-1}$, case 4: $c_{ACE}$ = 42.3 h$^{-1}$, case 5: $c_{ACE}$ = 54.1 h$^{-1}$, case 6: $c_{ACE}$ = 65.9 h$^{-1}$; *P*-value for case $i$ *vs*. case $j$ is denoted $P_{ij}$)

| Regimens | $P_{12}$ | $P_{13}$ | $P_{23}$ | $P_{14}$ | $P_{15}$ | $P_{16}$ | $P_{24}$ | $P_{25}$ | $P_{26}$ | $P_{34}$ | $P_{35}$ | $P_{36}$ | $P_{45}$ | $P_{46}$ | $P_{56}$ |
|---|---|---|---|---|---|---|---|---|---|---|---|---|---|---|---|
| Al300 | 0.21055 | 0.00386 | 0.36672 | 0.03663 | 0.11113 | 0.03663 | 0.81275 | 0.81275 | 0.00386 | 0.69937 | 0.46756 | 0.00136 | 0.96707 | 0.00386 | 0.00630 |
| E20 | 0.07832 | 0.00136 | 0.28093 | SS | SS | SS | 0.00004 | SS | SS | 0.00136 | 0.00013 | 0.00025 | 0.58062 | 0.58062 | 0.07832 |
| L100 | 0.15454 | 0.00232 | 0.28093 | 0.00045 | 0.00025 | 0.21055 | 0.11113 | 0.11113 | 0.36672 | 0.96707 | 0.90621 | 0.07832 | 0.96707 | 0.02431 | 0.00630 |
| Aml5 | 0.58062 | 0.58062 | 0.58062 | 0.15454 | 0.00386 | 0.03663 | 0.21055 | 0.03663 | 0.28093 | 0.36672 | 0.11113 | 0.36672 | 0.28093 | 0.81275 | 0.81275 |
| B5 | 0.15454 | 0.46756 | 0.81275 | 0.99376 | 0.99376 | 0.00386 | 0.07832 | 0.03663 | 0.00013 | 0.36672 | 0.36672 | 0.00025 | 0.96707 | 0.01581 | 0.02431 |
| H12.5 | 0.28093 | 0.90621 | 0.36672 | 0.96707 | 0.81275 | 0.58062 | 0.36672 | 0.07832 | 0.01581 | 0.96707 | 0.46756 | 0.21055 | 0.58062 | 0.46756 | 0.90621 |
| Al300 Aml5 | 0.58062 | 0.36672 | 0.90621 | 0.69937 | 0.58062 | 0.28093 | 0.81275 | 0.11113 | 0.03663 | 0.96707 | 0.05410 | 0.03663 | 0.11113 | 0.02431 | 0.36672 |
| Al300 B5 | 0.02431 | 0.00002 | 0.05410 | SS | SS | SS | SS | SS | SS | SS | SS | SS | 0.03663 | 0.28093 | 0.46756 |
| Al300 H12.5 | 0.81275 | 0.58062 | 0.46756 | 0.05410 | 0.00025 | SS | 0.07832 | 0.00232 | 0.00007 | 0.15454 | 0.00013 | 0.00013 | 0.05410 | 0.02431 | 0.15454 |
| E20 Aml5 | 0.28093 | 0.21055 | 0.36672 | SS | SS | SS | 0.00002 | 0.00002 | SS | 0.00004 | 0.00013 | SS | 0.96707 | 0.96707 | 0.90621 |
| E20 B5 | 0.36672 | 0.01008 | 0.15454 | SS | SS | SS | SS | SS | SS | 0.00007 | SS | SS | 0.01581 | 0.02431 | 0.58062 |
| E20 H12.5 | 0.99376 | 0.36672 | 0.58062 | 0.00386 | 0.05410 | 0.01581 | 0.01581 | 0.05410 | 0.07832 | 0.21055 | 0.11113 | 0.36672 | 0.46756 | 0.90621 | 0.90621 |
| L100 Aml5 | 0.46756 | 0.36672 | 0.90621 | 0.07832 | 0.28093 | 0.81275 | 0.46756 | 0.69937 | 0.69937 | 0.81275 | 0.36672 | 0.58062 | 0.21055 | 0.36672 | 0.69937 |
| L100 B5 | 0.02431 | SS | 0.03663 | SS | SS | SS | SS | SS | SS | SS | SS | SS | 0.03663 | 0.03663 | 0.58062 |
| L100 H12.5 | 0.90621 | 0.36672 | 0.46756 | 0.15454 | 0.00232 | 0.00045 | 0.21055 | 0.03663 | 0.00136 | 0.28093 | 0.00386 | 0.00079 | 0.07832 | 0.03663 | 0.46756 |
| Al300 Aml5/B5 | 0.02431 | 0.00045 | 0.11113 | SS | SS | SS | SS | SS | SS | SS | SS | SS | 0.03663 | 0.15454 | 0.58062 |
| Al300 Aml5/H12.5 | 0.69937 | 0.69937 | 0.96707 | 0.03663 | 0.00013 | 0.00025 | 0.11113 | 0.01008 | 0.01581 | 0.05410 | 0.00232 | 0.00386 | 0.21055 | 0.15454 | 0.99376 |
| Al300 B5/H12.5 | 0.36672 | 0.00630 | 0.07832 | SS | SS | SS | 0.00079 | 0.00004 | 0.00004 | 0.03663 | 0.00136 | 0.01581 | 0.69937 | 0.90621 | 0.58062 |
| E20 Aml5/B5 | 0.11113 | 0.01581 | 0.36672 | SS | SS | SS | SS | SS | SS | SS | SS | SS | 0.03663 | 0.00386 | 0.28093 |
| E20 Aml5/H12.5 | 0.81275 | 0.21055 | 0.69937 | 0.00232 | 0.03663 | 0.00079 | 0.03663 | 0.21055 | 0.01581 | 0.21055 | 0.58062 | 0.03663 | 0.58062 | 0.81275 | 0.46756 |
| E20 B5/H12.5 | 0.36672 | 0.00630 | 0.05410 | SS | SS | SS | SS | SS | SS | 0.00386 | 0.00004 | SS | 0.46756 | 0.11113 | 0.21055 |
| L100 Aml5/B5 | 0.03663 | 0.00007 | 0.05410 | SS | SS | SS | SS | SS | SS | SS | SS | SS | 0.02431 | 0.05410 | 0.28093 |
| L100 Aml5/H12.5 | 0.81275 | 0.58062 | 0.96707 | 0.03663 | 0.00232 | 0.01581 | 0.15454 | 0.01581 | 0.05410 | 0.15454 | 0.00386 | 0.02431 | 0.21055 | 0.36672 | 0.96707 |
| L100 B5/H12.5 | 0.15454 | 0.01008 | 0.03663 | SS | SS | SS | 0.00007 | SS | SS | 0.00386 | 0.00013 | 0.00136 | 0.58062 | 0.69937 | 0.46756 |

**Al300 =** aliskiren 300 mg; **Aml5** = amlodipine 5 mg; **B5** = bisoprolol 5 mg; **E20** = enalapril 20 mg; **H12.5** = hydrochlorothiazide 12.5 mg; **L100** = losartan 100 mg; **SS** = statistically significant ($P < 0.00001$)

**Figure S14.** Simulated change in systolic pulmonary arterial pressure from baseline to week 4 (mean ± SD, $n$ = 100)

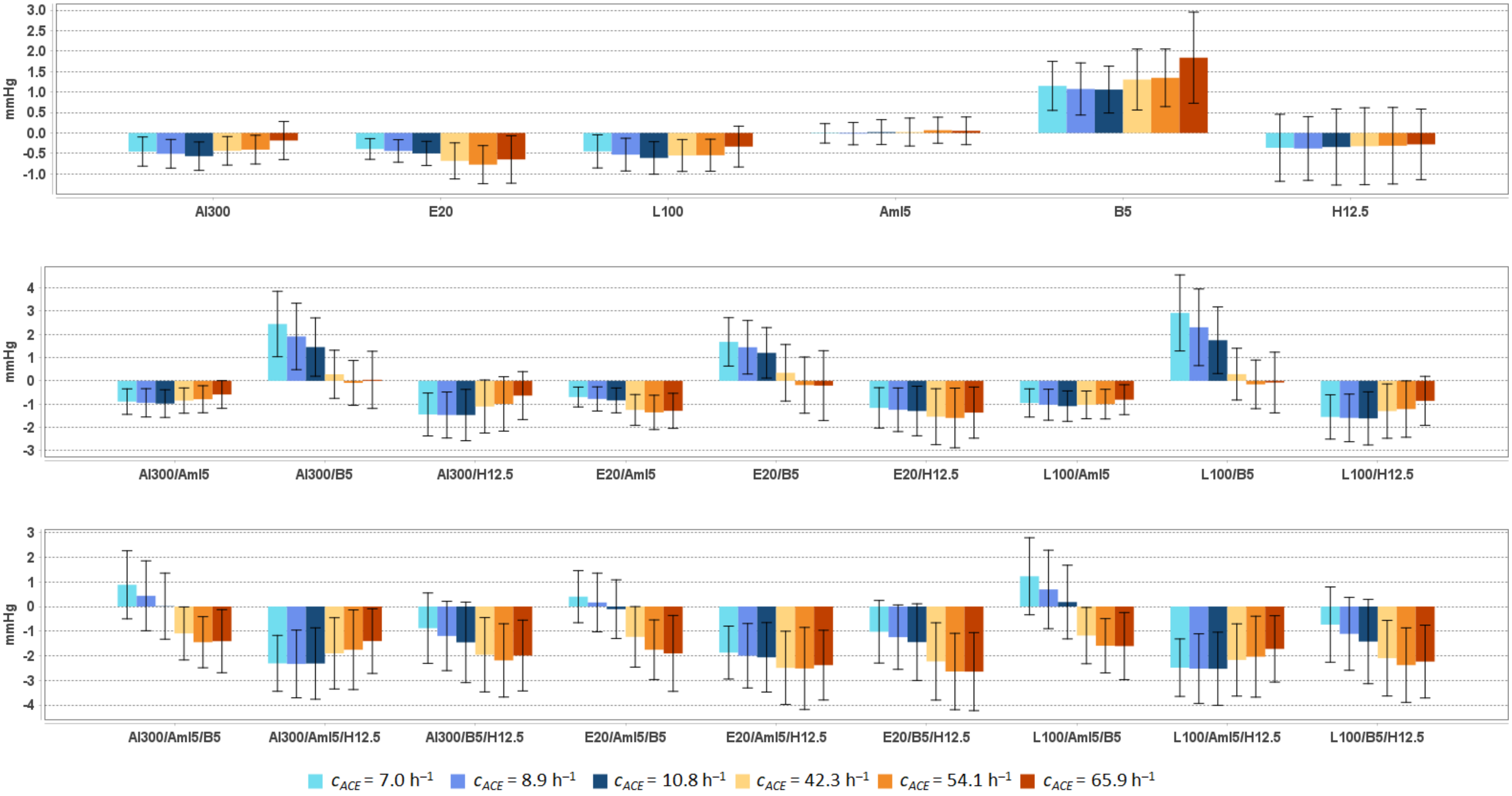


**Al300 =** aliskiren 300 mg; **Aml5** = amlodipine 5 mg; **B5** = bisoprolol 5 mg; **E20** = enalapril 20 mg; **H12.5** = hydrochlorothiazide 12.5 mg; **L100** = losartan 100 mg

**Table S14.** Simulated response of <u>systolic pulmonary arterial pressure</u> to antihypertensive therapy in virtual hypertensive populations ($n$ = 100) with different ACE activity, including $P$-values (Kolmogorov-Smirnov test) for endpoint *vs.* baseline; data are presented as mean ± SD in mmHg

| **Regimens** | **$c_{ACE}$ = 7.0 h$^{-1}$** | | | **$c_{ACE}$ = 8.9 h$^{-1}$** | | | **$c_{ACE}$ = 10.8 h$^{-1}$** | | | **$c_{ACE}$ = 42.3 h$^{-1}$** | | | **$c_{ACE}$ = 54.1 h$^{-1}$** | | | **$c_{ACE}$ = 65.9 h$^{-1}$** | | |
|---|---|---|---|---|---|---|---|---|---|---|---|---|---|---|---|---|---|---|
| | **Value** | **Change** | ***P*** | **Value** | **Change** | ***P*** | **Value** | **Change** | ***P*** | **Value** | **Change** | ***P*** | **Value** | **Change** | ***P*** | **Value** | **Change** | ***P*** |
| Baseline | 17.8 ± 1.5 | – | – | 17.6 ± 1.3 | – | – | 17.5 ± 1.4 | – | – | 17.4 ± 1.4 | – | – | 17.7 ± 1.6 | – | – | 17.8 ± 1.5 | – | – |
| Al300 | 17.3 ± 1.5 | -0.5 ± 0.4 | 0.11113 | 17.1 ± 1.2 | -0.5 ± 0.4 | 0.05410 | 17.0 ± 1.4 | -0.6 ± 0.3 | 0.05410 | 16.9 ± 1.3 | -0.4 ± 0.3 | 0.05410 | 17.3 ± 1.5 | -0.4 ± 0.4 | 0.15454 | 17.6 ± 1.5 | -0.2 ± 0.5 | 0.28093 |
| E20 | 17.4 ± 1.4 | -0.4 ± 0.3 | 0.21055 | 17.1 ± 1.2 | -0.4 ± 0.3 | 0.11113 | 17.0 ± 1.4 | -0.5 ± 0.3 | 0.11113 | 16.7 ± 1.3 | -0.7 ± 0.4 | 0.00136 | 16.9 ± 1.5 | -0.8 ± 0.5 | 0.01008 | 17.1 ± 1.4 | -0.6 ± 0.6 | 0.00630 |
| L100 | 17.3 ± 1.5 | -0.5 ± 0.4 | 0.11113 | 17.0 ± 1.3 | -0.5 ± 0.4 | 0.03663 | 16.9 ± 1.4 | -0.6 ± 0.4 | 0.01581 | 16.8 ± 1.3 | -0.6 ± 0.4 | 0.01008 | 17.2 ± 1.5 | -0.5 ± 0.4 | 0.07832 | 17.5 ± 1.4 | -0.3 ± 0.5 | 0.07832 |
| Aml5 | 17.7 ± 1.5 | -0.0 ± 0.2 | 0.99376 | 17.5 ± 1.3 | -0.0 ± 0.3 | 0.99376 | 17.6 ± 1.5 | 0.0 ± 0.3 | 0.99963 | 17.4 ± 1.4 | 0.0 ± 0.3 | 0.96707 | 17.8 ± 1.6 | 0.1 ± 0.3 | 0.99963 | 17.8 ± 1.6 | 0.1 ± 0.3 | 0.99376 |
| B5 | 18.9 ± 1.6 | 1.2 ± 0.6 | SS | 18.6 ± 1.5 | 1.1 ± 0.6 | SS | 18.6 ± 1.6 | 1.1 ± 0.6 | 0.00025 | 18.7 ± 1.5 | 1.3 ± 0.7 | SS | 19.1 ± 1.7 | 1.3 ± 0.7 | 0.00004 | 19.6 ± 1.8 | 1.8 ± 1.1 | SS |
| H12.5 | 17.4 ± 1.7 | -0.4 ± 0.8 | 0.21055 | 17.2 ± 1.3 | -0.4 ± 0.8 | 0.21055 | 17.2 ± 1.6 | -0.3 ± 0.9 | 0.21055 | 17.0 ± 1.6 | -0.3 ± 0.9 | 0.15454 | 17.4 ± 1.7 | -0.3 ± 0.9 | 0.36672 | 17.5 ± 1.7 | -0.3 ± 0.9 | 0.46756 |
| Al300<br>Aml5 | 16.9 ± 1.5 | -0.9 ± 0.5 | 0.00136 | 16.6 ± 1.3 | -0.9 ± 0.6 | 0.00007 | 16.6 ± 1.5 | -1.0 ± 0.6 | 0.00013 | 16.5 ± 1.3 | -0.8 ± 0.5 | 0.00025 | 16.9 ± 1.6 | -0.8 ± 0.6 | 0.01008 | 17.2 ± 1.4 | -0.6 ± 0.6 | 0.01581 |
| Al300<br>B5 | 20.2 ± 2.1 | 2.4 ± 1.4 | SS | 19.5 ± 2.0 | 1.9 ± 1.4 | SS | 19.0 ± 2.0 | 1.5 ± 1.3 | 0.00004 | 17.6 ± 1.5 | 0.3 ± 1.0 | 0.46756 | 17.6 ± 1.7 | -0.1 ± 1.0 | 0.90621 | 17.8 ± 1.7 | 0.0 ± 1.2 | 0.69937 |
| Al300<br>H12.5 | 16.3 ± 1.7 | -1.4 ± 0.9 | SS | 16.1 ± 1.4 | -1.5 ± 1.0 | SS | 16.1 ± 1.6 | -1.5 ± 1.1 | SS | 16.3 ± 1.6 | -1.1 ± 1.1 | SS | 16.7 ± 1.7 | -1.0 ± 1.2 | 0.00007 | 17.2 ± 1.7 | -0.6 ± 1.0 | 0.05410 |
| E20<br>Aml5 | 17.1 ± 1.5 | -0.7 ± 0.4 | 0.01581 | 16.8 ± 1.2 | -0.8 ± 0.5 | 0.00232 | 16.7 ± 1.5 | -0.8 ± 0.5 | 0.00232 | 16.1 ± 1.4 | -1.3 ± 0.7 | SS | 16.3 ± 1.6 | -1.4 ± 0.7 | 0.00004 | 16.5 ± 1.4 | -1.3 ± 0.8 | 0.00004 |
| E20<br>B5 | 19.4 ± 1.9 | 1.7 ± 1.0 | SS | 19.0 ± 1.8 | 1.4 ± 1.2 | SS | 18.7 ± 1.9 | 1.2 ± 1.1 | 0.00025 | 17.7 ± 1.7 | 0.3 ± 1.2 | 0.21055 | 17.5 ± 1.8 | -0.2 ± 1.2 | 0.46756 | 17.6 ± 1.8 | -0.2 ± 1.5 | 0.36672 |
| E20<br>H12.5 | 16.6 ± 1.6 | -1.2 ± 0.9 | 0.00002 | 16.3 ± 1.3 | -1.2 ± 0.9 | SS | 16.2 ± 1.6 | -1.3 ± 1.1 | 0.00004 | 15.8 ± 1.6 | -1.5 ± 1.2 | SS | 16.1 ± 1.8 | -1.6 ± 1.3 | SS | 16.4 ± 1.7 | -1.4 ± 1.1 | 0.00002 |
| L100<br>Aml5 | 16.8 ± 1.5 | -1.0 ± 0.6 | 0.00136 | 16.5 ± 1.3 | -1.0 ± 0.7 | 0.00007 | 16.4 ± 1.5 | -1.1 ± 0.7 | 0.00004 | 16.3 ± 1.4 | -1.0 ± 0.6 | SS | 16.7 ± 1.6 | -1.0 ± 0.6 | 0.00232 | 17.0 ± 1.4 | -0.8 ± 0.6 | 0.00136 |
| L100<br>B5 | 20.7 ± 2.3 | 2.9 ± 1.6 | SS | 19.9 ± 2.2 | 2.3 ± 1.6 | SS | 19.3 ± 2.1 | 1.7 ± 1.4 | SS | 17.7 ± 1.6 | 0.3 ± 1.1 | 0.28093 | 17.6 ± 1.7 | -0.2 ± 1.0 | 0.69937 | 17.7 ± 1.7 | -0.1 ± 1.3 | 0.46756 |
| L100<br>H12.5 | 16.2 ± 1.7 | -1.6 ± 1.0 | SS | 16.0 ± 1.4 | -1.6 ± 1.0 | SS | 15.9 ± 1.7 | -1.6 ± 1.1 | SS | 16.1 ± 1.6 | -1.3 ± 1.2 | SS | 16.5 ± 1.8 | -1.2 ± 1.2 | 0.00002 | 16.9 ± 1.7 | -0.9 ± 1.1 | 0.01581 |
| Al300<br>Aml5/B5 | 18.6 ± 2.1 | 0.9 ± 1.4 | 0.00045 | 18.0 ± 1.9 | 0.4 ± 1.4 | 0.07832 | 17.6 ± 2.0 | 0.0 ± 1.3 | 0.28093 | 16.3 ± 1.5 | -1.1 ± 1.1 | SS | 16.3 ± 1.7 | -1.4 ± 1.0 | 0.00002 | 16.4 ± 1.6 | -1.4 ± 1.3 | SS |
| Al300<br>Aml5/H12.5 | 15.4 ± 1.8 | -2.3 ± 1.1 | SS | 15.2 ± 1.6 | -2.3 ± 1.4 | SS | 15.2 ± 1.9 | -2.3 ± 1.4 | SS | 15.5 ± 1.8 | -1.9 ± 1.4 | SS | 15.9 ± 2.0 | -1.8 ± 1.6 | SS | 16.4 ± 1.9 | -1.4 ± 1.3 | SS |
| Al300<br>B5/H12.5 | 16.9 ± 2.0 | -0.9 ± 1.4 | 0.00079 | 16.4 ± 1.7 | -1.2 ± 1.4 | SS | 16.1 ± 2.0 | -1.5 ± 1.6 | SS | 15.4 ± 1.8 | -2.0 ± 1.5 | SS | 15.5 ± 1.8 | -2.2 ± 1.5 | SS | 15.8 ± 1.7 | -2.0 ± 1.4 | SS |
| E20<br>Aml5/B5 | 18.2 ± 1.8 | 0.4 ± 1.1 | 0.01581 | 17.7 ± 1.7 | 0.2 ± 1.2 | 0.46756 | 17.4 ± 1.9 | -0.1 ± 1.2 | 0.28093 | 16.1 ± 1.6 | -1.2 ± 1.2 | SS | 15.9 ± 1.8 | -1.8 ± 1.2 | SS | 15.9 ± 1.8 | -1.9 ± 1.5 | SS |
| E20<br>Aml5/H12.5 | 15.9 ± 1.8 | -1.9 ± 1.1 | SS | 15.6 ± 1.5 | -2.0 ± 1.3 | SS | 15.5 ± 1.9 | -2.1 ± 1.4 | SS | 14.9 ± 1.8 | -2.5 ± 1.5 | SS | 15.2 ± 2.0 | -2.5 ± 1.7 | SS | 15.4 ± 1.9 | -2.4 ± 1.4 | SS |
| E20<br>B5/H12.5 | 16.7 ± 1.9 | -1.0 ± 1.3 | 0.00025 | 16.3 ± 1.6 | -1.2 ± 1.3 | SS | 16.1 ± 2.0 | -1.4 ± 1.6 | SS | 15.1 ± 1.8 | -2.2 ± 1.6 | SS | 15.1 ± 1.8 | -2.6 ± 1.6 | SS | 15.1 ± 1.8 | -2.6 ± 1.6 | SS |
| L100<br>Aml5/B5 | 19.0 ± 2.2 | 1.2 ± 1.6 | SS | 18.3 ± 2.1 | 0.7 ± 1.6 | 0.01581 | 17.7 ± 2.1 | 0.2 ± 1.5 | 0.36672 | 16.2 ± 1.6 | -1.2 ± 1.1 | SS | 16.1 ± 1.7 | -1.6 ± 1.1 | SS | 16.2 ± 1.7 | -1.6 ± 1.4 | SS |
| L100<br>Aml5/H12.5 | 15.3 ± 1.8 | -2.5 ± 1.2 | SS | 15.0 ± 1.6 | -2.5 ± 1.4 | SS | 15.0 ± 1.9 | -2.5 ± 1.5 | SS | 15.2 ± 1.8 | -2.2 ± 1.5 | SS | 15.7 ± 2.0 | -2.0 ± 1.6 | SS | 16.1 ± 1.9 | -1.7 ± 1.3 | SS |
| L100<br>B5/H12.5 | 17.0 ± 2.1 | -0.7 ± 1.5 | 0.00386 | 16.5 ± 1.8 | -1.1 ± 1.5 | SS | 16.1 ± 2.1 | -1.4 ± 1.7 | SS | 15.3 ± 1.8 | -2.1 ± 1.5 | SS | 15.3 ± 1.8 | -2.4 ± 1.5 | SS | 15.6 ± 1.8 | -2.2 ± 1.5 | SS |

**Al300 =** aliskiren 300 mg; **Aml5** = amlodipine 5 mg; **B5** = bisoprolol 5 mg; **E20** = enalapril 20 mg; **H12.5** = hydrochlorothiazide 12.5 mg; **L100** = losartan 100 mg; **SS** = statistically significant ($P$ < 0.00001)

**Table S15.** *P*-values calculated using the Kolmogorov-Smirnov test for changes in systolic pulmonary arterial pressure in populations ($n$ = 100) with different ACE activity receiving the same regimens (case 1: $c_{ACE}$ = 7.0 $h^{-1}$, case 2: $c_{ACE}$ = 8.9 $h^{-1}$, case 3: $c_{ACE}$ = 10.8 $h^{-1}$, case 4: $c_{ACE}$ = 42.3 $h^{-1}$, case 5: $c_{ACE}$ = 54.1 $h^{-1}$, case 6: $c_{ACE}$ = 65.9 $h^{-1}$; *P*-value for case *i vs*. case *j* is denoted $P_{ij}$)

| Regimens | $P_{12}$ | $P_{13}$ | $P_{23}$ | $P_{14}$ | $P_{15}$ | $P_{16}$ | $P_{24}$ | $P_{25}$ | $P_{26}$ | $P_{34}$ | $P_{35}$ | $P_{36}$ | $P_{45}$ | $P_{46}$ | $P_{56}$ |
|---|---|---|---|---|---|---|---|---|---|---|---|---|---|---|---|
| Al300 | 0.36672 | 0.00630 | 0.11113 | 0.81275 | 0.58062 | 0.00013 | 0.58062 | 0.28093 | 0.00002 | 0.01581 | 0.00386 | SS | 0.90621 | 0.00136 | 0.00232 |
| E20 | 0.21055 | 0.00136 | 0.07832 | SS | SS | SS | 0.00007 | SS | 0.00004 | 0.00386 | 0.00004 | 0.00232 | 0.58062 | 0.58062 | 0.15454 |
| L100 | 0.28093 | 0.00386 | 0.11113 | 0.11113 | 0.11113 | 0.07832 | 0.81275 | 0.90621 | 0.00630 | 0.58062 | 0.36672 | 0.00079 | 0.99376 | 0.01008 | 0.00630 |
| Aml5 | 0.58062 | 0.69937 | 0.69937 | 0.15454 | 0.00630 | 0.11113 | 0.21055 | 0.02431 | 0.15454 | 0.81275 | 0.15454 | 0.69937 | 0.28093 | 0.90621 | 0.69937 |
| B5 | 0.28093 | 0.58062 | 0.69937 | 0.46756 | 0.28093 | 0.00079 | 0.01008 | 0.00630 | SS | 0.03663 | 0.01581 | SS | 0.90621 | 0.01008 | 0.00630 |
| H12.5 | 0.58062 | 0.69937 | 0.28093 | 0.69937 | 0.81275 | 0.69937 | 0.36672 | 0.11113 | 0.05410 | 0.99376 | 0.46756 | 0.36672 | 0.28093 | 0.69937 | 0.99963 |
| Al300 Aml5 | 0.58062 | 0.36672 | 0.96707 | 0.28093 | 0.21055 | 0.00386 | 0.15454 | 0.01581 | 0.00045 | 0.21055 | 0.00136 | 0.00079 | 0.11113 | 0.00386 | 0.11113 |
| Al300 B5 | 0.01581 | 0.00004 | 0.02431 | SS | SS | SS | SS | SS | SS | SS | SS | SS | 0.05410 | 0.21055 | 0.21055 |
| Al300 H12.5 | 0.90621 | 0.46756 | 0.36672 | 0.02431 | 0.00007 | SS | 0.05410 | 0.00136 | SS | 0.07832 | 0.00013 | SS | 0.07832 | 0.01008 | 0.36672 |
| E20 Aml5 | 0.15454 | 0.11113 | 0.58062 | SS | SS | SS | SS | SS | SS | 0.00013 | 0.00007 | SS | 0.69937 | 0.90621 | 0.69937 |
| E20 B5 | 0.36672 | 0.00232 | 0.01581 | SS | SS | SS | SS | SS | SS | 0.00007 | SS | SS | 0.02431 | 0.01008 | 0.69937 |
| E20 H12.5 | 0.69937 | 0.21055 | 0.28093 | 0.00386 | 0.11113 | 0.05410 | 0.00232 | 0.21055 | 0.03663 | 0.15454 | 0.15454 | 0.69937 | 0.46756 | 0.58062 | 0.81275 |
| L100 Aml5 | 0.58062 | 0.11113 | 0.81275 | 0.36672 | 0.81275 | 0.36672 | 0.81275 | 0.36672 | 0.07832 | 0.69937 | 0.03663 | 0.07832 | 0.28093 | 0.07832 | 0.21055 |
| L100 B5 | 0.02431 | SS | 0.01581 | SS | SS | SS | SS | SS | SS | SS | SS | SS | 0.01581 | 0.07832 | 0.28093 |
| L100 H12.5 | 0.90621 | 0.21055 | 0.36672 | 0.15454 | 0.00079 | 0.00002 | 0.11113 | 0.00630 | 0.00013 | 0.15454 | 0.00136 | 0.00013 | 0.07832 | 0.03663 | 0.28093 |
| Al300 Aml5/B5 | 0.02431 | 0.00045 | 0.11113 | SS | SS | SS | SS | SS | SS | SS | SS | SS | 0.01581 | 0.15454 | 0.81275 |
| Al300 Aml5/H12.5 | 0.58062 | 0.58062 | 0.96707 | 0.00386 | 0.00007 | SS | 0.07832 | 0.01008 | 0.00045 | 0.05410 | 0.00232 | 0.00045 | 0.28093 | 0.07832 | 0.69937 |
| Al300 B5/H12.5 | 0.36672 | 0.00630 | 0.28093 | 0.00007 | SS | SS | 0.00630 | 0.00136 | 0.00025 | 0.11113 | 0.00386 | 0.03663 | 0.58062 | 0.90621 | 0.69937 |
| E20 Aml5/B5 | 0.07832 | 0.01581 | 0.28093 | SS | SS | SS | SS | SS | SS | 0.00002 | SS | SS | 0.00630 | 0.02431 | 0.36672 |
| E20 Aml5/H12.5 | 0.46756 | 0.21055 | 0.81275 | 0.00079 | 0.01008 | 0.00232 | 0.05410 | 0.28093 | 0.07832 | 0.21055 | 0.46756 | 0.15454 | 0.58062 | 0.58062 | 0.69937 |
| E20 B5/H12.5 | 0.36672 | 0.00630 | 0.46756 | SS | SS | SS | 0.00013 | SS | SS | 0.01581 | 0.00013 | 0.00007 | 0.46756 | 0.11113 | 0.46756 |
| L100 Aml5/B5 | 0.03663 | 0.00025 | 0.05410 | SS | SS | SS | SS | SS | SS | SS | SS | SS | 0.01581 | 0.07832 | 0.81275 |
| L100 Aml5/H12.5 | 0.58062 | 0.58062 | 0.96707 | 0.00630 | 0.00045 | 0.00013 | 0.15454 | 0.02431 | 0.00630 | 0.07832 | 0.01008 | 0.00136 | 0.36672 | 0.11113 | 0.90621 |
| L100 B5/H12.5 | 0.36672 | 0.00630 | 0.28093 | SS | SS | SS | 0.00079 | 0.00007 | SS | 0.03663 | 0.00136 | 0.00630 | 0.58062 | 0.58062 | 0.69937 |

**Al300 =** aliskiren 300 mg; **Aml5** = amlodipine 5 mg; **B5** = bisoprolol 5 mg; **E20** = enalapril 20 mg; **H12.5** = hydrochlorothiazide 12.5 mg; **L100** = losartan 100 mg; **SS** = statistically significant ($P$ < 0.00001)

**Figure S15.** Simulated change in <u>stroke volume</u> from baseline to week 4 (mean ± SD, $n$ = 100)

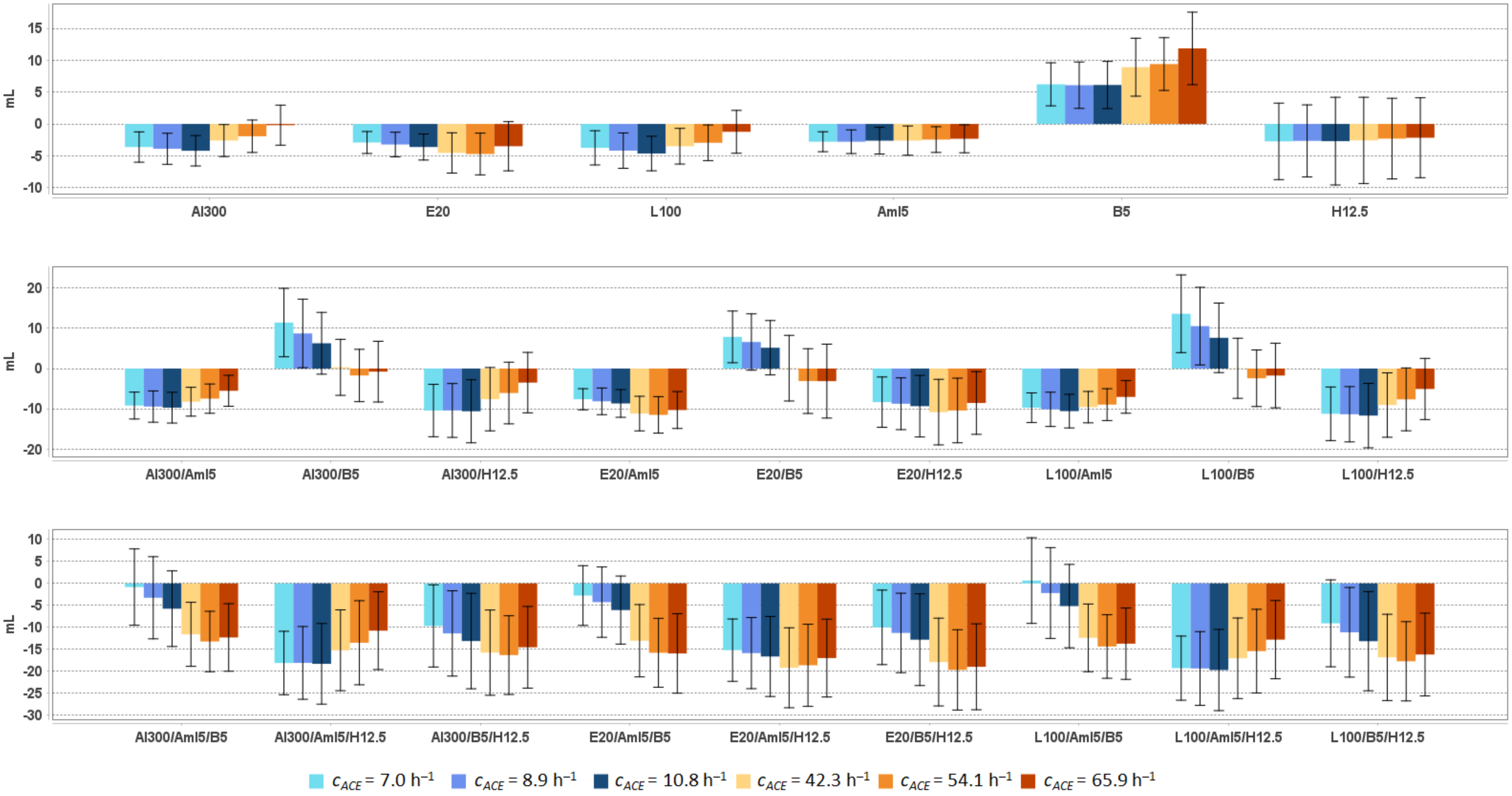


**Al300 =** aliskiren 300 mg; **Aml5** = amlodipine 5 mg; **B5** = bisoprolol 5 mg; **E20** = enalapril 20 mg; **H12.5** = hydrochlorothiazide 12.5 mg; **L100** = losartan 100 mg

**Table S16.** Simulated response of <u>stroke volume</u> to antihypertensive therapy in virtual hypertensive populations ($n$ = 100) with different ACE activity, including $P$-values (Kolmogorov-Smirnov test) for endpoint *vs*. baseline; data are presented as mean ± SD in mL

| Regimens | $c_{ACE}$ = 7.0 $h^{-1}$ | | | $c_{ACE}$ = 8.9 $h^{-1}$ | | | $c_{ACE}$ = 10.8 $h^{-1}$ | | | $c_{ACE}$ = 42.3 $h^{-1}$ | | | $c_{ACE}$ = 54.1 $h^{-1}$ | | | $c_{ACE}$ = 65.9 $h^{-1}$ | | |
|---|---|---|---|---|---|---|---|---|---|---|---|---|---|---|---|---|---|---|
| | **Value** | **Change** | ***P*** | **Value** | **Change** | ***P*** | **Value** | **Change** | ***P*** | **Value** | **Change** | ***P*** | **Value** | **Change** | ***P*** | **Value** | **Change** | ***P*** |
| Baseline | 73.2 ± 6.9 | – | – | 72.5 ± 7.6 | – | – | 73.5 ± 7.8 | – | – | 72.8 ± 7.3 | – | – | 73.4 ± 7.9 | – | – | 71.8 ± 7.0 | – | – |
| Al300 | 69.6 ± 7.0 | -3.6 ± 2.4 | 0.00079 | 68.6 ± 8.0 | -3.9 ± 2.4 | 0.02431 | 69.3 ± 8.0 | -4.2 ± 2.4 | 0.01008 | 70.2 ± 7.1 | -2.6 ± 2.5 | 0.02431 | 71.5 ± 8.1 | -1.9 ± 2.5 | 0.11113 | 71.7 ± 7.0 | -0.2 ± 3.1 | 0.99963 |
| E20 | 70.3 ± 6.9 | -2.9 ± 1.7 | 0.00630 | 69.3 ± 7.8 | -3.2 ± 1.9 | 0.07832 | 69.9 ± 7.9 | -3.6 ± 2.0 | 0.02431 | 68.2 ± 7.2 | -4.5 ± 3.2 | 0.00013 | 68.7 ± 8.1 | -4.7 ± 3.3 | 0.00045 | 68.4 ± 7.0 | -3.5 ± 3.9 | 0.01581 |
| L100 | 69.5 ± 7.2 | -3.8 ± 2.7 | 0.00079 | 68.3 ± 8.1 | -4.2 ± 2.8 | 0.01581 | 68.9 ± 8.0 | -4.6 ± 2.7 | 0.00386 | 69.3 ± 7.1 | -3.5 ± 2.8 | 0.00232 | 70.5 ± 8.1 | -3.0 ± 2.8 | 0.01581 | 70.6 ± 7.0 | -1.2 ± 3.4 | 0.46756 |
| Aml5 | 70.4 ± 7.1 | -2.8 ± 1.6 | 0.01581 | 69.7 ± 7.8 | -2.8 ± 1.9 | 0.07832 | 70.9 ± 7.8 | -2.6 ± 2.1 | 0.11113 | 70.2 ± 8.0 | -2.6 ± 2.3 | 0.11113 | 71.0 ± 8.0 | -2.5 ± 2.0 | 0.01008 | 69.5 ± 7.6 | -2.3 ± 2.2 | 0.05410 |
| B5 | 79.5 ± 7.6 | 6.2 ± 3.4 | SS | 78.6 ± 8.8 | 6.1 ± 3.6 | 0.00045 | 79.6 ± 8.9 | 6.1 ± 3.7 | SS | 81.7 ± 8.2 | 8.9 ± 4.6 | SS | 82.8 ± 9.8 | 9.4 ± 4.1 | SS | 83.7 ± 8.7 | 11.9 ± 5.7 | SS |
| H12.5 | 70.5 ± 9.1 | -2.7 ± 6.0 | 0.02431 | 69.9 ± 9.4 | -2.7 ± 5.7 | 0.07832 | 70.8 ± 10.5 | -2.7 ± 6.9 | 0.07832 | 70.2 ± 10.0 | -2.6 ± 6.8 | 0.11113 | 71.1 ± 10.5 | -2.3 ± 6.3 | 0.15454 | 69.7 ± 9.1 | -2.2 ± 6.3 | 0.11113 |
| Al300<br>Aml5 | 64.0 ± 7.2 | -9.2 ± 3.3 | SS | 63.1 ± 8.3 | -9.4 ± 3.9 | SS | 63.8 ± 8.1 | -9.7 ± 3.8 | SS | 64.6 ± 7.7 | -8.2 ± 3.6 | SS | 66.0 ± 8.2 | -7.5 ± 3.6 | SS | 66.3 ± 7.5 | -5.5 ± 3.9 | 0.00002 |
| Al300<br>B5 | 84.7 ± 11.2 | 11.4 ± 8.5 | SS | 81.3 ± 11.8 | 8.7 ± 8.5 | SS | 79.8 ± 10.8 | 6.3 ± 7.7 | SS | 73.1 ± 9.1 | 0.3 ± 7.0 | 0.96707 | 71.7 ± 10.1 | -1.7 ± 6.5 | 0.07832 | 71.1 ± 9.1 | -0.8 ± 7.5 | 0.58062 |
| Al300<br>H12.5 | 62.8 ± 9.0 | -10.4 ± 6.5 | SS | 62.1 ± 9.9 | -10.4 ± 6.7 | SS | 62.9 ± 10.8 | -10.6 ± 7.8 | SS | 65.2 ± 10.3 | -7.6 ± 7.9 | SS | 67.4 ± 11.2 | -6.1 ± 7.7 | 0.00002 | 68.4 ± 9.6 | -3.5 ± 7.5 | 0.01008 |
| E20<br>Aml5 | 65.6 ± 7.0 | -7.6 ± 2.6 | SS | 64.4 ± 8.1 | -8.1 ± 3.3 | SS | 64.8 ± 8.0 | -8.7 ± 3.4 | SS | 61.6 ± 7.8 | -11.2 ± 4.3 | SS | 61.9 ± 8.2 | -11.5 ± 4.5 | SS | 61.5 ± 7.4 | -10.3 ± 4.6 | SS |
| E20<br>B5 | 81.1 ± 9.6 | 7.9 ± 6.4 | SS | 79.1 ± 10.7 | 6.6 ± 7.0 | 0.00045 | 78.7 ± 10.3 | 5.2 ± 6.8 | 0.00004 | 72.9 ± 9.9 | 0.1 ± 8.2 | 0.81275 | 70.3 ± 11.1 | -3.1 ± 8.0 | 0.00630 | 68.7 ± 10.3 | -3.1 ± 9.2 | 0.02431 |
| E20<br>H12.5 | 64.9 ± 8.9 | -8.3 ± 6.3 | SS | 63.8 ± 9.7 | -8.7 ± 6.4 | SS | 64.2 ± 10.7 | -9.3 ± 7.7 | SS | 62.0 ± 10.3 | -10.8 ± 8.1 | SS | 63.0 ± 11.1 | -10.4 ± 8.0 | SS | 63.3 ± 9.4 | -8.5 ± 7.8 | SS |
| L100<br>Aml5 | 63.5 ± 7.4 | -9.7 ± 3.7 | SS | 62.4 ± 8.5 | -10.1 ± 4.3 | SS | 62.9 ± 8.3 | -10.6 ± 4.2 | SS | 63.2 ± 7.7 | -9.6 ± 3.9 | SS | 64.5 ± 8.2 | -9.0 ± 4.0 | SS | 64.8 ± 7.5 | -7.0 ± 4.1 | SS |
| L100<br>B5 | 86.9 ± 12.1 | 13.6 ± 9.7 | SS | 83.1 ± 12.6 | 10.5 ± 9.6 | SS | 81.1 ± 11.4 | 7.6 ± 8.6 | SS | 72.9 ± 9.4 | 0.1 ± 7.5 | 0.81275 | 71.0 ± 10.4 | -2.4 ± 7.0 | 0.05410 | 70.1 ± 9.4 | -1.7 ± 8.0 | 0.28093 |
| L100<br>H12.5 | 62.0 ± 9.1 | -11.2 ± 6.7 | SS | 61.2 ± 10.0 | -11.4 ± 6.9 | SS | 61.8 ± 10.9 | -11.7 ± 8.0 | SS | 63.7 ± 10.3 | -9.1 ± 8.0 | SS | 65.8 ± 11.2 | -7.7 ± 7.8 | SS | 66.8 ± 9.5 | -5.1 ± 7.6 | 0.00004 |
| Al300<br>Aml5/B5 | 72.4 ± 11.1 | -0.8 ± 8.7 | 0.07832 | 69.3 ± 12.3 | -3.3 ± 9.3 | 0.00232 | 67.8 ± 11.2 | -5.7 ± 8.6 | 0.00025 | 61.2 ± 9.2 | -11.6 ± 7.3 | SS | 60.2 ± 9.7 | -13.2 ± 6.9 | SS | 59.6 ± 9.0 | -12.3 ± 7.7 | SS |
| Al300<br>Aml5/H12.5 | 55.1 ± 9.5 | -18.1 ± 7.2 | SS | 54.5 ± 10.9 | -18.1 ± 8.3 | SS | 55.2 ± 11.5 | -18.3 ± 9.2 | SS | 57.6 ± 11.6 | -15.2 ± 9.2 | SS | 59.9 ± 12.2 | -13.5 ± 9.6 | SS | 61.1 ± 11.0 | -10.7 ± 8.8 | SS |
| Al300<br>B5/H12.5 | 63.6 ± 11.5 | -9.7 ± 9.4 | SS | 61.2 ± 12.3 | -11.4 ± 9.7 | SS | 60.4 ± 13.0 | -13.1 ± 10.8 | SS | 57.0 ± 11.2 | -15.7 ± 9.7 | SS | 57.1 ± 11.3 | -16.3 ± 9.0 | SS | 57.3 ± 10.0 | -14.5 ± 9.3 | SS |
| E20<br>Aml5/B5 | 70.5 ± 9.6 | -2.8 ± 6.8 | 0.00386 | 68.3 ± 11.3 | -4.3 ± 8.0 | 0.00045 | 67.4 ± 10.6 | -6.1 ± 7.7 | 0.00013 | 59.8 ± 9.8 | -13.0 ± 8.2 | SS | 57.7 ± 10.3 | -15.8 ± 7.8 | SS | 55.9 ± 9.8 | -15.9 ± 9.0 | SS |
| E20<br>Aml5/H12.5 | 58.1 ± 9.5 | -15.2 ± 7.1 | SS | 56.7 ± 10.8 | -15.8 ± 8.1 | SS | 56.9 ± 11.6 | -16.6 ± 9.1 | SS | 53.6 ± 11.3 | -19.2 ± 9.1 | SS | 54.8 ± 11.6 | -18.6 ± 9.3 | SS | 54.9 ± 10.7 | -17.0 ± 8.8 | SS |
| E20<br>B5/H12.5 | 63.2 ± 10.7 | -10.0 ± 8.5 | SS | 61.3 ± 11.7 | -11.3 ± 9.0 | SS | 60.7 ± 12.7 | -12.8 ± 10.4 | SS | 54.9 ± 11.3 | -17.9 ± 10.0 | SS | 53.8 ± 11.1 | -19.7 ± 9.1 | SS | 52.9 ± 10.2 | -19.0 ± 9.8 | SS |
| L100<br>Aml5/B5 | 73.9 ± 11.9 | 0.6 ± 9.7 | 0.21055 | 70.3 ± 13.1 | -2.2 ± 10.3 | 0.00232 | 68.3 ± 11.8 | -5.2 ± 9.5 | 0.00079 | 60.4 ± 9.4 | -12.4 ± 7.7 | SS | 59.1 ± 9.9 | -14.3 ± 7.2 | SS | 58.1 ± 9.2 | -13.7 ± 8.1 | SS |
| L100<br>Aml5/H12.5 | 54.0 ± 9.5 | -19.3 ± 7.3 | SS | 53.2 ± 11.0 | -19.3 ± 8.4 | SS | 53.8 ± 11.5 | -19.7 ± 9.2 | SS | 55.8 ± 11.4 | -17.0 ± 9.2 | SS | 58.0 ± 12.0 | -15.4 ± 9.5 | SS | 59.1 ± 11.0 | -12.8 ± 8.9 | SS |
| L100<br>B5/H12.5 | 64.2 ± 12.0 | -9.1 ± 9.9 | SS | 61.4 ± 12.7 | -11.1 ± 10.2 | SS | 60.4 ± 13.4 | -13.1 ± 11.3 | SS | 56.0 ± 11.2 | -16.8 ± 9.8 | SS | 55.7 ± 11.2 | -17.7 ± 9.0 | SS | 55.7 ± 10.0 | -16.2 ± 9.4 | SS |

**Al300 =** aliskiren 300 mg; **Aml5** = amlodipine 5 mg; **B5** = bisoprolol 5 mg; **E20** = enalapril 20 mg; **H12.5** = hydrochlorothiazide 12.5 mg; **L100** = losartan 100 mg; **SS** = statistically significant ($P$ < 0.00001)

**Table S17.** *P*-values calculated using the Kolmogorov-Smirnov test for changes in stroke volume in populations ($n$ = 100) with different ACE activity receiving the same regimens (case 1: $c_{ACE}$ = 7.0 $h^{-1}$, case 2: $c_{ACE}$ = 8.9 $h^{-1}$, case 3: $c_{ACE}$ = 10.8 $h^{-1}$, case 4: $c_{ACE}$ = 42.3 $h^{-1}$, case 5: $c_{ACE}$ = 54.1 $h^{-1}$, case 6: $c_{ACE}$ = 65.9 $h^{-1}$; *P*-value for case *i vs*. case *j* is denoted $P_{ij}$)

| **Regimens** | $P_{12}$ | $P_{13}$ | $P_{23}$ | $P_{14}$ | $P_{15}$ | $P_{16}$ | $P_{24}$ | $P_{25}$ | $P_{26}$ | $P_{34}$ | $P_{35}$ | $P_{36}$ | $P_{45}$ | $P_{46}$ | $P_{56}$ |
|---|---|---|---|---|---|---|---|---|---|---|---|---|---|---|---|
| Al300 | 0.58062 | 0.00630 | 0.36672 | 0.05410 | 0.00045 | SS | 0.03663 | 0.00045 | SS | 0.00045 | SS | SS | 0.15454 | 0.00007 | 0.00079 |
| E20 | 0.28093 | 0.00136 | 0.07832 | SS | SS | 0.00079 | 0.00013 | 0.00025 | 0.00386 | 0.00136 | 0.00232 | 0.03663 | 0.99376 | 0.21055 | 0.11113 |
| L100 | 0.36672 | 0.00386 | 0.36672 | 0.69937 | 0.11113 | SS | 0.36672 | 0.03663 | SS | 0.02431 | 0.00013 | SS | 0.28093 | 0.00079 | 0.01581 |
| Aml5 | 0.96707 | 0.21055 | 0.36672 | 0.21055 | 0.07832 | 0.11113 | 0.58062 | 0.15454 | 0.28093 | 0.96707 | 0.36672 | 0.46756 | 0.81275 | 0.69937 | 0.81275 |
| B5 | 0.69937 | 0.81275 | 0.96707 | 0.00002 | SS | SS | SS | SS | SS | SS | SS | SS | 0.46756 | 0.00386 | 0.00386 |
| H12.5 | 0.69937 | 0.58062 | 0.46756 | 0.36672 | 0.58062 | 0.81275 | 0.15454 | 0.15454 | 0.15454 | 0.90621 | 0.46756 | 0.46756 | 0.21055 | 0.69937 | 0.99376 |
| Al300 Aml5 | 0.81275 | 0.36672 | 0.69937 | 0.28093 | 0.00386 | SS | 0.05410 | 0.00045 | SS | 0.02431 | 0.00045 | SS | 0.15454 | 0.00002 | 0.02431 |
| Al300 B5 | 0.03663 | 0.00025 | 0.05410 | SS | SS | SS | SS | SS | SS | 0.00025 | SS | SS | 0.05410 | 0.21055 | 0.46756 |
| Al300 H12.5 | 0.99376 | 0.58062 | 0.46756 | 0.01581 | 0.00013 | SS | 0.01581 | 0.00013 | SS | 0.07832 | 0.00013 | SS | 0.21055 | 0.00386 | 0.11113 |
| E20 Aml5 | 0.28093 | 0.03663 | 0.36672 | SS | SS | SS | SS | SS | 0.00007 | 0.00025 | 0.00079 | 0.00630 | 0.90621 | 0.58062 | 0.46756 |
| E20 B5 | 0.46756 | 0.01008 | 0.28093 | SS | SS | SS | SS | SS | SS | 0.00025 | SS | SS | 0.02431 | 0.03663 | 0.81275 |
| E20 H12.5 | 0.81275 | 0.05410 | 0.46756 | 0.01581 | 0.05410 | 0.46756 | 0.05410 | 0.28093 | 0.81275 | 0.15454 | 0.58062 | 0.28093 | 0.58062 | 0.11113 | 0.58062 |
| L100 Aml5 | 0.46756 | 0.15454 | 0.21055 | 0.69937 | 0.21055 | 0.00232 | 0.69937 | 0.02431 | 0.00004 | 0.11113 | 0.00386 | SS | 0.15454 | 0.00045 | 0.05410 |
| L100 B5 | 0.03663 | 0.00013 | 0.05410 | SS | SS | SS | SS | SS | SS | SS | SS | SS | 0.03663 | 0.21055 | 0.58062 |
| L100 H12.5 | 0.96707 | 0.46756 | 0.46756 | 0.11113 | 0.00079 | SS | 0.03663 | 0.00079 | SS | 0.11113 | 0.00136 | SS | 0.28093 | 0.00630 | 0.15454 |
| Al300 Aml5/B5 | 0.05410 | 0.00136 | 0.28093 | SS | SS | SS | SS | SS | SS | 0.00013 | SS | SS | 0.05410 | 0.58062 | 0.69937 |
| Al300 Aml5/H12.5 | 0.58062 | 0.58062 | 0.81275 | 0.01008 | 0.00013 | SS | 0.01008 | 0.00079 | SS | 0.11113 | 0.00386 | SS | 0.28093 | 0.00232 | 0.11113 |
| Al300 B5/H12.5 | 0.46756 | 0.00630 | 0.11113 | 0.00136 | 0.00025 | 0.00045 | 0.01581 | 0.01581 | 0.03663 | 0.21055 | 0.01581 | 0.28093 | 0.81275 | 0.58062 | 0.58062 |
| E20 Aml5/B5 | 0.11113 | 0.00630 | 0.28093 | SS | SS | SS | SS | SS | SS | SS | SS | SS | 0.03663 | 0.11113 | 0.58062 |
| E20 Aml5/H12.5 | 0.46756 | 0.15454 | 0.69937 | 0.00232 | 0.00630 | 0.05410 | 0.01581 | 0.03663 | 0.46756 | 0.28093 | 0.58062 | 0.81275 | 0.81275 | 0.21055 | 0.46756 |
| E20 B5/H12.5 | 0.58062 | 0.00630 | 0.11113 | SS | SS | SS | 0.00013 | SS | SS | 0.00232 | 0.00013 | 0.00045 | 0.58062 | 0.28093 | 0.69937 |
| L100 Aml5/B5 | 0.05410 | 0.00079 | 0.15454 | SS | SS | SS | SS | SS | SS | 0.00002 | SS | SS | 0.05410 | 0.46756 | 0.90621 |
| L100 Aml5/H12.5 | 0.58062 | 0.36672 | 0.69937 | 0.01581 | 0.00045 | SS | 0.05410 | 0.00386 | 0.00002 | 0.11113 | 0.01008 | 0.00002 | 0.36672 | 0.00630 | 0.21055 |
| L100 B5/H12.5 | 0.36672 | 0.00630 | 0.21055 | 0.00004 | SS | SS | 0.01008 | 0.00232 | 0.00045 | 0.03663 | 0.00630 | 0.05410 | 0.69937 | 0.58062 | 0.58062 |

**Al300 =** aliskiren 300 mg; **Aml5** = amlodipine 5 mg; **B5** = bisoprolol 5 mg; **E20** = enalapril 20 mg; **H12.5** = hydrochlorothiazide 12.5 mg; **L100** = losartan 100 mg; **SS** = statistically significant ($P$ < 0.00001)

**Figure S16.** Simulated change in ejection fraction from baseline to week 4 (mean ± SD, *n* = 100)

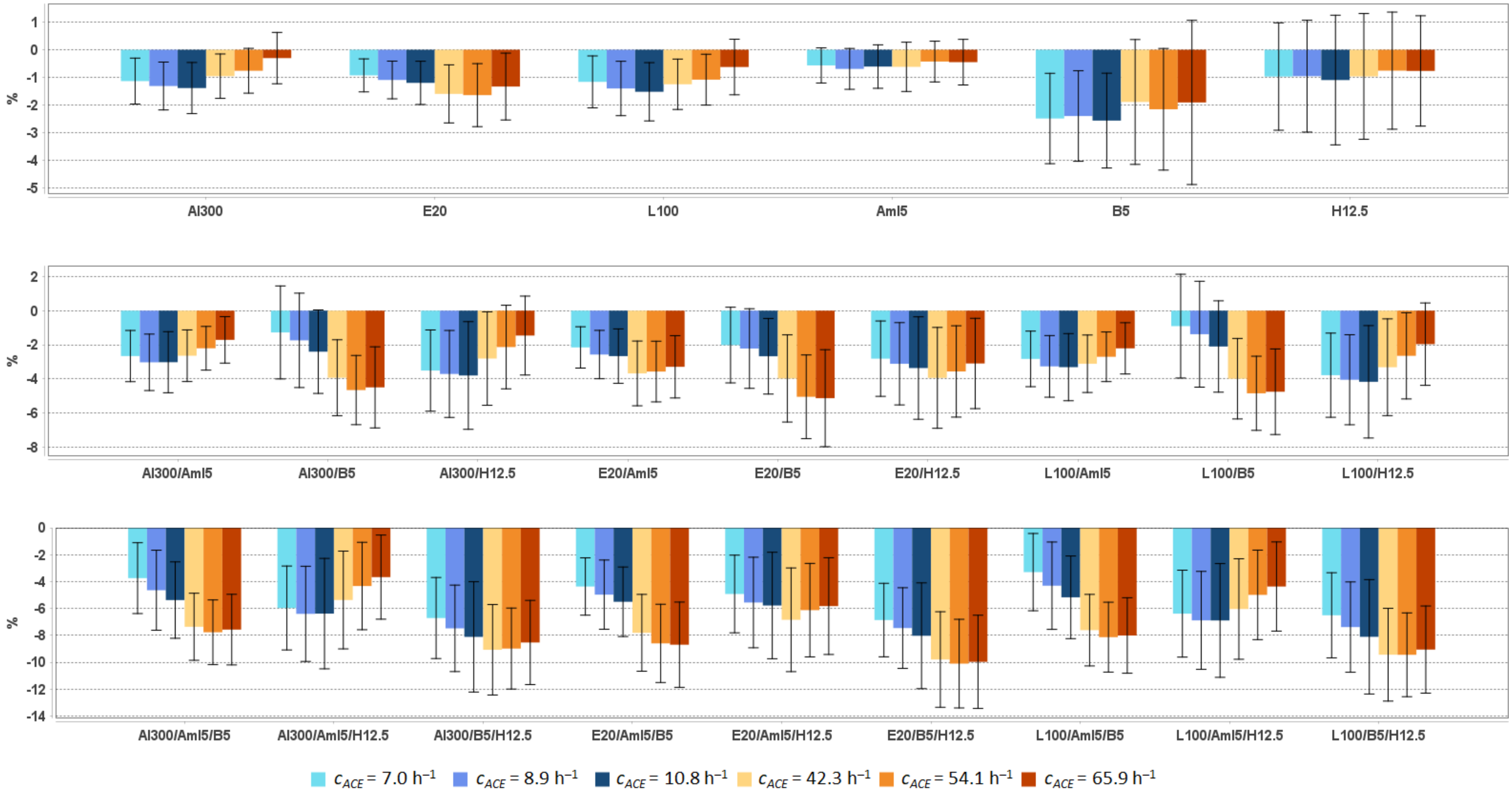


**Al300 =** aliskiren 300 mg; **Aml5** = amlodipine 5 mg; **B5** = bisoprolol 5 mg; **E20** = enalapril 20 mg; **H12.5** = hydrochlorothiazide 12.5 mg; **L100** = losartan 100 mg

**Table S18.** Simulated response of <u>ejection fraction</u> to antihypertensive therapy in virtual hypertensive populations ($n$ = 100) with different ACE activity, including $P$-values (Kolmogorov-Smirnov test) for endpoint *vs*. baseline; data are presented as mean ± SD in %

| **Regimens** | $c_{ACE}$ = 7.0 h$^{-1}$ | | | $c_{ACE}$ = 8.9 h$^{-1}$ | | | $c_{ACE}$ = 10.8 h$^{-1}$ | | | $c_{ACE}$ = 42.3 h$^{-1}$ | | | $c_{ACE}$ = 54.1 h$^{-1}$ | | | $c_{ACE}$ = 65.9 h$^{-1}$ | | |
|---|---|---|---|---|---|---|---|---|---|---|---|---|---|---|---|---|---|---|
| | **Value** | **Change** | ***P*** | **Value** | **Change** | ***P*** | **Value** | **Change** | ***P*** | **Value** | **Change** | ***P*** | **Value** | **Change** | ***P*** | **Value** | **Change** | ***P*** |
| Baseline | 66.6 ± 5.6 | – | – | 66.4 ± 6.2 | – | – | 66.7 ± 6.5 | – | – | 66.3 ± 6.8 | – | – | 66.5 ± 6.2 | – | – | 65.5 ± 6.6 | – | – |
| Al300 | 65.5 ± 5.7 | -1.1 ± 0.8 | 0.28093 | 65.1 ± 6.2 | -1.3 ± 0.9 | 0.46756 | 65.3 ± 6.5 | -1.4 ± 0.9 | 0.28093 | 65.4 ± 6.8 | -1.0 ± 0.8 | 0.90621 | 65.7 ± 6.3 | -0.8 ± 0.8 | 0.90621 | 65.2 ± 6.6 | -0.3 ± 0.9 | 0.99376 |
| E20 | 65.7 ± 5.7 | -0.9 ± 0.6 | 0.46756 | 65.3 ± 6.2 | -1.1 ± 0.7 | 0.58062 | 65.5 ± 6.5 | -1.2 ± 0.8 | 0.28093 | 64.7 ± 6.8 | -1.6 ± 1.1 | 0.36672 | 64.8 ± 6.4 | -1.6 ± 1.1 | 0.36672 | 64.2 ± 6.6 | -1.3 ± 1.2 | 0.36672 |
| L100 | 65.4 ± 5.8 | -1.2 ± 0.9 | 0.28093 | 65.0 ± 6.2 | -1.4 ± 1.0 | 0.28093 | 65.2 ± 6.5 | -1.5 ± 1.1 | 0.21055 | 65.1 ± 6.8 | -1.2 ± 0.9 | 0.69937 | 65.4 ± 6.4 | -1.1 ± 0.9 | 0.81275 | 64.9 ± 6.6 | -0.6 ± 1.0 | 0.90621 |
| Aml5 | 66.0 ± 5.7 | -0.6 ± 0.6 | 0.81275 | 65.7 ± 6.1 | -0.7 ± 0.7 | 0.90621 | 66.1 ± 6.3 | -0.6 ± 0.8 | 0.69937 | 65.7 ± 6.9 | -0.6 ± 0.9 | 0.81275 | 66.0 ± 6.3 | -0.4 ± 0.7 | 0.96707 | 65.1 ± 6.7 | -0.4 ± 0.8 | 0.96707 |
| B5 | 64.1 ± 5.4 | -2.5 ± 1.6 | 0.05410 | 64.0 ± 6.2 | -2.4 ± 1.6 | 0.02431 | 64.1 ± 6.5 | -2.6 ± 1.7 | 0.07832 | 64.5 ± 6.8 | -1.9 ± 2.3 | 0.28093 | 64.3 ± 5.9 | -2.2 ± 2.2 | 0.21055 | 63.6 ± 6.4 | -1.9 ± 3.0 | 0.28093 |
| H12.5 | 65.6 ± 6.0 | -1.0 ± 1.9 | 0.46756 | 65.4 ± 6.7 | -1.0 ± 2.0 | 0.28093 | 65.6 ± 6.6 | -1.1 ± 2.3 | 0.69937 | 65.4 ± 7.3 | -1.0 ± 2.3 | 0.69937 | 65.7 ± 6.5 | -0.8 ± 2.1 | 0.81275 | 64.8 ± 7.0 | -0.8 ± 2.0 | 0.58062 |
| Al300 Aml5 | 63.9 ± 6.0 | -2.7 ± 1.5 | 0.00386 | 63.4 ± 6.3 | -3.0 ± 1.7 | 0.00630 | 63.7 ± 6.5 | -3.0 ± 1.8 | 0.00630 | 63.7 ± 6.9 | -2.6 ± 1.5 | 0.05410 | 64.3 ± 6.5 | -2.2 ± 1.3 | 0.15454 | 63.8 ± 6.7 | -1.7 ± 1.4 | 0.15454 |
| Al300 B5 | 65.3 ± 5.9 | -1.3 ± 2.7 | 0.28093 | 64.6 ± 6.6 | -1.7 ± 2.8 | 0.21055 | 64.3 ± 6.7 | -2.4 ± 2.4 | 0.05410 | 62.4 ± 6.6 | -3.9 ± 2.2 | 0.01581 | 61.8 ± 6.1 | -4.6 ± 2.0 | 0.00025 | 61.0 ± 6.1 | -4.5 ± 2.4 | 0.00045 |
| Al300 H12.5 | 63.1 ± 6.2 | -3.5 ± 2.4 | 0.00007 | 62.7 ± 6.9 | -3.7 ± 2.5 | 0.00013 | 62.9 ± 6.8 | -3.8 ± 3.1 | 0.01008 | 63.5 ± 7.3 | -2.8 ± 2.7 | 0.05410 | 64.3 ± 6.7 | -2.1 ± 2.4 | 0.21055 | 64.1 ± 7.0 | -1.5 ± 2.3 | 0.11113 |
| E20 Aml5 | 64.5 ± 5.8 | -2.1 ± 1.2 | 0.02431 | 63.8 ± 6.3 | -2.6 ± 1.4 | 0.02431 | 64.0 ± 6.4 | -2.7 ± 1.6 | 0.01008 | 62.7 ± 7.0 | -3.7 ± 1.9 | 0.00386 | 62.9 ± 6.7 | -3.6 ± 1.8 | 0.00386 | 62.2 ± 6.7 | -3.3 ± 1.8 | 0.01008 |
| E20 B5 | 64.6 ± 5.7 | -2.0 ± 2.2 | 0.21055 | 64.2 ± 6.4 | -2.2 ± 2.3 | 0.05410 | 64.0 ± 6.6 | -2.7 ± 2.2 | 0.03663 | 62.4 ± 6.7 | -4.0 ± 2.6 | 0.01008 | 61.4 ± 6.3 | -5.0 ± 2.4 | 0.00013 | 60.4 ± 6.3 | -5.1 ± 2.8 | 0.00004 |
| E20 H12.5 | 63.8 ± 6.1 | -2.8 ± 2.2 | 0.00136 | 63.3 ± 6.8 | -3.1 ± 2.4 | 0.00079 | 63.3 ± 6.8 | -3.4 ± 3.0 | 0.01581 | 62.4 ± 7.4 | -3.9 ± 3.0 | 0.00136 | 62.9 ± 6.9 | -3.6 ± 2.7 | 0.00630 | 62.4 ± 7.1 | -3.1 ± 2.6 | 0.00386 |
| L100 Aml5 | 63.8 ± 6.0 | -2.8 ± 1.6 | 0.00386 | 63.1 ± 6.4 | -3.3 ± 1.8 | 0.00232 | 63.4 ± 6.5 | -3.3 ± 2.0 | 0.00630 | 63.2 ± 6.9 | -3.1 ± 1.7 | 0.02431 | 63.8 ± 6.6 | -2.7 ± 1.5 | 0.05410 | 63.3 ± 6.7 | -2.2 ± 1.5 | 0.11113 |
| L100 B5 | 65.7 ± 6.1 | -0.9 ± 3.0 | 0.69937 | 65.0 ± 6.8 | -1.4 ± 3.1 | 0.36672 | 64.6 ± 6.8 | -2.1 ± 2.7 | 0.07832 | 62.4 ± 6.7 | -4.0 ± 2.4 | 0.01581 | 61.6 ± 6.2 | -4.8 ± 2.2 | 0.00013 | 60.8 ± 6.2 | -4.7 ± 2.5 | 0.00007 |
| L100 H12.5 | 62.8 ± 6.3 | -3.8 ± 2.5 | 0.00007 | 62.3 ± 6.9 | -4.0 ± 2.6 | 0.00007 | 62.5 ± 6.9 | -4.2 ± 3.3 | 0.00386 | 63.0 ± 7.4 | -3.3 ± 2.8 | 0.01008 | 63.8 ± 6.8 | -2.6 ± 2.5 | 0.05410 | 63.6 ± 7.0 | -2.0 ± 2.4 | 0.05410 |
| Al300 Aml5/B5 | 62.9 ± 6.1 | -3.7 ± 2.6 | 0.00013 | 61.7 ± 6.5 | -4.6 ± 3.0 | SS | 61.3 ± 6.6 | -5.4 ± 2.8 | 0.00002 | 59.0 ± 6.6 | -7.4 ± 2.5 | SS | 58.7 ± 6.5 | -7.8 ± 2.4 | SS | 57.9 ± 6.3 | -7.6 ± 2.6 | SS |
| Al300 Aml5/H12.5 | 60.6 ± 6.7 | -6.0 ± 3.1 | SS | 60.0 ± 7.3 | -6.4 ± 3.5 | SS | 60.3 ± 7.1 | -6.4 ± 4.1 | SS | 61.0 ± 7.8 | -5.4 ± 3.6 | 0.00004 | 62.1 ± 7.2 | -4.3 ± 3.2 | 0.00079 | 61.9 ± 7.4 | -3.7 ± 3.1 | 0.00386 |
| Al300 B5/H12.5 | 59.9 ± 6.3 | -6.7 ± 3.0 | SS | 58.9 ± 6.8 | -7.5 ± 3.2 | SS | 58.6 ± 6.9 | -8.1 ± 4.1 | SS | 57.3 ± 7.1 | -9.1 ± 3.4 | SS | 57.5 ± 6.6 | -9.0 ± 3.0 | SS | 57.0 ± 6.6 | -8.5 ± 3.1 | SS |
| E20 Aml5/B5 | 62.2 ± 5.8 | -4.4 ± 2.1 | 0.00007 | 61.4 ± 6.3 | -5.0 ± 2.6 | SS | 61.2 ± 6.4 | -5.5 ± 2.6 | 0.00002 | 58.5 ± 6.7 | -7.8 ± 2.9 | SS | 57.9 ± 6.7 | -8.6 ± 2.9 | SS | 56.8 ± 6.5 | -8.7 ± 3.2 | SS |
| E20 Aml5/H12.5 | 61.7 ± 6.5 | -4.9 ± 2.9 | SS | 60.8 ± 7.2 | -5.6 ± 3.4 | SS | 60.9 ± 7.0 | -5.8 ± 4.0 | 0.00004 | 59.5 ± 7.9 | -6.8 ± 3.9 | SS | 60.3 ± 7.4 | -6.1 ± 3.5 | SS | 59.7 ± 7.7 | -5.8 ± 3.6 | SS |
| E20 B5/H12.5 | 59.7 ± 6.0 | -6.9 ± 2.7 | SS | 58.9 ± 6.7 | -7.5 ± 3.0 | SS | 58.7 ± 6.8 | -8.0 ± 3.9 | SS | 56.6 ± 7.2 | -9.8 ± 3.6 | SS | 56.4 ± 6.8 | -10.1 ± 3.3 | SS | 55.6 ± 6.8 | -10.0 ± 3.5 | SS |
| L100 Aml5/B5 | 63.3 ± 6.2 | -3.3 ± 2.9 | 0.00079 | 62.1 ± 6.6 | -4.3 ± 3.2 | SS | 61.5 ± 6.7 | -5.2 ± 3.1 | 0.00007 | 58.7 ± 6.7 | -7.6 ± 2.7 | SS | 58.3 ± 6.6 | -8.1 ± 2.6 | SS | 57.5 ± 6.3 | -8.0 ± 2.8 | SS |
| L100 Aml5/H12.5 | 60.2 ± 6.8 | -6.4 ± 3.2 | SS | 59.5 ± 7.3 | -6.9 ± 3.6 | SS | 59.8 ± 7.2 | -6.9 ± 4.2 | SS | 60.3 ± 7.8 | -6.0 ± 3.7 | SS | 61.5 ± 7.3 | -5.0 ± 3.3 | 0.00013 | 61.2 ± 7.5 | -4.4 ± 3.3 | 0.00045 |
| L100 B5/H12.5 | 60.1 ± 6.4 | -6.5 ± 3.2 | SS | 59.0 ± 6.9 | -7.4 ± 3.4 | SS | 58.6 ± 7.0 | -8.1 ± 4.3 | SS | 56.9 ± 7.1 | -9.4 ± 3.4 | SS | 57.0 ± 6.7 | -9.4 ± 3.1 | SS | 56.5 ± 6.7 | -9.1 ± 3.2 | SS |

**Al300 =** aliskiren 300 mg; **Aml5** = amlodipine 5 mg; **B5** = bisoprolol 5 mg; **E20** = enalapril 20 mg; **H12.5** = hydrochlorothiazide 12.5 mg; **L100** = losartan 100 mg; **SS** = statistically significant ($P$ < 0.00001)

**Table S19.** *P*-values calculated using the Kolmogorov-Smirnov test for changes in ejection fraction in populations ($n$ = 100) with different ACE activity receiving the same regimens (case 1: $c_{ACE}$ = 7.0 $h^{-1}$, case 2: $c_{ACE}$ = 8.9 $h^{-1}$, case 3: $c_{ACE}$ = 10.8 $h^{-1}$, case 4: $c_{ACE}$ = 42.3 $h^{-1}$, case 5: $c_{ACE}$ = 54.1 $h^{-1}$, case 6: $c_{ACE}$ = 65.9 $h^{-1}$; *P*-value for case *i vs*. case *j* is denoted $P_{ij}$)

| Regimens | $P_{12}$ | $P_{13}$ | $P_{23}$ | $P_{14}$ | $P_{15}$ | $P_{16}$ | $P_{24}$ | $P_{25}$ | $P_{26}$ | $P_{34}$ | $P_{35}$ | $P_{36}$ | $P_{45}$ | $P_{46}$ | $P_{56}$ |
|---|---|---|---|---|---|---|---|---|---|---|---|---|---|---|---|
| Al300 | 0.15454 | 0.05410 | 0.81275 | 0.28093 | 0.03663 | SS | 0.05410 | 0.00025 | SS | 0.07832 | 0.00004 | SS | 0.03663 | 0.00004 | 0.00630 |
| E20 | 0.03663 | 0.01581 | 0.69937 | SS | SS | 0.00025 | 0.00045 | 0.00630 | 0.00232 | 0.00045 | 0.01008 | 0.03663 | 0.58062 | 0.21055 | 0.28093 |
| L100 | 0.07832 | 0.03663 | 0.58062 | 0.15454 | 0.81275 | 0.00386 | 0.46756 | 0.02431 | 0.00045 | 0.28093 | 0.03663 | 0.00004 | 0.03663 | 0.00025 | 0.02431 |
| Aml5 | 0.46756 | 0.69937 | 0.36672 | 0.58062 | 0.21055 | 0.07832 | 0.36672 | 0.07832 | 0.02431 | 0.99376 | 0.21055 | 0.21055 | 0.15454 | 0.28093 | 0.99376 |
| B5 | 0.90621 | 0.96707 | 0.90621 | 0.15454 | 0.15454 | 0.01581 | 0.15454 | 0.28093 | 0.01581 | 0.03663 | 0.07832 | 0.00232 | 0.69937 | 0.36672 | 0.46756 |
| H12.5 | 0.81275 | 0.90621 | 0.81275 | 0.21055 | 0.58062 | 0.69937 | 0.21055 | 0.03663 | 0.07832 | 0.69937 | 0.28093 | 0.28093 | 0.11113 | 0.36672 | 0.90621 |
| Al300 Aml5 | 0.05410 | 0.11113 | 0.69937 | 0.90621 | 0.21055 | 0.00136 | 0.03663 | 0.00079 | SS | 0.15454 | 0.00630 | 0.00013 | 0.05410 | 0.00045 | 0.05410 |
| Al300 B5 | 0.69937 | 0.03663 | 0.03663 | SS | SS | SS | SS | SS | SS | 0.00045 | SS | SS | 0.15454 | 0.28093 | 0.90621 |
| Al300 H12.5 | 0.58062 | 0.58062 | 0.69937 | 0.07832 | 0.00079 | SS | 0.05410 | 0.00079 | SS | 0.21055 | 0.00045 | SS | 0.11113 | 0.00232 | 0.11113 |
| E20 Aml5 | 0.03663 | 0.05410 | 0.81275 | SS | SS | 0.00002 | 0.00045 | 0.00630 | 0.02431 | 0.00007 | 0.00232 | 0.02431 | 0.58062 | 0.15454 | 0.46756 |
| E20 B5 | 0.81275 | 0.05410 | 0.11113 | SS | SS | SS | 0.00002 | SS | SS | 0.00136 | SS | SS | 0.05410 | 0.01008 | 0.69937 |
| E20 H12.5 | 0.36672 | 0.11113 | 0.46756 | 0.00232 | 0.03663 | 0.46756 | 0.03663 | 0.69937 | 0.90621 | 0.07832 | 0.36672 | 0.58062 | 0.21055 | 0.15454 | 0.46756 |
| L100 Aml5 | 0.02431 | 0.07832 | 0.90621 | 0.28093 | 0.90621 | 0.03663 | 0.36672 | 0.01008 | 0.00045 | 0.69937 | 0.03663 | 0.00386 | 0.07832 | 0.00386 | 0.01581 |
| L100 B5 | 0.58062 | 0.05410 | 0.03663 | SS | SS | SS | SS | SS | SS | 0.00004 | SS | SS | 0.07832 | 0.15454 | 0.90621 |
| L100 H12.5 | 0.36672 | 0.46756 | 0.69937 | 0.21055 | 0.01008 | SS | 0.11113 | 0.00386 | SS | 0.58062 | 0.00386 | 0.00007 | 0.11113 | 0.00386 | 0.21055 |
| Al300 Aml5/B5 | 0.01581 | 0.00025 | 0.28093 | SS | SS | SS | SS | SS | SS | SS | SS | SS | 0.36672 | 0.96707 | 0.69937 |
| Al300 Aml5/H12.5 | 0.15454 | 0.69937 | 0.21055 | 0.28093 | 0.00136 | 0.00004 | 0.02431 | 0.00025 | SS | 0.28093 | 0.00386 | SS | 0.21055 | 0.01008 | 0.15454 |
| Al300 B5/H12.5 | 0.02431 | 0.03663 | 0.28093 | SS | SS | 0.00007 | 0.01581 | 0.02431 | 0.15454 | 0.02431 | 0.01008 | 0.21055 | 0.69937 | 0.69937 | 0.69937 |
| E20 Aml5/B5 | 0.03663 | 0.00079 | 0.21055 | SS | SS | SS | SS | SS | SS | SS | SS | SS | 0.21055 | 0.28093 | 0.58062 |
| E20 Aml5/H12.5 | 0.15454 | 0.28093 | 0.90621 | 0.00079 | 0.00386 | 0.05410 | 0.05410 | 0.46756 | 0.69937 | 0.01581 | 0.11113 | 0.69937 | 0.36672 | 0.11113 | 0.69937 |
| E20 B5/H12.5 | 0.05410 | 0.05410 | 0.28093 | SS | SS | SS | 0.00013 | 0.00004 | SS | 0.00079 | 0.00007 | 0.00025 | 0.69937 | 0.81275 | 0.58062 |
| L100 Aml5/B5 | 0.05410 | 0.00045 | 0.28093 | SS | SS | SS | SS | SS | SS | SS | SS | SS | 0.28093 | 0.69937 | 0.69937 |
| L100 Aml5/H12.5 | 0.15454 | 0.58062 | 0.36672 | 0.58062 | 0.01008 | 0.00013 | 0.07832 | 0.00045 | 0.00002 | 0.58062 | 0.01581 | 0.00013 | 0.15454 | 0.01008 | 0.21055 |
| L100 B5/H12.5 | 0.01008 | 0.02431 | 0.15454 | SS | SS | SS | 0.00386 | 0.00232 | 0.01581 | 0.00136 | 0.00232 | 0.01008 | 0.46756 | 0.90621 | 0.58062 |

**Al300 =** aliskiren 300 mg; **Aml5** = amlodipine 5 mg; **B5** = bisoprolol 5 mg; **E20** = enalapril 20 mg; **H12.5** = hydrochlorothiazide 12.5 mg; **L100** = losartan 100 mg; **SS** = statistically significant ($P$ < 0.00001)

**Figure S17.** Simulated change in <u>left ventricular end-diastolic pressure</u> from baseline to week 4 (mean ± SD, $n$ = 100)

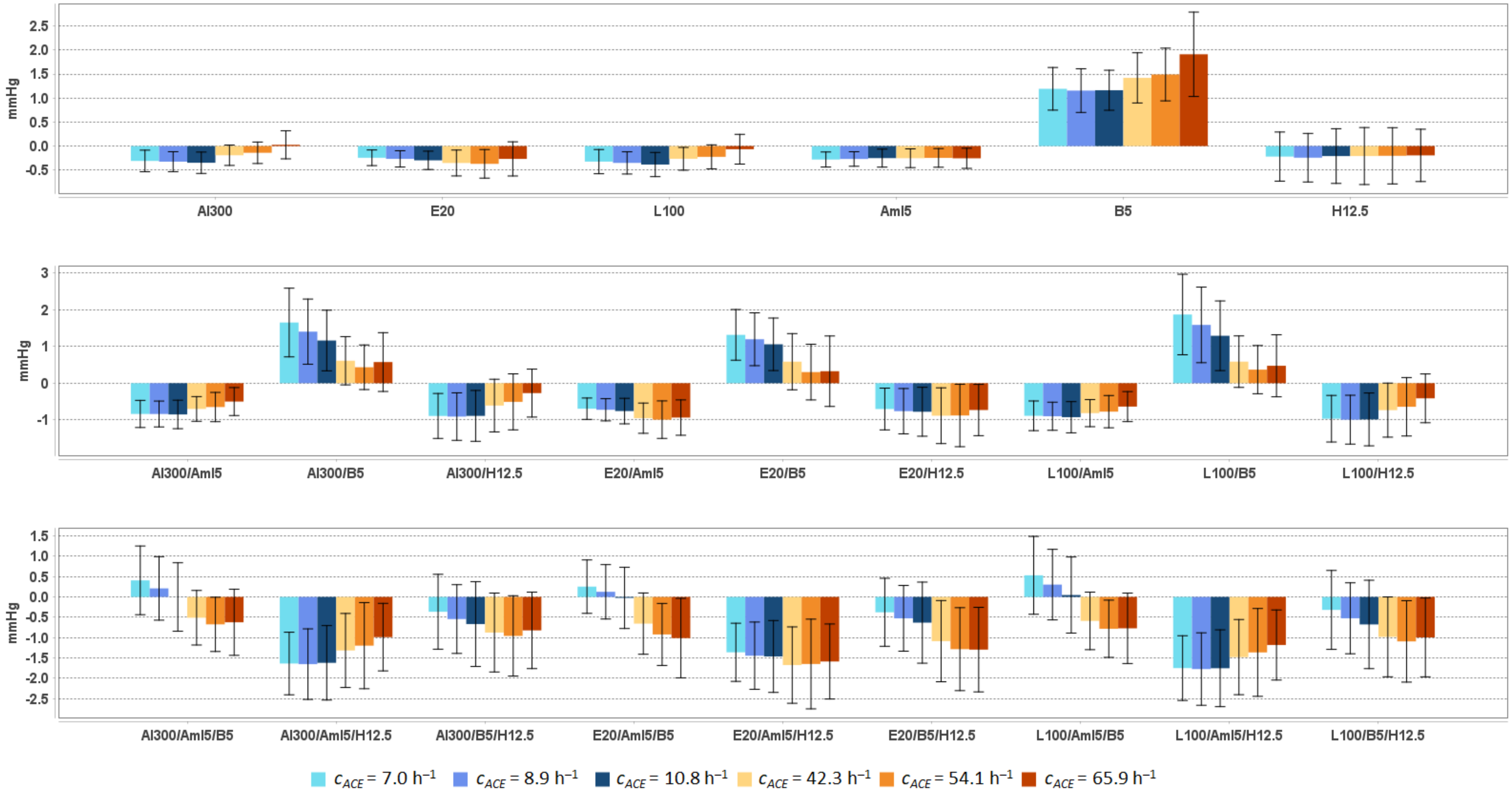


**Al300 =** aliskiren 300 mg; **Aml5** = amlodipine 5 mg; **B5** = bisoprolol 5 mg; **E20** = enalapril 20 mg; **H12.5** = hydrochlorothiazide 12.5 mg; **L100** = losartan 100 mg

**Table S20.** Simulated response of <u>left ventricular end-diastolic pressure</u> to antihypertensive therapy in virtual hypertensive populations ($n$ = 100) with different ACE activity, including $P$-values (Kolmogorov-Smirnov test) for endpoint *vs.* baseline; data are presented as mean ± SD in mmHg

| **Regimens** | **$c_{ACE}$ = 7.0 h$^{-1}$** | | | **$c_{ACE}$ = 8.9 h$^{-1}$** | | | **$c_{ACE}$ = 10.8 h$^{-1}$** | | | **$c_{ACE}$ = 42.3 h$^{-1}$** | | | **$c_{ACE}$ = 54.1 h$^{-1}$** | | | **$c_{ACE}$ = 65.9 h$^{-1}$** | | |
|---|---|---|---|---|---|---|---|---|---|---|---|---|---|---|---|---|---|---|
| | **Value** | **Change** | ***P*** | **Value** | **Change** | ***P*** | **Value** | **Change** | ***P*** | **Value** | **Change** | ***P*** | **Value** | **Change** | ***P*** | **Value** | **Change** | ***P*** |
| Baseline | 8.1 ± 1.3 | – | – | 8.0 ± 1.2 | – | – | 8.1 ± 1.1 | – | – | 8.0 ± 1.0 | – | – | 8.0 ± 1.1 | – | – | 8.2 ± 1.0 | – | – |
| Al300 | 7.7 ± 1.3 | -0.3 ± 0.2 | 0.11113 | 7.7 ± 1.2 | -0.3 ± 0.2 | 0.05410 | 7.7 ± 1.1 | -0.3 ± 0.2 | 0.03663 | 7.8 ± 1.0 | -0.2 ± 0.2 | 0.36672 | 7.9 ± 1.1 | -0.1 ± 0.2 | 0.69937 | 8.2 ± 1.1 | 0.0 ± 0.3 | 0.36672 |
| E20 | 7.8 ± 1.3 | -0.2 ± 0.2 | 0.15454 | 7.7 ± 1.2 | -0.3 ± 0.2 | 0.11113 | 7.8 ± 1.1 | -0.3 ± 0.2 | 0.05410 | 7.7 ± 1.0 | -0.4 ± 0.3 | 0.05410 | 7.6 ± 1.1 | -0.4 ± 0.3 | 0.03663 | 7.9 ± 1.1 | -0.3 ± 0.4 | 0.15454 |
| L100 | 7.7 ± 1.3 | -0.3 ± 0.3 | 0.05410 | 7.6 ± 1.2 | -0.4 ± 0.2 | 0.05410 | 7.7 ± 1.1 | -0.4 ± 0.3 | 0.01581 | 7.8 ± 1.0 | -0.3 ± 0.2 | 0.15454 | 7.8 ± 1.1 | -0.2 ± 0.2 | 0.21055 | 8.1 ± 1.1 | -0.1 ± 0.3 | 0.90621 |
| Aml5 | 7.8 ± 1.3 | -0.3 ± 0.2 | 0.11113 | 7.7 ± 1.2 | -0.3 ± 0.2 | 0.11113 | 7.8 ± 1.1 | -0.3 ± 0.2 | 0.11113 | 7.8 ± 1.0 | -0.3 ± 0.2 | 0.11113 | 7.7 ± 1.1 | -0.2 ± 0.2 | 0.15454 | 8.0 ± 1.0 | -0.3 ± 0.2 | 0.01581 |
| B5 | 9.2 ± 1.6 | 1.2 ± 0.4 | SS | 9.1 ± 1.4 | 1.2 ± 0.5 | SS | 9.2 ± 1.3 | 1.2 ± 0.4 | SS | 9.5 ± 1.3 | 1.4 ± 0.5 | SS | 9.5 ± 1.5 | 1.5 ± 0.5 | SS | 10.1 ± 1.5 | 1.9 ± 0.9 | SS |
| H12.5 | 7.8 ± 1.4 | -0.2 ± 0.5 | 0.28093 | 7.7 ± 1.2 | -0.2 ± 0.5 | 0.05410 | 7.8 ± 1.2 | -0.2 ± 0.6 | 0.21055 | 7.8 ± 1.2 | -0.2 ± 0.6 | 0.11113 | 7.8 ± 1.2 | -0.2 ± 0.6 | 0.11113 | 8.0 ± 1.1 | -0.2 ± 0.5 | 0.11113 |
| Al300<br>Aml5 | 7.2 ± 1.2 | -0.8 ± 0.4 | SS | 7.1 ± 1.1 | -0.8 ± 0.4 | SS | 7.2 ± 1.0 | -0.9 ± 0.4 | SS | 7.3 ± 0.9 | -0.7 ± 0.3 | SS | 7.3 ± 1.1 | -0.7 ± 0.4 | 0.00007 | 7.7 ± 1.0 | -0.5 ± 0.4 | 0.00007 |
| Al300<br>B5 | 9.7 ± 1.8 | 1.7 ± 0.9 | SS | 9.4 ± 1.7 | 1.4 ± 0.9 | SS | 9.2 ± 1.6 | 1.2 ± 0.8 | SS | 8.6 ± 1.3 | 0.6 ± 0.7 | 0.00013 | 8.4 ± 1.4 | 0.4 ± 0.6 | 0.07832 | 8.8 ± 1.4 | 0.6 ± 0.8 | SS |
| Al300<br>H12.5 | 7.2 ± 1.3 | -0.9 ± 0.6 | SS | 7.1 ± 1.1 | -0.9 ± 0.6 | SS | 7.2 ± 1.1 | -0.9 ± 0.7 | SS | 7.4 ± 1.2 | -0.6 ± 0.7 | 0.00025 | 7.5 ± 1.2 | -0.5 ± 0.8 | 0.00079 | 7.9 ± 1.2 | -0.3 ± 0.7 | 0.07832 |
| E20<br>Aml5 | 7.4 ± 1.2 | -0.7 ± 0.3 | SS | 7.3 ± 1.1 | -0.7 ± 0.3 | SS | 7.3 ± 1.0 | -0.8 ± 0.3 | SS | 7.1 ± 0.9 | -1.0 ± 0.4 | SS | 7.0 ± 1.1 | -1.0 ± 0.5 | SS | 7.3 ± 1.0 | -0.9 ± 0.5 | SS |
| E20<br>B5 | 9.4 ± 1.7 | 1.3 ± 0.7 | SS | 9.2 ± 1.6 | 1.2 ± 0.7 | SS | 9.1 ± 1.5 | 1.1 ± 0.7 | SS | 8.6 ± 1.3 | 0.6 ± 0.8 | 0.00025 | 8.3 ± 1.4 | 0.3 ± 0.8 | 0.15454 | 8.5 ± 1.4 | 0.3 ± 1.0 | 0.00386 |
| E20<br>H12.5 | 7.3 ± 1.3 | -0.7 ± 0.6 | 0.00025 | 7.2 ± 1.1 | -0.8 ± 0.6 | SS | 7.3 ± 1.2 | -0.8 ± 0.7 | SS | 7.1 ± 1.2 | -0.9 ± 0.8 | SS | 7.1 ± 1.2 | -0.9 ± 0.9 | SS | 7.5 ± 1.2 | -0.7 ± 0.7 | 0.00004 |
| L100<br>Aml5 | 7.2 ± 1.2 | -0.9 ± 0.4 | SS | 7.1 ± 1.1 | -0.9 ± 0.4 | SS | 7.1 ± 1.0 | -0.9 ± 0.4 | SS | 7.2 ± 0.9 | -0.8 ± 0.4 | SS | 7.2 ± 1.1 | -0.8 ± 0.4 | SS | 7.6 ± 1.0 | -0.6 ± 0.4 | 0.00002 |
| L100<br>B5 | 9.9 ± 1.9 | 1.9 ± 1.1 | SS | 9.6 ± 1.8 | 1.6 ± 1.0 | SS | 9.3 ± 1.7 | 1.3 ± 0.9 | SS | 8.6 ± 1.3 | 0.6 ± 0.7 | 0.00025 | 8.4 ± 1.4 | 0.4 ± 0.7 | 0.11113 | 8.7 ± 1.4 | 0.5 ± 0.8 | 0.00007 |
| L100<br>H12.5 | 7.1 ± 1.3 | -1.0 ± 0.6 | SS | 7.0 ± 1.1 | -1.0 ± 0.7 | SS | 7.1 ± 1.1 | -1.0 ± 0.7 | SS | 7.3 ± 1.2 | -0.7 ± 0.7 | 0.00002 | 7.3 ± 1.2 | -0.6 ± 0.8 | 0.00007 | 7.8 ± 1.2 | -0.4 ± 0.7 | 0.03663 |
| Al300<br>Aml5/B5 | 8.5 ± 1.6 | 0.4 ± 0.8 | 0.01581 | 8.2 ± 1.5 | 0.2 ± 0.8 | 0.21055 | 8.1 ± 1.4 | -0.0 ± 0.8 | 0.46756 | 7.5 ± 1.1 | -0.5 ± 0.7 | 0.00386 | 7.3 ± 1.2 | -0.7 ± 0.7 | 0.00013 | 7.6 ± 1.2 | -0.6 ± 0.8 | 0.00007 |
| Al300<br>Aml5/H12.5 | 6.4 ± 1.2 | -1.6 ± 0.8 | SS | 6.3 ± 1.1 | -1.7 ± 0.9 | SS | 6.4 ± 1.2 | -1.6 ± 0.9 | SS | 6.7 ± 1.2 | -1.3 ± 0.9 | SS | 6.8 ± 1.3 | -1.2 ± 1.1 | SS | 7.2 ± 1.2 | -1.0 ± 0.8 | SS |
| Al300<br>B5/H12.5 | 7.7 ± 1.5 | -0.4 ± 0.9 | 0.07832 | 7.4 ± 1.3 | -0.5 ± 0.8 | 0.00136 | 7.4 ± 1.4 | -0.7 ± 1.0 | 0.00002 | 7.2 ± 1.3 | -0.9 ± 1.0 | SS | 7.0 ± 1.3 | -1.0 ± 1.0 | SS | 7.4 ± 1.3 | -0.8 ± 0.9 | SS |
| E20<br>Aml5/B5 | 8.3 ± 1.5 | 0.3 ± 0.7 | 0.11113 | 8.1 ± 1.4 | 0.1 ± 0.7 | 0.46756 | 8.0 ± 1.4 | -0.0 ± 0.8 | 0.58062 | 7.4 ± 1.2 | -0.7 ± 0.8 | 0.00013 | 7.1 ± 1.2 | -0.9 ± 0.8 | SS | 7.2 ± 1.3 | -1.0 ± 1.0 | SS |
| E20<br>Aml5/H12.5 | 6.7 ± 1.2 | -1.4 ± 0.7 | SS | 6.5 ± 1.1 | -1.4 ± 0.8 | SS | 6.6 ± 1.2 | -1.5 ± 0.9 | SS | 6.4 ± 1.2 | -1.7 ± 0.9 | SS | 6.3 ± 1.3 | -1.6 ± 1.1 | SS | 6.6 ± 1.2 | -1.6 ± 0.9 | SS |
| E20<br>B5/H12.5 | 7.7 ± 1.5 | -0.4 ± 0.8 | 0.07832 | 7.5 ± 1.3 | -0.5 ± 0.8 | 0.00232 | 7.4 ± 1.4 | -0.6 ± 1.0 | 0.00013 | 6.9 ± 1.3 | -1.1 ± 1.0 | SS | 6.7 ± 1.3 | -1.3 ± 1.0 | SS | 6.9 ± 1.3 | -1.3 ± 1.0 | SS |
| L100<br>Aml5/B5 | 8.6 ± 1.7 | 0.5 ± 1.0 | 0.00232 | 8.3 ± 1.5 | 0.3 ± 0.9 | 0.07832 | 8.1 ± 1.5 | 0.0 ± 0.9 | 0.28093 | 7.4 ± 1.1 | -0.6 ± 0.7 | 0.00079 | 7.2 ± 1.2 | -0.8 ± 0.7 | SS | 7.4 ± 1.2 | -0.8 ± 0.9 | SS |
| L100<br>Aml5/H12.5 | 6.3 ± 1.2 | -1.8 ± 0.8 | SS | 6.2 ± 1.1 | -1.8 ± 0.9 | SS | 6.3 ± 1.2 | -1.8 ± 0.9 | SS | 6.6 ± 1.2 | -1.5 ± 0.9 | SS | 6.6 ± 1.3 | -1.4 ± 1.1 | SS | 7.0 ± 1.2 | -1.2 ± 0.9 | SS |
| L100<br>B5/H12.5 | 7.7 ± 1.6 | -0.3 ± 1.0 | 0.07832 | 7.5 ± 1.3 | -0.5 ± 0.9 | 0.00232 | 7.4 ± 1.5 | -0.7 ± 1.1 | 0.00004 | 7.1 ± 1.3 | -1.0 ± 1.0 | SS | 6.9 ± 1.3 | -1.1 ± 1.0 | SS | 7.2 ± 1.3 | -1.0 ± 1.0 | SS |

**Al300 =** aliskiren 300 mg; **Aml5** = amlodipine 5 mg; **B5** = bisoprolol 5 mg; **E20** = enalapril 20 mg; **H12.5** = hydrochlorothiazide 12.5 mg; **L100** = losartan 100 mg; **SS** = statistically significant ($P$ < 0.00001)

**Table S21.** *P*-values calculated using the Kolmogorov-Smirnov test for changes in left ventricular end-diastolic pressure in populations ($n$ = 100) with different ACE activity receiving the same regimens (case 1: $c_{ACE}$ = 7.0 h$^{-1}$, case 2: $c_{ACE}$ = 8.9 h$^{-1}$, case 3: $c_{ACE}$ = 10.8 h$^{-1}$, case 4: $c_{ACE}$ = 42.3 h$^{-1}$, case 5: $c_{ACE}$ = 54.1 h$^{-1}$, case 6: $c_{ACE}$ = 65.9 h$^{-1}$; *P*-value for case $i$ *vs*. case $j$ is denoted $P_{ij}$)

| **Regimens** | $P_{12}$ | $P_{13}$ | $P_{23}$ | $P_{14}$ | $P_{15}$ | $P_{16}$ | $P_{24}$ | $P_{25}$ | $P_{26}$ | $P_{34}$ | $P_{35}$ | $P_{36}$ | $P_{45}$ | $P_{46}$ | $P_{56}$ |
|---|---|---|---|---|---|---|---|---|---|---|---|---|---|---|---|
| Al300 | 0.58062 | 0.15454 | 0.58062 | 0.01008 | 0.00004 | SS | 0.00232 | SS | SS | 0.00045 | SS | SS | 0.05410 | 0.00004 | 0.00136 |
| E20 | 0.36672 | 0.02431 | 0.28093 | 0.00025 | 0.00025 | 0.00386 | 0.01008 | 0.00386 | 0.03663 | 0.05410 | 0.01008 | 0.02431 | 0.90621 | 0.15454 | 0.11113 |
| L100 | 0.28093 | 0.11113 | 0.46756 | 0.28093 | 0.02431 | SS | 0.15454 | 0.00386 | SS | 0.02431 | 0.00025 | SS | 0.36672 | 0.00045 | 0.00630 |
| Aml5 | 0.96707 | 0.46756 | 0.58062 | 0.69937 | 0.11113 | 0.28093 | 0.81275 | 0.15454 | 0.58062 | 0.81275 | 0.69937 | 0.90621 | 0.46756 | 0.90621 | 0.81275 |
| B5 | 0.28093 | 0.58062 | 0.90621 | 0.01008 | 0.00079 | SS | 0.00232 | 0.00004 | SS | 0.01008 | 0.00045 | SS | 0.46756 | 0.00025 | 0.00136 |
| H12.5 | 0.58062 | 0.81275 | 0.36672 | 0.69937 | 0.81275 | 0.90621 | 0.21055 | 0.15454 | 0.15454 | 0.81275 | 0.46756 | 0.46756 | 0.58062 | 0.81275 | 0.90621 |
| Al300 Aml5 | 0.90621 | 0.96707 | 0.81275 | 0.07832 | 0.00004 | SS | 0.01581 | SS | SS | 0.05410 | 0.00002 | SS | 0.05410 | 0.00386 | 0.02431 |
| Al300 B5 | 0.11113 | 0.00232 | 0.03663 | SS | SS | SS | SS | SS | SS | 0.00045 | SS | SS | 0.11113 | 0.28093 | 0.36672 |
| Al300 H12.5 | 0.81275 | 0.58062 | 0.36672 | 0.01008 | SS | SS | 0.00232 | SS | SS | 0.07832 | 0.00013 | SS | 0.21055 | 0.00630 | 0.36672 |
| E20 Aml5 | 0.58062 | 0.15454 | 0.69937 | SS | 0.00013 | 0.00004 | 0.00013 | 0.00386 | 0.00386 | 0.00045 | 0.01581 | 0.00386 | 0.58062 | 0.90621 | 0.90621 |
| E20 B5 | 0.69937 | 0.01581 | 0.07832 | SS | SS | SS | SS | SS | SS | 0.00013 | SS | SS | 0.02431 | 0.05410 | 0.58062 |
| E20 H12.5 | 0.90621 | 0.28093 | 0.36672 | 0.03663 | 0.21055 | 0.28093 | 0.07832 | 0.21055 | 0.46756 | 0.69937 | 0.69937 | 0.90621 | 0.46756 | 0.36672 | 0.90621 |
| L100 Aml5 | 0.69937 | 0.81275 | 0.90621 | 0.90621 | 0.02431 | 0.00232 | 0.15454 | 0.00079 | 0.00007 | 0.21055 | 0.00136 | 0.00025 | 0.07832 | 0.01581 | 0.21055 |
| L100 B5 | 0.07832 | 0.00079 | 0.05410 | SS | SS | SS | SS | SS | SS | 0.00007 | SS | SS | 0.03663 | 0.28093 | 0.36672 |
| L100 H12.5 | 0.96707 | 0.36672 | 0.46756 | 0.02431 | 0.00004 | SS | 0.01008 | 0.00004 | SS | 0.11113 | 0.00045 | SS | 0.21055 | 0.02431 | 0.36672 |
| Al300 Aml5/B5 | 0.07832 | 0.01581 | 0.21055 | SS | SS | SS | SS | SS | SS | 0.00079 | SS | 0.00002 | 0.15454 | 0.36672 | 0.58062 |
| Al300 Aml5/H12.5 | 0.96707 | 0.46756 | 0.81275 | 0.00232 | SS | SS | 0.02431 | 0.00007 | SS | 0.03663 | 0.00045 | 0.00007 | 0.28093 | 0.02431 | 0.46756 |
| Al300 B5/H12.5 | 0.21055 | 0.02431 | 0.21055 | 0.00079 | 0.00232 | 0.00136 | 0.07832 | 0.03663 | 0.03663 | 0.28093 | 0.03663 | 0.28093 | 0.81275 | 0.96707 | 0.69937 |
| E20 Aml5/B5 | 0.21055 | 0.01008 | 0.36672 | SS | SS | SS | SS | SS | SS | 0.00002 | SS | SS | 0.11113 | 0.02431 | 0.46756 |
| E20 Aml5/H12.5 | 0.96707 | 0.15454 | 0.58062 | 0.01008 | 0.05410 | 0.05410 | 0.05410 | 0.28093 | 0.15454 | 0.36672 | 0.69937 | 0.36672 | 0.81275 | 0.58062 | 0.69937 |
| E20 B5/H12.5 | 0.46756 | 0.01581 | 0.15454 | SS | SS | SS | 0.00045 | 0.00004 | SS | 0.02431 | 0.00079 | 0.00079 | 0.58062 | 0.28093 | 0.58062 |
| L100 Aml5/B5 | 0.05410 | 0.00232 | 0.21055 | SS | SS | SS | SS | SS | SS | 0.00007 | SS | SS | 0.15454 | 0.21055 | 0.58062 |
| L100 Aml5/H12.5 | 0.96707 | 0.46756 | 0.81275 | 0.01581 | 0.00007 | 0.00004 | 0.05410 | 0.00013 | 0.00007 | 0.07832 | 0.00136 | 0.00079 | 0.28093 | 0.05410 | 0.90621 |
| L100 B5/H12.5 | 0.11113 | 0.01581 | 0.21055 | 0.00002 | 0.00004 | 0.00007 | 0.01008 | 0.00630 | 0.00232 | 0.15454 | 0.01008 | 0.05410 | 0.69937 | 0.90621 | 0.81275 |

**Al300 =** aliskiren 300 mg; **Aml5** = amlodipine 5 mg; **B5** = bisoprolol 5 mg; **E20** = enalapril 20 mg; **H12.5** = hydrochlorothiazide 12.5 mg; **L100** = losartan 100 mg; **SS** = statistically significant ($P$ < 0.00001)

**Figure S18.** Simulated change in left ventricular peak systolic pressure from baseline to week 4 (mean ± SD, $n$ = 100)

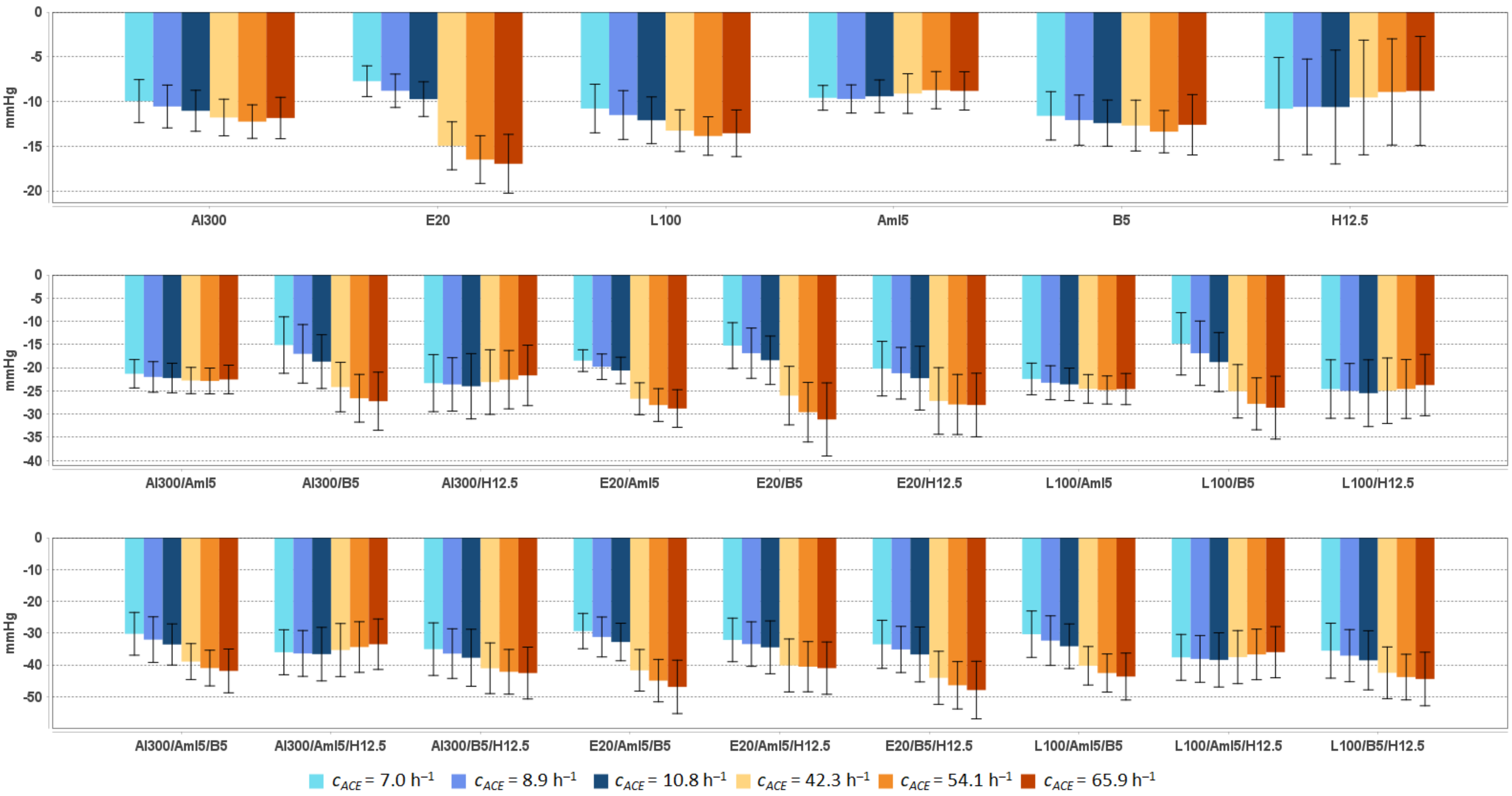


**Al300 =** aliskiren 300 mg; **Aml5** = amlodipine 5 mg; **B5** = bisoprolol 5 mg; **E20** = enalapril 20 mg; **H12.5** = hydrochlorothiazide 12.5 mg; **L100** = losartan 100 mg

**Table S22.** Simulated response of <u>left ventricular peak systolic pressure</u> to antihypertensive therapy in virtual hypertensive populations ($n$ = 100) with different ACE activity, including $P$-values (Kolmogorov-Smirnov test) for endpoint *vs*. baseline; data are presented as mean ± SD in mmHg

| Regimens | $c_{ACE}$ = 7.0 h$^{-1}$ | | | $c_{ACE}$ = 8.9 h$^{-1}$ | | | $c_{ACE}$ = 10.8 h$^{-1}$ | | | $c_{ACE}$ = 42.3 h$^{-1}$ | | | $c_{ACE}$ = 54.1 h$^{-1}$ | | | $c_{ACE}$ = 65.9 h$^{-1}$ | | |
|---|---|---|---|---|---|---|---|---|---|---|---|---|---|---|---|---|---|---|
| | **Value** | **Change** | ***P*** | **Value** | **Change** | ***P*** | **Value** | **Change** | ***P*** | **Value** | **Change** | ***P*** | **Value** | **Change** | ***P*** | **Value** | **Change** | ***P*** |
| Baseline | 174.0 ± 6.6 | – | – | 174.3 ± 6.9 | – | – | 173.8 ± 6.4 | – | – | 172.8 ± 6.7 | – | – | 172.6 ± 6.8 | – | – | 173.4 ± 7.0 | – | – |
| Al300 | 164.0 ± 6.6 | -10.0 ± 2.4 | SS | 163.7 ± 7.3 | -10.6 ± 2.4 | SS | 162.8 ± 6.9 | -11.0 ± 2.3 | SS | 161.0 ± 7.1 | -11.8 ± 2.0 | SS | 160.4 ± 7.1 | -12.3 ± 1.9 | SS | 161.5 ± 7.3 | -11.9 ± 2.3 | SS |
| E20 | 166.3 ± 6.5 | -7.7 ± 1.7 | SS | 165.5 ± 7.1 | -8.8 ± 1.9 | SS | 164.1 ± 6.8 | -9.7 ± 1.9 | SS | 157.9 ± 7.3 | -15.0 ± 2.7 | SS | 156.1 ± 7.3 | -16.5 ± 2.7 | SS | 156.4 ± 7.6 | -17.0 ± 3.3 | SS |
| L100 | 163.2 ± 6.7 | -10.8 ± 2.7 | SS | 162.8 ± 7.4 | -11.5 ± 2.7 | SS | 161.7 ± 7.0 | -12.1 ± 2.6 | SS | 159.6 ± 7.2 | -13.3 ± 2.3 | SS | 158.7 ± 7.1 | -13.9 ± 2.2 | SS | 159.8 ± 7.4 | -13.6 ± 2.6 | SS |
| Aml5 | 164.4 ± 6.5 | -9.6 ± 1.4 | SS | 164.6 ± 6.9 | -9.7 ± 1.6 | SS | 164.4 ± 6.5 | -9.4 ± 1.8 | SS | 163.7 ± 6.9 | -9.1 ± 2.2 | SS | 163.9 ± 6.4 | -8.7 ± 2.1 | SS | 164.6 ± 7.1 | -8.8 ± 2.1 | SS |
| B5 | 162.4 ± 6.5 | -11.6 ± 2.7 | SS | 162.2 ± 7.2 | -12.1 ± 2.8 | SS | 161.4 ± 6.7 | -12.4 ± 2.6 | SS | 160.1 ± 7.4 | -12.7 ± 2.8 | SS | 159.2 ± 7.0 | -13.4 ± 2.4 | SS | 160.8 ± 7.6 | -12.6 ± 3.4 | SS |
| H12.5 | 163.2 ± 8.2 | -10.8 ± 5.7 | SS | 163.7 ± 8.8 | -10.6 ± 5.3 | SS | 163.2 ± 8.8 | -10.6 ± 6.4 | SS | 163.3 ± 9.3 | -9.6 ± 6.4 | SS | 163.7 ± 8.3 | -8.9 ± 5.9 | SS | 164.6 ± 8.1 | -8.8 ± 6.1 | SS |
| Al300<br>Aml5 | 152.7 ± 6.8 | -21.3 ± 3.1 | SS | 152.3 ± 7.7 | -22.0 ± 3.3 | SS | 151.6 ± 7.4 | -22.2 ± 3.2 | SS | 150.1 ± 7.4 | -22.8 ± 2.8 | SS | 149.7 ± 7.0 | -22.9 ± 2.8 | SS | 150.8 ± 7.6 | -22.5 ± 3.1 | SS |
| Al300<br>B5 | 158.9 ± 8.9 | -15.1 ± 6.1 | SS | 157.3 ± 9.6 | -17.0 ± 6.3 | SS | 155.1 ± 8.8 | -18.7 ± 5.8 | SS | 148.7 ± 8.7 | -24.2 ± 5.3 | SS | 146.0 ± 8.4 | -26.6 ± 5.1 | SS | 146.2 ± 9.2 | -27.2 ± 6.3 | SS |
| Al300<br>H12.5 | 150.7 ± 8.2 | -23.3 ± 6.1 | SS | 150.7 ± 9.1 | -23.6 ± 5.8 | SS | 149.8 ± 9.5 | -24.0 ± 7.0 | SS | 149.7 ± 9.9 | -23.1 ± 7.0 | SS | 150.0 ± 8.7 | -22.6 ± 6.3 | SS | 151.7 ± 8.6 | -21.7 ± 6.5 | SS |
| E20<br>Aml5 | 155.5 ± 6.6 | -18.5 ± 2.3 | SS | 154.5 ± 7.4 | -19.8 ± 2.8 | SS | 153.2 ± 7.2 | -20.6 ± 2.9 | SS | 146.1 ± 7.6 | -26.7 ± 3.4 | SS | 144.6 ± 7.4 | -28.1 ± 3.5 | SS | 144.6 ± 8.0 | -28.8 ± 4.1 | SS |
| E20<br>B5 | 158.8 ± 7.9 | -15.2 ± 4.9 | SS | 157.4 ± 8.8 | -16.9 ± 5.4 | SS | 155.4 ± 8.4 | -18.4 ± 5.2 | SS | 146.8 ± 9.5 | -26.0 ± 6.3 | SS | 143.0 ± 9.3 | -29.6 ± 6.4 | SS | 142.2 ± 10.4 | -31.2 ± 7.9 | SS |
| E20<br>H12.5 | 153.8 ± 8.1 | -20.2 ± 5.9 | SS | 153.1 ± 9.0 | -21.2 ± 5.6 | SS | 151.5 ± 9.3 | -22.3 ± 6.9 | SS | 145.7 ± 10.0 | -27.2 ± 7.2 | SS | 144.7 ± 8.8 | -27.9 ± 6.5 | SS | 145.3 ± 8.8 | -28.0 ± 6.9 | SS |
| L100<br>Aml5 | 151.6 ± 7.0 | -22.4 ± 3.4 | SS | 151.1 ± 7.9 | -23.2 ± 3.7 | SS | 150.2 ± 7.6 | -23.6 ± 3.5 | SS | 148.3 ± 7.5 | -24.6 ± 3.1 | SS | 147.8 ± 7.1 | -24.8 ± 3.0 | SS | 148.8 ± 7.7 | -24.6 ± 3.4 | SS |
| L100<br>B5 | 159.1 ± 9.4 | -14.9 ± 6.7 | SS | 157.4 ± 10.1 | -16.9 ± 6.9 | SS | 155.0 ± 9.3 | -18.8 ± 6.4 | SS | 147.8 ± 9.0 | -25.1 ± 5.8 | SS | 144.8 ± 8.7 | -27.8 ± 5.6 | SS | 144.8 ± 9.6 | -28.6 ± 6.8 | SS |
| L100<br>H12.5 | 149.4 ± 8.3 | -24.6 ± 6.3 | SS | 149.3 ± 9.2 | -25.0 ± 5.9 | SS | 148.3 ± 9.6 | -25.5 ± 7.2 | SS | 147.9 ± 9.9 | -25.0 ± 7.1 | SS | 148.0 ± 8.7 | -24.6 ± 6.3 | SS | 149.6 ± 8.6 | -23.7 ± 6.6 | SS |
| Al300<br>Aml5/B5 | 143.8 ± 9.3 | -30.2 ± 6.7 | SS | 142.2 ± 10.4 | -32.1 ± 7.2 | SS | 140.2 ± 9.5 | -33.6 ± 6.5 | SS | 133.9 ± 8.8 | -39.0 ± 5.7 | SS | 131.6 ± 8.5 | -41.0 ± 5.6 | SS | 131.5 ± 9.6 | -41.9 ± 6.9 | SS |
| Al300<br>Aml5/H12.5 | 138.0 ± 8.8 | -36.0 ± 7.1 | SS | 137.9 ± 10.2 | -36.4 ± 7.2 | SS | 137.2 ± 10.7 | -36.6 ± 8.4 | SS | 137.5 ± 10.8 | -35.4 ± 8.3 | SS | 138.2 ± 9.7 | -34.4 ± 8.0 | SS | 139.9 ± 9.7 | -33.5 ± 7.9 | SS |
| Al300<br>B5/H12.5 | 138.9 ± 9.6 | -35.1 ± 8.3 | SS | 137.8 ± 10.6 | -36.5 ± 7.8 | SS | 136.0 ± 11.0 | -37.8 ± 9.0 | SS | 131.7 ± 10.2 | -41.1 ± 8.0 | SS | 130.4 ± 8.8 | -42.2 ± 7.0 | SS | 130.8 ± 9.6 | -42.6 ± 8.2 | SS |
| E20<br>Aml5/B5 | 144.6 ± 8.3 | -29.4 ± 5.6 | SS | 143.1 ± 9.6 | -31.2 ± 6.3 | SS | 141.0 ± 9.1 | -32.8 ± 5.9 | SS | 131.1 ± 9.4 | -41.7 ± 6.5 | SS | 127.6 ± 9.2 | -45.0 ± 6.7 | SS | 126.4 ± 10.7 | -46.9 ± 8.4 | SS |
| E20<br>Aml5/H12.5 | 141.8 ± 8.7 | -32.2 ± 6.8 | SS | 140.8 ± 10.1 | -33.5 ± 7.0 | SS | 139.3 ± 10.6 | -34.5 ± 8.3 | SS | 132.6 ± 10.6 | -40.2 ± 8.3 | SS | 132.0 ± 9.6 | -40.6 ± 7.9 | SS | 132.3 ± 9.8 | -41.0 ± 8.2 | SS |
| E20<br>B5/H12.5 | 140.4 ± 9.0 | -33.6 ± 7.6 | SS | 139.1 ± 10.1 | -35.2 ± 7.3 | SS | 137.1 ± 10.6 | -36.7 ± 8.6 | SS | 128.7 ± 10.5 | -44.1 ± 8.4 | SS | 126.2 ± 9.1 | -46.4 ± 7.5 | SS | 125.4 ± 10.3 | -47.9 ± 9.1 | SS |
| L100<br>Aml5/B5 | 143.6 ± 9.8 | -30.4 ± 7.3 | SS | 141.9 ± 10.9 | -32.4 ± 7.8 | SS | 139.6 ± 10.0 | -34.2 ± 7.0 | SS | 132.5 ± 9.1 | -40.3 ± 6.1 | SS | 130.0 ± 8.7 | -42.6 ± 6.0 | SS | 129.7 ± 9.9 | -43.7 ± 7.4 | SS |
| L100<br>Aml5/H12.5 | 136.3 ± 8.8 | -37.7 ± 7.2 | SS | 136.1 ± 10.3 | -38.1 ± 7.4 | SS | 135.3 ± 10.8 | -38.5 ± 8.5 | SS | 135.3 ± 10.7 | -37.6 ± 8.3 | SS | 135.9 ± 9.7 | -36.7 ± 8.0 | SS | 137.4 ± 9.7 | -36.0 ± 8.0 | SS |
| L100<br>B5/H12.5 | 138.5 ± 10.0 | -35.6 ± 8.6 | SS | 137.2 ± 10.9 | -37.1 ± 8.2 | SS | 135.2 ± 11.3 | -38.6 ± 9.3 | SS | 130.3 ± 10.3 | -42.5 ± 8.1 | SS | 128.7 ± 8.9 | -43.9 ± 7.2 | SS | 128.9 ± 9.8 | -44.5 ± 8.4 | SS |

**Al300 =** aliskiren 300 mg; **Aml5** = amlodipine 5 mg; **B5** = bisoprolol 5 mg; **E20** = enalapril 20 mg; **H12.5** = hydrochlorothiazide 12.5 mg; **L100** = losartan 100 mg; **SS** = statistically significant ($P$ < 0.00001)

**Table S23.** *P*-values calculated using the Kolmogorov-Smirnov test for changes in left ventricular peak systolic pressure in populations ($n$ = 100) with different ACE activity receiving the same regimens (case 1: $c_{ACE}$ = 7.0 h$^{-1}$, case 2: $c_{ACE}$ = 8.9 h$^{-1}$, case 3: $c_{ACE}$ = 10.8 h$^{-1}$, case 4: $c_{ACE}$ = 42.3 h$^{-1}$, case 5: $c_{ACE}$ = 54.1 h$^{-1}$, case 6: $c_{ACE}$ = 65.9 h$^{-1}$; *P*-value for case *i vs*. case *j* is denoted $P_{ij}$)

| Regimens | $P_{12}$ | $P_{13}$ | $P_{23}$ | $P_{14}$ | $P_{15}$ | $P_{16}$ | $P_{24}$ | $P_{25}$ | $P_{26}$ | $P_{34}$ | $P_{35}$ | $P_{36}$ | $P_{45}$ | $P_{46}$ | $P_{56}$ |
|---|---|---|---|---|---|---|---|---|---|---|---|---|---|---|---|
| Al300 | 0.15454 | 0.01581 | 0.36672 | SS | SS | SS | 0.00079 | SS | 0.00045 | 0.05410 | 0.00079 | 0.02431 | 0.21055 | 0.69937 | 0.28093 |
| E20 | 0.00079 | SS | 0.01581 | SS | SS | SS | SS | SS | SS | SS | SS | SS | 0.00079 | 0.00004 | 0.03663 |
| L100 | 0.07832 | 0.00630 | 0.36672 | SS | SS | SS | 0.00007 | SS | SS | 0.01008 | 0.00002 | 0.00136 | 0.11113 | 0.46756 | 0.36672 |
| Aml5 | 0.58062 | 0.69937 | 0.69937 | 0.15454 | 0.00045 | 0.01008 | 0.07832 | 0.00002 | 0.00045 | 0.36672 | 0.00045 | 0.01008 | 0.11113 | 0.21055 | 0.81275 |
| B5 | 0.28093 | 0.11113 | 0.11113 | 0.03663 | 0.00045 | 0.00386 | 0.03663 | 0.02431 | 0.01581 | 0.69937 | 0.05410 | 0.46756 | 0.21055 | 0.46756 | 0.02431 |
| H12.5 | 0.69937 | 0.69937 | 0.46756 | 0.46756 | 0.15454 | 0.07832 | 0.21055 | 0.01581 | 0.00232 | 0.28093 | 0.05410 | 0.02431 | 0.28093 | 0.36672 | 0.96707 |
| Al300 Aml5 | 0.03663 | 0.00630 | 0.69937 | 0.00045 | 0.00079 | 0.00232 | 0.07832 | 0.36672 | 0.11113 | 0.36672 | 0.36672 | 0.69937 | 0.81275 | 0.90621 | 0.96707 |
| Al300 B5 | 0.05410 | 0.00013 | 0.15454 | SS | SS | SS | SS | SS | SS | SS | SS | SS | 0.02431 | 0.00630 | 0.46756 |
| Al300 H12.5 | 0.81275 | 0.15454 | 0.46756 | 0.81275 | 0.69937 | 0.11113 | 0.46756 | 0.15454 | 0.02431 | 0.58062 | 0.03663 | 0.01008 | 0.46756 | 0.28093 | 0.58062 |
| E20 Aml5 | 0.00007 | SS | 0.02431 | SS | SS | SS | SS | SS | SS | SS | SS | SS | 0.07832 | 0.00079 | 0.15454 |
| E20 B5 | 0.07832 | SS | 0.07832 | SS | SS | SS | SS | SS | SS | SS | SS | SS | 0.00386 | 0.00007 | 0.15454 |
| E20 H12.5 | 0.36672 | 0.00386 | 0.07832 | SS | SS | SS | SS | SS | SS | 0.00002 | SS | SS | 0.81275 | 0.46756 | 0.90621 |
| L100 Aml5 | 0.03663 | 0.00136 | 0.36672 | SS | SS | 0.00002 | 0.01581 | 0.00386 | 0.00386 | 0.07832 | 0.03663 | 0.11113 | 0.90621 | 0.81275 | 0.81275 |
| L100 B5 | 0.05410 | 0.00007 | 0.11113 | SS | SS | SS | SS | SS | SS | SS | SS | SS | 0.02431 | 0.00232 | 0.36672 |
| L100 H12.5 | 0.58062 | 0.11113 | 0.15454 | 0.28093 | 0.96707 | 0.81275 | 0.36672 | 0.28093 | 0.15454 | 0.81275 | 0.07832 | 0.07832 | 0.36672 | 0.46756 | 0.81275 |
| Al300 Aml5/B5 | 0.05410 | 0.00136 | 0.28093 | SS | SS | SS | SS | SS | SS | SS | SS | SS | 0.07832 | 0.01581 | 0.28093 |
| Al300 Aml5/H12.5 | 0.58062 | 0.58062 | 0.46756 | 0.46756 | 0.15454 | 0.11113 | 0.36672 | 0.03663 | 0.03663 | 0.21055 | 0.07832 | 0.03663 | 0.69937 | 0.15454 | 0.36672 |
| Al300 B5/H12.5 | 0.21055 | 0.03663 | 0.28093 | SS | SS | SS | 0.00386 | 0.00007 | 0.00002 | 0.01581 | 0.00232 | 0.00630 | 0.58062 | 0.21055 | 0.81275 |
| E20 Aml5/B5 | 0.00630 | 0.00007 | 0.15454 | SS | SS | SS | SS | SS | SS | SS | SS | SS | 0.00386 | 0.00025 | 0.07832 |
| E20 Aml5/H12.5 | 0.21055 | 0.01008 | 0.21055 | SS | SS | SS | SS | SS | SS | 0.00045 | 0.00002 | SS | 0.46756 | 0.28093 | 0.36672 |
| E20 B5/H12.5 | 0.15454 | 0.01008 | 0.21055 | SS | SS | SS | SS | SS | SS | SS | SS | SS | 0.15454 | 0.00386 | 0.21055 |
| L100 Aml5/B5 | 0.05410 | 0.00136 | 0.28093 | SS | SS | SS | SS | SS | SS | SS | SS | SS | 0.03663 | 0.00630 | 0.28093 |
| L100 Aml5/H12.5 | 0.46756 | 0.46756 | 0.36672 | 0.58062 | 0.28093 | 0.36672 | 0.36672 | 0.05410 | 0.11113 | 0.36672 | 0.15454 | 0.05410 | 0.69937 | 0.21055 | 0.46756 |
| L100 B5/H12.5 | 0.21055 | 0.01581 | 0.15454 | SS | SS | SS | 0.00025 | SS | SS | 0.01008 | 0.00007 | 0.00136 | 0.46756 | 0.07832 | 0.69937 |

**Al300 =** aliskiren 300 mg; **Aml5** = amlodipine 5 mg; **B5** = bisoprolol 5 mg; **E20** = enalapril 20 mg; **H12.5** = hydrochlorothiazide 12.5 mg; **L100** = losartan 100 mg; **SS** = statistically significant ($P$ < 0.00001)

**Figure S19.** Simulated change in <u>left ventricular end-diastolic volume</u> from baseline to week 4 (mean ± SD, $n$ = 100)

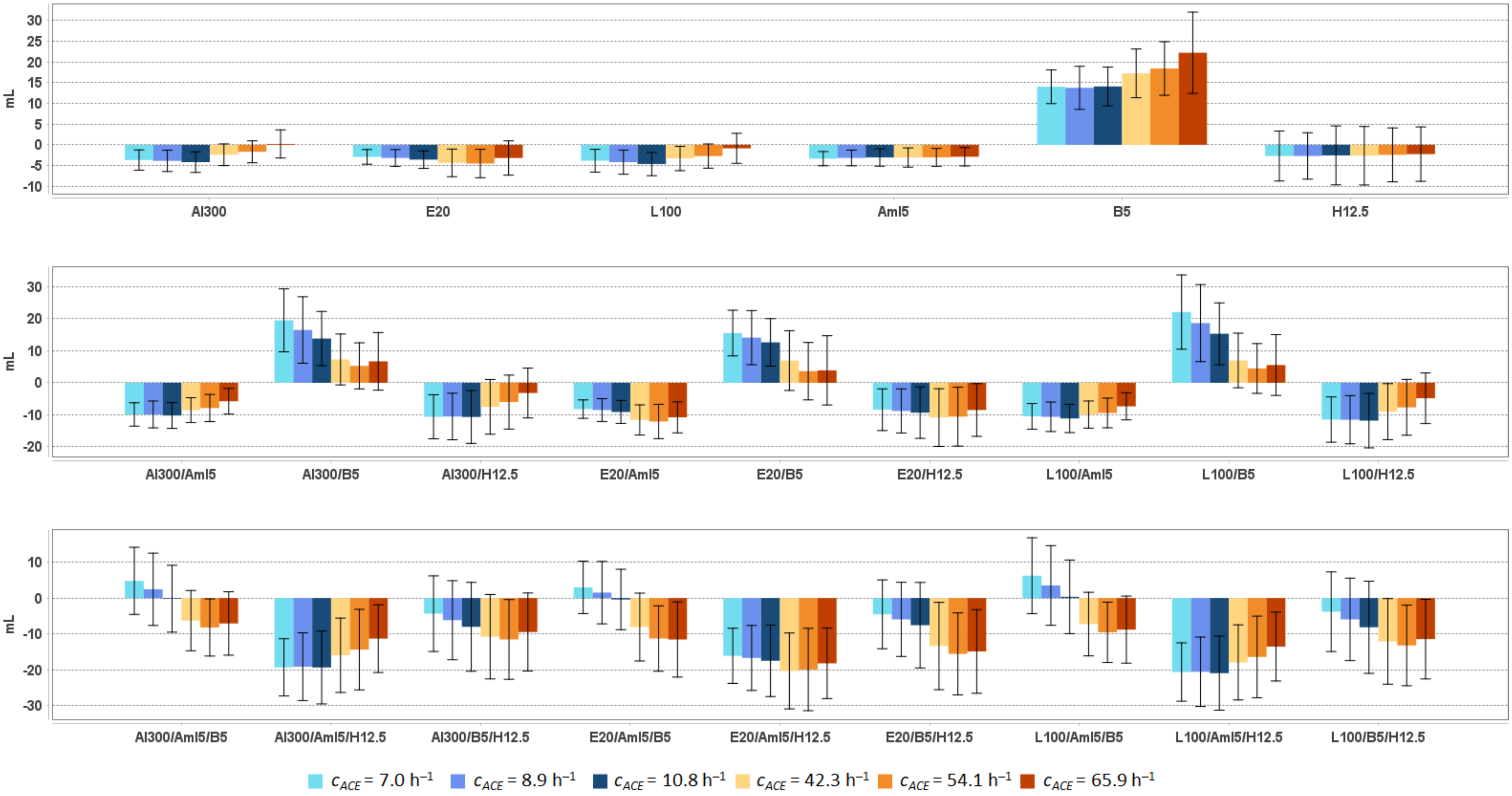


**Al300 =** aliskiren 300 mg; **Aml5** = amlodipine 5 mg; **B5** = bisoprolol 5 mg; **E20** = enalapril 20 mg; **H12.5** = hydrochlorothiazide 12.5 mg; **L100** = losartan 100 mg

**Table S24.** Simulated response of <u>left ventricular end-diastolic volume</u> to antihypertensive therapy in virtual hypertensive populations ($n$ = 100) with different ACE activity, including $P$-values (Kolmogorov-Smirnov test) for endpoint *vs*. baseline; data are presented as mean ± SD in mL

| Regimens | $c_{ACE}$ = 7.0 h$^{-1}$ | | | $c_{ACE}$ = 8.9 h$^{-1}$ | | | $c_{ACE}$ = 10.8 h$^{-1}$ | | | $c_{ACE}$ = 42.3 h$^{-1}$ | | | $c_{ACE}$ = 54.1 h$^{-1}$ | | | $c_{ACE}$ = 65.9 h$^{-1}$ | | |
|---|---|---|---|---|---|---|---|---|---|---|---|---|---|---|---|---|---|---|
| | **Value** | **Change** | ***P*** | **Value** | **Change** | ***P*** | **Value** | **Change** | ***P*** | **Value** | **Change** | ***P*** | **Value** | **Change** | ***P*** | **Value** | **Change** | ***P*** |
| Baseline | 110.6 ± 12.8 | – | – | 110.0 ± 14.2 | – | – | 111.0 ± 14.0 | – | – | 110.5 ± 13.3 | – | – | 111.2 ± 13.5 | – | – | 110.5 ± 13.4 | – | – |
| Al300 | 106.9 ± 12.6 | -3.7 ± 2.4 | 0.15454 | 106.2 ± 14.1 | -3.9 ± 2.6 | 0.15454 | 106.8 ± 13.8 | -4.2 ± 2.5 | 0.01008 | 108.1 ± 13.1 | -2.4 ± 2.6 | 0.15454 | 109.5 ± 13.7 | -1.7 ± 2.6 | 0.58062 | 110.7 ± 13.6 | 0.2 ± 3.4 | 0.96707 |
| E20 | 107.7 ± 12.6 | -2.9 ± 1.8 | 0.28093 | 106.9 ± 14.1 | -3.2 ± 2.0 | 0.36672 | 107.4 ± 13.8 | -3.6 ± 2.1 | 0.03663 | 106.1 ± 13.0 | -4.3 ± 3.3 | 0.01008 | 106.7 ± 13.5 | -4.5 ± 3.4 | 0.01581 | 107.3 ± 13.6 | -3.2 ± 4.1 | 0.11113 |
| L100 | 106.7 ± 12.6 | -3.8 ± 2.8 | 0.11113 | 105.8 ± 14.1 | -4.2 ± 2.9 | 0.11113 | 106.3 ± 13.8 | -4.6 ± 2.8 | 0.00630 | 107.2 ± 13.1 | -3.3 ± 2.9 | 0.03663 | 108.5 ± 13.6 | -2.7 ± 2.9 | 0.21055 | 109.6 ± 13.6 | -0.8 ± 3.6 | 0.90621 |
| Aml5 | 107.2 ± 12.6 | -3.3 ± 1.7 | 0.03663 | 106.9 ± 13.9 | -3.2 ± 1.9 | 0.36672 | 107.9 ± 13.8 | -3.0 ± 2.1 | 0.05410 | 107.4 ± 13.2 | -3.1 ± 2.3 | 0.11113 | 108.1 ± 13.5 | -3.0 ± 2.2 | 0.02431 | 107.6 ± 13.6 | -2.9 ± 2.2 | 0.15454 |
| B5 | 124.6 ± 14.3 | 14.0 ± 4.1 | SS | 123.8 ± 16.9 | 13.8 ± 5.2 | SS | 125.1 ± 16.3 | 14.1 ± 4.7 | SS | 127.8 ± 15.8 | 17.3 ± 5.9 | SS | 129.6 ± 17.2 | 18.4 ± 6.5 | SS | 132.7 ± 18.2 | 22.2 ± 9.8 | SS |
| H12.5 | 107.9 ± 14.3 | -2.7 ± 6.0 | 0.15454 | 107.3 ± 14.5 | -2.7 ± 5.6 | 0.58062 | 108.4 ± 15.9 | -2.5 ± 7.1 | 0.07832 | 107.9 ± 14.3 | -2.6 ± 7.1 | 0.28093 | 108.7 ± 15.6 | -2.4 ± 6.5 | 0.21055 | 108.2 ± 14.3 | -2.2 ± 6.6 | 0.46756 |
| Al300 Aml5 | 100.7 ± 12.4 | -9.9 ± 3.6 | 0.00004 | 100.1 ± 13.9 | -9.9 ± 4.2 | 0.00025 | 100.8 ± 13.5 | -10.2 ± 4.0 | SS | 102.0 ± 13.0 | -8.5 ± 3.9 | 0.00007 | 103.3 ± 13.7 | -7.9 ± 4.2 | 0.00045 | 104.7 ± 13.9 | -5.7 ± 4.0 | 0.00630 |
| Al300 B5 | 130.1 ± 16.6 | 19.5 ± 9.8 | SS | 126.5 ± 19.6 | 16.5 ± 10.4 | SS | 124.8 ± 17.1 | 13.8 ± 8.5 | SS | 117.8 ± 15.1 | 7.3 ± 7.9 | 0.00004 | 116.5 ± 15.4 | 5.3 ± 7.2 | 0.00630 | 117.2 ± 16.2 | 6.7 ± 9.0 | 0.00045 |
| Al300 H12.5 | 100.0 ± 14.1 | -10.6 ± 6.9 | 0.00002 | 99.5 ± 14.6 | -10.5 ± 7.3 | 0.00013 | 100.3 ± 15.9 | -10.7 ± 8.2 | SS | 103.0 ± 14.7 | -7.5 ± 8.5 | 0.00232 | 105.1 ± 16.6 | -6.0 ± 8.4 | 0.01008 | 107.3 ± 14.7 | -3.2 ± 7.8 | 0.28093 |
| E20 Aml5 | 102.4 ± 12.4 | -8.2 ± 2.9 | 0.00025 | 101.5 ± 13.8 | -8.5 ± 3.5 | 0.00232 | 101.9 ± 13.5 | -9.1 ± 3.6 | SS | 98.9 ± 13.0 | -11.6 ± 4.7 | SS | 99.1 ± 13.6 | -12.1 ± 5.4 | SS | 99.7 ± 14.0 | -10.8 ± 4.8 | SS |
| E20 B5 | 126.1 ± 15.2 | 15.5 ± 7.1 | SS | 124.1 ± 18.3 | 14.1 ± 8.4 | SS | 123.6 ± 16.6 | 12.6 ± 7.4 | SS | 117.4 ± 15.6 | 6.9 ± 9.3 | 0.00007 | 114.8 ± 15.9 | 3.6 ± 9.0 | 0.03663 | 114.4 ± 17.2 | 3.9 ± 10.8 | 0.03663 |
| E20 H12.5 | 102.2 ± 14.1 | -8.4 ± 6.5 | 0.00079 | 101.2 ± 14.5 | -8.8 ± 6.9 | 0.00136 | 101.6 ± 15.9 | -9.3 ± 8.0 | 0.00013 | 99.6 ± 14.7 | -10.8 ± 9.0 | 0.00004 | 100.6 ± 16.6 | -10.6 ± 9.2 | 0.00013 | 102.0 ± 14.8 | -8.5 ± 8.2 | 0.00045 |
| L100 Aml5 | 100.1 ± 12.4 | -10.5 ± 4.0 | 0.00002 | 99.4 ± 14.0 | -10.6 ± 4.6 | 0.00013 | 99.8 ± 13.6 | -11.1 ± 4.4 | SS | 100.6 ± 13.0 | -9.9 ± 4.2 | SS | 101.8 ± 13.7 | -9.4 ± 4.6 | 0.00004 | 103.1 ± 13.9 | -7.3 ± 4.2 | 0.00079 |
| L100 B5 | 132.7 ± 17.8 | 22.1 ± 11.6 | SS | 128.7 ± 20.8 | 18.6 ± 12.0 | SS | 126.3 ± 17.6 | 15.3 ± 9.6 | SS | 117.5 ± 15.3 | 7.0 ± 8.5 | 0.00004 | 115.6 ± 15.5 | 4.5 ± 7.8 | 0.01581 | 116.0 ± 16.4 | 5.6 ± 9.5 | 0.00630 |
| L100 H12.5 | 99.1 ± 14.1 | -11.5 ± 7.1 | SS | 98.5 ± 14.6 | -11.5 ± 7.5 | 0.00002 | 99.1 ± 16.0 | -11.8 ± 8.5 | SS | 101.5 ± 14.7 | -9.0 ± 8.7 | 0.00025 | 103.5 ± 16.6 | -7.7 ± 8.7 | 0.00386 | 105.7 ± 14.7 | -4.8 ± 7.9 | 0.11113 |
| Al300 Aml5/B5 | 115.4 ± 15.2 | 4.8 ± 9.3 | 0.00045 | 112.5 ± 17.9 | 2.5 ± 10.1 | 0.36672 | 110.8 ± 16.1 | -0.2 ± 9.3 | 0.28093 | 104.2 ± 14.7 | -6.3 ± 8.4 | 0.01581 | 103.0 ± 14.8 | -8.2 ± 8.0 | 0.00045 | 103.4 ± 15.9 | -7.1 ± 8.8 | 0.00232 |
| Al300 Aml5/H12.5 | 91.3 ± 14.5 | -19.3 ± 8.0 | SS | 90.9 ± 15.4 | -19.1 ± 9.4 | SS | 91.6 ± 16.7 | -19.3 ± 10.1 | SS | 94.6 ± 15.9 | -15.9 ± 10.4 | SS | 96.8 ± 17.8 | -14.3 ± 11.2 | SS | 99.2 ± 15.9 | -11.3 ± 9.4 | SS |
| Al300 B5/H12.5 | 106.2 ± 16.3 | -4.3 ± 10.5 | 0.05410 | 103.9 ± 17.2 | -6.1 ± 11.0 | 0.02431 | 103.0 ± 18.5 | -8.0 ± 12.4 | 0.00079 | 99.7 ± 16.4 | -10.8 ± 11.7 | 0.00004 | 99.6 ± 17.3 | -11.5 ± 11.1 | SS | 101.0 ± 16.3 | -9.4 ± 10.9 | 0.00079 |
| E20 Aml5/B5 | 113.6 ± 14.3 | 3.0 ± 7.3 | 0.00079 | 111.5 ± 17.1 | 1.5 ± 8.7 | 0.36672 | 110.6 ± 15.8 | -0.4 ± 8.4 | 0.36672 | 102.4 ± 15.0 | -8.1 ± 9.4 | 0.00232 | 99.9 ± 15.1 | -11.3 ± 9.1 | SS | 99.0 ± 16.7 | -11.5 ± 10.5 | SS |
| E20 Aml5/H12.5 | 94.5 ± 14.5 | -16.1 ± 7.7 | SS | 93.4 ± 15.3 | -16.7 ± 9.1 | SS | 93.5 ± 16.6 | -17.5 ± 10.0 | SS | 90.2 ± 15.9 | -20.3 ± 10.6 | SS | 91.3 ± 17.5 | -19.9 ± 11.5 | SS | 92.3 ± 16.0 | -18.2 ± 9.8 | SS |
| E20 B5/H12.5 | 106.1 ± 15.8 | -4.5 ± 9.6 | 0.07832 | 104.1 ± 16.8 | -5.9 ± 10.3 | 0.05410 | 103.4 ± 18.2 | -7.5 ± 11.9 | 0.00079 | 97.2 ± 16.5 | -13.3 ± 12.2 | SS | 95.6 ± 17.1 | -15.5 ± 11.4 | SS | 95.6 ± 16.7 | -14.8 ± 11.6 | SS |
| L100 Aml5/B5 | 116.8 ± 15.9 | 6.2 ± 10.6 | 0.00045 | 113.5 ± 18.6 | 3.5 ± 11.1 | 0.28093 | 111.3 ± 16.6 | 0.3 ± 10.2 | 0.21055 | 103.3 ± 14.8 | -7.2 ± 8.8 | 0.00630 | 101.6 ± 14.9 | -9.5 ± 8.4 | 0.00004 | 101.7 ± 16.2 | -8.8 ± 9.3 | 0.00013 |
| L100 Aml5/H12.5 | 90.0 ± 14.6 | -20.6 ± 8.1 | SS | 89.5 ± 15.5 | -20.5 ± 9.7 | SS | 90.1 ± 16.7 | -20.9 ± 10.3 | SS | 92.6 ± 15.9 | -17.9 ± 10.5 | SS | 94.8 ± 17.7 | -16.4 ± 11.4 | SS | 97.0 ± 16.0 | -13.5 ± 9.6 | SS |
| L100 B5/H12.5 | 106.8 ± 16.7 | -3.8 ± 11.1 | 0.11113 | 104.1 ± 17.6 | -5.9 ± 11.5 | 0.01581 | 102.8 ± 18.7 | -8.1 ± 12.8 | 0.00025 | 98.4 ± 16.4 | -12.1 ± 11.9 | SS | 98.0 ± 17.2 | -13.2 ± 11.2 | SS | 99.1 ± 16.4 | -11.4 ± 11.1 | 0.00007 |

**Al300 =** aliskiren 300 mg; **Aml5** = amlodipine 5 mg; **B5** = bisoprolol 5 mg; **E20** = enalapril 20 mg; **H12.5** = hydrochlorothiazide 12.5 mg; **L100** = losartan 100 mg; **SS** = statistically significant ($P$ < 0.00001)

**Table S25.** *P*-values calculated using the Kolmogorov-Smirnov test for changes in left ventricular end-diastolic volume in populations ($n$ = 100) with different ACE activity receiving the same regimens (case 1: $c_{ACE}$ = 7.0 h$^{-1}$, case 2: $c_{ACE}$ = 8.9 h$^{-1}$, case 3: $c_{ACE}$ = 10.8 h$^{-1}$, case 4: $c_{ACE}$ = 42.3 h$^{-1}$, case 5: $c_{ACE}$ = 54.1 h$^{-1}$, case 6: $c_{ACE}$ = 65.9 h$^{-1}$; *P*-value for case *i vs*. case *j* is denoted $P_{ij}$)

| Regimens | $P_{12}$ | $P_{13}$ | $P_{23}$ | $P_{14}$ | $P_{15}$ | $P_{16}$ | $P_{24}$ | $P_{25}$ | $P_{26}$ | $P_{34}$ | $P_{35}$ | $P_{36}$ | $P_{45}$ | $P_{46}$ | $P_{56}$ |
|---|---|---|---|---|---|---|---|---|---|---|---|---|---|---|---|
| Al300 | 0.28093 | 0.03663 | 0.15454 | 0.01581 | 0.00004 | SS | 0.00386 | 0.00002 | SS | 0.00013 | SS | SS | 0.21055 | 0.00002 | 0.00136 |
| E20 | 0.36672 | 0.00386 | 0.11113 | 0.00007 | 0.00013 | 0.00386 | 0.00045 | 0.00025 | 0.01008 | 0.00630 | 0.00232 | 0.02431 | 0.96707 | 0.15454 | 0.11113 |
| L100 | 0.28093 | 0.01581 | 0.21055 | 0.28093 | 0.03663 | SS | 0.15454 | 0.01581 | SS | 0.01581 | 0.00013 | SS | 0.46756 | 0.00025 | 0.00630 |
| Aml5 | 0.46756 | 0.15454 | 0.96707 | 0.58062 | 0.07832 | 0.11113 | 0.69937 | 0.46756 | 0.28093 | 0.96707 | 0.90621 | 0.90621 | 0.69937 | 0.90621 | 0.99963 |
| B5 | 0.15454 | 0.81275 | 0.11113 | 0.00232 | SS | SS | 0.00007 | SS | SS | 0.00136 | SS | SS | 0.11113 | 0.00079 | 0.01008 |
| H12.5 | 0.69937 | 0.58062 | 0.28093 | 0.69937 | 0.81275 | 0.81275 | 0.28093 | 0.15454 | 0.21055 | 0.99376 | 0.46756 | 0.36672 | 0.69937 | 0.90621 | 0.99963 |
| Al300 Aml5 | 0.99376 | 0.58062 | 0.58062 | 0.15454 | 0.00079 | SS | 0.07832 | 0.00045 | SS | 0.07832 | 0.00025 | SS | 0.28093 | 0.00025 | 0.00630 |
| Al300 B5 | 0.11113 | 0.00232 | 0.28093 | SS | SS | SS | SS | SS | SS | 0.00004 | SS | 0.00004 | 0.28093 | 0.58062 | 0.58062 |
| Al300 H12.5 | 0.99376 | 0.46756 | 0.36672 | 0.01008 | 0.00002 | SS | 0.01008 | 0.00004 | SS | 0.05410 | 0.00007 | SS | 0.11113 | 0.00232 | 0.28093 |
| E20 Aml5 | 0.46756 | 0.11113 | 0.21055 | SS | SS | SS | SS | SS | SS | 0.00136 | 0.00079 | 0.01008 | 0.90621 | 0.69937 | 0.36672 |
| E20 B5 | 0.15454 | 0.01008 | 0.58062 | SS | SS | SS | SS | SS | SS | 0.00013 | SS | SS | 0.03663 | 0.07832 | 0.46756 |
| E20 H12.5 | 0.90621 | 0.21055 | 0.36672 | 0.01581 | 0.11113 | 0.46756 | 0.02431 | 0.15454 | 0.81275 | 0.46756 | 0.69937 | 0.36672 | 0.58062 | 0.28093 | 0.69937 |
| L100 Aml5 | 0.96707 | 0.36672 | 0.36672 | 0.69937 | 0.05410 | 0.00079 | 0.58062 | 0.00630 | 0.00004 | 0.28093 | 0.00232 | SS | 0.21055 | 0.00045 | 0.01581 |
| L100 B5 | 0.11113 | 0.00232 | 0.28093 | SS | SS | SS | SS | SS | SS | SS | SS | SS | 0.11113 | 0.28093 | 0.58062 |
| L100 H12.5 | 0.90621 | 0.58062 | 0.36672 | 0.03663 | 0.00025 | SS | 0.02431 | 0.00045 | SS | 0.21055 | 0.00136 | SS | 0.21055 | 0.00232 | 0.28093 |
| Al300 Aml5/B5 | 0.07832 | 0.01008 | 0.28093 | SS | SS | SS | SS | SS | SS | 0.00079 | SS | SS | 0.15454 | 0.58062 | 0.81275 |
| Al300 Aml5/H12.5 | 0.69937 | 0.46756 | 0.90621 | 0.00136 | 0.00004 | SS | 0.02431 | 0.00079 | SS | 0.07832 | 0.00079 | SS | 0.28093 | 0.00630 | 0.28093 |
| Al300 B5/H12.5 | 0.58062 | 0.01581 | 0.21055 | 0.00136 | 0.00079 | 0.00136 | 0.03663 | 0.02431 | 0.05410 | 0.28093 | 0.03663 | 0.15454 | 0.90621 | 0.81275 | 0.69937 |
| E20 Aml5/B5 | 0.21055 | 0.02431 | 0.28093 | SS | SS | SS | SS | SS | SS | 0.00004 | SS | SS | 0.07832 | 0.07832 | 0.58062 |
| E20 Aml5/H12.5 | 0.69937 | 0.11113 | 0.69937 | 0.00232 | 0.01581 | 0.07832 | 0.00232 | 0.05410 | 0.46756 | 0.11113 | 0.58062 | 0.81275 | 0.69937 | 0.15454 | 0.58062 |
| E20 B5/H12.5 | 0.36672 | 0.02431 | 0.11113 | SS | SS | SS | 0.00004 | 0.00002 | SS | 0.02431 | 0.00045 | 0.00136 | 0.58062 | 0.46756 | 0.81275 |
| L100 Aml5/B5 | 0.11113 | 0.00630 | 0.15454 | SS | SS | SS | SS | SS | SS | 0.00004 | SS | SS | 0.15454 | 0.21055 | 0.90621 |
| L100 Aml5/H12.5 | 0.81275 | 0.46756 | 0.81275 | 0.00630 | 0.00013 | SS | 0.07832 | 0.00232 | 0.00004 | 0.05410 | 0.00232 | 0.00004 | 0.36672 | 0.01581 | 0.36672 |
| L100 B5/H12.5 | 0.46756 | 0.02431 | 0.15454 | 0.00004 | 0.00002 | 0.00004 | 0.00386 | 0.00386 | 0.00232 | 0.07832 | 0.01008 | 0.03663 | 0.81275 | 0.96707 | 0.81275 |

**Al300 =** aliskiren 300 mg; **Aml5** = amlodipine 5 mg; **B5** = bisoprolol 5 mg; **E20** = enalapril 20 mg; **H12.5** = hydrochlorothiazide 12.5 mg; **L100** = losartan 100 mg; **SS** = statistically significant ($P < 0.00001$)

**Figure S20.** Simulated change in <u>left ventricular end-systolic volume</u> from baseline to week 4 (mean ± SD, $n$ = 100)

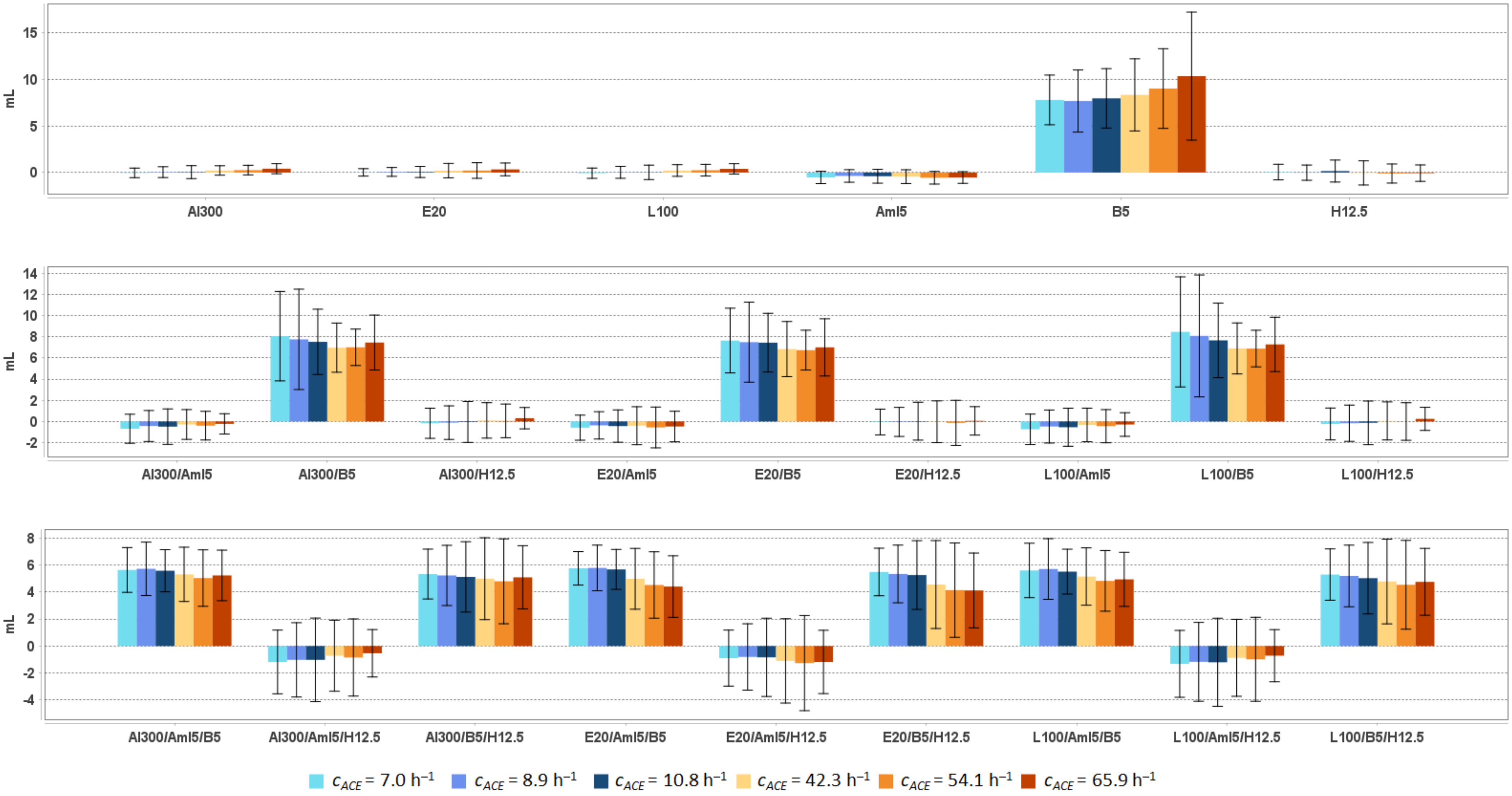


**Al300 =** aliskiren 300 mg; **Aml5** = amlodipine 5 mg; **B5** = bisoprolol 5 mg; **E20** = enalapril 20 mg; **H12.5** = hydrochlorothiazide 12.5 mg; **L100** = losartan 100 mg

**Table S26.** Simulated response of left ventricular end-systolic volume to antihypertensive therapy in virtual hypertensive populations ($n$ = 100) with different ACE activity, including $P$-values (Kolmogorov-Smirnov test) for endpoint *vs.* baseline; data are presented as mean ± SD in mL

| Regimens | $c_{ACE}$ = 7.0 h$^{-1}$ | | | $c_{ACE}$ = 8.9 h$^{-1}$ | | | $c_{ACE}$ = 10.8 h$^{-1}$ | | | $c_{ACE}$ = 42.3 h$^{-1}$ | | | $c_{ACE}$ = 54.1 h$^{-1}$ | | | $c_{ACE}$ = 65.9 h$^{-1}$ | | |
|---|---|---|---|---|---|---|---|---|---|---|---|---|---|---|---|---|---|---|
| | **Value** | **Change** | ***P*** | **Value** | **Change** | ***P*** | **Value** | **Change** | ***P*** | **Value** | **Change** | ***P*** | **Value** | **Change** | ***P*** | **Value** | **Change** | ***P*** |
| Baseline | 37.3 ± 9.4 | – | – | 37.5 ± 10.2 | – | – | 37.5 ± 10.5 | – | – | 37.7 ± 10.9 | – | – | 37.7 ± 10.1 | – | – | 38.6 ± 10.7 | – | – |
| Al300 | 37.3 ± 9.2 | -0.0 ± 0.5 | 1.00000 | 37.5 ± 10.0 | 0.0 ± 0.6 | 1.00000 | 37.5 ± 10.3 | 0.0 ± 0.7 | 0.99963 | 37.9 ± 10.7 | 0.2 ± 0.5 | 0.99963 | 38.0 ± 10.1 | 0.3 ± 0.5 | 0.99963 | 39.0 ± 10.7 | 0.4 ± 0.5 | 0.99376 |
| E20 | 37.4 ± 9.2 | 0.0 ± 0.4 | 0.99963 | 37.6 ± 10.1 | 0.1 ± 0.5 | 1.00000 | 37.5 ± 10.3 | 0.1 ± 0.6 | 0.99963 | 37.9 ± 10.5 | 0.2 ± 0.8 | 0.99963 | 37.9 ± 10.0 | 0.2 ± 0.8 | 0.99963 | 39.0 ± 10.6 | 0.3 ± 0.7 | 0.99963 |
| L100 | 37.3 ± 9.2 | -0.1 ± 0.6 | 1.00000 | 37.5 ± 10.0 | 0.0 ± 0.6 | 1.00000 | 37.5 ± 10.2 | 0.0 ± 0.8 | 0.99963 | 37.9 ± 10.6 | 0.2 ± 0.6 | 0.99963 | 38.0 ± 10.1 | 0.3 ± 0.6 | 0.99963 | 39.0 ± 10.7 | 0.4 ± 0.6 | 0.99376 |
| Aml5 | 36.8 ± 9.1 | -0.5 ± 0.7 | 0.99376 | 37.1 ± 9.9 | -0.4 ± 0.7 | 0.99376 | 37.1 ± 10.1 | -0.4 ± 0.7 | 0.99376 | 37.3 ± 10.5 | -0.4 ± 0.7 | 0.96707 | 37.2 ± 9.9 | -0.6 ± 0.7 | 0.99963 | 38.1 ± 10.5 | -0.5 ± 0.6 | 0.96707 |
| B5 | 45.1 ± 10.7 | 7.8 ± 2.7 | SS | 45.2 ± 12.4 | 7.7 ± 3.3 | 0.00136 | 45.4 ± 12.6 | 8.0 ± 3.2 | 0.00079 | 46.1 ± 13.1 | 8.3 ± 3.9 | 0.00025 | 46.7 ± 12.2 | 9.0 ± 4.3 | SS | 49.0 ± 14.0 | 10.4 ± 6.9 | 0.00004 |
| H12.5 | 37.4 ± 9.4 | 0.0 ± 0.8 | 1.00000 | 37.5 ± 10.1 | -0.0 ± 0.8 | 1.00000 | 37.6 ± 10.3 | 0.2 ± 1.2 | 0.99963 | 37.7 ± 10.6 | -0.0 ± 1.3 | 0.99963 | 37.6 ± 10.1 | -0.1 ± 1.0 | 1.00000 | 38.6 ± 10.6 | -0.1 ± 0.9 | 1.00000 |
| Al300 Aml5 | 36.7 ± 9.0 | -0.7 ± 1.4 | 0.90621 | 37.1 ± 9.5 | -0.4 ± 1.5 | 0.96707 | 37.0 ± 9.7 | -0.5 ± 1.7 | 0.81275 | 37.4 ± 10.2 | -0.3 ± 1.4 | 0.69937 | 37.3 ± 9.9 | -0.4 ± 1.4 | 0.99963 | 38.4 ± 10.4 | -0.2 ± 1.0 | 0.99963 |
| Al300 B5 | 45.4 ± 11.4 | 8.1 ± 4.2 | SS | 45.3 ± 13.2 | 7.8 ± 4.7 | 0.00136 | 45.0 ± 12.3 | 7.5 ± 3.1 | 0.00079 | 44.7 ± 11.6 | 7.0 ± 2.3 | 0.00045 | 44.7 ± 10.4 | 7.0 ± 1.7 | 0.00025 | 46.1 ± 11.5 | 7.5 ± 2.6 | 0.00079 |
| Al300 H12.5 | 37.2 ± 9.2 | -0.2 ± 1.4 | 0.99963 | 37.4 ± 9.7 | -0.1 ± 1.6 | 0.99963 | 37.4 ± 9.9 | -0.0 ± 1.9 | 0.96707 | 37.8 ± 10.3 | 0.1 ± 1.7 | 0.99963 | 37.8 ± 10.2 | 0.1 ± 1.6 | 1.00000 | 38.9 ± 10.6 | 0.3 ± 1.0 | 0.99963 |
| E20 Aml5 | 36.7 ± 9.0 | -0.6 ± 1.2 | 0.90621 | 37.1 ± 9.6 | -0.4 ± 1.3 | 0.96707 | 37.0 ± 9.8 | -0.4 ± 1.5 | 0.90621 | 37.3 ± 10.0 | -0.4 ± 1.8 | 0.58062 | 37.2 ± 9.8 | -0.6 ± 1.9 | 0.96707 | 38.2 ± 10.3 | -0.5 ± 1.4 | 0.96707 |
| E20 B5 | 45.0 ± 10.8 | 7.6 ± 3.1 | 0.00004 | 45.0 ± 12.6 | 7.5 ± 3.8 | 0.00136 | 44.9 ± 12.1 | 7.4 ± 2.8 | 0.00079 | 44.6 ± 11.6 | 6.8 ± 2.6 | 0.00079 | 44.5 ± 10.2 | 6.7 ± 1.9 | 0.00045 | 45.6 ± 11.4 | 7.0 ± 2.7 | 0.00079 |
| E20 H12.5 | 37.3 ± 9.3 | -0.1 ± 1.2 | 1.00000 | 37.4 ± 9.8 | -0.0 ± 1.4 | 0.99963 | 37.5 ± 9.9 | 0.0 ± 1.8 | 0.99376 | 37.7 ± 10.2 | -0.0 ± 2.0 | 0.81275 | 37.6 ± 10.1 | -0.1 ± 2.1 | 0.99376 | 38.7 ± 10.5 | 0.1 ± 1.3 | 1.00000 |
| L100 Aml5 | 36.6 ± 9.0 | -0.7 ± 1.4 | 0.90621 | 37.0 ± 9.5 | -0.5 ± 1.6 | 0.90621 | 36.9 ± 9.7 | -0.5 ± 1.8 | 0.81275 | 37.4 ± 10.1 | -0.3 ± 1.6 | 0.69937 | 37.3 ± 9.8 | -0.4 ± 1.6 | 0.99963 | 38.3 ± 10.4 | -0.3 ± 1.1 | 0.96707 |
| L100 B5 | 45.8 ± 11.9 | 8.5 ± 5.2 | SS | 45.6 ± 14.0 | 8.1 ± 5.8 | 0.00136 | 45.1 ± 12.5 | 7.7 ± 3.5 | 0.00079 | 44.6 ± 11.6 | 6.9 ± 2.4 | 0.00045 | 44.6 ± 10.3 | 6.9 ± 1.7 | 0.00045 | 45.9 ± 11.4 | 7.3 ± 2.6 | 0.00079 |
| L100 H12.5 | 37.1 ± 9.2 | -0.2 ± 1.5 | 0.99376 | 37.3 ± 9.6 | -0.2 ± 1.7 | 0.99376 | 37.3 ± 9.8 | -0.1 ± 2.1 | 0.90621 | 37.8 ± 10.3 | 0.1 ± 1.8 | 0.99376 | 37.7 ± 10.1 | -0.0 ± 1.8 | 1.00000 | 38.9 ± 10.6 | 0.2 ± 1.1 | 1.00000 |
| Al300 Aml5/B5 | 43.0 ± 9.6 | 5.6 ± 1.7 | 0.00025 | 43.2 ± 10.8 | 5.7 ± 2.0 | 0.01581 | 43.0 ± 10.5 | 5.6 ± 1.6 | 0.01008 | 43.0 ± 10.4 | 5.3 ± 2.0 | 0.00630 | 42.8 ± 9.8 | 5.0 ± 2.1 | 0.00630 | 43.9 ± 10.8 | 5.2 ± 1.9 | 0.00386 |
| Al300 Aml5/H12.5 | 36.2 ± 9.2 | -1.2 ± 2.4 | 0.81275 | 36.5 ± 9.2 | -1.0 ± 2.8 | 0.69937 | 36.4 ± 9.6 | -1.0 ± 3.1 | 0.69937 | 37.0 ± 10.0 | -0.7 ± 2.6 | 0.46756 | 36.9 ± 10.2 | -0.8 ± 2.9 | 0.90621 | 38.1 ± 10.4 | -0.5 ± 1.8 | 0.96707 |
| Al300 B5/H12.5 | 42.7 ± 9.6 | 5.3 ± 1.8 | 0.00079 | 42.7 ± 10.2 | 5.2 ± 2.2 | 0.02431 | 42.6 ± 10.3 | 5.1 ± 2.6 | 0.00630 | 42.7 ± 10.3 | 5.0 ± 3.0 | 0.01008 | 42.5 ± 10.2 | 4.8 ± 3.1 | 0.02431 | 43.7 ± 10.7 | 5.1 ± 2.3 | 0.01008 |
| E20 Aml5/B5 | 43.1 ± 9.6 | 5.8 ± 1.2 | 0.00025 | 43.3 ± 10.7 | 5.8 ± 1.7 | 0.01581 | 43.1 ± 10.5 | 5.7 ± 1.5 | 0.01008 | 42.7 ± 10.2 | 5.0 ± 2.2 | 0.01008 | 42.2 ± 9.7 | 4.5 ± 2.5 | 0.01008 | 43.0 ± 10.8 | 4.4 ± 2.3 | 0.02431 |
| E20 Aml5/H12.5 | 36.4 ± 9.2 | -0.9 ± 2.1 | 0.81275 | 36.7 ± 9.3 | -0.8 ± 2.5 | 0.81275 | 36.6 ± 9.6 | -0.8 ± 2.9 | 0.69937 | 36.6 ± 9.8 | -1.1 ± 3.1 | 0.21055 | 36.5 ± 10.2 | -1.3 ± 3.5 | 0.81275 | 37.5 ± 10.3 | -1.2 ± 2.3 | 0.69937 |
| E20 B5/H12.5 | 42.8 ± 9.6 | 5.5 ± 1.8 | 0.00079 | 42.8 ± 10.1 | 5.3 ± 2.1 | 0.02431 | 42.7 ± 10.3 | 5.3 ± 2.5 | 0.00630 | 42.3 ± 10.2 | 4.6 ± 3.3 | 0.01581 | 41.9 ± 10.1 | 4.1 ± 3.5 | 0.03663 | 42.7 ± 10.7 | 4.1 ± 2.8 | 0.03663 |
| L100 Aml5/B5 | 42.9 ± 9.7 | 5.6 ± 2.0 | 0.00045 | 43.2 ± 11.0 | 5.7 ± 2.2 | 0.01581 | 43.0 ± 10.5 | 5.5 ± 1.7 | 0.01581 | 42.9 ± 10.3 | 5.1 ± 2.1 | 0.00630 | 42.5 ± 9.7 | 4.8 ± 2.2 | 0.01008 | 43.6 ± 10.8 | 4.9 ± 2.0 | 0.00630 |
| L100 Aml5/H12.5 | 36.0 ± 9.2 | -1.3 ± 2.5 | 0.58062 | 36.3 ± 9.2 | -1.2 ± 2.9 | 0.58062 | 36.3 ± 9.6 | -1.2 ± 3.3 | 0.69937 | 36.8 ± 9.9 | -0.9 ± 2.8 | 0.36672 | 36.7 ± 10.2 | -1.0 ± 3.1 | 0.81275 | 37.9 ± 10.3 | -0.7 ± 1.9 | 0.90621 |
| L100 B5/H12.5 | 42.6 ± 9.6 | 5.3 ± 1.9 | 0.00079 | 42.7 ± 10.2 | 5.2 ± 2.3 | 0.02431 | 42.5 ± 10.3 | 5.0 ± 2.6 | 0.00630 | 42.5 ± 10.2 | 4.8 ± 3.1 | 0.01581 | 42.3 ± 10.2 | 4.5 ± 3.3 | 0.02431 | 43.4 ± 10.7 | 4.8 ± 2.5 | 0.01581 |

**Al300** = aliskiren 300 mg; **Aml5** = amlodipine 5 mg; **B5** = bisoprolol 5 mg; **E20** = enalapril 20 mg; **H12.5** = hydrochlorothiazide 12.5 mg; **L100** = losartan 100 mg; **SS** = statistically significant ($P$ < 0.00001)

**Table S27.** *P*-values calculated using the Kolmogorov-Smirnov test for changes in left ventricular end-systolic volume in populations ($n$ = 100) with different ACE activity receiving the same regimens (case 1: $c_{ACE}$ = 7.0 h$^{-1}$, case 2: $c_{ACE}$ = 8.9 h$^{-1}$, case 3: $c_{ACE}$ = 10.8 h$^{-1}$, case 4: $c_{ACE}$ = 42.3 h$^{-1}$, case 5: $c_{ACE}$ = 54.1 h$^{-1}$, case 6: $c_{ACE}$ = 65.9 h$^{-1}$; *P*-value for case *i vs*. case *j* is denoted $P_{ij}$)

| **Regimens** | $\boldsymbol{P_{12}}$ | $\boldsymbol{P_{13}}$ | $\boldsymbol{P_{23}}$ | $\boldsymbol{P_{14}}$ | $\boldsymbol{P_{15}}$ | $\boldsymbol{P_{16}}$ | $\boldsymbol{P_{24}}$ | $\boldsymbol{P_{25}}$ | $\boldsymbol{P_{26}}$ | $\boldsymbol{P_{34}}$ | $\boldsymbol{P_{35}}$ | $\boldsymbol{P_{36}}$ | $\boldsymbol{P_{45}}$ | $\boldsymbol{P_{46}}$ | $\boldsymbol{P_{56}}$ |
|---|---|---|---|---|---|---|---|---|---|---|---|---|---|---|---|
| Al300 | 0.21055 | 0.58062 | 0.46756 | SS | 0.00013 | SS | 0.02431 | 0.07832 | SS | 0.00079 | 0.00079 | SS | 0.96707 | 0.01008 | 0.00386 |
| E20 | 0.28093 | 0.46756 | 0.46756 | 0.00007 | 0.00007 | SS | 0.01008 | 0.00136 | 0.00002 | 0.00232 | 0.00232 | 0.00004 | 0.69937 | 0.11113 | 0.28093 |
| L100 | 0.15454 | 0.36672 | 0.58062 | 0.00002 | 0.00013 | SS | 0.01581 | 0.05410 | SS | 0.00045 | 0.00136 | SS | 0.96707 | 0.01581 | 0.03663 |
| Aml5 | 0.02431 | 0.46756 | 0.03663 | 0.81275 | 0.69937 | 0.69937 | 0.21055 | 0.01581 | 0.00386 | 0.90621 | 0.28093 | 0.28093 | 0.46756 | 0.46756 | 0.90621 |
| B5 | 0.11113 | 0.58062 | 0.03663 | 0.07832 | 0.11113 | 0.00025 | 0.07832 | 0.00079 | 0.00079 | 0.58062 | 0.21055 | 0.01581 | 0.21055 | 0.21055 | 0.28093 |
| H12.5 | 0.96707 | 0.21055 | 0.21055 | 0.58062 | 0.36672 | 0.21055 | 0.46756 | 0.11113 | 0.11113 | 0.69937 | 0.21055 | 0.21055 | 0.81275 | 0.81275 | 0.99376 |
| Al300<br>Aml5 | 0.15454 | 0.15454 | 0.46756 | 0.03663 | 0.05410 | 0.00025 | 0.58062 | 0.90621 | 0.07832 | 0.11113 | 0.15454 | 0.00232 | 0.69937 | 0.28093 | 0.36672 |
| Al300<br>B5 | 0.03663 | 0.81275 | 0.28093 | 0.02431 | 0.03663 | 0.21055 | 0.21055 | 0.05410 | 0.01581 | 0.07832 | 0.15454 | 0.28093 | 0.28093 | 0.11113 | 0.81275 |
| Al300<br>H12.5 | 0.36672 | 0.36672 | 0.69937 | 0.07832 | 0.05410 | 0.00045 | 0.69937 | 0.69937 | 0.01008 | 0.21055 | 0.21055 | 0.00136 | 0.81275 | 0.11113 | 0.28093 |
| E20<br>Aml5 | 0.21055 | 0.11113 | 0.36672 | 0.07832 | 0.21055 | 0.58062 | 0.46756 | 0.69937 | 0.58062 | 0.58062 | 0.58062 | 0.58062 | 0.90621 | 0.46756 | 0.69937 |
| E20<br>B5 | 0.02431 | 0.81275 | 0.36672 | 0.00386 | 0.05410 | 0.00630 | 0.36672 | 0.21055 | 0.46756 | 0.02431 | 0.05410 | 0.05410 | 0.58062 | 0.69937 | 0.90621 |
| E20<br>H12.5 | 0.46756 | 0.28093 | 0.46756 | 0.15454 | 0.00630 | 0.21055 | 0.46756 | 0.15454 | 0.58062 | 0.81275 | 0.21055 | 0.69937 | 0.69937 | 0.58062 | 0.46756 |
| L100<br>Aml5 | 0.15454 | 0.15454 | 0.36672 | 0.05410 | 0.03663 | 0.00136 | 0.69937 | 0.99376 | 0.15454 | 0.11113 | 0.28093 | 0.00630 | 0.90621 | 0.46756 | 0.58062 |
| L100<br>B5 | 0.05410 | 0.69937 | 0.21055 | 0.00386 | 0.01581 | 0.03663 | 0.11113 | 0.03663 | 0.01008 | 0.05410 | 0.07832 | 0.28093 | 0.58062 | 0.15454 | 0.96707 |
| L100<br>H12.5 | 0.36672 | 0.46756 | 0.81275 | 0.07832 | 0.02431 | 0.00232 | 0.58062 | 0.36672 | 0.02431 | 0.21055 | 0.11113 | 0.01581 | 0.69937 | 0.36672 | 0.46756 |
| Al300<br>Aml5/B5 | 0.46756 | 0.15454 | 0.46756 | 0.69937 | 0.21055 | 0.36672 | 0.58062 | 0.07832 | 0.07832 | 0.81275 | 0.36672 | 0.21055 | 0.81275 | 0.46756 | 0.90621 |
| Al300<br>Aml5/H12.5 | 0.36672 | 0.46756 | 0.96707 | 0.02431 | 0.00386 | 0.00386 | 0.58062 | 0.36672 | 0.03663 | 0.58062 | 0.21055 | 0.02431 | 0.46756 | 0.36672 | 0.36672 |
| Al300<br>B5/H12.5 | 0.58062 | 0.15454 | 0.69937 | 0.05410 | 0.03663 | 0.21055 | 0.46756 | 0.15454 | 0.46756 | 0.46756 | 0.15454 | 0.58062 | 0.81275 | 0.58062 | 0.21055 |
| E20<br>Aml5/B5 | 0.81275 | 0.15454 | 0.69937 | 0.01008 | 0.00002 | SS | 0.01581 | 0.00004 | SS | 0.28093 | 0.00386 | 0.00013 | 0.36672 | 0.01581 | 0.36672 |
| E20<br>Aml5/H12.5 | 0.28093 | 0.46756 | 0.99376 | 0.28093 | 0.11113 | 0.36672 | 0.69937 | 0.81275 | 0.36672 | 0.96707 | 0.90621 | 0.69937 | 0.90621 | 0.58062 | 0.15454 |
| E20<br>B5/H12.5 | 0.46756 | 0.21055 | 0.58062 | 0.00630 | 0.00013 | 0.00013 | 0.02431 | 0.01008 | 0.00079 | 0.21055 | 0.11113 | 0.00232 | 0.96707 | 0.02431 | 0.11113 |
| L100<br>Aml5/B5 | 0.46756 | 0.15454 | 0.36672 | 0.46756 | 0.15454 | 0.28093 | 0.46756 | 0.02431 | 0.03663 | 0.58062 | 0.28093 | 0.07832 | 0.69937 | 0.28093 | 0.69937 |
| L100<br>Aml5/H12.5 | 0.36672 | 0.69937 | 0.99376 | 0.03663 | 0.00386 | 0.03663 | 0.69937 | 0.28093 | 0.15454 | 0.58062 | 0.15454 | 0.05410 | 0.69937 | 0.46756 | 0.28093 |
| L100<br>B5/H12.5 | 0.58062 | 0.15454 | 0.58062 | 0.03663 | 0.01581 | 0.05410 | 0.28093 | 0.07832 | 0.28093 | 0.81275 | 0.36672 | 0.58062 | 0.99376 | 0.36672 | 0.21055 |

**Al300 =** aliskiren 300 mg; **Aml5** = amlodipine 5 mg; **B5** = bisoprolol 5 mg; **E20** = enalapril 20 mg; **H12.5** = hydrochlorothiazide 12.5 mg; **L100** = losartan 100 mg; **SS** = statistically significant ($P$ < 0.00001)

**Figure S21.** Simulated change in right ventricular end-diastolic pressure from baseline to week 4 (mean ± SD, $n$ = 100)

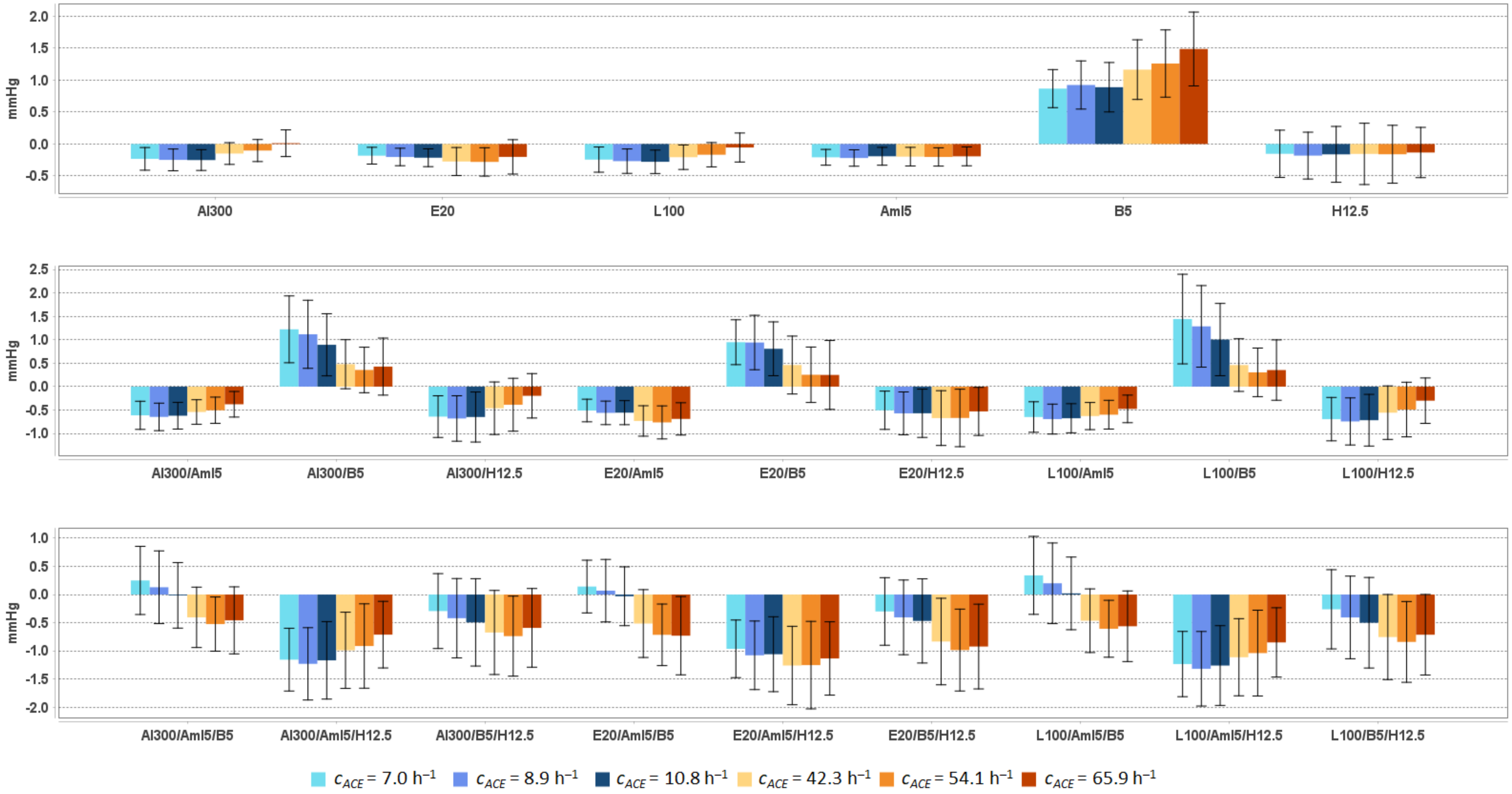


**Al300 =** aliskiren 300 mg; **Aml5** = amlodipine 5 mg; **B5** = bisoprolol 5 mg; **E20** = enalapril 20 mg; **H12.5** = hydrochlorothiazide 12.5 mg; **L100** = losartan 100 mg

**Table S28.** Simulated response of right ventricular end-diastolic pressure to antihypertensive therapy in virtual hypertensive populations ($n$ = 100) with different ACE activity, including $P$-values (Kolmogorov-Smirnov test) for endpoint *vs.* baseline; data are presented as mean ± SD in mmHg

| **Regimens** | $c_{ACE}$ = 7.0 h$^{-1}$ | | | $c_{ACE}$ = 8.9 h$^{-1}$ | | | $c_{ACE}$ = 10.8 h$^{-1}$ | | | $c_{ACE}$ = 42.3 h$^{-1}$ | | | $c_{ACE}$ = 54.1 h$^{-1}$ | | | $c_{ACE}$ = 65.9 h$^{-1}$ | | |
|---|---|---|---|---|---|---|---|---|---|---|---|---|---|---|---|---|---|---|
| | **Value** | **Change** | ***P*** | **Value** | **Change** | ***P*** | **Value** | **Change** | ***P*** | **Value** | **Change** | ***P*** | **Value** | **Change** | ***P*** | **Value** | **Change** | ***P*** |
| Baseline | 5.4 ± 1.2 | – | – | 5.8 ± 1.1 | – | – | 5.2 ± 1.2 | – | – | 5.6 ± 1.1 | – | – | 5.6 ± 1.1 | – | – | 5.7 ± 1.0 | – | – |
| Al300 | 5.2 ± 1.1 | -0.2 ± 0.2 | 0.36672 | 5.5 ± 1.1 | -0.2 ± 0.2 | 0.05410 | 5.0 ± 1.2 | -0.3 ± 0.2 | 0.21055 | 5.5 ± 1.1 | -0.1 ± 0.2 | 0.69937 | 5.5 ± 1.1 | -0.1 ± 0.2 | 0.90621 | 5.7 ± 1.0 | 0.0 ± 0.2 | 1.00000 |
| E20 | 5.2 ± 1.1 | -0.2 ± 0.1 | 0.69937 | 5.6 ± 1.1 | -0.2 ± 0.1 | 0.11113 | 5.0 ± 1.2 | -0.2 ± 0.1 | 0.28093 | 5.4 ± 1.0 | -0.3 ± 0.2 | 0.28093 | 5.3 ± 1.0 | -0.3 ± 0.2 | 0.07832 | 5.5 ± 1.0 | -0.2 ± 0.3 | 0.15454 |
| L100 | 5.2 ± 1.1 | -0.2 ± 0.2 | 0.28093 | 5.5 ± 1.1 | -0.3 ± 0.2 | 0.03663 | 4.9 ± 1.2 | -0.3 ± 0.2 | 0.15454 | 5.4 ± 1.1 | -0.2 ± 0.2 | 0.46756 | 5.4 ± 1.1 | -0.2 ± 0.2 | 0.58062 | 5.6 ± 1.0 | -0.1 ± 0.2 | 0.90621 |
| Aml5 | 5.2 ± 1.1 | -0.2 ± 0.1 | 0.58062 | 5.5 ± 1.1 | -0.2 ± 0.1 | 0.21055 | 5.0 ± 1.2 | -0.2 ± 0.1 | 0.28093 | 5.4 ± 1.1 | -0.2 ± 0.1 | 0.58062 | 5.4 ± 1.1 | -0.2 ± 0.1 | 0.15454 | 5.5 ± 1.0 | -0.2 ± 0.1 | 0.36672 |
| B5 | 6.3 ± 1.3 | 0.9 ± 0.3 | 0.00013 | 6.7 ± 1.3 | 0.9 ± 0.4 | SS | 6.1 ± 1.5 | 0.9 ± 0.4 | 0.00007 | 6.8 ± 1.3 | 1.2 ± 0.5 | SS | 6.9 ± 1.4 | 1.3 ± 0.5 | SS | 7.2 ± 1.2 | 1.5 ± 0.6 | SS |
| H12.5 | 5.2 ± 1.2 | -0.2 ± 0.4 | 0.36672 | 5.6 ± 1.1 | -0.2 ± 0.4 | 0.21055 | 5.1 ± 1.2 | -0.2 ± 0.4 | 0.69937 | 5.5 ± 1.2 | -0.2 ± 0.5 | 0.46756 | 5.4 ± 1.1 | -0.2 ± 0.5 | 0.58062 | 5.6 ± 1.1 | -0.1 ± 0.4 | 0.90621 |
| Al300<br>Aml5 | 4.8 ± 1.0 | -0.6 ± 0.3 | 0.00045 | 5.1 ± 1.0 | -0.6 ± 0.3 | 0.00007 | 4.6 ± 1.1 | -0.6 ± 0.3 | 0.00079 | 5.1 ± 1.0 | -0.5 ± 0.3 | 0.00386 | 5.1 ± 1.0 | -0.5 ± 0.3 | 0.00232 | 5.3 ± 1.0 | -0.4 ± 0.3 | 0.02431 |
| Al300<br>B5 | 6.6 ± 1.5 | 1.2 ± 0.7 | SS | 6.9 ± 1.5 | 1.1 ± 0.7 | SS | 6.1 ± 1.6 | 0.9 ± 0.7 | 0.00045 | 6.1 ± 1.2 | 0.5 ± 0.5 | 0.01581 | 5.9 ± 1.2 | 0.4 ± 0.5 | 0.00630 | 6.1 ± 1.2 | 0.4 ± 0.6 | 0.03663 |
| Al300<br>H12.5 | 4.8 ± 1.1 | -0.6 ± 0.4 | 0.00004 | 5.1 ± 1.0 | -0.7 ± 0.5 | 0.00004 | 4.6 ± 1.2 | -0.6 ± 0.5 | 0.00025 | 5.2 ± 1.2 | -0.5 ± 0.6 | 0.00386 | 5.2 ± 1.1 | -0.4 ± 0.6 | 0.07832 | 5.5 ± 1.1 | -0.2 ± 0.5 | 0.46756 |
| E20<br>Aml5 | 4.9 ± 1.1 | -0.5 ± 0.2 | 0.00386 | 5.2 ± 1.0 | -0.6 ± 0.3 | 0.00013 | 4.7 ± 1.1 | -0.6 ± 0.3 | 0.00630 | 4.9 ± 1.0 | -0.7 ± 0.3 | 0.00013 | 4.8 ± 1.0 | -0.8 ± 0.4 | SS | 5.0 ± 1.0 | -0.7 ± 0.3 | 0.00013 |
| E20<br>B5 | 6.3 ± 1.4 | 0.9 ± 0.5 | 0.00025 | 6.7 ± 1.4 | 0.9 ± 0.6 | SS | 6.0 ± 1.6 | 0.8 ± 0.6 | 0.00079 | 6.1 ± 1.3 | 0.5 ± 0.6 | 0.01581 | 5.8 ± 1.3 | 0.3 ± 0.6 | 0.07832 | 5.9 ± 1.3 | 0.3 ± 0.7 | 0.11113 |
| E20<br>H12.5 | 4.9 ± 1.1 | -0.5 ± 0.4 | 0.00386 | 5.2 ± 1.0 | -0.6 ± 0.5 | 0.00025 | 4.6 ± 1.2 | -0.6 ± 0.5 | 0.00386 | 5.0 ± 1.1 | -0.7 ± 0.6 | 0.00013 | 4.9 ± 1.1 | -0.7 ± 0.6 | 0.00025 | 5.2 ± 1.1 | -0.5 ± 0.5 | 0.00386 |
| L100<br>Aml5 | 4.7 ± 1.0 | -0.6 ± 0.3 | 0.00013 | 5.1 ± 1.0 | -0.7 ± 0.3 | 0.00002 | 4.5 ± 1.1 | -0.7 ± 0.3 | 0.00045 | 5.0 ± 1.0 | -0.6 ± 0.3 | 0.00079 | 5.0 ± 1.0 | -0.6 ± 0.3 | 0.00025 | 5.2 ± 1.0 | -0.5 ± 0.3 | 0.01008 |
| L100<br>B5 | 6.8 ± 1.7 | 1.4 ± 1.0 | SS | 7.1 ± 1.6 | 1.3 ± 0.9 | SS | 6.2 ± 1.7 | 1.0 ± 0.8 | 0.00007 | 6.1 ± 1.3 | 0.5 ± 0.6 | 0.01581 | 5.9 ± 1.2 | 0.3 ± 0.5 | 0.01581 | 6.0 ± 1.2 | 0.4 ± 0.6 | 0.07832 |
| L100<br>H12.5 | 4.7 ± 1.1 | -0.7 ± 0.5 | 0.00002 | 5.0 ± 1.0 | -0.7 ± 0.5 | SS | 4.5 ± 1.2 | -0.7 ± 0.6 | 0.00013 | 5.1 ± 1.2 | -0.6 ± 0.6 | 0.00045 | 5.1 ± 1.1 | -0.5 ± 0.6 | 0.01581 | 5.4 ± 1.1 | -0.3 ± 0.5 | 0.07832 |
| Al300<br>Aml5/B5 | 5.6 ± 1.3 | 0.3 ± 0.6 | 0.28093 | 5.9 ± 1.3 | 0.1 ± 0.6 | 0.21055 | 5.2 ± 1.4 | -0.0 ± 0.6 | 0.69937 | 5.2 ± 1.1 | -0.4 ± 0.5 | 0.01581 | 5.1 ± 1.1 | -0.5 ± 0.5 | 0.00386 | 5.2 ± 1.1 | -0.5 ± 0.6 | 0.01008 |
| Al300<br>Aml5/H12.5 | 4.2 ± 1.0 | -1.2 ± 0.6 | SS | 4.5 ± 1.0 | -1.2 ± 0.6 | SS | 4.0 ± 1.2 | -1.2 ± 0.7 | SS | 4.6 ± 1.2 | -1.0 ± 0.7 | SS | 4.7 ± 1.1 | -0.9 ± 0.7 | SS | 5.0 ± 1.1 | -0.7 ± 0.6 | 0.00025 |
| Al300<br>B5/H12.5 | 5.1 ± 1.2 | -0.3 ± 0.7 | 0.05410 | 5.3 ± 1.2 | -0.4 ± 0.7 | 0.00386 | 4.7 ± 1.4 | -0.5 ± 0.8 | 0.00386 | 5.0 ± 1.2 | -0.7 ± 0.7 | 0.00013 | 4.9 ± 1.1 | -0.7 ± 0.7 | 0.00002 | 5.1 ± 1.2 | -0.6 ± 0.7 | 0.00232 |
| E20<br>Aml5/B5 | 5.5 ± 1.2 | 0.1 ± 0.5 | 0.90621 | 5.8 ± 1.2 | 0.1 ± 0.6 | 0.36672 | 5.2 ± 1.4 | -0.0 ± 0.5 | 0.69937 | 5.1 ± 1.2 | -0.5 ± 0.6 | 0.00079 | 4.9 ± 1.1 | -0.7 ± 0.5 | 0.00013 | 5.0 ± 1.2 | -0.7 ± 0.7 | 0.00007 |
| E20<br>Aml5/H12.5 | 4.4 ± 1.1 | -1.0 ± 0.5 | SS | 4.7 ± 1.0 | -1.1 ± 0.6 | SS | 4.2 ± 1.2 | -1.1 ± 0.7 | SS | 4.4 ± 1.1 | -1.3 ± 0.7 | SS | 4.3 ± 1.1 | -1.3 ± 0.8 | SS | 4.6 ± 1.1 | -1.1 ± 0.7 | SS |
| E20<br>B5/H12.5 | 5.1 ± 1.2 | -0.3 ± 0.6 | 0.02431 | 5.4 ± 1.2 | -0.4 ± 0.7 | 0.00630 | 4.7 ± 1.3 | -0.5 ± 0.7 | 0.00386 | 4.8 ± 1.2 | -0.8 ± 0.8 | SS | 4.6 ± 1.1 | -1.0 ± 0.7 | SS | 4.8 ± 1.2 | -0.9 ± 0.8 | 0.00002 |
| L100<br>Aml5/B5 | 5.7 ± 1.4 | 0.3 ± 0.7 | 0.15454 | 6.0 ± 1.3 | 0.2 ± 0.7 | 0.07832 | 5.2 ± 1.5 | 0.0 ± 0.6 | 0.58062 | 5.2 ± 1.1 | -0.5 ± 0.6 | 0.01581 | 5.0 ± 1.1 | -0.6 ± 0.5 | 0.00079 | 5.1 ± 1.2 | -0.6 ± 0.6 | 0.00232 |
| L100<br>Aml5/H12.5 | 4.2 ± 1.0 | -1.2 ± 0.6 | SS | 4.4 ± 1.0 | -1.3 ± 0.7 | SS | 4.0 ± 1.1 | -1.3 ± 0.7 | SS | 4.5 ± 1.1 | -1.1 ± 0.7 | SS | 4.6 ± 1.1 | -1.0 ± 0.8 | SS | 4.8 ± 1.1 | -0.8 ± 0.6 | SS |
| L100<br>B5/H12.5 | 5.1 ± 1.3 | -0.3 ± 0.7 | 0.07832 | 5.4 ± 1.2 | -0.4 ± 0.7 | 0.00630 | 4.7 ± 1.4 | -0.5 ± 0.8 | 0.00386 | 4.9 ± 1.2 | -0.8 ± 0.8 | 0.00007 | 4.8 ± 1.1 | -0.8 ± 0.7 | SS | 5.0 ± 1.2 | -0.7 ± 0.7 | 0.00045 |

**Al300 =** aliskiren 300 mg; **Aml5** = amlodipine 5 mg; **B5** = bisoprolol 5 mg; **E20** = enalapril 20 mg; **H12.5** = hydrochlorothiazide 12.5 mg; **L100** = losartan 100 mg; **SS** = statistically significant ($P$ < 0.00001)

**Table S29.** *P*-values calculated using the Kolmogorov-Smirnov test for changes in right ventricular end-diastolic pressure in populations ($n$ = 100) with different ACE activity receiving the same regimens (case 1: $c_{ACE}$ = 7.0 h$^{-1}$, case 2: $c_{ACE}$ = 8.9 h$^{-1}$, case 3: $c_{ACE}$ = 10.8 h$^{-1}$, case 4: $c_{ACE}$ = 42.3 h$^{-1}$, case 5: $c_{ACE}$ = 54.1 h$^{-1}$, case 6: $c_{ACE}$ = 65.9 h$^{-1}$; *P*-value for case *i* *vs*. case *j* is denoted $P_{ij}$)

| Regimens | $P_{12}$ | $P_{13}$ | $P_{23}$ | $P_{14}$ | $P_{15}$ | $P_{16}$ | $P_{24}$ | $P_{25}$ | $P_{26}$ | $P_{34}$ | $P_{35}$ | $P_{36}$ | $P_{45}$ | $P_{46}$ | $P_{56}$ |
|---|---|---|---|---|---|---|---|---|---|---|---|---|---|---|---|
| Al300 | 0.46756 | 0.05410 | 0.36672 | 0.02431 | 0.00004 | SS | 0.00386 | SS | SS | 0.00232 | SS | SS | 0.15454 | SS | 0.00386 |
| E20 | 0.28093 | 0.02431 | 0.15454 | 0.00025 | 0.00136 | 0.01008 | 0.00630 | 0.00232 | 0.02431 | 0.01008 | 0.00630 | 0.03663 | 0.90621 | 0.15454 | 0.11113 |
| L100 | 0.28093 | 0.01008 | 0.28093 | 0.28093 | 0.05410 | SS | 0.28093 | 0.00630 | SS | 0.01581 | 0.00004 | SS | 0.28093 | 0.00025 | 0.01008 |
| Aml5 | 0.07832 | 0.58062 | 0.21055 | 0.28093 | 0.36672 | 0.36672 | 0.21055 | 0.28093 | 0.11113 | 0.28093 | 0.69937 | 0.81275 | 0.69937 | 0.81275 | 0.99376 |
| B5 | 0.46756 | 0.81275 | 0.46756 | 0.00007 | SS | SS | 0.00079 | 0.00002 | SS | 0.00136 | SS | SS | 0.46756 | 0.00232 | 0.05410 |
| H12.5 | 0.58062 | 0.81275 | 0.69937 | 0.28093 | 0.69937 | 0.81275 | 0.58062 | 0.07832 | 0.21055 | 0.96707 | 0.36672 | 0.46756 | 0.81275 | 0.36672 | 0.81275 |
| Al300 Aml5 | 0.15454 | 0.58062 | 0.69937 | 0.46756 | 0.11113 | 0.00045 | 0.03663 | 0.00136 | SS | 0.15454 | 0.01008 | 0.00002 | 0.58062 | 0.00630 | 0.11113 |
| Al300 B5 | 0.46756 | 0.01008 | 0.11113 | SS | SS | SS | SS | SS | SS | 0.00232 | SS | 0.00002 | 0.11113 | 0.36672 | 0.81275 |
| Al300 H12.5 | 0.81275 | 0.28093 | 0.28093 | 0.01581 | 0.00002 | SS | 0.01008 | SS | SS | 0.15454 | 0.00136 | SS | 0.28093 | 0.00386 | 0.21055 |
| E20 Aml5 | 0.05410 | 0.05410 | 0.81275 | 0.00002 | SS | 0.00004 | 0.00079 | 0.00232 | 0.01008 | 0.00386 | 0.00136 | 0.00630 | 0.99376 | 0.58062 | 0.69937 |
| E20 B5 | 0.69937 | 0.02431 | 0.21055 | SS | SS | SS | 0.00002 | SS | SS | 0.00079 | SS | SS | 0.05410 | 0.07832 | 0.90621 |
| E20 H12.5 | 0.46756 | 0.21055 | 0.46756 | 0.01008 | 0.11113 | 0.36672 | 0.21055 | 0.58062 | 0.81275 | 0.58062 | 0.69937 | 0.90621 | 0.46756 | 0.46756 | 0.46756 |
| L100 Aml5 | 0.11113 | 0.46756 | 0.69937 | 0.96707 | 0.81275 | 0.01581 | 0.21055 | 0.02431 | 0.00013 | 0.81275 | 0.11113 | 0.00232 | 0.69937 | 0.02431 | 0.21055 |
| L100 B5 | 0.36672 | 0.00630 | 0.07832 | SS | SS | SS | SS | SS | SS | 0.00045 | SS | SS | 0.05410 | 0.21055 | 0.81275 |
| L100 H12.5 | 0.81275 | 0.21055 | 0.28093 | 0.03663 | 0.00013 | SS | 0.01581 | 0.00004 | SS | 0.36672 | 0.00630 | 0.00013 | 0.21055 | 0.01008 | 0.36672 |
| Al300 Aml5/B5 | 0.07832 | 0.01008 | 0.36672 | SS | SS | SS | SS | SS | SS | 0.00136 | SS | 0.00002 | 0.11113 | 0.58062 | 0.58062 |
| Al300 Aml5/H12.5 | 0.15454 | 0.36672 | 0.69937 | 0.05410 | 0.00136 | SS | 0.07832 | 0.00025 | SS | 0.36672 | 0.01008 | 0.00079 | 0.21055 | 0.02431 | 0.36672 |
| Al300 B5/H12.5 | 0.15454 | 0.07832 | 0.36672 | 0.00136 | 0.00079 | 0.00232 | 0.11113 | 0.03663 | 0.28093 | 0.21055 | 0.05410 | 0.36672 | 0.81275 | 0.58062 | 0.36672 |
| E20 Aml5/B5 | 0.15454 | 0.01581 | 0.58062 | SS | SS | SS | SS | SS | SS | SS | SS | SS | 0.05410 | 0.05410 | 0.58062 |
| E20 Aml5/H12.5 | 0.03663 | 0.07832 | 0.81275 | 0.00013 | 0.00630 | 0.01581 | 0.21055 | 0.46756 | 0.28093 | 0.11113 | 0.58062 | 0.11113 | 0.69937 | 0.46756 | 0.81275 |
| E20 B5/H12.5 | 0.58062 | 0.05410 | 0.46756 | 0.00002 | SS | SS | 0.00079 | 0.00002 | SS | 0.00386 | 0.00007 | 0.00079 | 0.69937 | 0.28093 | 0.58062 |
| L100 Aml5/B5 | 0.11113 | 0.00630 | 0.28093 | SS | SS | SS | SS | SS | SS | 0.00007 | SS | SS | 0.07832 | 0.36672 | 0.81275 |
| L100 Aml5/H12.5 | 0.21055 | 0.36672 | 0.69937 | 0.28093 | 0.00386 | 0.00013 | 0.11113 | 0.00136 | 0.00013 | 0.58062 | 0.01008 | 0.00386 | 0.36672 | 0.03663 | 0.46756 |
| L100 B5/H12.5 | 0.07832 | 0.05410 | 0.15454 | 0.00004 | SS | 0.00004 | 0.01581 | 0.00232 | 0.00630 | 0.07832 | 0.01008 | 0.07832 | 0.69937 | 0.69937 | 0.58062 |

**Al300 =** aliskiren 300 mg; **Aml5** = amlodipine 5 mg; **B5** = bisoprolol 5 mg; **E20** = enalapril 20 mg; **H12.5** = hydrochlorothiazide 12.5 mg; **L100** = losartan 100 mg; **SS** = statistically significant ($P$ < 0.00001)

**Figure S22.** Simulated change in right ventricular peak systolic pressure from baseline to week 4 (mean ± SD, $n$ = 100)

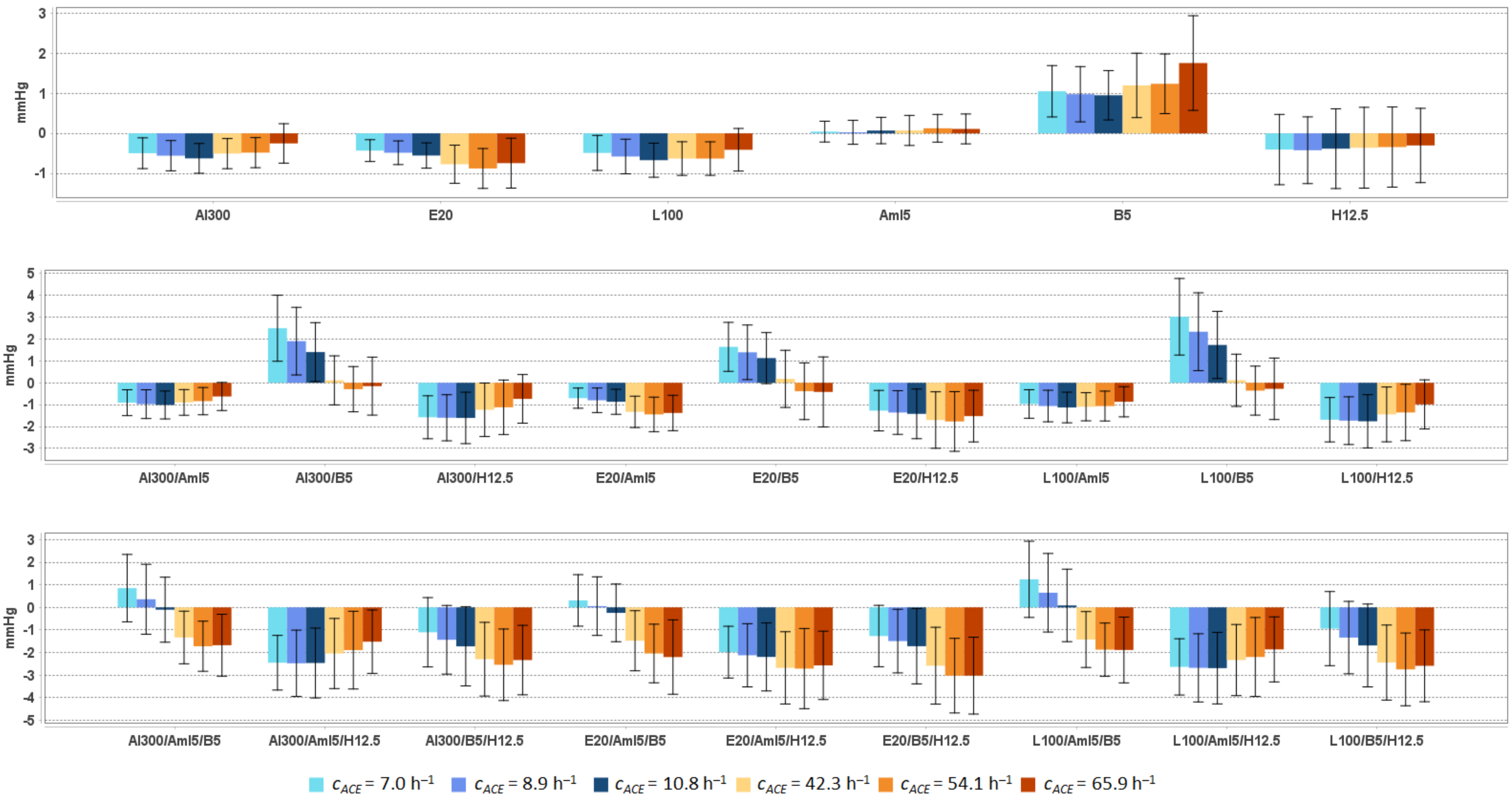


**Al300** = aliskiren 300 mg; **Aml5** = amlodipine 5 mg; **B5** = bisoprolol 5 mg; **E20** = enalapril 20 mg; **H12.5** = hydrochlorothiazide 12.5 mg; **L100** = losartan 100 mg

**Table S30.** Simulated response of right ventricular peak systolic pressure to antihypertensive therapy in virtual hypertensive populations ($n$ = 100) with different ACE activity, including $P$-values (Kolmogorov-Smirnov test) for endpoint *vs.* baseline; data are presented as mean ± SD in mmHg

| **Regimens** | $c_{ACE}$ **= 7.0 h$^{-1}$** | | | $c_{ACE}$ **= 8.9 h$^{-1}$** | | | $c_{ACE}$ **= 10.8 h$^{-1}$** | | | $c_{ACE}$ **= 42.3 h$^{-1}$** | | | $c_{ACE}$ **= 54.1 h$^{-1}$** | | | $c_{ACE}$ **= 65.9 h$^{-1}$** | | |
|---|---|---|---|---|---|---|---|---|---|---|---|---|---|---|---|---|---|---|
| | **Value** | **Change** | ***P*** | **Value** | **Change** | ***P*** | **Value** | **Change** | ***P*** | **Value** | **Change** | ***P*** | **Value** | **Change** | ***P*** | **Value** | **Change** | ***P*** |
| Baseline | 20.2 ± 1.6 | – | – | 19.9 ± 1.4 | – | – | 19.9 ± 1.6 | – | – | 19.7 ± 1.5 | – | – | 20.1 ± 1.6 | – | – | 20.1 ± 1.7 | – | – |
| Al300 | 19.7 ± 1.6 | -0.5 ± 0.4 | 0.15454 | 19.4 ± 1.4 | -0.6 ± 0.4 | 0.02431 | 19.3 ± 1.5 | -0.6 ± 0.4 | 0.05410 | 19.2 ± 1.4 | -0.5 ± 0.4 | 0.05410 | 19.6 ± 1.6 | -0.5 ± 0.4 | 0.15454 | 19.9 ± 1.6 | -0.3 ± 0.5 | 0.36672 |
| E20 | 19.8 ± 1.6 | -0.4 ± 0.3 | 0.21055 | 19.4 ± 1.4 | -0.5 ± 0.3 | 0.05410 | 19.4 ± 1.5 | -0.6 ± 0.3 | 0.07832 | 19.0 ± 1.4 | -0.8 ± 0.5 | 0.00232 | 19.2 ± 1.6 | -0.9 ± 0.5 | 0.00630 | 19.4 ± 1.6 | -0.7 ± 0.6 | 0.01581 |
| L100 | 19.7 ± 1.6 | -0.5 ± 0.4 | 0.15454 | 19.3 ± 1.4 | -0.6 ± 0.4 | 0.02431 | 19.2 ± 1.5 | -0.7 ± 0.4 | 0.02431 | 19.1 ± 1.4 | -0.6 ± 0.4 | 0.01008 | 19.5 ± 1.6 | -0.6 ± 0.4 | 0.07832 | 19.7 ± 1.6 | -0.4 ± 0.5 | 0.36672 |
| Aml5 | 20.2 ± 1.6 | 0.0 ± 0.3 | 0.99376 | 19.9 ± 1.4 | 0.0 ± 0.3 | 0.90621 | 20.0 ± 1.6 | 0.1 ± 0.3 | 0.99376 | 19.8 ± 1.6 | 0.1 ± 0.4 | 0.90621 | 20.2 ± 1.7 | 0.1 ± 0.3 | 0.90621 | 20.2 ± 1.7 | 0.1 ± 0.4 | 0.96707 |
| B5 | 21.2 ± 1.7 | 1.0 ± 0.6 | 0.00013 | 20.9 ± 1.5 | 1.0 ± 0.7 | 0.00007 | 20.9 ± 1.7 | 1.0 ± 0.6 | 0.00079 | 20.9 ± 1.6 | 1.2 ± 0.8 | SS | 21.3 ± 1.8 | 1.2 ± 0.7 | 0.00013 | 21.9 ± 1.9 | 1.8 ± 1.2 | SS |
| H12.5 | 19.8 ± 1.8 | -0.4 ± 0.9 | 0.15454 | 19.5 ± 1.5 | -0.4 ± 0.8 | 0.05410 | 19.5 ± 1.8 | -0.4 ± 1.0 | 0.36672 | 19.4 ± 1.7 | -0.4 ± 1.0 | 0.21055 | 19.8 ± 1.8 | -0.3 ± 1.0 | 0.15454 | 19.8 ± 1.9 | -0.3 ± 0.9 | 0.58062 |
| Al300 Aml5 | 19.3 ± 1.7 | -0.9 ± 0.6 | 0.00386 | 18.9 ± 1.4 | -1.0 ± 0.7 | 0.00025 | 18.9 ± 1.6 | -1.0 ± 0.6 | 0.00079 | 18.8 ± 1.4 | -0.9 ± 0.6 | 0.00079 | 19.3 ± 1.7 | -0.8 ± 0.6 | 0.00630 | 19.5 ± 1.6 | -0.6 ± 0.6 | 0.07832 |
| Al300 B5 | 22.7 ± 2.3 | 2.5 ± 1.5 | SS | 21.8 ± 2.1 | 1.9 ± 1.5 | SS | 21.3 ± 2.1 | 1.4 ± 1.3 | 0.00007 | 19.8 ± 1.6 | 0.1 ± 1.1 | 0.81275 | 19.8 ± 1.8 | -0.3 ± 1.0 | 0.69937 | 20.0 ± 1.8 | -0.2 ± 1.3 | 0.58062 |
| Al300 H12.5 | 18.6 ± 1.8 | -1.6 ± 1.0 | SS | 18.3 ± 1.5 | -1.6 ± 1.1 | SS | 18.3 ± 1.8 | -1.6 ± 1.2 | SS | 18.5 ± 1.7 | -1.2 ± 1.2 | 0.00002 | 19.0 ± 1.9 | -1.1 ± 1.2 | 0.00013 | 19.4 ± 1.8 | -0.7 ± 1.1 | 0.01008 |
| E20 Aml5 | 19.5 ± 1.6 | -0.7 ± 0.5 | 0.02431 | 19.1 ± 1.4 | -0.8 ± 0.6 | 0.00232 | 19.0 ± 1.6 | -0.9 ± 0.6 | 0.00136 | 18.4 ± 1.5 | -1.3 ± 0.7 | SS | 18.7 ± 1.7 | -1.4 ± 0.8 | 0.00004 | 18.8 ± 1.6 | -1.4 ± 0.8 | SS |
| E20 B5 | 21.8 ± 2.0 | 1.6 ± 1.1 | SS | 21.3 ± 1.9 | 1.4 ± 1.2 | SS | 21.0 ± 2.0 | 1.1 ± 1.2 | 0.00136 | 19.9 ± 1.7 | 0.2 ± 1.3 | 0.36672 | 19.7 ± 1.9 | -0.4 ± 1.3 | 0.28093 | 19.7 ± 2.0 | -0.4 ± 1.6 | 0.28093 |
| E20 H12.5 | 18.9 ± 1.8 | -1.3 ± 0.9 | 0.00004 | 18.6 ± 1.5 | -1.4 ± 1.0 | SS | 18.5 ± 1.8 | -1.4 ± 1.1 | SS | 18.0 ± 1.7 | -1.7 ± 1.3 | SS | 18.3 ± 1.9 | -1.8 ± 1.4 | SS | 18.6 ± 1.8 | -1.5 ± 1.2 | SS |
| L100 Aml5 | 19.2 ± 1.7 | -1.0 ± 0.7 | 0.00232 | 18.9 ± 1.4 | -1.1 ± 0.7 | 0.00013 | 18.8 ± 1.6 | -1.1 ± 0.7 | 0.00013 | 18.6 ± 1.5 | -1.1 ± 0.6 | 0.00004 | 19.0 ± 1.7 | -1.1 ± 0.7 | 0.00136 | 19.3 ± 1.6 | -0.9 ± 0.7 | 0.00386 |
| L100 B5 | 23.2 ± 2.5 | 3.0 ± 1.7 | SS | 22.2 ± 2.3 | 2.3 ± 1.8 | SS | 21.6 ± 2.2 | 1.7 ± 1.5 | SS | 19.8 ± 1.7 | 0.1 ± 1.2 | 0.58062 | 19.7 ± 1.8 | -0.4 ± 1.1 | 0.46756 | 19.9 ± 1.9 | -0.3 ± 1.4 | 0.36672 |
| L100 H12.5 | 18.5 ± 1.8 | -1.7 ± 1.0 | SS | 18.2 ± 1.5 | -1.7 ± 1.1 | SS | 18.1 ± 1.8 | -1.8 ± 1.2 | SS | 18.3 ± 1.7 | -1.4 ± 1.3 | SS | 18.7 ± 1.9 | -1.4 ± 1.3 | SS | 19.1 ± 1.8 | -1.0 ± 1.1 | 0.00232 |
| Al300 Aml5/B5 | 21.0 ± 2.2 | 0.9 ± 1.5 | 0.00045 | 20.3 ± 2.1 | 0.4 ± 1.5 | 0.07832 | 19.8 ± 2.1 | -0.1 ± 1.4 | 0.21055 | 18.4 ± 1.6 | -1.3 ± 1.2 | SS | 18.4 ± 1.8 | -1.7 ± 1.1 | SS | 18.5 ± 1.8 | -1.7 ± 1.4 | SS |
| Al300 Aml5/H12.5 | 17.7 ± 2.0 | -2.4 ± 1.2 | SS | 17.4 ± 1.8 | -2.5 ± 1.5 | SS | 17.4 ± 2.1 | -2.5 ± 1.5 | SS | 17.7 ± 1.9 | -2.0 ± 1.6 | SS | 18.2 ± 2.2 | -1.9 ± 1.7 | SS | 18.6 ± 2.0 | -1.5 ± 1.4 | SS |
| Al300 B5/H12.5 | 19.1 ± 2.2 | -1.1 ± 1.5 | 0.00004 | 18.5 ± 1.8 | -1.4 ± 1.5 | SS | 18.2 ± 2.2 | -1.7 ± 1.8 | SS | 17.4 ± 1.9 | -2.3 ± 1.6 | SS | 17.6 ± 2.0 | -2.5 ± 1.6 | SS | 17.8 ± 1.9 | -2.3 ± 1.5 | SS |
| E20 Aml5/B5 | 20.5 ± 2.0 | 0.3 ± 1.1 | 0.11113 | 20.0 ± 1.9 | 0.1 ± 1.3 | 0.58062 | 19.7 ± 2.0 | -0.2 ± 1.3 | 0.21055 | 18.3 ± 1.7 | -1.5 ± 1.3 | SS | 18.1 ± 1.9 | -2.0 ± 1.3 | SS | 17.9 ± 2.0 | -2.2 ± 1.6 | SS |
| E20 Aml5/H12.5 | 18.2 ± 1.9 | -2.0 ± 1.1 | SS | 17.8 ± 1.8 | -2.1 ± 1.4 | SS | 17.7 ± 2.0 | -2.2 ± 1.5 | SS | 17.1 ± 1.9 | -2.7 ± 1.6 | SS | 17.4 ± 2.2 | -2.7 ± 1.8 | SS | 17.6 ± 2.1 | -2.6 ± 1.5 | SS |
| E20 B5/H12.5 | 18.9 ± 2.0 | -1.3 ± 1.4 | SS | 18.4 ± 1.7 | -1.5 ± 1.4 | SS | 18.2 ± 2.1 | -1.7 ± 1.7 | SS | 17.1 ± 1.9 | -2.6 ± 1.7 | SS | 17.1 ± 2.0 | -3.0 ± 1.7 | SS | 17.1 ± 2.0 | -3.0 ± 1.7 | SS |
| L100 Aml5/B5 | 21.4 ± 2.4 | 1.2 ± 1.7 | SS | 20.6 ± 2.2 | 0.6 ± 1.7 | 0.00630 | 20.0 ± 2.2 | 0.1 ± 1.6 | 0.36672 | 18.3 ± 1.7 | -1.4 ± 1.2 | SS | 18.2 ± 1.8 | -1.9 ± 1.2 | SS | 18.2 ± 1.8 | -1.9 ± 1.5 | SS |
| L100 Aml5/H12.5 | 17.6 ± 2.0 | -2.6 ± 1.2 | SS | 17.2 ± 1.8 | -2.7 ± 1.5 | SS | 17.2 ± 2.1 | -2.7 ± 1.6 | SS | 17.4 ± 1.9 | -2.3 ± 1.6 | SS | 17.9 ± 2.2 | -2.2 ± 1.7 | SS | 18.3 ± 2.1 | -1.9 ± 1.4 | SS |
| L100 B5/H12.5 | 19.3 ± 2.3 | -0.9 ± 1.6 | 0.00025 | 18.6 ± 1.9 | -1.3 ± 1.6 | SS | 18.2 ± 2.3 | -1.7 ± 1.8 | SS | 17.3 ± 1.9 | -2.4 ± 1.7 | SS | 17.3 ± 2.0 | -2.7 ± 1.6 | SS | 17.5 ± 1.9 | -2.6 ± 1.6 | SS |

**Al300 =** aliskiren 300 mg; **Aml5** = amlodipine 5 mg; **B5** = bisoprolol 5 mg; **E20** = enalapril 20 mg; **H12.5** = hydrochlorothiazide 12.5 mg; **L100** = losartan 100 mg; **SS** = statistically significant ($P$ < 0.00001)

**Table S31.** *P*-values calculated using the Kolmogorov-Smirnov test for changes in right ventricular peak systolic pressure in populations ($n$ = 100) with different ACE activity receiving the same regimens (case 1: $c_{ACE}$ = 7.0 $h^{-1}$, case 2: $c_{ACE}$ = 8.9 $h^{-1}$, case 3: $c_{ACE}$ = 10.8 $h^{-1}$, case 4: $c_{ACE}$ = 42.3 $h^{-1}$, case 5: $c_{ACE}$ = 54.1 $h^{-1}$, case 6: $c_{ACE}$ = 65.9 $h^{-1}$; *P*-value for case *i vs*. case *j* is denoted $P_{ij}$)

| Regimens | $P_{12}$ | $P_{13}$ | $P_{23}$ | $P_{14}$ | $P_{15}$ | $P_{16}$ | $P_{24}$ | $P_{25}$ | $P_{26}$ | $P_{34}$ | $P_{35}$ | $P_{36}$ | $P_{45}$ | $P_{46}$ | $P_{56}$ |
|---|---|---|---|---|---|---|---|---|---|---|---|---|---|---|---|
| Al300 | 0.36672 | 0.00386 | 0.11113 | 0.58062 | 0.96707 | 0.00045 | 0.81275 | 0.58062 | 0.00007 | 0.03663 | 0.01581 | SS | 0.96707 | 0.00232 | 0.00232 |
| E20 | 0.28093 | 0.00045 | 0.07832 | SS | SS | SS | 0.00004 | SS | SS | 0.00136 | 0.00002 | 0.00079 | 0.69937 | 0.36672 | 0.07832 |
| L100 | 0.36672 | 0.00232 | 0.15454 | 0.03663 | 0.05410 | 0.21055 | 0.69937 | 0.81275 | 0.01008 | 0.69937 | 0.69937 | 0.00386 | 0.99376 | 0.00630 | 0.01008 |
| Aml5 | 0.46756 | 0.90621 | 0.69937 | 0.28093 | 0.02431 | 0.11113 | 0.15454 | 0.00630 | 0.11113 | 0.69937 | 0.15454 | 0.46756 | 0.21055 | 0.69937 | 0.81275 |
| B5 | 0.28093 | 0.81275 | 0.46756 | 0.36672 | 0.36672 | 0.00045 | 0.02431 | 0.01008 | SS | 0.03663 | 0.03663 | 0.00002 | 0.90621 | 0.01008 | 0.00232 |
| H12.5 | 0.58062 | 0.69937 | 0.28093 | 0.81275 | 0.81275 | 0.69937 | 0.28093 | 0.07832 | 0.05410 | 0.99376 | 0.46756 | 0.28093 | 0.28093 | 0.58062 | 0.99963 |
| Al300 Aml5 | 0.69937 | 0.21055 | 0.96707 | 0.46756 | 0.36672 | 0.03663 | 0.15454 | 0.01581 | 0.00045 | 0.28093 | 0.00232 | 0.00079 | 0.07832 | 0.00630 | 0.15454 |
| Al300 B5 | 0.02431 | 0.00002 | 0.01581 | SS | SS | SS | SS | SS | SS | SS | SS | SS | 0.01581 | 0.28093 | 0.21055 |
| Al300 H12.5 | 0.90621 | 0.36672 | 0.36672 | 0.05410 | 0.00013 | SS | 0.05410 | 0.00232 | SS | 0.15454 | 0.00025 | SS | 0.07832 | 0.01581 | 0.28093 |
| E20 Aml5 | 0.11113 | 0.07832 | 0.58062 | SS | SS | SS | SS | SS | SS | 0.00007 | 0.00007 | SS | 0.58062 | 0.90621 | 0.69937 |
| E20 B5 | 0.36672 | 0.00079 | 0.01008 | SS | SS | SS | SS | SS | SS | 0.00007 | SS | SS | 0.01008 | 0.01008 | 0.58062 |
| E20 H12.5 | 0.81275 | 0.11113 | 0.21055 | 0.00232 | 0.07832 | 0.02431 | 0.00630 | 0.21055 | 0.03663 | 0.15454 | 0.15454 | 0.58062 | 0.58062 | 0.58062 | 0.69937 |
| L100 Aml5 | 0.36672 | 0.07832 | 0.69937 | 0.15454 | 0.46756 | 0.46756 | 0.90621 | 0.69937 | 0.21055 | 0.81275 | 0.15454 | 0.15454 | 0.36672 | 0.07832 | 0.21055 |
| L100 B5 | 0.03663 | SS | 0.02431 | SS | SS | SS | SS | SS | SS | SS | SS | SS | 0.01581 | 0.05410 | 0.28093 |
| L100 H12.5 | 0.90621 | 0.36672 | 0.46756 | 0.15454 | 0.00136 | 0.00004 | 0.15454 | 0.00630 | 0.00045 | 0.21055 | 0.00386 | 0.00045 | 0.07832 | 0.03663 | 0.28093 |
| Al300 Aml5/B5 | 0.01581 | 0.00025 | 0.15454 | SS | SS | SS | SS | SS | SS | SS | SS | SS | 0.01581 | 0.15454 | 0.69937 |
| Al300 Aml5/H12.5 | 0.46756 | 0.46756 | 0.96707 | 0.00386 | 0.00007 | SS | 0.11113 | 0.01008 | 0.00079 | 0.07832 | 0.00630 | 0.00136 | 0.21055 | 0.11113 | 0.58062 |
| Al300 B5/H12.5 | 0.46756 | 0.00630 | 0.36672 | SS | SS | SS | 0.00386 | 0.00079 | 0.00045 | 0.07832 | 0.00386 | 0.02431 | 0.58062 | 0.90621 | 0.81275 |
| E20 Aml5/B5 | 0.05410 | 0.01581 | 0.21055 | SS | SS | SS | SS | SS | SS | SS | SS | SS | 0.01008 | 0.00630 | 0.21055 |
| E20 Aml5/H12.5 | 0.46756 | 0.28093 | 0.81275 | 0.00079 | 0.00386 | 0.00079 | 0.01581 | 0.15454 | 0.05410 | 0.15454 | 0.36672 | 0.11113 | 0.58062 | 0.58062 | 0.81275 |
| E20 B5/H12.5 | 0.28093 | 0.00630 | 0.28093 | SS | SS | SS | 0.00002 | SS | SS | 0.01581 | 0.00025 | 0.00002 | 0.36672 | 0.21055 | 0.46756 |
| L100 Aml5/B5 | 0.03663 | 0.00025 | 0.05410 | SS | SS | SS | SS | SS | SS | SS | SS | SS | 0.01008 | 0.11113 | 0.69937 |
| L100 Aml5/H12.5 | 0.69937 | 0.81275 | 0.99376 | 0.01008 | 0.00136 | 0.00045 | 0.28093 | 0.03663 | 0.01008 | 0.15454 | 0.00630 | 0.00232 | 0.28093 | 0.15454 | 0.81275 |
| L100 B5/H12.5 | 0.46756 | 0.00386 | 0.28093 | SS | SS | SS | 0.00079 | 0.00002 | SS | 0.02431 | 0.00136 | 0.00630 | 0.58062 | 0.58062 | 0.46756 |

**Al300 =** aliskiren 300 mg; **Aml5** = amlodipine 5 mg; **B5** = bisoprolol 5 mg; **E20** = enalapril 20 mg; **H12.5** = hydrochlorothiazide 12.5 mg; **L100** = losartan 100 mg; **SS** = statistically significant ($P$ < 0.00001)

**Figure S23.** Simulated change in right ventricular end-diastolic volume from baseline to week 4 (mean ± SD, $n$ = 100)

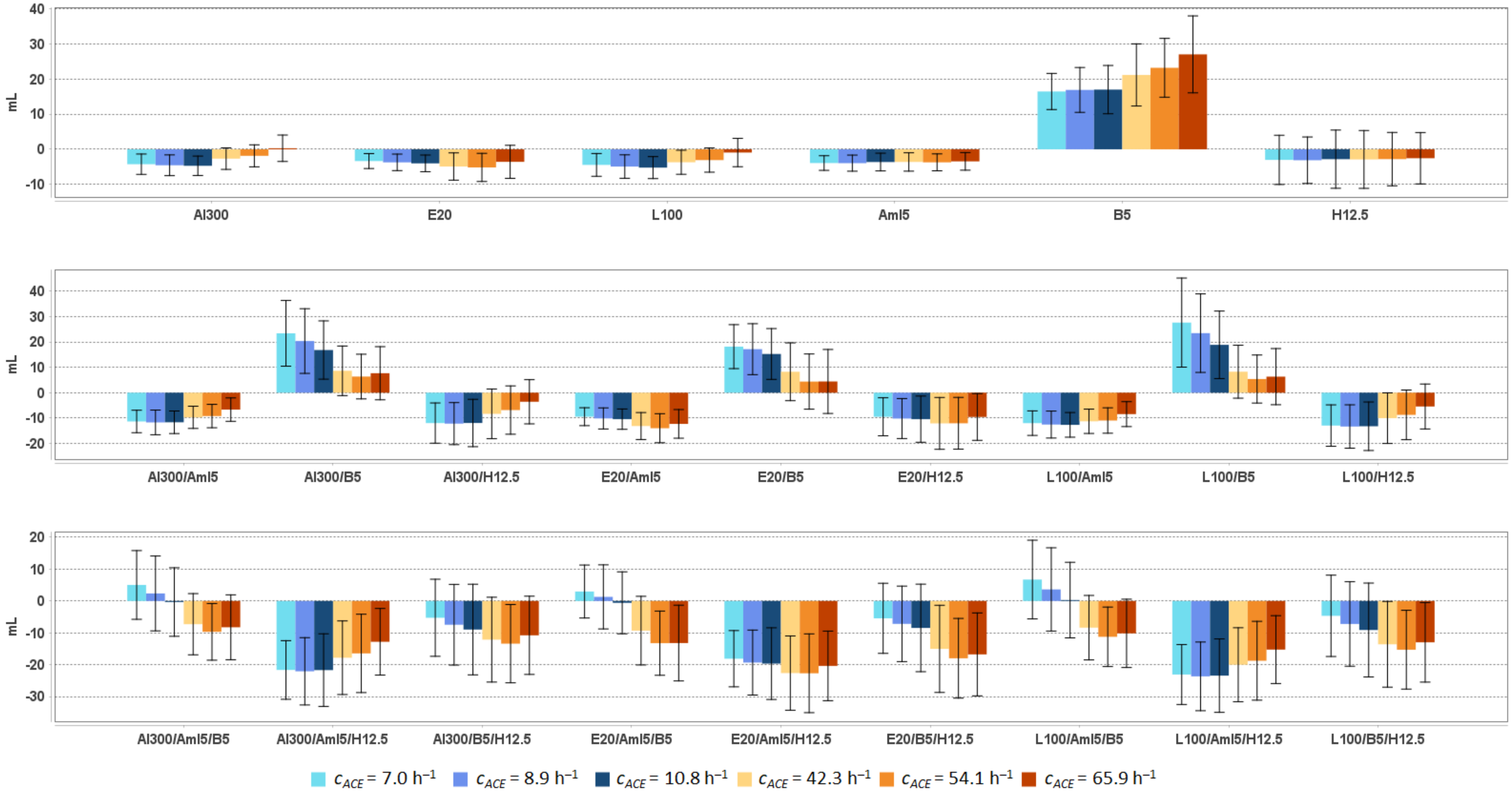


**Al300 =** aliskiren 300 mg; **Aml5** = amlodipine 5 mg; **B5** = bisoprolol 5 mg; **E20** = enalapril 20 mg; **H12.5** = hydrochlorothiazide 12.5 mg; **L100** = losartan 100 mg

**Table S32.** Simulated response of <u>right ventricular end-diastolic volume</u> to antihypertensive therapy in virtual hypertensive populations ($n$ = 100) with different ACE activity, including $P$-values (Kolmogorov-Smirnov test) for endpoint *vs*. baseline; data are presented as mean ± SD in mL

| Regimens | $c_{ACE}$ = 7.0 h$^{-1}$ | | | $c_{ACE}$ = 8.9 h$^{-1}$ | | | $c_{ACE}$ = 10.8 h$^{-1}$ | | | $c_{ACE}$ = 42.3 h$^{-1}$ | | | $c_{ACE}$ = 54.1 h$^{-1}$ | | | $c_{ACE}$ = 65.9 h$^{-1}$ | | |
|---|---|---|---|---|---|---|---|---|---|---|---|---|---|---|---|---|---|---|
| | **Value** | **Change** | ***P*** | **Value** | **Change** | ***P*** | **Value** | **Change** | ***P*** | **Value** | **Change** | ***P*** | **Value** | **Change** | ***P*** | **Value** | **Change** | ***P*** |
| Baseline | 116.8 ± 14.7 | – | – | 118.1 ± 14.0 | – | – | 117.4 ± 14.9 | – | – | 117.2 ± 14.3 | – | – | 119.2 ± 12.4 | – | – | 117.1 ± 13.1 | – | – |
| Al300 | 112.5 ± 14.1 | -4.2 ± 2.9 | 0.01008 | 113.5 ± 13.8 | -4.5 ± 3.0 | 0.00630 | 112.7 ± 14.8 | -4.7 ± 2.8 | 0.00079 | 114.5 ± 14.1 | -2.7 ± 3.1 | 0.07832 | 117.3 ± 12.6 | -1.9 ± 3.1 | 0.36672 | 117.4 ± 13.2 | 0.3 ± 3.8 | 0.81275 |
| E20 | 113.4 ± 14.2 | -3.4 ± 2.1 | 0.03663 | 114.3 ± 13.8 | -3.7 ± 2.4 | 0.01008 | 113.4 ± 14.8 | -4.0 ± 2.4 | 0.00630 | 112.2 ± 14.0 | -4.9 ± 3.9 | 0.00136 | 114.0 ± 12.5 | -5.2 ± 4.0 | 0.00007 | 113.5 ± 13.2 | -3.6 ± 4.7 | 0.01581 |
| L100 | 112.3 ± 14.1 | -4.4 ± 3.2 | 0.00630 | 113.1 ± 13.9 | -4.9 ± 3.3 | 0.00630 | 112.2 ± 14.9 | -5.2 ± 3.1 | 0.00045 | 113.5 ± 14.0 | -3.7 ± 3.4 | 0.01581 | 116.1 ± 12.5 | -3.1 ± 3.4 | 0.07832 | 116.2 ± 13.2 | -0.9 ± 4.0 | 0.46756 |
| Aml5 | 112.9 ± 14.3 | -3.9 ± 2.1 | 0.03663 | 114.1 ± 13.6 | -3.9 ± 2.3 | 0.00630 | 113.8 ± 14.6 | -3.6 ± 2.5 | 0.01008 | 113.5 ± 14.2 | -3.6 ± 2.6 | 0.01581 | 115.5 ± 12.7 | -3.7 ± 2.4 | 0.00386 | 113.6 ± 13.3 | -3.4 ± 2.5 | 0.01581 |
| B5 | 133.2 ± 17.2 | 16.5 ± 5.1 | SS | 135.0 ± 17.5 | 16.9 ± 6.4 | SS | 134.4 ± 19.0 | 17.0 ± 6.9 | SS | 138.3 ± 18.8 | 21.2 ± 8.9 | SS | 142.4 ± 16.4 | 23.2 ± 8.4 | SS | 144.1 ± 18.2 | 27.1 ± 11.0 | SS |
| H12.5 | 113.8 ± 16.1 | -3.0 ± 7.0 | 0.07832 | 114.9 ± 14.4 | -3.1 ± 6.6 | 0.03663 | 114.6 ± 17.5 | -2.8 ± 8.3 | 0.03663 | 114.3 ± 15.6 | -2.9 ± 8.2 | 0.03663 | 116.4 ± 15.2 | -2.8 ± 7.6 | 0.11113 | 114.5 ± 14.1 | -2.5 ± 7.3 | 0.11113 |
| Al300 Aml5 | 105.4 ± 13.7 | -11.3 ± 4.4 | SS | 106.3 ± 13.6 | -11.7 ± 4.9 | SS | 105.8 ± 14.5 | -11.7 ± 4.4 | SS | 107.4 ± 14.0 | -9.7 ± 4.4 | SS | 110.0 ± 13.0 | -9.2 ± 4.6 | SS | 110.4 ± 13.4 | -6.7 ± 4.6 | 0.00079 |
| Al300 B5 | 140.2 ± 20.8 | 23.4 ± 13.0 | SS | 138.4 ± 20.8 | 20.4 ± 12.8 | SS | 134.3 ± 20.7 | 16.8 ± 11.5 | SS | 125.8 ± 17.2 | 8.6 ± 9.8 | 0.00630 | 125.5 ± 14.7 | 6.4 ± 8.8 | 0.01008 | 124.8 ± 16.4 | 7.7 ± 10.5 | 0.00045 |
| Al300 H12.5 | 104.8 ± 15.4 | -12.0 ± 7.9 | SS | 105.9 ± 14.5 | -12.2 ± 8.3 | SS | 105.5 ± 17.5 | -12.0 ± 9.3 | 0.00002 | 108.8 ± 16.0 | -8.3 ± 9.8 | 0.00045 | 112.3 ± 16.4 | -6.9 ± 9.5 | 0.00136 | 113.5 ± 14.5 | -3.6 ± 8.7 | 0.03663 |
| E20 Aml5 | 107.3 ± 13.7 | -9.5 ± 3.5 | SS | 107.9 ± 13.5 | -10.2 ± 4.2 | SS | 107.0 ± 14.4 | -10.4 ± 4.0 | SS | 104.0 ± 14.0 | -13.2 ± 5.3 | SS | 105.2 ± 13.1 | -14.0 ± 5.7 | SS | 104.8 ± 13.6 | -12.3 ± 5.7 | SS |
| E20 B5 | 134.9 ± 18.1 | 18.2 ± 8.7 | SS | 135.2 ± 18.9 | 17.2 ± 10.1 | SS | 132.7 ± 19.8 | 15.3 ± 10.0 | SS | 125.4 ± 18.0 | 8.3 ± 11.4 | 0.01008 | 123.6 ± 15.7 | 4.4 ± 10.9 | 0.01581 | 121.5 ± 17.7 | 4.4 ± 12.6 | 0.01581 |
| E20 H12.5 | 107.3 ± 15.5 | -9.5 ± 7.5 | 0.00007 | 107.9 ± 14.4 | -10.2 ± 7.9 | SS | 107.0 ± 17.4 | -10.4 ± 9.1 | 0.00002 | 105.0 ± 16.1 | -12.1 ± 10.2 | SS | 107.1 ± 16.5 | -12.1 ± 10.2 | SS | 107.5 ± 14.7 | -9.6 ± 9.2 | SS |
| L100 Aml5 | 104.8 ± 13.7 | -12.0 ± 4.8 | SS | 105.5 ± 13.7 | -12.6 ± 5.3 | SS | 104.7 ± 14.5 | -12.7 ± 4.9 | SS | 105.9 ± 14.0 | -11.3 ± 4.8 | SS | 108.2 ± 13.1 | -11.0 ± 5.0 | SS | 108.6 ± 13.5 | -8.5 ± 4.9 | SS |
| L100 B5 | 144.4 ± 24.3 | 27.7 ± 17.6 | SS | 141.6 ± 22.9 | 23.5 ± 15.5 | SS | 136.3 ± 21.9 | 18.9 ± 13.3 | SS | 125.4 ± 17.5 | 8.3 ± 10.4 | 0.00630 | 124.6 ± 15.0 | 5.4 ± 9.5 | 0.01008 | 123.4 ± 16.7 | 6.3 ± 11.1 | 0.00136 |
| L100 H12.5 | 103.8 ± 15.4 | -13.0 ± 8.2 | SS | 104.7 ± 14.6 | -13.3 ± 8.6 | SS | 104.2 ± 17.5 | -13.2 ± 9.6 | SS | 107.1 ± 16.0 | -10.0 ± 10.0 | SS | 110.4 ± 16.4 | -8.7 ± 9.8 | 0.00002 | 111.6 ± 14.5 | -5.4 ± 8.9 | 0.00079 |
| Al300 Aml5/B5 | 121.8 ± 17.3 | 5.0 ± 10.8 | 0.03663 | 120.4 ± 18.2 | 2.4 ± 11.7 | 0.07832 | 117.1 ± 18.2 | -0.3 ± 10.8 | 0.21055 | 109.9 ± 16.2 | -7.3 ± 9.6 | 0.00025 | 109.5 ± 14.6 | -9.7 ± 8.9 | 0.00002 | 108.8 ± 16.0 | -8.2 ± 10.2 | 0.00004 |
| Al300 Aml5/H12.5 | 95.1 ± 15.5 | -21.6 ± 9.2 | SS | 96.0 ± 15.5 | -22.1 ± 10.6 | SS | 95.7 ± 18.1 | -21.7 ± 11.4 | SS | 99.4 ± 17.4 | -17.8 ± 11.6 | SS | 102.7 ± 18.2 | -16.4 ± 12.3 | SS | 104.3 ± 15.8 | -12.8 ± 10.5 | SS |
| Al300 B5/H12.5 | 111.5 ± 18.0 | -5.3 ± 12.1 | 0.00136 | 110.6 ± 17.6 | -7.5 ± 12.7 | 0.00136 | 108.5 ± 21.0 | -9.0 ± 14.2 | 0.00136 | 105.0 ± 18.1 | -12.1 ± 13.3 | SS | 105.8 ± 17.5 | -13.4 ± 12.3 | SS | 106.3 ± 16.7 | -10.8 ± 12.3 | SS |
| E20 Aml5/B5 | 119.7 ± 16.1 | 3.0 ± 8.3 | 0.11113 | 119.3 ± 17.1 | 1.3 ± 10.1 | 0.15454 | 116.8 ± 17.7 | -0.6 ± 9.7 | 0.28093 | 107.9 ± 16.6 | -9.3 ± 10.8 | SS | 105.9 ± 15.0 | -13.2 ± 10.1 | SS | 103.9 ± 17.0 | -13.2 ± 11.9 | SS |
| E20 Aml5/H12.5 | 98.7 ± 15.7 | -18.1 ± 8.8 | SS | 98.7 ± 15.3 | -19.3 ± 10.2 | SS | 97.8 ± 18.1 | -19.7 ± 11.2 | SS | 94.5 ± 17.2 | -22.6 ± 11.7 | SS | 96.5 ± 17.9 | -22.7 ± 12.4 | SS | 96.7 ± 16.1 | -20.4 ± 10.9 | SS |
| E20 B5/H12.5 | 111.3 ± 17.4 | -5.4 ± 11.0 | 0.00079 | 110.9 ± 17.0 | -7.2 ± 11.8 | 0.00232 | 109.0 ± 20.7 | -8.5 ± 13.7 | 0.00136 | 102.1 ± 18.2 | -15.0 ± 13.7 | SS | 101.2 ± 17.3 | -18.0 ± 12.5 | SS | 100.3 ± 17.1 | -16.8 ± 13.0 | SS |
| L100 Aml5/B5 | 123.5 ± 18.2 | 6.7 ± 12.3 | 0.01008 | 121.7 ± 19.0 | 3.6 ± 13.1 | 0.05410 | 117.7 ± 18.8 | 0.3 ± 11.8 | 0.15454 | 108.8 ± 16.4 | -8.4 ± 10.1 | 0.00002 | 107.9 ± 14.7 | -11.2 ± 9.3 | SS | 106.9 ± 16.3 | -10.2 ± 10.7 | SS |
| L100 Aml5/H12.5 | 93.7 ± 15.5 | -23.1 ± 9.4 | SS | 94.4 ± 15.6 | -23.6 ± 10.8 | SS | 94.0 ± 18.1 | -23.4 ± 11.5 | SS | 97.2 ± 17.3 | -20.0 ± 11.6 | SS | 100.4 ± 18.1 | -18.8 ± 12.4 | SS | 101.8 ± 16.0 | -15.3 ± 10.7 | SS |
| L100 B5/H12.5 | 112.1 ± 18.4 | -4.7 ± 12.8 | 0.00136 | 110.8 ± 18.0 | -7.2 ± 13.3 | 0.00232 | 108.3 ± 21.3 | -9.1 ± 14.7 | 0.00079 | 103.6 ± 18.1 | -13.6 ± 13.5 | SS | 103.9 ± 17.4 | -15.3 ± 12.4 | SS | 104.1 ± 16.8 | -13.0 ± 12.5 | SS |

**Al300 =** aliskiren 300 mg; **Aml5** = amlodipine 5 mg; **B5** = bisoprolol 5 mg; **E20** = enalapril 20 mg; **H12.5** = hydrochlorothiazide 12.5 mg; **L100** = losartan 100 mg; **SS** = statistically significant ($P$ < 0.00001)

**Table S33.** *P*-values calculated using the Kolmogorov-Smirnov test for changes in <u>right ventricular end-diastolic volume</u> in populations ($n$ = 100) with different ACE activity receiving the same regimens (case 1: $c_{ACE}$ = 7.0 h$^{-1}$, case 2: $c_{ACE}$ = 8.9 h$^{-1}$, case 3: $c_{ACE}$ = 10.8 h$^{-1}$, case 4: $c_{ACE}$ = 42.3 h$^{-1}$, case 5: $c_{ACE}$ = 54.1 h$^{-1}$, case 6: $c_{ACE}$ = 65.9 h$^{-1}$; *P*-value for case *i vs*. case *j* is denoted $P_{ij}$)

| Regimens | $P_{12}$ | $P_{13}$ | $P_{23}$ | $P_{14}$ | $P_{15}$ | $P_{16}$ | $P_{24}$ | $P_{25}$ | $P_{26}$ | $P_{34}$ | $P_{35}$ | $P_{36}$ | $P_{45}$ | $P_{46}$ | $P_{56}$ |
|---|---|---|---|---|---|---|---|---|---|---|---|---|---|---|---|
| Al300 | 0.36672 | 0.02431 | 0.36672 | 0.01581 | 0.00002 | SS | 0.00232 | SS | SS | 0.00007 | SS | SS | 0.05410 | SS | 0.00630 |
| E20 | 0.28093 | 0.00630 | 0.15454 | 0.00013 | 0.00025 | 0.00630 | 0.00136 | 0.00136 | 0.02431 | 0.00630 | 0.00136 | 0.02431 | 0.36672 | 0.21055 | 0.07832 |
| L100 | 0.28093 | 0.01008 | 0.21055 | 0.21055 | 0.03663 | SS | 0.15454 | 0.01581 | SS | 0.00630 | SS | SS | 0.21055 | 0.00025 | 0.00630 |
| Aml5 | 0.28093 | 0.28093 | 0.81275 | 0.58062 | 0.21055 | 0.15454 | 0.81275 | 0.90621 | 0.58062 | 0.81275 | 0.99376 | 0.36672 | 0.90621 | 0.81275 | 0.69937 |
| B5 | 0.90621 | 0.69937 | 0.96707 | 0.00079 | SS | SS | 0.01581 | SS | SS | 0.00232 | SS | SS | 0.05410 | 0.00045 | 0.02431 |
| H12.5 | 0.69937 | 0.58062 | 0.46756 | 0.69937 | 0.69937 | 0.81275 | 0.36672 | 0.15454 | 0.21055 | 0.90621 | 0.36672 | 0.58062 | 0.81275 | 0.81275 | 0.96707 |
| Al300<br>Aml5 | 0.90621 | 0.81275 | 0.90621 | 0.21055 | 0.00232 | SS | 0.02431 | 0.00007 | SS | 0.02431 | 0.00045 | SS | 0.15454 | 0.00013 | 0.01581 |
| Al300<br>B5 | 0.15454 | 0.00386 | 0.15454 | SS | SS | SS | SS | SS | SS | 0.00007 | SS | 0.00004 | 0.36672 | 0.69937 | 0.58062 |
| Al300<br>H12.5 | 0.90621 | 0.58062 | 0.46756 | 0.02431 | 0.00007 | SS | 0.01008 | 0.00002 | SS | 0.05410 | 0.00045 | SS | 0.28093 | 0.00232 | 0.11113 |
| E20<br>Aml5 | 0.58062 | 0.15454 | 0.46756 | SS | SS | 0.00004 | SS | 0.00002 | 0.00013 | 0.00136 | 0.00013 | 0.01581 | 0.69937 | 0.69937 | 0.21055 |
| E20<br>B5 | 0.46756 | 0.02431 | 0.36672 | SS | SS | SS | SS | SS | SS | 0.00013 | SS | SS | 0.11113 | 0.11113 | 0.90621 |
| E20<br>H12.5 | 0.69937 | 0.28093 | 0.36672 | 0.05410 | 0.07832 | 0.46756 | 0.15454 | 0.21055 | 0.81275 | 0.69937 | 0.46756 | 0.28093 | 0.90621 | 0.21055 | 0.46756 |
| L100<br>Aml5 | 0.36672 | 0.46756 | 0.69937 | 0.69937 | 0.21055 | 0.00136 | 0.28093 | 0.01008 | SS | 0.11113 | 0.05410 | SS | 0.58062 | 0.00232 | 0.05410 |
| L100<br>B5 | 0.21055 | 0.00232 | 0.11113 | SS | SS | SS | SS | SS | SS | 0.00002 | SS | SS | 0.21055 | 0.36672 | 0.69937 |
| L100<br>H12.5 | 0.69937 | 0.81275 | 0.69937 | 0.05410 | 0.00025 | SS | 0.02431 | 0.00045 | SS | 0.11113 | 0.00386 | SS | 0.28093 | 0.00386 | 0.15454 |
| Al300<br>Aml5/B5 | 0.07832 | 0.01008 | 0.21055 | SS | SS | SS | SS | SS | SS | 0.00079 | SS | 0.00002 | 0.11113 | 0.58062 | 0.69937 |
| Al300<br>Aml5/H12.5 | 0.58062 | 0.58062 | 0.96707 | 0.01008 | 0.00025 | SS | 0.01008 | 0.00025 | SS | 0.05410 | 0.00232 | SS | 0.28093 | 0.05410 | 0.21055 |
| Al300<br>B5/H12.5 | 0.58062 | 0.01581 | 0.21055 | 0.00630 | 0.00232 | 0.00232 | 0.11113 | 0.02431 | 0.11113 | 0.28093 | 0.05410 | 0.28093 | 0.81275 | 0.81275 | 0.58062 |
| E20<br>Aml5/B5 | 0.15454 | 0.01581 | 0.36672 | SS | SS | SS | SS | SS | SS | SS | SS | SS | 0.05410 | 0.01581 | 0.36672 |
| E20<br>Aml5/H12.5 | 0.28093 | 0.21055 | 0.96707 | 0.00630 | 0.00630 | 0.05410 | 0.05410 | 0.11113 | 0.58062 | 0.07832 | 0.28093 | 0.69937 | 0.69937 | 0.46756 | 0.58062 |
| E20<br>B5/H12.5 | 0.28093 | 0.00630 | 0.36672 | SS | SS | SS | 0.00045 | 0.00002 | SS | 0.01008 | 0.00025 | 0.00232 | 0.58062 | 0.36672 | 0.69937 |
| L100<br>Aml5/B5 | 0.07832 | 0.00386 | 0.21055 | SS | SS | SS | SS | SS | SS | 0.00007 | SS | SS | 0.07832 | 0.36672 | 0.81275 |
| L100<br>Aml5/H12.5 | 0.58062 | 0.46756 | 0.99376 | 0.03663 | 0.00079 | SS | 0.02431 | 0.00079 | SS | 0.11113 | 0.00630 | 0.00007 | 0.36672 | 0.03663 | 0.36672 |
| L100<br>B5/H12.5 | 0.15454 | 0.01581 | 0.28093 | 0.00013 | 0.00004 | 0.00013 | 0.01581 | 0.00232 | 0.01008 | 0.07832 | 0.01008 | 0.05410 | 0.69937 | 0.96707 | 0.46756 |

**Al300 =** aliskiren 300 mg; **Aml5** = amlodipine 5 mg; **B5** = bisoprolol 5 mg; **E20** = enalapril 20 mg; **H12.5** = hydrochlorothiazide 12.5 mg; **L100** = losartan 100 mg; **SS** = statistically significant ($P < 0.00001$)

**Figure S24.** Simulated change in <u>right ventricular end-systolic volume</u> from baseline to week 4 (mean ± SD, $n$ = 100)

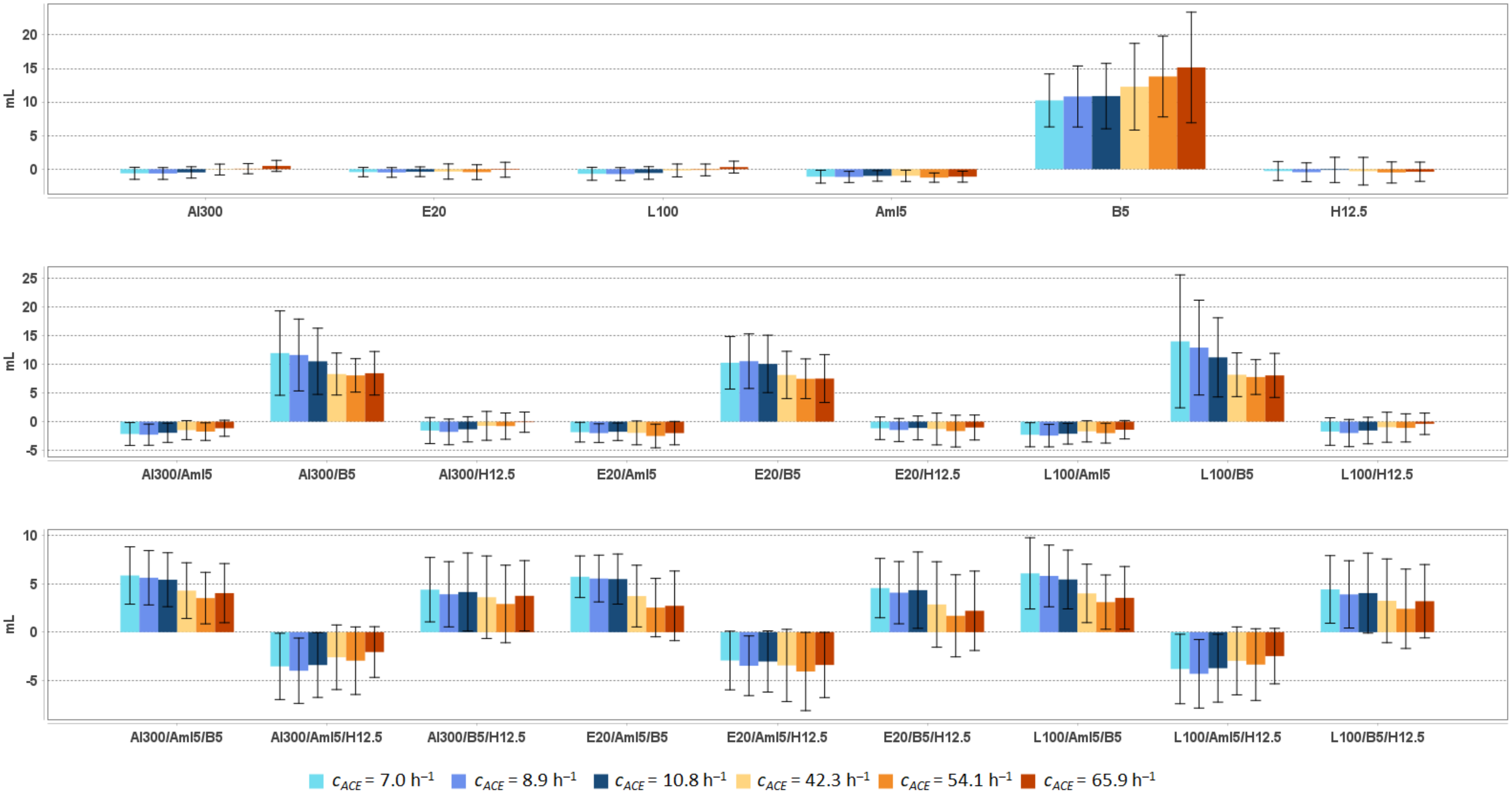


**Al300 =** aliskiren 300 mg; **Aml5** = amlodipine 5 mg; **B5** = bisoprolol 5 mg; **E20** = enalapril 20 mg; **H12.5** = hydrochlorothiazide 12.5 mg; **L100** = losartan 100 mg

**Table S34.** Simulated response of right ventricular end-systolic volume to antihypertensive therapy in virtual hypertensive populations ($n$ = 100) with different ACE activity, including $P$-values (Kolmogorov-Smirnov test) for endpoint *vs.* baseline; data are presented as mean ± SD in mL

| Regimens | $c_{ACE}$ = 7.0 h$^{-1}$ | | | $c_{ACE}$ = 8.9 h$^{-1}$ | | | $c_{ACE}$ = 10.8 h$^{-1}$ | | | $c_{ACE}$ = 42.3 h$^{-1}$ | | | $c_{ACE}$ = 54.1 h$^{-1}$ | | | $c_{ACE}$ = 65.9 h$^{-1}$ | | |
|---|---|---|---|---|---|---|---|---|---|---|---|---|---|---|---|---|---|---|
| | **Value** | **Change** | ***P*** | **Value** | **Change** | ***P*** | **Value** | **Change** | ***P*** | **Value** | **Change** | ***P*** | **Value** | **Change** | ***P*** | **Value** | **Change** | ***P*** |
| Baseline | 43.5 ± 12.2 | – | – | 45.5 ± 10.5 | – | – | 43.9 ± 11.4 | – | – | 44.4 ± 11.5 | – | – | 45.7 ± 9.7 | – | – | 45.2 ± 11.4 | – | – |
| Al300 | 42.9 ± 11.8 | -0.6 ± 0.9 | 0.96707 | 44.9 ± 10.1 | -0.6 ± 0.9 | 0.96707 | 43.4 ± 11.0 | -0.5 ± 0.8 | 0.99376 | 44.3 ± 11.3 | -0.1 ± 0.8 | 1.00000 | 45.8 ± 9.7 | 0.1 ± 0.8 | 0.90621 | 45.7 ± 11.6 | 0.5 ± 0.8 | 0.99963 |
| E20 | 43.1 ± 11.9 | -0.4 ± 0.7 | 0.99376 | 45.0 ± 10.2 | -0.5 ± 0.7 | 0.96707 | 43.5 ± 11.1 | -0.4 ± 0.7 | 0.99963 | 44.0 ± 11.1 | -0.4 ± 1.1 | 0.99376 | 45.3 ± 9.6 | -0.4 ± 1.1 | 0.96707 | 45.1 ± 11.3 | -0.1 ± 1.1 | 0.99963 |
| L100 | 42.8 ± 11.8 | -0.7 ± 1.0 | 0.96707 | 44.8 ± 10.1 | -0.7 ± 1.0 | 0.90621 | 43.3 ± 11.0 | -0.6 ± 0.9 | 0.99376 | 44.2 ± 11.2 | -0.2 ± 1.0 | 0.99963 | 45.6 ± 9.6 | -0.1 ± 0.9 | 0.99376 | 45.5 ± 11.5 | 0.3 ± 0.9 | 0.99963 |
| Aml5 | 42.4 ± 11.7 | -1.1 ± 1.0 | 0.69937 | 44.4 ± 10.1 | -1.2 ± 0.8 | 0.81275 | 42.9 ± 11.0 | -1.0 ± 0.8 | 0.96707 | 43.4 ± 11.0 | -1.0 ± 0.8 | 0.90621 | 44.5 ± 9.7 | -1.3 ± 0.7 | 0.81275 | 44.1 ± 11.2 | -1.1 ± 0.8 | 0.81275 |
| B5 | 53.8 ± 14.8 | 10.2 ± 3.9 | SS | 56.4 ± 13.2 | 10.8 ± 4.5 | SS | 54.8 ± 14.5 | 10.9 ± 4.9 | SS | 56.6 ± 15.2 | 12.3 ± 6.5 | SS | 59.6 ± 11.9 | 13.8 ± 6.0 | SS | 60.4 ± 16.0 | 15.2 ± 8.2 | SS |
| H12.5 | 43.3 ± 12.0 | -0.3 ± 1.4 | 0.99963 | 45.1 ± 10.2 | -0.5 ± 1.4 | 0.99376 | 43.8 ± 11.6 | -0.1 ± 1.9 | 0.99376 | 44.1 ± 11.1 | -0.3 ± 2.1 | 0.90621 | 45.2 ± 9.8 | -0.5 ± 1.6 | 0.99376 | 44.9 ± 11.3 | -0.4 ± 1.4 | 0.90621 |
| Al300 Aml5 | 41.4 ± 11.2 | -2.2 ± 2.0 | 0.36672 | 43.3 ± 9.6 | -2.3 ± 1.9 | 0.28093 | 42.0 ± 10.5 | -2.0 ± 1.7 | 0.46756 | 42.9 ± 10.7 | -1.5 ± 1.7 | 0.36672 | 44.0 ± 9.7 | -1.7 ± 1.5 | 0.36672 | 44.1 ± 11.1 | -1.2 ± 1.4 | 0.69937 |
| Al300 B5 | 55.5 ± 16.4 | 12.0 ± 7.4 | SS | 57.2 ± 14.1 | 11.7 ± 6.3 | SS | 54.5 ± 14.6 | 10.6 ± 5.8 | SS | 52.7 ± 12.7 | 8.3 ± 3.7 | 0.00013 | 53.8 ± 9.9 | 8.1 ± 2.9 | SS | 53.7 ± 12.7 | 8.5 ± 3.8 | 0.00013 |
| Al300 H12.5 | 42.0 ± 11.4 | -1.6 ± 2.3 | 0.46756 | 43.7 ± 9.7 | -1.8 ± 2.2 | 0.58062 | 42.6 ± 11.0 | -1.3 ± 2.2 | 0.81275 | 43.6 ± 10.8 | -0.7 ± 2.5 | 0.81275 | 45.0 ± 10.0 | -0.8 ± 2.3 | 0.81275 | 45.1 ± 11.3 | -0.1 ± 1.8 | 0.90621 |
| E20 Aml5 | 41.7 ± 11.3 | -1.8 ± 1.7 | 0.36672 | 43.5 ± 9.7 | -2.0 ± 1.6 | 0.36672 | 42.2 ± 10.6 | -1.8 ± 1.5 | 0.46756 | 42.4 ± 10.5 | -2.0 ± 2.1 | 0.15454 | 43.2 ± 9.6 | -2.5 ± 2.1 | 0.28093 | 43.2 ± 10.9 | -2.0 ± 2.0 | 0.28093 |
| E20 B5 | 53.8 ± 14.9 | 10.3 ± 4.6 | SS | 56.1 ± 13.1 | 10.6 ± 4.8 | SS | 54.0 ± 14.1 | 10.1 ± 5.0 | SS | 52.5 ± 12.8 | 8.2 ± 4.1 | 0.00079 | 53.2 ± 9.8 | 7.5 ± 3.5 | SS | 52.8 ± 12.6 | 7.5 ± 4.2 | 0.00136 |
| E20 H12.5 | 42.4 ± 11.6 | -1.2 ± 2.0 | 0.69937 | 44.1 ± 9.8 | -1.5 ± 2.0 | 0.58062 | 42.8 ± 11.1 | -1.1 ± 2.1 | 0.90621 | 43.1 ± 10.6 | -1.3 ± 2.8 | 0.36672 | 44.1 ± 10.0 | -1.7 ± 2.8 | 0.46756 | 44.2 ± 11.1 | -1.0 ± 2.2 | 0.69937 |
| L100 Aml5 | 41.2 ± 11.1 | -2.3 ± 2.1 | 0.28093 | 43.1 ± 9.6 | -2.4 ± 2.0 | 0.21055 | 41.8 ± 10.5 | -2.1 ± 1.8 | 0.46756 | 42.7 ± 10.6 | -1.7 ± 1.9 | 0.15454 | 43.7 ± 9.7 | -2.0 ± 1.7 | 0.36672 | 43.8 ± 11.0 | -1.4 ± 1.6 | 0.58062 |
| L100 B5 | 57.6 ± 19.2 | 14.0 ± 11.6 | SS | 58.5 ± 15.4 | 12.9 ± 8.3 | SS | 55.2 ± 15.3 | 11.2 ± 6.9 | SS | 52.6 ± 12.7 | 8.2 ± 3.8 | 0.00079 | 53.5 ± 9.8 | 7.8 ± 3.0 | SS | 53.3 ± 12.6 | 8.1 ± 3.9 | 0.00013 |
| L100 H12.5 | 41.8 ± 11.4 | -1.7 ± 2.4 | 0.36672 | 43.5 ± 9.6 | -2.0 ± 2.4 | 0.36672 | 42.4 ± 11.0 | -1.6 ± 2.3 | 0.69937 | 43.4 ± 10.7 | -1.0 ± 2.6 | 0.69937 | 44.6 ± 10.0 | -1.1 ± 2.5 | 0.69937 | 44.9 ± 11.2 | -0.4 ± 1.9 | 0.81275 |
| Al300 Aml5/B5 | 49.4 ± 12.7 | 5.9 ± 3.0 | 0.00079 | 51.2 ± 10.7 | 5.6 ± 2.8 | 0.01581 | 49.3 ± 11.7 | 5.4 ± 2.8 | 0.00136 | 48.7 ± 11.1 | 4.3 ± 2.9 | 0.05410 | 49.3 ± 9.6 | 3.5 ± 2.7 | 0.01008 | 49.3 ± 11.5 | 4.0 ± 3.1 | 0.11113 |
| Al300 Aml5/H12.5 | 40.0 ± 10.9 | -3.5 ± 3.4 | 0.03663 | 41.5 ± 9.3 | -4.0 ± 3.4 | 0.02431 | 40.5 ± 10.7 | -3.4 ± 3.3 | 0.07832 | 41.8 ± 10.6 | -2.6 ± 3.3 | 0.05410 | 42.8 ± 10.4 | -3.0 ± 3.5 | 0.28093 | 43.2 ± 10.9 | -2.1 ± 2.6 | 0.28093 |
| Al300 B5/H12.5 | 47.9 ± 12.1 | 4.4 ± 3.3 | 0.02431 | 49.5 ± 10.1 | 3.9 ± 3.4 | 0.05410 | 48.1 ± 12.0 | 4.1 ± 4.0 | 0.07832 | 48.0 ± 11.1 | 3.6 ± 4.3 | 0.15454 | 48.7 ± 10.3 | 2.9 ± 4.0 | 0.00630 | 49.0 ± 11.4 | 3.8 ± 3.6 | 0.07832 |
| E20 Aml5/B5 | 49.3 ± 12.4 | 5.7 ± 2.2 | 0.00136 | 51.1 ± 10.6 | 5.5 ± 2.4 | 0.01581 | 49.4 ± 11.6 | 5.5 ± 2.6 | 0.00136 | 48.1 ± 11.0 | 3.7 ± 3.2 | 0.07832 | 48.3 ± 9.6 | 2.5 ± 3.0 | 0.05410 | 48.0 ± 11.5 | 2.7 ± 3.6 | 0.36672 |
| E20 Aml5/H12.5 | 40.6 ± 11.1 | -2.9 ± 3.0 | 0.21055 | 42.1 ± 9.4 | -3.5 ± 3.1 | 0.07832 | 40.9 ± 10.7 | -3.0 ± 3.2 | 0.11113 | 40.9 ± 10.4 | -3.4 ± 3.7 | 0.02431 | 41.7 ± 10.4 | -4.1 ± 4.0 | 0.15454 | 41.8 ± 10.8 | -3.4 ± 3.4 | 0.07832 |
| E20 B5/H12.5 | 48.1 ± 12.1 | 4.6 ± 3.1 | 0.01008 | 49.6 ± 10.1 | 4.1 ± 3.2 | 0.03663 | 48.3 ± 12.0 | 4.3 ± 3.9 | 0.05410 | 47.2 ± 11.0 | 2.9 ± 4.4 | 0.36672 | 47.4 ± 10.2 | 1.7 ± 4.2 | 0.05410 | 47.4 ± 11.4 | 2.2 ± 4.1 | 0.46756 |
| L100 Aml5/B5 | 49.6 ± 13.0 | 6.1 ± 3.7 | 0.00136 | 51.3 ± 10.9 | 5.8 ± 3.2 | 0.01581 | 49.4 ± 11.7 | 5.4 ± 3.0 | 0.00136 | 48.4 ± 11.1 | 4.0 ± 3.0 | 0.05410 | 48.8 ± 9.6 | 3.1 ± 2.8 | 0.02431 | 48.8 ± 11.5 | 3.5 ± 3.2 | 0.15454 |
| L100 Aml5/H12.5 | 39.7 ± 10.9 | -3.8 ± 3.6 | 0.01581 | 41.2 ± 9.3 | -4.3 ± 3.5 | 0.01581 | 40.2 ± 10.6 | -3.7 ± 3.5 | 0.07832 | 41.4 ± 10.5 | -3.0 ± 3.5 | 0.03663 | 42.4 ± 10.4 | -3.4 ± 3.7 | 0.28093 | 42.8 ± 10.9 | -2.5 ± 2.9 | 0.21055 |
| L100 B5/H12.5 | 48.0 ± 12.2 | 4.4 ± 3.5 | 0.01581 | 49.4 ± 10.1 | 3.9 ± 3.5 | 0.05410 | 48.0 ± 12.0 | 4.0 ± 4.1 | 0.11113 | 47.6 ± 11.0 | 3.2 ± 4.3 | 0.28093 | 48.2 ± 10.3 | 2.4 ± 4.1 | 0.01581 | 48.4 ± 11.4 | 3.2 ± 3.8 | 0.21055 |

**Al300 =** aliskiren 300 mg; **Aml5** = amlodipine 5 mg; **B5** = bisoprolol 5 mg; **E20** = enalapril 20 mg; **H12.5** = hydrochlorothiazide 12.5 mg; **L100** = losartan 100 mg; **SS** = statistically significant ($P$ < 0.00001)

**Table S35.** *P*-values calculated using the Kolmogorov-Smirnov test for changes in right ventricular end-systolic volume in populations ($n$ = 100) with different ACE activity receiving the same regimens (case 1: $c_{ACE}$ = 7.0 h$^{-1}$, case 2: $c_{ACE}$ = 8.9 h$^{-1}$, case 3: $c_{ACE}$ = 10.8 h$^{-1}$, case 4: $c_{ACE}$ = 42.3 h$^{-1}$, case 5: $c_{ACE}$ = 54.1 h$^{-1}$, case 6: $c_{ACE}$ = 65.9 h$^{-1}$; *P*-value for case *i vs*. case *j* is denoted $P_{ij}$)

| Regimens | $P_{12}$ | $P_{13}$ | $P_{23}$ | $P_{14}$ | $P_{15}$ | $P_{16}$ | $P_{24}$ | $P_{25}$ | $P_{26}$ | $P_{34}$ | $P_{35}$ | $P_{36}$ | $P_{45}$ | $P_{46}$ | $P_{56}$ |
|---|---|---|---|---|---|---|---|---|---|---|---|---|---|---|---|
| Al300 | 0.90621 | 0.36672 | 0.46756 | SS | SS | SS | SS | SS | SS | SS | SS | SS | 0.15454 | SS | 0.01581 |
| E20 | 0.81275 | 0.28093 | 0.46756 | 0.00002 | 0.00045 | SS | 0.00002 | 0.00045 | SS | 0.00232 | 0.02431 | 0.00002 | 0.46756 | 0.01581 | 0.02431 |
| L100 | 0.69937 | 0.36672 | 0.36672 | SS | SS | SS | SS | SS | SS | 0.00002 | 0.00007 | SS | 0.58062 | 0.00079 | 0.02431 |
| Aml5 | 0.96707 | 0.96707 | 0.81275 | 0.90621 | 0.07832 | 0.99376 | 0.58062 | 0.28093 | 0.99376 | 0.96707 | 0.11113 | 0.99376 | 0.01581 | 0.81275 | 0.11113 |
| B5 | 0.58062 | 0.69937 | 0.81275 | 0.00630 | 0.00013 | 0.00004 | 0.01581 | 0.00386 | 0.00013 | 0.11113 | 0.00079 | 0.00013 | 0.07832 | 0.05410 | 0.69937 |
| H12.5 | 0.28093 | 0.69937 | 0.21055 | 0.81275 | 0.28093 | 0.90621 | 0.15454 | 0.21055 | 0.21055 | 0.81275 | 0.90621 | 0.90621 | 0.81275 | 0.99963 | 0.46756 |
| Al300 Aml5 | 0.81275 | 0.99376 | 0.46756 | 0.00232 | 0.15454 | 0.00004 | 0.01581 | 0.07832 | 0.00002 | 0.01581 | 0.11113 | 0.00004 | 0.46756 | 0.11113 | 0.00630 |
| Al300 B5 | 0.99376 | 0.15454 | 0.21055 | 0.00025 | 0.00013 | 0.00079 | 0.00013 | 0.00013 | 0.00025 | 0.00386 | 0.02431 | 0.11113 | 0.46756 | 0.58062 | 0.90621 |
| Al300 H12.5 | 0.69937 | 0.28093 | 0.46756 | 0.00045 | 0.00079 | SS | 0.00013 | 0.00007 | SS | 0.01581 | 0.01581 | 0.00004 | 0.58062 | 0.15454 | 0.05410 |
| E20 Aml5 | 0.58062 | 0.90621 | 0.69937 | 0.15454 | 0.05410 | 0.21055 | 0.46756 | 0.15454 | 0.46756 | 0.15454 | 0.07832 | 0.21055 | 0.07832 | 0.96707 | 0.15454 |
| E20 B5 | 0.99963 | 0.46756 | 0.36672 | 0.00013 | SS | 0.00002 | 0.00045 | SS | 0.00002 | 0.00136 | 0.00045 | 0.00232 | 0.58062 | 0.28093 | 0.99376 |
| E20 H12.5 | 0.46756 | 0.69937 | 0.28093 | 0.21055 | 0.11113 | 0.11113 | 0.02431 | 0.15454 | 0.00630 | 0.81275 | 0.46756 | 0.58062 | 0.69937 | 0.69937 | 0.28093 |
| L100 Aml5 | 0.69937 | 0.96707 | 0.46756 | 0.00630 | 0.36672 | 0.00079 | 0.03663 | 0.28093 | 0.00079 | 0.03663 | 0.15454 | 0.00136 | 0.36672 | 0.28093 | 0.02431 |
| L100 B5 | 0.99376 | 0.11113 | 0.11113 | SS | SS | SS | SS | SS | 0.00002 | 0.00232 | 0.00232 | 0.02431 | 0.69937 | 0.69937 | 0.96707 |
| L100 H12.5 | 0.46756 | 0.46756 | 0.46756 | 0.00045 | 0.00232 | SS | 0.00025 | 0.00045 | SS | 0.01008 | 0.03663 | 0.00045 | 0.96707 | 0.21055 | 0.07832 |
| Al300 Aml5/B5 | 0.90621 | 0.58062 | 0.69937 | 0.01581 | SS | 0.00136 | 0.07832 | 0.00013 | 0.00630 | 0.11113 | 0.00045 | 0.02431 | 0.05410 | 0.58062 | 0.36672 |
| Al300 Aml5/H12.5 | 0.46756 | 0.90621 | 0.69937 | 0.01008 | 0.03663 | 0.00013 | 0.01008 | 0.03663 | 0.00025 | 0.11113 | 0.36672 | 0.00630 | 0.58062 | 0.28093 | 0.28093 |
| Al300 B5/H12.5 | 0.21055 | 0.46756 | 0.99376 | 0.21055 | 0.03663 | 0.28093 | 0.69937 | 0.15454 | 0.90621 | 0.69937 | 0.36672 | 0.81275 | 0.36672 | 0.90621 | 0.58062 |
| E20 Aml5/B5 | 0.69937 | 0.36672 | 0.81275 | 0.00013 | SS | SS | 0.00386 | SS | SS | 0.00232 | SS | SS | 0.00630 | 0.02431 | 0.69937 |
| E20 Aml5/H12.5 | 0.21055 | 0.36672 | 0.58062 | 0.15454 | 0.03663 | 0.21055 | 0.36672 | 0.58062 | 0.81275 | 0.46756 | 0.21055 | 0.90621 | 0.28093 | 0.81275 | 0.90621 |
| E20 B5/H12.5 | 0.28093 | 0.36672 | 0.90621 | 0.01008 | SS | 0.00002 | 0.28093 | 0.00025 | 0.00630 | 0.28093 | 0.00079 | 0.00630 | 0.11113 | 0.46756 | 0.90621 |
| L100 Aml5/B5 | 0.99376 | 0.58062 | 0.46756 | 0.02431 | SS | 0.00025 | 0.02431 | SS | 0.00045 | 0.07832 | 0.00004 | 0.00232 | 0.03663 | 0.28093 | 0.36672 |
| L100 Aml5/H12.5 | 0.36672 | 0.81275 | 0.58062 | 0.03663 | 0.15454 | 0.00232 | 0.01581 | 0.05410 | 0.00136 | 0.21055 | 0.58062 | 0.01581 | 0.46756 | 0.36672 | 0.36672 |
| L100 B5/H12.5 | 0.36672 | 0.46756 | 0.99376 | 0.15454 | 0.00136 | 0.02431 | 0.46756 | 0.07832 | 0.36672 | 0.69937 | 0.02431 | 0.36672 | 0.21055 | 0.96707 | 0.58062 |

**Al300 =** aliskiren 300 mg; **Aml5** = amlodipine 5 mg; **B5** = bisoprolol 5 mg; **E20** = enalapril 20 mg; **H12.5** = hydrochlorothiazide 12.5 mg; **L100** = losartan 100 mg; **SS** = statistically significant ($P$ < 0.00001)

**Figure S25.** Simulated change in glomerular filtration rate from baseline to week 4 (mean ± SD, $n$ = 100)

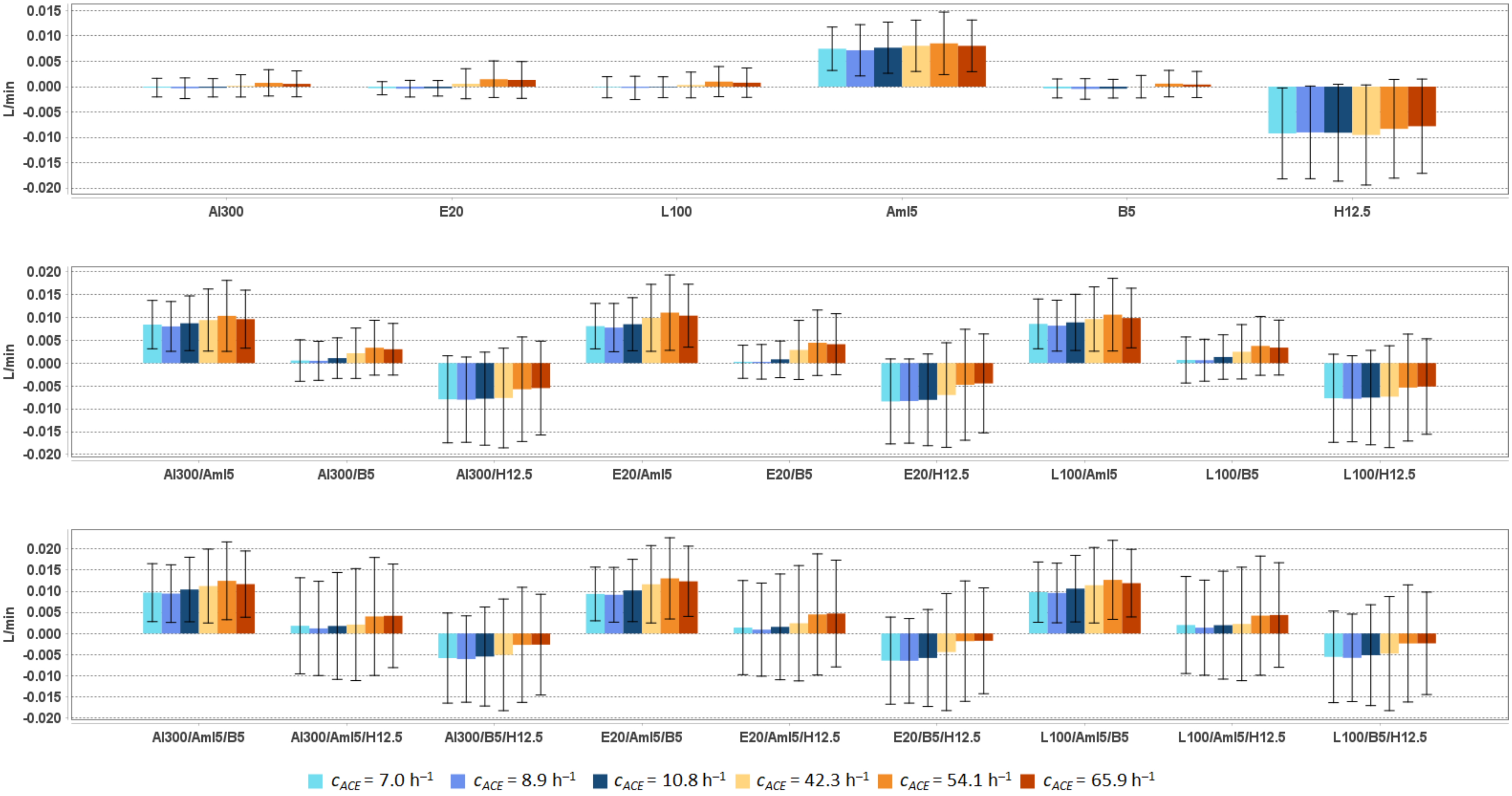


**Al300 =** aliskiren 300 mg; **Aml5** = amlodipine 5 mg; **B5** = bisoprolol 5 mg; **E20** = enalapril 20 mg; **H12.5** = hydrochlorothiazide 12.5 mg; **L100** = losartan 100 mg

**Table S36.** Simulated response of glomerular filtration rate to antihypertensive therapy in virtual hypertensive populations ($n$ = 100) with different ACE activity, including $P$-values (Kolmogorov-Smirnov test) for endpoint *vs*. baseline; data are presented as mean ± SD in L/min

| Regimens | $c_{ACE}$ = 7.0 h$^{-1}$ | | | $c_{ACE}$ = 8.9 h$^{-1}$ | | | $c_{ACE}$ = 10.8 h$^{-1}$ | | | $c_{ACE}$ = 42.3 h$^{-1}$ | | | $c_{ACE}$ = 54.1 h$^{-1}$ | | | $c_{ACE}$ = 65.9 h$^{-1}$ | | |
|---|---|---|---|---|---|---|---|---|---|---|---|---|---|---|---|---|---|---|
| | **Value** | **Change** | ***P*** | **Value** | **Change** | ***P*** | **Value** | **Change** | ***P*** | **Value** | **Change** | ***P*** | **Value** | **Change** | ***P*** | **Value** | **Change** | ***P*** |
| Baseline | 0.0945 ± 0.0192 | – | – | 0.0948 ± 0.0197 | – | – | 0.0916 ± 0.0197 | – | – | 0.0922 ± 0.0186 | – | – | 0.0936 ± 0.0181 | – | – | 0.0950 ± 0.0196 | – | – |
| Al300 | 0.0943 ± 0.0197 | -0.0002 ± 0.0018 | 0.99963 | 0.0945 ± 0.0203 | -0.0003 ± 0.0020 | 1.00000 | 0.0913 ± 0.0201 | -0.0002 ± 0.0018 | 0.99963 | 0.0923 ± 0.0188 | 0.0002 ± 0.0022 | 0.99963 | 0.0943 ± 0.0181 | 0.0007 ± 0.0026 | 0.96707 | 0.0955 ± 0.0200 | 0.0006 ± 0.0026 | 0.99376 |
| E20 | 0.0942 ± 0.0196 | -0.0003 ± 0.0013 | 1.00000 | 0.0944 ± 0.0202 | -0.0004 ± 0.0016 | 1.00000 | 0.0913 ± 0.0200 | -0.0003 ± 0.0015 | 1.00000 | 0.0928 ± 0.0189 | 0.0006 ± 0.0030 | 0.96707 | 0.0950 ± 0.0181 | 0.0015 ± 0.0036 | 0.96707 | 0.0963 ± 0.0201 | 0.0013 ± 0.0036 | 0.90621 |
| L100 | 0.0944 ± 0.0198 | -0.0001 ± 0.0021 | 0.96707 | 0.0946 ± 0.0204 | -0.0002 ± 0.0023 | 1.00000 | 0.0915 ± 0.0201 | -0.0001 ± 0.0021 | 0.99963 | 0.0925 ± 0.0188 | 0.0003 ± 0.0025 | 0.99963 | 0.0946 ± 0.0181 | 0.0010 ± 0.0030 | 0.96707 | 0.0957 ± 0.0200 | 0.0008 ± 0.0029 | 0.99376 |
| Aml5 | 0.1020 ± 0.0182 | 0.0075 ± 0.0043 | 0.01008 | 0.1020 ± 0.0185 | 0.0072 ± 0.0051 | 0.03663 | 0.0993 ± 0.0189 | 0.0077 ± 0.0050 | 0.00025 | 0.1002 ± 0.0178 | 0.0081 ± 0.0051 | 0.00136 | 0.1021 ± 0.0165 | 0.0085 ± 0.0062 | 0.00045 | 0.1030 ± 0.0181 | 0.0080 ± 0.0051 | 0.00630 |
| B5 | 0.0941 ± 0.0198 | -0.0004 ± 0.0019 | 0.99963 | 0.0944 ± 0.0204 | -0.0005 ± 0.0020 | 0.99963 | 0.0912 ± 0.0201 | -0.0004 ± 0.0018 | 0.99963 | 0.0922 ± 0.0188 | -0.0000 ± 0.0022 | 1.00000 | 0.0942 ± 0.0181 | 0.0006 ± 0.0026 | 0.99376 | 0.0954 ± 0.0200 | 0.0004 ± 0.0026 | 0.99963 |
| H12.5 | 0.0853 ± 0.0237 | -0.0092 ± 0.0090 | 0.00136 | 0.0858 ± 0.0236 | -0.0090 ± 0.0091 | 0.01008 | 0.0825 ± 0.0239 | -0.0091 ± 0.0096 | 0.00386 | 0.0827 ± 0.0230 | -0.0095 ± 0.0099 | 0.00232 | 0.0853 ± 0.0223 | -0.0083 ± 0.0097 | 0.00630 | 0.0872 ± 0.0239 | -0.0078 ± 0.0093 | 0.00630 |
| Al300 Aml5 | 0.1029 ± 0.0186 | 0.0084 ± 0.0053 | 0.00630 | 0.1029 ± 0.0187 | 0.0080 ± 0.0054 | 0.00630 | 0.1003 ± 0.0192 | 0.0087 ± 0.0060 | 0.00025 | 0.1016 ± 0.0182 | 0.0094 ± 0.0068 | 0.00025 | 0.1039 ± 0.0166 | 0.0103 ± 0.0078 | SS | 0.1046 ± 0.0183 | 0.0096 ± 0.00630 | 0.00136 |
| Al300 B5 | 0.0950 ± 0.0208 | 0.0006 ± 0.0045 | 0.81275 | 0.0953 ± 0.0210 | 0.0005 ± 0.0043 | 0.99376 | 0.0926 ± 0.0206 | 0.0011 ± 0.0044 | 0.58062 | 0.0943 ± 0.0193 | 0.0021 ± 0.0055 | 0.46756 | 0.0969 ± 0.0183 | 0.0034 ± 0.0060 | 0.36672 | 0.0980 ± 0.0204 | 0.0030 ± 0.0056 | 0.21055 |
| Al300 H12.5 | 0.0866 ± 0.0242 | -0.0079 ± 0.0095 | 0.00630 | 0.0868 ± 0.0241 | -0.0080 ± 0.0093 | 0.02431 | 0.0838 ± 0.0245 | -0.0078 ± 0.0102 | 0.01008 | 0.0846 ± 0.0234 | -0.0076 ± 0.0109 | 0.03663 | 0.0879 ± 0.0227 | -0.0057 ± 0.0114 | 0.03663 | 0.0895 ± 0.0243 | -0.0055 ± 0.0103 | 0.03663 |
| E20 Aml5 | 0.1026 ± 0.0185 | 0.0081 ± 0.0050 | 0.01008 | 0.1026 ± 0.0187 | 0.0078 ± 0.0053 | 0.01008 | 0.1001 ± 0.0192 | 0.0085 ± 0.0058 | 0.00025 | 0.1021 ± 0.0183 | 0.0099 ± 0.0073 | 0.00013 | 0.1046 ± 0.0167 | 0.0110 ± 0.0082 | SS | 0.1053 ± 0.0183 | 0.0104 ± 0.0069 | 0.00013 |
| E20 B5 | 0.0948 ± 0.0204 | 0.0003 ± 0.0036 | 0.99376 | 0.0951 ± 0.0209 | 0.0003 ± 0.0038 | 0.99376 | 0.0924 ± 0.0205 | 0.0008 ± 0.0040 | 0.69937 | 0.0951 ± 0.0195 | 0.0029 ± 0.0065 | 0.28093 | 0.0980 ± 0.0184 | 0.0044 ± 0.0072 | 0.07832 | 0.0991 ± 0.0205 | 0.0041 ± 0.0067 | 0.03663 |
| E20 H12.5 | 0.0861 ± 0.0240 | -0.0084 ± 0.0093 | 0.00386 | 0.0865 ± 0.0240 | -0.0083 ± 0.0092 | 0.02431 | 0.0835 ± 0.0244 | -0.0080 ± 0.0100 | 0.01008 | 0.0852 ± 0.0237 | -0.0070 ± 0.0114 | 0.05410 | 0.0888 ± 0.0229 | -0.0047 ± 0.0121 | 0.05410 | 0.0905 ± 0.0246 | -0.0044 ± 0.0108 | 0.07832 |
| L100 Aml5 | 0.1030 ± 0.0186 | 0.0086 ± 0.0054 | 0.00232 | 0.1030 ± 0.0187 | 0.0082 ± 0.0056 | 0.00386 | 0.1005 ± 0.0193 | 0.0089 ± 0.0061 | 0.00025 | 0.1018 ± 0.0182 | 0.0096 ± 0.0070 | 0.00013 | 0.1042 ± 0.0166 | 0.0106 ± 0.0079 | SS | 0.1048 ± 0.0183 | 0.0098 ± 0.0065 | 0.00079 |
| L100 B5 | 0.0952 ± 0.0210 | 0.0007 ± 0.0050 | 0.81275 | 0.0955 ± 0.0211 | 0.0006 ± 0.0046 | 0.99376 | 0.0929 ± 0.0207 | 0.0013 ± 0.0049 | 0.36672 | 0.0947 ± 0.0194 | 0.0025 ± 0.0059 | 0.36672 | 0.0973 ± 0.0184 | 0.0037 ± 0.0064 | 0.15454 | 0.0983 ± 0.0204 | 0.0034 ± 0.0060 | 0.07832 |
| L100 H12.5 | 0.0868 ± 0.0243 | -0.0077 ± 0.0096 | 0.00630 | 0.0870 ± 0.0242 | -0.0078 ± 0.0094 | 0.03663 | 0.0840 ± 0.0246 | -0.0075 ± 0.0103 | 0.01008 | 0.0848 ± 0.0235 | -0.0073 ± 0.0111 | 0.03663 | 0.0882 ± 0.0228 | -0.0054 ± 0.0117 | 0.03663 | 0.0898 ± 0.0244 | -0.0051 ± 0.0104 | 0.07832 |
| Al300 Aml5/B5 | 0.1042 ± 0.0192 | 0.0097 ± 0.0069 | 0.00025 | 0.1043 ± 0.0190 | 0.0094 ± 0.0068 | 0.00232 | 0.1020 ± 0.0195 | 0.0104 ± 0.0076 | 0.00025 | 0.1034 ± 0.0187 | 0.0112 ± 0.0088 | SS | 0.1060 ± 0.0167 | 0.0125 ± 0.0092 | SS | 0.1066 ± 0.0182 | 0.0117 ± 0.0078 | SS |
| Al300 Aml5/H12.5 | 0.0963 ± 0.0237 | 0.0018 ± 0.0114 | 0.15454 | 0.0960 ± 0.0236 | 0.0012 ± 0.0112 | 0.15454 | 0.0933 ± 0.0249 | 0.0018 ± 0.0127 | 0.03663 | 0.0943 ± 0.0239 | 0.0021 ± 0.0133 | 0.11113 | 0.0976 ± 0.0225 | 0.0040 ± 0.0140 | 0.02431 | 0.0991 ± 0.0236 | 0.0042 ± 0.0123 | 0.01008 |
| Al300 B5/H12.5 | 0.0886 ± 0.0249 | -0.0058 ± 0.0107 | 0.02431 | 0.0888 ± 0.0248 | -0.0061 ± 0.0102 | 0.15454 | 0.0861 ± 0.0254 | -0.0055 ± 0.0117 | 0.01008 | 0.0871 ± 0.0245 | -0.0051 ± 0.0132 | 0.05410 | 0.0909 ± 0.0235 | -0.0027 ± 0.0136 | 0.21055 | 0.0923 ± 0.0250 | -0.0026 ± 0.0119 | 0.21055 |
| E20 Aml5/B5 | 0.1038 ± 0.0190 | 0.0094 ± 0.0064 | 0.00079 | 0.1040 ± 0.0189 | 0.0092 ± 0.0065 | 0.00232 | 0.1018 ± 0.0194 | 0.0102 ± 0.0074 | 0.00025 | 0.1038 ± 0.0188 | 0.0117 ± 0.0092 | SS | 0.1066 ± 0.0168 | 0.0131 ± 0.0096 | SS | 0.1073 ± 0.0182 | 0.0124 ± 0.0083 | SS |
| E20 Aml5/H12.5 | 0.0959 ± 0.0236 | 0.0014 ± 0.0112 | 0.21055 | 0.0957 ± 0.0235 | 0.0009 ± 0.0110 | 0.21055 | 0.0931 ± 0.0248 | 0.0016 ± 0.0125 | 0.05410 | 0.0946 ± 0.0241 | 0.0024 ± 0.0137 | 0.07832 | 0.0981 ± 0.0226 | 0.0045 ± 0.0144 | 0.01008 | 0.0997 ± 0.0237 | 0.0047 ± 0.0126 | 0.00630 |
| E20 B5/H12.5 | 0.0880 ± 0.0247 | -0.0064 ± 0.0103 | 0.02431 | 0.0884 ± 0.0247 | -0.0065 ± 0.0100 | 0.11113 | 0.0858 ± 0.0253 | -0.0058 ± 0.0115 | 0.01008 | 0.0878 ± 0.0248 | -0.0044 ± 0.0139 | 0.05410 | 0.0917 ± 0.0238 | -0.0018 ± 0.0143 | 0.21055 | 0.0932 ± 0.0252 | -0.0017 ± 0.0125 | 0.21055 |
| L100 Aml5/B5 | 0.1043 ± 0.0193 | 0.0098 ± 0.0071 | 0.00025 | 0.1044 ± 0.0190 | 0.0096 ± 0.0071 | 0.00232 | 0.1022 ± 0.0195 | 0.0106 ± 0.0079 | 0.00025 | 0.1036 ± 0.0187 | 0.0114 ± 0.0089 | SS | 0.1063 ± 0.0167 | 0.0127 ± 0.0094 | SS | 0.1069 ± 0.0182 | 0.0119 ± 0.0080 | SS |
| L100 Aml5/H12.5 | 0.0965 ± 0.0238 | 0.0020 ± 0.0115 | 0.15454 | 0.0962 ± 0.0236 | 0.0014 ± 0.0113 | 0.15454 | 0.0935 ± 0.0250 | 0.0020 ± 0.0128 | 0.02431 | 0.0944 ± 0.0240 | 0.0023 ± 0.0134 | 0.11113 | 0.0978 ± 0.0225 | 0.0042 ± 0.0141 | 0.01581 | 0.0993 ± 0.0236 | 0.0044 ± 0.0124 | 0.00630 |
| L100 B5/H12.5 | 0.0889 ± 0.0251 | -0.0055 ± 0.0108 | 0.02431 | 0.0891 ± 0.0249 | -0.0058 ± 0.0104 | 0.15454 | 0.0864 ± 0.0255 | -0.0051 ± 0.0120 | 0.01008 | 0.0874 ± 0.0247 | -0.0048 ± 0.0135 | 0.05410 | 0.0912 ± 0.0236 | -0.0024 ± 0.0139 | 0.21055 | 0.0926 ± 0.0251 | -0.0023 ± 0.0121 | 0.21055 |

**Al300 =** aliskiren 300 mg; **Aml5** = amlodipine 5 mg; **B5** = bisoprolol 5 mg; **E20** = enalapril 20 mg; **H12.5** = hydrochlorothiazide 12.5 mg; **L100** = losartan 100 mg; **SS** = statistically significant ($P$ < 0.00001)

**Table S37.** $P$-values calculated using the Kolmogorov-Smirnov test for changes in glomerular filtration rate in populations ($n$ = 100) with different ACE activity receiving the same regimens (case 1: $c_{ACE}$ = 7.0 $h^{-1}$, case 2: $c_{ACE}$ = 8.9 $h^{-1}$, case 3: $c_{ACE}$ = 10.8 $h^{-1}$, case 4: $c_{ACE}$ = 42.3 $h^{-1}$, case 5: $c_{ACE}$ = 54.1 $h^{-1}$, case 6: $c_{ACE}$ = 65.9 $h^{-1}$; $P$-value for case $i$ *vs.* case $j$ is denoted $P_{ij}$)

| Regimens | $P_{12}$ | $P_{13}$ | $P_{23}$ | $P_{14}$ | $P_{15}$ | $P_{16}$ | $P_{24}$ | $P_{25}$ | $P_{26}$ | $P_{34}$ | $P_{35}$ | $P_{36}$ | $P_{45}$ | $P_{46}$ | $P_{56}$ |
|---|---|---|---|---|---|---|---|---|---|---|---|---|---|---|---|
| Al300 | 0.99376 | 0.58062 | 0.90621 | 0.58062 | 0.05410 | 0.07832 | 0.58062 | 0.01581 | 0.00630 | 0.28093 | 0.01581 | 0.01581 | 0.21055 | 0.36672 | 0.99376 |
| E20 | 0.90621 | 0.90621 | 0.99963 | 0.00232 | SS | SS | 0.00386 | SS | SS | 0.00136 | SS | SS | 0.11113 | 0.28093 | 0.99376 |
| L100 | 0.90621 | 0.46756 | 0.81275 | 0.46756 | 0.02431 | 0.05410 | 0.46756 | 0.00630 | 0.01008 | 0.28093 | 0.01008 | 0.02431 | 0.21055 | 0.36672 | 0.99963 |
| Aml5 | 0.36672 | 0.81275 | 0.81275 | 0.81275 | 0.81275 | 0.46756 | 0.21055 | 0.15454 | 0.00630 | 0.46756 | 0.36672 | 0.05410 | 0.96707 | 0.36672 | 0.69937 |
| B5 | 0.90621 | 0.69937 | 0.99963 | 0.58062 | 0.07832 | 0.05410 | 0.36672 | 0.05410 | 0.01008 | 0.28093 | 0.05410 | 0.02431 | 0.28093 | 0.11113 | 0.96707 |
| H12.5 | 0.96707 | 0.46756 | 0.46756 | 0.69937 | 0.36672 | 0.21055 | 0.58062 | 0.36672 | 0.02431 | 0.81275 | 0.21055 | 0.36672 | 0.46756 | 0.15454 | 0.81275 |
| Al300<br>Aml5 | 0.46756 | 0.69937 | 0.58062 | 0.81275 | 0.46756 | 0.36672 | 0.21055 | 0.07832 | 0.05410 | 0.28093 | 0.07832 | 0.07832 | 0.69937 | 0.81275 | 0.81275 |
| Al300<br>B5 | 0.69937 | 0.58062 | 0.69937 | 0.21055 | 0.00630 | 0.01008 | 0.21055 | 0.00232 | 0.00630 | 0.58062 | 0.01581 | 0.03663 | 0.28093 | 0.28093 | 0.96707 |
| Al300<br>H12.5 | 0.90621 | 0.58062 | 0.69937 | 0.90621 | 0.28093 | 0.11113 | 0.58062 | 0.21055 | 0.03663 | 0.69937 | 0.15454 | 0.15454 | 0.58062 | 0.28093 | 0.99376 |
| E20<br>Aml5 | 0.46756 | 0.58062 | 0.69937 | 0.46756 | 0.07832 | 0.05410 | 0.03663 | 0.00136 | 0.00232 | 0.11113 | 0.00630 | 0.00630 | 0.69937 | 0.46756 | 0.81275 |
| E20<br>B5 | 0.96707 | 0.58062 | 0.58062 | 0.00630 | SS | 0.00002 | 0.00630 | SS | SS | 0.05410 | 0.00013 | 0.00025 | 0.15454 | 0.15454 | 0.99376 |
| E20<br>H12.5 | 0.90621 | 0.58062 | 0.46756 | 0.46756 | 0.07832 | 0.05410 | 0.28093 | 0.01581 | 0.00386 | 0.46756 | 0.03663 | 0.02431 | 0.46756 | 0.15454 | 0.96707 |
| L100<br>Aml5 | 0.46756 | 0.69937 | 0.69937 | 0.81275 | 0.46756 | 0.46756 | 0.21055 | 0.05410 | 0.05410 | 0.36672 | 0.07832 | 0.11113 | 0.58062 | 0.81275 | 0.81275 |
| L100<br>B5 | 0.46756 | 0.46756 | 0.58062 | 0.15454 | 0.00630 | 0.00630 | 0.15454 | 0.00136 | 0.00630 | 0.46756 | 0.03663 | 0.05410 | 0.28093 | 0.28093 | 0.96707 |
| L100<br>H12.5 | 0.96707 | 0.58062 | 0.69937 | 0.96707 | 0.36672 | 0.07832 | 0.58062 | 0.15454 | 0.07832 | 0.69937 | 0.21055 | 0.15454 | 0.46756 | 0.21055 | 0.99376 |
| Al300<br>Aml5/B5 | 0.58062 | 0.36672 | 0.58062 | 0.58062 | 0.15454 | 0.15454 | 0.28093 | 0.01581 | 0.02431 | 0.36672 | 0.15454 | 0.07832 | 0.28093 | 0.28093 | 0.96707 |
| Al300<br>Aml5/H12.5 | 0.46756 | 0.36672 | 0.81275 | 0.90621 | 0.69937 | 0.46756 | 0.28093 | 0.07832 | 0.05410 | 0.69937 | 0.36672 | 0.21055 | 0.58062 | 0.36672 | 0.90621 |
| Al300<br>B5/H12.5 | 0.69937 | 0.58062 | 0.81275 | 0.69937 | 0.36672 | 0.46756 | 0.28093 | 0.07832 | 0.05410 | 0.46756 | 0.11113 | 0.03663 | 0.36672 | 0.28093 | 0.96707 |
| E20<br>Aml5/B5 | 0.69937 | 0.46756 | 0.69937 | 0.46756 | 0.03663 | 0.05410 | 0.07832 | 0.00386 | 0.01008 | 0.21055 | 0.01008 | 0.01008 | 0.21055 | 0.21055 | 0.99376 |
| E20<br>Aml5/H12.5 | 0.36672 | 0.46756 | 0.81275 | 0.81275 | 0.36672 | 0.21055 | 0.15454 | 0.02431 | 0.01008 | 0.46756 | 0.28093 | 0.05410 | 0.69937 | 0.46756 | 0.81275 |
| E20<br>B5/H12.5 | 0.96707 | 0.90621 | 0.81275 | 0.21055 | 0.05410 | 0.07832 | 0.05410 | 0.01008 | 0.00630 | 0.36672 | 0.02431 | 0.01581 | 0.36672 | 0.28093 | 0.81275 |
| L100<br>Aml5/B5 | 0.69937 | 0.36672 | 0.36672 | 0.36672 | 0.11113 | 0.11113 | 0.28093 | 0.02431 | 0.02431 | 0.36672 | 0.15454 | 0.07832 | 0.28093 | 0.28093 | 0.96707 |
| L100<br>Aml5/H12.5 | 0.46756 | 0.36672 | 0.81275 | 0.96707 | 0.58062 | 0.46756 | 0.28093 | 0.07832 | 0.03663 | 0.81275 | 0.36672 | 0.21055 | 0.69937 | 0.46756 | 0.81275 |
| L100<br>B5/H12.5 | 0.58062 | 0.69937 | 0.81275 | 0.69937 | 0.28093 | 0.36672 | 0.28093 | 0.07832 | 0.05410 | 0.58062 | 0.07832 | 0.03663 | 0.36672 | 0.28093 | 0.90621 |

**Al300 =** aliskiren 300 mg; **Aml5** = amlodipine 5 mg; **B5** = bisoprolol 5 mg; **E20** = enalapril 20 mg; **H12.5** = hydrochlorothiazide 12.5 mg; **L100** = losartan 100 mg; **SS** = statistically significant ($P < 0.00001$)

**Figure S26.** Simulated change in <u>renal blood flow</u> from baseline to week 4 (mean ± SD, $n$ = 100)

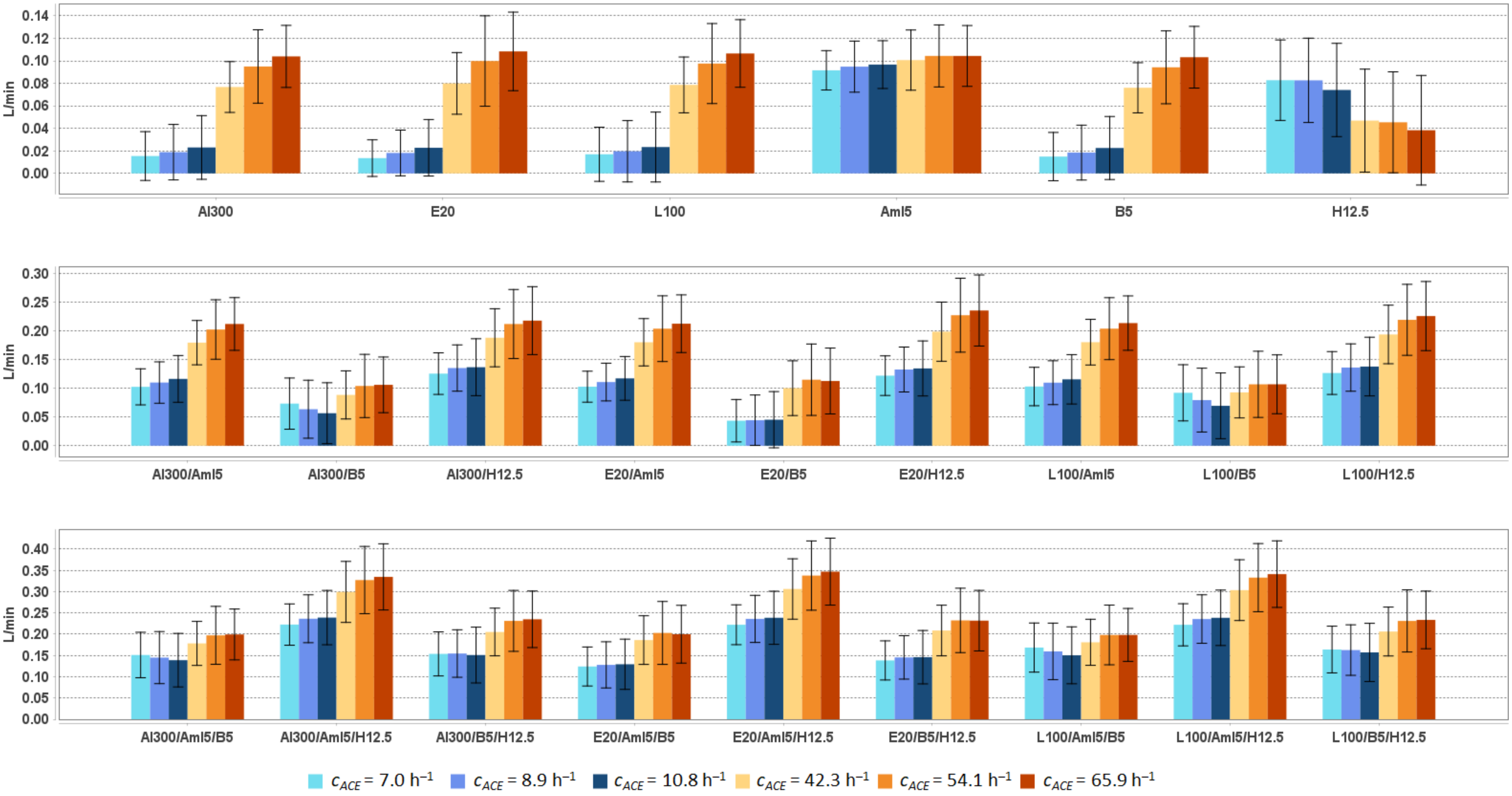


**Al300 =** aliskiren 300 mg; **Aml5** = amlodipine 5 mg; **B5** = bisoprolol 5 mg; **E20** = enalapril 20 mg; **H12.5** = hydrochlorothiazide 12.5 mg; **L100** = losartan 100 mg

**Table S38.** Simulated response of renal blood flow to antihypertensive therapy in virtual hypertensive populations ($n$ = 100) with different ACE activity, including $P$-values (Kolmogorov-Smirnov test) for endpoint *vs*. baseline; data are presented as mean ± SD in L/min

| **Regimens** | $c_{ACE}$ = 7.0 h$^{-1}$ | | | $c_{ACE}$ = 8.9 h$^{-1}$ | | | $c_{ACE}$ = 10.8 h$^{-1}$ | | | $c_{ACE}$ = 42.3 h$^{-1}$ | | | $c_{ACE}$ = 54.1 h$^{-1}$ | | | $c_{ACE}$ = 65.9 h$^{-1}$ | | |
|---|---|---|---|---|---|---|---|---|---|---|---|---|---|---|---|---|---|---|
| | **Value** | **Change** | ***P*** | **Value** | **Change** | ***P*** | **Value** | **Change** | ***P*** | **Value** | **Change** | ***P*** | **Value** | **Change** | ***P*** | **Value** | **Change** | ***P*** |
| Baseline | 1.082 ± 0.147 | – | – | 1.108 ± 0.168 | – | – | 1.114 ± 0.161 | – | – | 1.106 ± 0.153 | – | – | 1.127 ± 0.152 | – | – | 1.142 ± 0.164 | – | – |
| Al300 | 1.098 ± 0.147 | 0.015 ± 0.022 | 0.69937 | 1.127 ± 0.165 | 0.019 ± 0.025 | 0.81275 | 1.137 ± 0.162 | 0.023 ± 0.028 | 0.90621 | 1.183 ± 0.159 | 0.077 ± 0.023 | 0.00232 | 1.222 ± 0.165 | 0.095 ± 0.033 | 0.00232 | 1.246 ± 0.175 | 0.104 ± 0.028 | 0.00025 |
| E20 | 1.096 ± 0.147 | 0.014 ± 0.016 | 0.81275 | 1.126 ± 0.166 | 0.018 ± 0.020 | 0.90621 | 1.137 ± 0.162 | 0.023 ± 0.025 | 0.90621 | 1.186 ± 0.159 | 0.080 ± 0.027 | 0.00079 | 1.227 ± 0.166 | 0.100 ± 0.040 | 0.00136 | 1.251 ± 0.175 | 0.108 ± 0.035 | 0.00025 |
| L100 | 1.099 ± 0.147 | 0.017 ± 0.024 | 0.69937 | 1.128 ± 0.164 | 0.020 ± 0.027 | 0.81275 | 1.138 ± 0.162 | 0.023 ± 0.031 | 0.81275 | 1.185 ± 0.159 | 0.079 ± 0.025 | 0.00232 | 1.225 ± 0.165 | 0.098 ± 0.036 | 0.00136 | 1.249 ± 0.175 | 0.107 ± 0.030 | 0.00025 |
| Aml5 | 1.174 ± 0.156 | 0.092 ± 0.017 | 0.00136 | 1.203 ± 0.181 | 0.095 ± 0.023 | 0.00232 | 1.211 ± 0.171 | 0.097 ± 0.021 | 0.00079 | 1.207 ± 0.166 | 0.101 ± 0.027 | 0.00079 | 1.232 ± 0.166 | 0.104 ± 0.028 | 0.00025 | 1.246 ± 0.177 | 0.104 ± 0.027 | 0.00007 |
| B5 | 1.097 ± 0.147 | 0.015 ± 0.021 | 0.69937 | 1.127 ± 0.165 | 0.018 ± 0.024 | 0.81275 | 1.137 ± 0.162 | 0.023 ± 0.028 | 0.90621 | 1.183 ± 0.159 | 0.076 ± 0.022 | 0.00232 | 1.222 ± 0.165 | 0.094 ± 0.032 | 0.00232 | 1.245 ± 0.175 | 0.103 ± 0.027 | 0.00045 |
| H12.5 | 1.165 ± 0.158 | 0.083 ± 0.036 | 0.00232 | 1.191 ± 0.185 | 0.083 ± 0.037 | 0.02431 | 1.188 ± 0.174 | 0.074 ± 0.041 | 0.01581 | 1.153 ± 0.171 | 0.047 ± 0.046 | 0.28093 | 1.173 ± 0.165 | 0.045 ± 0.045 | 0.21055 | 1.180 ± 0.179 | 0.038 ± 0.049 | 0.46756 |
| Al300 Aml5 | 1.185 ± 0.156 | 0.102 ± 0.031 | 0.00004 | 1.218 ± 0.177 | 0.110 ± 0.036 | 0.00025 | 1.230 ± 0.172 | 0.116 ± 0.041 | 0.00013 | 1.286 ± 0.172 | 0.179 ± 0.039 | SS | 1.329 ± 0.179 | 0.202 ± 0.052 | SS | 1.354 ± 0.188 | 0.212 ± 0.046 | SS |
| Al300 B5 | 1.156 ± 0.155 | 0.073 ± 0.045 | 0.00232 | 1.172 ± 0.168 | 0.063 ± 0.050 | 0.03663 | 1.171 ± 0.164 | 0.057 ± 0.053 | 0.11113 | 1.195 ± 0.158 | 0.088 ± 0.042 | 0.00045 | 1.231 ± 0.166 | 0.104 ± 0.055 | 0.00045 | 1.248 ± 0.174 | 0.106 ± 0.048 | 0.00013 |
| Al300 H12.5 | 1.208 ± 0.159 | 0.125 ± 0.036 | SS | 1.243 ± 0.184 | 0.135 ± 0.040 | 0.00004 | 1.251 ± 0.180 | 0.136 ± 0.049 | 0.00002 | 1.294 ± 0.182 | 0.188 ± 0.050 | SS | 1.339 ± 0.185 | 0.212 ± 0.060 | SS | 1.360 ± 0.197 | 0.217 ± 0.059 | SS |
| E20 Aml5 | 1.185 ± 0.156 | 0.103 ± 0.027 | 0.00004 | 1.219 ± 0.178 | 0.111 ± 0.033 | 0.00025 | 1.231 ± 0.172 | 0.117 ± 0.038 | 0.00007 | 1.286 ± 0.170 | 0.180 ± 0.041 | SS | 1.331 ± 0.179 | 0.203 ± 0.057 | SS | 1.354 ± 0.187 | 0.212 ± 0.050 | SS |
| E20 B5 | 1.126 ± 0.151 | 0.044 ± 0.037 | 0.11113 | 1.153 ± 0.166 | 0.044 ± 0.044 | 0.15454 | 1.159 ± 0.163 | 0.045 ± 0.049 | 0.28093 | 1.207 ± 0.159 | 0.100 ± 0.048 | 0.00013 | 1.242 ± 0.167 | 0.115 ± 0.062 | 0.00013 | 1.255 ± 0.176 | 0.113 ± 0.057 | 0.00007 |
| E20 H12.5 | 1.204 ± 0.159 | 0.122 ± 0.034 | SS | 1.241 ± 0.185 | 0.133 ± 0.039 | 0.00007 | 1.249 ± 0.180 | 0.134 ± 0.048 | 0.00002 | 1.305 ± 0.181 | 0.198 ± 0.051 | SS | 1.354 ± 0.186 | 0.227 ± 0.064 | SS | 1.377 ± 0.198 | 0.235 ± 0.062 | SS |
| L100 Aml5 | 1.185 ± 0.156 | 0.103 ± 0.033 | 0.00002 | 1.218 ± 0.176 | 0.110 ± 0.038 | 0.00013 | 1.230 ± 0.172 | 0.115 ± 0.043 | 0.00013 | 1.286 ± 0.171 | 0.180 ± 0.040 | SS | 1.331 ± 0.179 | 0.204 ± 0.054 | SS | 1.355 ± 0.188 | 0.213 ± 0.047 | SS |
| L100 B5 | 1.174 ± 0.157 | 0.092 ± 0.049 | 0.00013 | 1.188 ± 0.171 | 0.079 ± 0.055 | 0.01008 | 1.184 ± 0.165 | 0.069 ± 0.057 | 0.05410 | 1.199 ± 0.158 | 0.093 ± 0.044 | 0.00025 | 1.234 ± 0.166 | 0.107 ± 0.058 | 0.00045 | 1.249 ± 0.175 | 0.107 ± 0.051 | 0.00013 |
| L100 H12.5 | 1.209 ± 0.159 | 0.126 ± 0.037 | SS | 1.244 ± 0.184 | 0.136 ± 0.041 | 0.00004 | 1.252 ± 0.180 | 0.138 ± 0.051 | 0.00002 | 1.300 ± 0.182 | 0.193 ± 0.051 | SS | 1.346 ± 0.185 | 0.219 ± 0.062 | SS | 1.368 ± 0.198 | 0.225 ± 0.060 | SS |
| Al300 Aml5/B5 | 1.233 ± 0.162 | 0.151 ± 0.053 | SS | 1.253 ± 0.178 | 0.145 ± 0.061 | SS | 1.253 ± 0.173 | 0.139 ± 0.063 | 0.00007 | 1.285 ± 0.168 | 0.178 ± 0.052 | SS | 1.325 ± 0.177 | 0.197 ± 0.068 | SS | 1.341 ± 0.184 | 0.199 ± 0.060 | SS |
| Al300 Aml5/H12.5 | 1.305 ± 0.170 | 0.223 ± 0.049 | SS | 1.344 ± 0.198 | 0.236 ± 0.056 | SS | 1.353 ± 0.191 | 0.239 ± 0.064 | SS | 1.406 ± 0.198 | 0.299 ± 0.072 | SS | 1.455 ± 0.199 | 0.327 ± 0.079 | SS | 1.477 ± 0.213 | 0.335 ± 0.078 | SS |
| Al300 B5/H12.5 | 1.236 ± 0.162 | 0.154 ± 0.052 | SS | 1.263 ± 0.182 | 0.155 ± 0.056 | SS | 1.265 ± 0.179 | 0.151 ± 0.066 | SS | 1.312 ± 0.178 | 0.205 ± 0.056 | SS | 1.359 ± 0.185 | 0.231 ± 0.072 | SS | 1.377 ± 0.196 | 0.235 ± 0.066 | SS |
| E20 Aml5/B5 | 1.206 ± 0.159 | 0.124 ± 0.046 | SS | 1.236 ± 0.176 | 0.128 ± 0.054 | SS | 1.243 ± 0.172 | 0.129 ± 0.059 | 0.00013 | 1.293 ± 0.169 | 0.186 ± 0.057 | SS | 1.330 ± 0.177 | 0.203 ± 0.074 | SS | 1.342 ± 0.185 | 0.200 ± 0.068 | SS |
| E20 Aml5/H12.5 | 1.304 ± 0.170 | 0.222 ± 0.047 | SS | 1.344 ± 0.200 | 0.236 ± 0.055 | SS | 1.353 ± 0.192 | 0.239 ± 0.062 | SS | 1.413 ± 0.197 | 0.306 ± 0.071 | SS | 1.465 ± 0.200 | 0.338 ± 0.081 | SS | 1.489 ± 0.212 | 0.347 ± 0.079 | SS |
| E20 B5/H12.5 | 1.221 ± 0.160 | 0.138 ± 0.046 | SS | 1.254 ± 0.181 | 0.145 ± 0.051 | SS | 1.260 ± 0.179 | 0.146 ± 0.063 | SS | 1.315 ± 0.178 | 0.209 ± 0.059 | SS | 1.360 ± 0.184 | 0.232 ± 0.076 | SS | 1.374 ± 0.195 | 0.232 ± 0.071 | SS |
| L100 Aml5/B5 | 1.251 ± 0.165 | 0.169 ± 0.058 | SS | 1.268 ± 0.181 | 0.160 ± 0.066 | SS | 1.264 ± 0.174 | 0.150 ± 0.067 | SS | 1.287 ± 0.168 | 0.181 ± 0.054 | SS | 1.326 ± 0.177 | 0.198 ± 0.070 | SS | 1.340 ± 0.184 | 0.198 ± 0.062 | SS |
| L100 Aml5/H12.5 | 1.304 ± 0.169 | 0.222 ± 0.050 | SS | 1.344 ± 0.197 | 0.236 ± 0.057 | SS | 1.353 ± 0.191 | 0.238 ± 0.065 | SS | 1.410 ± 0.198 | 0.304 ± 0.071 | SS | 1.460 ± 0.200 | 0.333 ± 0.080 | SS | 1.483 ± 0.213 | 0.341 ± 0.078 | SS |
| L100 B5/H12.5 | 1.246 ± 0.164 | 0.164 ± 0.055 | SS | 1.271 ± 0.183 | 0.163 ± 0.059 | SS | 1.271 ± 0.180 | 0.157 ± 0.068 | SS | 1.313 ± 0.178 | 0.206 ± 0.057 | SS | 1.359 ± 0.185 | 0.231 ± 0.073 | SS | 1.376 ± 0.196 | 0.234 ± 0.068 | SS |

**Al300 =** aliskiren 300 mg; **Aml5** = amlodipine 5 mg; **B5** = bisoprolol 5 mg; **E20** = enalapril 20 mg; **H12.5** = hydrochlorothiazide 12.5 mg; **L100** = losartan 100 mg; **SS** = statistically significant ($P$ < 0.00001)

**Table S39.** *P*-values calculated using the Kolmogorov-Smirnov test for changes in renal blood flow in populations ($n$ = 100) with different ACE activity receiving the same regimens (case 1: $c_{ACE}$ = 7.0 h$^{-1}$, case 2: $c_{ACE}$ = 8.9 h$^{-1}$, case 3: $c_{ACE}$ = 10.8 h$^{-1}$, case 4: $c_{ACE}$ = 42.3 h$^{-1}$, case 5: $c_{ACE}$ = 54.1 h$^{-1}$, case 6: $c_{ACE}$ = 65.9 h$^{-1}$; *P*-value for case *i* *vs*. case *j* is denoted $P_{ij}$)

| Regimens | $P_{12}$ | $P_{13}$ | $P_{23}$ | $P_{14}$ | $P_{15}$ | $P_{16}$ | $P_{24}$ | $P_{25}$ | $P_{26}$ | $P_{34}$ | $P_{35}$ | $P_{36}$ | $P_{45}$ | $P_{46}$ | $P_{56}$ |
|---|---|---|---|---|---|---|---|---|---|---|---|---|---|---|---|
| Al300 | 0.58062 | 0.03663 | 0.69937 | SS | SS | SS | SS | SS | SS | SS | SS | SS | 0.00079 | SS | 0.01581 |
| E20 | 0.11113 | 0.00386 | 0.36672 | SS | SS | SS | SS | SS | SS | SS | SS | SS | 0.00232 | 0.00002 | 0.07832 |
| L100 | 0.69937 | 0.15454 | 0.69937 | SS | SS | SS | SS | SS | SS | SS | SS | SS | 0.00136 | SS | 0.02431 |
| Aml5 | 0.36672 | 0.46756 | 0.58062 | 0.05410 | 0.00232 | 0.00045 | 0.36672 | 0.05410 | 0.01581 | 0.90621 | 0.07832 | 0.07832 | 0.28093 | 0.36672 | 0.99376 |
| B5 | 0.58062 | 0.03663 | 0.69937 | SS | SS | SS | SS | SS | SS | SS | SS | SS | 0.00045 | SS | 0.02431 |
| H12.5 | 0.81275 | 0.07832 | 0.15454 | SS | SS | SS | SS | SS | SS | 0.00045 | 0.00136 | 0.00013 | 0.81275 | 0.46756 | 0.36672 |
| Al300 Aml5 | 0.58062 | 0.07832 | 0.36672 | SS | SS | SS | SS | SS | SS | SS | SS | SS | 0.03663 | 0.00013 | 0.11113 |
| Al300 B5 | 0.36672 | 0.21055 | 0.58062 | 0.21055 | 0.00630 | 0.00630 | 0.01008 | 0.00045 | 0.00007 | 0.00232 | 0.00002 | SS | 0.07832 | 0.21055 | 0.90621 |
| Al300 H12.5 | 0.15454 | 0.11113 | 0.90621 | SS | SS | SS | SS | SS | SS | SS | SS | SS | 0.03663 | 0.00136 | 0.36672 |
| E20 Aml5 | 0.36672 | 0.00630 | 0.21055 | SS | SS | SS | SS | SS | SS | SS | SS | SS | 0.02431 | 0.00232 | 0.28093 |
| E20 B5 | 0.99376 | 0.58062 | 0.69937 | SS | SS | SS | SS | SS | SS | SS | SS | SS | 0.07832 | 0.21055 | 0.69937 |
| E20 H12.5 | 0.07832 | 0.05410 | 0.96707 | SS | SS | SS | SS | SS | SS | SS | SS | SS | 0.05410 | 0.00025 | 0.21055 |
| L100 Aml5 | 0.58062 | 0.15454 | 0.28093 | SS | SS | SS | SS | SS | SS | SS | SS | SS | 0.03663 | 0.00079 | 0.11113 |
| L100 B5 | 0.07832 | 0.05410 | 0.58062 | 0.69937 | 0.15454 | 0.36672 | 0.28093 | 0.02431 | 0.02431 | 0.03663 | 0.01008 | 0.00232 | 0.07832 | 0.28093 | 0.99376 |
| L100 H12.5 | 0.21055 | 0.21055 | 0.69937 | SS | SS | SS | SS | SS | SS | SS | SS | SS | 0.07832 | 0.00079 | 0.36672 |
| Al300 Aml5/B5 | 0.58062 | 0.28093 | 0.81275 | 0.00630 | 0.00007 | 0.00013 | 0.00079 | 0.00002 | SS | 0.00025 | SS | SS | 0.21055 | 0.21055 | 0.46756 |
| Al300 Aml5/H12.5 | 0.05410 | 0.03663 | 0.58062 | SS | SS | SS | SS | SS | SS | SS | SS | SS | 0.15454 | 0.00232 | 0.21055 |
| Al300 B5/H12.5 | 0.90621 | 0.58062 | 0.81275 | SS | SS | SS | SS | SS | SS | SS | SS | SS | 0.11113 | 0.03663 | 0.81275 |
| E20 Aml5/B5 | 0.90621 | 0.28093 | 0.90621 | SS | SS | SS | SS | SS | SS | SS | SS | SS | 0.28093 | 0.46756 | 0.21055 |
| E20 Aml5/H12.5 | 0.03663 | 0.02431 | 0.69937 | SS | SS | SS | SS | SS | SS | SS | SS | SS | 0.11113 | 0.00079 | 0.28093 |
| E20 B5/H12.5 | 0.90621 | 0.21055 | 0.46756 | SS | SS | SS | SS | SS | SS | SS | SS | SS | 0.28093 | 0.11113 | 0.96707 |
| L100 Aml5/B5 | 0.46756 | 0.11113 | 0.81275 | 0.46756 | 0.03663 | 0.07832 | 0.07832 | 0.00232 | 0.00386 | 0.01581 | 0.00025 | 0.00007 | 0.28093 | 0.36672 | 0.81275 |
| L100 Aml5/H12.5 | 0.07832 | 0.05410 | 0.69937 | SS | SS | SS | SS | SS | SS | SS | SS | SS | 0.11113 | 0.00079 | 0.21055 |
| L100 B5/H12.5 | 0.81275 | 0.46756 | 0.58062 | 0.00045 | SS | SS | SS | SS | SS | SS | SS | SS | 0.11113 | 0.07832 | 0.90621 |

**Al300 =** aliskiren 300 mg; **Aml5** = amlodipine 5 mg; **B5** = bisoprolol 5 mg; **E20** = enalapril 20 mg; **H12.5** = hydrochlorothiazide 12.5 mg; **L100** = losartan 100 mg; **SS** = statistically significant ($P$ < 0.00001)

**Figure S27.** Simulated change in <u>renal vascular resistance</u> from baseline to week 4 (mean ± SD, $n$ = 100)

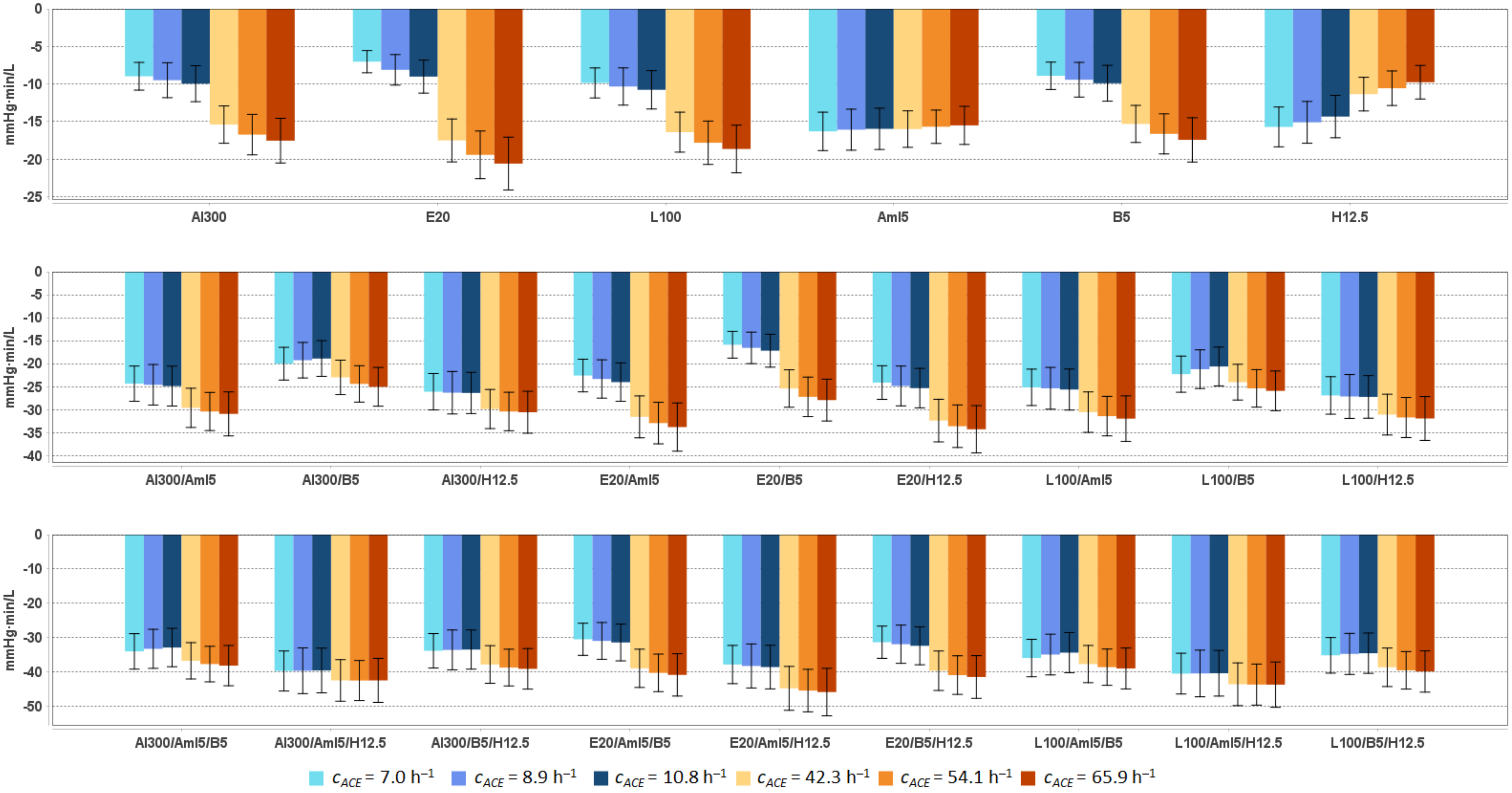


**Al300 =** aliskiren 300 mg; **Aml5** = amlodipine 5 mg; **B5** = bisoprolol 5 mg; **E20** = enalapril 20 mg; **H12.5** = hydrochlorothiazide 12.5 mg; **L100** = losartan 100 mg

**Table S40.** Simulated response of <u>renal vascular resistance</u> to antihypertensive therapy in virtual hypertensive populations ($n$ = 100) with different ACE activity, including $P$-values (Kolmogorov-Smirnov test) for endpoint *vs*. baseline; data are presented as mean ± SD in mmHg·min/L

| Regimens | $c_{ACE}$ = 7.0 h$^{-1}$ | | | $c_{ACE}$ = 8.9 h$^{-1}$ | | | $c_{ACE}$ = 10.8 h$^{-1}$ | | | $c_{ACE}$ = 42.3 h$^{-1}$ | | | $c_{ACE}$ = 54.1 h$^{-1}$ | | | $c_{ACE}$ = 65.9 h$^{-1}$ | | |
|---|---|---|---|---|---|---|---|---|---|---|---|---|---|---|---|---|---|---|
| | **Value** | **Change** | ***P*** | **Value** | **Change** | ***P*** | **Value** | **Change** | ***P*** | **Value** | **Change** | ***P*** | **Value** | **Change** | ***P*** | **Value** | **Change** | ***P*** |
| Baseline | 107.4 ± 14.3 | – | – | 106.0 ± 15.9 | – | – | 105.1 ± 15.3 | – | – | 104.7 ± 13.7 | – | – | 102.7 ± 12.8 | – | – | 102.0 ± 13.9 | – | – |
| Al300 | 98.4 ± 13.3 | -9.0 ± 1.8 | 0.00079 | 96.5 ± 14.3 | -9.5 ± 2.3 | 0.00025 | 95.1 ± 14.0 | -10.0 ± 2.4 | 0.00232 | 89.3 ± 12.0 | -15.4 ± 2.5 | SS | 85.9 ± 11.4 | -16.7 ± 2.7 | SS | 84.5 ± 11.9 | -17.5 ± 3.0 | SS |
| E20 | 100.4 ± 13.5 | -7.0 ± 1.5 | 0.00136 | 97.9 ± 14.5 | -8.1 ± 2.0 | 0.00079 | 96.0 ± 14.1 | -9.0 ± 2.2 | 0.00386 | 87.2 ± 11.8 | -17.5 ± 2.9 | SS | 83.2 ± 11.3 | -19.4 ± 3.2 | SS | 81.4 ± 11.6 | -20.6 ± 3.5 | SS |
| L100 | 97.5 ± 13.2 | -9.9 ± 2.0 | 0.00025 | 95.6 ± 14.2 | -10.3 ± 2.5 | 0.00007 | 94.3 ± 13.9 | -10.8 ± 2.6 | 0.00025 | 88.3 ± 11.9 | -16.4 ± 2.7 | SS | 84.9 ± 11.4 | -17.8 ± 2.9 | SS | 83.4 ± 11.7 | -18.7 ± 3.2 | SS |
| Aml5 | 91.1 ± 11.9 | -16.3 ± 2.6 | SS | 89.9 ± 13.4 | -16.1 ± 2.7 | SS | 89.1 ± 12.7 | -16.0 ± 2.7 | SS | 88.7 ± 11.5 | -16.0 ± 2.4 | SS | 87.0 ± 10.9 | -15.7 ± 2.2 | SS | 86.5 ± 11.6 | -15.5 ± 2.5 | SS |
| B5 | 98.5 ± 13.3 | -8.9 ± 1.8 | 0.00079 | 96.5 ± 14.3 | -9.4 ± 2.3 | 0.00025 | 95.2 ± 14.0 | -9.9 ± 2.4 | 0.00232 | 89.4 ± 12.0 | -15.3 ± 2.5 | SS | 86.0 ± 11.4 | -16.6 ± 2.7 | SS | 84.6 ± 11.9 | -17.4 ± 2.9 | SS |
| H12.5 | 91.7 ± 12.0 | -15.7 ± 2.7 | SS | 90.9 ± 13.4 | -15.1 ± 2.8 | SS | 90.7 ± 12.9 | -14.3 ± 2.8 | SS | 93.4 ± 12.1 | -11.3 ± 2.2 | SS | 92.1 ± 11.3 | -10.6 ± 2.3 | 0.00004 | 92.2 ± 12.7 | -9.8 ± 2.2 | 0.00002 |
| Al300<br>Aml5 | 83.1 ± 11.1 | -24.4 ± 3.8 | SS | 81.4 ± 12.1 | -24.6 ± 4.4 | SS | 80.2 ± 11.7 | -24.9 ± 4.3 | SS | 75.1 ± 10.0 | -29.6 ± 4.3 | SS | 72.3 ± 9.6 | -30.4 ± 4.1 | SS | 71.1 ± 9.8 | -30.9 ± 4.8 | SS |
| Al300<br>B5 | 87.4 ± 12.0 | -20.0 ± 3.6 | SS | 86.7 ± 13.1 | -19.3 ± 3.9 | SS | 86.2 ± 12.7 | -18.9 ± 3.9 | SS | 81.7 ± 11.4 | -23.0 ± 3.7 | SS | 78.3 ± 11.0 | -24.4 ± 4.0 | SS | 77.0 ± 11.2 | -25.0 ± 4.2 | SS |
| Al300<br>H12.5 | 81.3 ± 10.9 | -26.1 ± 4.0 | SS | 79.7 ± 11.7 | -26.3 ± 4.6 | SS | 78.7 ± 11.3 | -26.4 ± 4.5 | SS | 74.8 ± 9.8 | -29.9 ± 4.3 | SS | 72.3 ± 9.2 | -30.4 ± 4.2 | SS | 71.4 ± 9.8 | -30.6 ± 4.6 | SS |
| E20<br>Aml5 | 84.8 ± 11.3 | -22.6 ± 3.6 | SS | 82.7 ± 12.3 | -23.3 ± 4.2 | SS | 81.0 ± 11.8 | -24.0 ± 4.2 | SS | 73.1 ± 9.9 | -31.6 ± 4.6 | SS | 69.8 ± 9.5 | -32.9 ± 4.5 | SS | 68.2 ± 9.6 | -33.8 ± 5.2 | SS |
| E20<br>B5 | 91.5 ± 12.5 | -15.9 ± 2.9 | SS | 89.4 ± 13.4 | -16.6 ± 3.4 | SS | 87.9 ± 13.0 | -17.2 ± 3.6 | SS | 79.3 ± 11.1 | -25.4 ± 4.1 | SS | 75.5 ± 10.8 | -27.2 ± 4.3 | SS | 74.1 ± 11.0 | -27.9 ± 4.5 | SS |
| E20<br>H12.5 | 83.3 ± 11.1 | -24.1 ± 3.7 | SS | 81.1 ± 11.9 | -24.9 ± 4.3 | SS | 79.7 ± 11.4 | -25.3 ± 4.3 | SS | 72.3 ± 9.6 | -32.4 ± 4.6 | SS | 69.1 ± 9.0 | -33.6 ± 4.6 | SS | 67.7 ± 9.5 | -34.3 ± 5.1 | SS |
| L100<br>Aml5 | 82.3 ± 11.0 | -25.2 ± 4.0 | SS | 80.6 ± 12.0 | -25.4 ± 4.5 | SS | 79.4 ± 11.6 | -25.6 ± 4.5 | SS | 74.2 ± 10.0 | -30.6 ± 4.4 | SS | 71.3 ± 9.6 | -31.4 ± 4.3 | SS | 70.1 ± 9.7 | -32.0 ± 5.0 | SS |
| L100<br>B5 | 85.1 ± 11.8 | -22.3 ± 3.9 | SS | 84.8 ± 12.9 | -21.2 ± 4.2 | SS | 84.4 ± 12.5 | -20.6 ± 4.2 | SS | 80.7 ± 11.3 | -24.0 ± 3.9 | SS | 77.3 ± 11.0 | -25.4 ± 4.1 | SS | 76.1 ± 11.2 | -25.9 ± 4.3 | SS |
| L100<br>H12.5 | 80.5 ± 10.9 | -26.9 ± 4.1 | SS | 78.8 ± 11.6 | -27.1 ± 4.8 | SS | 77.8 ± 11.2 | -27.2 ± 4.6 | SS | 73.6 ± 9.7 | -31.1 ± 4.4 | SS | 71.0 ± 9.1 | -31.7 ± 4.4 | SS | 70.1 ± 9.7 | -31.9 ± 4.8 | SS |
| Al300<br>Aml5/B5 | 73.5 ± 10.1 | -33.9 ± 5.2 | SS | 72.8 ± 11.1 | -33.2 ± 5.7 | SS | 72.2 ± 10.7 | -32.8 ± 5.6 | SS | 68.0 ± 9.5 | -36.7 ± 5.3 | SS | 65.0 ± 9.3 | -37.6 ± 5.2 | SS | 63.9 ± 9.3 | -38.1 ± 5.9 | SS |
| Al300<br>Aml5/H12.5 | 67.8 ± 9.1 | -39.6 ± 5.8 | SS | 66.4 ± 9.7 | -39.6 ± 6.6 | SS | 65.6 ± 9.3 | -39.5 ± 6.5 | SS | 62.3 ± 8.1 | -42.4 ± 6.1 | SS | 60.3 ± 7.6 | -42.4 ± 5.8 | SS | 59.6 ± 8.0 | -42.4 ± 6.4 | SS |
| Al300<br>B5/H12.5 | 73.6 ± 10.2 | -33.8 ± 5.0 | SS | 72.5 ± 10.8 | -33.5 ± 5.8 | SS | 71.6 ± 10.5 | -33.4 ± 5.7 | SS | 67.0 ± 9.2 | -37.8 ± 5.5 | SS | 64.0 ± 8.8 | -38.7 ± 5.3 | SS | 63.0 ± 9.1 | -39.0 ± 5.9 | SS |
| E20<br>Aml5/B5 | 77.0 ± 10.5 | -30.4 ± 4.7 | SS | 75.1 ± 11.3 | -30.9 ± 5.3 | SS | 73.7 ± 10.9 | -31.3 ± 5.3 | SS | 65.9 ± 9.3 | -38.9 ± 5.6 | SS | 62.5 ± 9.1 | -40.2 ± 5.5 | SS | 61.2 ± 9.1 | -40.8 ± 6.2 | SS |
| E20<br>Aml5/H12.5 | 69.6 ± 9.2 | -37.8 ± 5.6 | SS | 67.8 ± 9.9 | -38.2 ± 6.4 | SS | 66.6 ± 9.4 | -38.5 ± 6.3 | SS | 60.0 ± 7.9 | -44.7 ± 6.4 | SS | 57.3 ± 7.4 | -45.3 ± 6.2 | SS | 56.2 ± 7.7 | -45.8 ± 6.9 | SS |
| E20<br>B5/H12.5 | 76.1 ± 10.5 | -31.3 ± 4.7 | SS | 74.1 ± 11.0 | -31.9 ± 5.6 | SS | 72.7 ± 10.7 | -32.3 ± 5.5 | SS | 65.2 ± 9.0 | -39.6 ± 5.7 | SS | 61.8 ± 8.6 | -40.8 ± 5.6 | SS | 60.6 ± 8.9 | -41.4 ± 6.2 | SS |
| L100<br>Aml5/B5 | 71.5 ± 9.9 | -35.9 ± 5.4 | SS | 71.1 ± 10.9 | -34.9 ± 5.9 | SS | 70.7 ± 10.5 | -34.3 ± 5.8 | SS | 67.1 ± 9.4 | -37.6 ± 5.4 | SS | 64.1 ± 9.2 | -38.5 ± 5.3 | SS | 63.1 ± 9.2 | -38.9 ± 6.0 | SS |
| L100<br>Aml5/H12.5 | 67.0 ± 9.0 | -40.4 ± 5.9 | SS | 65.6 ± 9.6 | -40.4 ± 6.8 | SS | 64.8 ± 9.2 | -40.3 ± 6.6 | SS | 61.2 ± 8.0 | -43.5 ± 6.2 | SS | 59.1 ± 7.5 | -43.6 ± 6.0 | SS | 58.4 ± 7.9 | -43.6 ± 6.6 | SS |
| L100<br>B5/H12.5 | 72.3 ± 10.1 | -35.1 ± 5.2 | SS | 71.3 ± 10.7 | -34.7 ± 6.0 | SS | 70.6 ± 10.4 | -34.5 ± 5.9 | SS | 66.2 ± 9.1 | -38.6 ± 5.6 | SS | 63.2 ± 8.7 | -39.5 ± 5.4 | SS | 62.2 ± 9.0 | -39.8 ± 6.0 | SS |

**Al300 =** aliskiren 300 mg; **Aml5** = amlodipine 5 mg; **B5** = bisoprolol 5 mg; **E20** = enalapril 20 mg; **H12.5** = hydrochlorothiazide 12.5 mg; **L100** = losartan 100 mg; **SS** = statistically significant ($P$ < 0.00001)

**Table S41.** *P*-values calculated using the Kolmogorov-Smirnov test for changes in <u>renal vascular resistance</u> in populations (*n* = 100) with different ACE activity receiving the same regimens (case 1: $c_{ACE}$ = 7.0 h$^{-1}$, case 2: $c_{ACE}$ = 8.9 h$^{-1}$, case 3: $c_{ACE}$ = 10.8 h$^{-1}$, case 4: $c_{ACE}$ = 42.3 h$^{-1}$, case 5: $c_{ACE}$ = 54.1 h$^{-1}$, case 6: $c_{ACE}$ = 65.9 h$^{-1}$; *P*-value for case *i vs.* case *j* is denoted $P_{ij}$)

| Regimens | $P_{12}$ | $P_{13}$ | $P_{23}$ | $P_{14}$ | $P_{15}$ | $P_{16}$ | $P_{24}$ | $P_{25}$ | $P_{26}$ | $P_{34}$ | $P_{35}$ | $P_{36}$ | $P_{45}$ | $P_{46}$ | $P_{56}$ |
|---|---|---|---|---|---|---|---|---|---|---|---|---|---|---|---|
| Al300 | 0.28093 | 0.00232 | 0.15454 | SS | SS | SS | SS | SS | SS | SS | SS | SS | 0.00136 | 0.00002 | 0.28093 |
| E20 | 0.01008 | SS | 0.00630 | SS | SS | SS | SS | SS | SS | SS | SS | SS | SS | SS | 0.21055 |
| L100 | 0.46756 | 0.00386 | 0.21055 | SS | SS | SS | SS | SS | SS | SS | SS | SS | 0.00136 | 0.00002 | 0.28093 |
| Aml5 | 0.81275 | 0.11113 | 0.69937 | 0.69937 | 0.36672 | 0.11113 | 0.69937 | 0.46756 | 0.11113 | 0.36672 | 0.11113 | 0.28093 | 0.81275 | 0.28093 | 0.36672 |
| B5 | 0.28093 | 0.00232 | 0.11113 | SS | SS | SS | SS | SS | SS | SS | SS | SS | 0.00079 | 0.00002 | 0.28093 |
| H12.5 | 0.11113 | 0.00386 | 0.15454 | SS | SS | SS | SS | SS | SS | SS | SS | SS | 0.11113 | 0.00045 | 0.02431 |
| Al300 Aml5 | 0.90621 | 0.69937 | 0.90621 | SS | SS | SS | SS | SS | SS | SS | SS | SS | 0.21055 | 0.15454 | 0.69937 |
| Al300 B5 | 0.58062 | 0.01581 | 0.28093 | SS | SS | SS | SS | SS | SS | SS | SS | SS | 0.03663 | 0.00232 | 0.81275 |
| Al300 H12.5 | 0.69937 | 0.96707 | 0.90621 | SS | SS | SS | SS | SS | SS | SS | SS | SS | 0.36672 | 0.81275 | 0.90621 |
| E20 Aml5 | 0.46756 | 0.07832 | 0.46756 | SS | SS | SS | SS | SS | SS | SS | SS | SS | 0.03663 | 0.01008 | 0.58062 |
| E20 B5 | 0.28093 | 0.03663 | 0.15454 | SS | SS | SS | SS | SS | SS | SS | SS | SS | 0.01581 | 0.00386 | 0.81275 |
| E20 H12.5 | 0.36672 | 0.11113 | 0.81275 | SS | SS | SS | SS | SS | SS | SS | SS | SS | 0.11113 | 0.07832 | 0.69937 |
| L100 Aml5 | 0.90621 | 0.81275 | 0.81275 | SS | SS | SS | SS | SS | SS | SS | SS | SS | 0.15454 | 0.11113 | 0.81275 |
| L100 B5 | 0.46756 | 0.01581 | 0.28093 | 0.00025 | SS | SS | SS | SS | SS | SS | SS | SS | 0.03663 | 0.02431 | 0.81275 |
| L100 H12.5 | 0.58062 | 0.90621 | 0.81275 | SS | SS | SS | SS | SS | SS | SS | SS | SS | 0.28093 | 0.58062 | 0.81275 |
| Al300 Aml5/B5 | 0.69937 | 0.05410 | 0.46756 | 0.00079 | 0.00002 | 0.00002 | 0.00013 | SS | SS | 0.00002 | SS | SS | 0.36672 | 0.28093 | 0.81275 |
| Al300 Aml5/H12.5 | 0.81275 | 0.46756 | 0.81275 | 0.00386 | 0.01581 | 0.02431 | 0.00136 | 0.00386 | 0.01008 | 0.00045 | 0.00045 | 0.00232 | 0.90621 | 0.90621 | 0.96707 |
| Al300 B5/H12.5 | 0.81275 | 0.46756 | 0.96707 | SS | SS | SS | SS | SS | SS | SS | SS | SS | 0.21055 | 0.28093 | 0.90621 |
| E20 Aml5/B5 | 0.81275 | 0.58062 | 0.96707 | SS | SS | SS | SS | SS | SS | SS | SS | SS | 0.21055 | 0.07832 | 0.81275 |
| E20 Aml5/H12.5 | 0.69937 | 0.46756 | 0.90621 | SS | SS | SS | SS | SS | SS | SS | SS | SS | 0.46756 | 0.69937 | 0.96707 |
| E20 B5/H12.5 | 0.46756 | 0.46756 | 0.90621 | SS | SS | SS | SS | SS | SS | SS | SS | SS | 0.15454 | 0.21055 | 0.90621 |
| L100 Aml5/B5 | 0.69937 | 0.05410 | 0.46756 | 0.07832 | 0.00232 | 0.00136 | 0.00386 | 0.00025 | 0.00045 | 0.00007 | SS | 0.00002 | 0.46756 | 0.36672 | 0.81275 |
| L100 Aml5/H12.5 | 0.81275 | 0.46756 | 0.69937 | 0.00232 | 0.01008 | 0.01581 | 0.00079 | 0.00232 | 0.00630 | 0.00013 | 0.00007 | 0.00136 | 0.81275 | 0.90621 | 0.96707 |
| L100 B5/H12.5 | 0.81275 | 0.28093 | 0.90621 | 0.00013 | SS | SS | 0.00002 | SS | SS | SS | SS | SS | 0.15454 | 0.36672 | 0.90621 |

**Al300 =** aliskiren 300 mg; **Aml5** = amlodipine 5 mg; **B5** = bisoprolol 5 mg; **E20** = enalapril 20 mg; **H12.5** = hydrochlorothiazide 12.5 mg; **L100** = losartan 100 mg; **SS** = statistically significant ($P < 0.00001$)

**Figure S28.** Simulated change in afferent arteriolar resistance from baseline to week 4 (mean ± SD, $n$ = 100)

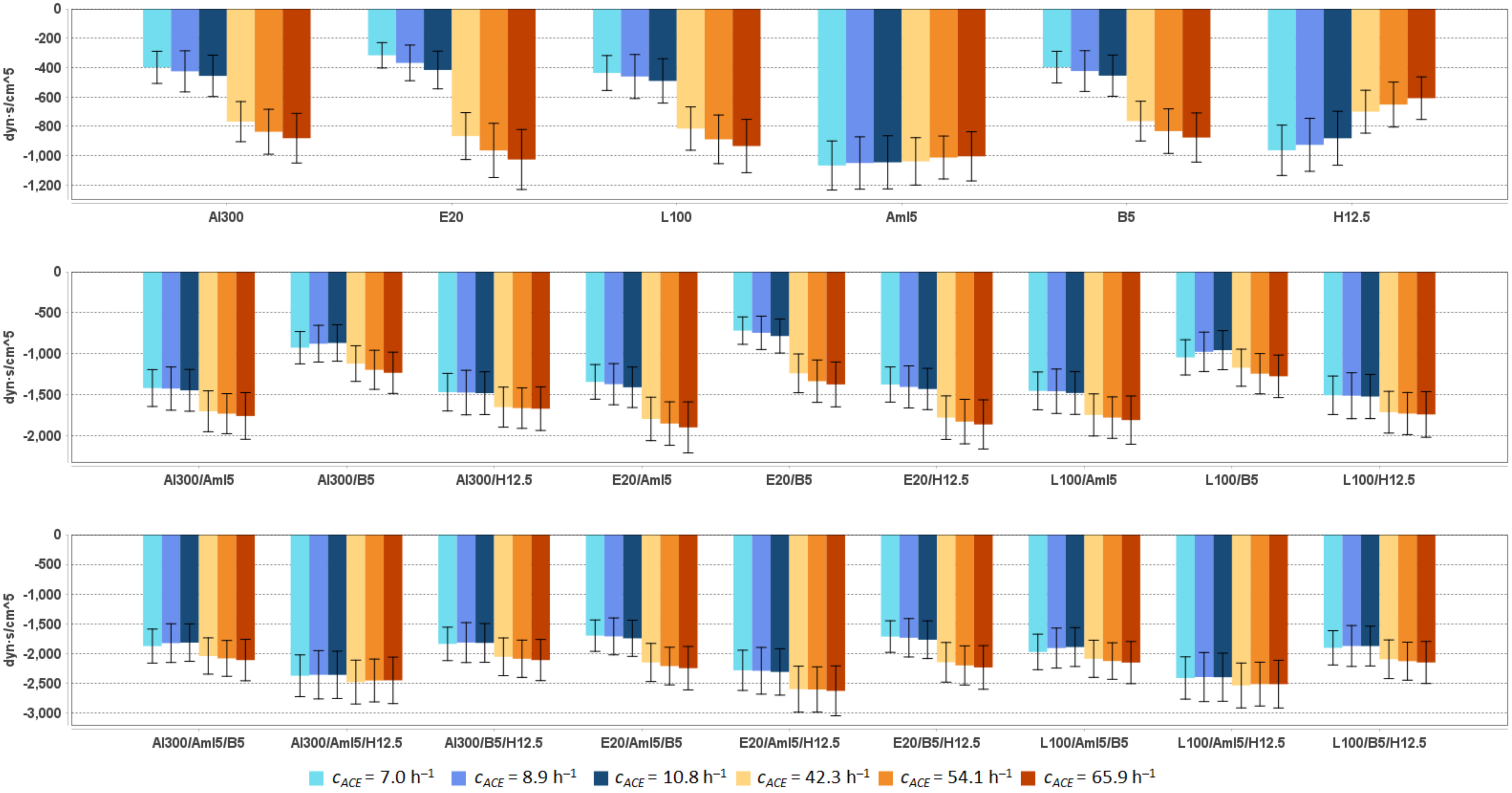


**Al300 =** aliskiren 300 mg; **Aml5** = amlodipine 5 mg; **B5** = bisoprolol 5 mg; **E20** = enalapril 20 mg; **H12.5** = hydrochlorothiazide 12.5 mg; **L100** = losartan 100 mg

**Table S42.** Simulated response of <u>afferent arteriolar resistance</u> to antihypertensive therapy in virtual hypertensive populations ($n$ = 100) with different ACE activity, including $P$-values (Kolmogorov-Smirnov test) for endpoint *vs*. baseline; data are presented as mean ± SD in dyn·s/cm$^5$

| Regimens | $c_{ACE}$ = 7.0 h$^{-1}$ | | | $c_{ACE}$ = 8.9 h$^{-1}$ | | | $c_{ACE}$ = 10.8 h$^{-1}$ | | | $c_{ACE}$ = 42.3 h$^{-1}$ | | | $c_{ACE}$ = 54.1 h$^{-1}$ | | | $c_{ACE}$ = 65.9 h$^{-1}$ | | |
|---|---|---|---|---|---|---|---|---|---|---|---|---|---|---|---|---|---|---|
| | **Value** | **Change** | ***P*** | **Value** | **Change** | ***P*** | **Value** | **Change** | ***P*** | **Value** | **Change** | ***P*** | **Value** | **Change** | ***P*** | **Value** | **Change** | ***P*** |
| Baseline | 5011 ± 814 | – | – | 4936 ± 864 | – | – | 4886 ± 859 | – | – | 4773 ± 756 | – | – | 4613 ± 739 | – | – | 4586 ± 778 | – | – |
| Al300 | 4613 ± 774 | -398 ± 109 | 0.01581 | 4511 ± 792 | -425 ± 140 | 0.00630 | 4430 ± 810 | -456 ± 141 | 0.00630 | 4006 ± 687 | -768 ± 136 | SS | 3776 ± 680 | -837 ± 153 | SS | 3706 ± 678 | -881 ± 169 | SS |
| E20 | 4695 ± 780 | -316 ± 86 | 0.05410 | 4569 ± 800 | -368 ± 121 | 0.01008 | 4470 ± 814 | -416 ± 128 | 0.01581 | 3907 ± 684 | -866 ± 160 | SS | 3649 ± 680 | -964 ± 185 | SS | 3561 ± 670 | -1026 ± 204 | SS |
| L100 | 4575 ± 771 | -437 ± 119 | 0.01008 | 4476 ± 787 | -461 ± 150 | 0.00232 | 4396 ± 807 | -491 ± 151 | 0.00386 | 3958 ± 685 | -815 ± 147 | SS | 3724 ± 680 | -888 ± 165 | SS | 3652 ± 675 | -934 ± 182 | SS |
| Aml5 | 3945 ± 665 | -1066 ± 167 | SS | 3887 ± 702 | -1049 ± 178 | SS | 3841 ± 697 | -1045 ± 181 | SS | 3735 ± 619 | -1038 ± 161 | SS | 3601 ± 617 | -1012 ± 146 | SS | 3582 ± 631 | -1004 ± 168 | SS |
| B5 | 4614 ± 774 | -397 ± 108 | 0.01581 | 4513 ± 792 | -424 ± 139 | 0.00630 | 4432 ± 810 | -455 ± 140 | 0.00630 | 4009 ± 687 | -764 ± 135 | SS | 3780 ± 679 | -833 ± 152 | SS | 3710 ± 678 | -876 ± 167 | SS |
| H12.5 | 4049 ± 671 | -963 ± 171 | SS | 4011 ± 710 | -926 ± 180 | SS | 4005 ± 707 | -881 ± 183 | SS | 4073 ± 653 | -700 ± 146 | SS | 3961 ± 628 | -652 ± 153 | 0.00002 | 3978 ± 696 | -608 ± 145 | SS |
| Al300<br>Aml5 | 3592 ± 639 | -1420 ± 225 | SS | 3510 ± 649 | -1426 ± 264 | SS | 3439 ± 662 | -1448 ± 255 | SS | 3070 ± 563 | -1703 ± 250 | SS | 2881 ± 566 | -1732 ± 244 | SS | 2826 ± 549 | -1760 ± 285 | SS |
| Al300<br>B5 | 4084 ± 727 | -927 ± 197 | SS | 4058 ± 738 | -879 ± 224 | SS | 4017 ± 761 | -869 ± 223 | SS | 3653 ± 678 | -1121 ± 217 | SS | 3415 ± 681 | -1197 ± 238 | SS | 3353 ± 666 | -1234 ± 252 | SS |
| Al300<br>H12.5 | 3541 ± 629 | -1470 ± 229 | SS | 3462 ± 626 | -1475 ± 272 | SS | 3404 ± 640 | -1482 ± 261 | SS | 3121 ± 549 | -1652 ± 244 | SS | 2948 ± 532 | -1664 ± 247 | SS | 2915 ± 550 | -1672 ± 266 | SS |
| E20<br>Aml5 | 3667 ± 643 | -1344 ± 211 | SS | 3564 ± 655 | -1373 ± 251 | SS | 3476 ± 664 | -1410 ± 247 | SS | 2977 ± 561 | -1796 ± 265 | SS | 2760 ± 566 | -1853 ± 265 | SS | 2687 ± 544 | -1900 ± 311 | SS |
| E20<br>B5 | 4293 ± 746 | -719 ± 167 | SS | 4191 ± 752 | -746 ± 203 | SS | 4102 ± 772 | -785 ± 208 | SS | 3533 ± 672 | -1240 ± 237 | SS | 3277 ± 678 | -1336 ± 258 | SS | 3211 ± 663 | -1375 ± 274 | SS |
| E20<br>H12.5 | 3635 ± 633 | -1377 ± 215 | SS | 3531 ± 634 | -1406 ± 257 | SS | 3455 ± 643 | -1431 ± 252 | SS | 2992 ± 542 | -1781 ± 266 | SS | 2784 ± 526 | -1829 ± 272 | SS | 2723 ± 535 | -1864 ± 300 | SS |
| L100<br>Aml5 | 3557 ± 638 | -1454 ± 231 | SS | 3478 ± 646 | -1459 ± 271 | SS | 3406 ± 659 | -1480 ± 261 | SS | 3026 ± 562 | -1748 ± 257 | SS | 2832 ± 566 | -1780 ± 253 | SS | 2776 ± 546 | -1811 ± 294 | SS |
| L100<br>B5 | 3966 ± 717 | -1045 ± 215 | SS | 3959 ± 729 | -978 ± 240 | SS | 3929 ± 751 | -957 ± 238 | SS | 3602 ± 676 | -1172 ± 226 | SS | 3368 ± 680 | -1244 ± 246 | SS | 3310 ± 665 | -1276 ± 259 | SS |
| L100<br>H12.5 | 3503 ± 628 | -1508 ± 235 | SS | 3423 ± 622 | -1513 ± 280 | SS | 3363 ± 637 | -1523 ± 270 | SS | 3059 ± 545 | -1715 ± 255 | SS | 2881 ± 529 | -1731 ± 257 | SS | 2844 ± 544 | -1742 ± 278 | SS |
| Al300<br>Aml5/B5 | 3137 ± 611 | -1875 ± 286 | SS | 3112 ± 613 | -1825 ± 325 | SS | 3070 ± 627 | -1816 ± 315 | SS | 2732 ± 557 | -2041 ± 305 | SS | 2531 ± 568 | -2081 ± 302 | SS | 2476 ± 543 | -2110 ± 348 | SS |
| Al300<br>Aml5/H12.5 | 2638 ± 516 | -2373 ± 351 | SS | 2579 ± 503 | -2358 ± 406 | SS | 2528 ± 510 | -2358 ± 398 | SS | 2294 ± 435 | -2480 ± 369 | SS | 2160 ± 425 | -2453 ± 360 | SS | 2136 ± 428 | -2451 ± 391 | SS |
| Al300<br>B5/H12.5 | 3174 ± 615 | -1838 ± 281 | SS | 3121 ± 597 | -1816 ± 336 | SS | 3065 ± 617 | -1821 ± 326 | SS | 2719 ± 537 | -2054 ± 317 | SS | 2525 ± 526 | -2087 ± 313 | SS | 2477 ± 527 | -2109 ± 346 | SS |
| E20<br>Aml5/B5 | 3312 ± 623 | -1699 ± 264 | SS | 3225 ± 623 | -1711 ± 311 | SS | 3143 ± 634 | -1743 ± 304 | SS | 2623 ± 553 | -2150 ± 320 | SS | 2402 ± 565 | -2211 ± 316 | SS | 2340 ± 542 | -2246 ± 366 | SS |
| E20<br>Aml5/H12.5 | 2729 ± 518 | -2282 ± 338 | SS | 2646 ± 509 | -2290 ± 393 | SS | 2577 ± 512 | -2310 ± 390 | SS | 2174 ± 429 | -2599 ± 387 | SS | 2008 ± 418 | -2605 ± 381 | SS | 1959 ± 417 | -2628 ± 420 | SS |
| E20<br>B5/H12.5 | 3297 ± 622 | -1715 ± 266 | SS | 3202 ± 603 | -1734 ± 323 | SS | 3119 ± 622 | -1767 ± 317 | SS | 2626 ± 536 | -2147 ± 333 | SS | 2412 ± 525 | -2200 ± 329 | SS | 2352 ± 526 | -2234 ± 367 | SS |
| L100<br>Aml5/B5 | 3038 ± 604 | -1973 ± 299 | SS | 3029 ± 606 | -1908 ± 336 | SS | 2995 ± 619 | -1891 ± 327 | SS | 2686 ± 555 | -2088 ± 312 | SS | 2486 ± 567 | -2126 ± 307 | SS | 2435 ± 543 | -2152 ± 354 | SS |
| L100<br>Aml5/H12.5 | 2601 ± 515 | -2411 ± 356 | SS | 2541 ± 501 | -2395 ± 413 | SS | 2488 ± 509 | -2399 ± 405 | SS | 2236 ± 432 | -2537 ± 377 | SS | 2099 ± 422 | -2514 ± 368 | SS | 2072 ± 423 | -2515 ± 401 | SS |
| L100<br>B5/H12.5 | 3107 ± 610 | -1904 ± 289 | SS | 3063 ± 593 | -1873 ± 344 | SS | 3012 ± 613 | -1874 ± 335 | SS | 2678 ± 537 | -2096 ± 325 | SS | 2484 ± 526 | -2129 ± 319 | SS | 2436 ± 526 | -2150 ± 353 | SS |

**Al300 =** aliskiren 300 mg; **Aml5** = amlodipine 5 mg; **B5** = bisoprolol 5 mg; **E20** = enalapril 20 mg; **H12.5** = hydrochlorothiazide 12.5 mg; **L100** = losartan 100 mg; **SS** = statistically significant ($P$ < 0.00001)

**Table S43.** *P*-values calculated using the Kolmogorov-Smirnov test for changes in afferent arteriolar resistance in populations ($n$ = 100) with different ACE activity receiving the same regimens (case 1: $c_{ACE}$ = 7.0 h$^{-1}$, case 2: $c_{ACE}$ = 8.9 h$^{-1}$, case 3: $c_{ACE}$ = 10.8 h$^{-1}$, case 4: $c_{ACE}$ = 42.3 h$^{-1}$, case 5: $c_{ACE}$ = 54.1 h$^{-1}$, case 6: $c_{ACE}$ = 65.9 h$^{-1}$; *P*-value for case *i* *vs*. case *j* is denoted $P_{ij}$)

| Regimens | $P_{12}$ | $P_{13}$ | $P_{23}$ | $P_{14}$ | $P_{15}$ | $P_{16}$ | $P_{24}$ | $P_{25}$ | $P_{26}$ | $P_{34}$ | $P_{35}$ | $P_{36}$ | $P_{45}$ | $P_{46}$ | $P_{56}$ |
|---|---|---|---|---|---|---|---|---|---|---|---|---|---|---|---|
| Al300 | 0.07832 | 0.00079 | 0.21055 | SS | SS | SS | SS | SS | SS | SS | SS | SS | 0.00079 | 0.00004 | 0.28093 |
| E20 | 0.01008 | SS | 0.01581 | SS | SS | SS | SS | SS | SS | SS | SS | SS | 0.00007 | SS | 0.11113 |
| L100 | 0.15454 | 0.00386 | 0.21055 | SS | SS | SS | SS | SS | SS | SS | SS | SS | 0.00232 | 0.00004 | 0.28093 |
| Aml5 | 0.69937 | 0.07832 | 0.58062 | 0.58062 | 0.28093 | 0.01008 | 0.46756 | 0.21055 | 0.11113 | 0.81275 | 0.15454 | 0.28093 | 0.58062 | 0.11113 | 0.46756 |
| B5 | 0.11113 | 0.00136 | 0.28093 | SS | SS | SS | SS | SS | SS | SS | SS | SS | 0.00136 | 0.00007 | 0.21055 |
| H12.5 | 0.21055 | 0.01008 | 0.15454 | SS | SS | SS | SS | SS | SS | SS | SS | SS | 0.07832 | 0.00079 | 0.05410 |
| Al300<br>Aml5 | 0.81275 | 0.69937 | 0.81275 | SS | SS | SS | SS | SS | SS | SS | SS | SS | 0.69937 | 0.58062 | 0.81275 |
| Al300<br>B5 | 0.46756 | 0.21055 | 0.96707 | SS | SS | SS | SS | SS | SS | SS | SS | SS | 0.01008 | 0.00136 | 0.69937 |
| Al300<br>H12.5 | 0.81275 | 0.81275 | 0.99376 | SS | SS | 0.00004 | 0.00004 | 0.00004 | 0.00002 | 0.00004 | 0.00007 | 0.00004 | 0.96707 | 0.58062 | 0.58062 |
| E20<br>Aml5 | 0.28093 | 0.11113 | 0.46756 | SS | SS | SS | SS | SS | SS | SS | SS | SS | 0.21055 | 0.21055 | 0.58062 |
| E20<br>B5 | 0.46756 | 0.03663 | 0.46756 | SS | SS | SS | SS | SS | SS | SS | SS | SS | 0.00386 | 0.00079 | 0.69937 |
| E20<br>H12.5 | 0.58062 | 0.28093 | 0.96707 | SS | SS | SS | SS | SS | SS | SS | SS | SS | 0.36672 | 0.46756 | 0.46756 |
| L100<br>Aml5 | 0.81275 | 0.69937 | 0.81275 | SS | SS | SS | SS | SS | SS | SS | SS | SS | 0.69937 | 0.58062 | 0.81275 |
| L100<br>B5 | 0.21055 | 0.05410 | 0.69937 | 0.00013 | SS | SS | SS | SS | SS | SS | SS | SS | 0.03663 | 0.00386 | 0.81275 |
| L100<br>H12.5 | 0.81275 | 0.69937 | 0.99376 | SS | SS | SS | SS | SS | SS | SS | 0.00002 | SS | 0.81275 | 0.90621 | 0.69937 |
| Al300<br>Aml5/B5 | 0.28093 | 0.21055 | 0.96707 | 0.00025 | SS | 0.00002 | 0.00004 | SS | SS | 0.00002 | SS | SS | 0.28093 | 0.46756 | 0.69937 |
| Al300<br>Aml5/H12.5 | 0.58062 | 0.28093 | 0.90621 | 0.11113 | 0.28093 | 0.46756 | 0.05410 | 0.21055 | 0.21055 | 0.00630 | 0.07832 | 0.05410 | 0.69937 | 0.21055 | 0.69937 |
| Al300<br>B5/H12.5 | 0.36672 | 0.46756 | 0.90621 | 0.00004 | SS | 0.00002 | SS | SS | SS | SS | SS | SS | 0.81275 | 0.81275 | 0.90621 |
| E20<br>Aml5/B5 | 0.69937 | 0.36672 | 0.58062 | SS | SS | SS | SS | SS | SS | SS | SS | SS | 0.21055 | 0.46756 | 0.81275 |
| E20<br>Aml5/H12.5 | 0.69937 | 0.58062 | 0.69937 | SS | SS | SS | SS | 0.00007 | 0.00002 | SS | 0.00002 | SS | 0.96707 | 0.99376 | 0.69937 |
| E20<br>B5/H12.5 | 0.58062 | 0.58062 | 0.36672 | SS | SS | SS | SS | SS | SS | SS | SS | SS | 0.46756 | 0.58062 | 0.81275 |
| L100<br>Aml5/B5 | 0.21055 | 0.15454 | 0.81275 | 0.00630 | 0.00079 | 0.00079 | 0.00136 | 0.00002 | 0.00013 | 0.00013 | 0.00002 | 0.00002 | 0.46756 | 0.46756 | 0.81275 |
| L100<br>Aml5/H12.5 | 0.69937 | 0.28093 | 0.90621 | 0.07832 | 0.21055 | 0.36672 | 0.01581 | 0.11113 | 0.15454 | 0.00386 | 0.03663 | 0.03663 | 0.81275 | 0.36672 | 0.69937 |
| L100<br>B5/H12.5 | 0.36672 | 0.28093 | 0.96707 | 0.00013 | 0.00002 | 0.00025 | SS | 0.00004 | SS | SS | SS | SS | 0.81275 | 0.81275 | 0.90621 |

**Al300 =** aliskiren 300 mg; **Aml5** = amlodipine 5 mg; **B5** = bisoprolol 5 mg; **E20** = enalapril 20 mg; **H12.5** = hydrochlorothiazide 12.5 mg; **L100** = losartan 100 mg; **SS** = statistically significant ($P$ < 0.00001)

**Figure S29.** Simulated change in <u>efferent arteriolar resistance</u> from baseline to week 4 (mean ± SD, $n$ = 100)

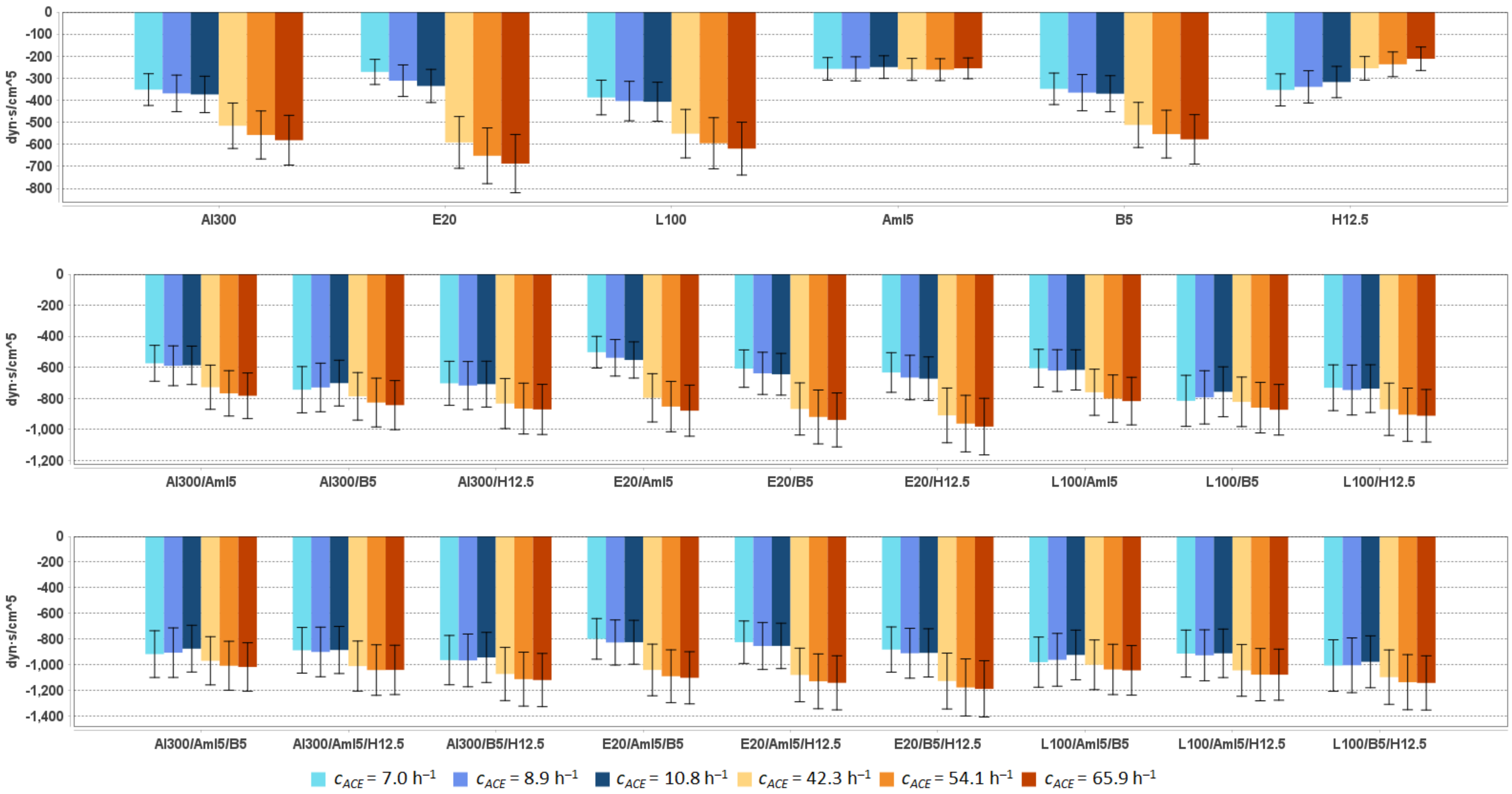


**Al300 =** aliskiren 300 mg; **Aml5** = amlodipine 5 mg; **B5** = bisoprolol 5 mg; **E20** = enalapril 20 mg; **H12.5** = hydrochlorothiazide 12.5 mg; **L100** = losartan 100 mg

**Table S44.** Simulated response of <u>efferent arteriolar resistance</u> to antihypertensive therapy in virtual hypertensive populations ($n$ = 100) with different ACE activity, including $P$-values (Kolmogorov-Smirnov test) for endpoint *vs*. baseline; data are presented as mean ± SD in dyn·s/cm$^5$

| Regimens | $c_{ACE}$ = 7.0 h$^{-1}$ | | | $c_{ACE}$ = 8.9 h$^{-1}$ | | | $c_{ACE}$ = 10.8 h$^{-1}$ | | | $c_{ACE}$ = 42.3 h$^{-1}$ | | | $c_{ACE}$ = 54.1 h$^{-1}$ | | | $c_{ACE}$ = 65.9 h$^{-1}$ | | |
|---|---|---|---|---|---|---|---|---|---|---|---|---|---|---|---|---|---|---|
| | **Value** | **Change** | ***P*** | **Value** | **Change** | ***P*** | **Value** | **Change** | ***P*** | **Value** | **Change** | ***P*** | **Value** | **Change** | ***P*** | **Value** | **Change** | ***P*** |
| Baseline | 2485 ± 493 | – | – | 2489 ± 530 | – | – | 2419 ± 498 | – | – | 2559 ± 491 | – | – | 2581 ± 491 | – | – | 2528 ± 466 | – | – |
| Al300 | 2133 ± 422 | -352 ± 72 | 0.00013 | 2120 ± 450 | -369 ± 83 | 0.00079 | 2045 ± 422 | -374 ± 82 | 0.00025 | 2043 ± 392 | -516 ± 103 | SS | 2023 ± 392 | -557 ± 109 | SS | 1947 ± 364 | -581 ± 113 | SS |
| E20 | 2213 ± 438 | -272 ± 57 | 0.00386 | 2178 ± 463 | -311 ± 72 | 0.00630 | 2083 ± 430 | -335 ± 75 | 0.00079 | 1967 ± 378 | -591 ± 117 | SS | 1929 ± 375 | -652 ± 126 | SS | 1841 ± 345 | -687 ± 132 | SS |
| L100 | 2097 ± 415 | -388 ± 79 | SS | 2085 ± 443 | -404 ± 90 | 0.00013 | 2012 ± 415 | -407 ± 89 | 0.00007 | 2007 ± 385 | -552 ± 110 | SS | 1986 ± 385 | -595 ± 116 | SS | 1909 ± 357 | -619 ± 120 | SS |
| Aml5 | 2227 ± 441 | -258 ± 51 | 0.00630 | 2232 ± 475 | -257 ± 55 | 0.02431 | 2169 ± 446 | -250 ± 51 | 0.01581 | 2299 ± 441 | -260 ± 50 | 0.00232 | 2319 ± 441 | -261 ± 50 | 0.00386 | 2273 ± 419 | -255 ± 47 | 0.00232 |
| B5 | 2137 ± 422 | -348 ± 71 | 0.00013 | 2124 ± 451 | -366 ± 82 | 0.00079 | 2048 ± 423 | -370 ± 82 | 0.00025 | 2047 ± 393 | -512 ± 102 | SS | 2027 ± 393 | -554 ± 108 | SS | 1951 ± 365 | -577 ± 112 | SS |
| H12.5 | 2132 ± 423 | -353 ± 73 | 0.00025 | 2150 ± 460 | -339 ± 73 | 0.00232 | 2101 ± 432 | -317 ± 71 | 0.00136 | 2304 ± 444 | -255 ± 54 | 0.00232 | 2344 ± 444 | -237 ± 56 | 0.01581 | 2316 ± 425 | -212 ± 53 | 0.01581 |
| Al300 Aml5 | 1911 ± 378 | -574 ± 116 | SS | 1899 ± 403 | -590 ± 128 | SS | 1832 ± 378 | -587 ± 124 | SS | 1831 ± 351 | -728 ± 142 | SS | 1813 ± 351 | -768 ± 146 | SS | 1745 ± 326 | -783 ± 147 | SS |
| Al300 B5 | 1741 ± 349 | -744 ± 149 | SS | 1759 ± 377 | -730 ± 156 | SS | 1717 ± 352 | -702 ± 147 | SS | 1772 ± 340 | -787 ± 153 | SS | 1754 ± 340 | -827 ± 157 | SS | 1685 ± 316 | -843 ± 158 | SS |
| Al300 H12.5 | 1782 ± 351 | -703 ± 142 | SS | 1772 ± 377 | -717 ± 155 | SS | 1710 ± 353 | -708 ± 148 | SS | 1725 ± 331 | -833 ± 161 | SS | 1715 ± 331 | -866 ± 164 | SS | 1657 ± 308 | -871 ± 161 | SS |
| E20 Aml5 | 1982 ± 392 | -502 ± 102 | SS | 1951 ± 414 | -539 ± 118 | SS | 1866 ± 385 | -552 ± 117 | SS | 1762 ± 338 | -796 ± 156 | SS | 1728 ± 336 | -853 ± 162 | SS | 1649 ± 309 | -879 ± 165 | SS |
| E20 B5 | 1877 ± 373 | -608 ± 121 | SS | 1850 ± 395 | -639 ± 136 | SS | 1774 ± 364 | -645 ± 134 | SS | 1691 ± 324 | -868 ± 168 | SS | 1661 ± 321 | -920 ± 174 | SS | 1589 ± 297 | -939 ± 174 | SS |
| E20 H12.5 | 1852 ± 365 | -633 ± 128 | SS | 1823 ± 388 | -666 ± 143 | SS | 1746 ± 360 | -673 ± 140 | SS | 1649 ± 316 | -910 ± 176 | SS | 1618 ± 313 | -962 ± 182 | SS | 1546 ± 289 | -982 ± 182 | SS |
| L100 Aml5 | 1879 ± 371 | -606 ± 122 | SS | 1868 ± 397 | -621 ± 135 | SS | 1802 ± 372 | -616 ± 129 | SS | 1798 ± 345 | -761 ± 149 | SS | 1779 ± 345 | -802 ± 153 | SS | 1710 ± 320 | -818 ± 153 | SS |
| L100 B5 | 1669 ± 337 | -816 ± 164 | SS | 1696 ± 366 | -793 ± 172 | SS | 1661 ± 341 | -758 ± 161 | SS | 1736 ± 333 | -822 ± 160 | SS | 1721 ± 333 | -860 ± 163 | SS | 1655 ± 310 | -873 ± 163 | SS |
| L100 H12.5 | 1753 ± 346 | -732 ± 147 | SS | 1743 ± 371 | -746 ± 161 | SS | 1681 ± 347 | -737 ± 154 | SS | 1688 ± 324 | -870 ± 169 | SS | 1676 ± 324 | -905 ± 171 | SS | 1616 ± 301 | -912 ± 169 | SS |
| Al300 Aml5/B5 | 1566 ± 313 | -919 ± 182 | SS | 1582 ± 339 | -907 ± 193 | SS | 1542 ± 317 | -876 ± 182 | SS | 1588 ± 304 | -971 ± 188 | SS | 1571 ± 305 | -1009 ± 191 | SS | 1509 ± 283 | -1019 ± 189 | SS |
| Al300 Aml5/H12.5 | 1596 ± 315 | -889 ± 178 | SS | 1587 ± 337 | -902 ± 193 | SS | 1532 ± 316 | -886 ± 184 | SS | 1547 ± 297 | -1012 ± 195 | SS | 1538 ± 296 | -1043 ± 197 | SS | 1486 ± 276 | -1042 ± 192 | SS |
| Al300 B5/H12.5 | 1519 ± 301 | -966 ± 192 | SS | 1521 ± 324 | -968 ± 206 | SS | 1474 ± 303 | -945 ± 195 | SS | 1485 ± 285 | -1074 ± 208 | SS | 1467 ± 285 | -1114 ± 211 | SS | 1407 ± 264 | -1121 ± 208 | SS |
| E20 Aml5/B5 | 1685 ± 335 | -800 ± 159 | SS | 1661 ± 354 | -828 ± 176 | SS | 1592 ± 327 | -826 ± 171 | SS | 1516 ± 290 | -1042 ± 201 | SS | 1489 ± 287 | -1091 ± 206 | SS | 1425 ± 266 | -1103 ± 203 | SS |
| E20 Aml5/H12.5 | 1659 ± 327 | -826 ± 166 | SS | 1634 ± 347 | -855 ± 183 | SS | 1564 ± 322 | -855 ± 177 | SS | 1477 ± 283 | -1081 ± 209 | SS | 1450 ± 281 | -1131 ± 214 | SS | 1385 ± 259 | -1143 ± 211 | SS |
| E20 B5/H12.5 | 1602 ± 316 | -883 ± 176 | SS | 1577 ± 336 | -912 ± 195 | SS | 1510 ± 310 | -909 ± 188 | SS | 1430 ± 274 | -1129 ± 218 | SS | 1402 ± 272 | -1179 ± 223 | SS | 1338 ± 251 | -1190 ± 220 | SS |
| L100 Aml5/B5 | 1504 ± 303 | -981 ± 196 | SS | 1526 ± 329 | -963 ± 206 | SS | 1493 ± 307 | -925 ± 194 | SS | 1557 ± 298 | -1002 ± 194 | SS | 1542 ± 299 | -1038 ± 196 | SS | 1483 ± 278 | -1045 ± 193 | SS |
| L100 Aml5/H12.5 | 1570 ± 310 | -915 ± 183 | SS | 1561 ± 332 | -928 ± 199 | SS | 1506 ± 311 | -913 ± 189 | SS | 1513 ± 290 | -1046 ± 202 | SS | 1502 ± 290 | -1079 ± 204 | SS | 1449 ± 270 | -1079 ± 199 | SS |
| L100 B5/H12.5 | 1477 ± 293 | -1008 ± 201 | SS | 1483 ± 317 | -1006 ± 214 | SS | 1440 ± 295 | -978 ± 202 | SS | 1460 ± 280 | -1098 ± 212 | SS | 1443 ± 280 | -1138 ± 215 | SS | 1384 ± 260 | -1144 ± 211 | SS |

**Al300 =** aliskiren 300 mg; **Aml5** = amlodipine 5 mg; **B5** = bisoprolol 5 mg; **E20** = enalapril 20 mg; **H12.5** = hydrochlorothiazide 12.5 mg; **L100** = losartan 100 mg; **SS** = statistically significant ($P$ < 0.00001)

**Table S45.** *P*-values calculated using the Kolmogorov-Smirnov test for changes in efferent arteriolar resistance in populations ($n$ = 100) with different ACE activity receiving the same regimens (case 1: $c_{ACE}$ = 7.0 $h^{-1}$, case 2: $c_{ACE}$ = 8.9 $h^{-1}$, case 3: $c_{ACE}$ = 10.8 $h^{-1}$, case 4: $c_{ACE}$ = 42.3 $h^{-1}$, case 5: $c_{ACE}$ = 54.1 $h^{-1}$, case 6: $c_{ACE}$ = 65.9 $h^{-1}$; *P*-value for case *i vs.* case *j* is denoted $P_{ij}$)

| **Regimens** | $P_{12}$ | $P_{13}$ | $P_{23}$ | $P_{14}$ | $P_{15}$ | $P_{16}$ | $P_{24}$ | $P_{25}$ | $P_{26}$ | $P_{34}$ | $P_{35}$ | $P_{36}$ | $P_{45}$ | $P_{46}$ | $P_{56}$ |
|---|---|---|---|---|---|---|---|---|---|---|---|---|---|---|---|
| Al300 | 0.28093 | 0.36672 | 0.90621 | SS | SS | SS | SS | SS | SS | SS | SS | SS | 0.01581 | 0.00007 | 0.28093 |
| E20 | 0.01008 | SS | 0.11113 | SS | SS | SS | SS | SS | SS | SS | SS | SS | 0.00386 | SS | 0.21055 |
| L100 | 0.36672 | 0.58062 | 0.99376 | SS | SS | SS | SS | SS | SS | SS | SS | SS | 0.01581 | 0.00013 | 0.36672 |
| Aml5 | 0.90621 | 0.28093 | 0.69937 | 0.90621 | 0.69937 | 0.69937 | 0.46756 | 0.36672 | 0.58062 | 0.07832 | 0.07832 | 0.11113 | 0.90621 | 0.69937 | 0.81275 |
| B5 | 0.28093 | 0.28093 | 0.90621 | SS | SS | SS | SS | SS | SS | SS | SS | SS | 0.01581 | 0.00007 | 0.28093 |
| H12.5 | 0.46756 | 0.01008 | 0.15454 | SS | SS | SS | SS | SS | SS | SS | SS | SS | 0.02431 | SS | 0.00386 |
| Al300<br>Aml5 | 0.46756 | 0.90621 | 0.96707 | SS | SS | SS | SS | SS | SS | SS | SS | SS | 0.05410 | 0.01008 | 0.81275 |
| Al300<br>B5 | 0.69937 | 0.05410 | 0.46756 | 0.15454 | 0.00136 | 0.00007 | 0.07832 | 0.00045 | 0.00079 | 0.00232 | SS | SS | 0.07832 | 0.01581 | 0.90621 |
| Al300<br>H12.5 | 0.69937 | 0.90621 | 0.96707 | SS | SS | SS | 0.00025 | SS | SS | 0.00004 | SS | SS | 0.21055 | 0.07832 | 0.96707 |
| E20<br>Aml5 | 0.11113 | 0.07832 | 0.96707 | SS | SS | SS | SS | SS | SS | SS | SS | SS | 0.01581 | 0.00025 | 0.69937 |
| E20<br>B5 | 0.21055 | 0.21055 | 0.96707 | SS | SS | SS | SS | SS | SS | SS | SS | SS | 0.02431 | 0.00136 | 0.81275 |
| E20<br>H12.5 | 0.21055 | 0.36672 | 0.96707 | SS | SS | SS | SS | SS | SS | SS | SS | SS | 0.03663 | 0.00232 | 0.90621 |
| L100<br>Aml5 | 0.69937 | 0.90621 | 0.96707 | SS | SS | SS | SS | SS | SS | SS | SS | SS | 0.05410 | 0.01008 | 0.81275 |
| L100<br>B5 | 0.46756 | 0.02431 | 0.46756 | 0.81275 | 0.07832 | 0.01008 | 0.46756 | 0.00630 | 0.01008 | 0.01581 | 0.00004 | 0.00013 | 0.07832 | 0.02431 | 0.90621 |
| L100<br>H12.5 | 0.81275 | 0.90621 | 0.96707 | SS | SS | SS | 0.00025 | SS | SS | 0.00004 | SS | SS | 0.21055 | 0.07832 | 0.96707 |
| Al300<br>Aml5/B5 | 0.69937 | 0.11113 | 0.58062 | 0.15454 | 0.00386 | 0.00025 | 0.05410 | 0.00136 | 0.00630 | 0.00386 | SS | 0.00004 | 0.15454 | 0.02431 | 0.96707 |
| Al300<br>Aml5/H12.5 | 0.81275 | 0.81275 | 0.90621 | 0.00079 | SS | SS | 0.00232 | 0.00007 | 0.00045 | 0.00013 | SS | 0.00002 | 0.28093 | 0.28093 | 0.96707 |
| Al300<br>B5/H12.5 | 0.90621 | 0.36672 | 0.69937 | 0.01008 | 0.00004 | 0.00002 | 0.00630 | 0.00025 | 0.00025 | 0.00045 | SS | SS | 0.21055 | 0.05410 | 0.96707 |
| E20<br>Aml5/B5 | 0.36672 | 0.46756 | 0.96707 | SS | SS | SS | SS | SS | SS | SS | SS | SS | 0.05410 | 0.02431 | 0.99376 |
| E20<br>Aml5/H12.5 | 0.46756 | 0.58062 | 0.96707 | SS | SS | SS | SS | SS | SS | SS | SS | SS | 0.07832 | 0.03663 | 0.99376 |
| E20<br>B5/H12.5 | 0.46756 | 0.58062 | 0.96707 | SS | SS | SS | SS | SS | SS | SS | SS | SS | 0.11113 | 0.05410 | 0.99376 |
| L100<br>Aml5/B5 | 0.69937 | 0.03663 | 0.36672 | 0.81275 | 0.05410 | 0.02431 | 0.21055 | 0.00630 | 0.02431 | 0.01581 | 0.00007 | 0.00079 | 0.15454 | 0.11113 | 0.96707 |
| L100<br>Aml5/H12.5 | 0.81275 | 0.81275 | 0.90621 | 0.00079 | SS | SS | 0.00232 | 0.00007 | 0.00025 | 0.00013 | SS | SS | 0.28093 | 0.15454 | 0.99376 |
| L100<br>B5/H12.5 | 0.81275 | 0.28093 | 0.69937 | 0.03663 | 0.00025 | 0.00007 | 0.01581 | 0.00045 | 0.00232 | 0.00045 | SS | 0.00007 | 0.21055 | 0.11113 | 0.99376 |

**Al300 =** aliskiren 300 mg; **Aml5** = amlodipine 5 mg; **B5** = bisoprolol 5 mg; **E20** = enalapril 20 mg; **H12.5** = hydrochlorothiazide 12.5 mg; **L100** = losartan 100 mg; **SS** = statistically significant ($P$ < 0.00001)

**Figure S30.** Simulated change in glomerular hydrostatic pressure from baseline to week 4 (mean ± SD, $n$ = 100)

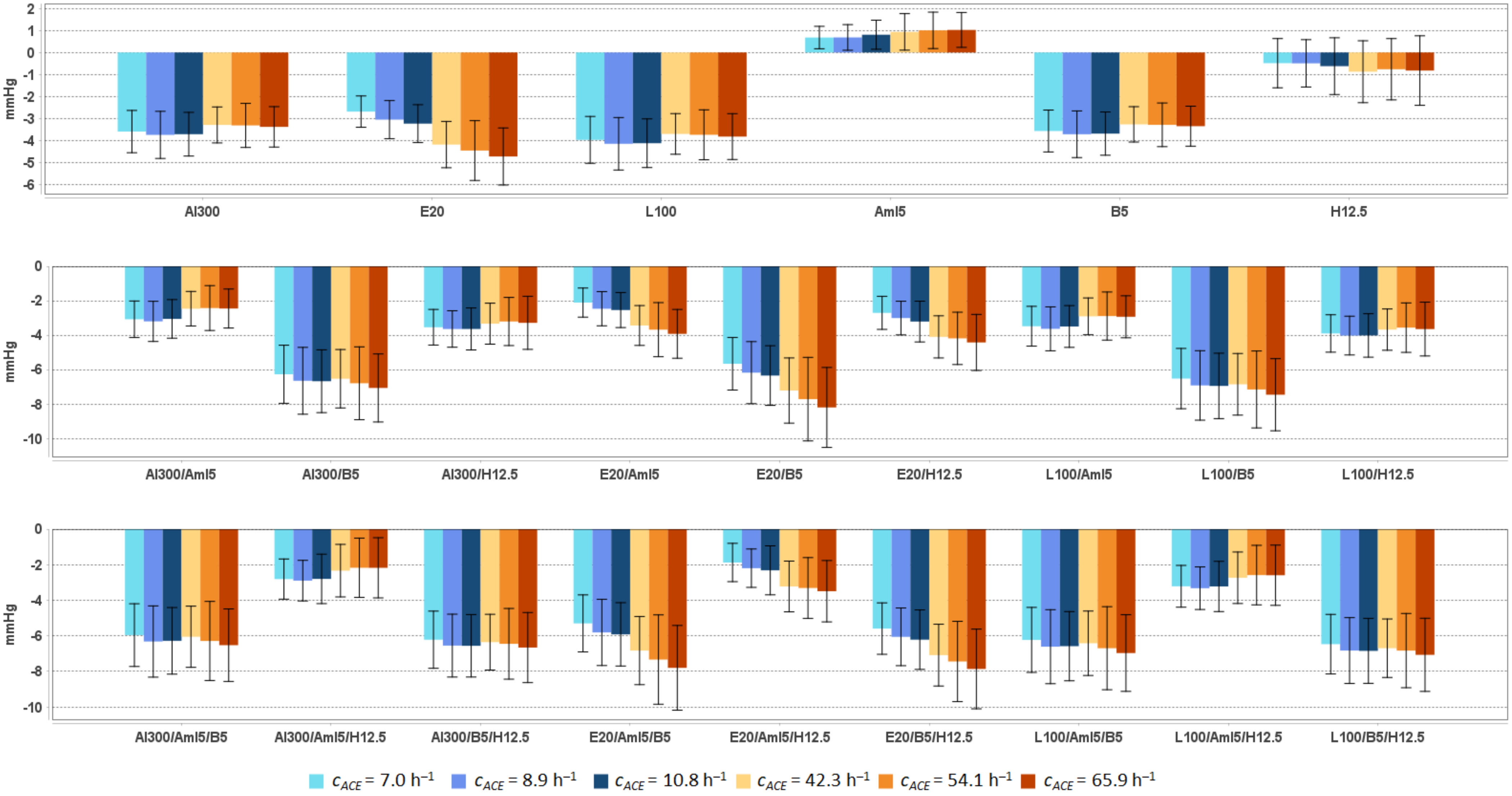


**Al300 =** aliskiren 300 mg; **Aml5** = amlodipine 5 mg; **B5** = bisoprolol 5 mg; **E20** = enalapril 20 mg; **H12.5** = hydrochlorothiazide 12.5 mg; **L100** = losartan 100 mg

**Table S46.** Simulated response of glomerular hydrostatic pressure to antihypertensive therapy in virtual hypertensive populations (*n* = 100) with different ACE activity, including *P*-values (Kolmogorov-Smirnov test) for endpoint *vs*. baseline; data are presented as mean ± SD in mmHg

| Regimens | $c_{ACE}$ = 7.0 h$^{-1}$ | | | $c_{ACE}$ = 8.9 h$^{-1}$ | | | $c_{ACE}$ = 10.8 h$^{-1}$ | | | $c_{ACE}$ = 42.3 h$^{-1}$ | | | $c_{ACE}$ = 54.1 h$^{-1}$ | | | $c_{ACE}$ = 65.9 h$^{-1}$ | | |
|---|---|---|---|---|---|---|---|---|---|---|---|---|---|---|---|---|---|---|
| | **Value** | **Change** | ***P*** | **Value** | **Change** | ***P*** | **Value** | **Change** | ***P*** | **Value** | **Change** | ***P*** | **Value** | **Change** | ***P*** | **Value** | **Change** | ***P*** |
| Baseline | 51.5 ± 2.8 | – | – | 51.9 ± 3.0 | – | – | 52.0 ± 3.0 | – | – | 52.9 ± 3.3 | – | – | 53.7 ± 4.0 | – | – | 54.0 ± 3.6 | – | – |
| Al300 | 47.9 ± 2.4 | -3.6 ± 1.0 | SS | 48.2 ± 2.4 | -3.7 ± 1.1 | SS | 48.3 ± 2.6 | -3.7 ± 1.0 | SS | 49.6 ± 2.9 | -3.3 ± 0.8 | SS | 50.4 ± 3.4 | -3.3 ± 1.0 | SS | 50.6 ± 3.2 | -3.4 ± 0.9 | SS |
| E20 | 48.8 ± 2.5 | -2.7 ± 0.7 | SS | 48.9 ± 2.5 | -3.1 ± 0.9 | SS | 48.8 ± 2.7 | -3.2 ± 0.9 | SS | 48.7 ± 2.8 | -4.2 ± 1.1 | SS | 49.2 ± 3.3 | -4.5 ± 1.4 | SS | 49.3 ± 3.0 | -4.7 ± 1.3 | SS |
| L100 | 47.5 ± 2.4 | -4.0 ± 1.1 | SS | 47.8 ± 2.4 | -4.2 ± 1.2 | SS | 47.9 ± 2.6 | -4.1 ± 1.1 | SS | 49.2 ± 2.9 | -3.7 ± 0.9 | SS | 50.0 ± 3.4 | -3.7 ± 1.1 | SS | 50.2 ± 3.1 | -3.8 ± 1.0 | SS |
| Aml5 | 52.2 ± 2.9 | 0.7 ± 0.5 | 0.07832 | 52.6 ± 3.1 | 0.7 ± 0.6 | 0.21055 | 52.8 ± 3.2 | 0.8 ± 0.7 | 0.28093 | 53.9 ± 3.4 | 0.9 ± 0.8 | 0.07832 | 54.7 ± 4.1 | 1.0 ± 0.8 | 0.15454 | 55.0 ± 3.8 | 1.0 ± 0.8 | 0.15454 |
| B5 | 48.0 ± 2.4 | -3.6 ± 1.0 | SS | 48.2 ± 2.4 | -3.7 ± 1.1 | SS | 48.3 ± 2.6 | -3.7 ± 1.0 | SS | 49.7 ± 2.9 | -3.3 ± 0.8 | SS | 50.4 ± 3.4 | -3.3 ± 1.0 | SS | 50.6 ± 3.2 | -3.4 ± 0.9 | SS |
| H12.5 | 51.0 ± 3.1 | -0.5 ± 1.1 | 0.11113 | 51.4 ± 3.4 | -0.5 ± 1.1 | 0.15454 | 51.4 ± 3.4 | -0.6 ± 1.3 | 0.05410 | 52.0 ± 3.6 | -0.9 ± 1.4 | 0.07832 | 52.9 ± 4.2 | -0.8 ± 1.4 | 0.28093 | 53.2 ± 3.9 | -0.8 ± 1.6 | 0.15454 |
| Al300 Aml5 | 48.4 ± 2.5 | -3.1 ± 1.1 | SS | 48.7 ± 2.5 | -3.2 ± 1.2 | SS | 49.0 ± 2.8 | -3.1 ± 1.1 | SS | 50.5 ± 3.0 | -2.5 ± 1.0 | SS | 51.3 ± 3.6 | -2.4 ± 1.3 | 0.00079 | 51.5 ± 3.3 | -2.5 ± 1.1 | 0.00045 |
| Al300 B5 | 45.3 ± 2.3 | -6.3 ± 1.7 | SS | 45.3 ± 2.2 | -6.6 ± 1.9 | SS | 45.3 ± 2.6 | -6.7 ± 1.8 | SS | 46.4 ± 2.7 | -6.5 ± 1.7 | SS | 46.9 ± 3.0 | -6.8 ± 2.1 | SS | 46.9 ± 2.8 | -7.1 ± 2.0 | SS |
| Al300 H12.5 | 48.0 ± 2.6 | -3.5 ± 1.0 | SS | 48.3 ± 2.7 | -3.6 ± 1.1 | SS | 48.4 ± 2.9 | -3.6 ± 1.2 | SS | 49.6 ± 3.2 | -3.3 ± 1.2 | SS | 50.5 ± 3.7 | -3.2 ± 1.4 | SS | 50.7 ± 3.5 | -3.3 ± 1.5 | SS |
| E20 Aml5 | 49.4 ± 2.6 | -2.1 ± 0.8 | SS | 49.5 ± 2.6 | -2.5 ± 1.0 | SS | 49.5 ± 2.9 | -2.5 ± 1.0 | SS | 49.5 ± 2.9 | -3.4 ± 1.2 | SS | 50.0 ± 3.4 | -3.7 ± 1.6 | SS | 50.1 ± 3.1 | -3.9 ± 1.4 | SS |
| E20 B5 | 45.9 ± 2.3 | -5.7 ± 1.5 | SS | 45.7 ± 2.2 | -6.2 ± 1.8 | SS | 45.7 ± 2.5 | -6.3 ± 1.7 | SS | 45.7 ± 2.7 | -7.2 ± 1.9 | SS | 46.0 ± 2.9 | -7.7 ± 2.4 | SS | 45.8 ± 2.8 | -8.2 ± 2.3 | SS |
| E20 H12.5 | 48.8 ± 2.7 | -2.7 ± 1.0 | SS | 48.9 ± 2.8 | -3.0 ± 1.0 | SS | 48.8 ± 3.0 | -3.2 ± 1.2 | SS | 48.8 ± 3.1 | -4.1 ± 1.2 | SS | 49.5 ± 3.6 | -4.2 ± 1.5 | SS | 49.6 ± 3.4 | -4.4 ± 1.6 | SS |
| L100 Aml5 | 48.0 ± 2.4 | -3.5 ± 1.2 | SS | 48.3 ± 2.4 | -3.6 ± 1.3 | SS | 48.5 ± 2.8 | -3.5 ± 1.2 | SS | 50.0 ± 3.0 | -2.9 ± 1.1 | SS | 50.8 ± 3.6 | -2.9 ± 1.4 | 0.00007 | 51.1 ± 3.3 | -2.9 ± 1.2 | SS |
| L100 B5 | 45.0 ± 2.3 | -6.5 ± 1.8 | SS | 45.0 ± 2.2 | -6.9 ± 2.0 | SS | 45.1 ± 2.6 | -6.9 ± 1.9 | SS | 46.1 ± 2.7 | -6.9 ± 1.8 | SS | 46.5 ± 2.9 | -7.2 ± 2.2 | SS | 46.5 ± 2.8 | -7.5 ± 2.1 | SS |
| L100 H12.5 | 47.6 ± 2.5 | -3.9 ± 1.1 | SS | 47.9 ± 2.6 | -4.0 ± 1.1 | SS | 48.0 ± 2.9 | -4.0 ± 1.3 | SS | 49.2 ± 3.1 | -3.7 ± 1.2 | SS | 50.1 ± 3.7 | -3.6 ± 1.4 | SS | 50.3 ± 3.5 | -3.6 ± 1.6 | SS |
| Al300 Aml5/B5 | 45.6 ± 2.3 | -6.0 ± 1.8 | SS | 45.6 ± 2.2 | -6.3 ± 2.0 | SS | 45.7 ± 2.6 | -6.3 ± 1.9 | SS | 46.9 ± 2.7 | -6.0 ± 1.7 | SS | 47.4 ± 3.1 | -6.3 ± 2.2 | SS | 47.5 ± 2.9 | -6.5 ± 2.0 | SS |
| Al300 Aml5/H12.5 | 48.7 ± 2.7 | -2.8 ± 1.1 | SS | 49.0 ± 2.8 | -2.9 ± 1.1 | SS | 49.2 ± 3.1 | -2.8 ± 1.4 | SS | 50.6 ± 3.3 | -2.3 ± 1.5 | SS | 51.5 ± 4.0 | -2.2 ± 1.7 | 0.00136 | 51.8 ± 3.7 | -2.2 ± 1.7 | 0.00136 |
| Al300 B5/H12.5 | 45.3 ± 2.3 | -6.2 ± 1.6 | SS | 45.4 ± 2.3 | -6.5 ± 1.8 | SS | 45.4 ± 2.7 | -6.6 ± 1.8 | SS | 46.6 ± 2.8 | -6.4 ± 1.6 | SS | 47.2 ± 3.2 | -6.4 ± 2.0 | SS | 47.3 ± 3.1 | -6.7 ± 2.0 | SS |
| E20 Aml5/B5 | 46.2 ± 2.3 | -5.3 ± 1.6 | SS | 46.1 ± 2.2 | -5.8 ± 1.9 | SS | 46.1 ± 2.6 | -5.9 ± 1.8 | SS | 46.1 ± 2.7 | -6.8 ± 1.9 | SS | 46.4 ± 3.0 | -7.3 ± 2.5 | SS | 46.2 ± 2.9 | -7.8 ± 2.4 | SS |
| E20 Aml5/H12.5 | 49.6 ± 2.8 | -1.9 ± 1.1 | SS | 49.7 ± 2.9 | -2.2 ± 1.1 | SS | 49.7 ± 3.2 | -2.3 ± 1.4 | SS | 49.7 ± 3.2 | -3.2 ± 1.4 | SS | 50.4 ± 3.7 | -3.3 ± 1.7 | SS | 50.5 ± 3.5 | -3.5 ± 1.7 | SS |
| E20 B5/H12.5 | 45.9 ± 2.4 | -5.6 ± 1.4 | SS | 45.9 ± 2.3 | -6.1 ± 1.6 | SS | 45.8 ± 2.7 | -6.2 ± 1.7 | SS | 45.8 ± 2.7 | -7.1 ± 1.7 | SS | 46.3 ± 3.0 | -7.4 ± 2.3 | SS | 46.1 ± 3.0 | -7.9 ± 2.2 | SS |
| L100 Aml5/B5 | 45.3 ± 2.3 | -6.2 ± 1.8 | SS | 45.3 ± 2.2 | -6.6 ± 2.1 | SS | 45.4 ± 2.6 | -6.6 ± 1.9 | SS | 46.5 ± 2.7 | -6.4 ± 1.8 | SS | 47.0 ± 3.0 | -6.7 ± 2.3 | SS | 47.0 ± 2.9 | -7.0 ± 2.2 | SS |
| L100 Aml5/H12.5 | 48.3 ± 2.6 | -3.2 ± 1.2 | SS | 48.6 ± 2.7 | -3.3 ± 1.2 | SS | 48.8 ± 3.1 | -3.2 ± 1.4 | SS | 50.2 ± 3.3 | -2.7 ± 1.4 | SS | 51.1 ± 3.9 | -2.6 ± 1.7 | 0.00007 | 51.4 ± 3.6 | -2.6 ± 1.7 | 0.00025 |
| L100 B5/H12.5 | 45.1 ± 2.3 | -6.5 ± 1.7 | SS | 45.1 ± 2.3 | -6.8 ± 1.9 | SS | 45.2 ± 2.7 | -6.8 ± 1.8 | SS | 46.2 ± 2.7 | -6.7 ± 1.6 | SS | 46.9 ± 3.1 | -6.8 ± 2.1 | SS | 46.9 ± 3.0 | -7.1 ± 2.1 | SS |

**Al300 =** aliskiren 300 mg; **Aml5** = amlodipine 5 mg; **B5** = bisoprolol 5 mg; **E20** = enalapril 20 mg; **H12.5** = hydrochlorothiazide 12.5 mg; **L100** = losartan 100 mg; **SS** = statistically significant (*P* < 0.00001)

**Table S47.** $P$-values calculated using the Kolmogorov-Smirnov test for changes in glomerular hydrostatic pressure in populations ($n$ = 100) with different ACE activity receiving the same regimens (case 1: $c_{ACE}$ = 7.0 h$^{-1}$, case 2: $c_{ACE}$ = 8.9 h$^{-1}$, case 3: $c_{ACE}$ = 10.8 h$^{-1}$, case 4: $c_{ACE}$ = 42.3 h$^{-1}$, case 5: $c_{ACE}$ = 54.1 h$^{-1}$, case 6: $c_{ACE}$ = 65.9 h$^{-1}$; $P$-value for case $i$ *vs*. case $j$ is denoted $P_{ij}$)

| **Regimens** | $P_{12}$ | $P_{13}$ | $P_{23}$ | $P_{14}$ | $P_{15}$ | $P_{16}$ | $P_{24}$ | $P_{25}$ | $P_{26}$ | $P_{34}$ | $P_{35}$ | $P_{36}$ | $P_{45}$ | $P_{46}$ | $P_{56}$ |
|---|---|---|---|---|---|---|---|---|---|---|---|---|---|---|---|
| Al300 | 0.46756 | 0.69937 | 0.96707 | 0.15454 | 0.15454 | 0.36672 | 0.01581 | 0.07832 | 0.21055 | 0.01008 | 0.02431 | 0.11113 | 0.90621 | 0.46756 | 0.81275 |
| E20 | 0.03663 | 0.00013 | 0.15454 | SS | SS | SS | SS | SS | SS | SS | SS | SS | 0.46756 | 0.01581 | 0.21055 |
| L100 | 0.46756 | 0.58062 | 0.96707 | 0.36672 | 0.21055 | 0.46756 | 0.02431 | 0.21055 | 0.36672 | 0.03663 | 0.05410 | 0.28093 | 0.90621 | 0.46756 | 0.81275 |
| Aml5 | 0.96707 | 0.36672 | 0.15454 | 0.21055 | 0.01008 | 0.00079 | 0.15454 | 0.00232 | 0.00079 | 0.58062 | 0.05410 | 0.00630 | 0.36672 | 0.11113 | 0.58062 |
| B5 | 0.58062 | 0.69937 | 0.90621 | 0.15454 | 0.11113 | 0.28093 | 0.01581 | 0.07832 | 0.21055 | 0.01008 | 0.02431 | 0.11113 | 0.90621 | 0.46756 | 0.90621 |
| H12.5 | 0.46756 | 0.01581 | 0.15454 | 0.01008 | 0.15454 | 0.15454 | 0.05410 | 0.28093 | 0.03663 | 0.36672 | 0.81275 | 0.58062 | 0.46756 | 0.11113 | 0.90621 |
| Al300 Aml5 | 0.46756 | 0.90621 | 0.90621 | 0.00079 | 0.00013 | 0.00079 | 0.00004 | 0.00007 | 0.00045 | 0.00007 | 0.00013 | 0.00045 | 0.69937 | 0.90621 | 0.96707 |
| Al300 B5 | 0.21055 | 0.21055 | 0.58062 | 0.46756 | 0.15454 | 0.01581 | 0.69937 | 0.69937 | 0.03663 | 0.58062 | 0.96707 | 0.21055 | 0.69937 | 0.11113 | 0.28093 |
| Al300 H12.5 | 0.46756 | 0.46756 | 0.81275 | 0.28093 | 0.21055 | 0.01008 | 0.11113 | 0.05410 | 0.00136 | 0.15454 | 0.05410 | 0.00630 | 0.69937 | 0.15454 | 0.46756 |
| E20 Aml5 | 0.01581 | 0.00079 | 0.46756 | SS | SS | SS | SS | SS | SS | SS | SS | SS | 0.28093 | 0.00232 | 0.02431 |
| E20 B5 | 0.15454 | 0.02431 | 0.28093 | SS | SS | SS | 0.00025 | 0.00002 | SS | 0.01008 | 0.00079 | SS | 0.46756 | 0.01008 | 0.21055 |
| E20 H12.5 | 0.00386 | 0.00045 | 0.36672 | SS | SS | SS | SS | SS | SS | SS | SS | SS | 0.46756 | 0.58062 | 0.46756 |
| L100 Aml5 | 0.28093 | 0.90621 | 0.96707 | 0.00232 | 0.00136 | 0.01008 | 0.00025 | 0.00079 | 0.00386 | 0.00079 | 0.00079 | 0.00386 | 0.69937 | 0.81275 | 0.58062 |
| L100 B5 | 0.21055 | 0.21055 | 0.58062 | 0.28093 | 0.07832 | 0.00386 | 0.58062 | 0.58062 | 0.03663 | 0.58062 | 0.99376 | 0.11113 | 0.58062 | 0.11113 | 0.28093 |
| L100 H12.5 | 0.46756 | 0.58062 | 0.90621 | 0.58062 | 0.28093 | 0.01581 | 0.11113 | 0.11113 | 0.00232 | 0.15454 | 0.07832 | 0.02431 | 0.96707 | 0.21055 | 0.58062 |
| Al300 Aml5/B5 | 0.28093 | 0.28093 | 0.99376 | 0.96707 | 0.69937 | 0.03663 | 0.58062 | 0.90621 | 0.15454 | 0.46756 | 0.90621 | 0.36672 | 0.58062 | 0.07832 | 0.28093 |
| Al300 Aml5/H12.5 | 0.46756 | 0.58062 | 0.58062 | 0.05410 | 0.01581 | 0.00630 | 0.00136 | 0.00025 | 0.00025 | 0.05410 | 0.00630 | 0.00232 | 0.46756 | 0.21055 | 0.96707 |
| Al300 B5/H12.5 | 0.46756 | 0.21055 | 0.90621 | 0.90621 | 0.46756 | 0.15454 | 0.69937 | 0.90621 | 0.81275 | 0.15454 | 0.90621 | 0.90621 | 0.81275 | 0.15454 | 0.90621 |
| E20 Aml5/B5 | 0.05410 | 0.01008 | 0.81275 | SS | SS | SS | 0.00079 | 0.00013 | SS | 0.00386 | 0.00079 | SS | 0.36672 | 0.00630 | 0.11113 |
| E20 Aml5/H12.5 | 0.00232 | SS | 0.28093 | SS | SS | SS | SS | SS | SS | 0.00007 | SS | SS | 0.81275 | 0.69937 | 0.81275 |
| E20 B5/H12.5 | 0.21055 | 0.01008 | 0.69937 | SS | SS | SS | 0.00007 | 0.00002 | SS | 0.00386 | 0.00232 | 0.00004 | 0.36672 | 0.00630 | 0.46756 |
| L100 Aml5/B5 | 0.28093 | 0.21055 | 0.90621 | 0.69937 | 0.46756 | 0.01581 | 0.58062 | 0.96707 | 0.07832 | 0.69937 | 0.96707 | 0.15454 | 0.58062 | 0.07832 | 0.21055 |
| L100 Aml5/H12.5 | 0.36672 | 0.58062 | 0.46756 | 0.02431 | 0.02431 | 0.00630 | 0.00045 | 0.00079 | 0.00025 | 0.05410 | 0.01581 | 0.00136 | 0.90621 | 0.36672 | 0.96707 |
| L100 B5/H12.5 | 0.36672 | 0.11113 | 0.90621 | 0.58062 | 0.28093 | 0.05410 | 0.58062 | 0.69937 | 0.36672 | 0.15454 | 0.81275 | 0.58062 | 0.58062 | 0.11113 | 0.81275 |

**Al300 =** aliskiren 300 mg; **Aml5** = amlodipine 5 mg; **B5** = bisoprolol 5 mg; **E20** = enalapril 20 mg; **H12.5** = hydrochlorothiazide 12.5 mg; **L100** = losartan 100 mg; **SS** = statistically significant ($P$ < 0.00001)

**Figure S31.** Simulated change in plasma sodium from baseline to week 4 (mean ± SD, $n$ = 100)

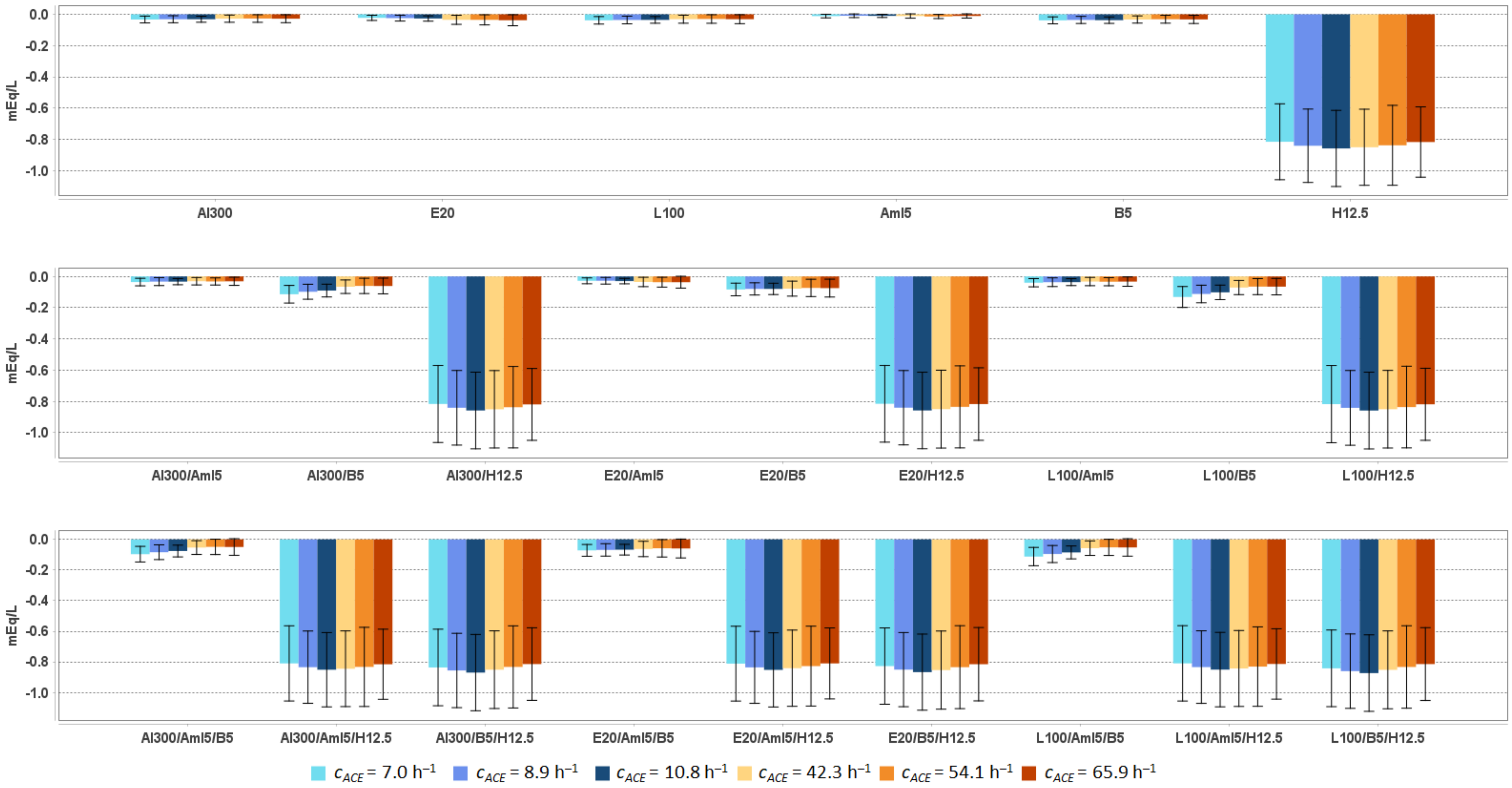


**Al300 =** aliskiren 300 mg; **Aml5** = amlodipine 5 mg; **B5** = bisoprolol 5 mg; **E20** = enalapril 20 mg; **H12.5** = hydrochlorothiazide 12.5 mg; **L100** = losartan 100 mg

**Table S48.** Simulated response of plasma sodium to antihypertensive therapy in virtual hypertensive populations (*n* = 100) with different ACE activity, including *P*-values (Kolmogorov-Smirnov test) for endpoint *vs*. baseline; data are presented as mean ± SD in mEq/L

| Regimens | $c_{ACE}$ = 7.0 h$^{-1}$ | | | $c_{ACE}$ = 8.9 h$^{-1}$ | | | $c_{ACE}$ = 10.8 h$^{-1}$ | | | $c_{ACE}$ = 42.3 h$^{-1}$ | | | $c_{ACE}$ = 54.1 h$^{-1}$ | | | $c_{ACE}$ = 65.9 h$^{-1}$ | | |
|---|---|---|---|---|---|---|---|---|---|---|---|---|---|---|---|---|---|---|
| | **Value** | **Change** | ***P*** | **Value** | **Change** | ***P*** | **Value** | **Change** | ***P*** | **Value** | **Change** | ***P*** | **Value** | **Change** | ***P*** | **Value** | **Change** | ***P*** |
| Baseline | 143.62 ± 1.29 | – | – | 143.65 ± 1.24 | – | – | 143.72 ± 1.27 | – | – | 143.47 ± 1.37 | – | – | 143.38 ± 1.34 | – | – | 143.45 ± 1.40 | – | – |
| Al300 | 143.59 ± 1.28 | -0.03 ± 0.02 | 0.99963 | 143.62 ± 1.24 | -0.03 ± 0.02 | 1.00000 | 143.69 ± 1.27 | -0.03 ± 0.02 | 0.99963 | 143.44 ± 1.37 | -0.03 ± 0.02 | 1.00000 | 143.35 ± 1.34 | -0.03 ± 0.02 | 0.99963 | 143.43 ± 1.40 | -0.03 ± 0.03 | 1.00000 |
| E20 | 143.60 ± 1.28 | -0.02 ± 0.02 | 1.00000 | 143.62 ± 1.24 | -0.02 ± 0.02 | 1.00000 | 143.69 ± 1.27 | -0.03 ± 0.02 | 1.00000 | 143.43 ± 1.37 | -0.04 ± 0.03 | 0.99963 | 143.34 ± 1.34 | -0.04 ± 0.03 | 0.99376 | 143.42 ± 1.40 | -0.04 ± 0.04 | 1.00000 |
| L100 | 143.58 ± 1.28 | -0.04 ± 0.02 | 0.99963 | 143.61 ± 1.24 | -0.04 ± 0.02 | 0.99963 | 143.68 ± 1.27 | -0.04 ± 0.02 | 0.99376 | 143.44 ± 1.37 | -0.03 ± 0.03 | 0.99963 | 143.35 ± 1.34 | -0.03 ± 0.03 | 0.99963 | 143.42 ± 1.40 | -0.03 ± 0.03 | 1.00000 |
| Aml5 | 143.61 ± 1.29 | -0.01 ± 0.01 | 1.00000 | 143.64 ± 1.24 | -0.01 ± 0.01 | 1.00000 | 143.71 ± 1.28 | -0.01 ± 0.01 | 1.00000 | 143.46 ± 1.38 | -0.01 ± 0.01 | 1.00000 | 143.36 ± 1.34 | -0.01 ± 0.01 | 1.00000 | 143.44 ± 1.40 | -0.01 ± 0.01 | 1.00000 |
| B5 | 143.58 ± 1.28 | -0.04 ± 0.02 | 0.99963 | 143.61 ± 1.24 | -0.04 ± 0.02 | 1.00000 | 143.68 ± 1.27 | -0.04 ± 0.02 | 0.99963 | 143.44 ± 1.37 | -0.03 ± 0.02 | 0.99963 | 143.35 ± 1.34 | -0.03 ± 0.03 | 0.99963 | 143.42 ± 1.40 | -0.03 ± 0.03 | 1.00000 |
| H12.5 | 142.81 ± 1.28 | -0.82 ± 0.24 | 0.00013 | 142.81 ± 1.30 | -0.84 ± 0.23 | 0.00025 | 142.86 ± 1.25 | -0.86 ± 0.24 | SS | 142.62 ± 1.36 | -0.85 ± 0.24 | 0.00045 | 142.54 ± 1.41 | -0.84 ± 0.26 | 0.00232 | 142.64 ± 1.39 | -0.82 ± 0.22 | 0.00025 |
| Al300 Aml5 | 143.59 ± 1.28 | -0.04 ± 0.02 | 1.00000 | 143.62 ± 1.25 | -0.03 ± 0.03 | 1.00000 | 143.69 ± 1.27 | -0.03 ± 0.02 | 0.99963 | 143.44 ± 1.37 | -0.03 ± 0.02 | 1.00000 | 143.35 ± 1.34 | -0.03 ± 0.02 | 1.00000 | 143.42 ± 1.40 | -0.03 ± 0.03 | 1.00000 |
| Al300 B5 | 143.51 ± 1.28 | -0.11 ± 0.06 | 0.90621 | 143.55 ± 1.24 | -0.10 ± 0.05 | 0.81275 | 143.63 ± 1.27 | -0.09 ± 0.04 | 0.90621 | 143.40 ± 1.36 | -0.07 ± 0.04 | 0.99376 | 143.32 ± 1.34 | -0.06 ± 0.05 | 0.96707 | 143.39 ± 1.40 | -0.06 ± 0.05 | 0.99963 |
| Al300 H12.5 | 142.80 ± 1.27 | -0.82 ± 0.25 | 0.00013 | 142.81 ± 1.30 | -0.84 ± 0.24 | 0.00025 | 142.86 ± 1.25 | -0.86 ± 0.25 | SS | 142.62 ± 1.36 | -0.85 ± 0.25 | 0.00045 | 142.54 ± 1.41 | -0.84 ± 0.26 | 0.00232 | 142.64 ± 1.39 | -0.82 ± 0.23 | 0.00045 |
| E20 Aml5 | 143.59 ± 1.28 | -0.03 ± 0.02 | 1.00000 | 143.62 ± 1.24 | -0.03 ± 0.02 | 1.00000 | 143.69 ± 1.27 | -0.03 ± 0.02 | 0.99963 | 143.43 ± 1.37 | -0.04 ± 0.03 | 0.99963 | 143.34 ± 1.34 | -0.04 ± 0.03 | 0.99963 | 143.42 ± 1.40 | -0.04 ± 0.04 | 1.00000 |
| E20 B5 | 143.54 ± 1.28 | -0.08 ± 0.04 | 0.96707 | 143.57 ± 1.24 | -0.08 ± 0.04 | 0.81275 | 143.64 ± 1.27 | -0.08 ± 0.04 | 0.96707 | 143.39 ± 1.36 | -0.08 ± 0.05 | 0.96707 | 143.30 ± 1.34 | -0.07 ± 0.06 | 0.96707 | 143.38 ± 1.40 | -0.07 ± 0.06 | 0.99963 |
| E20 H12.5 | 142.81 ± 1.27 | -0.81 ± 0.25 | 0.00013 | 142.81 ± 1.30 | -0.84 ± 0.24 | 0.00025 | 142.86 ± 1.25 | -0.86 ± 0.24 | SS | 142.62 ± 1.36 | -0.85 ± 0.25 | 0.00045 | 142.54 ± 1.41 | -0.83 ± 0.26 | 0.00232 | 142.64 ± 1.39 | -0.82 ± 0.23 | 0.00045 |
| L100 Aml5 | 143.58 ± 1.28 | -0.04 ± 0.03 | 0.99963 | 143.61 ± 1.25 | -0.04 ± 0.03 | 1.00000 | 143.68 ± 1.27 | -0.04 ± 0.02 | 0.99376 | 143.44 ± 1.37 | -0.03 ± 0.03 | 1.00000 | 143.34 ± 1.34 | -0.03 ± 0.03 | 1.00000 | 143.42 ± 1.40 | -0.03 ± 0.03 | 1.00000 |
| L100 B5 | 143.49 ± 1.28 | -0.13 ± 0.07 | 0.81275 | 143.54 ± 1.24 | -0.11 ± 0.06 | 0.81275 | 143.62 ± 1.26 | -0.10 ± 0.05 | 0.90621 | 143.40 ± 1.36 | -0.07 ± 0.05 | 0.99376 | 143.31 ± 1.34 | -0.07 ± 0.05 | 0.96707 | 143.39 ± 1.40 | -0.07 ± 0.05 | 0.99963 |
| L100 H12.5 | 142.80 ± 1.27 | -0.82 ± 0.25 | 0.00013 | 142.81 ± 1.30 | -0.84 ± 0.24 | 0.00025 | 142.86 ± 1.25 | -0.86 ± 0.25 | SS | 142.62 ± 1.36 | -0.85 ± 0.25 | 0.00045 | 142.54 ± 1.41 | -0.84 ± 0.26 | 0.00232 | 142.64 ± 1.39 | -0.82 ± 0.23 | 0.00045 |
| Al300 Aml5/B5 | 143.52 ± 1.28 | -0.10 ± 0.05 | 0.96707 | 143.57 ± 1.25 | -0.08 ± 0.05 | 0.81275 | 143.64 ± 1.27 | -0.08 ± 0.04 | 0.96707 | 143.42 ± 1.36 | -0.05 ± 0.05 | 0.99963 | 143.33 ± 1.34 | -0.05 ± 0.05 | 0.99376 | 143.40 ± 1.40 | -0.05 ± 0.05 | 0.99963 |
| Al300 Aml5/H12.5 | 142.81 ± 1.28 | -0.81 ± 0.25 | 0.00025 | 142.81 ± 1.31 | -0.83 ± 0.24 | 0.00025 | 142.87 ± 1.25 | -0.85 ± 0.24 | SS | 142.62 ± 1.36 | -0.84 ± 0.25 | 0.00045 | 142.55 ± 1.41 | -0.83 ± 0.26 | 0.00232 | 142.64 ± 1.39 | -0.82 ± 0.23 | 0.00045 |
| Al300 B5/H12.5 | 142.78 ± 1.27 | -0.84 ± 0.25 | 0.00004 | 142.79 ± 1.31 | -0.86 ± 0.24 | 0.00025 | 142.85 ± 1.25 | -0.87 ± 0.25 | SS | 142.62 ± 1.36 | -0.85 ± 0.25 | 0.00045 | 142.55 ± 1.41 | -0.83 ± 0.27 | 0.00232 | 142.64 ± 1.40 | -0.81 ± 0.24 | 0.00045 |
| E20 Aml5/B5 | 143.55 ± 1.28 | -0.07 ± 0.04 | 0.96707 | 143.58 ± 1.25 | -0.07 ± 0.04 | 0.81275 | 143.65 ± 1.27 | -0.07 ± 0.03 | 0.96707 | 143.41 ± 1.36 | -0.06 ± 0.05 | 0.99376 | 143.32 ± 1.34 | -0.06 ± 0.06 | 0.99376 | 143.39 ± 1.40 | -0.06 ± 0.06 | 0.99963 |
| E20 Aml5/H12.5 | 142.81 ± 1.28 | -0.81 ± 0.24 | 0.00025 | 142.81 ± 1.31 | -0.84 ± 0.23 | 0.00025 | 142.87 ± 1.25 | -0.85 ± 0.24 | SS | 142.63 ± 1.36 | -0.84 ± 0.25 | 0.00045 | 142.55 ± 1.41 | -0.83 ± 0.26 | 0.00232 | 142.64 ± 1.40 | -0.81 ± 0.23 | 0.00079 |
| E20 B5/H12.5 | 142.79 ± 1.27 | -0.83 ± 0.25 | 0.00007 | 142.80 ± 1.31 | -0.85 ± 0.24 | 0.00025 | 142.85 ± 1.25 | -0.87 ± 0.25 | SS | 142.62 ± 1.36 | -0.85 ± 0.26 | 0.00045 | 142.54 ± 1.41 | -0.83 ± 0.27 | 0.00232 | 142.64 ± 1.40 | -0.82 ± 0.24 | 0.00045 |
| L100 Aml5/B5 | 143.51 ± 1.28 | -0.11 ± 0.06 | 0.90621 | 143.55 ± 1.25 | -0.10 ± 0.06 | 0.81275 | 143.63 ± 1.27 | -0.09 ± 0.04 | 0.96707 | 143.41 ± 1.36 | -0.06 ± 0.05 | 0.99963 | 143.33 ± 1.34 | -0.05 ± 0.05 | 0.99376 | 143.40 ± 1.40 | -0.05 ± 0.06 | 0.99963 |
| L100 Aml5/H12.5 | 142.81 ± 1.28 | -0.81 ± 0.25 | 0.00025 | 142.81 ± 1.31 | -0.83 ± 0.24 | 0.00025 | 142.87 ± 1.26 | -0.85 ± 0.24 | SS | 142.63 ± 1.36 | -0.84 ± 0.25 | 0.00045 | 142.55 ± 1.41 | -0.83 ± 0.26 | 0.00232 | 142.64 ± 1.39 | -0.81 ± 0.23 | 0.00045 |
| L100 B5/H12.5 | 142.78 ± 1.28 | -0.84 ± 0.25 | 0.00004 | 142.79 ± 1.31 | -0.86 ± 0.24 | 0.00025 | 142.85 ± 1.25 | -0.87 ± 0.25 | SS | 142.62 ± 1.36 | -0.85 ± 0.25 | 0.00045 | 142.54 ± 1.41 | -0.83 ± 0.27 | 0.00232 | 142.64 ± 1.40 | -0.81 ± 0.24 | 0.00045 |

**Al300** = aliskiren 300 mg; **Aml5** = amlodipine 5 mg; **B5** = bisoprolol 5 mg; **E20** = enalapril 20 mg; **H12.5** = hydrochlorothiazide 12.5 mg; **L100** = losartan 100 mg; **SS** = statistically significant (*P* < 0.00001)

**Table S49.** $P$-values calculated using the Kolmogorov-Smirnov test for changes in plasma sodium in populations ($n$ = 100) with different ACE activity receiving the same regimens (case 1: $c_{ACE}$ = 7.0 h$^{-1}$, case 2: $c_{ACE}$ = 8.9 h$^{-1}$, case 3: $c_{ACE}$ = 10.8 h$^{-1}$, case 4: $c_{ACE}$ = 42.3 h$^{-1}$, case 5: $c_{ACE}$ = 54.1 h$^{-1}$, case 6: $c_{ACE}$ = 65.9 h$^{-1}$; $P$-value for case $i$ *vs*. case $j$ is denoted $P_{ij}$)

| **Regimens** | $P_{12}$ | $P_{13}$ | $P_{23}$ | $P_{14}$ | $P_{15}$ | $P_{16}$ | $P_{24}$ | $P_{25}$ | $P_{26}$ | $P_{34}$ | $P_{35}$ | $P_{36}$ | $P_{45}$ | $P_{46}$ | $P_{56}$ |
|---|---|---|---|---|---|---|---|---|---|---|---|---|---|---|---|
| Al300 | 0.90621 | 0.58062 | 0.90621 | 0.11113 | 0.01581 | 0.21055 | 0.21055 | 0.07832 | 0.46756 | 0.07832 | 0.01581 | 0.28093 | 0.81275 | 0.69937 | 0.28093 |
| E20 | 0.58062 | 0.28093 | 0.58062 | 0.00079 | 0.00007 | 0.00002 | 0.02431 | 0.00232 | 0.00025 | 0.03663 | 0.00232 | 0.00079 | 0.81275 | 0.69937 | 0.15454 |
| L100 | 0.96707 | 0.46756 | 0.81275 | 0.05410 | 0.00386 | 0.15454 | 0.15454 | 0.05410 | 0.28093 | 0.05410 | 0.00630 | 0.28093 | 0.81275 | 0.69937 | 0.28093 |
| Aml5 | 0.21055 | 0.69937 | 0.69937 | 0.46756 | 0.28093 | 0.58062 | 0.46756 | 0.07832 | 0.46756 | 0.58062 | 0.07832 | 0.90621 | 0.28093 | 0.96707 | 0.11113 |
| B5 | 0.81275 | 0.81275 | 0.90621 | 0.21055 | 0.01581 | 0.15454 | 0.36672 | 0.15454 | 0.58062 | 0.21055 | 0.02431 | 0.11113 | 0.81275 | 0.81275 | 0.58062 |
| H12.5 | 0.36672 | 0.15454 | 0.69937 | 0.46756 | 0.69937 | 0.81275 | 0.81275 | 0.99376 | 0.81275 | 0.90621 | 0.58062 | 0.28093 | 0.90621 | 0.28093 | 0.90621 |
| Al300<br>Aml5 | 0.58062 | 0.69937 | 0.58062 | 0.15454 | 0.15454 | 0.28093 | 0.58062 | 0.69937 | 0.81275 | 0.21055 | 0.15454 | 0.81275 | 0.96707 | 0.69937 | 0.90621 |
| Al300<br>B5 | 0.11113 | 0.11113 | 0.28093 | SS | SS | SS | 0.00025 | SS | 0.00013 | 0.00136 | SS | 0.00025 | 0.36672 | 0.96707 | 0.28093 |
| Al300<br>H12.5 | 0.58062 | 0.11113 | 0.69937 | 0.36672 | 0.69937 | 0.81275 | 0.69937 | 0.96707 | 0.90621 | 0.81275 | 0.58062 | 0.36672 | 0.69937 | 0.46756 | 0.90621 |
| E20<br>Aml5 | 0.90621 | 0.36672 | 0.36672 | 0.00386 | 0.00232 | 0.00045 | 0.03663 | 0.01581 | 0.00232 | 0.01008 | 0.01008 | 0.00136 | 0.96707 | 0.58062 | 0.69937 |
| E20<br>B5 | 0.58062 | 0.90621 | 0.90621 | 0.11113 | 0.00386 | 0.05410 | 0.58062 | 0.03663 | 0.36672 | 0.21055 | 0.00232 | 0.07832 | 0.46756 | 0.90621 | 0.46756 |
| E20<br>H12.5 | 0.58062 | 0.15454 | 0.69937 | 0.36672 | 0.69937 | 0.81275 | 0.81275 | 0.96707 | 0.81275 | 0.81275 | 0.58062 | 0.28093 | 0.69937 | 0.46756 | 0.81275 |
| L100<br>Aml5 | 0.58062 | 0.58062 | 0.58062 | 0.02431 | 0.05410 | 0.21055 | 0.36672 | 0.21055 | 0.46756 | 0.07832 | 0.05410 | 0.46756 | 0.99376 | 0.58062 | 0.58062 |
| L100<br>B5 | 0.07832 | 0.03663 | 0.21055 | SS | SS | SS | 0.00013 | SS | SS | 0.00025 | SS | 0.00004 | 0.28093 | 0.81275 | 0.36672 |
| L100<br>H12.5 | 0.58062 | 0.11113 | 0.69937 | 0.36672 | 0.69937 | 0.81275 | 0.81275 | 0.96707 | 0.90621 | 0.81275 | 0.58062 | 0.28093 | 0.69937 | 0.46756 | 0.81275 |
| Al300<br>Aml5/B5 | 0.05410 | 0.01008 | 0.36672 | SS | SS | SS | 0.00079 | SS | 0.00136 | 0.00013 | SS | 0.00025 | 0.58062 | 0.96707 | 0.28093 |
| Al300<br>Aml5/H12.5 | 0.46756 | 0.07832 | 0.69937 | 0.36672 | 0.58062 | 0.81275 | 0.90621 | 0.90621 | 0.96707 | 0.81275 | 0.58062 | 0.36672 | 0.69937 | 0.58062 | 0.69937 |
| Al300<br>B5/H12.5 | 0.69937 | 0.11113 | 0.46756 | 0.81275 | 0.96707 | 0.69937 | 0.96707 | 0.81275 | 0.36672 | 0.58062 | 0.36672 | 0.11113 | 0.69937 | 0.36672 | 0.81275 |
| E20<br>Aml5/B5 | 0.58062 | 0.90621 | 0.90621 | 0.00630 | 0.00013 | 0.00630 | 0.07832 | 0.00386 | 0.11113 | 0.02431 | 0.00045 | 0.02431 | 0.46756 | 0.96707 | 0.36672 |
| E20<br>Aml5/H12.5 | 0.36672 | 0.07832 | 0.69937 | 0.46756 | 0.69937 | 0.90621 | 0.90621 | 0.96707 | 0.81275 | 0.81275 | 0.58062 | 0.28093 | 0.58062 | 0.46756 | 0.81275 |
| E20<br>B5/H12.5 | 0.58062 | 0.07832 | 0.58062 | 0.36672 | 0.90621 | 0.81275 | 0.81275 | 0.81275 | 0.58062 | 0.58062 | 0.36672 | 0.11113 | 0.69937 | 0.46756 | 0.90621 |
| L100<br>Aml5/B5 | 0.03663 | 0.00386 | 0.46756 | SS | SS | SS | 0.00013 | SS | 0.00013 | 0.00007 | SS | 0.00013 | 0.46756 | 0.96707 | 0.36672 |
| L100<br>Aml5/H12.5 | 0.46756 | 0.07832 | 0.69937 | 0.46756 | 0.58062 | 0.81275 | 0.90621 | 0.96707 | 0.90621 | 0.90621 | 0.58062 | 0.36672 | 0.69937 | 0.58062 | 0.81275 |
| L100<br>B5/H12.5 | 0.81275 | 0.11113 | 0.36672 | 0.81275 | 0.96707 | 0.69937 | 0.96707 | 0.81275 | 0.36672 | 0.46756 | 0.28093 | 0.11113 | 0.69937 | 0.36672 | 0.81275 |

**Al300 =** aliskiren 300 mg; **Aml5** = amlodipine 5 mg; **B5** = bisoprolol 5 mg; **E20** = enalapril 20 mg; **H12.5** = hydrochlorothiazide 12.5 mg; **L100** = losartan 100 mg; **SS** = statistically significant ($P$ < 0.00001)

**Figure S32.** Simulated change in plasma renin activity from baseline to week 4 (mean ± SD, $n$ = 100)

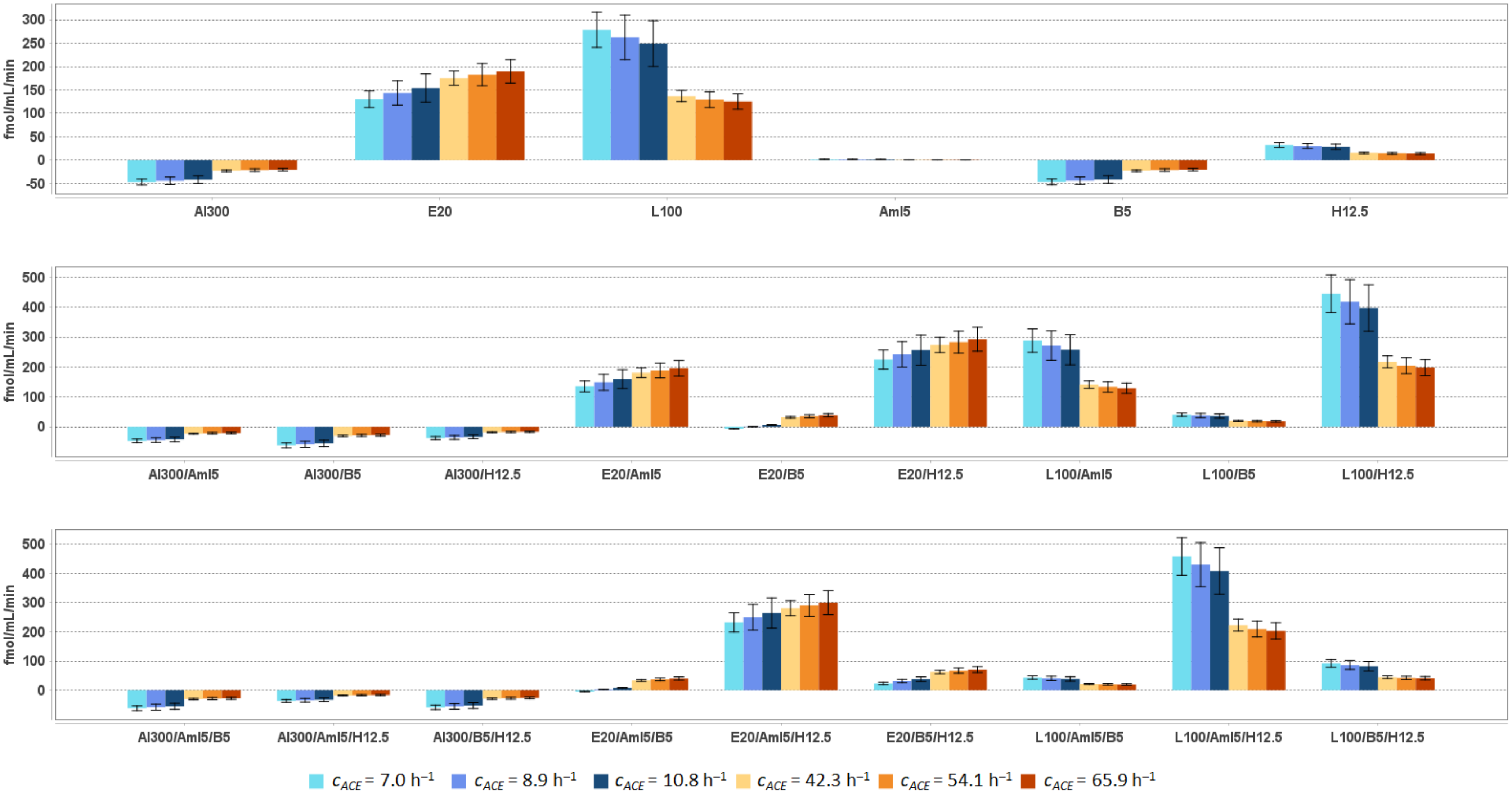


**Al300** = aliskiren 300 mg; **Aml5** = amlodipine 5 mg; **B5** = bisoprolol 5 mg; **E20** = enalapril 20 mg; **H12.5** = hydrochlorothiazide 12.5 mg; **L100** = losartan 100 mg

**Table S50.** Simulated response of plasma renin activity to antihypertensive therapy in virtual hypertensive populations (*n* = 100) with different ACE activity, including *P*-values (Kolmogorov-Smirnov test) for endpoint *vs*. baseline; data are presented as mean ± SD in fmol/mL/min

| Regimens | $c_{ACE}$ = 7.0 h$^{-1}$ | | | $c_{ACE}$ = 8.9 h$^{-1}$ | | | $c_{ACE}$ = 10.8 h$^{-1}$ | | | $c_{ACE}$ = 42.3 h$^{-1}$ | | | $c_{ACE}$ = 54.1 h$^{-1}$ | | | $c_{ACE}$ = 65.9 h$^{-1}$ | | |
|---|---|---|---|---|---|---|---|---|---|---|---|---|---|---|---|---|---|---|
| | **Value** | **Change** | ***P*** | **Value** | **Change** | ***P*** | **Value** | **Change** | ***P*** | **Value** | **Change** | ***P*** | **Value** | **Change** | ***P*** | **Value** | **Change** | ***P*** |
| Baseline | 68.1 ± 9.1 | – | – | 64.1 ± 11.6 | – | – | 60.7 ± 11.8 | – | – | 33.2 ± 2.8 | – | – | 31.3 ± 4.0 | – | – | 30.3 ± 3.9 | – | – |
| Al300 | 21.0 ± 2.8 | -47.1 ± 6.3 | SS | 19.8 ± 3.6 | -44.3 ± 8.0 | SS | 18.8 ± 3.7 | -42.0 ± 8.2 | SS | 10.3 ± 0.9 | -22.9 ± 1.9 | SS | 9.7 ± 1.3 | -21.6 ± 2.8 | SS | 9.4 ± 1.2 | -20.8 ± 2.7 | SS |
| E20 | 198.3 ± 26.7 | 130.2 ± 17.6 | SS | 207.7 ± 37.6 | 143.7 ± 26.0 | SS | 215.0 ± 42.0 | 154.3 ± 30.2 | SS | 208.6 ± 18.2 | 175.4 ± 15.4 | SS | 214.1 ± 27.8 | 182.8 ± 23.8 | SS | 220.1 ± 29.1 | 189.8 ± 25.1 | SS |
| L100 | 346.9 ± 46.9 | 278.8 ± 37.8 | SS | 326.6 ± 59.1 | 262.6 ± 47.6 | SS | 310.1 ± 60.7 | 249.3 ± 48.9 | SS | 170.2 ± 14.8 | 137.0 ± 12.0 | SS | 160.6 ± 20.8 | 129.4 ± 16.8 | SS | 155.5 ± 20.4 | 125.3 ± 16.5 | SS |
| Aml5 | 69.6 ± 9.3 | 1.6 ± 0.3 | 0.69937 | 65.5 ± 11.8 | 1.4 ± 0.3 | 0.90621 | 62.1 ± 12.1 | 1.3 ± 0.3 | 0.58062 | 33.9 ± 2.9 | 0.7 ± 0.1 | 0.21055 | 32.0 ± 4.1 | 0.7 ± 0.1 | 0.11113 | 30.9 ± 4.0 | 0.6 ± 0.1 | 0.58062 |
| B5 | 21.3 ± 2.9 | -46.8 ± 6.3 | SS | 20.1 ± 3.6 | -44.0 ± 7.9 | SS | 19.0 ± 3.7 | -41.7 ± 8.1 | SS | 10.4 ± 0.9 | -22.8 ± 1.9 | SS | 9.9 ± 1.3 | -21.4 ± 2.7 | SS | 9.5 ± 1.3 | -20.7 ± 2.7 | SS |
| H12.5 | 100.3 ± 13.9 | 32.2 ± 5.1 | SS | 94.2 ± 16.5 | 30.1 ± 5.2 | SS | 89.3 ± 17.4 | 28.6 ± 5.7 | SS | 48.7 ± 4.4 | 15.5 ± 1.7 | SS | 45.8 ± 5.8 | 14.5 ± 2.0 | SS | 44.2 ± 5.9 | 14.0 ± 2.1 | SS |
| Al300<br>Aml5 | 21.6 ± 2.9 | -46.5 ± 6.2 | SS | 20.3 ± 3.7 | -43.8 ± 7.9 | SS | 19.3 ± 3.8 | -41.5 ± 8.1 | SS | 10.6 ± 0.9 | -22.6 ± 1.9 | SS | 10.0 ± 1.3 | -21.3 ± 2.7 | SS | 9.6 ± 1.3 | -20.6 ± 2.7 | SS |
| Al300<br>B5 | 6.6 ± 0.9 | -61.5 ± 8.2 | SS | 6.2 ± 1.1 | -57.9 ± 10.4 | SS | 5.9 ± 1.2 | -54.9 ± 10.7 | SS | 3.2 ± 0.3 | -30.0 ± 2.6 | SS | 3.1 ± 0.4 | -28.2 ± 3.6 | SS | 3.0 ± 0.4 | -27.3 ± 3.5 | SS |
| Al300<br>H12.5 | 31.0 ± 4.3 | -37.0 ± 4.9 | SS | 29.2 ± 5.2 | -34.9 ± 6.5 | SS | 27.7 ± 5.4 | -33.0 ± 6.5 | SS | 15.2 ± 1.4 | -18.1 ± 1.5 | SS | 14.3 ± 1.8 | -17.0 ± 2.2 | SS | 13.8 ± 1.9 | -16.5 ± 2.1 | SS |
| E20<br>Aml5 | 203.3 ± 27.4 | 135.3 ± 18.3 | SS | 213.0 ± 38.5 | 149.0 ± 26.9 | SS | 220.5 ± 43.0 | 159.8 ± 31.2 | SS | 214.1 ± 18.7 | 180.9 ± 15.9 | SS | 219.6 ± 28.4 | 188.3 ± 24.4 | SS | 225.7 ± 29.9 | 195.5 ± 25.9 | SS |
| E20<br>B5 | 62.0 ± 8.4 | -6.0 ± 0.9 | 0.00045 | 65.0 ± 11.7 | 0.9 ± 0.5 | 0.99376 | 67.3 ± 13.2 | 6.6 ± 1.4 | 0.01008 | 65.4 ± 5.8 | 32.2 ± 3.0 | SS | 67.2 ± 8.8 | 35.9 ± 4.8 | SS | 69.2 ± 9.3 | 38.9 ± 5.4 | SS |
| E20<br>H12.5 | 292.7 ± 40.9 | 224.7 ± 31.9 | SS | 306.1 ± 54.0 | 242.1 ± 42.6 | SS | 317.1 ± 61.9 | 256.3 ± 50.1 | SS | 306.9 ± 28.1 | 273.7 ± 25.4 | SS | 314.1 ± 40.5 | 282.8 ± 36.6 | SS | 322.8 ± 43.8 | 292.6 ± 39.9 | SS |
| L100<br>Aml5 | 356.2 ± 48.1 | 288.1 ± 39.0 | SS | 335.3 ± 60.6 | 271.2 ± 49.0 | SS | 318.2 ± 62.2 | 257.5 ± 50.3 | SS | 174.5 ± 15.2 | 141.3 ± 12.4 | SS | 164.7 ± 21.3 | 133.4 ± 17.3 | SS | 159.5 ± 21.0 | 129.2 ± 17.1 | SS |
| L100<br>B5 | 108.8 ± 14.6 | 40.8 ± 5.5 | SS | 102.3 ± 18.4 | 38.3 ± 6.9 | SS | 97.2 ± 19.0 | 36.4 ± 7.2 | SS | 53.4 ± 4.7 | 20.2 ± 1.9 | SS | 50.5 ± 6.6 | 19.2 ± 2.6 | SS | 48.9 ± 6.5 | 18.7 ± 2.6 | SS |
| L100<br>H12.5 | 512.4 ± 71.8 | 444.3 ± 62.8 | SS | 481.6 ± 85.2 | 417.6 ± 73.7 | SS | 457.4 ± 89.4 | 396.6 ± 77.6 | SS | 250.3 ± 22.9 | 217.1 ± 20.2 | SS | 235.6 ± 30.3 | 204.3 ± 26.4 | SS | 228.1 ± 30.8 | 197.9 ± 27.0 | SS |
| Al300<br>Aml5/B5 | 6.8 ± 0.9 | -61.3 ± 8.2 | SS | 6.4 ± 1.1 | -57.7 ± 10.4 | SS | 6.0 ± 1.2 | -54.7 ± 10.7 | SS | 3.3 ± 0.3 | -29.9 ± 2.6 | SS | 3.1 ± 0.4 | -28.2 ± 3.6 | SS | 3.0 ± 0.4 | -27.2 ± 3.5 | SS |
| Al300<br>Aml5/H12.5 | 31.8 ± 4.4 | -36.2 ± 4.8 | SS | 29.9 ± 5.3 | -34.1 ± 6.3 | SS | 28.4 ± 5.5 | -32.4 ± 6.4 | SS | 15.5 ± 1.4 | -17.7 ± 1.5 | SS | 14.6 ± 1.9 | -16.7 ± 2.2 | SS | 14.1 ± 1.9 | -16.1 ± 2.1 | SS |
| Al300<br>B5/H12.5 | 9.7 ± 1.4 | -58.4 ± 7.8 | SS | 9.1 ± 1.6 | -54.9 ± 9.9 | SS | 8.7 ± 1.7 | -52.1 ± 10.2 | SS | 4.7 ± 0.4 | -28.5 ± 2.4 | SS | 4.5 ± 0.6 | -26.8 ± 3.4 | SS | 4.3 ± 0.6 | -25.9 ± 3.3 | SS |
| E20<br>Aml5/B5 | 63.7 ± 8.6 | -4.3 ± 0.7 | 0.00232 | 66.8 ± 12.0 | 2.7 ± 0.7 | 0.46756 | 69.1 ± 13.5 | 8.4 ± 1.8 | 0.00136 | 67.1 ± 5.9 | 33.9 ± 3.1 | SS | 68.9 ± 9.0 | 37.6 ± 5.0 | SS | 70.8 ± 9.5 | 40.6 ± 5.6 | SS |
| E20<br>Aml5/H12.5 | 300.2 ± 41.9 | 232.1 ± 32.9 | SS | 313.9 ± 55.3 | 249.8 ± 43.8 | SS | 325.0 ± 63.2 | 264.2 ± 51.5 | SS | 314.0 ± 28.5 | 280.8 ± 25.8 | SS | 321.2 ± 41.3 | 289.9 ± 37.4 | SS | 330.1 ± 44.7 | 299.8 ± 40.9 | SS |
| E20<br>B5/H12.5 | 91.5 ± 12.8 | 23.4 ± 4.1 | SS | 95.7 ± 17.0 | 31.7 ± 5.6 | SS | 99.2 ± 19.4 | 38.4 ± 7.8 | SS | 96.0 ± 8.8 | 62.8 ± 6.1 | SS | 98.4 ± 12.8 | 67.1 ± 8.8 | SS | 101.2 ± 13.9 | 71.0 ± 10.0 | SS |
| L100<br>Aml5/B5 | 111.8 ± 14.9 | 43.7 ± 5.9 | SS | 105.1 ± 18.9 | 41.1 ± 7.3 | SS | 99.8 ± 19.5 | 39.0 ± 7.6 | SS | 54.8 ± 4.8 | 21.5 ± 2.0 | SS | 51.7 ± 6.7 | 20.4 ± 2.7 | SS | 50.1 ± 6.7 | 19.9 ± 2.8 | SS |
| L100<br>Aml5/H12.5 | 525.4 ± 73.3 | 457.4 ± 64.3 | SS | 493.8 ± 87.2 | 429.8 ± 75.7 | SS | 468.8 ± 91.3 | 408.0 ± 79.6 | SS | 256.2 ± 23.3 | 223.0 ± 20.5 | SS | 241.1 ± 31.0 | 209.8 ± 27.0 | SS | 233.4 ± 31.6 | 203.1 ± 27.7 | SS |
| L100<br>B5/H12.5 | 160.0 ± 22.4 | 92.0 ± 13.4 | SS | 150.5 ± 26.7 | 86.4 ± 15.2 | SS | 143.0 ± 28.0 | 82.2 ± 16.3 | SS | 78.4 ± 7.2 | 45.2 ± 4.5 | SS | 73.9 ± 9.6 | 42.6 ± 5.7 | SS | 71.7 ± 9.8 | 41.4 ± 6.0 | SS |

**Al300 =** aliskiren 300 mg; **Aml5** = amlodipine 5 mg; **B5** = bisoprolol 5 mg; **E20** = enalapril 20 mg; **H12.5** = hydrochlorothiazide 12.5 mg; **L100** = losartan 100 mg; **SS** = statistically significant ($P < 0.00001$)

**Table S51.** *P*-values calculated using the Kolmogorov-Smirnov test for changes in plasma renin activity in populations ($n$ = 100) with different ACE activity receiving the same regimens (case 1: $c_{ACE} = 7.0$ h$^{-1}$, case 2: $c_{ACE} = 8.9$ h$^{-1}$, case 3: $c_{ACE} = 10.8$ h$^{-1}$, case 4: $c_{ACE} = 42.3$ h$^{-1}$, case 5: $c_{ACE} = 54.1$ h$^{-1}$, case 6: $c_{ACE} = 65.9$ h$^{-1}$; *P*-value for case *i vs.* case *j* is denoted $P_{ij}$)

| **Regimens** | $P_{12}$ | $P_{13}$ | $P_{23}$ | $P_{14}$ | $P_{15}$ | $P_{16}$ | $P_{24}$ | $P_{25}$ | $P_{26}$ | $P_{34}$ | $P_{35}$ | $P_{36}$ | $P_{45}$ | $P_{46}$ | $P_{56}$ |
|---|---|---|---|---|---|---|---|---|---|---|---|---|---|---|---|
| Al300 | 0.02431 | 0.00004 | 0.36672 | SS | SS | SS | SS | SS | SS | SS | SS | SS | 0.00025 | SS | 0.07832 |
| E20 | 0.00004 | SS | 0.05410 | SS | SS | SS | SS | SS | SS | SS | SS | SS | 0.00002 | SS | 0.02431 |
| L100 | 0.02431 | 0.00007 | 0.36672 | SS | SS | SS | SS | SS | SS | SS | SS | SS | 0.00025 | SS | 0.11113 |
| Aml5 | 0.03663 | SS | 0.02431 | SS | SS | SS | SS | SS | SS | SS | SS | SS | 0.00007 | SS | 0.11113 |
| B5 | 0.02431 | 0.00007 | 0.36672 | SS | SS | SS | SS | SS | SS | SS | SS | SS | 0.00025 | SS | 0.07832 |
| H12.5 | 0.15454 | 0.00025 | 0.11113 | SS | SS | SS | SS | SS | SS | SS | SS | SS | 0.00045 | SS | 0.21055 |
| Al300<br>Aml5 | 0.02431 | 0.00007 | 0.36672 | SS | SS | SS | SS | SS | SS | SS | SS | SS | 0.00025 | SS | 0.07832 |
| Al300<br>B5 | 0.02431 | 0.00007 | 0.36672 | SS | SS | SS | SS | SS | SS | SS | SS | SS | 0.00025 | SS | 0.07832 |
| Al300<br>H12.5 | 0.01008 | 0.00007 | 0.28093 | SS | SS | SS | SS | SS | SS | SS | SS | SS | 0.00025 | SS | 0.03663 |
| E20<br>Aml5 | 0.00004 | SS | 0.05410 | SS | SS | SS | SS | SS | SS | SS | SS | SS | 0.00002 | SS | 0.02431 |
| E20<br>B5 | SS | SS | SS | SS | SS | SS | SS | SS | SS | SS | SS | SS | SS | SS | 0.00045 |
| E20<br>H12.5 | 0.00045 | 0.00002 | 0.07832 | SS | SS | SS | SS | SS | SS | 0.00025 | 0.00013 | 0.00004 | 0.00045 | 0.00004 | 0.11113 |
| L100<br>Aml5 | 0.03663 | 0.00007 | 0.36672 | SS | SS | SS | SS | SS | SS | SS | SS | SS | 0.00025 | SS | 0.15454 |
| L100<br>B5 | 0.01581 | 0.00002 | 0.28093 | SS | SS | SS | SS | SS | SS | SS | SS | SS | 0.00079 | 0.00045 | 0.28093 |
| L100<br>H12.5 | 0.03663 | 0.00004 | 0.28093 | SS | SS | SS | SS | SS | SS | SS | SS | SS | 0.00045 | SS | 0.11113 |
| Al300<br>Aml5/B5 | 0.02431 | 0.00007 | 0.36672 | SS | SS | SS | SS | SS | SS | SS | SS | SS | 0.00025 | SS | 0.07832 |
| Al300<br>Aml5/H12.5 | 0.01008 | 0.00007 | 0.28093 | SS | SS | SS | SS | SS | SS | SS | SS | SS | 0.00025 | 0.00002 | 0.03663 |
| Al300<br>B5/H12.5 | 0.02431 | 0.00004 | 0.36672 | SS | SS | SS | SS | SS | SS | SS | SS | SS | 0.00025 | SS | 0.05410 |
| E20<br>Aml5/B5 | SS | SS | SS | SS | SS | SS | SS | SS | SS | SS | SS | SS | SS | SS | 0.00045 |
| E20<br>Aml5/H12.5 | 0.00045 | 0.00007 | 0.07832 | SS | SS | SS | SS | SS | SS | 0.00025 | 0.00045 | 0.00013 | 0.00045 | 0.00004 | 0.11113 |
| E20<br>B5/H12.5 | SS | SS | SS | SS | SS | SS | SS | SS | SS | SS | SS | SS | 0.00002 | SS | 0.01581 |
| L100<br>Aml5/B5 | 0.01008 | SS | 0.28093 | SS | SS | SS | SS | SS | SS | SS | SS | SS | 0.00136 | 0.00013 | 0.36672 |
| L100<br>Aml5/H12.5 | 0.02431 | SS | 0.21055 | SS | SS | SS | SS | SS | SS | SS | SS | SS | 0.00013 | SS | 0.11113 |
| L100<br>B5/H12.5 | 0.05410 | 0.00004 | 0.21055 | SS | SS | SS | SS | SS | SS | SS | SS | SS | 0.00136 | 0.00004 | 0.36672 |

**Al300 =** aliskiren 300 mg; **Aml5** = amlodipine 5 mg; **B5** = bisoprolol 5 mg; **E20** = enalapril 20 mg; **H12.5** = hydrochlorothiazide 12.5 mg; **L100** = losartan 100 mg; **SS** = statistically significant ($P < 0.00001$)

**Figure S33.** Simulated change in plasma angiotensin I from baseline to week 4 (mean ± SD, $n$ = 100)

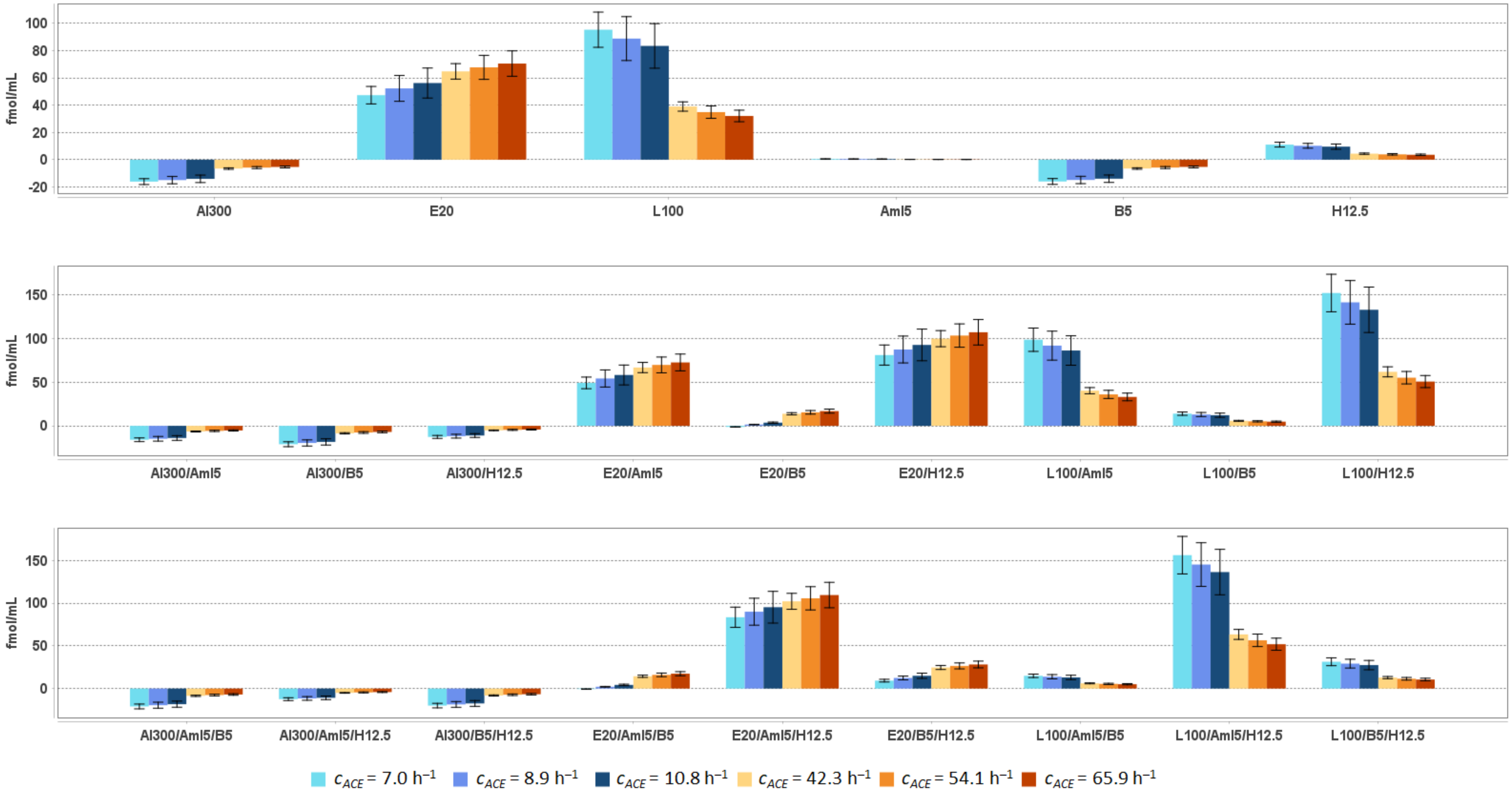


**Al300** = aliskiren 300 mg; **Aml5** = amlodipine 5 mg; **B5** = bisoprolol 5 mg; **E20** = enalapril 20 mg; **H12.5** = hydrochlorothiazide 12.5 mg; **L100** = losartan 100 mg

**Table S52.** Simulated response of <u>plasma angiotensin I</u> to antihypertensive therapy in virtual hypertensive populations ($n$ = 100) with different ACE activity, including $P$-values (Kolmogorov-Smirnov test) for endpoint *vs*. baseline; data are presented as mean ± SD in fmol/mL

| **Regimens** | $c_{ACE}$ = 7.0 h$^{-1}$ | | | $c_{ACE}$ = 8.9 h$^{-1}$ | | | $c_{ACE}$ = 10.8 h$^{-1}$ | | | $c_{ACE}$ = 42.3 h$^{-1}$ | | | $c_{ACE}$ = 54.1 h$^{-1}$ | | | $c_{ACE}$ = 65.9 h$^{-1}$ | | |
|---|---|---|---|---|---|---|---|---|---|---|---|---|---|---|---|---|---|---|
| | **Value** | **Change** | ***P*** | **Value** | **Change** | ***P*** | **Value** | **Change** | ***P*** | **Value** | **Change** | ***P*** | **Value** | **Change** | ***P*** | **Value** | **Change** | ***P*** |
| Baseline | 23.3 ± 3.1 | – | – | 21.7 ± 3.9 | – | – | 20.3 ± 4.0 | – | – | 9.4 ± 0.8 | – | – | 8.4 ± 1.1 | – | – | 7.7 ± 1.0 | – | – |
| Al300 | 7.2 ± 1.0 | -16.1 ± 2.1 | SS | 6.7 ± 1.2 | -15.0 ± 2.7 | SS | 6.3 ± 1.2 | -14.0 ± 2.7 | SS | 2.9 ± 0.3 | -6.5 ± 0.6 | SS | 2.6 ± 0.3 | -5.8 ± 0.7 | SS | 2.4 ± 0.3 | -5.3 ± 0.7 | SS |
| E20 | 70.6 ± 9.5 | 47.3 ± 6.4 | SS | 73.9 ± 13.4 | 52.3 ± 9.5 | SS | 76.5 ± 15.0 | 56.2 ± 11.0 | SS | 74.2 ± 6.5 | 64.7 ± 5.7 | SS | 76.1 ± 9.9 | 67.7 ± 8.8 | SS | 78.2 ± 10.3 | 70.5 ± 9.3 | SS |
| L100 | 118.6 ± 16.0 | 95.3 ± 12.9 | SS | 110.4 ± 20.0 | 88.8 ± 16.1 | SS | 103.7 ± 20.3 | 83.4 ± 16.3 | SS | 48.4 ± 4.2 | 39.0 ± 3.4 | SS | 43.3 ± 5.6 | 34.9 ± 4.5 | SS | 39.8 ± 5.2 | 32.1 ± 4.2 | SS |
| Aml5 | 23.8 ± 3.2 | 0.5 ± 0.1 | 0.69937 | 22.1 ± 4.0 | 0.5 ± 0.1 | 0.90621 | 20.8 ± 4.0 | 0.4 ± 0.1 | 0.58062 | 9.7 ± 0.8 | 0.2 ± 0.0 | 0.21055 | 8.6 ± 1.1 | 0.2 ± 0.0 | 0.11113 | 7.9 ± 1.0 | 0.2 ± 0.0 | 0.58062 |
| B5 | 7.3 ± 1.0 | -16.0 ± 2.1 | SS | 6.8 ± 1.2 | -14.9 ± 2.7 | SS | 6.4 ± 1.2 | -14.0 ± 2.7 | SS | 3.0 ± 0.3 | -6.5 ± 0.6 | SS | 2.7 ± 0.3 | -5.8 ± 0.7 | SS | 2.4 ± 0.3 | -5.3 ± 0.7 | SS |
| H12.5 | 34.3 ± 4.8 | 11.0 ± 1.7 | SS | 31.8 ± 5.6 | 10.2 ± 1.8 | SS | 29.9 ± 5.8 | 9.6 ± 1.9 | SS | 13.9 ± 1.2 | 4.4 ± 0.5 | SS | 12.3 ± 1.6 | 3.9 ± 0.5 | SS | 11.3 ± 1.5 | 3.6 ± 0.5 | SS |
| Al300 Aml5 | 7.4 ± 1.0 | -15.9 ± 2.1 | SS | 6.9 ± 1.2 | -14.8 ± 2.7 | SS | 6.4 ± 1.3 | -13.9 ± 2.7 | SS | 3.0 ± 0.3 | -6.4 ± 0.5 | SS | 2.7 ± 0.3 | -5.7 ± 0.7 | SS | 2.5 ± 0.3 | -5.3 ± 0.7 | SS |
| Al300 B5 | 2.3 ± 0.3 | -21.0 ± 2.8 | SS | 2.1 ± 0.4 | -19.6 ± 3.5 | SS | 2.0 ± 0.4 | -18.4 ± 3.6 | SS | 0.9 ± 0.1 | -8.5 ± 0.7 | SS | 0.8 ± 0.1 | -7.6 ± 1.0 | SS | 0.8 ± 0.1 | -7.0 ± 0.9 | SS |
| Al300 H12.5 | 10.6 ± 1.5 | -12.7 ± 1.7 | SS | 9.9 ± 1.7 | -11.8 ± 2.2 | SS | 9.3 ± 1.8 | -11.1 ± 2.2 | SS | 4.3 ± 0.4 | -5.1 ± 0.4 | SS | 3.8 ± 0.5 | -4.6 ± 0.6 | SS | 3.5 ± 0.5 | -4.2 ± 0.5 | SS |
| E20 Aml5 | 72.4 ± 9.8 | 49.1 ± 6.6 | SS | 75.8 ± 13.7 | 54.2 ± 9.8 | SS | 78.5 ± 15.3 | 58.1 ± 11.4 | SS | 76.1 ± 6.7 | 66.7 ± 5.8 | SS | 78.1 ± 10.1 | 69.6 ± 9.0 | SS | 80.2 ± 10.6 | 72.5 ± 9.6 | SS |
| E20 B5 | 22.1 ± 3.0 | -1.2 ± 0.2 | 0.05410 | 23.1 ± 4.2 | 1.5 ± 0.3 | 0.15454 | 24.0 ± 4.7 | 3.6 ± 0.7 | 0.00007 | 23.3 ± 2.1 | 13.8 ± 1.3 | SS | 23.9 ± 3.1 | 15.5 ± 2.0 | SS | 24.6 ± 3.3 | 16.8 ± 2.3 | SS |
| E20 H12.5 | 104.2 ± 14.6 | 80.9 ± 11.5 | SS | 108.9 ± 19.2 | 87.3 ± 15.4 | SS | 112.8 ± 22.0 | 92.5 ± 18.1 | SS | 109.1 ± 10.0 | 99.7 ± 9.2 | SS | 111.7 ± 14.4 | 103.2 ± 13.3 | SS | 114.7 ± 15.6 | 107.0 ± 14.6 | SS |
| L100 Aml5 | 121.7 ± 16.5 | 98.5 ± 13.3 | SS | 113.4 ± 20.5 | 91.7 ± 16.6 | SS | 106.5 ± 20.8 | 86.1 ± 16.8 | SS | 49.7 ± 4.3 | 40.2 ± 3.5 | SS | 44.4 ± 5.7 | 36.0 ± 4.7 | SS | 40.8 ± 5.4 | 33.1 ± 4.4 | SS |
| L100 B5 | 37.2 ± 5.0 | 13.9 ± 1.9 | SS | 34.6 ± 6.2 | 12.9 ± 2.3 | SS | 32.5 ± 6.3 | 12.2 ± 2.4 | SS | 15.2 ± 1.3 | 5.7 ± 0.5 | SS | 13.6 ± 1.8 | 5.2 ± 0.7 | SS | 12.5 ± 1.7 | 4.8 ± 0.7 | SS |
| L100 H12.5 | 175.1 ± 24.5 | 151.9 ± 21.5 | SS | 162.8 ± 28.8 | 141.2 ± 24.9 | SS | 153.0 ± 29.9 | 132.7 ± 26.0 | SS | 71.2 ± 6.5 | 61.8 ± 5.7 | SS | 63.5 ± 8.2 | 55.1 ± 7.1 | SS | 58.4 ± 7.9 | 50.6 ± 6.9 | SS |
| Al300 Aml5/B5 | 2.3 ± 0.3 | -21.0 ± 2.8 | SS | 2.2 ± 0.4 | -19.5 ± 3.5 | SS | 2.0 ± 0.4 | -18.3 ± 3.6 | SS | 0.9 ± 0.1 | -8.5 ± 0.7 | SS | 0.8 ± 0.1 | -7.6 ± 1.0 | SS | 0.8 ± 0.1 | -7.0 ± 0.9 | SS |
| Al300 Aml5/H12.5 | 10.9 ± 1.5 | -12.4 ± 1.6 | SS | 10.1 ± 1.8 | -11.5 ± 2.1 | SS | 9.5 ± 1.8 | -10.8 ± 2.1 | SS | 4.4 ± 0.4 | -5.0 ± 0.4 | SS | 3.9 ± 0.5 | -4.5 ± 0.6 | SS | 3.6 ± 0.5 | -4.1 ± 0.5 | SS |
| Al300 B5/H12.5 | 3.3 ± 0.5 | -20.0 ± 2.7 | SS | 3.1 ± 0.5 | -18.6 ± 3.4 | SS | 2.9 ± 0.6 | -17.4 ± 3.4 | SS | 1.4 ± 0.1 | -8.1 ± 0.7 | SS | 1.2 ± 0.2 | -7.2 ± 0.9 | SS | 1.1 ± 0.2 | -6.6 ± 0.9 | SS |
| E20 Aml5/B5 | 22.7 ± 3.1 | -0.6 ± 0.2 | 0.46756 | 23.8 ± 4.3 | 2.1 ± 0.4 | 0.01581 | 24.6 ± 4.8 | 4.3 ± 0.9 | SS | 23.9 ± 2.1 | 14.4 ± 1.3 | SS | 24.5 ± 3.2 | 16.1 ± 2.1 | SS | 25.2 ± 3.4 | 17.4 ± 2.4 | SS |
| E20 Aml5/H12.5 | 106.8 ± 14.9 | 83.6 ± 11.8 | SS | 111.7 ± 19.7 | 90.1 ± 15.8 | SS | 115.6 ± 22.5 | 95.3 ± 18.6 | SS | 111.7 ± 10.1 | 102.2 ± 9.4 | SS | 114.2 ± 14.7 | 105.8 ± 13.6 | SS | 117.3 ± 15.9 | 109.6 ± 14.9 | SS |
| E20 B5/H12.5 | 32.6 ± 4.6 | 9.3 ± 1.6 | SS | 34.1 ± 6.0 | 12.4 ± 2.2 | SS | 35.3 ± 6.9 | 15.0 ± 3.0 | SS | 34.1 ± 3.1 | 24.7 ± 2.4 | SS | 35.0 ± 4.5 | 26.6 ± 3.5 | SS | 36.0 ± 4.9 | 28.2 ± 3.9 | SS |
| L100 Aml5/B5 | 38.2 ± 5.1 | 14.9 ± 2.0 | SS | 35.5 ± 6.4 | 13.9 ± 2.5 | SS | 33.4 ± 6.5 | 13.1 ± 2.6 | SS | 15.6 ± 1.4 | 6.1 ± 0.6 | SS | 13.9 ± 1.8 | 5.5 ± 0.7 | SS | 12.8 ± 1.7 | 5.1 ± 0.7 | SS |
| L100 Aml5/H12.5 | 179.6 ± 25.1 | 156.3 ± 22.0 | SS | 167.0 ± 29.5 | 145.3 ± 25.6 | SS | 156.8 ± 30.5 | 136.5 ± 26.6 | SS | 72.9 ± 6.6 | 63.4 ± 5.8 | SS | 65.0 ± 8.3 | 56.5 ± 7.3 | SS | 59.7 ± 8.1 | 52.0 ± 7.1 | SS |
| L100 B5/H12.5 | 54.7 ± 7.6 | 31.4 ± 4.6 | SS | 50.9 ± 9.0 | 29.2 ± 5.2 | SS | 47.8 ± 9.4 | 27.5 ± 5.5 | SS | 22.3 ± 2.0 | 12.9 ± 1.3 | SS | 19.9 ± 2.6 | 11.5 ± 1.5 | SS | 18.3 ± 2.5 | 10.6 ± 1.5 | SS |

**Al300 =** aliskiren 300 mg; **Aml5** = amlodipine 5 mg; **B5** = bisoprolol 5 mg; **E20** = enalapril 20 mg; **H12.5** = hydrochlorothiazide 12.5 mg; **L100** = losartan 100 mg; **SS** = statistically significant ($p$ < 0.00001)

**Table S53.** *P*-values calculated using the Kolmogorov-Smirnov test for changes in plasma angiotensin I in populations ($n$ = 100) with different ACE activity receiving the same regimens (case 1: $c_{ACE} = 7.0\ h^{-1}$, case 2: $c_{ACE} = 8.9\ h^{-1}$, case 3: $c_{ACE} = 10.8\ h^{-1}$, case 4: $c_{ACE} = 42.3\ h^{-1}$, case 5: $c_{ACE} = 54.1\ h^{-1}$, case 6: $c_{ACE} = 65.9\ h^{-1}$; *P*-value for case *i vs.* case *j* is denoted $P_{ij}$)

| **Regimens** | $P_{12}$ | $P_{13}$ | $P_{23}$ | $P_{14}$ | $P_{15}$ | $P_{16}$ | $P_{24}$ | $P_{25}$ | $P_{26}$ | $P_{34}$ | $P_{35}$ | $P_{36}$ | $P_{45}$ | $P_{46}$ | $P_{56}$ |
|---|---|---|---|---|---|---|---|---|---|---|---|---|---|---|---|
| Al300 | 0.01008 | SS | 0.21055 | SS | SS | SS | SS | SS | SS | SS | SS | SS | SS | SS | 0.00004 |
| E20 | 0.00002 | SS | 0.05410 | SS | SS | SS | SS | SS | SS | SS | SS | SS | SS | SS | 0.02431 |
| L100 | 0.01581 | SS | 0.28093 | SS | SS | SS | SS | SS | SS | SS | SS | SS | SS | SS | 0.00007 |
| Aml5 | 0.01581 | SS | 0.01008 | SS | SS | SS | SS | SS | SS | SS | SS | SS | SS | SS | 0.00136 |
| B5 | 0.01008 | SS | 0.21055 | SS | SS | SS | SS | SS | SS | SS | SS | SS | SS | SS | 0.00004 |
| H12.5 | 0.05410 | 0.00004 | 0.05410 | SS | SS | SS | SS | SS | SS | SS | SS | SS | SS | SS | 0.00232 |
| Al300<br>Aml5 | 0.01008 | SS | 0.21055 | SS | SS | SS | SS | SS | SS | SS | SS | SS | SS | SS | 0.00004 |
| Al300<br>B5 | 0.01008 | SS | 0.28093 | SS | SS | SS | SS | SS | SS | SS | SS | SS | SS | SS | 0.00004 |
| Al300<br>H12.5 | 0.00232 | SS | 0.28093 | SS | SS | SS | SS | SS | SS | SS | SS | SS | SS | SS | 0.00007 |
| E20<br>Aml5 | 0.00004 | SS | 0.05410 | SS | SS | SS | SS | SS | SS | SS | SS | SS | 0.00002 | SS | 0.02431 |
| E20<br>B5 | SS | SS | SS | SS | SS | SS | SS | SS | SS | SS | SS | SS | SS | SS | 0.00025 |
| E20<br>H12.5 | 0.00045 | SS | 0.07832 | SS | SS | SS | SS | SS | SS | 0.00007 | 0.00007 | SS | 0.00013 | 0.00002 | 0.07832 |
| L100<br>Aml5 | 0.01581 | SS | 0.21055 | SS | SS | SS | SS | SS | SS | SS | SS | SS | SS | SS | 0.00007 |
| L100<br>B5 | 0.00630 | SS | 0.21055 | SS | SS | SS | SS | SS | SS | SS | SS | SS | SS | SS | 0.00136 |
| L100<br>H12.5 | 0.01008 | SS | 0.15454 | SS | SS | SS | SS | SS | SS | SS | SS | SS | SS | SS | 0.00136 |
| Al300<br>Aml5/B5 | 0.01008 | SS | 0.28093 | SS | SS | SS | SS | SS | SS | SS | SS | SS | SS | SS | 0.00004 |
| Al300<br>Aml5/H12.5 | 0.00232 | SS | 0.28093 | SS | SS | SS | SS | SS | SS | SS | SS | SS | SS | SS | 0.00007 |
| Al300<br>B5/H12.5 | 0.01008 | SS | 0.21055 | SS | SS | SS | SS | SS | SS | SS | SS | SS | SS | SS | 0.00004 |
| E20<br>Aml5/B5 | SS | SS | SS | SS | SS | SS | SS | SS | SS | SS | SS | SS | SS | SS | 0.00025 |
| E20<br>Aml5/H12.5 | 0.00045 | SS | 0.07832 | SS | SS | SS | SS | SS | SS | 0.00013 | 0.00013 | SS | 0.00025 | 0.00004 | 0.07832 |
| E20<br>B5/H12.5 | SS | SS | SS | SS | SS | SS | SS | SS | SS | SS | SS | SS | SS | SS | 0.01008 |
| L100<br>Aml5/B5 | 0.00630 | SS | 0.11113 | SS | SS | SS | SS | SS | SS | SS | SS | SS | SS | SS | 0.00079 |
| L100<br>Aml5/H12.5 | 0.01008 | SS | 0.11113 | SS | SS | SS | SS | SS | SS | SS | SS | SS | SS | SS | 0.00136 |
| L100<br>B5/H12.5 | 0.01581 | SS | 0.07832 | SS | SS | SS | SS | SS | SS | SS | SS | SS | SS | SS | 0.01008 |

**Al300 =** aliskiren 300 mg; **Aml5** = amlodipine 5 mg; **B5** = bisoprolol 5 mg; **E20** = enalapril 20 mg; **H12.5** = hydrochlorothiazide 12.5 mg; **L100** = losartan 100 mg; **SS** = statistically significant ($P < 0.00001$)

**Figure S34.** Simulated change in plasma angiotensin II from baseline to week 4 (mean ± SD, $n$ = 100)

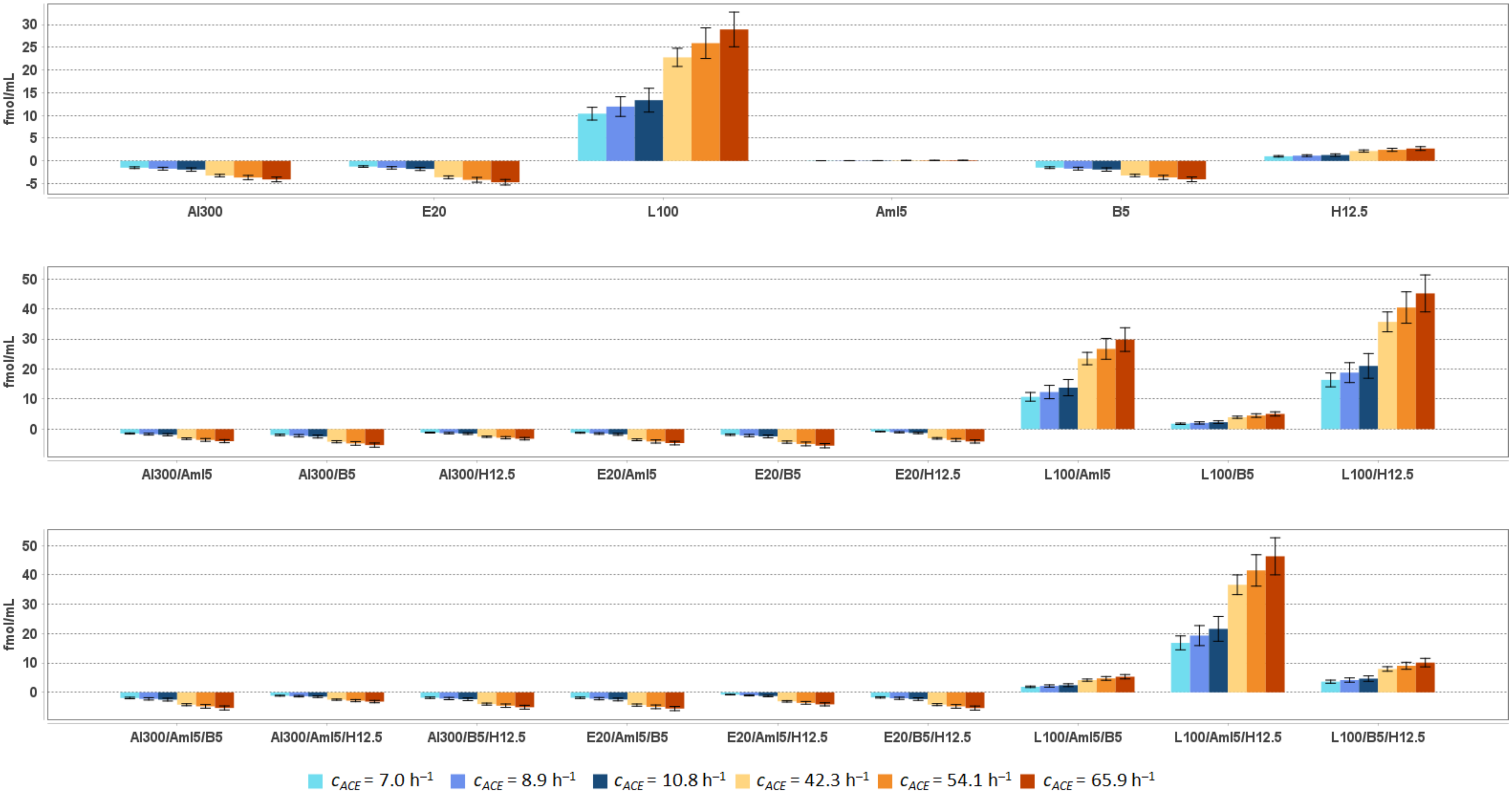


**Al300** = aliskiren 300 mg; **Aml5** = amlodipine 5 mg; **B5** = bisoprolol 5 mg; **E20** = enalapril 20 mg; **H12.5** = hydrochlorothiazide 12.5 mg; **L100** = losartan 100 mg

**Table S54.** Simulated response of <u>plasma angiotensin II</u> to antihypertensive therapy in virtual hypertensive populations ($n$ = 100) with different ACE activity, including $P$-values (Kolmogorov-Smirnov test) for endpoint *vs*. baseline; data are presented as mean ± SD in fmol/mL

| **Regimens** | $c_{ACE}$ = 7.0 h$^{-1}$ | | | $c_{ACE}$ = 8.9 h$^{-1}$ | | | $c_{ACE}$ = 10.8 h$^{-1}$ | | | $c_{ACE}$ = 42.3 h$^{-1}$ | | | $c_{ACE}$ = 54.1 h$^{-1}$ | | | $c_{ACE}$ = 65.9 h$^{-1}$ | | |
|---|---|---|---|---|---|---|---|---|---|---|---|---|---|---|---|---|---|---|
| | **Value** | **Change** | ***P*** | **Value** | **Change** | ***P*** | **Value** | **Change** | ***P*** | **Value** | **Change** | ***P*** | **Value** | **Change** | ***P*** | **Value** | **Change** | ***P*** |
| Baseline | 2.1 ± 0.3 | – | – | 2.5 ± 0.4 | – | – | 2.8 ± 0.5 | – | – | 4.7 ± 0.4 | – | – | 5.3 ± 0.7 | – | – | 5.9 ± 0.8 | – | – |
| Al300 | 0.7 ± 0.1 | -1.5 ± 0.2 | SS | 0.8 ± 0.1 | -1.7 ± 0.3 | SS | 0.9 ± 0.2 | -1.9 ± 0.4 | SS | 1.4 ± 0.1 | -3.2 ± 0.3 | SS | 1.6 ± 0.2 | -3.7 ± 0.5 | SS | 1.8 ± 0.2 | -4.1 ± 0.5 | SS |
| E20 | 0.9 ± 0.1 | -1.2 ± 0.2 | SS | 1.0 ± 0.2 | -1.5 ± 0.3 | SS | 1.0 ± 0.2 | -1.8 ± 0.3 | SS | 1.1 ± 0.1 | -3.6 ± 0.3 | SS | 1.1 ± 0.1 | -4.2 ± 0.5 | SS | 1.2 ± 0.2 | -4.7 ± 0.6 | SS |
| L100 | 12.5 ± 1.7 | 10.4 ± 1.4 | SS | 14.4 ± 2.6 | 12.0 ± 2.2 | SS | 16.1 ± 3.2 | 13.4 ± 2.6 | SS | 27.5 ± 2.4 | 22.8 ± 2.0 | SS | 31.2 ± 4.0 | 25.9 ± 3.4 | SS | 34.8 ± 4.6 | 28.9 ± 3.8 | SS |
| Aml5 | 2.2 ± 0.3 | 0.0 ± 0.0 | 0.69937 | 2.5 ± 0.5 | 0.1 ± 0.0 | 0.90621 | 2.8 ± 0.5 | 0.1 ± 0.0 | 0.58062 | 4.8 ± 0.4 | 0.1 ± 0.0 | 0.21055 | 5.4 ± 0.7 | 0.1 ± 0.0 | 0.11113 | 6.0 ± 0.8 | 0.1 ± 0.0 | 0.58062 |
| B5 | 0.7 ± 0.1 | -1.5 ± 0.2 | SS | 0.8 ± 0.1 | -1.7 ± 0.3 | SS | 0.9 ± 0.2 | -1.9 ± 0.4 | SS | 1.5 ± 0.1 | -3.2 ± 0.3 | SS | 1.7 ± 0.2 | -3.6 ± 0.5 | SS | 1.9 ± 0.2 | -4.0 ± 0.5 | SS |
| H12.5 | 3.2 ± 0.4 | 1.0 ± 0.2 | SS | 3.6 ± 0.6 | 1.2 ± 0.2 | SS | 4.1 ± 0.8 | 1.3 ± 0.3 | SS | 6.8 ± 0.6 | 2.2 ± 0.2 | SS | 7.8 ± 1.0 | 2.5 ± 0.3 | SS | 8.6 ± 1.2 | 2.7 ± 0.4 | SS |
| Al300<br>Aml5 | 0.7 ± 0.1 | -1.5 ± 0.2 | SS | 0.8 ± 0.1 | -1.7 ± 0.3 | SS | 0.9 ± 0.2 | -1.9 ± 0.4 | SS | 1.5 ± 0.1 | -3.2 ± 0.3 | SS | 1.7 ± 0.2 | -3.6 ± 0.5 | SS | 1.9 ± 0.2 | -4.0 ± 0.5 | SS |
| Al300<br>B5 | 0.2 ± 0.0 | -1.9 ± 0.3 | SS | 0.2 ± 0.0 | -2.2 ± 0.4 | SS | 0.3 ± 0.1 | -2.5 ± 0.5 | SS | 0.5 ± 0.0 | -4.2 ± 0.4 | SS | 0.5 ± 0.1 | -4.8 ± 0.6 | SS | 0.6 ± 0.1 | -5.3 ± 0.7 | SS |
| Al300<br>H12.5 | 1.0 ± 0.1 | -1.2 ± 0.2 | SS | 1.1 ± 0.2 | -1.3 ± 0.2 | SS | 1.3 ± 0.2 | -1.5 ± 0.3 | SS | 2.1 ± 0.2 | -2.5 ± 0.2 | SS | 2.4 ± 0.3 | -2.9 ± 0.4 | SS | 2.7 ± 0.4 | -3.2 ± 0.4 | SS |
| E20<br>Aml5 | 0.9 ± 0.1 | -1.2 ± 0.2 | SS | 1.0 ± 0.2 | -1.5 ± 0.3 | SS | 1.0 ± 0.2 | -1.7 ± 0.3 | SS | 1.1 ± 0.1 | -3.6 ± 0.3 | SS | 1.2 ± 0.2 | -4.1 ± 0.5 | SS | 1.2 ± 0.2 | -4.7 ± 0.6 | SS |
| E20<br>B5 | 0.3 ± 0.0 | -1.9 ± 0.2 | SS | 0.3 ± 0.1 | -2.2 ± 0.4 | SS | 0.3 ± 0.1 | -2.4 ± 0.5 | SS | 0.3 ± 0.0 | -4.3 ± 0.4 | SS | 0.4 ± 0.0 | -4.9 ± 0.6 | SS | 0.4 ± 0.1 | -5.5 ± 0.7 | SS |
| E20<br>H12.5 | 1.3 ± 0.2 | -0.8 ± 0.1 | SS | 1.4 ± 0.2 | -1.1 ± 0.2 | SS | 1.5 ± 0.3 | -1.3 ± 0.3 | SS | 1.6 ± 0.1 | -3.1 ± 0.3 | SS | 1.7 ± 0.2 | -3.6 ± 0.5 | SS | 1.8 ± 0.2 | -4.1 ± 0.5 | SS |
| L100<br>Aml5 | 12.9 ± 1.7 | 10.7 ± 1.5 | SS | 14.8 ± 2.7 | 12.3 ± 2.2 | SS | 16.5 ± 3.2 | 13.8 ± 2.7 | SS | 28.2 ± 2.5 | 23.5 ± 2.1 | SS | 32.0 ± 4.1 | 26.7 ± 3.5 | SS | 35.7 ± 4.7 | 29.8 ± 3.9 | SS |
| L100<br>B5 | 3.9 ± 0.5 | 1.8 ± 0.2 | SS | 4.5 ± 0.8 | 2.1 ± 0.4 | SS | 5.1 ± 1.0 | 2.3 ± 0.5 | SS | 8.6 ± 0.8 | 3.9 ± 0.4 | SS | 9.8 ± 1.3 | 4.5 ± 0.6 | SS | 11.0 ± 1.5 | 5.0 ± 0.7 | SS |
| L100<br>H12.5 | 18.5 ± 2.6 | 16.4 ± 2.3 | SS | 21.3 ± 3.8 | 18.8 ± 3.3 | SS | 23.8 ± 4.6 | 21.0 ± 4.1 | SS | 40.4 ± 3.7 | 35.7 ± 3.3 | SS | 45.8 ± 5.9 | 40.5 ± 5.2 | SS | 51.1 ± 6.9 | 45.2 ± 6.2 | SS |
| Al300<br>Aml5/B5 | 0.2 ± 0.0 | -1.9 ± 0.3 | SS | 0.2 ± 0.0 | -2.2 ± 0.4 | SS | 0.3 ± 0.1 | -2.5 ± 0.5 | SS | 0.5 ± 0.0 | -4.2 ± 0.4 | SS | 0.5 ± 0.1 | -4.8 ± 0.6 | SS | 0.6 ± 0.1 | -5.3 ± 0.7 | SS |
| Al300<br>Aml5/H12.5 | 1.0 ± 0.1 | -1.1 ± 0.2 | SS | 1.2 ± 0.2 | -1.3 ± 0.2 | SS | 1.3 ± 0.3 | -1.5 ± 0.3 | SS | 2.2 ± 0.2 | -2.5 ± 0.2 | SS | 2.5 ± 0.3 | -2.8 ± 0.4 | SS | 2.8 ± 0.4 | -3.1 ± 0.4 | SS |
| Al300<br>B5/H12.5 | 0.3 ± 0.0 | -1.8 ± 0.2 | SS | 0.4 ± 0.1 | -2.1 ± 0.4 | SS | 0.4 ± 0.1 | -2.4 ± 0.5 | SS | 0.7 ± 0.1 | -4.0 ± 0.3 | SS | 0.8 ± 0.1 | -4.5 ± 0.6 | SS | 0.8 ± 0.1 | -5.1 ± 0.7 | SS |
| E20<br>Aml5/B5 | 0.3 ± 0.0 | -1.9 ± 0.2 | SS | 0.3 ± 0.1 | -2.2 ± 0.4 | SS | 0.3 ± 0.1 | -2.4 ± 0.5 | SS | 0.3 ± 0.0 | -4.3 ± 0.4 | SS | 0.4 ± 0.0 | -4.9 ± 0.6 | SS | 0.4 ± 0.1 | -5.5 ± 0.7 | SS |
| E20<br>Aml5/H12.5 | 1.4 ± 0.2 | -0.8 ± 0.1 | SS | 1.4 ± 0.3 | -1.0 ± 0.2 | SS | 1.5 ± 0.3 | -1.2 ± 0.2 | SS | 1.6 ± 0.1 | -3.1 ± 0.3 | SS | 1.7 ± 0.2 | -3.6 ± 0.5 | SS | 1.8 ± 0.2 | -4.1 ± 0.5 | SS |
| E20<br>B5/H12.5 | 0.4 ± 0.1 | -1.7 ± 0.2 | SS | 0.4 ± 0.1 | -2.0 ± 0.4 | SS | 0.5 ± 0.1 | -2.3 ± 0.4 | SS | 0.5 ± 0.0 | -4.2 ± 0.4 | SS | 0.5 ± 0.1 | -4.8 ± 0.6 | SS | 0.6 ± 0.1 | -5.3 ± 0.7 | SS |
| L100<br>Aml5/B5 | 4.0 ± 0.5 | 1.9 ± 0.3 | SS | 4.6 ± 0.8 | 2.2 ± 0.4 | SS | 5.2 ± 1.0 | 2.4 ± 0.5 | SS | 8.8 ± 0.8 | 4.2 ± 0.4 | SS | 10.1 ± 1.3 | 4.8 ± 0.6 | SS | 11.2 ± 1.5 | 5.3 ± 0.7 | SS |
| L100<br>Aml5/H12.5 | 19.0 ± 2.7 | 16.9 ± 2.4 | SS | 21.8 ± 3.8 | 19.3 ± 3.4 | SS | 24.4 ± 4.7 | 21.6 ± 4.2 | SS | 41.3 ± 3.8 | 36.7 ± 3.4 | SS | 46.8 ± 6.0 | 41.5 ± 5.3 | SS | 52.3 ± 7.1 | 46.4 ± 6.3 | SS |
| L100<br>B5/H12.5 | 5.8 ± 0.8 | 3.6 ± 0.5 | SS | 6.6 ± 1.2 | 4.2 ± 0.7 | SS | 7.4 ± 1.5 | 4.7 ± 0.9 | SS | 12.6 ± 1.2 | 8.0 ± 0.8 | SS | 14.4 ± 1.9 | 9.1 ± 1.2 | SS | 16.0 ± 2.2 | 10.1 ± 1.4 | SS |

**Al300 =** aliskiren 300 mg; **Aml5** = amlodipine 5 mg; **B5** = bisoprolol 5 mg; **E20** = enalapril 20 mg; **H12.5** = hydrochlorothiazide 12.5 mg; **L100** = losartan 100 mg; **SS** = statistically significant ($P$ < 0.00001)

**Table S55.** *P*-values calculated using the Kolmogorov-Smirnov test for changes in plasma angiotensin II in populations ($n$ = 100) with different ACE activity receiving the same regimens (case 1: $c_{ACE}$ = 7.0 $h^{-1}$, case 2: $c_{ACE}$ = 8.9 $h^{-1}$, case 3: $c_{ACE}$ = 10.8 $h^{-1}$, case 4: $c_{ACE}$ = 42.3 $h^{-1}$, case 5: $c_{ACE}$ = 54.1 $h^{-1}$, case 6: $c_{ACE}$ = 65.9 $h^{-1}$; *P*-value for case *i* *vs*. case *j* is denoted $P_{ij}$)

| Regimens | $P_{12}$ | $P_{13}$ | $P_{23}$ | $P_{14}$ | $P_{15}$ | $P_{16}$ | $P_{24}$ | $P_{25}$ | $P_{26}$ | $P_{34}$ | $P_{35}$ | $P_{36}$ | $P_{45}$ | $P_{46}$ | $P_{56}$ |
|---|---|---|---|---|---|---|---|---|---|---|---|---|---|---|---|
| Al300 | SS | SS | 0.00136 | SS | SS | SS | SS | SS | SS | SS | SS | SS | SS | SS | 0.00002 |
| E20 | SS | SS | 0.00013 | SS | SS | SS | SS | SS | SS | SS | SS | SS | SS | SS | SS |
| L100 | SS | SS | 0.00386 | SS | SS | SS | SS | SS | SS | SS | SS | SS | SS | SS | 0.00004 |
| Aml5 | 0.00025 | SS | 0.00136 | SS | SS | SS | SS | SS | SS | SS | SS | SS | 0.00025 | SS | 0.00232 |
| B5 | SS | SS | 0.00136 | SS | SS | SS | SS | SS | SS | SS | SS | SS | SS | SS | 0.00002 |
| H12.5 | SS | SS | 0.00630 | SS | SS | SS | SS | SS | SS | SS | SS | SS | SS | SS | 0.00079 |
| Al300<br>Aml5 | SS | SS | 0.00136 | SS | SS | SS | SS | SS | SS | SS | SS | SS | SS | SS | 0.00002 |
| Al300<br>B5 | SS | SS | 0.00136 | SS | SS | SS | SS | SS | SS | SS | SS | SS | SS | SS | 0.00002 |
| Al300<br>H12.5 | SS | SS | 0.00386 | SS | SS | SS | SS | SS | SS | SS | SS | SS | SS | SS | 0.00007 |
| E20<br>Aml5 | SS | SS | 0.00013 | SS | SS | SS | SS | SS | SS | SS | SS | SS | SS | SS | SS |
| E20<br>B5 | SS | SS | 0.00079 | SS | SS | SS | SS | SS | SS | SS | SS | SS | SS | SS | 0.00002 |
| E20<br>H12.5 | SS | SS | SS | SS | SS | SS | SS | SS | SS | SS | SS | SS | SS | SS | SS |
| L100<br>Aml5 | SS | SS | 0.00386 | SS | SS | SS | SS | SS | SS | SS | SS | SS | SS | SS | 0.00004 |
| L100<br>B5 | SS | SS | 0.00386 | SS | SS | SS | SS | SS | SS | SS | SS | SS | SS | SS | SS |
| L100<br>H12.5 | SS | SS | 0.00386 | SS | SS | SS | SS | SS | SS | SS | SS | SS | SS | SS | 0.00002 |
| Al300<br>Aml5/B5 | SS | SS | 0.00136 | SS | SS | SS | SS | SS | SS | SS | SS | SS | SS | SS | 0.00002 |
| Al300<br>Aml5/H12.5 | SS | SS | 0.00630 | SS | SS | SS | SS | SS | SS | SS | SS | SS | SS | SS | 0.00007 |
| Al300<br>B5/H12.5 | SS | SS | 0.00232 | SS | SS | SS | SS | SS | SS | SS | SS | SS | SS | SS | 0.00002 |
| E20<br>Aml5/B5 | SS | SS | 0.00079 | SS | SS | SS | SS | SS | SS | SS | SS | SS | SS | SS | 0.00002 |
| E20<br>Aml5/H12.5 | SS | SS | SS | SS | SS | SS | SS | SS | SS | SS | SS | SS | SS | SS | SS |
| E20<br>B5/H12.5 | SS | SS | 0.00079 | SS | SS | SS | SS | SS | SS | SS | SS | SS | SS | SS | 0.00002 |
| L100<br>Aml5/B5 | SS | SS | 0.00386 | SS | SS | SS | SS | SS | SS | SS | SS | SS | SS | SS | SS |
| L100<br>Aml5/H12.5 | SS | SS | 0.00386 | SS | SS | SS | SS | SS | SS | SS | SS | SS | SS | SS | 0.00002 |
| L100<br>B5/H12.5 | SS | SS | 0.00630 | SS | SS | SS | SS | SS | SS | SS | SS | SS | SS | SS | 0.00004 |

**Al300 =** aliskiren 300 mg; **Aml5** = amlodipine 5 mg; **B5** = bisoprolol 5 mg; **E20** = enalapril 20 mg; **H12.5** = hydrochlorothiazide 12.5 mg; **L100** = losartan 100 mg; **SS** = statistically significant ($P$ < 0.00001)

**Figure S35.** Simulated change in plasma aldosterone from baseline to week 4 (mean ± SD, $n$ = 100)

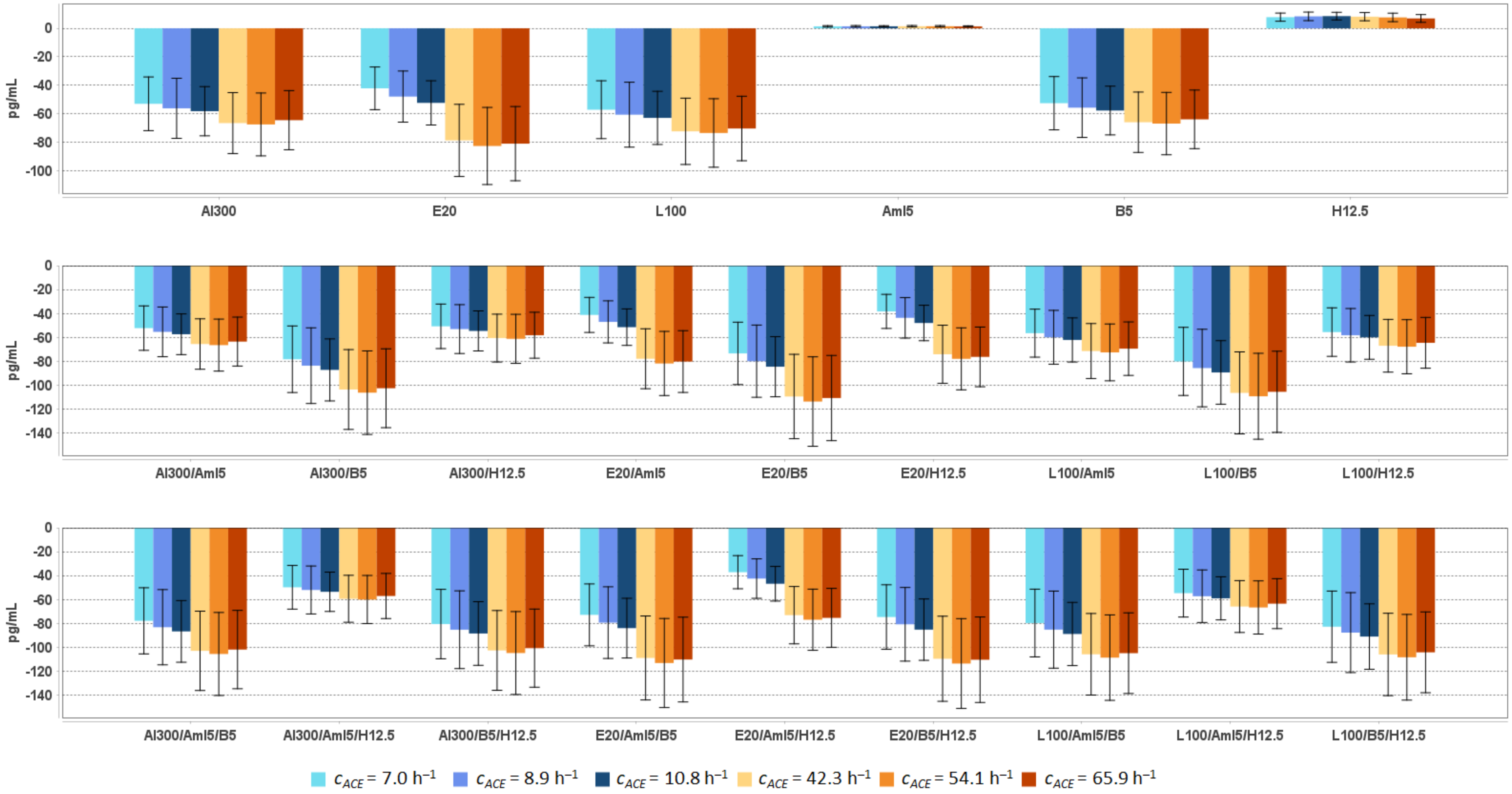


**Al300** = aliskiren 300 mg; **Aml5** = amlodipine 5 mg; **B5** = bisoprolol 5 mg; **E20** = enalapril 20 mg; **H12.5** = hydrochlorothiazide 12.5 mg; **L100** = losartan 100 mg

**Table S56.** Simulated response of <u>plasma aldosterone</u> to antihypertensive therapy in virtual hypertensive populations ($n$ = 100) with different ACE activity, including $P$-values (Kolmogorov-Smirnov test) for endpoint *vs*. baseline; data are presented as mean ± SD in pg/mL

| Regimens | $c_{ACE}$ = 7.0 h$^{-1}$ | | | $c_{ACE}$ = 8.9 h$^{-1}$ | | | $c_{ACE}$ = 10.8 h$^{-1}$ | | | $c_{ACE}$ = 42.3 h$^{-1}$ | | | $c_{ACE}$ = 54.1 h$^{-1}$ | | | $c_{ACE}$ = 65.9 h$^{-1}$ | | |
|---|---|---|---|---|---|---|---|---|---|---|---|---|---|---|---|---|---|---|
| | **Value** | **Change** | ***P*** | **Value** | **Change** | ***P*** | **Value** | **Change** | ***P*** | **Value** | **Change** | ***P*** | **Value** | **Change** | ***P*** | **Value** | **Change** | ***P*** |
| Baseline | 166 ± 59 | – | – | 172 ± 63 | – | – | 176 ± 52 | – | – | 193 ± 62 | – | – | 196 ± 64 | – | – | 188 ± 60 | – | – |
| Al300 | 113 ± 40 | -53 ± 19 | SS | 116 ± 42 | -56 ± 21 | SS | 118 ± 35 | -58 ± 17 | SS | 127 ± 41 | -67 ± 21 | SS | 129 ± 42 | -68 ± 22 | SS | 123 ± 40 | -65 ± 21 | SS |
| E20 | 124 ± 44 | -42 ± 15 | SS | 124 ± 45 | -48 ± 18 | 0.00013 | 124 ± 37 | -52 ± 16 | SS | 114 ± 37 | -79 ± 25 | SS | 113 ± 37 | -83 ± 27 | SS | 107 ± 34 | -81 ± 26 | SS |
| L100 | 109 ± 39 | -57 ± 20 | SS | 111 ± 41 | -61 ± 23 | SS | 113 ± 34 | -63 ± 19 | SS | 121 ± 39 | -72 ± 23 | SS | 123 ± 40 | -74 ± 24 | SS | 118 ± 38 | -70 ± 23 | SS |
| Aml5 | 167 ± 60 | 1 ± 1 | 1.00000 | 173 ± 64 | 1 ± 1 | 0.99963 | 177 ± 53 | 1 ± 0 | 0.99963 | 195 ± 63 | 1 ± 1 | 1.00000 | 198 ± 65 | 1 ± 1 | 0.99376 | 189 ± 61 | 1 ± 0 | 1.00000 |
| B5 | 113 ± 41 | -53 ± 19 | SS | 116 ± 42 | -56 ± 21 | SS | 118 ± 35 | -58 ± 17 | SS | 127 ± 41 | -66 ± 21 | SS | 129 ± 42 | -67 ± 22 | SS | 124 ± 40 | -64 ± 21 | SS |
| H12.5 | 174 ± 61 | 8 ± 3 | 0.81275 | 180 ± 65 | 8 ± 3 | 0.81275 | 185 ± 54 | 9 ± 3 | 0.69937 | 201 ± 64 | 8 ± 3 | 0.58062 | 204 ± 66 | 8 ± 3 | 0.81275 | 195 ± 62 | 7 ± 3 | 0.96707 |
| Al300 Aml5 | 114 ± 41 | -52 ± 19 | SS | 117 ± 43 | -55 ± 21 | SS | 119 ± 35 | -57 ± 17 | SS | 128 ± 41 | -66 ± 21 | SS | 130 ± 42 | -66 ± 22 | SS | 125 ± 40 | -64 ± 20 | SS |
| Al300 B5 | 88 ± 32 | -78 ± 28 | SS | 88 ± 32 | -84 ± 32 | SS | 89 ± 27 | -87 ± 26 | SS | 90 ± 29 | -104 ± 33 | SS | 90 ± 29 | -106 ± 35 | SS | 86 ± 28 | -103 ± 33 | SS |
| Al300 H12.5 | 115 ± 41 | -51 ± 19 | SS | 119 ± 43 | -53 ± 20 | 0.00002 | 121 ± 35 | -55 ± 17 | SS | 133 ± 42 | -61 ± 20 | SS | 135 ± 44 | -61 ± 21 | SS | 130 ± 41 | -58 ± 19 | SS |
| E20 Aml5 | 125 ± 45 | -41 ± 15 | 0.00002 | 125 ± 46 | -47 ± 18 | 0.00013 | 125 ± 37 | -51 ± 15 | SS | 115 ± 37 | -78 ± 25 | SS | 114 ± 37 | -82 ± 27 | SS | 108 ± 35 | -80 ± 26 | SS |
| E20 B5 | 93 ± 33 | -73 ± 26 | SS | 92 ± 33 | -80 ± 30 | SS | 92 ± 27 | -85 ± 25 | SS | 84 ± 27 | -109 ± 35 | SS | 83 ± 27 | -114 ± 37 | SS | 77 ± 25 | -111 ± 36 | SS |
| E20 H12.5 | 128 ± 45 | -38 ± 14 | 0.00013 | 128 ± 46 | -44 ± 17 | 0.00045 | 128 ± 37 | -48 ± 15 | SS | 119 ± 38 | -74 ± 24 | SS | 118 ± 38 | -78 ± 26 | SS | 112 ± 36 | -76 ± 25 | SS |
| L100 Aml5 | 110 ± 39 | -56 ± 20 | SS | 112 ± 41 | -60 ± 23 | SS | 114 ± 34 | -62 ± 18 | SS | 122 ± 39 | -71 ± 23 | SS | 124 ± 40 | -73 ± 24 | SS | 119 ± 38 | -69 ± 22 | SS |
| L100 B5 | 86 ± 31 | -80 ± 28 | SS | 86 ± 31 | -86 ± 32 | SS | 87 ± 26 | -89 ± 27 | SS | 87 ± 28 | -106 ± 34 | SS | 87 ± 28 | -109 ± 36 | SS | 83 ± 27 | -106 ± 34 | SS |
| L100 H12.5 | 110 ± 39 | -56 ± 20 | SS | 114 ± 41 | -58 ± 22 | SS | 116 ± 34 | -60 ± 18 | SS | 126 ± 40 | -67 ± 22 | SS | 128 ± 42 | -68 ± 23 | SS | 123 ± 39 | -65 ± 21 | SS |
| Al300 Aml5/B5 | 88 ± 32 | -78 ± 28 | SS | 89 ± 32 | -83 ± 31 | SS | 89 ± 27 | -87 ± 26 | SS | 90 ± 29 | -103 ± 33 | SS | 91 ± 30 | -106 ± 35 | SS | 86 ± 28 | -102 ± 33 | SS |
| Al300 Aml5/H12.5 | 116 ± 41 | -50 ± 18 | SS | 120 ± 43 | -52 ± 20 | 0.00004 | 122 ± 36 | -54 ± 16 | SS | 134 ± 43 | -59 ± 20 | SS | 136 ± 44 | -60 ± 20 | SS | 131 ± 42 | -57 ± 19 | SS |
| Al300 B5/H12.5 | 85 ± 30 | -81 ± 29 | SS | 87 ± 31 | -85 ± 33 | SS | 88 ± 26 | -89 ± 27 | SS | 91 ± 29 | -103 ± 33 | SS | 91 ± 30 | -105 ± 35 | SS | 87 ± 28 | -101 ± 33 | SS |
| E20 Aml5/B5 | 93 ± 34 | -73 ± 26 | SS | 93 ± 33 | -79 ± 30 | SS | 92 ± 28 | -84 ± 25 | SS | 84 ± 27 | -109 ± 35 | SS | 83 ± 27 | -113 ± 37 | SS | 78 ± 25 | -110 ± 35 | SS |
| E20 Aml5/H12.5 | 129 ± 45 | -37 ± 14 | 0.00025 | 129 ± 47 | -43 ± 17 | 0.00136 | 129 ± 38 | -47 ± 15 | SS | 120 ± 38 | -73 ± 24 | SS | 119 ± 39 | -77 ± 26 | SS | 113 ± 36 | -75 ± 25 | SS |
| E20 B5/H12.5 | 91 ± 32 | -75 ± 27 | SS | 91 ± 33 | -81 ± 31 | SS | 91 ± 27 | -85 ± 26 | SS | 84 ± 26 | -110 ± 36 | SS | 83 ± 27 | -114 ± 38 | SS | 78 ± 25 | -110 ± 36 | SS |
| L100 Aml5/B5 | 86 ± 31 | -80 ± 28 | SS | 87 ± 31 | -85 ± 32 | SS | 87 ± 26 | -89 ± 26 | SS | 87 ± 28 | -106 ± 34 | SS | 88 ± 29 | -109 ± 36 | SS | 83 ± 27 | -105 ± 34 | SS |
| L100 Aml5/H12.5 | 111 ± 39 | -55 ± 20 | SS | 115 ± 41 | -57 ± 22 | SS | 117 ± 34 | -59 ± 18 | SS | 127 ± 40 | -66 ± 22 | SS | 130 ± 42 | -67 ± 22 | SS | 125 ± 40 | -64 ± 21 | SS |
| L100 B5/H12.5 | 83 ± 30 | -83 ± 30 | SS | 84 ± 30 | -88 ± 33 | SS | 85 ± 25 | -91 ± 27 | SS | 87 ± 28 | -106 ± 34 | SS | 88 ± 28 | -108 ± 36 | SS | 84 ± 27 | -104 ± 34 | SS |

**Al300 =** aliskiren 300 mg; **Aml5** = amlodipine 5 mg; **B5** = bisoprolol 5 mg; **E20** = enalapril 20 mg; **H12.5** = hydrochlorothiazide 12.5 mg; **L100** = losartan 100 mg; **SS** = statistically significant ($P$ < 0.00001)

**Table S57.** *P*-values calculated using the Kolmogorov-Smirnov test for changes in plasma aldosterone in populations ($n$ = 100) with different ACE activity receiving the same regimens (case 1: $c_{ACE}$ = 7.0 $h^{-1}$, case 2: $c_{ACE}$ = 8.9 $h^{-1}$, case 3: $c_{ACE}$ = 10.8 $h^{-1}$, case 4: $c_{ACE}$ = 42.3 $h^{-1}$, case 5: $c_{ACE}$ = 54.1 $h^{-1}$, case 6: $c_{ACE}$ = 65.9 $h^{-1}$; *P*-value for case *i* *vs*. case *j* is denoted $P_{ij}$)

| Regimens | $P_{12}$ | $P_{13}$ | $P_{23}$ | $P_{14}$ | $P_{15}$ | $P_{16}$ | $P_{24}$ | $P_{25}$ | $P_{26}$ | $P_{34}$ | $P_{35}$ | $P_{36}$ | $P_{45}$ | $P_{46}$ | $P_{56}$ |
|---|---|---|---|---|---|---|---|---|---|---|---|---|---|---|---|
| Al300 | 0.58062 | 0.02431 | 0.05410 | 0.00007 | SS | 0.00136 | 0.00232 | 0.00079 | 0.01581 | 0.00045 | 0.00013 | 0.02431 | 0.90621 | 0.46756 | 0.46756 |
| E20 | 0.11113 | 0.00002 | 0.01008 | SS | SS | SS | SS | SS | SS | SS | SS | SS | 0.46756 | 0.81275 | 0.81275 |
| L100 | 0.58062 | 0.02431 | 0.05410 | 0.00007 | SS | 0.00136 | 0.00136 | 0.00045 | 0.01581 | 0.00025 | 0.00007 | 0.02431 | 0.90621 | 0.46756 | 0.46756 |
| Aml5 | 0.81275 | 0.58062 | 0.28093 | 0.15454 | 0.21055 | 0.81275 | 0.15454 | 0.28093 | 0.46756 | 0.58062 | 0.69937 | 0.69937 | 0.90621 | 0.11113 | 0.05410 |
| B5 | 0.58062 | 0.02431 | 0.05410 | 0.00007 | SS | 0.00136 | 0.00232 | 0.00079 | 0.01581 | 0.00045 | 0.00013 | 0.02431 | 0.90621 | 0.46756 | 0.46756 |
| H12.5 | 0.46756 | 0.15454 | 0.81275 | 0.90621 | 0.81275 | 0.11113 | 0.90621 | 0.21055 | 0.00386 | 0.46756 | 0.03663 | 0.00025 | 0.28093 | 0.01008 | 0.21055 |
| Al300 Aml5 | 0.58062 | 0.02431 | 0.05410 | 0.00007 | SS | 0.00136 | 0.00232 | 0.00079 | 0.01581 | 0.00045 | 0.00013 | 0.02431 | 0.90621 | 0.46756 | 0.36672 |
| Al300 B5 | 0.46756 | 0.01008 | 0.02431 | SS | SS | 0.00002 | 0.00025 | 0.00004 | 0.00232 | 0.00004 | SS | 0.00136 | 0.90621 | 0.69937 | 0.69937 |
| Al300 H12.5 | 0.69937 | 0.05410 | 0.05410 | 0.00025 | 0.00013 | 0.03663 | 0.00630 | 0.00630 | 0.07832 | 0.00232 | 0.00630 | 0.15454 | 0.99376 | 0.46756 | 0.28093 |
| E20 Aml5 | 0.11113 | 0.00002 | 0.01008 | SS | SS | SS | SS | SS | SS | SS | SS | SS | 0.46756 | 0.81275 | 0.81275 |
| E20 B5 | 0.36672 | 0.00232 | 0.01581 | SS | SS | SS | SS | SS | SS | SS | SS | SS | 0.58062 | 0.96707 | 0.81275 |
| E20 H12.5 | 0.15454 | SS | 0.00630 | SS | SS | SS | SS | SS | SS | SS | SS | SS | 0.36672 | 0.69937 | 0.69937 |
| L100 Aml5 | 0.58062 | 0.02431 | 0.05410 | 0.00007 | SS | 0.00136 | 0.00136 | 0.00045 | 0.01581 | 0.00025 | 0.00007 | 0.02431 | 0.90621 | 0.46756 | 0.46756 |
| L100 B5 | 0.46756 | 0.01008 | 0.02431 | SS | SS | SS | 0.00025 | 0.00004 | 0.00232 | 0.00004 | SS | 0.00079 | 0.90621 | 0.69937 | 0.69937 |
| L100 H12.5 | 0.69937 | 0.05410 | 0.05410 | 0.00025 | 0.00007 | 0.02431 | 0.00630 | 0.00386 | 0.05410 | 0.00136 | 0.00232 | 0.15454 | 0.96707 | 0.46756 | 0.36672 |
| Al300 Aml5/B5 | 0.46756 | 0.01008 | 0.02431 | SS | SS | 0.00002 | 0.00025 | 0.00004 | 0.00386 | 0.00004 | SS | 0.00136 | 0.90621 | 0.69937 | 0.69937 |
| Al300 Aml5/H12.5 | 0.69937 | 0.05410 | 0.05410 | 0.00025 | 0.00013 | 0.03663 | 0.00630 | 0.00630 | 0.07832 | 0.00232 | 0.00386 | 0.15454 | 0.99376 | 0.46756 | 0.28093 |
| Al300 B5/H12.5 | 0.69937 | 0.03663 | 0.05410 | 0.00004 | SS | 0.00045 | 0.00079 | 0.00025 | 0.01008 | 0.00025 | 0.00007 | 0.00630 | 0.90621 | 0.69937 | 0.46756 |
| E20 Aml5/B5 | 0.36672 | 0.00232 | 0.01581 | SS | SS | SS | SS | SS | SS | SS | SS | SS | 0.58062 | 0.96707 | 0.81275 |
| E20 Aml5/H12.5 | 0.15454 | SS | 0.00630 | SS | SS | SS | SS | SS | SS | SS | SS | SS | 0.36672 | 0.81275 | 0.69937 |
| E20 B5/H12.5 | 0.36672 | 0.00630 | 0.03663 | SS | SS | SS | 0.00002 | SS | SS | SS | SS | SS | 0.81275 | 0.90621 | 0.81275 |
| L100 Aml5/B5 | 0.46756 | 0.01008 | 0.02431 | SS | SS | SS | 0.00025 | 0.00004 | 0.00232 | 0.00004 | SS | 0.00079 | 0.90621 | 0.81275 | 0.69937 |
| L100 Aml5/H12.5 | 0.69937 | 0.05410 | 0.05410 | 0.00025 | 0.00007 | 0.01581 | 0.00630 | 0.00386 | 0.05410 | 0.00136 | 0.00232 | 0.11113 | 0.96707 | 0.46756 | 0.36672 |
| L100 B5/H12.5 | 0.58062 | 0.03663 | 0.05410 | 0.00004 | SS | 0.00045 | 0.00079 | 0.00025 | 0.01008 | 0.00013 | 0.00004 | 0.00630 | 0.90621 | 0.69937 | 0.58062 |

**Al300 =** aliskiren 300 mg; **Aml5** = amlodipine 5 mg; **B5** = bisoprolol 5 mg; **E20** = enalapril 20 mg; **H12.5** = hydrochlorothiazide 12.5 mg; **L100** = losartan 100 mg; **SS** = statistically significant ($P$ < 0.00001)

**Table S58.** Pearson correlation coefficients of RAAS parameters with a decrease in diastolic blood pressure during simulated treatment of virtual populations (*n* = 100) with aliskiren 300 mg (A300), amlodipine 5 mg (A5), bisoprolol 5 mg (B5), enalapril 20 mg (E20), HCTZ 12.5 mg (H12.5), losartan 100 mg (L100), and combinations of these drugs.

| $c_{ACE}$ ($h^{-1}$) | A300 | A5 | B5 | E20 | H12.5 | L100 | A300 A5 | A300 B5 | A300 H12.5 | E20 A5 | E20 B5 | E20 H12.5 | L100 A5 | L100 B5 | L100 H12.5 | A300 A5 B5 | A300 A5 H12.5 | A300 B5 H12.5 | E20 A5 B5 | E20 A5 H12.5 | E20 B5 H12.5 | L100 A5 B5 | L100 A5 H12.5 | L100 B5 H12.5 |
|---|---|---|---|---|---|---|---|---|---|---|---|---|---|---|---|---|---|---|---|---|---|---|---|---|
| ***Plasma renin activity*** | | | | | | | | | | | | | | | | | | | | | | | | |
| **7.0** | 0.25 | -0.07 | 0.18 | 0.32 | 0.05 | 0.21 | 0.15 | -0.35 | 0.20 | 0.16 | -0.15 | 0.19 | 0.13 | -0.43 | 0.20 | -0.34 | 0.15 | 0.04 | -0.14 | 0.14 | 0.12 | -0.43 | 0.15 | -0.01 |
| **8.9** | 0.42 | -0.15 | 0.37 | 0.47 | -0.27 | 0.38 | 0.25 | -0.23 | 0.07 | 0.25 | -0.03 | 0.05 | 0.24 | -0.35 | 0.08 | -0.14 | 0.00 | 0.00 | 0.02 | -0.02 | 0.04 | -0.26 | 0.01 | -0.05 |
| **10.8** | 0.40 | -0.13 | 0.45 | 0.44 | -0.22 | 0.35 | 0.18 | -0.13 | 0.04 | 0.20 | 0.00 | 0.04 | 0.15 | -0.25 | 0.04 | -0.15 | -0.06 | -0.02 | -0.05 | -0.06 | 0.01 | -0.24 | -0.05 | -0.06 |
| **42.3** | 0.49 | -0.08 | 0.41 | 0.45 | 0.01 | 0.47 | 0.31 | 0.35 | 0.29 | 0.29 | 0.30 | 0.30 | 0.31 | 0.33 | 0.29 | 0.28 | 0.19 | 0.29 | 0.24 | 0.19 | 0.28 | 0.26 | 0.19 | 0.29 |
| **54.1** | 0.68 | -0.08 | 0.63 | 0.62 | -0.21 | 0.66 | 0.51 | 0.48 | 0.29 | 0.48 | 0.42 | 0.32 | 0.50 | 0.46 | 0.31 | 0.41 | 0.19 | 0.30 | 0.36 | 0.21 | 0.30 | 0.39 | 0.20 | 0.30 |
| **65.9** | 0.71 | -0.01 | 0.56 | 0.67 | -0.11 | 0.70 | 0.65 | 0.53 | 0.43 | 0.61 | 0.46 | 0.46 | 0.64 | 0.51 | 0.44 | 0.49 | 0.40 | 0.43 | 0.43 | 0.42 | 0.42 | 0.47 | 0.42 | 0.43 |
| ***Plasma angiotensin I*** | | | | | | | | | | | | | | | | | | | | | | | | |
| **7.0** | 0.25 | -0.07 | 0.18 | 0.32 | 0.05 | 0.21 | 0.15 | -0.35 | 0.20 | 0.16 | -0.15 | 0.19 | 0.13 | -0.43 | 0.20 | -0.34 | 0.15 | 0.04 | -0.14 | 0.14 | 0.12 | -0.43 | 0.15 | -0.01 |
| **8.9** | 0.42 | -0.15 | 0.37 | 0.47 | -0.27 | 0.38 | 0.25 | -0.23 | 0.07 | 0.25 | -0.03 | 0.05 | 0.24 | -0.35 | 0.08 | -0.14 | 0.00 | 0.00 | 0.02 | -0.02 | 0.04 | -0.26 | 0.01 | -0.05 |
| **10.8** | 0.40 | -0.13 | 0.45 | 0.44 | -0.22 | 0.35 | 0.18 | -0.13 | 0.04 | 0.20 | 0.00 | 0.04 | 0.15 | -0.25 | 0.04 | -0.15 | -0.06 | -0.02 | -0.05 | -0.06 | 0.01 | -0.24 | -0.05 | -0.06 |
| **42.3** | 0.49 | -0.08 | 0.41 | 0.45 | 0.01 | 0.47 | 0.31 | 0.35 | 0.29 | 0.29 | 0.30 | 0.30 | 0.31 | 0.33 | 0.29 | 0.28 | 0.19 | 0.29 | 0.24 | 0.19 | 0.28 | 0.26 | 0.19 | 0.29 |
| **54.1** | 0.68 | -0.08 | 0.63 | 0.62 | -0.21 | 0.66 | 0.51 | 0.48 | 0.29 | 0.48 | 0.42 | 0.32 | 0.50 | 0.46 | 0.31 | 0.41 | 0.19 | 0.30 | 0.36 | 0.21 | 0.30 | 0.39 | 0.20 | 0.30 |
| **65.9** | 0.71 | -0.01 | 0.56 | 0.67 | -0.11 | 0.70 | 0.65 | 0.53 | 0.43 | 0.61 | 0.46 | 0.46 | 0.64 | 0.51 | 0.44 | 0.49 | 0.40 | 0.43 | 0.43 | 0.42 | 0.42 | 0.47 | 0.42 | 0.43 |
| ***Plasma angiotensin II*** | | | | | | | | | | | | | | | | | | | | | | | | |
| **7.0** | 0.25 | -0.07 | 0.18 | 0.32 | 0.05 | 0.21 | 0.15 | -0.35 | 0.20 | 0.16 | -0.15 | 0.19 | 0.13 | -0.43 | 0.20 | -0.34 | 0.15 | 0.04 | -0.14 | 0.14 | 0.12 | -0.43 | 0.15 | -0.01 |
| **8.9** | 0.42 | -0.15 | 0.37 | 0.47 | -0.27 | 0.38 | 0.25 | -0.23 | 0.07 | 0.25 | -0.03 | 0.05 | 0.24 | -0.35 | 0.08 | -0.14 | 0.00 | 0.00 | 0.02 | -0.02 | 0.04 | -0.26 | 0.01 | -0.05 |
| **10.8** | 0.40 | -0.13 | 0.45 | 0.44 | -0.22 | 0.35 | 0.18 | -0.13 | 0.04 | 0.20 | 0.00 | 0.04 | 0.15 | -0.25 | 0.04 | -0.15 | -0.06 | -0.02 | -0.05 | -0.06 | 0.01 | -0.24 | -0.05 | -0.06 |
| **42.3** | 0.49 | -0.08 | 0.41 | 0.45 | 0.01 | 0.47 | 0.31 | 0.35 | 0.29 | 0.29 | 0.30 | 0.30 | 0.31 | 0.33 | 0.29 | 0.28 | 0.19 | 0.29 | 0.24 | 0.19 | 0.28 | 0.26 | 0.19 | 0.29 |
| **54.1** | 0.68 | -0.08 | 0.63 | 0.62 | -0.21 | 0.66 | 0.51 | 0.48 | 0.29 | 0.48 | 0.42 | 0.32 | 0.50 | 0.46 | 0.31 | 0.41 | 0.19 | 0.30 | 0.36 | 0.21 | 0.30 | 0.39 | 0.20 | 0.30 |
| **65.9** | 0.71 | -0.01 | 0.56 | 0.67 | -0.11 | 0.70 | 0.65 | 0.53 | 0.43 | 0.61 | 0.46 | 0.46 | 0.64 | 0.51 | 0.44 | 0.49 | 0.40 | 0.43 | 0.43 | 0.42 | 0.42 | 0.47 | 0.42 | 0.43 |
| ***Plasma aldosterone*** | | | | | | | | | | | | | | | | | | | | | | | | |
| **7.0** | -0.02 | 0.02 | -0.12 | -0.03 | 0.09 | -0.01 | 0.03 | -0.04 | 0.09 | 0.02 | -0.07 | 0.09 | 0.03 | -0.02 | 0.09 | -0.03 | 0.10 | 0.06 | -0.04 | 0.09 | 0.06 | -0.02 | 0.10 | 0.06 |
| **8.9** | -0.17 | -0.06 | -0.17 | -0.15 | 0.05 | -0.18 | -0.11 | -0.27 | 0.00 | -0.10 | -0.26 | 0.01 | -0.12 | -0.27 | -0.01 | -0.22 | 0.05 | -0.08 | -0.20 | 0.05 | -0.06 | -0.23 | 0.05 | -0.09 |
| **10.8** | -0.01 | 0.02 | -0.01 | -0.01 | 0.05 | -0.02 | 0.01 | -0.08 | 0.06 | 0.01 | -0.07 | 0.06 | 0.00 | -0.10 | 0.06 | -0.05 | 0.05 | 0.03 | -0.04 | 0.05 | 0.03 | -0.06 | 0.05 | 0.02 |
| **42.3** | 0.04 | -0.12 | 0.04 | 0.03 | -0.07 | 0.03 | -0.03 | 0.04 | 0.01 | -0.02 | 0.03 | 0.01 | -0.02 | 0.04 | 0.01 | 0.01 | -0.03 | 0.04 | 0.01 | -0.03 | 0.04 | 0.01 | -0.03 | 0.04 |
| **54.1** | 0.08 | 0.02 | 0.03 | 0.09 | -0.01 | 0.08 | 0.10 | 0.08 | 0.06 | 0.12 | 0.08 | 0.08 | 0.11 | 0.08 | 0.07 | 0.13 | 0.08 | 0.11 | 0.13 | 0.10 | 0.12 | 0.13 | 0.08 | 0.12 |
| **65.9** | 0.01 | -0.05 | 0.03 | 0.00 | 0.05 | 0.00 | -0.02 | 0.01 | 0.06 | -0.02 | 0.01 | 0.05 | -0.02 | 0.01 | 0.05 | -0.02 | 0.05 | 0.04 | -0.03 | 0.05 | 0.03 | -0.02 | 0.05 | 0.04 |

**Table S59.** Pearson correlation coefficients of RAAS parameters with a decrease in systolic blood pressure during simulated treatment of virtual populations ($n$ = 100) with aliskiren 300 mg (A300), amlodipine 5 mg (A5), bisoprolol 5 mg (B5), enalapril 20 mg (E20), HCTZ 12.5 mg (H12.5), losartan 100 mg (L100), and combinations of these drugs.

| $c_{ACE}$ ($h^{-1}$) | A300 | A5 | B5 | E20 | H12.5 | L100 | A300 A5 | A300 B5 | A300 H12.5 | E20 A5 | E20 B5 | E20 H12.5 | L100 A5 | L100 B5 | L100 H12.5 | A300 A5 B5 | A300 A5 H12.5 | A300 B5 H12.5 | E20 A5 B5 | E20 A5 H12.5 | E20 B5 H12.5 | L100 A5 B5 | L100 A5 H12.5 | L100 B5 H12.5 |
|---|---|---|---|---|---|---|---|---|---|---|---|---|---|---|---|---|---|---|---|---|---|---|---|---|
| ***Plasma renin activity*** | | | | | | | | | | | | | | | | | | | | | | | | |
| **7.0** | 0.23 | -0.05 | 0.20 | 0.19 | 0.15 | 0.24 | 0.15 | 0.27 | 0.20 | 0.10 | 0.28 | 0.18 | 0.17 | 0.25 | 0.21 | 0.22 | 0.16 | 0.26 | 0.23 | 0.14 | 0.25 | 0.22 | 0.17 | 0.26 |
| **8.9** | 0.24 | -0.13 | 0.19 | 0.19 | -0.26 | 0.26 | 0.13 | 0.37 | -0.16 | 0.07 | 0.35 | -0.20 | 0.16 | 0.37 | -0.14 | 0.29 | -0.16 | 0.06 | 0.27 | -0.19 | 0.02 | 0.29 | -0.13 | 0.08 |
| **10.8** | 0.17 | -0.14 | 0.05 | 0.14 | -0.11 | 0.20 | 0.04 | 0.28 | -0.06 | -0.01 | 0.27 | -0.08 | 0.07 | 0.29 | -0.04 | 0.20 | -0.10 | 0.07 | 0.19 | -0.11 | 0.05 | 0.21 | -0.08 | 0.08 |
| **42.3** | 0.18 | 0.02 | 0.10 | 0.24 | 0.12 | 0.21 | 0.16 | 0.31 | 0.14 | 0.22 | 0.33 | 0.17 | 0.19 | 0.32 | 0.15 | 0.30 | 0.10 | 0.26 | 0.31 | 0.13 | 0.28 | 0.30 | 0.12 | 0.27 |
| **54.1** | 0.01 | -0.11 | -0.04 | 0.13 | -0.11 | 0.06 | -0.10 | 0.26 | -0.16 | 0.01 | 0.29 | -0.12 | -0.05 | 0.27 | -0.15 | 0.17 | -0.19 | 0.02 | 0.21 | -0.16 | 0.08 | 0.19 | -0.18 | 0.04 |
| **65.9** | -0.01 | -0.07 | -0.05 | 0.15 | 0.06 | 0.05 | 0.00 | 0.29 | 0.02 | 0.15 | 0.34 | 0.10 | 0.06 | 0.31 | 0.05 | 0.31 | 0.03 | 0.26 | 0.36 | 0.09 | 0.32 | 0.33 | 0.05 | 0.29 |
| ***Plasma angiotensin I*** | | | | | | | | | | | | | | | | | | | | | | | | |
| **7.0** | 0.23 | -0.05 | 0.20 | 0.19 | 0.15 | 0.24 | 0.15 | 0.27 | 0.20 | 0.10 | 0.28 | 0.18 | 0.17 | 0.25 | 0.21 | 0.22 | 0.16 | 0.26 | 0.23 | 0.14 | 0.25 | 0.22 | 0.17 | 0.26 |
| **8.9** | 0.24 | -0.13 | 0.19 | 0.19 | -0.26 | 0.26 | 0.13 | 0.37 | -0.16 | 0.07 | 0.35 | -0.20 | 0.16 | 0.37 | -0.14 | 0.29 | -0.16 | 0.06 | 0.27 | -0.19 | 0.02 | 0.29 | -0.13 | 0.08 |
| **10.8** | 0.17 | -0.14 | 0.05 | 0.14 | -0.11 | 0.20 | 0.04 | 0.28 | -0.06 | -0.01 | 0.27 | -0.08 | 0.07 | 0.29 | -0.04 | 0.20 | -0.10 | 0.07 | 0.19 | -0.11 | 0.05 | 0.21 | -0.08 | 0.08 |
| **42.3** | 0.18 | 0.02 | 0.10 | 0.24 | 0.12 | 0.21 | 0.16 | 0.31 | 0.14 | 0.22 | 0.33 | 0.17 | 0.19 | 0.32 | 0.15 | 0.30 | 0.10 | 0.26 | 0.31 | 0.13 | 0.28 | 0.30 | 0.12 | 0.27 |
| **54.1** | 0.01 | -0.11 | -0.04 | 0.13 | -0.11 | 0.06 | -0.10 | 0.26 | -0.16 | 0.01 | 0.29 | -0.12 | -0.05 | 0.27 | -0.15 | 0.17 | -0.19 | 0.02 | 0.21 | -0.16 | 0.08 | 0.19 | -0.18 | 0.04 |
| **65.9** | -0.01 | -0.07 | -0.05 | 0.15 | 0.06 | 0.05 | 0.00 | 0.29 | 0.02 | 0.15 | 0.34 | 0.10 | 0.06 | 0.31 | 0.05 | 0.31 | 0.03 | 0.26 | 0.36 | 0.09 | 0.32 | 0.33 | 0.05 | 0.29 |
| ***Plasma angiotensin II*** | | | | | | | | | | | | | | | | | | | | | | | | |
| **7.0** | 0.23 | -0.05 | 0.20 | 0.19 | 0.15 | 0.24 | 0.15 | 0.27 | 0.20 | 0.10 | 0.28 | 0.18 | 0.17 | 0.25 | 0.21 | 0.22 | 0.16 | 0.26 | 0.23 | 0.14 | 0.25 | 0.22 | 0.17 | 0.26 |
| **8.9** | 0.24 | -0.13 | 0.19 | 0.19 | -0.26 | 0.26 | 0.13 | 0.37 | -0.16 | 0.07 | 0.35 | -0.20 | 0.16 | 0.37 | -0.14 | 0.29 | -0.16 | 0.06 | 0.27 | -0.19 | 0.02 | 0.29 | -0.13 | 0.08 |
| **10.8** | 0.17 | -0.14 | 0.05 | 0.14 | -0.11 | 0.20 | 0.04 | 0.28 | -0.06 | -0.01 | 0.27 | -0.08 | 0.07 | 0.29 | -0.04 | 0.20 | -0.10 | 0.07 | 0.19 | -0.11 | 0.05 | 0.21 | -0.08 | 0.08 |
| **42.3** | 0.18 | 0.02 | 0.10 | 0.24 | 0.12 | 0.21 | 0.16 | 0.31 | 0.14 | 0.22 | 0.33 | 0.17 | 0.19 | 0.32 | 0.15 | 0.30 | 0.10 | 0.26 | 0.31 | 0.13 | 0.28 | 0.30 | 0.12 | 0.27 |
| **54.1** | 0.01 | -0.11 | -0.04 | 0.13 | -0.11 | 0.06 | -0.10 | 0.26 | -0.16 | 0.01 | 0.29 | -0.12 | -0.05 | 0.27 | -0.15 | 0.17 | -0.19 | 0.02 | 0.21 | -0.16 | 0.08 | 0.19 | -0.18 | 0.04 |
| **65.9** | -0.01 | -0.07 | -0.05 | 0.15 | 0.06 | 0.05 | 0.00 | 0.29 | 0.02 | 0.15 | 0.34 | 0.10 | 0.06 | 0.31 | 0.05 | 0.31 | 0.03 | 0.26 | 0.36 | 0.09 | 0.32 | 0.33 | 0.05 | 0.29 |
| ***Plasma aldosterone*** | | | | | | | | | | | | | | | | | | | | | | | | |
| **7.0** | 0.01 | 0.02 | 0.08 | 0.00 | 0.14 | 0.01 | 0.04 | 0.07 | 0.14 | 0.03 | 0.07 | 0.14 | 0.04 | 0.07 | 0.14 | 0.09 | 0.12 | 0.16 | 0.09 | 0.12 | 0.15 | 0.09 | 0.12 | 0.16 |
| **8.9** | -0.13 | 0.05 | -0.10 | -0.14 | 0.17 | -0.12 | -0.03 | -0.06 | 0.13 | -0.03 | -0.07 | 0.14 | -0.03 | -0.05 | 0.12 | -0.03 | 0.14 | 0.08 | -0.03 | 0.15 | 0.09 | -0.02 | 0.13 | 0.07 |
| **10.8** | -0.12 | 0.11 | -0.08 | -0.13 | 0.06 | -0.11 | -0.04 | -0.02 | 0.02 | -0.03 | -0.04 | 0.03 | -0.04 | -0.01 | 0.02 | -0.02 | 0.03 | 0.01 | -0.03 | 0.04 | 0.01 | -0.02 | 0.03 | 0.01 |
| **42.3** | 0.08 | -0.07 | 0.05 | 0.11 | -0.05 | 0.10 | 0.04 | 0.12 | -0.01 | 0.07 | 0.12 | 0.01 | 0.06 | 0.12 | 0.00 | 0.11 | -0.02 | 0.07 | 0.12 | 0.01 | 0.09 | 0.12 | -0.01 | 0.08 |
| **54.1** | 0.01 | -0.12 | 0.04 | 0.01 | -0.07 | 0.01 | -0.09 | 0.04 | -0.07 | -0.07 | 0.05 | -0.07 | -0.09 | 0.04 | -0.07 | -0.02 | -0.08 | -0.05 | -0.01 | -0.08 | -0.04 | -0.02 | -0.08 | -0.05 |
| **65.9** | -0.04 | -0.11 | -0.05 | -0.04 | 0.04 | -0.04 | -0.11 | -0.04 | 0.03 | -0.09 | -0.04 | 0.03 | -0.10 | -0.04 | 0.03 | -0.07 | -0.01 | 0.02 | -0.05 | -0.02 | 0.02 | -0.06 | -0.01 | 0.02 |
