## Supplementary File 2 for "Mathematical modeling of the influence of *ACE I/D* polymorphism on blood pressure and antihypertensive therapy"

*Supplementary File 2 – Model Description*

### Mathematical modeling of the influence of *ACE I/D* polymorphism on blood pressure and antihypertensive therapy

**Elena Kutumova*, Anna Kovaleva, Ruslan Sharipov, Galina Lifshits, Fedor Kolpakov**

**Table S1**. Mathematical functions of the model

| Functions | Description |
|---|---|
| $mass_{elasticity}(e,m) = \frac{29.97 \cdot e}{m^{0.75}}$ | Allometric scaling of cardiac and vascular elasticity as a function of body mass *m* and the average normal elasticity *e*. |
| $mass_{volume}(v,m) = \frac{m}{70} \cdot v$ | Allometric scaling relative to body mass *m*, considering the average human weight of 70 kg. |
| $mass_{conductivity}(c,m) = 0.042 \cdot m^{0.75} \cdot c$ | Allometric scaling of cardiac and vascular conductivity as a function of body mass *m* and the average normal conductivity *c*. |
| $r_{plus}(a,b,x,x0) = \frac{1-e^{-a\cdot(x-x0)}}{1+b\cdot e^{-a\cdot(x-x0)}}$ | Receptor activity function. |
| $r_{minus}(a,b,x,x_0) = 1 - r_{plus}(a,b,x,x_0)$ | Receptor activity function. |
| $atrium_{pulse}(T,T_S,P_{AL},t) = \begin{cases} 0, & t \le T - 0.2\cdot(T-T_S) \\ 0.1\cdot P_{AL}\cdot \lvert T-t \rvert^{1/2}, & otherwise \end{cases}$ | Function for calculating pulse waves of the left and right atrium depending on the duration of the cardiac cycle *T*, the actual duration of left or right ventricular systole $T_S$, systemic arterial pressure $P_{AL}$, and the current time of the cardiac cycle *t*. |
| $valve(P_{in},P_{out},R_{factor},YY) = \begin{cases} YY\cdot(P_{in}-P_{out}), & P_{in} \ge P_{out} \\ R_{factor}\cdot YY\cdot(P_{in}-P_{out}), & otherwise \end{cases}$ | Function for calculating blood flow through heart valves taking into account inlet pressure $P_{in}$, outlet pressure $P_{out}$, conductivity *YY* and regurgitation coefficient $R_{factor}$. The second part of the formula describes the reverse flow through the valve (regurgitation, $0 < R_{factor} < 0.3$). Under normal conditions, $R_{factor} = 0$. |
| $sigm(x) = \frac{1}{1+e^{-x}}$ | Logistic function. |
| $\mathrm{sgn}(x,y) = \begin{cases} 0, & x \le 0 \ and \ y \le 0 \\ y, & otherwise \end{cases}$ | Function used to calculate oxygen debt. |

**Table S2**. Model equations

| № | Equations | Description | Modification[1] |
|---|---|---|---|
| **Cardiovascular module, systemic circulation** | | | |
| 001 | $\frac{dV_{AL}}{dt} = F_{HLAL} - F_{ALVL}$ | Change in blood volume in the systemic arteries ($V_{AL}$) is the difference between incoming ($F_{HLAL}$) and outgoing ($F_{ALVL}$) blood flows. | – |
| 002 | $V_{AL}(0) = 0.13 \cdot V$ | Starting value[2] of the blood volume in the systemic arteries is 13% of the total blood volume *V*. | – |
| 003 | $G_{AL} = mass_{elasticity}(G_{AL0} + A_9 \cdot H, m) \cdot (1 - B_{blocker\_st})$ | $G_{AL}$ – systemic arterial elasticity, $G_{AL0}$ – basic elasticity, $A_9$ – systemic arterial tone, $H$ – neurohumoral factor, $m$ – body mass, $B_{blocker\_st}$ – effect of the β-blocker bisoprolol on arterial stiffness. | Reduction of arterial stiffness with bisoprolol (Asmar et al., 1991; Kahonen et al., 2000; Palmieri et al., 2004; Ong et al., 2011; Zhou et al., 2013; Eguchi et al., 2015) was added. |
| 004 | $\omega_{AL} = \omega_{AL\_nom} - mass_{volume}(A_8, m) \cdot H$ | $\omega_{AL}$ – unstressed volume of the systemic arteries, $\omega_{AL\_nom}$ – nominal unstressed volume, $A_8$ – sympathetic sensitivity of the systemic arteries, $m$ – body mass, $H$ – neurohumoral factor. | – |
| 005 | $\omega_{AL\_nom}(0) = k_{AL} \cdot V_{AL}$ | Starting value of the nominal unstressed volume of the systemic arteries without nervous and hormonal influences is a fraction $k_{AL}$ of $V_{AL}$. | – |
| 006 | $P_{AL} = G_{AL} \cdot (V_{AL} - \omega_{AL})$ | $P_{AL}$ – systemic arterial pressure, $G_{AL}$ – systemic arterial elasticity, $V_{AL}$ – systemic arterial volume, $\omega_{AL}$ – unstressed volume of the systemic arteries. | – |
| 007 | $Vis = 1.23 \cdot \left(1 - \frac{Hct}{99}\right)^{-n}, \; n = 1.7 + 9.86 \cdot \exp(-0.0607 \cdot Hct)$ | Blood viscosity *Vis* is calculated as a function of hematocrit *Hct* at a plasma viscosity of 1.23 cP. | – |
| 008 | $F_{ALVL} = \frac{(P_{AL} - P_{VL})}{\frac{Vis}{Vis_{norm}} \cdot R_{ALVL}}$ | $F_{ALVL}$ – blood flow through the systemic microvessels, $R_{ALVL}$ – resistance of the systemic microvessels, $P_{AL}$ – systemic arterial pressure, $P_{VL}$ – systemic venous pressure, $Vis/Vis_{norm}$ – normalized blood viscosity. | – |
| 009 | $R_{ALVL} = \frac{1}{Y_{ALVL}} \cdot \psi_{AT1\_ALVL} \cdot (1 - CCB_{sys}) \cdot (1 - Diuretic_{sys})$ | $R_{ALVL}$ – resistance of the systemic microvessels, $\psi_{AT1\_ALVL}$ – effect of angiotensin II bound to AT1 receptors in the vascular smooth muscle, $CCB_{sys}$ – effect of the calcium channel blocker amlodipine, $Diuretic_{sys}$ – effect of the thiazide diuretic hydrochlorothiazide (HCTZ). | The vasodilatory effects of amlodipine and HCTZ were taken from the model extension (Kutumova et al., 2022) |
| 010 | $\psi_{AT1\_ALVL} = A_{AT1\_ALVL} + B_{AT1\_ALVL} \cdot AT1\_ANGII - \frac{C_{AT1\_ALVL}}{AT1\_ANGII}$ | $\psi_{AT1\_ALVL}$ – effect of AT1-bound angiotensin II (*AT1_ANGII*) on $R_{ALVL}$. $A_{AT1\_ALVL}$, $B_{AT1\_ALVL}$, and $C_{AT1\_ALVL}$ are constant. | – |

[1] Modification from the basic model by Kutumova et al. (2021)
[2] Formulas for calculating starting values are used to generate virtual patients and are not taken into account when considering the model in equilibrium.

| | | | |
|---|---|---|---|
| 011 | $Y_{ALVL} = mass_{conductivity}\left(Y_{ALVL0} - A_3 \cdot H + A_4 \cdot DO_2, m\right)$ | $Y_{ALVL}$ – conductivity of the systemic microvessels, $Y_{ALVL0}$ – basic conductivity, $A_3$ – sympathetic sensitivity of the systemic microvessels (dependence on nervous and hormonal influences), $H$ – neurohumoral factor, $A_4$ – oxygen-deficient sensitivity of the systemic microvessels (dependence on oxygen debt $DO_2$), $m$ – body mass. | – |
| 012 | $V_{VL} = V - V_{AL} - V_{AR} - V_{VR} - V_{HL} - V_{HR}$ | Blood volume in the systemic veins $V_{VL}$ is calculated by reducing the total circulating blood volume $V$ by the volumes of all other parts of the cardiovascular system, including systemic arteries ($V_{AL}$), pulmonary arteries ($V_{AR}$), pulmonary veins ($V_{VR}$), left ventricle ($V_{HL}$) and right ventricle ($V_{HR}$). | – |
| 013 | $G_{VL} = mass_{elasticity}\left(G_{VL0} + A_{11} \cdot H, m\right)$ | $G_{VL}$ – systemic venous elasticity, $G_{VL0}$ – basic elasticity, $A_{11}$ – venous tone, $H$ – neurohumoral factor, $m$ – body mass. | – |
| 014 | $\omega_{VL}(0) = k_{VL} \cdot V_{VL}$ | Starting value of the unstressed volume of the systemic veins, calculated as a fraction $k_{VL}$ of $V_{VL}$. | – |
| 015 | $P_{VL} = P_0 + RA_{PULSE} + G_{VL} \cdot \left(V_{VL} - \omega_{VL}\right)$ | $P_{VL}$ – systemic venous pressure, $P_0$ – basic pressure, $RA_{PULSE}$ – pulse wave of the right atrium, $G_{VL}$ – systemic venous elasticity, $V_{VL}$ – blood volume in the systemic veins, $\omega_{VL}$ – unstressed volume of the systemic veins. | – |
| 016 | $MAP = \dfrac{P_S + 2 \cdot P_D}{3}$ | $MAP$ – mean arterial pressure, $P_S$ – systolic blood pressure, $P_D$ – diastolic blood pressure. | – |
| 017 | $TPR = MAP / CO$ | Total peripheral resistance $TPR$ is the ratio of mean arterial pressure $MAP$ to cardiac output $CO$. | – |
| 018 | $SVR = \dfrac{1}{Y_{HLAL}} + R_{ALVL} + \dfrac{1}{Y_{VLHR}}$ | $SVR$ – systemic vascular resistance, $Y_{HLAL}$ – conductivity (reciprocal of resistance) of the aortic valve and systemic arteries, $R_{ALVL}$ – resistance of the systemic microvessels, $Y_{VLHR}$ – conductivity of the tricuspid valve and systemic veins. | – |
| **Cardiovascular module, pulmonary circulation** | | | |
| 019 | $\dfrac{dV_{AR}}{dt} = F_{HRAR} - F_{ARVR}$ | Change in blood volume in the pulmonary arteries ($V_{AR}$) is the difference between incoming ($F_{HRAR}$) and outgoing ($F_{ARVR}$) blood flows. | – |
| 020 | $V_{AR}(0) = 0.035 \cdot V$ | Starting value of the blood volume in the pulmonary arteries is 3.5% of the total blood volume $V$. | – |
| 021 | $G_{AR} = mass_{elasticity}\left(G_{AR0} + A_{19} \cdot H, m\right)$ | $G_{AR}$ – pulmonary arterial elasticity, $G_{AR0}$ – basic elasticity, $A_{19}$ – pulmonary arterial tone, $H$ – neurohumoral factor, $m$ – body mass. | – |
| 022 | $\omega_{AR} = \omega_{AR\_nom} - mass_{volume}\left(A_{18}, m\right) \cdot H$ | $\omega_{AR}$ – unstressed volume of the pulmonary arteries, $\omega_{AR\_nom}$ – nominal unstressed volume, $A_{18}$ – sympathetic sensitivity of the pulmonary arteries, $m$ – body mass, $H$ – neurohumoral factor. | – |
| 023 | $\omega_{AR\_nom}(0) = k_{AR} \cdot V_{AR}$ | Starting value of the nominal unstressed volume of the pulmonary arteries without nervous and hormonal influences is a fraction $k_{AR}$ of $V_{AR}$. | – |

| | | | |
|---|---|---|---|
| 024 | $P_{AR} = G_{AR} \cdot (V_{AR} - \omega_{AR})$ | $P_{AR}$ – pulmonary arterial pressure, $G_{AR}$ – pulmonary arterial elasticity, $V_{AR}$ – blood volume in pulmonary arteries, $\omega_{AR}$ – unstressed volume of the pulmonary arteries. | – |
| 025 | $F_{ARVR} = \frac{Y_{ARVR}}{Vis/Vis_{norm}} \cdot (P_{AR} - P_{VR})$ | $F_{ARVR}$ – blood flow through the pulmonary microvessels, $Y_{ARVR}$ – conductivity of the pulmonary microvessels, $P_{AR}$ – pulmonary arterial pressure, $P_{VR}$ – pulmonary venous pressure, $Vis/Vis_{norm}$ – normalized blood viscosity. | – |
| 026 | $Y_{ARVR} = mass_{conductivity}(Y_{ARVR0} - A_{13} \cdot H + A_{14} \cdot DO_2, m)$ | $Y_{ARVR}$ – conductivity of the pulmonary microvessels, $Y_{ARVR0}$ – basic conductivity, $A_{13}$ – sympathetic sensitivity of the pulmonary microvessels (dependence on nervous and hormonal influences), $H$ – neurohumoral factor, $A_{14}$ – oxygen-deficient sensitivity of the pulmonary microvessels (dependence on oxygen debt $DO_2$), $m$ – body mass. | – |
| 027 | $\frac{dV_{VR}}{dt} = F_{ARVR} - F_{VRHL}$ | Change in blood volume in the pulmonary veins ($V_{VR}$) is the difference between incoming ($F_{ARVR}$) and outgoing ($F_{VRHL}$) blood flows. | – |
| 028 | $V_{VR}(0) = 0.065 \cdot V$ | Starting value of the blood volume in the pulmonary veins is 6.5% of the total blood volume $V$. | – |
| 029 | $G_{VR} = mass_{elasticity}(G_{VR0} + A_{11} \cdot H, m)$ | $G_{VR}$ – pulmonary venous elasticity, $G_{VR0}$ – basic elasticity, $A_{11}$ – venous tone, $H$ – neurohumoral factor, $m$ – body mass. | – |
| 030 | $\omega_{VR}(0) = k_{VR} \cdot V_{VR}$ | Starting value of the unstressed volume of the pulmonary veins, calculated as a fraction $k_{VR}$ of $V_{VR}$. | – |
| 031 | $P_{VR} = P_0 + LA_{PULSE} + G_{VR} \cdot (V_{VR} - \omega_{VR})$ | $P_{VR}$ – pulmonary venous pressure, $P_0$ – basic pressure, $LA_{PULSE}$ – pulse wave of the left atrium, $G_{VR}$ – pulmonary venous elasticity, $V_{VR}$ – blood volume in the pulmonary veins, $\omega_{VR}$ – unstressed volume of the pulmonary veins. | – |
| 032 | $PVR = \frac{1}{Y_{HRAR}} + \frac{1}{Y_{ARVR}} + \frac{1}{Y_{VRHL}}$ | $PVR$ – pulmonary vascular resistance, $Y_{HRAR}$ – conductivity of the pulmonary valve and pulmonary arteries, $Y_{ARVR}$ – conductivity of the pulmonary microvessels, $Y_{VRHL}$ – conductivity of the mitral valve and pulmonary veins. | – |
| **Cardiovascular module, heart (LV – left ventricle, RV – right ventricle)** | | | |
| 033 | $\frac{dV_{HL}}{dt} = F_{VRHL} - F_{HLAL}$ | Change in blood volume in the LV ($V_{HL}$) due to the difference between incoming ($F_{VRHL}$) and outgoing ($F_{HLAL}$) blood flows. | – |
| 034 | $V_{HL}(0) = 0.03 \cdot V$ | Starting value of blood volume in the LV is 3% of the total blood volume $V$. | – |
| 035 | $\omega_{HL}(0) = k_{HL} \cdot V_{HL}$ | Starting value of the unstressed LV volume is a fraction $k_{HL}$ of $V_{HL}$. | – |
| 036 | $\frac{dV_{HR}}{dt} = F_{VLHR} - F_{HRAR}$ | Change in blood volume in the RV ($V_{HR}$) due to the difference between incoming ($F_{VLHR}$) and outgoing ($F_{HRAR}$) blood flows. | – |
| 037 | $V_{HR}(0) = 0.03 \cdot V$ | Starting value of blood volume in the RV is 3% of the total blood volume $V$. | – |
| 038 | $\omega_{HR}(0) = k_{HR} \cdot V_{HR}$ | Starting value of the unstressed RV volume is a fraction $k_{HR}$ of $V_{HR}$. | – |

| | | | |
|---|---|---|---|
| 039 | $K_L = \left(K_{L0} + 0.25 \cdot nH \cdot sigm\left(20 \cdot \left(K_{L0} - 0.4\right)\right) - \right.$ $\left. -0.25 \cdot nH \cdot sigm\left(20 \cdot \left(K_{L0} - 0.7\right)\right)\right) \cdot \left(1 - B_{blocker\_in}\right)$ | $K_L$ – inotropic factor of the LV, $K_{L0}$ – inotropic status of the LV, $nH$ – sympathetic inotropic sensitivity of the myocardium, $B_{blocker\_in}$ – inotropic effect of the β-blocker bisoprolol. | The negative inotropic effect of bisoprolol (Bazroon and Alrashidi, 2022) was added. |
| 040 | $K_R = \left(K_{R0} + 0.25 \cdot nH \cdot sigm\left(20 \cdot \left(K_{R0} - 0.4\right)\right) - \right.$ $\left. -0.25 \cdot nH \cdot sigm\left(20 \cdot \left(K_{R0} - 0.7\right)\right)\right) \cdot \left(1 - B_{blocker\_in}\right)$ | $K_R$ – inotropic factor of the RV, $K_{R0}$ – inotropic status of the RV, $nH$ – sympathetic inotropic sensitivity of the myocardium, $B_{blocker\_in}$ – inotropic effect of the β-blocker bisoprolol. | The negative inotropic effect of bisoprolol (Bazroon and Alrashidi, 2022) was added. |
| 041 | If: $Cycle_{Time} \geq Cycle_{Length}$<br>Then:<br>1. $Cycle_{Length} = 1/H$<br>2. $Cycle_{Time} = 0$<br>3. $V_{HL\_KD} = V_{HL}$<br>4. $V_{HR\_KD} = V_{HR}$<br>5. $SV = K_L \cdot SV_{\max} \cdot \left[ sigm\left(0.03 \cdot \left(V_{HL} - FS_{threshold} - 80\right)\right) - \right.$ $\left. -sigm\left(0.03 \cdot \left(V_{HL} - FS_{threshold} - 260\right)\right)\right]$<br>6. $V_{HL\_KS} = V_{HL\_KD} - SV$<br>7. $V_{HR\_KS} = V_{HR} - K_R \cdot SV_{\max} \cdot \left[ sigm\left(0.03 \cdot \left(V_{HR} - \right.\right.\right.$ $\left.\left.\left. -FS_{threshold} - 80\right)\right) - sigm\left(0.03 \cdot \left(V_{HR} - FS_{threshold} - 260\right)\right)\right]$<br>8. $Systole_{Length\_L\_Exp} = \frac{0.25}{H} + 0.2 \cdot \left(1 - K_L\right)$<br>9. $Systole_{Length\_R\_Exp} = \frac{0.25}{H} + 0.2 \cdot \left(1 - K_R\right)$<br>10. $Systole_{Length\_L} = \frac{0.25}{H} + 0.2 \cdot \left(1 - K_L\right)$<br>11. $Systole_{Length\_R} = \frac{0.25}{H} + 0.2 \cdot \left(1 - K_R\right)$<br>12. $Systole_L = 1$<br>13. $Systole_R = 1$<br>14. $P_D = P_{AL}$<br>15. $P_{AR\_D} = P_{AR}$<br>16. $P_{HL\_KD} = P_{HL\_D}$<br>17. $P_{HR\_KD} = P_{HR\_D}$ | Transition "diastole – systole". A discrete event defined by an instantaneous change in the model parameters at the beginning of the cardiac cycle. The event is triggered when the current cycle time $Cycle_{Time}$ reaches the cycle length $Cycle_{Length}$.<br><br>$H$ – neurohumoral factor, $SV_{\max}$ – theoretical maximum stroke volume, $FS_{threshold}$ – the Frank-Starling law threshold, $P_{AL}$ – systemic arterial pressure, $P_D$ – diastolic blood pressure, $P_{AR}$ – pulmonary arterial pressure, $P_{AR\_D}$ – diastolic pulmonary arterial pressure.<br><br>$V_{HL}$ – current LV volume, $V_{HL\_KD}$ – LV end-diastolic volume, $SV$ – LV stroke volume, $V_{HL\_KS}$ – LV end-systolic volume, $K_L$ – inotropic factor of the LV, $Systole_{Length\_L\_Exp}$ – nominal (expected) duration of the LV systole, $Systole_{Length\_L}$ – actual duration of the LV systole, $Systole_L$ – indicator of the actual LV systole, $P_{HL\_KD}$ – LV end-diastolic pressure, $P_{HL\_D}$ – LV diastolic pressure.<br><br>$V_{HR}$ – current RV volume, $V_{HR\_KD}$ – RV end-diastolic volume, $V_{HR\_KS}$ – RV end-systolic volume, $K_R$ – inotropic factor of the RV, $Systole_{Length\_R\_Exp}$ – nominal (expected) duration of the RV systole, $Systole_{Length\_R}$ – actual duration of the RV systole, $Systole_R$ – indicator of the actual RV systole, $P_{HR\_KD}$ – RV end-diastolic pressure, $P_{HR\_D}$ – RV diastolic pressure. | Equations fixing the values of $P_{HL\_KD}$ and $P_{HR\_KD}$ were added. |

| | | | |
|---|---|---|---|
| 042 | $FS_{threshold}(0) = mass_{volume}(FS_{threshold0}, m)$ | Starting value of the Frank-Starling law threshold, $FS_{threshold0}$ – the average normal value of the threshold, $m$ – body mass. | – |
| 043 | $SV_{\max}(0) = mass_{volume}(SV_{\max 0}, m)$ | Starting value of the theoretical maximum stroke volume, $SV_{\max 0}$ – the average value of $SV_{\max}$, $m$ – body mass. | – |
| 044 | If: $Systole = 1$<br>Then:<br>1. $F_{HLAL\_p} = F_{HLAL}$<br>2. $F_{HRAR\_p} = F_{HRAR}$ | A discrete event is triggered after the transition "diastole – systole" (*Systole* = 1). At this point, blood flow through the aortic valve $F_{HLAL}$ reaches its peak $F_{HLAL\_p}$, and blood flow through the pulmonary valve $F_{HRAR}$ reaches its peak $F_{HRAR\_p}$. | – |
| 045 | If: $V_{HL} < V_{HL\_KS}$ and $Systole_L = 1$<br>Then:<br>1. $Systole_L = 0$<br>2. $Systole_{Length\_L} = Cycle_{Time}$<br>3. $P_S = P_{AL}$ | Transition "LV systole – LV diastole". A discrete event is triggered if LV volume $V_{HL}$ reaches LV end-systolic volume $V_{HL\_KS}$ and the LV is in systole. At this point, the LV switches to diastole ($Systole_L = 0$), and the actual duration of the LV systole $Systole_{Length\_L}$ and systolic blood pressure $P_S$ are determined as the current values of cycle time $Cycle_{Time}$ and systemic arterial pressure $P_{AL}$, respectively. | – |
| 046 | If: $V_{HR} < V_{HR\_KS}$ and $Systole_R = 1$<br>Then:<br>1. $Systole_R = 0$<br>2. $Systole_{Length\_R} = Cycle_{Time}$<br>3. $P_{AR\_S} = P_{AR}$ | Transition "RV systole – RV diastole". A discrete event is triggered if RV volume $V_{HR}$ reaches RV end-systolic volume $V_{HR\_KS}$ and the RV is in systole. At this point, the RV switches to diastole ($Systole_R = 0$), and the actual duration of the RV systole $Systole_{Length\_R}$ and systolic pulmonary arterial pressure $P_{AR\_S}$ are determined as the current values of cycle time $Cycle_{Time}$ and pulmonary arterial pressure $P_{AR}$, respectively. | – |
| 047 | If: $P_{VR} > P_{HL}$<br>Then: $F_{VRHL\_ep} = F_{VRHL}$ | Opening of the mitral valve. A discrete event that occurs as the LV enters diastole when pulmonary venous pressure $P_{VR}$ becomes greater than LV pressure $P_{HL}$. In healthy people, the LV filling rate $F_{VRHL}$ reaches its maximum value (early peak) $F_{VRHL\_ep}$ immediately after the valve opens. | – |
| 048 | If: $P_{VL} > P_{HR}$<br>Then: $F_{VLHR\_ep} = F_{VLHR}$ | Opening of the tricuspid valve. A discrete event that occurs as the RV enters diastole when pressure in the systemic veins $P_{VL}$ becomes greater than pressure in the RV $P_{HR}$. $F_{VLHR\_ep}$ – early peak of the RV filling. | – |
| 049 | $P_{HL\_D} = mass_{elasticity}(G_{HL}, m) \cdot (V_{HL} - \omega_{HL})$ | $P_{HL\_D}$ – LV diastolic pressure, $V_{HL}$ – LV volume, $\omega_{HL}$ – unstressed LV volume, $G_{HL}$ – LV wall elasticity, $m$ – body mass. | – |
| 050 | $P_{HR\_D} = mass_{elasticity}(G_{HR}, m) \cdot (V_{HR} - \omega_{HR})$ | $P_{HR\_D}$ – RV diastolic pressure, $V_{HR}$ – RV volume, $\omega_{HR}$ – unstressed RV volume, $G_{HR}$ – RV wall elasticity, $m$ – body mass. | – |
| 051 | $Systole_{L\_Exp} = \begin{cases} 0, & Cycle_{Time} \geq Systole_{Length\_L\_Exp} \\ 1, & otherwise \end{cases}$ | Nominal LV systole indicator. | – |

| | | | |
|---|---|---|---|
| 052 | $Systole_{R\_Exp} = \begin{cases} 0, & Cycle_{Time} \ge Systole_{Length\_R\_Exp} \\ 1, & otherwise \end{cases}$ | Nominal RV systole indicator. | – |
| 053 | $Systole = \begin{cases} 0, & Systole_L = 0 \text{ and } Systole_R = 0 \\ 1, & otherwise \end{cases}$ | *Systole* is an indicator of the total actual systole. It is equal to 0 if the actual LV and RV systole indicators are simultaneously equal to 0, and 1, otherwise. | – |
| 054 | $DTS_L = Systole_L - Systole_{L\_Exp}$ | LV systolic mismatch. | – |
| 055 | $DTS_R = Systole_R - Systole_{R\_Exp}$ | RV systolic mismatch. | – |
| 056 | $\frac{dP_{HL\_S}}{dt} = A_5 \cdot DTS_L$ | $P_{HL\_S}$ – LV systolic pressure, $DTS_L$ – LV systolic mismatch, $A_5$ – sensitivity factor. | – |
| 057 | $\frac{dP_{HR\_S}}{dt} = A_{15} \cdot DTS_R$ | $P_{HR\_S}$ – RV systolic pressure, $DTS_R$ – RV systolic mismatch, $A_{15}$ – sensitivity factor. | – |
| 058 | $P_{HL} = (Systole - Systole_L) \cdot P_{AL} + (1 - Systole) \cdot P_{HL\_D} + P_{HL\_S} \cdot Systole_L$ | LV pressure $P_{HL}$ is equal to LV systolic pressure $P_{HL\_S}$ if the LV is in systole; systemic arterial pressure $P_{AL}$ if the LV is in diastole, and the RV is in systole; and LV diastolic pressure $P_{HL\_D}$ if both the LV and RV are in diastole. | – |
| 059 | $P_{HR} = Systole_R \cdot P_{HR\_S} + P_{AR} \cdot (Systole - Systole_R) + (1 - Systole) \cdot P_{HR\_D}$ | RV pressure $P_{HR}$ is equal to RV systolic pressure $P_{HR\_S}$ if the RV is in systole; pulmonary arterial pressure $P_{AR}$ if the RV is in diastole, and the LV is in systole; and RV diastolic pressure $P_{HR\_D}$ if both the RV and LV are in diastole. | – |
| 060 | $Y_{VRHL} = mass_{conductivity}(Y_{VRHL0} + A_{16} \cdot P_{VR}, m)$ | $Y_{VRHL}$ – conductivity of the mitral valve and pulmonary veins, $Y_{VRHL0}$ – basic conductivity, $P_{VR}$ – pressure in the pulmonary vein and left atrium. $A_{16}$ – constant, *m* – body mass. | – |
| 061 | $Y_{VLHR} = mass_{conductivity}(Y_{VLHR0} + P_{VL} \cdot A_6 + RO_{20} \cdot A_7 - P_{HR\_S} \cdot A_{10}, m)$ | $Y_{VLHR}$ – conductivity of the tricuspid valve and systemic veins, $Y_{VLHR0}$ – basic conductivity, $P_{VL}$ – pressure in the inferior vena cava and right atrium, $RO_{20}$ – the average normal value of oxygen demand, $P_{HR\_S}$ – systolic pressure in the RV. $A_6$, $A_7$, $A_{10}$ – constants. *m* – body mass. | – |
| 062 | $F_{VRHL} = valve(P_{VR}, P_{HL}, K_{VRHL}, Y_{VRHL})$ | Blood flow through the mitral valve $F_{VRHL}$ is described by the difference between pressure in the pulmonary vein and left atrium $P_{VR}$ and LV pressure $P_{HL}$. $Y_{VRHL}$ – conductivity of the mitral valve and pulmonary veins. $K_{VRHL}$ – regurgitation coefficient. | – |
| 063 | $F_{HLAL} = valve(P_{HL}, P_{AL}, K_{HLAL}, mass_{conductivity}(Y_{HLAL}, m))$ | Blood flow through the aortic valve $F_{HLAL}$ is characterized by the difference between LV pressure $P_{HL}$ and systemic arterial pressure $P_{AL}$. $Y_{HLAL}$ – conductivity of the aortic valve and systemic arteries, allometrically scaled by body mass *m*. $K_{HLAL}$ – regurgitation coefficient. | – |

| | | | |
|---|---|---|---|
| 064 | $F_{VLHR} = valve\left(P_{VL}, P_{HR}, K_{VLHR}, Y_{VLHR}\right)$ | Blood flow through the tricuspid valve $F_{VLHR}$ is described by the difference between pressure in the inferior vena cava and right atrium $P_{VL}$ and RV pressure $P_{HR}$. $Y_{VLHR}$ – conductivity of the tricuspid valve and systemic veins. $K_{VLHR}$ – regurgitation coefficient. | – |
| 065 | $F_{HRAR} = valve\left(P_{HR}, P_{AR}, K_{HRAR},\ mass_{conductivity}\left(Y_{HRAR}, m\right)\right)$ | Blood flow through the pulmonary valve $F_{HRAR}$ is characterized by the difference between RV pressure $P_{HR}$ and pulmonary arterial pressure $P_{AR}$. $Y_{HRAR}$ – conductivity of the pulmonary valve and pulmonary arteries, allometrically scaled by body mass *m*. $K_{HRAR}$ – regurgitation coefficient. | – |
| 066 | $LA_{PULSE} = atrium_{pulse}\left(Cycle_{Length}, Systole_{Length\_L}, P_{AL}, Cycle_{Time}\right)$ | Left atrium pulse wave depends on the cardiac cycle length $Cycle_{Length}$, the actual duration of the LV systole $Systole_{Length\_L}$, systemic arterial pressure $P_{AL}$, and the current cycle time $Cycle_{Time}$. | – |
| 067 | $RA_{PULSE} = atrium_{pulse}\left(Cycle_{Length}, Systole_{Length\_R}, P_{AL}, Cycle_{Time}\right)$ | Right atrium pulse wave depends on the cardiac cycle length $Cycle_{Length}$, the actual duration of the RV systole $Systole_{Length\_R}$, pulmonary arterial pressure $P_{AL}$ and the current cycle time $Cycle_{Time}$. | – |
| 068 | If: $Cycle_{Time} > Cycle_{Length} - 0.2 \cdot \left(Cycle_{Length} - Systole_{Length\_L}\right)$<br>Then: $F_{VRHL\_ap} = F_{VRHL}$ | A discrete event that determines the active peak $F_{VRHL\_ap}$ of blood flow through the mitral valve $F_{VRHL}$. This peak is reached by contraction of the left atrium at the time when $LA_{PULSE}$ becomes positive. | – |
| 069 | If: $Cycle_{Time} > Cycle_{Length} - 0.2 \cdot \left(Cycle_{Length} - Systole_{Length\_R}\right)$<br>Then: $F_{VLHR\_ap} = F_{VLHR}$ | A discrete event that defines the active peak $F_{VLHR\_ap}$ of blood flow through the tricuspid valve $F_{VLHR}$. This peak is reached by contraction of the right atrium at the time when $RA_{PULSE}$ becomes positive. | – |
| 070 | $V_{HL\_KS}(0) = V_{HL\_KD} - K_L \cdot SV_{\max} \cdot \left[sigm\left(0.03 \cdot \left(V_{HL\_KD} -\right.\right.\right.$<br>$\left.\left.\left.- FS_{threshold} - 80\right)\right) - sigm\left(0.03 \cdot \left(V_{HL\_KD} - FS_{threshold} - 260\right)\right)\right];$<br>$Systole_{Length\_L\_Exp}(0) = 0.25 \cdot Cycle_{Length} + 0.2 \cdot \left(1 - K_L\right);$<br>$Systole_{Length\_L}(0) = 0.25 \cdot Cycle_{Length} + 0.2 \cdot \left(1 - K_L\right);$<br>$V_{HR\_KS}(0) = V_{HR\_KD} - K_R \cdot SV_{\max} \cdot \left[sigm\left(0.03 \cdot \left(V_{HR\_KD} -\right.\right.\right.$<br>$\left.\left.\left.- FS_{threshold} - 80\right)\right) - sigm\left(0.03 \cdot \left(V_{HR\_KD} - FS_{threshold} - 260\right)\right)\right];$<br>$Systole_{Length\_R\_Exp}(0) = 0.25 \cdot Cycle_{Length} + 0.2 \cdot \left(1 - K_R\right);$<br>$Systole_{Length\_R}(0) = 0.25 \cdot Cycle_{Length} + 0.2 \cdot \left(1 - K_R\right).$ | Initialization of starting values. See Equations 041 for explanation. | – |
| 071 | $\frac{dCycle_{Time}}{dt} = 1$ | Linking the current time of the cardiac cycle $Cycle_{Time}$ to the model time. | – |

| | | | |
|---|---|---|---|
| 072 | $Heart_{Rate}=\dfrac{60}{Cycle_{Length}}$ | The duration of the cardiac cycle $Cycle_{Length}$, by definition, is the time of one heartbeat. Heart rate $Heart_{Rate}$ is the number of heart beats per minute. | – |
| 073 | $CO=\dfrac{SV\cdot Heart_{Rate}}{1000}$ | Cardiac output $CO$ (L/min) is equal to stroke volume $SV$ (mL) times heart rate $Heart_{Rate}$ (bpm). | – |
| 074 | $EF=\dfrac{SV}{V_{HL\_KD}}\cdot 100$ | Ejection fraction $EF$ is stroke volume $SV$ divided by the LV end-diastolic volume $V_{HL\_KD}$ and multiplied by 100%. | – |
| **Cardiovascular module, tissue metabolism** | | | |
| 075 | $gO_2=F_{ALVL}\cdot\min\left(AO_2-VO_2,1\right)$ | Oxygen consumption $gO_2$ in tissues is equal to the product of blood flow through the tissues $F_{ALVL}$ and the arteriovenous oxygen difference $AO_2-VO_2$. | – |
| 076 | $AO_2=\dfrac{He\cdot C_H\cdot SpO_2}{1000}$ | Arterial oxygen content $AO_2$ is calculated as the product of the total amount of hemoglobin $He$ (g/L), the oxygen capacity of hemoglobin $C_H$ (mg/mL), and the arterial oxygen saturation $SpO_2$. | – |
| 077 | $\dfrac{dDO_2}{dt}=\mathrm{sgn}\left(DO_2,A_2\cdot\left(RO_2-gO_2\right)\right)$ | The rate of change in oxygen debt $DO_2$ is proportional to the difference between oxygen demand $RO_2$ and oxygen consumption $gO_2$. $A_2$ is the functional status of the body. | – |
| 078 | $\dfrac{dVO_2}{dt}=A_1\cdot\left(gO_2-RO_2\right)$ | The rate of change in venous oxygen content $VO_2$ is proportional to the difference between oxygen consumption $gO_2$ and oxygen demand $RO_2$. $A_1$ – total metabolic intensity. | – |
| 079 | $RO_2\left(0\right)=mass_{volume}\left(RO_{20},m\right)$ | Starting value of the oxygen demand, $RO_{20}$ – the average normal value of $RO_2$, $m$ – body mass. | – |
| **Cardiovascular module, neurohumoral control** | | | |
| 080 | $nB=\min\left(r_{minus}\left(0.07,30,P_{AL},50\cdot\psi_{AT1\_Baro}\right),1\right)$ | $nB$ – baroreceptor activity, $P_{AL}$ – systemic arterial pressure. $P_0=50$ mmHg is the baroreceptor sensitivity threshold, i.e., the lower bound for $P_{AL}$ at which $nB$ = 0. $\psi_{AT1\_Baro}$ determines the effect of AT1-bound angiotensin II on blood pressure without affecting heart rate (by changing the $P_0$ threshold). | – |
| 081 | $\psi_{AT1\_Baro}=sl_{baro}\cdot\dfrac{AT1\_ANGII}{AT1\_ANGII_{norm}}+1-sl_{baro}$ | Linear function with a slope $0<sl_{baro}<1$, which defines the influence of the normalized level of AT1-bound angiotensin II $\left(AT1\_ANGII/AT1\_ANGII_{norm}\right)$ on the baroreceptor sensitivity threshold. | – |
| 082 | $nS=\min\left(r_{plus}\left(10,50,st\cdot\psi_{AT1\_Stress},0\right),1\right)$ | $nS$ – stress receptor activity. Parameter $st$ is a stress factor that describes the level of steroid hormones in the blood (adrenaline, norepinephrine) and taking nominal values from 0 (absolute rest) to 1 (absolute stress). $\psi_{AT1\_Stress}$ determines the effect of AT1-bound angiotensin II on the norepinephrine release from the atria. | – |

| | | | |
|---|---|---|---|
| 083 | $\Psi_{AT1\_Stress} = sl_{stress} \cdot \frac{AT1\_ANGII}{AT1\_ANGII_{norm}} + 1 - sl_{stress}$ | Linear function with a slope $0 < sl_{stress} < 1$, which defines the influence of the normalized level of AT1-bound angiotensin II $(AT1\_ANGII / AT1\_ANGII_{norm})$ on the stress factor. | – |
| 084 | $nV = \min\left(r_{minus}\left(30,30,VO_2,0\right),1\right)$ | $nV$ – respiratory receptor activity, $VO_2$ – venous oxygen content. | – |
| 085 | $nD = \min\left(r_{plus}\left(0.1,10,DO_2,0\right),1\right)$ | $nD$ – fatigue receptor activity, $DO_2$ – oxygen debt. | – |
| 086 | $nSum = Heart_{Stress} \cdot nS \cdot \left(1 - B_{blocker}\right) \cdot \left(1 - Diuretic_{stress}\right) + $ $+ Heart_{Baro} \cdot nB + Heart_{Oxygen} \cdot nD + Heart_{VO2} \cdot nV + Heart_{Base}$ | The activity of the cardiac center (*nSum*) is the sum of the activities of stress (*nS*), fatigue (*nD*), and respiratory receptors (*nV*), as well as baroreceptors (*nB*). $Heart_{Stress}$ – stress sensitivity, $Heart_{Baro}$ – baroreceptor sensitivity, $Heart_{Oxygen}$ – sensitivity to fatigue (a lower value corresponds to the greater exercise tolerance), $Heart_{VO2}$ – respiratory sensitivity, $Heart_{Base}$ – basic activity of the cardiac center, $B_{blocker}$ – negative chronotropic effect of the β-blocker bisoprolol, $Diuretic_{stress}$ – effect of the thiazide diuretic hydrochlorothiazide (HCTZ) on the pressor response of norepinephrine. | The effects of bisoprolol and HCTZ on stress receptor activity were taken from the model extension (Kutumova et al., 2022). |
| 087 | $nH = \begin{cases} 0, & H < 0 \\ \frac{1 - \exp(-3H)}{1 + 100 \cdot \exp(-3H)}, & otherwise \end{cases}$ | Sigmoid function of sympathetic inotropic sensitivity of the myocardium depending on the neurohumoral factor *H*. | – |
| 088 | $\frac{dH}{dt} = A_{12} \cdot \left(3.2 \cdot \min\left(r_{plus}\left(2,18,nSum,0\right),1\right) - H\right)$ | Calculation of the neurohumoral factor *H*, considering the reactivity of the cardiac center $A_{12}$, which characterizes its adaptive ability. | – |
| **Renal module, nervous system** | | | |
| 089 | $RSNA = N_{rsna} \cdot \alpha_{map} \cdot \alpha_{rap}$ | *RSNA* – renal sympathetic nerve activity, $N_{rsna}$ – normalized *RSNA* value. $\alpha_{map}$ and $\alpha_{rap}$ represent the effects of mean arterial pressure and right atrial pressure on *RSNA*, respectively. | – |
| 090 | $\alpha_{map} = 0.5 + 1.1 \cdot \left(1 + \exp\left(\frac{MAP - 100}{15}\right)\right)^{-1}$ | Effect of mean arterial pressure *MAP* on *RSNA*. | – |
| 091 | $\alpha_{rap} = 1 - 0.008 \cdot P_{ra}$ | Effect of right atrial pressure $P_{ra}$ on *RSNA*. | – |
| 092 | $P_{ra} = 0.2787 \cdot \exp\left(0.2281 \cdot CO\right)$ | Normalized right atrial pressure $P_{ra}$ is calculated using the Frank-Starling law as a function of cardiac output *CO*. | – |
| **Renal module, sodium transport and reabsorption along the nephron** | | | |
| 093 | $\Phi_{filsod} = GFR \cdot C_{sod}$ | $\Phi_{filsod}$ – amount of sodium filtered from the glomerulus to the proximal tubule per minute (filtered sodium load), *GFR* – glomerular filtration rate, $C_{sod}$ – serum sodium concentration. | – |
| 094 | $\eta_{pt\_sodreab} = \min\left(1, n_{\eta\_pt} \cdot \gamma_{filsod} \cdot \gamma_{at} \cdot \gamma_{rsna}\right)$ | Fractional proximal sodium reabsorption is defined as the normalized value $n_{\eta\_pt}$ multiplied by the influence functions described below. Cannot exceed 100%. | – |

| | | | |
|---|---|---|---|
| 095 | $\gamma_{at} = 0.95 + \frac{0.12}{1+\exp\left(2.6 - 1.8 \cdot \log AT1\_ANGII\right)}$ | An increase in the level of AT1-bound angiotensin II (*AT1_ANGII*) leads to an increase in fractional proximal sodium reabsorption. | – |
| 096 | $\gamma_{filsod} = 0.8 + \frac{0.3}{1+138^{-1} \cdot \exp\left(\Phi_{filsod} - 14\right)}$ | An increase in the filtered sodium load $\Phi_{filsod}$ leads to a decrease in fractional proximal sodium reabsorption. | – |
| 097 | $\gamma_{rsna} = 0.5 + \frac{0.7}{1+2.18^{-1} \cdot \exp\left(1 - RSNA\right)}$ | Increased renal sympathetic nerve activity *RSNA* results in increased fractional proximal sodium reabsorption. | – |
| 098 | $\Phi_{pt\_sodreab} = \Phi_{filsod} \cdot \eta_{pt\_sodreab}$ | $\Phi_{pt\_sodreab}$ – absolute proximal sodium reabsorption rate, $\Phi_{filsod}$ – filtered sodium load, $\eta_{pt\_sodreab}$ – fractional proximal sodium reabsorption. | – |
| 099 | $\Phi_{md\_sod} = \Phi_{filsod} - \Phi_{pt\_sodreab}$ | $\Phi_{md\_sod}$ – macula densa sodium flow rate, $\Phi_{filsod}$ – filtered sodium load, $\Phi_{pt\_sodreab}$ – absolute proximal sodium reabsorption rate. | – |
| 100 | $\eta_{dt\_sodreab} = \min\left(1, n_{\varepsilon\_dt} \cdot \psi_{al} \cdot \left(1 - Diuretic_{Inhibition}\right)\right)$ | Fractional distal sodium reabsorption is defined as the normalized value $n_{\varepsilon\_dt}$ multiplied by the aldosterone influence function $\psi_{al}$. Cannot exceed 100%. $Diuretic_{Inhibition}$ – inhibition of distal tubular sodium reabsorption by the thiazide diuretic hydrochlorothiazide (HCTZ). | The effect of HCTZ on fractional distal sodium reabsorption was taken from the model extension (Kutumova et al., 2022). |
| 101 | $\Psi_{al} = 0.17 + 0.94 \cdot \left(1 + \exp\left(\frac{0.48 - 1.2 \cdot \log C_{al}}{0.88}\right)\right)^{-1}$ | Effect of aldosterone concentration $C_{al}$ on fractional distal sodium reabsorption. | – |
| 102 | $\Phi_{dt\_sodreab} = \Phi_{md\_sod} \cdot \eta_{dt\_sodreab}$ | $\Phi_{dt\_sodreab}$ – absolute distal sodium reabsorption rate, $\Phi_{md\_sod}$ – macula densa sodium flow rate, $\eta_{dt\_sodreab}$ – fractional distal sodium reabsorption. | – |
| 103 | $\Phi_{dt\_sod} = \Phi_{md\_sod} - \Phi_{dt\_sodreab}$ | $\Phi_{dt\_sod}$ – rate of the distal sodium outflow, $\Phi_{md\_sod}$ – macula densa sodium flow rate, $\Phi_{dt\_sodreab}$ – absolute distal sodium reabsorption rate. | – |
| 104 | $\eta_{cd\_sodreab} = \min\left(1, n_{\eta\_cd} \cdot \lambda_{dt} \cdot \lambda_{anp}\right)$ | Fractional collecting duct sodium reabsorption is defined as the normalized value $n_{\eta\_cd}$ multiplied by the influence functions described below. Cannot exceed 100%. | – |
| 105 | $\lambda_{dt} = 0.82 + \frac{0.39}{1+\exp\left(0.5 \cdot \left(\Phi_{dt\_sod} - 1.6\right)\right)}$ | Effect of distal sodium outflow rate $\Phi_{dt\_sod}$ on fractional collecting duct sodium reabsorption. | – |
| 106 | $\lambda_{anp} = -\ 0.1 \cdot \frac{C_{anp}}{C_{anp\_norm}} + 1.1199$ | Effect of normalized natriuretic peptide concentration $C_{anp}/C_{anp\_norm}$ on fractional collecting duct sodium reabsorption. | – |
| 107 | $\Phi_{cd\_sodreab} = \Phi_{dt\_sod} \cdot \eta_{cd\_sodreab}$ | $\Phi_{cd\_sodreab}$ – absolute collecting duct sodium reabsorption rate, $\Phi_{dt\_sod}$ – rate of the distal sodium outflow, $\eta_{cd\_sodreab}$ – fractional collecting duct sodium reabsorption. | – |

| | | | |
|---|---|---|---|
| 108 | $\Phi_{u\_sod} = \Phi_{dt\_sod} - \Phi_{cd\_sodreab}$ | $\Phi_{u\_sod}$ – urine sodium excretion rate, $\Phi_{dt\_sod}$ – rate of the distal sodium outflow, $\Phi_{cd\_sodreab}$ – absolute collecting duct sodium reabsorption rate. | – |
| 109 | $\frac{dM_{sod}}{dt} = \Phi_{sodin} - \Phi_{u\_sod}$ | The rate of formation of total exchangeable sodium ( $M_{sod}$ ) is determined by the difference between the rate of sodium intake $\Phi_{sodin}$ and the rate of sodium excretion $\Phi_{u\_sod}$. | – |
| 110 | $C_{sod} = 1.03 \cdot \frac{(M_{sod} - 410)}{TBW} - 0.29 \cdot glucose + 78.44$ | Serum sodium concentration $C_{sod}$ as a function of total exchangeable sodium $M_{sod}$, total body water *TBW*, and plasma glucose (*glucose*). | – |
| **Renal module, renin-angiotensin-aldosterone system** | | | |
| 111 | $R_{sec} = \frac{\ln 2}{h_{renin}} \cdot PRC_{nom} \cdot \nu_{MD\_sod} \cdot \nu_{RSNA} \cdot \nu_{AT1\_ANGII} \cdot \cdot (1 + Diuretic_{Stimulation}) \cdot (1 - B_{blocker\_rs})$ | Renin secretion rate $R_{sec}$ is defined by the nominal value of plasma renin concentration (PRC) $PRC_{nom}$ multiplied by the influence functions described below. $h_{renin}$ – half-life of PRC, $Diuretic_{Stimulation}$ – stimulation of renin secretion by the thiazide diuretic hydrochlorothiazide (HCTZ), $B_{blocker\_rs}$ – suppression of renin secretion by the β-blocker bisoprolol. | The effects of HCTZ and bisoprolol on renin secretion rate were taken from the model extension (Kutumova et al., 2022). |
| 112 | $\nu_{MD\_sod} = exp(-\tau_{MD\_renin} \cdot (\Phi_{md\_sod} - \Phi_{md\_sod\_0}))$ | Effect of macula densa sodium flow rate $\Phi_{md\_sod}$ on the rate of renin secretion. $\Phi_{md\_sod\_0}$ – nominal value of $\Phi_{md\_sod}$, $\tau_{MD\_renin}$ – constant. | – |
| 113 | $\nu_{RSNA} = 1.89 - \frac{2.056}{1.358 + \exp(RSNA - 0.8667)}$ | Effect of renal sympathetic nerve activity *RSNA* on the rate of renin secretion. | – |
| 114 | $\nu_{AT1\_ANGII} = \left(\frac{AT1\_ANGII_{norm}}{AT1\_ANGII}\right)^{slope_{AT1\_PRC}}$ | Effect of the normalized value of AT1-bound angiotensin II ( $AT1\_ANGII_{norm} / AT1\_ANGII$ ) on the rate of renin secretion. $slope_{AT1\_PRC}$ – constant. | – |
| 115 | $\frac{dPRC}{dt} = R_{sec} - \frac{\ln 2}{h_{renin}} \cdot PRC$ | Plasma renin concentration (*PRC*) is described by the rate of renin secretion $R_{sec}$ and the rate of renin clearance, with a half-life of $h_{renin}$. | – |
| 116 | $PRC(0) = PRC_{nom}$ | Starting value of *PRC* value is the nominal value $PRC_{nom}$. | – |
| 117 | $PRA = PRC \cdot X_{PRC\_PRA} \cdot (1 - DRI)$ | *PRA* – plasma renin activity, *PRC* – plasma renin concentration, $X_{PRC\_PRA}$ – equilibrium ratio of *PRA* to *PRC*, *DRI* – direct renin inhibition by aliskiren. | The effect of aliskiren on *PRC* was taken from the model extension (Kutumova et al., 2022). |
| 118 | $\frac{dANGI}{dt} = PRA - (c_{ACE} \cdot (1 - ACEi) + c_{chym} + c_{nep}) \cdot ANGI - - \frac{\ln 2}{h_{ANGI}} \cdot ANGI$ | *ANGI* – plasma angiotensin I, *PRA* – plasma renin activity, $c_{ACE}$ and $c_{chym}$ – rates of conversion of *ANGI* to angiotensin II by ACE and chymase, $c_{nep}$ – rate of conversion of *ANTI* to angiotensin-(1-7) by neprilisin, $h_{ANGI}$ – half-life of *ANGI*, *ACEi* – inhibition of angiotensin-converting enzyme by enalapril. | The effect of enalapril on $c_{ACE}$ was taken from the model extension (Kutumova et al., 2022). |

| | | | |
|---|---|---|---|
| 119 | $$\frac{dANGII}{dt} = \left(c_{ACE}\cdot(1-ACEi)+c_{chym}\right)\cdot ANGI - \left(c_{ACE2}+c_{ANGII\_ANGIV}+c_{AT1}\cdot(1-ARB)+c_{AT2}\right)\cdot ANGII - \frac{\ln 2}{h_{ANGII}}\cdot ANGII$$ | *ANGII* – plasma angiotensin II, *ANGI* – plasma angiotensin I, $c_{ACE}$ and $c_{chym}$ – rates of conversion of *ANGI* to *ANGII* by ACE and chymase, $c_{ACE2}$ – rate of conversion of *ANGII* to angiotensin-(1-7) by ACE2, $c_{ANGII\_ANGIV}$ – rate of conversion of *ANGII* to angiotensin IV, $c_{AT1}$ and $c_{AT2}$ – rates of *ANGII* binding to AT1 and AT2 receptors, $h_{ANGII}$ – half-life of *ANGII*, $ACEi$ – inhibition of angiotensin-converting enzyme by enalapril, *ARB* – blocking angiotensin receptors with losartan. | The effects of enalapril on $c_{ACE}$ and losartan on $c_{AT1}$ were taken from the model extension (Kutumova et al., 2022). |
| 120 | $$\frac{dANG17}{dt} = c_{nep}\cdot ANGI + c_{ACE2}\cdot ANGII - \frac{\ln 2}{h_{ANG17}}\cdot ANG17$$ | *ANG*17 – plasma angiotensin-(1-7), *ANGI* – plasma angiotensin I, *ANGII* – plasma angiotensin II, $c_{nep}$ – rate of conversion of *ANTI* to *ANG*17 by neprilisin, $c_{ACE2}$ – rate of conversion of *ANGII* to *ANG*17 by ACE2, $h_{ANG17}$ – half-life of *ANG*17. | – |
| 121 | $$\frac{dANGIV}{dt} = c_{ANGII\_ANGIV}\cdot ANGII - \frac{\ln 2}{h_{ANGIV}}\cdot ANGIV$$ | *ANGIV* – plasma angiotensin IV, *ANGII* – plasma angiotensin II, $c_{ANGII\_ANGIV}$ – rate of conversion of *ANGII* to *ANGIV*, $h_{ANGIV}$ – half-life of *ANGIV*. | – |
| 122 | $$\frac{dAT1\_ANGII}{dt} = c_{AT1}\cdot(1-ARB)\cdot ANGII - \frac{\ln 2}{h_{AT1}}\cdot AT1\_ANGII$$ | *AT1_ANGII* – concentration of AT1-bound angiotensin II, *ANGII* – plasma angiotensin II, $c_{AT1}$ – rate of *ANGII* binding to AT1 receptors, $h_{AT1}$ – half-life of *AT1_ANGII*, *ARB* – blocking angiotensin receptors with losartan. | The effect of losartan on $c_{AT1}$ was taken from the model extension (Kutumova et al., 2022) |
| 123 | $$\frac{dAT2\_ANGII}{dt} = c_{AT2}\cdot ANGII - \frac{\ln 2}{h_{AT2}}\cdot AT2\_ANGII$$ | *AT2_ANGII* – concentration of AT2-bound angiotensin II, *ANGII* – plasma angiotensin II, $c_{AT2}$ – rate of *ANGII* binding to AT2 receptors, $h_{AT2}$ – half-life of *AT2_ANGII*. | – |
| 124 | $N_{als} = \xi_{k\_sod}\cdot\xi_{map}\cdot\xi_{at}$ | The normalized aldosterone secretion rate $N_{als}$ is calculated as the product of the influence functions described below. $\xi_{map}$ – effect of mean arterial pressure *MAP*. $\xi_{map} = 1$ (no effect) if *MAP* is normal or above normal. | – |
| 125 | $\xi_{k\_sod} = 2^{power}, \quad power = C_K\cdot\left(1-Diuretic_{potassium}\right)-4.5$ | Effect of potassium concentration ($C_K$) on the rate of aldosterone secretion. $Diuretic_{potassium}$ – reducing the level of potassium in the blood with the thiazide diuretic hydrochlorothiazide (HCTZ). | The effect of HCTZ on potassium was taken from the model extension (Kutumova et al., 2022). |
| 126 | $$\xi_{at} = 0.4 + 2.4\cdot\left(1+\exp\left(2.82-1.5\cdot\frac{\log AT1\_ANGII}{0.8}\right)\right)^{-1}$$ | Effect of AT1-bound angiotensin II (*AT1_ANGII*) on the rate of aldosterone secretion. | – |
| 127 | $$\frac{dN_{al}}{dt} = \frac{N_{als}-N_{al}}{T_{al}}$$ | $N_{al}$ – normalized aldosterone concentration, $N_{als}$ – normalized aldosterone secretion rate, $T_{al}$ – time constant. | – |
| 128 | $C_{al} = C_{al\_norm}\cdot N_{al}$ | Plasma aldosterone $C_{al}$ is equal to the product of the normalized aldosterone concentration $N_{al}$ and its normal value $C_{al\_norm}$. | – |

| | | | |
|---|---|---|---|
| **Renal module, hormonal system** | | | |
| 129 | $C_{anp} = C_{anp\_norm} \cdot \left(7.427 - \dfrac{6.554}{1+\exp\left(P_{ra} - 3.762\right)}\right)$ | Natriuretic peptide concentration $C_{anp}$ depending on pressure in the right atrium $P_{ra}$ and the normal hormone level $C_{anp\_norm}$. | – |
| 130 | $osmolality = 1.86 \cdot C_{sod} + glucose + urea \cdot \left(1 + Diuretic_{urea}\right) + 9$ | $osmolality$ – serum osmolality, $C_{sod}$ – serum sodium concentration, $glucose$ – plasma glucose, $urea$ – plasma urea, $Diuretic_{urea}$ – increasing the level of urea in the blood with the thiazide diuretic hydrochlorothiazide (HCTZ). | The effect of HCTZ on urea was taken from the model extension (Kutumova et al., 2022). |
| 131 | $C_{adh} = \max\left(0, 0.23 \cdot \left(osmolality - 271\right)\right)$ | Concentration of antidiuretic hormone $C_{adh}$ depending on serum osmolality. | – |
| **Renal module, diuresis** | | | |
| 132 | $\Phi_{t\_wreab} = \eta_{pt\_sodreab} \cdot GFR + \left(1 - \eta_{pt\_sodreab}\right) \cdot GFR \cdot \mu_{adh}$ | $\Phi_{t\_wreab}$ – tubular water reabsorption rate, $GFR$ – glomerular filtration rate, $\eta_{pt\_sodreab}$ – fractional proximal sodium reabsorption, $\mu_{adh}$ – effect of plasma antidiuretic hormone concentration. | – |
| 133 | $\mu_{adh} = \begin{cases} 0.0, & C_{adh} \le 0.765 \\ 0.383 \cdot C_{adh} - 0.293, & 0.765 < C_{adh} \le 3 \\ -0.0383 \cdot C_{adh}^2 + 0.364 \cdot C_{adh} + 0.109, & 3 < C_{adh} \le 5 \\ 0.0012 \cdot C_{adh} + 0.9653, & C_{adh} > 5 \end{cases}$ | Effect of plasma antidiuretic hormone concentration $C_{adh}$ on the rate of tubular water reabsorption. | – |
| 134 | $\Phi_u = GFR - \Phi_{t\_wreab}$ | $\Phi_u$ – urine flow rate, $GFR$ – glomerular filtration rate, $\Phi_{t\_wreab}$ – tubular water reabsorption rate. | – |
| **Renal module, body fluids** | | | |
| 135 | $\Phi_{win} = \Phi_{win\_norm} \cdot \left(0.25 + 1.5 \cdot \left(1 + exp\left(2 - 2 \cdot \dfrac{C_{adh}}{C_{adh\_norm}}\right)\right)^{-1}\right)$ | $\Phi_{win}$ – rate of water intake, $\Phi_{win\_norm}$ – nominal value of water consumption rate, $C_{adh}/C_{adh\_norm}$ – normalized antidiuretic hormone concentration. | – |
| 136 | $\dfrac{dTBW}{dt} = \Phi_{win} - \Phi_u$ | $TBW$ – total body water, $\Phi_{win}$ – rate of water intake, $\Phi_u$ – urine flow rate. | – |
| 137 | $V = 111.5 \cdot TBW + 650$ | Total blood volume $V$ as a linear function of total body water $TBW$. | – |
| 138 | $V_{ecf} = 0.37 \cdot TBW + 2.7$ | Extracellular fluid volume $V_{ecf}$ as a linear function of total body water $TBW$. | – |
| **Renal module, glomerular filtration** | | | |
| 139 | $R_{aa} = R_{aa\_0} \cdot \beta_{rsna} \cdot \Sigma_{tgf} \cdot \Sigma_{myo} \cdot \Psi_{AT1\_aa} \cdot \left(1 - CCB_{aa}\right) \cdot \left(1 - Diuretic_{aa}\right)$ | The resistance of a single afferent arteriole $R_{aa}$ is equal to the nominal value $R_{aa\_0}$ multiplied by the influence functions described below. $CCB_{aa}$ – effect of the calcium channel blocker amlodipine, $Diuretic_{aa}$ – effect of the thiazide diuretic hydrochlorothiazide (HCTZ). | The vasodilatory effects of amlodipine and HCTZ were taken from the model extension (Kutumova et al., 2022) |

| | | | |
|---|---|---|---|
| 140 | $R_{aa\_0} = 1.25E8 \cdot \frac{128 \cdot Vis \cdot L_{aa}}{\pi \cdot d_{aa}^4}$ | The nominal value of the resistance of a single afferent arteriole $R_{aa\_0}$ can be determined by Poiseuille's law. $d_{aa}$ and $L_{aa}$ – diameter and length of the arteriole, *Vis* – blood viscosity. The coefficient 1.25E8 is used to convert units from cP · μm$^{-3}$ to mmHg · min · L$^{-1}$. | – |
| 141 | $R_{preglom} = R_{preglom\_0} \cdot \beta_{rsna} \cdot \Sigma_{myo} \cdot \Psi_{AT1\_preglom} \cdot \left(1 - CCB_{preglom}\right) \cdot \left(1 - Diuretic_{preglom}\right)$ | The resistance of interlobar, arcuate, and interlobular arteries $R_{preglom}$ is equal to the nominal value $R_{preglom\_0}$ multiplied by the influence functions described below. $CCB_{preglom}$ – effect of the calcium channel blocker amlodipine, $Diuretic_{preglom}$ – effect of the thiazide diuretic hydrochlorothiazide (HCTZ). | The vasodilatory effects of amlodipine and HCTZ were taken from the model extension (Kutumova et al., 2022) |
| 142 | $\beta_{rsna} = 1.5 \cdot \left(RSNA - 1\right) + 1$ | Effect of renal sympathetic nerve activity *RSNA* on the resistance of single afferent arterioles ($R_{aa}$) and the resistance of interlobar, arcuate, and interlobular arteries ($R_{preglom}$). | – |
| 143 | $\frac{d\Sigma_{tgf}}{dt} = 0.3408 + 3.449 \cdot \left(3.88 + \exp\left(\frac{\Phi_{md\_sod} - 3.859}{-0.9617}\right)\right)^{-1} - \Sigma_{tgf}$ | $\Sigma_{tgf}$ – tubuloglomerular feedback signal, $\Phi_{md\_sod}$ – macula densa sodium flow rate. | – |
| 144 | $\frac{d\Sigma_{myo}}{dt} = sl_{Pgh} \cdot \frac{P_{gh}}{P_{gh\_norm}} + 1 - sl_{Pgh} - \Sigma_{myo}$ | $\Sigma_{myo}$ – myogenic autoregulation signal, $P_{gh}/P_{gh\_norm}$ – the normalized value of glomerular hydrostatic pressure, $sl_{Pgh}$ – constant. | – |
| 145 | $\Psi_{AT1\_aa} = A_{AT1\_aa} + B_{AT1\_aa} \cdot AT1\_ANGII - \frac{C_{AT1\_aa}}{AT1\_ANGII}$ | Effect of AT1-bound angiotensin II (*AT1_ANGII*) on the resistance of a single afferent arteriole. $A_{AT1\_aa}$, $B_{AT1\_aa}$, and $C_{AT1\_aa}$ – constants. | – |
| 146 | $\Psi_{AT1\_preglom} = A_{AT1\_preglom} + B_{AT1\_preglom} \cdot AT1\_ANGII - \frac{C_{AT1\_preglom}}{AT1\_ANGII}$ | Effect of AT1-bound angiotensin II (*AT1_ANGII*) on the resistance of interlobar, arcuate, and interlobular arteries. $A_{AT1\_preglom}$, $B_{AT1\_preglom}$, and $C_{AT1\_preglom}$ – constants. | – |
| 147 | $R_a = \frac{R_{aa}}{N_{nephrons}} + R_{preglom}$ | $R_a$ – resistance of afferent vessels, $R_{aa}$ – resistance of a single afferent arteriole, $N_{nephrons}$ – number of nephrons in the kidneys, $R_{preglom}$ – resistance of interlobar, arcuate, and interlobular arteries. | – |
| 148 | $R_{a\_dyne} = 79.68 \cdot R_a$ | Conversion units of $R_a$ from mmHg · min · L$^{-1}$ to dyn · s · cm$^{-5}$. | – |
| 149 | $R_{ea} = R_{ea\_0} \cdot \Psi_{AT1\_ea} \cdot \left(1 - CCB_{ea}\right) \cdot \left(1 - Diuretic_{ea}\right)$ | The resistance of a single efferent arteriole $R_{ea}$ is equal to the nominal value $R_{ea\_0}$ multiplied by the effect function of AT1-bound angiotensin II ($\Psi_{AT1\_ea}$). $CCB_{ea}$ – effect of the calcium channel blocker amlodipine, $Diuretic_{ea}$ – effect of the thiazide diuretic hydrochlorothiazide (HCTZ). | The vasodilatory effects of amlodipine and HCTZ were taken from the model extension (Kutumova et al., 2022) |
| 150 | $R_{ea\_0} = 1.25E8 \cdot \frac{128 \cdot Vis \cdot L_{ea}}{\pi \cdot d_{ea}^4}$ | The nominal value of the resistance of a single efferent arteriole $R_{ea\_0}$ can be determined by Poiseuille's law. $d_{ea}$ and $L_{ea}$ – diameter and length of the arteriole, *Vis* – blood viscosity. The coefficient 1.25E8 is used to convert units from cP · μm$^{-3}$ to mmHg · min · L$^{-1}$. | – |

| | | | |
|---|---|---|---|
| 151 | $\Psi_{AT1\_ea} = A_{AT1\_ea} + B_{AT1\_ea} \cdot AT1\_ANGII - \dfrac{C_{AT1\_ea}}{AT1\_ANGII}$ | Effect of AT1-bound angiotensin II (*AT1_ANGII*) on the resistance of a single efferent arteriole. $A_{AT1\_ea}$, $B_{AT1\_ea}$, and $C_{AT1\_ea}$ – constants. | – |
| 152 | $R_e = \dfrac{R_{ea}}{N_{nephrons}}$ | $R_e$ – resistance of all efferent arterioles, $R_{ea}$ – resistance of a single efferent arteriole, $N_{nephrons}$ – number of nephrons in the kidneys. | – |
| 153 | $R_{e\_dyne} = 79.68 \cdot R_e$ | Conversion units of $R_e$ from mmHg · min · L$^{-1}$ to dyn · s · cm$^{-5}$. | – |
| 154 | $RVR = R_a + \dfrac{RBF - GFR}{RBF} \cdot R_e + R_v$ | *RVR* – renal vascular resistance, $R_a$ – resistance of afferent vessels, $R_e$ – resistance of all efferent arterioles, $R_v$ – renal venous resistance, *GFR* – glomerular filtration rate, *RBF* – renal blood flow. | – |
| 155 | $RBF = \dfrac{MAP - P_v + K_{FG} \cdot R_e \cdot \left(MAP - P_B - P_{go}\right)}{R_a + R_e + R_v + K_{FG} \cdot R_e \cdot R_a}$ | *RBF* – renal blood flow, *MAP* – mean arterial pressure, $P_v$ – renal venous pressure, $P_B$ – hydrostatic pressure in Bowman's space, $P_{go}$ – glomerular capillary oncotic pressure, $K_{FG}$ – glomerular filtration coefficient, $R_a$ – resistance of afferent vessels, $R_e$ – resistance of all efferent arterioles, $R_v$ – renal venous resistance. | – |
| 156 | $P_{gh} = MAP - RBF \cdot R_a$ | Calculation of glomerular hydrostatic pressure $P_{gh}$ using Ohm's law. *MAP* – mean arterial pressure, *RBF* – renal blood flow, $R_a$ – resistance of afferent vessels. | – |
| 157 | $RPF = RBF \cdot \left(1 - 0.01 \cdot Hct\right)$ | *RPF* – renal plasma flow, *RBF* – renal blood flow, *Hct* – hematocrit. | – |
| 158 | $FF = \dfrac{GFR}{RPF}$ | Filtration fraction *FF* is the ratio of the glomerular filtration rate *GFR* to the renal plasma flow *RPF*. | – |
| 159 | $C_M = 0.1 \cdot \dfrac{TP}{FF} \cdot \ln\left(\dfrac{1}{1 - FF}\right)$ | $C_M$ – plasma protein mean concentration within the glomerular capillaries, *TP* – total protein, *FF* – filtration fraction. | – |
| 160 | $\dfrac{dP_{go}}{dt} = 5 \cdot \left(C_M - 2\right) - P_{go}$ | $P_{go}$ – glomerular capillary oncotic pressure, $C_M$ – plasma protein mean concentration within the glomerular capillaries. | – |
| 161 | $K_{FG} = K_{FG\_0} \cdot \left(sl_{KFG} \cdot \dfrac{AT1\_ANGII_{norm}}{AT1\_ANGII} + 1 - sl_{KFG}\right)$ | The glomerular filtration coefficient $K_{FG}$ is calculated as the product of the normal value $K_{FG\_0}$ and a linear function (with slope $0 < sl_{KFG} < 1$) expressing the inverse relationship between $K_{FG}$ and the normalized concentration of *AT1_ANGII*. The function is chosen so that the value $AT1\_ANGII = AT1\_ANGII_{norm}$ gives the result $K_{FG} = K_{FG\_0}$. | – |
| 162 | $GFR = K_{FG} \cdot \left(P_{gh} - P_B - P_{go}\right)$ | *GFR* – glomerular filtration rate, $K_{FG}$ – glomerular filtration coefficient, $P_{gh}$ – glomerular hydrostatic pressure, $P_B$ – hydrostatic pressure in Bowman's space, $P_{go}$ – glomerular capillary oncotic pressure. | – |

**Table S3**. Model parameters with values for a normotensive person[3]

| № | Notation | Description | Initial value | Primary source | Normal range[4] | Units |
|---|---|---|---|---|---|---|
| 001 | $A_1$ | Total metabolic intensity | 0.00076 | Fitted | 0.00032 – 0.00128 (Kutumova et al., 2021) | $mL^{-1}$ |
| 002 | $A_{10}$ | Constant for calculating the conductivity of the tricuspid valve and systemic veins | 0.8 | (Proshin and Solodyannikov, 2006) | – | $mL \cdot s^{-1} \cdot mmHg^{-2}$ |
| 003 | $A_{11}$ | Systemic and pulmonary venous tone | 0.0325 | (Proshin and Solodyannikov, 2006) | – | $mL \cdot mmHg^{-1}$ |
| 004 | $A_{12}$ | Reactivity of the cardiac center | 0.19336 | (Proshin and Solodyannikov, 2006) | – | $s^{-1}$ |
| 005 | $A_{13}$ | Sympathetic sensitivity of the pulmonary microvessels | 0.65 | (Proshin and Solodyannikov, 2006) | – | $mL \cdot mmHg^{-1}$ |
| 006 | $A_{14}$ | Sensitivity of pulmonary microvessels to oxygen debt | 0.08265 | (Proshin and Solodyannikov, 2006) | – | $s^{-1} \cdot mmHg^{-1}$ |
| 007 | $A_{15}$ | Sensitivity factor for calculating RV systolic pressure | 22.0 | (Proshin and Solodyannikov, 2006) | – | $s^{-1} \cdot mmHg^{-1}$ |
| 008 | $A_{16}$ | Constant for calculating the conductivity of the mitral valve and pulmonary veins | 3.3 | (Proshin and Solodyannikov, 2006) | – | $mL \cdot s^{-1} \cdot mmHg^{-2}$ |
| 009 | $A_{18}$ | Sympathetic sensitivity of the pulmonary arteries | 40.0 | (Proshin and Solodyannikov, 2006) | – | $mL \cdot s$ |
| 010 | $A_{19}$ | Pulmonary arterial tone | 0.02 | (Kutumova et al., 2021) | – | $mmHg \cdot s \cdot mL^{-1}$ |
| 011 | $A_2$ | Functional status of the body | 0.3752 | (Proshin and Solodyannikov, 2006) | – | – |
| 012 | $A_3$ | Sympathetic sensitivity of the systemic microvessels | 0.1 | (Proshin and Solodyannikov, 2006) | – | $mL \cdot mmHg^{-1}$ |
| 013 | $A_4$ | Sensitivity of systemic microvessels to oxygen debt | 0.031537 | (Proshin and Solodyannikov, 2006) | – | $s^{-1} \cdot mmHg^{-1}$ |
| 014 | $A_5$ | Sensitivity factor for calculating LV systolic pressure | 22.0 | (Proshin and Solodyannikov, 2006) | – | $s \cdot mmHg^{-1}$ |
| 015 | $A_6$ | Constant for calculating the conductivity of the tricuspid valve and systemic veins | 3.3 | (Proshin and Solodyannikov, 2006) | – | $mL \cdot s^{-1} \cdot mmHg^{-2}$ |
| 016 | $A_7$ | Constant for calculating the conductivity of the tricuspid valve and systemic veins | 4.0 | (Proshin and Solodyannikov, 2006) | – | $mmHg^{-1}$ |
| 017 | $A_8$ | Sympathetic sensitivity of the systemic arteries | 45.0 | (Proshin and Solodyannikov, 2006) | – | $mL \cdot s$ |
| 018 | $A_9$ | Systemic arterial tone | 0.07 | (Proshin and Solodyannikov, 2006) | – | $mmHg \cdot s \cdot mL^{-1}$ |
| 019 | $A_{AT1\_aa}$ | Constant for AT1-bound angiotensin II effect on afferent arteriole resistance | 1.3754 | (Kutumova et al., 2021) | – | – |
| 020 | $A_{AT1\_ALVL}$ | Constant for AT1-bound angiotensin II effect on blood flow through systemic microvessels | 1.7323 | (Kutumova et al., 2021) | – | – |
| 021 | $A_{AT1\_ea}$ | Constant for AT1-bound angiotensin II effect on efferent arteriole resistance | 1.6601 | (Kutumova et al., 2021) | – | – |
| 022 | $A_{AT1\_preglom}$ | Constant for AT1-bound angiotensin II effect on resistance of interlobar, arcuate, and interlobular arteries | 0.4464 | (Kutumova et al., 2021) | – | – |
| 023 | $AO_2$ | Arterial oxygen content | 0.2 | Calculated as $0.001 \cdot He \cdot C_H \cdot SpO_2$ | 0.145 – 0.244 (Hattori et al., 2004) | – |
| 024 | $AT1\_ANGII_{norm}$ | Normal concentration of AT1-bound angiotensin II | 16.63 | (Hallow and Gebremichael , 2017) | – | $fmol \cdot mL^{-1}$ |

[3] All parameters of the model, except for the pharmacokinetic parameters of drugs, which are given in Table S5 below
[4] Data range or mean ± SD

| | | | | | | |
|---|---|---|---|---|---|---|
| 025 | $B_{AT1\_aa}$ | Constant for AT1-bound angiotensin II effect on afferent arteriole resistance | 0.0549 | (Kutumova et al., 2021) | – | – |
| 026 | $B_{AT1\_ALVL}$ | Constant for AT1-bound angiotensin II effect on blood flow through systemic microvessels | 0.0032 | (Kutumova et al., 2021) | – | – |
| 027 | $B_{AT1\_ea}$ | Constant for AT1-bound angiotensin II effect on efferent arteriole resistance | 0.0383 | (Kutumova et al., 2021) | – | – |
| 028 | $B_{AT1\_preglom}$ | Constant for AT1-bound angiotensin II effect on resistance of interlobar, arcuate, and interlobular arteries | 0.0661 | (Kutumova et al., 2021) | – | – |
| 029 | $c_{ACE}$ | Rate of conversion of angiotensin I to angiotensin II by ACE | 0.90167 | (Hallow et al., 2014) | – | $min^{-1}$ |
| 030 | $c_{ACE2}$ | Rate of conversion of angiotensin II to angiotensin-(1-7) by ACE2 | 0.04 | (Hallow et al., 2014) | – | $min^{-1}$ |
| 031 | $C_{adh\_norm}$ | Normal plasma concentration of antidiuretic hormone | 4.97 | Fitted | 1.0 – 13.3 (Yarmohammadi et al., 2015) | $pg \cdot mL^{-1}$ |
| 032 | $C_{al\_norm}$ | Normal plasma concentration of aldosterone | 255.04 | Fitted | 70 – 300 (Fischbach, 2003) | $pg \cdot mL^{-1}$ |
| 033 | $c_{ANGII\_ANGIV}$ | Rate of conversion of angiotensin II to angiotensin IV | 0.39167 | (Hallow et al., 2014) | – | $min^{-1}$ |
| 034 | $C_{anp\_norm}$ | Normal plasma concentration of atrial natriuretic peptide | 35.05 | Fitted | 7.4 – 152.0 (Cannone et al., 2018; Nozuki et al., 1986) | $ng \cdot L^{-1}$ |
| 035 | $c_{AT1}$ | Rate of angiotensin II binding to AT1 receptors | 0.19667 | (Hallow et al., 2014) | – | $min^{-1}$ |
| 036 | $C_{AT1\_aa}$ | Constant for AT1-bound angiotensin II effect on afferent arteriole resistance | 0.1437 | (Kutumova et al., 2021) | – | – |
| 037 | $C_{AT1\_ALVL}$ | Constant for AT1-bound angiotensin II effect on blood flow through systemic microvessels | 0.2170 | (Kutumova et al., 2021) | – | – |
| 038 | $C_{AT1\_ea}$ | Constant for AT1-bound angiotensin II effect on efferent arteriole resistance | 0.2441 | (Kutumova et al., 2021) | – | – |
| 039 | $C_{AT1\_preglom}$ | Constant for AT1-bound angiotensin II effect on resistance of interlobar, arcuate, and interlobular arteries | 0.2133 | (Kutumova et al., 2021) | – | – |
| 040 | $c_{AT2}$ | Rate of angiotensin II binding to AT2 receptors | 0.065 | (Hallow et al., 2014) | – | $min^{-1}$ |
| 041 | $c_{chym}$ | Rate of conversion of angiotensin I to angiotensin II by chymase | 0.01833 | (Hallow et al., 2014) | – | $min^{-1}$ |
| 042 | $C_H$ | Oxygen capacity of hemoglobin | 1.35 | Fitted | 1.32 – 1.39 (Dijkhuizen et al., 1977) | $mL \cdot g^{-1}$ |
| 043 | $C_K$ | Serum potassium concentration | 3.88 | Fitted | 3.5 – 5.5 (Rastegar, 1990) | $mEq \cdot L^{-1}$ |
| 044 | $c_{nep}$ | Rate of conversion of angiotensin I to angiotensin-(1-7) by neprilysin | 0.01833 | (Hallow et al., 2014) | – | $min^{-1}$ |
| 045 | $d_{aa}$ | Afferent arteriolar diameter | 19.45 | Fitted | 8.7 – 23.9 (Neal et al., 2019; Hill et al., 2006) | μm |
| 046 | $d_{ea}$ | Efferent arteriolar diameter | 17.55 | Fitted | 13.5 – 18.3 (Neal et al., 2019) | μm |
| 047 | $FS_{threshold}$ | Frank-Starling law threshold | 45.55 | Calculated as $m/70 \cdot FS_{threshold0}$ | – | mL |
| 048 | $FS_{threshold0}$ | Average normal value of the Frank-Starling law threshold | 39.85 | Fitted | – | mL |

| | | | | | | |
|---|---|---|---|---|---|---|
| 049 | $G_{AL0}$ | Basic systemic arterial elasticity | 0.76 | Fitted | 0.33 – 1.00 (Laskey et al., 1990) | mmHg·mL$^{-1}$ |
| 050 | $G_{AR0}$ | Basic pulmonary arterial elasticity | 0.15 | Fitted | 0.08 – 0.26 (Thenappan et al., 2016) | mmHg·mL$^{-1}$ |
| 051 | $G_{HL}$ | LV wall elasticity | 0.079 | Fitted | 0.01 – 0.43 (Zhang and Kovács, 2008) | mmHg·mL$^{-1}$ |
| 052 | $G_{HR}$ | RV wall elasticity | 0.048 | Fitted | 0.01 – 0.43 (Zhang and Kovács, 2008) | mmHg·mL$^{-1}$ |
| 053 | $G_{VL0}$ | Basic elasticity of the systemic veins | 0.028 | Fitted | – | mmHg·mL$^{-1}$ |
| 054 | $G_{VR0}$ | Basic elasticity of the pulmonary veins | 0.046 | Fitted | – | mmHg·mL$^{-1}$ |
| 055 | *glucose* | Plasma glucose | 5.01 | Fitted | 3.9 – 6.1 (Dedov et al., 2017) | mmol·L$^{-1}$ |
| 056 | $h_{ANG17}$ | Half-life of angiotensin (1-7) | 30.0 | (Hallow et al., 2014) | 19.2 – 51.6 (Rodgers et al., 2006) | min |
| 057 | $h_{ANGI}$ | Half-life of angiotensin I | 0.25 | (Kutumova et al., 2021) | 0.25 ± 0.08 (Admiraal et al., 1993) | min |
| 058 | $h_{ANGII}$ | Half-life of angiotensin II | 0.9 | (Kutumova et al., 2021) | Men: 1.0; Women: 0.8 (Magness et al., 1994; Donato et al., 1972) | min |
| 059 | $h_{ANGIV}$ | Half-life of angiotensin IV | 0.5 | (Hallow et al., 2014) | – | min |
| 060 | $h_{AT1}$ | Half-life of AT1-bound angiotensin II | 12.0 | (Hallow et al., 2014) | 12.0 (Inada et al., 1999) | min |
| 061 | $h_{AT2}$ | Half-life of AT2-bound angiotensin II | 12.0 | (Hallow et al., 2014) | – | min |
| 062 | $h_{renin}$ | Half-life of circulating renin | 12.0 | (Hallow et al., 2014) | 10.0 – 15.0 (Skrabal, 1974) | min |
| 063 | *Hct* | Hematocrit | 42.62 | Fitted | Men: 40 – 54 Women: 36 – 48 (Billett, 1990) | % |
| 064 | *He* | Hemoglobin | 154.3 | Fitted | Men:140–180 Women:120–160 (Billett, 1990) | g·L$^{-1}$ |
| 065 | $Heart_{Base}$ | Basic activity of the cardiac center | 0.17 | Fitted | – | – |
| 066 | $Heart_{Baro}$ | Baroreceptor sensitivity of the cardiac center | 0.6 | (Proshin and Solodyannikov, 2006) | – | – |
| 067 | $Heart_{Oxygen}$ | Sensitivity of the cardiac center to fatigue | 1.75 | (Proshin and Solodyannikov, 2006) | – | – |
| 068 | $Heart_{Stress}$ | Stress sensitivity of the cardiac center | 1.5 | (Proshin and Solodyannikov, 2006) | – | – |
| 069 | $Heart_{VO2}$ | Respiratory sensitivity of the cardiac center | 1.0 | (Proshin and Solodyannikov, 2006) | – | – |
| 070 | $k_{AL}$ | Ratio of unstressed volume to stressed volume of the systemic arteries | 0.92 | Fitted | 0.7 – 1.0 (Magder, 2016) | – |
| 071 | $k_{AR}$ | Ratio of unstressed volume to stressed volume of the pulmonary arteries | 0.85 | Fitted | 0.7 – 1.0 (Magder, 2016) | – |

| | | | | | | |
|---|---|---|---|---|---|---|
| 072 | $K_{FG\_0}$ | Normal glomerular filtration coefficient | 0.0051 | Fitted | 0.0039–0.0162 (Hoang et al., 2003) | $L \cdot min^{-1} \cdot mmHg^{-1}$ |
| 073 | $k_{HL}$ | Ratio of unstressed volume to stressed volume of the left ventricle | 0.13 | Fitted | 0.0 – 0.3 (Kutumova et al., 2021) | – |
| 074 | $K_{HLAL}$ | Aortic valve regurgitation coefficient | 0.0 | (Proshin and Solodyannikov, 2006) | – | – |
| 075 | $k_{HR}$ | Ratio of unstressed volume to stressed volume of the right ventricle | 0.15 | Fitted | 0.0 – 0.3 (Kutumova et al., 2021) | – |
| 076 | $K_{HRAR}$ | Pulmonary valve regurgitation coefficient | 0.0 | (Proshin and Solodyannikov, 2006) | – | – |
| 077 | $k_{VL}$ | Ratio of unstressed volume to stressed volume of the systemic veins | 1.0 | Fitted | 0.7 – 1.0 (Magder, 2016) | – |
| 078 | $K_{VLHR}$ | Tricuspid valve regurgitation coefficient | 0.0 | (Proshin and Solodyannikov, 2006) | – | – |
| 079 | $k_{VR}$ | Ratio of unstressed volume to stressed volume of the pulmonary veins | 0.98 | Fitted | 0.7 – 1.0 (Magder, 2016) | – |
| 080 | $K_{VRHL}$ | Mitral valve regurgitation coefficient | 0.0 | (Proshin and Solodyannikov, 2006) | – | – |
| 081 | $K_{L0}$ | Inotropic status of the LV | 0.56 | Fitted | 0.50 – 0.80 (Solodyannikov, 1994) | – |
| 082 | $K_{R0}$ | Inotropic status of the RV | 0.52 | Fitted | 0.50 – 0.80 (Solodyannikov, 1994) | – |
| 083 | $L_{aa}$ | Afferent arteriolar length | 120.8 | Fitted | 112.0 (Neal et al., 2019) | μm |
| 084 | $L_{ea}$ | Efferent arteriolar length | 127.5 | Fitted | 138.0 (Neal et al., 2019) | μm |
| 085 | $m$ | Body mass | 80.0 | Fitted | – | kg |
| 086 | $N_{nephrons}$ | Number of nephrons in the kidneys | 2702905 | Fitted | 1.50E6 – 3.00E6 (Bertram et al., 2011) | – |
| 087 | $N_{rsna}$ | Normalized renal sympathetic nerve activity | 1.0 | (Karaaslan et al., 2005) | – | – |
| 088 | $n_{\varepsilon\_dt}$ | Normal fractional distal sodium reabsorption | 0.44 | Fitted | – | – |
| 089 | $n_{\eta\_cd}$ | Normal fractional collecting duct sodium reabsorption | 0.82 | Fitted | – | – |
| 090 | $n_{\eta\_pt}$ | Normal fractional sodium reabsorption in the proximal tubule and the loop of Henle | 0.85 | Fitted | 0.67 – 0.97 (Fliser et al., 1997; Bochud et al., 2009; Jin et al., 2009) | – |
| 091 | $n_{\varepsilon\_dt} + n_{\eta\_cd} - n_{\eta\_cd} \cdot n_{\varepsilon\_dt}$ | Normal fractional sodium reabsorption in the distal tubule and subsequent parts of the nephron | 0.90 | Fitted | 0.78 – 0.98 (Fliser et al., 1997; Bochud et al., 2009; Jin et al., 2009) | – |
| 092 | $P_0$ | Baroreceptor sensitivity threshold | 2.0 | (Proshin and Solodyannikov, 2006) | – | mmHg |
| 093 | $P_B$ | Hydrostatic pressure in the Bowman's space | 13.35 | Fitted | 10.0 – 15.0 (Digne-Malcolm et al., 2016) | mmHg |
| 094 | $P_{gh\_norm}$ | Normal glomerular hydrostatic pressure | 56.84 | Fitted | 48.0 – 63.0 (Guberina et al., 2013) | mmHg |

| | | | | | | |
|---|---|---|---|---|---|---|
| 095 | $P_v$ | Renal venous pressure | 6.0 | (Digne-Malcolm et al., 2016) | – | mmHg |
| 096 | $PRC_{nom}$ | Nominal value of plasma renin concentration | 24.62 | Fitted | 3.42 – 69.4 (Perschel et al., 2004) | $pg \cdot mL^{-1}$ |
| 097 | $R_{preglom\_0}$ | Nominal resistance of interlobar, arcuate, and interlobular arteries | 10.08 | Fitted | 10.0 – 20.0 (Hallow and Gebremichael , 2017) | $mmHg \cdot min \cdot L^{-1}$ |
| 098 | $R_v$ | Renal venous resistance | 18.10 | Fitted | 11.3 – 20.1 (Kutumova et al., 2021) | $mmHg \cdot min \cdot L^{-1}$ |
| 099 | $RO_2$ | Oxygen demand | 6.26 | Calculated as $m/70 \cdot RO_{20}$ | – | $mL \cdot s^{-1}$ |
| 100 | $RO_{20}$ | Average normal value of oxygen demand | 5.48 | Fitted | ≈ 4.2 (Treacher and Leach, 1998) | $mL \cdot s^{-1}$ |
| 101 | $sl_{baro}$ | Slope of the linear function of the effect of AT1-bound angiotensin II on baroreceptor activity | 0.0499 | (Kutumova et al., 2022) | – | – |
| 102 | $sl_{KFG}$ | Slope of the linear function of the effect of AT1-bound angiotensin II on filtration coefficient | 0.2015 | (Kutumova et al., 2022) | – | – |
| 103 | $sl_{Pgh}$ | Slope of the linear function of the effect of glomerular hydrostatic pressure on myogenic autoregulation signal | 0.8294 | (Kutumova et al., 2021) | – | – |
| 104 | $sl_{stress}$ | Slope of the linear function of the effect of AT1-bound angiotensin II on stress receptor activity | 0.2133 | (Kutumova et al., 2022) | – | – |
| 105 | $slope_{AT1\_PRC}$ | Constant for AT1-bound angiotensin II effect on renin secretion rate | 1.2 | (Hallow and Gebremichael , 2017) | – | – |
| 106 | $SpO_2$ | Arterial oxygen saturation | 0.96 | Fitted | 0.95 – 0.99 (Goldberg et al., 2017) | – |
| 107 | $st$ | Stress factor, level of steroid hormones in the blood | 0.25 | (Proshin and Solodyannikov, 2006) | – | – |
| 108 | $SV_{max}$ | Theoretical maximum stroke volume | 228.57 | Calculated as $m/70 \cdot SV_{max0}$ | – | mL |
| 109 | $SV_{max0}$ | Average value of theoretical maximum stroke volume | 200.0 | (Proshin and Solodyannikov, 2006) | – | mL |
| 110 | $T_{al}$ | Time constant for calculating the aldosterone secretion | 30.0 | (Karaaslan et al., 2005) | – | min |
| 111 | $TP$ | Total protein | 75.72 | Fitted | 60.0 – 86.0 (Busher, 1990; Gardner and Scott, 1980) | $g \cdot L^{-1}$ |
| 112 | $urea$ | Plasma urea concentration | 2.02 | Fitted | 1.8 – 7.1 (Hosten, 1990) | $mmol \cdot L^{-1}$ |
| 113 | $Vis$ | Blood viscosity | 4.86 | Calculated as $Vis = 1.23 \cdot \left(1 - \frac{Hct}{99}\right)^{-n}$, $n = 1.7 + 9.86 \cdot \exp(-0.0607 \cdot Hct)$ | 4.66 ± 0.72 (Furukawa et al., 2016) | cP |
| 114 | $Vis_{norm}$ | Average normal blood viscosity | 4.65 | (Hund et al., 2017) | 4.66 ± 0.72 (Furukawa et al., 2016) | cP |
| 115 | $X_{PRC\_PRA}$ | Equilibrium ratio of plasma renin activity to plasma renin concentration | 0.87 | Fitted | – | $fmol \cdot min^{-1} \cdot pg^{-1}$ |
| 116 | $Y_{ALVL0}$ | Basic conductivity of the systemic microvessels | 1.363 | (Kutumova et al., 2021) | – | $mL \cdot s^{-1} \cdot mmHg^{-1}$ |
| 117 | $Y_{ARVR0}$ | Basic conductivity of the pulmonary microvessels | 13.89 | Fitted | – | $mL \cdot s^{-1} \cdot mmHg^{-1}$ |

| | | | | | | |
|---|---|---|---|---|---|---|
| 118 | $Y_{HLAL}$ | Conductivity of the aortic valve and systemic arteries | 7.0 | (Kutumova et al., 2021) | – | mL·s$^{-1}$·mmHg$^{-1}$ |
| 119 | $Y_{HRAR}$ | Conductivity of the pulmonary valve and pulmonary arteries | 57.0 | (Kutumova et al., 2021) | – | mL·s$^{-1}$·mmHg$^{-1}$ |
| 120 | $Y_{VLHR0}$ | Basic conductivity of the tricuspid valve and systemic veins | 90.0 | (Kutumova et al., 2021) | – | mL·s$^{-1}$·mmHg$^{-1}$ |
| 121 | $Y_{VRHL0}$ | Basic conductivity of the mitral valve and pulmonary veins | 57.0 | (Kutumova et al., 2021) | – | mL·s$^{-1}$·mmHg$^{-1}$ |
| 122 | $\xi_{map}$ | Effect of mean arterial pressure on aldosterone secretion | 1.0 | (Karaaslan et al., 2005) | – | – |
| 123 | $\tau_{MD\_renin}$ | Constant to calculate the effect of macula densa sodium flow on renin secretion rate | 0.0959 | (Kutumova et al., 2021) | – | – |
| 124 | $\Phi_{md\_sod\_0}$ | Nominal value of macula densa sodium flow rate | 2.80 | Fitted | – | mEq·min$^{-1}$ |
| 125 | $\Phi_{sodin}$ | Sodium intake | 0.048 | Fitted | 0.035 – 0.174<br>(Luft et al., 1982) | mEq·min$^{-1}$ |
| 126 | $\Phi_{win\_norm}$ | Normal value of water intake | 0.0016 | Fitted | 0.0019 ± 0.0007<br>(Malisova et al., 2016) | L·min$^{-1}$ |
| 127 | $\omega_{AL\_nom}$ | Nominal unstressed volume of the systemic arteries | 551.2 | $\approx k_{AL} \cdot V_{AL}(0)$ | – | mL |
| 128 | $\omega_{AR\_nom}$ | Nominal unstressed volume of the pulmonary arteries | 136.7 | $\approx k_{AR} \cdot V_{AR}(0)$ | – | mL |
| 129 | $\omega_{HL}$ | Unstressed LV volume | 18.1 | $\approx k_{HL} \cdot V_{HL}(0)$ | 0.0 – 42.0<br>(Kutumova et al., 2021) | mL |
| 130 | $\omega_{HR}$ | Unstressed RV volume | 20.6 | $\approx k_{HR} \cdot V_{HR}(0)$ | 0.0 – 42.0<br>(Kutumova et al., 2021) | mL |
| 131 | $\omega_{VL}$ | Unstressed volume of the systemic veins | 3260.3 | $\approx k_{VL} \cdot V_{VL}(0)$ | – | mL |
| 132 | $\omega_{VR}$ | Unstressed volume of the pulmonary veins | 292.1 | $\approx k_{VR} \cdot V_{VR}(0)$ | – | mL |

**Table S4**. Model variables with equilibrium values for a normotensive person

| № | Variables | Description | Values | Normal range[5] | Units |
|---|---|---|---|---|---|
| 001 | $ANG17$ | Plasma angiotensin-(1-7) concentration | 14.792 | 14.1 – 31.7 (Ferrario et al., 1998) | fmol·mL$^{-1}$ |
| 002 | $ANGI$ | Plasma angiotensin I concentration | 7.860 | 2.8 – 28.5 (Lawrence et al., 1990; Nussberger et al., 1992) | fmol·mL$^{-1}$ |
| 003 | $ANGII$ | Plasma angiotensin II concentration | 4.941 | 0.0 – 21.4 (Lawrence et al., 1990; Nussberger et al., 1992; Duggan et al., 1993) | fmol·mL$^{-1}$ |
| 004 | $ANGIV$ | Plasma angiotensin IV concentration | 1.396 | – | fmol·mL$^{-1}$ |
| 005 | $AT1\_ANGII$ | Concentration of AT1-bound angiotensin II | 16.824 | – | fmol·mL$^{-1}$ |
| 006 | $AT2\_ANGII$ | Concentration of AT2-bound angiotensin II | 5.561 | – | fmol·mL$^{-1}$ |
| 007 | $C_{adh}$ | Plasma concentration of antidiuretic hormone | 2.785 | 1.0 – 13.3 (Yarmohammadi et al., 2015) | pg·mL$^{-1}$ |
| 008 | $C_{al}$ | Plasma aldosterone concentration | 215.385 | 70 – 300 (Fischbach, 2003) | pg·mL$^{-1}$ |
| 009 | $C_{anp}$ | Plasma concentration of atrial natriuretic peptide | 42.736 | 7.4 – 152.0 (Cannone et al., 2018; Nozuki et al., 1986) | ng·L$^{-1}$ |
| 010 | $C_M$ | Plasma protein mean concentration within the glomerular capillaries | 8.035 | – | g·L$^{-1}$ |
| 011 | $C_{sod}$ | Serum sodium concentration | 143.594 | 137 – 147 (Payne and Levell, 1968) | mEq·L$^{-1}$ |
| 012 | $CO$ | Cardiac output | 5.019 | 2.51 – 9.00 (Cattermole et al., 2017) | L·min$^{-1}$ |
| 013 | $Cycle_{Length}$ | Cardiac cycle length (RR-interval) | 0.812 | Men: 0.624 – 1.284 Women: 0.600 – 1.213 (Mason et al., 2007) | s |
| 014 | $Cycle_{Time}$ | Current cardiac cycle time | 0.492 | – | s |
| 015 | $DO_2$ | Oxygen debt | 6.054 | – | mL |
| 016 | $DTS_L$ | LV systolic mismatch | 0.0 | – | – |
| 017 | $DTS_R$ | RV systolic mismatch | 0.0 | – | – |
| 018 | $EF$ | Ejection fraction | 55.199 | 50 – 80 (Pfisterer et al., 1985; Saghiv and Sagiv, 2017) | % |
| 019 | $F_{ALVL}$ | Blood flow through the systemic microvessels | 84.176 | – | mL·s$^{-1}$ |
| 020 | $F_{ARVR}$ | Blood flow through the pulmonary microvessels | 83.486 | – | mL·s$^{-1}$ |
| 021 | $F_{HLAL}$ | Blood flow through the aortic valve | 0.0 | – | mL·s$^{-1}$ |
| 022 | $F_{HLAL\_p}$ | Peak flow rate of $F_{HLAL}$ | 461.530 | 347.0 – 677.0 (Kyhl et al., 2013) | mL·s$^{-1}$ |

[5] Data range or mean ± SD

| | | | | | |
|---|---|---|---|---|---|
| 023 | $F_{HRAR}$ | Blood flow through the pulmonary valve | 0.0 | – | mL·s$^{-1}$ |
| 024 | $F_{HRAR\_p}$ | Peak flow rate of $F_{HRAR}$ | 589.434 | 264.5 – 793.0 (Kyhl et al., 2013; Macedo et al., 2007) | mL·s$^{-1}$ |
| 025 | $F_{VLHR}$ | Blood flow through the tricuspid valve | 68.810 | – | mL·s$^{-1}$ |
| 026 | $F_{VLHR\_ap}$ | Active peak of $F_{VLHR}$ | 433.925 | 29 – 770 (Maceira et al., 2006a) | mL·s$^{-1}$ |
| 027 | $F_{VLHR\_ep}$ | Early peak of $F_{VLHR}$ | 599.566 | 105 – 753 (Maceira et al., 2006a) | mL·s$^{-1}$ |
| 028 | $F_{VRHL}$ | Blood flow through the mitral valve | 67.323 | – | mL·s$^{-1}$ |
| 029 | $F_{VRHL\_ap}$ | Active peak of $F_{VRHL}$ | 332.964 | 77 – 497 (Maceira et al., 2006b) | mL·s$^{-1}$ |
| 030 | $F_{VRHL\_ep}$ | Early peak of $F_{VRHL}$ | 798.271 | 178 – 1000 (Maceira et al., 2006b) | mL·s$^{-1}$ |
| 031 | $FF$ | Filtration fraction | 0.113 | – | – |
| 032 | $G_{AL}$ | Systemic arterial elasticity | 0.949 | 0.33 – 1.00 (Laskey et al., 1990) | mmHg·mL$^{-1}$ |
| 033 | $G_{AR}$ | Pulmonary arterial elasticity | 0.194 | 0.08 – 0.26 (Thenappan et al., 2016) | mmHg·mL$^{-1}$ |
| 034 | $G_{VL}$ | Systemic venous elasticity | 0.075 | – | mmHg·mL$^{-1}$ |
| 035 | $G_{VR}$ | Pulmonary venous elasticity | 0.096 | – | mmHg·mL$^{-1}$ |
| 036 | $GFR$ | Glomerular filtration rate | 0.077 | 0.060 – 0.135 (Levin and Stevens, 2013; Cachat et al., 2015) | L·min$^{-1}$ |
| 037 | $gO_2$ | Oxygen consumption | 6.291 | ≈ 4.2 (Treacher and Leach, 1998) | mL·s$^{-1}$ |
| 038 | $H$ | Neurohumoral factor | 1.220 | – | s$^{-1}$ |
| 039 | $Heart_{Rate}$ | Heart rate | 73.890 | 60 – 100 (Ostchega et al., 2011) | beats·min$^{-1}$ |
| 040 | $K_{FG}$ | Glomerular filtration coefficient | 0.0051 | 0.0039–0.0162 (Hoang et al., 2003) | L·min$^{-1}$·mmHg$^{-1}$ |
| 041 | $K_L$ | Inotropic factor of the LV | 0.621 | – | – |
| 042 | $K_R$ | Inotropic factor of the RV | 0.583 | – | – |
| 043 | $LA_{PULSE}$ | Pulse wave of the left atrium | 0.0 | – | mmHg |
| 044 | $M_{sod}$ | Total exchangeable sodium | 2707.016 | 2040 – 3950 (Farber and Soberman, 1956) | mEq |
| 045 | $MAP$ | Mean arterial pressure | 93.799 | 70 – 105 (Doenyas-Barak et al., 2019) | mmHg |
| 046 | $N_{al}$ | Normalized aldosterone concentration | 0.845 | – | – |
| 047 | $N_{als}$ | Normalized aldosterone secretion rate | 0.845 | – | – |
| 048 | $nB$ | Baroreceptor activity | 0.457 | – | – |
| 049 | $nD$ | Fatigue receptor activity | 0.070 | – | – |

| | | | | | |
|---|---|---|---|---|---|
| 050 | *nH* | Sympathetic inotropic sensitivity of the myocardium | 0.273 | – | – |
| 051 | *nS* | Stress receptor activity | 0.181 | – | – |
| 052 | *nSum* | Activity of the cardiac center | 1.265 | – | – |
| 053 | *nV* | Respiratory receptor activity | 0.426 | – | – |
| 054 | *osmolality* | Blood osmolality | 283.108 | 275 – 295 (Fogarty and Loughrey, 2016) | $mOsm \cdot kg^{-1}$ |
| 055 | $P_{AL}$ | Systemic arterial pressure | 101.943 | – | mmHg |
| 056 | $P_{AR}$ | Pulmonary arterial pressure | 14.016 | – | mmHg |
| 057 | $P_{AR\_D}$ | Diastolic pulmonary arterial pressure | 11.160 | 4 – 12 (Marini and Leatherman, 2005; Pagani et al., 1988) | mmHg |
| 058 | $P_{AR\_S}$ | Systolic pulmonary arterial pressure | 18.358 | 15 – 30 (Marini and Leatherman, 2005; Pagani et al., 1988) | mmHg |
| 059 | $P_D$ | Diastolic blood pressure | 80.228 | < 90 (Oparil et al., 2018) | mmHg |
| 060 | $P_{gh}$ | Glomerular hydrostatic pressure | 58.557 | 48.0 – 63.0 (Guberina et al., 2013) | mmHg |
| 061 | $P_{go}$ | Glomerular capillary oncotic pressure | 30.173 | 23.6 – 34.0 (Škrtić et al., 2015; Chagnac et al., 2000; Guasch et al., 1997) | mmHg |
| 062 | $P_{HL}$ | LV pressure | 7.587 | – | mmHg |
| 063 | $P_{HL\_D}$ | LV diastolic pressure | 7.587 | 3 – 12 (Pagani et al., 1988) | mmHg |
| 064 | $P_{HL\_S}$ | LV systolic pressure | 138.914 | 100 – 140 (Pagani et al., 1988) | mmHg |
| 065 | $P_{HR}$ | RV pressure | 4.338 | – | mmHg |
| 066 | $P_{HR\_D}$ | RV diastolic pressure | 4.338 | 2 – 8 (Pagani et al., 1988) | mmHg |
| 067 | $P_{HR\_S}$ | RV systolic pressure | 20.364 | 15 – 30 (Pagani et al., 1988) | mmHg |
| 068 | $P_{ra}$ | Normalized right atrial pressure | 0.876 | – | – |
| 069 | $P_S$ | Systolic blood pressure | 120.942 | < 140 (Oparil et al., 2018) | mmHg |
| 070 | $P_{VL}$ | Systemic venous pressure | 4.886 | 2 – 8 (Klingensmith et al., 2016) | mmHg |
| 071 | $P_{VR}$ | Pulmonary venous pressure | 8.297 | 3 – 20 (Kutumova et al., 2021) | mmHg |
| 072 | *PRA* | Plasma renin activity | 29.170 | 15.0 – 31.7 (Valabhji et al., 2001) | $fmol \cdot mL^{-1} \cdot min^{-1}$ |
| 073 | *PRC* | Plasma renin concentration | 33.414 | 3.42 – 69.4 (Perschel et al., 2004) | $pg \cdot mL^{-1}$ |
| 074 | *PVR* | Pulmonary vascular resistance | 0.093 | 0.0151 – 0.1353 (Klingensmith et al., 2016; Stefanadis et al., 2001) | $s \cdot mmHg \cdot mL^{-1}$ |

| | | | | | |
|---|---|---|---|---|---|
| 075 | $R_a$ | Resistance of the afferent arterioles, and interlobar, arcuate, and interlobular arteries | 29.827 | – | mmHg·min·mL$^{-1}$ |
| 076 | $R_{a\_dyne}$ | $R_a$ in dyn·s·cm$^{-5}$ | 2376.597 | 753 – 6863 (Tsuda et al., 2018; Kutumova et al., 2021) | dyn·s·cm$^{-5}$ |
| 077 | $R_{aa}$ | Single afferent arteriole resistance | 2.746E7 | – | mmHg·min·L$^{-1}$ |
| 078 | $R_{aa\_0}$ | Nominal single afferent arteriole resistance | 2.089E7 | – | mmHg·min·L$^{-1}$ |
| 079 | $R_e$ | Resistance of all efferent arterioles | 28.212 | – | mmHg·min·mL$^{-1}$ |
| 080 | $R_{e\_dyne}$ | $R_e$ in dyn·s·cm$^{-5}$ | 2247.921 | 1669 – 2843 (Kutumova et al., 2021) | dyn·s·cm$^{-5}$ |
| 081 | $R_{ea}$ | Single efferent arteriole resistance | 7.625E7 | – | mmHg·min·L$^{-1}$ |
| 082 | $R_{ea\_0}$ | Nominal single efferent arteriole resistance | 3.330E7 | – | mmHg·min·L$^{-1}$ |
| 083 | $R_{preglom}$ | Resistance of interlobar, arcuate, and interlobular arteries | 19.668 | 10.0 – 20.0 (Hallow and Gebremichael , 2017) | mmHg·min·L$^{-1}$ |
| 084 | $R_{sec}$ | Renin secretion rate | 1.930 | – | pg·mL$^{-1}$·min$^{-1}$ |
| 085 | $RA_{PULSE}$ | Pulse wave of the right atrium | 0.0 | – | mmHg |
| 086 | $RBF$ | Renal blood flow | 1.182 | 0.623 – 1.730 (Bax et al., 2005) | L·min$^{-1}$ |
| 087 | $RPF$ | Renal plasma flow | 0.678 | 0.628 ± 0.162 (Škrtić et al., 2015) | L·min$^{-1}$ |
| 088 | $RSNA$ | Renal sympathetic nerve activity | 1.154 | – | – |
| 089 | $RVR$ | Renal vascular resistance | 74.308 | 55.1 – 83.6 (Kutumova et al., 2021) | mmHg·min·L$^{-1}$ |
| 090 | $SV$ | LV stroke volume | 67.927 | 39.1 – 115.3 (Cattermole et al., 2017) | mL |
| 091 | $SVR$ | Systemic vascular resistance (can be estimated by total peripheral resistance $TPR$) | 1.253 | 0.5271 – 1.2048 (Klingensmith et al., 2016) | s·mmHg·mL$^{-1}$ |
| 092 | $Systole$ | Indicator of the total actual systole | 0.0 | – | – |
| 093 | $Systole_L$ | Indicator of the actual LV systole | 0.0 | – | – |
| 094 | $Systole_{L\_Exp}$ | Indicator of the nominal LV systole | 0.0 | – | – |
| 095 | $Systole_{Length\_L}$ | Duration of the actual LV systole | 0.278 | – | s |
| 096 | $Systole_{Length\_L\_Exp}$ | Duration of the nominal LV systole | 0.278 | – | s |
| 097 | $Systole_{Length\_R}$ | Duration of the actual RV systole | 0.286 | – | s |
| 098 | $Systole_{Length\_R\_Exp}$ | Duration of the nominal RV systole | 0.286 | – | s |
| 099 | $Systole_R$ | Indicator of the actual RV systole | 0.0 | – | – |
| 100 | $Systole_{R\_Exp}$ | Indicator of the nominal RV systole | 0.0 | – | – |
| 101 | $TBW$ | Total body water | 35.521 | 24.45 – 56.63 (Hoffer et al., 1969) | L |
| 102 | $TPR$ | Total peripheral resistance | 18.688 | 12.5 – 22.5 (Daly and Bondurant, 1962) | mmHg·min·L$^{-1}$ |
| 103 | $V$ | Total blood volume | 4610.630 | 3061 – 6092 (Wennesland et al., 1959) | mL |

| | | | | | |
|---|---|---|---|---|---|
| 104 | $V_{AL}$ | Systemic arterial blood volume | 595.910 | – | mL |
| 105 | $V_{AR}$ | Pulmonary arterial blood volume | 153.369 | – | mL |
| 106 | $V_{ecf}$ | Extracellular fluid volume | 15.843 | – | L |
| 107 | $V_{HL}$ | LV blood volume | 104.098 | – | mL |
| 108 | $V_{HL\_KD}$ | LV end-diastolic volume | 123.058 | Men: 67 – 155; Women: 56 – 104 (Lang et al., 2006) | mL |
| 109 | $V_{HL\_KS}$ | LV end-systolic volume | 55.130 | Men: 22 – 58; Women: 19 – 49 (Lang et al., 2006) | mL |
| 110 | $V_{HR}$ | RV blood volume | 101.156 | – | mL |
| 111 | $V_{HR\_KD}$ | RV end-diastolic volume | 127.237 | Men: 124 – 256; Women: 78 – 218 (Hudsmith et al., 2005) | mL |
| 112 | $V_{HR\_KS}$ | RV end-systolic volume | 59.310 | Men: 38 – 118; Women: 20 – 92 (Hudsmith et al., 2005) | mL |
| 113 | $V_{VL}$ | Systemic venous blood volume | 3298.698 | ~ 2000 – 3500 (Hall, 2011) | mL |
| 114 | $V_{VR}$ | Pulmonary venous blood volume | 357.400 | – | mL |
| 115 | $V_{AR}+V_{VR}$ | Pulmonary blood volume | 510.769 | ~ $0.09 \cdot V$ – $0.10 \cdot V$ (Hall, 2011; Gazioglu and Yu, 1967) | mL |
| 116 | $V_{AR}+V_{VR}+V_{HR}+V_{HL}$ | Cardiopulmonary blood volume | 716.023 | ~ $0.153 \cdot V$ (Levinson et al., 1996) | mL |
| 117 | $VO_2$ | Venous oxygen content | 0.125 | 0.095 – 0.168 (Hattori et al., 2004) | – |
| 118 | $Y_{ALVL}$ | Conductivity of the systemic microvessels | 1.609 | – | $mL \cdot s^{-1} \cdot mmHg^{-1}$ |
| 119 | $Y_{ARVR}$ | Conductivity of the pulmonary microvessels | 15.273 | – | $mL \cdot s^{-1} \cdot mmHg^{-1}$ |
| 120 | $Y_{VLHR}$ | Conductivity of the tricuspid valve and systemic veins | 125.542 | – | $mL \cdot s^{-1} \cdot mmHg^{-1}$ |
| 121 | $Y_{VRHL}$ | Conductivity of the mitral valve and pulmonary veins | 94.801 | – | $mL \cdot s^{-1} \cdot mmHg^{-1}$ |
| 122 | $\alpha_{map}$ | Effect of mean arterial pressure on renal sympathetic nerve activity | 1.162 | – | – |
| 123 | $\alpha_{rap}$ | Effect of right atrial pressure on renal sympathetic nerve activity | 0.993 | – | – |
| 124 | $\beta_{rsna}$ | Effect of renal sympathetic nerve activity on afferent arteriole resistance and resistance of interlobar, arcuate, and interlobular arteries | 1.231 | – | – |
| 125 | $\gamma_{at}$ | Effect of AT1-bound angiotensin II on fractional sodium reabsorption in the proximal tubule and loop of Henle | 0.998 | – | – |
| 126 | $\gamma_{filsod}$ | Effect of filtered sodium load on fractional sodium reabsorption in the proximal tubule and loop of Henle | 1.100 | – | – |
| 127 | $\gamma_{rsna}$ | Effect of renal sympathetic nerve activity on fractional sodium reabsorption in the proximal tubule and loop of Henle | 1.002 | – | – |
| 128 | $\eta_{cd\_sodreab}$ | Fractional collecting duct sodium reabsorption | 0.876 | – | – |
| 129 | $\eta_{dt\_sodreab}$ | Fractional distal sodium reabsorption | 0.460 | – | – |
| 130 | $\eta_{pt\_sodreab}$ | Fractional sodium reabsorption in the proximal tubule and loop of Henle | 0.935 | 0.67 – 0.97 (Fliser et al., 1997; Bochud et al., 2009; Jin et al., 2009) | – |

| | | | | | |
|---|---|---|---|---|---|
| 131 | $\eta_{dt\_sodreab} + \eta_{cd\_sodreab} - -\eta_{dt\_sodreab} \cdot \eta_{cd\_sodreab}$ | Fractional sodium reabsorption in the distal tubule and subsequent parts of the nephron | 0.933 | 0.78 – 0.98 (Fliser et al., 1997; Bochud et al., 2009; Jin et al., 2009) | – |
| 132 | $\lambda_{anp}$ | Effect of natriuretic peptide on fractional collecting duct sodium reabsorption | 0.998 | – | – |
| 133 | $\lambda_{dt}$ | Effect of distal sodium outflow on fractional collecting duct sodium reabsorption | 1.073 | – | – |
| 134 | $\mu_{adh}$ | Effect of antidiuretic hormone concentration on tubular water reabsorption rate | 0.774 | – | – |
| 135 | $\nu_{AT1\_ANGII}$ | Effect of AT1-bound angiotensin II on renin secretion rate | 0.986 | – | – |
| 136 | $\nu_{MD\_sod}$ | Effect of macula densa sodium flow on renin secretion rate | 1.222 | – | – |
| 137 | $\nu_{RSNA}$ | Effect of renal sympathetic nerve activity on renin secretion rate | 1.126 | – | – |
| 138 | $\xi_{at}$ | Effect of angiotensin hormone on aldosterone secretion rate | 1.294 | – | – |
| 139 | $\xi_{k\_sod}$ | Effect of potassium concentration on aldosterone secretion rate | 0.653 | – | – |
| 140 | $\Sigma_{myo}$ | Myogenic autoregulation signal | 1.025 | – | – |
| 141 | $\Sigma_{tgf}$ | Tubuloglomerular feedback signal | 0.455 | – | – |
| 142 | $\Phi_{cd\_sodreab}$ | Absolute collecting duct sodium reabsorption rate | 0.337 | – | $mEq \cdot min^{-1}$ |
| 143 | $\Phi_{dt\_sod}$ | Distal sodium outflow | 0.384 | – | $mEq \cdot min^{-1}$ |
| 144 | $\Phi_{dt\_sodreab}$ | Absolute distal sodium reabsorption rate | 0.327 | – | $mEq \cdot min^{-1}$ |
| 145 | $\Phi_{filsod}$ | Amount of sodium filtered from the glomerulus to the proximal tubule per minute (filtered sodium load) | 11.002 | 10.0 – 23.0 (Natarajan et al., 2016) | $mEq \cdot min^{-1}$ |
| 146 | $\Phi_{md\_sod}$ | Macula densa sodium flow rate | 0.712 | – | $mEq \cdot min^{-1}$ |
| 147 | $\Phi_{pt\_sodreab}$ | Absolute proximal sodium reabsorption rate | 10.290 | – | $mEq \cdot min^{-1}$ |
| 148 | $\Phi_{t\_wreab}$ | Tubular water reabsorption rate | 0.075 | – | $L \cdot min^{-1}$ |
| 149 | $\Phi_{u}$ | Urine flow rate | 0.0011 | 0.0011 ± 0.0005 (Malisova et al., 2016) | $L \cdot min^{-1}$ |
| 150 | $\Phi_{u\_sod}$ | Urine sodium flow rate | 0.048 | 0.097 ± 0.049 (Letcher et al., 1981) | $mEq \cdot min^{-1}$ |
| 151 | $\Phi_{win}$ | Water intake | 0.0011 | 0.0019 ± 0.0007 (Malisova et al., 2016) | $L \cdot min^{-1}$ |
| 152 | $\psi_{al}$ | Effect of aldosterone on fractional distal sodium reabsorption | 1.047 | – | – |
| 153 | $\Psi_{AT1\_aa}$ | Effect of AT1-bound angiotensin II on afferent arteriole resistance | 2.291 | – | – |
| 154 | $\Psi_{AT1\_ALVL}$ | Effect of AT1-bound angiotensin II on blood flow through the systemic microvessels | 1.773 | – | – |
| 155 | $\Psi_{AT1\_Baro}$ | Effect of AT1-bound angiotensin II on baroreceptor activity | 1.001 | – | – |
| 156 | $\Psi_{AT1\_ea}$ | Effect of AT1-bound angiotensin II on efferent arteriole resistance | 2.290 | – | – |
| 157 | $\Psi_{AT1\_preglom}$ | Effect of AT1-bound angiotensin II on resistance of interlobar, arcuate, interlobular arteries | 1.546 | – | – |
| 158 | $\Psi_{AT1\_Stress}$ | Effect of AT1-bound angiotensin II on stress receptor activity | 1.002 | – | – |
| 159 | $\omega_{AL}$ | Unstressed volume of the systemic arteries | 488.449 | – | mL |
| 160 | $\omega_{AR}$ | Unstressed volume of the pulmonary arteries | 80.947 | – | mL |

**Table S5.** Target variables of antihypertensive drugs in the model[6]

| Therapy | Target variables | | | Drug effect | | Value ($E_v$) | References |
|---|---|---|---|---|---|---|---|
| | **Module** | **Symbol** | **Definition** | **Sign[7]** | **Parameter** | | |
| Aliskiren, 300 mg/day | Renal system | $PRC$ | Plasma renin concentration | – | $DRI$ | 0.928 | (Kutumova et al., 2022) |
| Amlodipine, 5 mg/day | Renal system | $R_{aa}$ | Single afferent arteriole resistance | – | $CCB_{aa}$ | 0.413 | (Kutumova et al., 2022) |
| | | $R_{ea}$ | Single efferent arteriole resistance | – | $CCB_{ea}$ | 0.107 | |
| | | $R_{preglom}$ | Resistance of the interlobar, arcuate, and interlobular arteries | – | $CCB_{preglom}$ | 0.413 | |
| | Cardiovascular system | $R_{ALVL}$ | Resistance of the systemic microvessels | – | $CCB_{sys}$ | 0.107 | |
| Bisoprolol, 5 mg/day | Cardiovascular system | $nS$ | Activity of the stress receptors | – | $B_{blocker}$ | 0.467 | Values were fitted to SBP, DBP, and HR response in clinical trials (Porthan et al., 2009). |
| | | $K_L$, $K_R$ | Inotropic factors of the left and right ventricles | – | $B_{blocker\_in}$ | 0.092 | |
| | | $G_{AL}$ | Systemic arterial elasticity | – | $B_{blocker\_st}$ | 0.149 | |
| | Renal system | $R_{sec}$ | Renin secretion rate | – | $B_{blocker\_rs}$ | 0.926 | |
| Enalapril, 20 mg/day | Renal system | $c_{ACE}$ | Rate of conversion of angiotensin I to angiotensin II by ACE | – | $ACEi$ | 0.996 | (Kutumova et al., 2022) |
| HCTZ, 12.5 mg/day | Renal system | $\eta_{dt\_sodreab}$ | Fractional distal sodium reabsorption | – | $Diuretic_{Inhibition}$ | 0.292 | Values were adjusted to better fit SBP and DBP response in clinical trials (MacKay et al., 1996), taking into account the following long-term dynamics:<br>- PRA increases by 45% (Villamil et al., 2007);<br>- HR, ECFV, GFR, and CO do not change significantly (van Brummelen et al., 1980; Shah et al., 1978; Scaglione et al., 1992; Scaglione et al., 1995; Duarte and Cooper-DeHoff, 2010; Rapoport and Soleimani, 2019; Leth, 1970; Digne-Malcolm et al., 2016). |
| | | $R_{sec}$ | Renin secretion rate | + | $Diuretic_{Stimulation}$ | 1.095 | |
| | | $R_{aa}$ | Single afferent arteriole resistance | – | $Diuretic_{aa}$ | 0.358 | |
| | | $R_{ea}$ | Single efferent arteriole resistance | – | $Diuretic_{ea}$ | 0.202 | |
| | | $R_{preglom}$ | Resistance of the interlobar, arcuate, and interlobular arteries | – | $Diuretic_{preglom}$ | 0.419 | |
| | Cardiovascular system | $R_{ALVL}$ | Resistance of the systemic microvessels | – | $Diuretic_{sys}$ | $\min\left(\frac{time}{1950000}\cdot 0.063, 0.063\right)$ | |
| | | $nS$ | Activity of the stress receptors | – | $Diuretic_{stress}$ | 0.355 | |
| | Renal system | $C_K$ | Potassium level in the blood | – | $Diuretic_{potassium}$ | 0.030 | (Kutumova et al., 2022) |
| | | $urea$ | Urea level in the blood | + | $Diuretic_{urea}$ | 0.100 | |
| Losartan, 100 mg/day | Renal system | $c_{AT1}$ | Rate of angiotensin II binding to the AT1 receptors | – | $ARB$ | 0.954 | (Kutumova et al., 2022) |

[6] ACE = angiotensin-converting enzyme; CO = cardiac output; DBP = diastolic blood pressure; ECFV = extracellular fluid volume; GFR = glomerular filtration rate; HCTZ = hydrochlorothiazide; HR = heart rate; PRA = plasma renin activity; SBP = systolic blood pressure.

[7] Stimulation: "+"; inhibition: "–".

**Table S6.** List of parameter ranges used to create virtual patients

| № | Parameters | Notations | Ranges | | | Units |
|---|---|---|---|---|---|---|
| | | | **Norm** | **Hypertension** | **References** | |
| 01 | Total metabolic intensity | $A_1$ | 0.00032 – 0.00128 | | (Kutumova et al., 2021) | mL$^{-1}$ |
| 02 | Sympathetic sensitivity of the systemic microvessels | $A_3$ | 0.1 | 0.1 – 0.6 | (Proshin and Solodyannikov, 2006; Kutumova et al., 2021) | mL·mmHg$^{-1}$ |
| 03 | Systemic arterial tone | $A_9$ | 0.07 | 0.07 – 0.09 | (Proshin and Solodyannikov, 2006; Kutumova et al., 2021) | mmHg·s·mL$^{-1}$ |
| 04 | Normal concentration of antidiuretic hormone | $C_{adh\_norm}$ | 1.0 – 13.3 | | (Yarmohammadi et al., 2015) | pg·mL$^{-1}$ |
| 05 | Normal concentration of aldosterone | $C_{al\_norm}$ | 70 – 300 | | (Fischbach, 2003) | pg·mL$^{-1}$ |
| 06 | Normal concentration of natriuretic peptide | $C_{anp\_norm}$ | 7.4 – 152.0 | | (Cannone et al., 2018; Nozuki et al., 1986) | ng·L$^{-1}$ |
| 07 | Oxygen capacity of hemoglobin | $C_H$ | 1.32 – 1.39 | | (Dijkhuizen et al., 1977) | mL·g$^{-1}$ |
| 08 | Serum potassium | $C_K$ | 3.5 – 5.5 | | (Rastegar, 1990) | mEq·L$^{-1}$ |
| 09 | Cardiac output (in the renal submodel) | $CO$ | 2.51 – 9.00 | | (Cattermole et al., 2017) | L·min$^{-1}$ |
| 10 | Afferent arteriolar diameter | $d_{aa}$ | 8.7 – 23.9 | | (Neal et al., 2019; Hill et al., 2006) | µm |
| 11 | Efferent arteriolar diameter | $d_{ea}$ | 12.2 – 20.1 | | (Kutumova et al., 2021) | µm |
| 12 | Average normal value of the Frank-Starling law threshold | $FS_{threshold0}$ | 0.0 – 40.0 | | (Kutumova et al., 2021) | mL |
| 13 | Basic elasticity of the systemic arteries | $G_{AL0}$ | 0.33 – 1.00 | 0.33 – 1.67 | (Laskey et al., 1990; Haluska et al., 2010) | mmHg·mL$^{-1}$ |
| 14 | Basic elasticity of the pulmonary arteries | $G_{AR0}$ | 0.08 – 0.26 | | (Thenappan et al., 2016) | mmHg·mL$^{-1}$ |
| 15 | Left ventricular wall elasticity | $G_{HL}$ | 0.01 – 0.43 | 0.02 – 0.72 | (Zhang and Kovács, 2008; Kutumova et al., 2021) | mmHg·mL$^{-1}$ |
| 16 | Right ventricular wall elasticity | $G_{HR}$ | 0.01 – 0.43 | 0.02 – 0.72 | (Zhang and Kovács, 2008; Kutumova et al., 2021) | mmHg·mL$^{-1}$ |
| 17 | Basic elasticity of the systemic veins | $G_{VL0}$ | 0.01 – 0.05 | | (Kutumova et al., 2021) | mmHg·mL$^{-1}$ |
| 18 | Basic elasticity of the pulmonary veins | $G_{VR0}$ | 0.01 – 0.05 | | (Kutumova et al., 2021) | mmHg·mL$^{-1}$ |
| 19 | Plasma glucose | *glucose* | 3.9 – 6.1 | | (Dedov et al., 2017) | mmol·L$^{-1}$ |
| 20 | Hematocrit | *Hct* | Men: 40 – 54<br>Women: 36 – 48 | | (Billett, 1990) | % |
| 21 | Hemoglobin | *He* | Men: 140 – 180<br>Women: 120 – 160 | | (Billett, 1990) | g·L$^{-1}$ |
| 22 | Basic activity of the cardiac center | $Heart_{Base}$ | 0.01 – 1.00 | | (Kutumova et al., 2021) | – |
| 23 | Ratio of unstressed volume to stressed volume of the systemic arteries | $k_{AL}$ | 0.7 – 1.0 | | (Magder, 2016) | – |
| 24 | Ratio of unstressed volume to stressed volume of the pulmonary arteries | $k_{AR}$ | 0.7 – 1.0 | | (Magder, 2016) | – |
| 25 | Ratio of unstressed volume to stressed volume of the systemic veins | $k_{VL}$ | 0.7 – 1.0 | | (Magder, 2016) | – |
| 26 | Ratio of unstressed volume to stressed volume of the pulmonary veins | $k_{VR}$ | 0.7 – 1.0 | | (Magder, 2016) | – |
| 27 | Ratio of unstressed volume to stressed volume of the left ventricle | $k_{HL}$ | 0.0 – 0.3 | | (Kutumova et al., 2021) | – |
| 28 | Ratio of unstressed volume to stressed volume of the right ventricle | $k_{HR}$ | 0.0 – 0.3 | | (Kutumova et al., 2021) | – |

| | | | | | | |
|---|---|---|---|---|---|---|
| 29 | Normal glomerular filtration coefficient | $K_{FG\_0}$ | 0.0039 – 0.0162 | | (Hoang et al., 2003) | L·min$^{-1}$·mmHg$^{-1}$ |
| 30 | Inotropic status of the left ventricle | $K_{L0}$ | 0.5 – 0.8 | | (Solodyannikov, 1994) | – |
| 31 | Inotropic status of the right ventricle | $K_{R0}$ | 0.5 – 0.8 | | (Solodyannikov, 1994) | – |
| 32 | Afferent arteriolar length | $L_{aa}$ | 101 – 123 | | (Kutumova et al., 2021) | µm |
| 33 | Efferent arteriolar length | $L_{ea}$ | 124 – 152 | | (Kutumova et al., 2021) | µm |
| 34 | Initial value of the total exchangeable sodium | $M_{sod}$ | 2040 – 3950 | | (Farber and Soberman, 1956) | mEq |
| 35 | Number of nephrons in the kidneys | $N_{nephrons}$ | 1.50$E$6 – 3.00$E$6 | 0.80$E$6 – 2.75$E$6 | (Bertram et al., 2011; Hoy et al., 2006) | – |
| 36 | Normal fractional distal sodium reabsorption | $n_{\varepsilon\_dt}$ | 0.3 – 0.7 | | (Kutumova et al., 2021) | – |
| 37 | Normal fractional collecting duct sodium reabsorption | $n_{\eta\_cd}$ | 0.6 – 1.0 | | (Kutumova et al., 2021) | – |
| 38 | Normal fractional proximal sodium reabsorption | $n_{\eta\_pt}$ | 0.67 – 0.97 | | (Fliser et al., 1997; Bochud et al., 2009; Jin et al., 2009) | – |
| 39 | Hydrostatic pressure in the Bowman's space | $P_B$ | 10.0 – 15.0 | | (Digne-Malcolm et al., 2016) | mmHg |
| 40 | Normal value of the glomerular hydrostatic pressure | $P_{gh\_norm}$ | 48.0 – 63.0 | | (Guberina et al., 2013) | mmHg |
| 41 | Initial value of the glomerular capillary oncotic pressure | $P_{go}$ | 23.6 – 34.0 | | (Škrtić et al., 2015; Chagnac et al., 2000; Guasch et al., 1997) | mmHg |
| 42 | Renal venous pressure | $P_v$ | 6.0 | 2.0 – 6.0 | (Digne-Malcolm et al., 2016; Kutumova et al., 2021) | mmHg |
| 43 | Normal plasma renin concentration | $PRC_{nom}$ | 3.42 – 69.4 | | (Perschel et al., 2004) | pg·mL$^{-1}$ |
| 44 | Nominal resistance of interlobar, arcuate, and interlobular arteries | $R_{preglom\_0}$ | 7.0 – 20.0 | 7.0 – 28.0 | (Kutumova et al., 2021) | mmHg·min·l$^{-1}$ |
| 45 | Renal venous resistance | $R_v$ | 11.3 – 20.1 | | (Kutumova et al., 2021) | mmHg·min·l$^{-1}$ |
| 46 | Nominal body oxygen demand | $RO_{20}$ | 2.52 – 5.88 | | (Kutumova et al., 2021) | mL·s$^{-1}$ |
| 47 | Arterial oxygen saturation | $SpO_2$ | 0.95 – 0.99 | 0.92 – 0.99 | (Goldberg et al., 2017; Kutumova et al., 2021) | – |
| 48 | Total protein | $TP$ | 60.0 – 86.0 | | (Busher, 1990; Gardner and Scott, 1980) | g·L$^{-1}$ |
| 49 | Plasma urea concentration | $urea$ | 1.8 – 7.1 | | (Hosten, 1990) | mmol·L$^{-1}$ |
| 50 | Initial value of the total body water | $TBW$ | $f(0.9 \cdot N_{BV}) – f(1.1 \cdot N_{BV})$[8] | | (Kutumova et al., 2021) | L |
| 51 | Initial value of the venous oxygen content | $VO_2$ | 0.0855 – 0.1848 | | (Kutumova et al., 2021) | – |
| 52 | Equilibrium ratio of plasma renin activity to plasma renin concentration | $X_{PRC\_PRA}$ | 0.61 – 1.42 | | (Kutumova et al., 2021) | fmol·min$^{-1}$·pg$^{-1}$ |
| 53 | Basic conductivity of the systemic microvessels | $Y_{ALVL0}$ | 1.363 | 0.5 – 2.0 | (Kutumova et al., 2021) | mL·s$^{-1}$·mmHg$^{-1}$ |
| 54 | Basic conductivity of the pulmonary microvessels | $Y_{ARVR0}$ | 9.0 – 21.0 | | (Kutumova et al., 2021) | mL·s$^{-1}$·mmHg$^{-1}$ |
| 55 | Nominal value of the macula densa sodium flow rate | $\Phi_{md\_sod\_0}$ | 1.0 – 4.0 | | (Kutumova et al., 2021) | mEq·min$^{-1}$ |
| 56 | Sodium intake | $\Phi_{sodin}$ | 0.0280 – 0.2088 | | (Kutumova et al., 2021) | mEq·min$^{-1}$ |
| 57 | Normal value of water intake | $\Phi_{win\_norm}$ | 0.00096 – 0.00312 | | (Kutumova et al., 2021) | L·min$^{-1}$ |

[8] To calculate *TBW*, we used the formula $f(N_{BV}) = (N_{BV} – 650)/111.5$ (Moore, 1967), where $N_{BV}$ is an estimate defined by Nadler et al. (1962). We considered $N_{BV}$ ± 10% as the range of total blood volume in normal humans.

**Table S7.** List of variable ranges used to create virtual patients

| № | Variables | Notations/formulas | Ranges | | | Units |
|---|---|---|---|---|---|---|
| | | | Norm | Hypertension | References | |
| 01 | Arterial oxygen content | $AO_2$ | 0.145 – 0.244 | | (Hattori et al., 2004) | – |
| 02 | Plasma angiotensin (1-7) | $ANG17$ | 14.1 – 31.7 | 12.7 – 34.9 | (Ferrario et al., 1998; Kutumova et al., 2021) | fmol·mL$^{-1}$ |
| 03 | Plasma angiotensin I | $ANGI$ | 2.8 – 28.5 | | (Lawrence et al., 1990; Nussberger et al., 1992) | fmol·mL$^{-1}$ |
| 04 | Plasma angiotensin II | $ANGII$ | 0.0 – 21.4 | | (Lawrence et al., 1990; Nussberger et al., 1992; Duggan et al., 1993) | fmol·mL$^{-1}$ |
| 05 | Plasma antidiuretic hormone | $C_{adh}$ | 1.0 – 13.3 | | (Yarmohammadi et al., 2015) | pg·mL$^{-1}$ |
| 06 | Plasma aldosterone | $C_{al}$ | 70 – 300 | | (Fischbach, 2003) | pg·mL$^{-1}$ |
| 07 | Plasma natriuretic peptide | $C_{anp}$ | 7.4 – 152.0 | | (Cannone et al., 2018; Nozuki et al., 1986) | ng·L$^{-1}$ |
| 08 | Plasma sodium | $C_{sod}$ | 137 – 147 | | (Payne and Levell, 1968) | mEq·L$^{-1}$ |
| 09 | Cardiac output (in the heart submodel) | $CO$ | 2.51 – 9.00 | | (Cattermole et al., 2017) | L·min$^{-1}$ |
| 10 | Ejection fraction | $EF$ | 50 – 80 | | (Pfisterer et al., 1985; Saghiv and Sagiv, 2017) | % |
| 11 | Blood flow in the systemic microvessels | $F_{ALVL}$ | > 0.0 | | (Kutumova et al., 2021) | mL·s$^{-1}$ |
| 12 | Blood flow in the pulmonary microvessels | $F_{ARVR}$ | > 0.0 | | (Kutumova et al., 2021) | mL·s$^{-1}$ |
| 13 | Peak rate of the transaortic flow | $F_{HLAL\_p}$ | 347.0 – 677.0 | | (Kyhl et al., 2013) | mL·s$^{-1}$ |
| 14 | Peak rate of the transpulmonary flow | $F_{HRAR\_p}$ | 264.5 – 793.0 | | (Kyhl et al., 2013; Macedo et al., 2007) | mL·s$^{-1}$ |
| 15 | Active peak filling rate of the right ventricle | $F_{VLHR\_ap}$ | Men: 23 – 947; Women: 54 – 680 | | (Maceira et al., 2006a) | mL·s$^{-1}$ |
| 16 | Early peak filling rate of the right ventricle | $F_{VLHR\_ep}$ | Men: 8 – 814; Women: -17 – 701 | | (Maceira et al., 2006a) | mL·s$^{-1}$ |
| 17 | Ratio of early to active peak filling rates of the right ventricle | $F_{VLHR\_ep}/F_{VLHR\_ap}$ | Men: -0.5 – 2.5; Women: -0.4 – 2.5 | | (Maceira et al., 2006a) | – |
| 18 | Active peak filling rate of the left ventricle | $F_{VRHL\_ap}$ | Men: 99 – 647; Women: 58 – 508 | | (Maceira et al., 2006b) | mL·s$^{-1}$ |
| 19 | Early peak filling rate of the left ventricle | $F_{VRHL\_ep}$ | Men: 21 – 1034; Women: -13 – 967 | | (Maceira et al., 2006b) | mL·s$^{-1}$ |
| 20 | Ratio of early to active peak filling rates of the left ventricle | $F_{VRHL\_ep}/F_{VRHL\_ap}$ | Men: 0.3 – 5.9; Women: 0.3 – 6.6 | | (Maceira et al., 2006b) | – |
| 21 | Systemic arterial elasticity | $G_{AL}$ | 0.33 – 1.00 | 0.33 – 1.67 | (Laskey et al., 1990; Haluska et al., 2010) | mmHg·mL$^{-1}$ |
| 22 | Pulmonary arterial elasticity | $G_{AR}$ | 0.08 – 0.26 | | (Thenappan et al., 2016) | mmHg·mL$^{-1}$ |
| 23 | Glomerular filtration rate | $GFR$ | 0.060 – 0.135 | | (Levin and Stevens, 2013; Cachat et al., 2015) | L·min$^{-1}$ |
| 24 | Total exchangeable sodium | $M_{sod}$ | 2040 – 3950 | | (Farber and Soberman, 1956) | mEq |
| 25 | Normal fractional sodium reabsorption in the distal tubule and subsequent parts of the nephron | $n_{\varepsilon\_dt} + n_{\eta\_cd} - n_{\eta\_cd} \cdot n_{\varepsilon\_dt}$ | 0.78 – 0.98 | | (Fliser et al., 1997; Bochud et al., 2009; Jin et al., 2009) | – |
| 26 | Plasma osmolality | $osmolality$ | 275 – 295 | | (Fogarty and Loughrey, 2016) | mOsmol·kg$^{-1}$ |
| 27 | Diastolic pulmonary arterial pressure | $P_{AR\_D}$ | 4.0 – 12.0 | | (Marini and Leatherman, 2005; Pagani et al., 1988) | mmHg |

| | | | | | | |
|---|---|---|---|---|---|---|
| 28 | Systolic pulmonary arterial pressure | $P_{AR\_S}$ | 15.0 – 30.0 | | (Marini and Leatherman, 2005; Pagani et al., 1988) | mmHg |
| 29 | Glomerular hydrostatic pressure | $P_{gh}$ | 48.0 – 63.0 | | (Guberina et al., 2013) | mmHg |
| 30 | Glomerular capillary oncotic pressure | $P_{go}$ | 23.6 – 34.0 | | (Škrtić et al., 2015; Chagnac et al., 2000; Guasch et al., 1997) | mmHg |
| 31 | Left ventricular diastolic pressure | $P_{HL\_D}$ | 3.0 – 12.0 | 1.0 – 18.0 | (Pagani et al., 1988; Kasner et al., 2007; Antony et al., 1993) | mmHg |
| 32 | Left ventricular systolic pressure | $P_{HL\_S}$ | 100 – 140 | 100 – 186 | (Pagani et al., 1988; Antony et al., 1993) | mmHg |
| 33 | Right ventricular diastolic pressure | $P_{HR\_D}$ | 2.0 – 8.0 | 0.0 – 10.0 | (Pagani et al., 1988; Ferlinz, 1980; Kasner et al., 2012) | mmHg |
| 34 | Right ventricular systolic pressure | $P_{HR\_S}$ | 15.0 – 30.0 | | (Pagani et al., 1988) | mmHg |
| 35 | Systemic venous pressure | $P_{VL}$ | 2.0 –8.0 | 1.0 – 10.0 | (Klingensmith et al., 2016; Ferlinz, 1980) | mmHg |
| 36 | Pulmonary venous pressure | $P_{VR}$ | 3.0 – 20.0 | | (Kutumova et al., 2021) | mmHg |
| 37 | Plasma renin concentration | $PRC$ | 3.42 – 69.4 | | (Perschel et al., 2004) | $pg \cdot mL^{-1}$ |
| 38 | Resistance of the afferent vessels | $R_{a\_dyne}$ | 753 – 6863 | 3000 – 25000 | (Tsuda et al., 2018; Gomez, 1951; Kutumova et al., 2021) | $dyn \cdot s \cdot cm^{-5}$ |
| 39 | Resistance of the efferent arterioles | $R_{e\_dyne}$ | 1350 – 3400 | | (Kutumova et al., 2021) | $dyn \cdot s \cdot cm^{-5}$ |
| 40 | Resistance of the interlobar, arcuate, and interlobular arteries | $R_{preglom}$ | 7.0 – 20.0 | 7.0 – 28.0 | (Kutumova et al., 2021) | $mmHg \cdot min \cdot L^{-1}$ |
| 41 | Renal blood flow | $RBF$ | 0.623 – 1.730 | | (Bax et al., 2005) | $L \cdot min^{-1}$ |
| 42 | Renal vascular resistance | $RVR$ | 55.0 – 84.0 | 55.0 – 190.0 | (Bauer et al., 1982; Kutumova et al., 2021) | $mmHg \cdot min \cdot L^{-1}$ |
| 43 | Total blood volume | $V$ | $0.9 \cdot N_{BV} - 1.1 \cdot N_{BV}$[9] | | (Kutumova et al., 2021) | mL |
| 44 | Left ventricular end-diastolic volume | $V_{HL\_KD}$ | Men: 67 – 155; Women: 56 – 104 | | (Lang et al., 2006) | mL |
| 45 | Left ventricular end-systolic volume | $V_{HL\_KS}$ | Men: 22 – 58; Women: 19 – 49 | | (Lang et al., 2006) | mL |
| 46 | Right ventricular end-diastolic volume | $V_{HR\_KD}$ | Men: 124 – 256; Women: 78 – 218 | | (Hudsmith et al., 2005) | mL |
| 47 | Right ventricular end-systolic volume | $V_{HR\_KS}$ | Men: 38 – 118; Women: 20 – 92 | | (Hudsmith et al., 2005) | mL |
| 48 | Venous oxygen content | $VO_2$ | 0.0855 – 0.1848 | | (Kutumova et al., 2021) | – |
| 49 | Systemic vascular resistance | $SVR$ | 0.5271 – 1.2048 | 0.5271 – 1.9608 | (Klingensmith et al., 2016; Prys-Roberts et al., 1971) | $s \cdot mmHg \cdot mL^{-1}$ |
| 50 | Pulmonary vascular resistance | $PVR$ | 0.0151 – 0.1353 | | (Klingensmith et al., 2016; Stefanadis et al., 2001) | $s \cdot mmHg \cdot mL^{-1}$ |
| 51 | Fractional proximal sodium reabsorption | $\eta_{pt\_sodreab}$ | 0.67 – 0.97 | | (Fliser et al., 1997; Bochud et al., 2009; Jin et al., 2009) | – |
| 52 | Fractional sodium reabsorption in the distal tubule and subsequent parts of the nephron | $\eta_{dt\_sodreab} + \eta_{cd\_sodreab} - \eta_{dt\_sodreab} \cdot \eta_{cd\_sodreab}$ | 0.78 – 0.98 | | (Fliser et al., 1997; Bochud et al., 2009; Jin et al., 2009) | – |
| 53 | Water intake | $\Phi_{win}$ | 0.00096 – 0.00312 | | (Kutumova et al., 2021) | $L \cdot min^{-1}$ |

[9] $N_{BV}$ is an estimate defined by Nadler et al. (1962). We considered $N_{BV} \pm 10\%$ as the range of total blood volume in normal humans.
